# Assessing Assembly Errors in Immunoglobulin Loci: A Comprehensive Evaluation of Long-read Genome Assemblies Across Vertebrates

**DOI:** 10.1101/2024.07.19.604360

**Authors:** Yixin Zhu, Corey Watson, Yana Safonova, Matt Pennell, Anton Bankevich

## Abstract

Long-read sequencing technologies have revolutionized genome assembly producing near-complete chromosome assemblies for numerous organisms, which are invaluable to research in many fields. However, regions with complex repetitive structure continue to represent a challenge for genome assembly algorithms, particularly in areas with high heterozygosity. Robust and comprehensive solutions for the assessment of assembly accuracy and completeness in these regions do not exist. In this study we focus on the assembly of biomedically important antibody-encoding immunoglobulin (IG) loci, which are characterized by complex duplications and repeat structures. High-quality full-length assemblies for these loci are critical for resolving haplotype-level annotations of IG genes, without which, functional and evolutionary studies of antibody immunity across vertebrates are not tractable. To address these challenges, we developed a pipeline, “CloseRead”, that generates multiple assembly verification metrics for analysis and visualization. These metrics expand upon those of existing quality assessment tools and specifically target complex and highly heterozygous regions. Using CloseRead, we systematically assessed the accuracy and completeness of IG loci in publicly available assemblies of 74 vertebrate species, identifying problematic regions. We also demonstrated that inspecting assembly graphs for problematic regions can both identify the root cause of assembly errors and illuminate solutions for improving erroneous assemblies. For a subset of species, we were able to correct assembly errors through targeted reassembly. Together, our analysis demonstrated the utility of assembly assessment in improving the completeness and accuracy of IG loci across species.

## Introduction

Long-read sequencing technologies have been instrumental in overcoming the limitations of short reads, as they can span large, repetitive, and complex genomic regions that were previously difficult to assemble (1–5). For example, highly abundant LINE repeats that represent a challenge for short read-based assemblies are easily spanned by long reads (5,6). Moreover, the increasing read accuracy of long-read platforms now allows even the most complex repeats to be resolved (like centromeric tandem repeats) as minor paralogous sequence variants between repeat copies can be detected at high-resolution (7). These advances have reduced gaps and ambiguities in genome assemblies, enabling a more complete and accurate depiction of genomic architecture (8–10). And consequently, many research teams and consortia have been able to produce near-complete full-chromosome assemblies for an ever-growing number of organisms, including both model and non-model species; prominent examples include the Telomere-to-Telomere (T2T) consortia (7,11), the Vertebrate Genome Project (VGP) (12) and the California Conservation Genomics Project (CCGP) (13). In addition to being of interest to evolutionary biologists and genome researchers, having complete assemblies for many species has the potential to be transformational for medical genetics, agriculture, conservation biology, and many other disciplines (14–25).

While modern long-read assemblies have made it possible to assemble repetitive and structurally complex regions of the genome, challenges remain in our ability to assess the accuracy and completeness of assemblies spanning such loci (2,26,27). Ensuring that these regions are accurately represented within whole-genome assemblies is essential not just for the sake of completeness, but also because these regions often harbor critically important genes and functional sequences. Arguably the foremost example of these are the Immunoglobulin (IG) loci, which harbor large and expanded families of antibody-encoding genes. These regions are known for their structural complexity (28–35), which has long thwarted efforts to reconstruct them at the genomic level. As a result, our understanding of intra- and inter-species IG genetic diversity remains limited.

Within mammalian and reptilian genomes, IG genes are most often localized to three primary loci: the IG heavy chain (IGH), and the kappa (IGK) and lambda (IGL) light chain loci (36). Each of these loci contain expanded families of variable genes (V), diversity genes (D; in IGH only) and joining genes (J) that represent templates of one of two antibody chains, the heavy chain or the light chain. During B cell development, the IGH loci and one of the light chain loci undergo a somatic process called V(D)J recombination, in which one V gene, one D gene (in IGH only), and joining (J) gene are concatenated into a V(D)J sequence that forms the variable region of either a heavy or light chain transcript (37). Importantly, this combinatorial diversity allows for the generation of a diverse array of antibodies, enabling the immune system to recognize and neutralize a wide range of pathogens.

The organization and counts of germline IG genes vary within and across vertebrate species (25,38). For example, cows have only 11 reported IGH functional V genes, whereas murine IGH loci contain over 100 functional V genes, and the little brown bat is estimated to have 236 V genes in its IGH loci (39–41). The number of genes observed within a locus can also vary among individuals within a species; for instance, humans have been characterized to have many structural variants between different individuals (42). These variations are primarily due to gene duplication and mutation events (43).

Given their pivotal role in immunity, it is essential to characterize IG genes across species at haplotype-level to understand the evolution and functional diversity of immune responses. To ensure that these genes are effectively curated from whole-genome assemblies, the evaluation of assembly accuracy within the IG loci is crucial. Accurate assembly of these loci is vital not only for developing standardized nomenclature across different studies and databases but also serves as a critical test case for our ability to assemble complex genomic regions (42,44,45). This is essential for achieving a complete diploid representation of any genome, which in turn is crucial for fully understanding the diversity and functional dynamics of the immune system (46).

Released assemblies invariably have been subjected to extensive quality control, and a number of methods exist for evaluating the support for a proposed assembly. Some common metrics such as N50, NG50 and L50 are used to assess assembly continuity and fragmentation (47). QUAST is a tool used to calculate various genome assembly metrics, including these continuity metrics and misassemblies—structural differences between the assembly and the reference genome (48). However, for mammalian genomes, the misassembly metric often loses significance due to numerous structural differences between individuals, making it necessary to use the genome of the same biological individual (i.e., not just the same species) as a reference for interpretable results. Thus, the effectiveness of QUAST is limited when there is no suitable reference genome available for comparison. Moreover, for many species, the lack of a reference genome necessitates the use of reference-free methods to evaluate the assembly. Furthermore, QUAST is not particularly suitable for evaluating diploid genomes. The Benchmarking Universal Single-Copy Orthologs (BUSCO) program is another key tool in this domain (49,50). BUSCO evaluates genome completeness using information inferred by highly-conserved genes, and is employed by both VGP and CCGP in their quality assessments. Other advanced techniques further refine the evaluation process, some of which are reference-free. The Long Terminal Repeat (LTR) Assembly Index (LAI) is one such reference-free method, assessing repeat space to refine base-level inaccuracies (51). K-mer-based methods, including JASPER and Merqury, are also reference-free. Merqury refines base-level inaccuracies by assessing k-mer frequencies, while JASPER provides a range of statistical measures for assembly evaluation (52,53). Notably, Merqury is utilized by CCGP. Tools such as Pilon and ntEdit are also commonly used for assembly evaluation (54,55). With the advancement of long-read sequencing, correction methods for long-read assembly, such as Inspector, CONSENT, POLCA, Medaka (https://github.com/nanoporetech/medaka), GCpp (https://github.com/PacificBiosciences/gcpp), are also available (56,57). For detailed curation, CRAQ and Inspector are able to perform error detection at the single-nucleotide resolution and to correct significant structural inaccuracies (47,58).

However, even with existing curation and quality checks, complex and highly polymorphic genomic regions such as the IG loci still represent hot spots for assembly errors. Given the biological importance, it is particularly important to determine whether an assembly accurately represents IG genes and their organization in the genome. Therefore, single nucleotide resolution for quality assessment is essential. However, existing assembly evaluators, such as Inspector and CRAQ, are inadequate for our needs due to the unique challenges posed by IG loci, which exhibit high heterozygosity and often have incomplete diploid assemblies (47,58). Additionally, publicly available data provides only original HiFi raw sequences without distinguishing which reads correspond to which haplotype, making it impossible to meet the evaluators’ input requirements.

To allow us to gain a more fine scale understanding of the quality of the IG assemblies across vertebrate species we developed our own, targeted approach, which we call CloseRead. Our pipeline includes intuitive visualizations to highlight assembly quality in the IG loci, thereby enabling researchers to easily identify and rectify potential errors. These visualizations and corresponding metrics are designed to serve as a complement to the existing quality checks. We applied our pipeline to conduct a systematic evaluation of existing assemblies of 74 vertebrate species (61 mammals and 13 reptiles) to assess their accuracy and completeness. Finally, we conducted three case studies in which we demonstrated manual assessment of problematic regions, confirmed the presence of genuine errors and facilitated correction of the erroneous assembly.

## Results and Discussion

### Overview of Methods

The CloseRead pipeline takes a genome assembly and sequencing reads corresponding to it as an input and aligns the read to the assembly (**Figure 1A**). CloseRead then runs IgDetective tool (59) to identify boundaries of IGH, IGK, and IGL loci in the assembly (**Figure 1B**). Within the IG loci, CloseRead reviews general assembly statistics and identifies mismatches and coverage breaks (**Figure 1C**). Finally, the identified errors are visualized and reported in a user-friendly format (**Figure 1D**). The problematic regions revealed by CloseRead can be further manually inspected or reassembled (**Figure 1E**).

**Figure 1.**
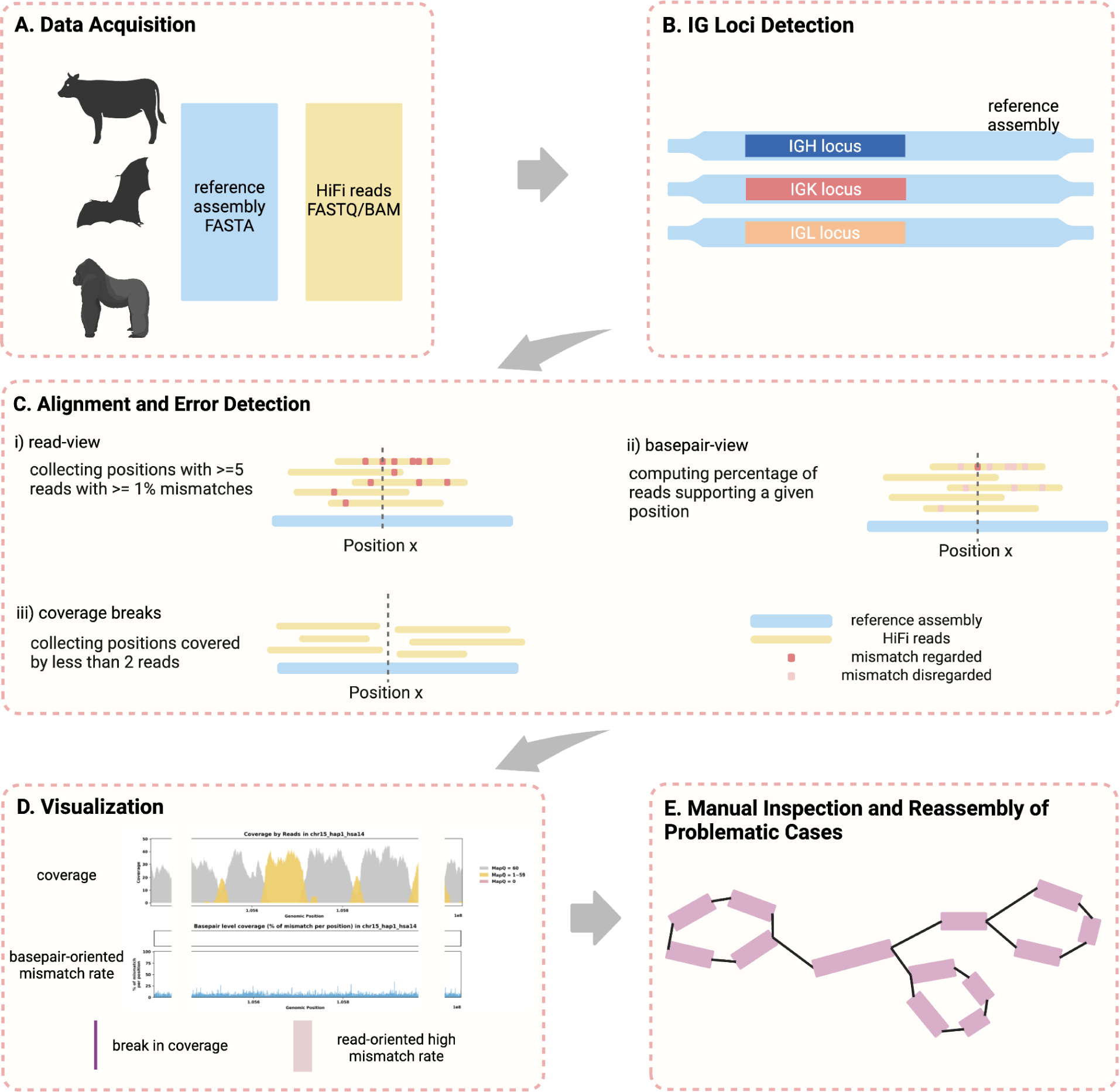
Error Identification and Curation Analysis Overview. A. Publicly available assemblies and the corresponding HiFi sequencing data is downloaded and used as input. B. IGH, IGK and IGL loci location is identified using IgDetective (59) for all species. C. Alignment using minimap2 (60) is done to identify mismatches and breaks in coverage, which could be an indication of potential assembly error. i) low confidence positions are captured from a read view by collecting positions with ≥5 reads that have >1% mismatch rate. ii) mismatch rate is calculated for every position, considering only mismatches occurred exactly at the given position (*basepair-oriented*). iii) coverage breaks are found by collecting positions with ≤2 reads mapped. D. Visualizing assembly evaluation by coverage and basepair-oriented mismatch rate. Purple bars indicate break in coverage. Red highlights indicate positions with high read-oriented mismatch rate computed in C-ii. Shown example is of case with no significant errors found. E. Assembly errors and inaccuracies identified at step C can be further manually investigated and curated.

### Alignment to Detect Assembly Error

The vast majority of near-complete vertebrate assemblies are based on PacBio HiFi reads (5). HiFi reads combine high accuracy (<0.5% error rate) with significant length (15-25 kbp). Thus alignments of HiFi reads to the resulting assembly represent an efficient tool for assembly validation: in case of an error-free assembly, we expect that most reads have almost perfect alignments. Reads with no alignments to the assembly may represent sample contaminations, but reads that have poor alignment to the assembly are very likely to indicate assembly errors or missing sequences.

### Types of Errors Identified

In our comprehensive analysis of IG loci assemblies, we identified two primary types of errors that are possible indications of an inaccurate assembly: i) mismatches, and ii) breaks in coverage (61). We also often observe reads that map to multiple locations in the assembly related to repetitive regions in the genome. Although it presents significant challenges in read placement, as we discuss below, such observations are not necessarily an indication of assembly errors. To further investigate the likelihood of alignment errors, we conducted extensive simulations (**Supplementary Figure 1**). In these simulations, we generated synthetic HiFi reads from human, bovine, and rhesus assemblies, with a virtual diploid human genome created as a combination of haploid GRCh38 and T2T references (7,62). By performing the same alignment analysis, we showed that alignment errors were minimal in the simulated case. These findings confirm that the observed errors are primarily due to assembly inaccuracies rather than misalignments. Below, we also provide a detailed examination of each error type and its implications.

Beyond these errors, the quality of an assembly can also be significantly influenced by the accuracy of its haplotype resolution. Depending on the input data and assembly pipeline, the result of genome assembly of a diploid organism may have different representations. The most complete representation includes a haplotype-resolved assembly capturing both parental haplotypes of a diploid genome and providing detailed genetic insights crucial for precise studies. However, in some assembly projects, available data is not sufficient to reconstruct chromosome-scale phased haplotypes. In such cases, genome assembly output represents a mixture of both haplotypes into one assembly (often referred to as a primary assembly), leaving alternate haplotypes fragmented and/or compressing regions with high heterozygosity, which can obscure important genetic variation (12). We refer to such assemblies as haplotype-unresolved.

### Mismatches

A prevalent type of assembly error in our results is missing sequences. This error is flagged when a significant number of reads align to a specific region of the reference assembly – referred to as region *i* – with a high percentage of mismatches while having increased coverage. These mismatches arise due to failure of the alignment algorithm to find perfect placement of these reads, and it identifies region *i* as the best match for these reads, despite the high number of mismatches and often results in abnormally increased coverage. This often suggests that the reference assembly lacks the correct sequence to accurately represent these reads, indicating the complete absence of a large contig or sequence within the assembly. From our case studies, we often find these mismatch errors are resolved by reassembly and inclusion of the missing contigs.

To note, existing single-nucleotide resolution assembly evaluators, such as CRAQ and Inspector, assume that the assembly being evaluated is relatively complete, meaning all sequences are present but may not be in the correct order or position (47,58). Under this assumption, these tools interpret approximately 50% of reads with high errors or ratio of clipping as normal due to heterozygosity and report assembly errors only when there is a near 100% error rate (over 75% for CRAQ). This assumption fails in cases where the second haplotype is incomplete, leading to consistent error rates around 50%, as observed in the later case study 1 section. Consequently, these tools likely would not detect the missing sequences, as they would not classify the ∼50% error rate as an indicator of such errors and, at least for IG, miss important problems. Unlike CRAQ and Inspector, CloseRead does not assume a relatively complete assembly and uses two approaches to better locate these regions with many mismatches in alignment, one from a *read-oriented view* and one from *basepair-oriented* view.

First, we adopt a *read-oriented* view to analyze genome assembly quality (**Figure 1C i**). We consider *R* to be set of all reads associated with an assembly *G* with a nucleotide at every position 1 through *l* (i.e., *G* = *G*_1_, *G*_2_, …, *G_l_*). For every position *j* in the genome, there is a set of reads *J_j_* ⊆ *R* that covers the position. Each individual read *r* ∈ *J_j_* is of length *n*(*r*) and covers a set of positions *P* where *j* ∈ *P*. We can compute the distance *d* between *g* = *G_P_* and *r* by

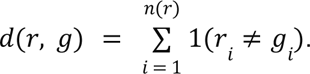

As such, the mismatch percentage *e* for each read can be computed as:

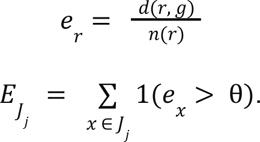

We set θ = 0. 01 to account for any sequencing errors. The overall quality of each genomic position from a *read-oriented* view is subsequently assessed using a threshold condition. For each position *j* in the genome assembly, if *E_J_*_*j*_, the count of reads in *J_J_* whose mismatch percentage exceeds the threshold θ, is greater than 5, then position *j* is deemed poorly supported. This condition can be expressed formally as:

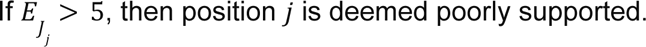

This process is repeated for every position in the reference genome to quantify the *read-oriented* mismatch error rate across the entire genome.

Second, we adopt a *basepair-oriented* view to evaluate the mismatch error rate at specific positions within a reference genome (**Figure 1C ii**). This approach focuses exclusively on each individual position, independent of adjacent base pairs. Again, for every position *j* in the genome, there is a set of reads *J_j_* ⊆ *R* that covers the position. We can compute δ*j*, the counts of errors occurred exactly at position *j*, by

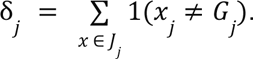

This process is repeated for every position in the reference genome to quantify the *basepair-oriented* mismatch error rate across the entire genome.

### Breaks in Coverage

Another critical indication of assembly error CloseRead identifies is a break in read coverage (**Figure 1C iii**). Continuity of coverage is essential for the accuracy of genomic assemblies; every base pair in the genome is expected to be uniformly (or nearly uniformly) covered by multiple reads (63). Positions with low coverage, which we define as fewer than two reads aligned to a base pair, indicate minimal support and might represent erroneous sequences. For instance, reassembling of the Greenland Wolf (*Canis lupus orion,* hereafter we remove the subspecies designation) individual 2’s IGH region, where we discovered a break in coverage, revealed that a large (1.5 Mb) inversion in the original genome was actually assembled in the reverse orientation (**Figure 6,7**). This finding highlights that breaks in coverage may correspond to substantial structural errors within the assembly, which can be corrected through reassembly.

### Impact of Repetitive Sequences on Read Alignment

In addition to the aforementioned errors, we also encountered cases where repetitive sequences caused ambiguity in read placement. Specifically, when a read maps perfectly to two regions in the genome – regions *Y* and *Z* – the aligner is unable to determine the correct origin of the read due to the identical sequences in the reference. This issue is commonly seen in phased genomes, where similarities between haplotypes complicate accurate read placement, and it is even more prevalent in regions with repetitive sequences, such as IG loci. In these scenarios, the aligner assigns a mapping quality of 0 to these reads and arbitrarily places them in either region *Y* or region *Z*. It is important to note that this situation, characterized by perfect matches and no breaks in coverage, is not considered an assembly error. Instead, it reflects the inherent complexity of repetitive sequences and the challenges they pose for accurate genome alignment and assembly (64). This is also illustrated in the simulations we conducted (**Supplementary Figure 1**).

### Summary of IG Assembly Quality in Vertebrate Genomes

Since haplotype-resolved and haplotype-unresolved assemblies often show significant quality differences in their second haplotype (65), we consider them separately in the following results. Overall, there were no significant disparities in the overall error rates between haplotype-resolved and haplotype-unresolved assemblies. However, the prevalence of each error type was distinct in both assembly types.

We were able to identify the IGH loci for all 74 species using IgDetective (59). The assembly quality results for these loci indicate significant variability. **Table 1** shows that, in the 26 haplotype-resolved assemblies, one species’ assembly (3.8%) exhibited mismatch errors, 15 species’ assemblies (57.7%) had breaks in coverage, and 11 species’ assemblies (42.3%) had no significant errors. In the 48 haplotype-unresolved assemblies, mismatch errors were found in 24 species’ assemblies (50%), 15 species’ assemblies (31.3%) had breaks in coverage, and 20 species’ assemblies (41.7%) had no significant errors. Overall, out of the 74 species, 43 species’ assembly (58.1%) showed some indication of assembly error, either through mismatches or breaks in coverage.

**Table 1:** Summary of Assembly Quality for IGH, IGK, and IGL Loci. This table provides a summary of the number of species’ assemblies that exhibit mismatches, breaks in coverage, and no significant errors. These data are categorized by haplotype-resolved and haplotype-unresolved assemblies across the IGH, IGK, and IGL loci.

| Summary of Assembly Quality for IG loci |  |  |  |  |
| --- | --- | --- | --- | --- |
| Locus | Assembly Type | Mismatch | Breaks in Coverage | Good/No Significant Error |
| IGH | Haplotype Resolved | 1 (3.8%) | 15 (57.7%) | 11 (42.3%) |
|  | Haplotype Unresolved | 24 (50%) | 15 (31.3%) | 20 (41.7%) |
| IGK | Haplotype Resolved | 1 (4.8%) | 3 (14.3%) | 18 (85.7%) |
|  | Haplotype Unresolved | 9 (25%) | 4 (11.1%) | 26 (72.2%) |
| IGL | Haplotype Resolved | 2 (8.7%) | 7 (30.4%) | 16 (69.6%) |
|  | Haplotype Unresolved | 17 (44.7%) | 12 (31.6%) | 16 (42.1%) |

We identified IGK loci in 57 out of 74 species and IGL loci in 61 out of 74 species. Some species naturally lack IGK loci, possessing only IGL loci, and vice versa, either due to inherent absence or high sequence divergence (66,67). For IGK/IGL loci, there were 21/23 haplotype-resolved and 36/38 haplotype-unresolved assemblies. In the haplotype-resolved group, 1/2 assembly (4.8%/8.7%) displayed mismatch errors, 3/7 assemblies (14.3%/30.4%) had coverage breaks, and 18/16 assemblies (85.7%/69.6%) exhibited no significant errors. In the haplotype-unresolved group, mismatch errors were observed in 9/17 assemblies (25%/44.7%), 4/12 assemblies (11.1%/31.6%) had coverage breaks, and 26/16 assemblies (72.2%/42.1%) showed no significant errors. In total, 13 of the 57 species (22.8%) and 29 out of the 61 species (47.5%) presented some form of assembly error in IGK and IGL loci, respectively.

To further understand genome assembly challenges, we categorize breaks based on their coverage and the characteristics of the reference sequences as it could imply different things about the assembly (6,68) (**Supplementary Figure 2**). Overall, our analysis indicates that haplotype-unresolved assemblies have higher mismatch errors, suggesting a systematic issue with representing haplotypes that cannot be fully resolved despite sufficient data. Additionally, we observe a higher rate of coverage breaks in the IGH loci, highlighting a particular challenge in accurately assembling these regions.

### Case Studies

To verify the errors identified by CloseRead and better understand their causes, we present case studies that are emblematic of some of the broader features discovered across the datasets we examined. In the supplementary material, we repeat the same detailed analyses for each of the other 71 genomes we evaluated in this study.

#### Case 1: Mismatches in Alignment Indicating Missing IG Sequences

The assemblies of the Greenland wolf individual 1 (*C. lupus*; GCA_905319855.2) and the Philippine flying lemur (*Cynocephalus volans*; GCA_027409185.1) were constructed as part of the Vertebrate Genome Project (VGP) and are haplotype-unresolved. Specifically, the primary IGH loci of the *C. lupus* individual 1 is 2.5 Mbp, while the alternate IGH loci is about 1.3 Mbp. Similarly, the primary IGH loci of *C. volans* is about 2.3 Mbp, and the alternate IGH loci sums up to 0.8 Mbp. In both case studies, the majority of the reads showed great mapping quality when aligned to the assembly (**Figure 2A i, 3A i**). However, there are a portion of the reads (6.4% and 3.7% respectively) that were soft-clipped in both case studies (**Figure 2A ii, 3A ii**). In addition, both species show significant numbers of reads (7.9% and 14.5% respectively) aligned with high mismatch rates and indels (insertions and deletions greater than 2 consecutive base pairs) (**Figure 2A iii-iv, 3A iii-iv**). Given that the two haplotypes of the IGH loci in both species exhibit substantial differences in length, the mismatches identified likely suggest missing contigs in the alternate assemblies (**Figure 2B-C, 3B-C**).

**Figure 2.**
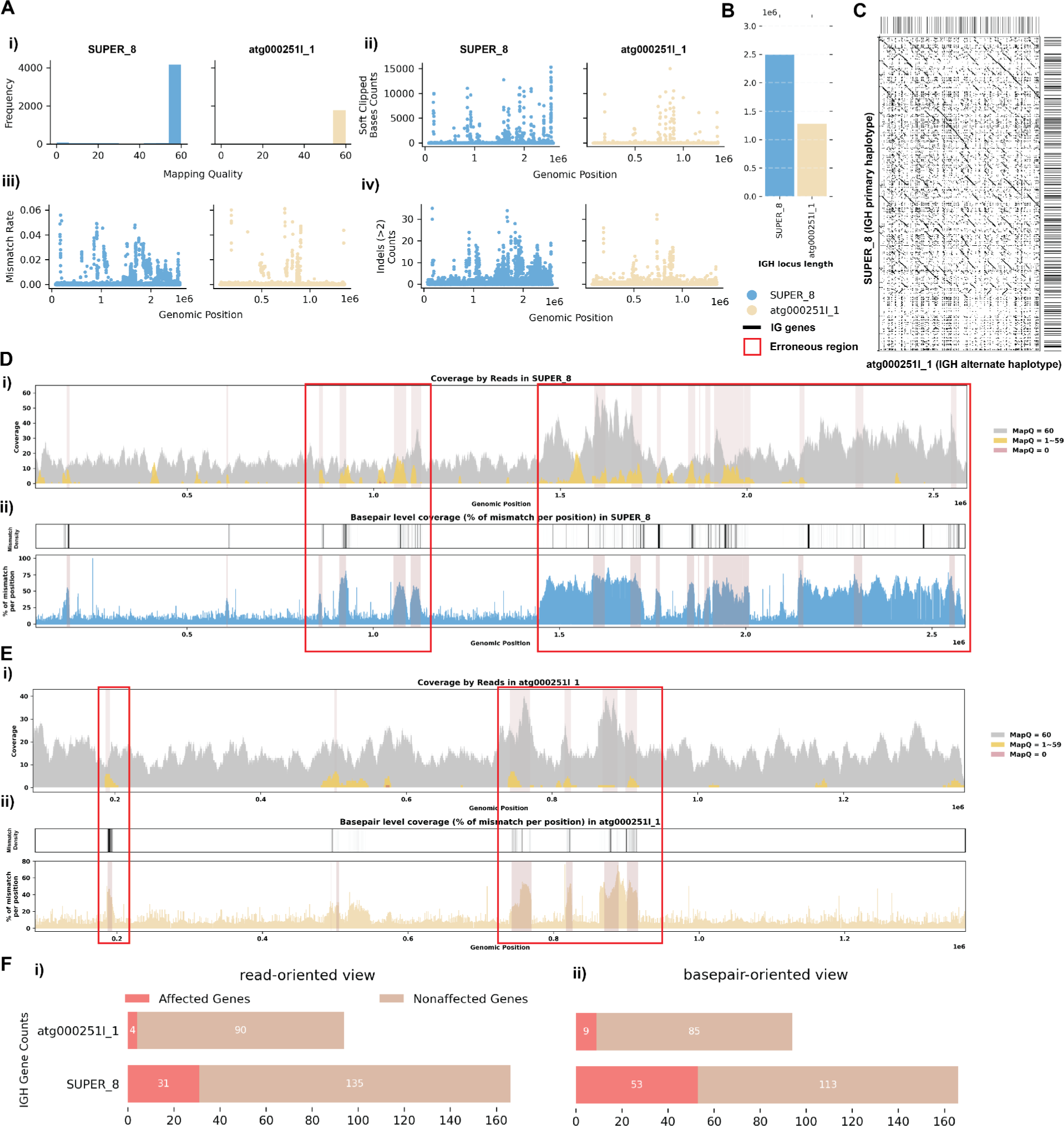
Detailed Analysis of IGH Locus Assembly Errors in Greenland wolf (*C. lupus*) individual 1. A. Summary statistics of the read alignment situation are depicted, showing mapping quality across IGH loci for both haplotypes, with blue representing the primary assembly and yellow the alternate. i) The plots illustrate the read mapping quality frequency, ii) count of soft clipped bases, iii) mismatch rates of reads, and iv) number of indels (consecutive length of at least 2 bp) for both haplotypes across IGH loci. B. IGH locus length in both the primary and alternate assemblies are compared using bar charts. C. Dotplots comparing gene locations and alignments are shown for primary vs alternate haplotypes. We generated dotplots using Gepard(v2.1) (69). D. A detailed analysis of alignment mismatch in the primary IGH haplotype includes i) read coverage across the entire IGH loci, color-coded by mapping quality, and ii) *basepair-oriented* mismatch rate, a heatmap above indicating the frequency of high mismatch rate base pairs, with darker colors denoting more frequent occurrences. Light red highlights positions covered by ≥5 reads with an error rate >1%, and purple bars indicate coverage breaks (coverage ≤2). E. Similar analysis to D for the alternate IGH haplotype. F. IG gene quality from i) *read-oriented* view, affected genes are IGH genes covered by > 5 poorly aligned reads; and ii) *basepair-oriented* view, affected genes are IGH genes without perfect support (>80% confidence at every position)

**Figure 3.**
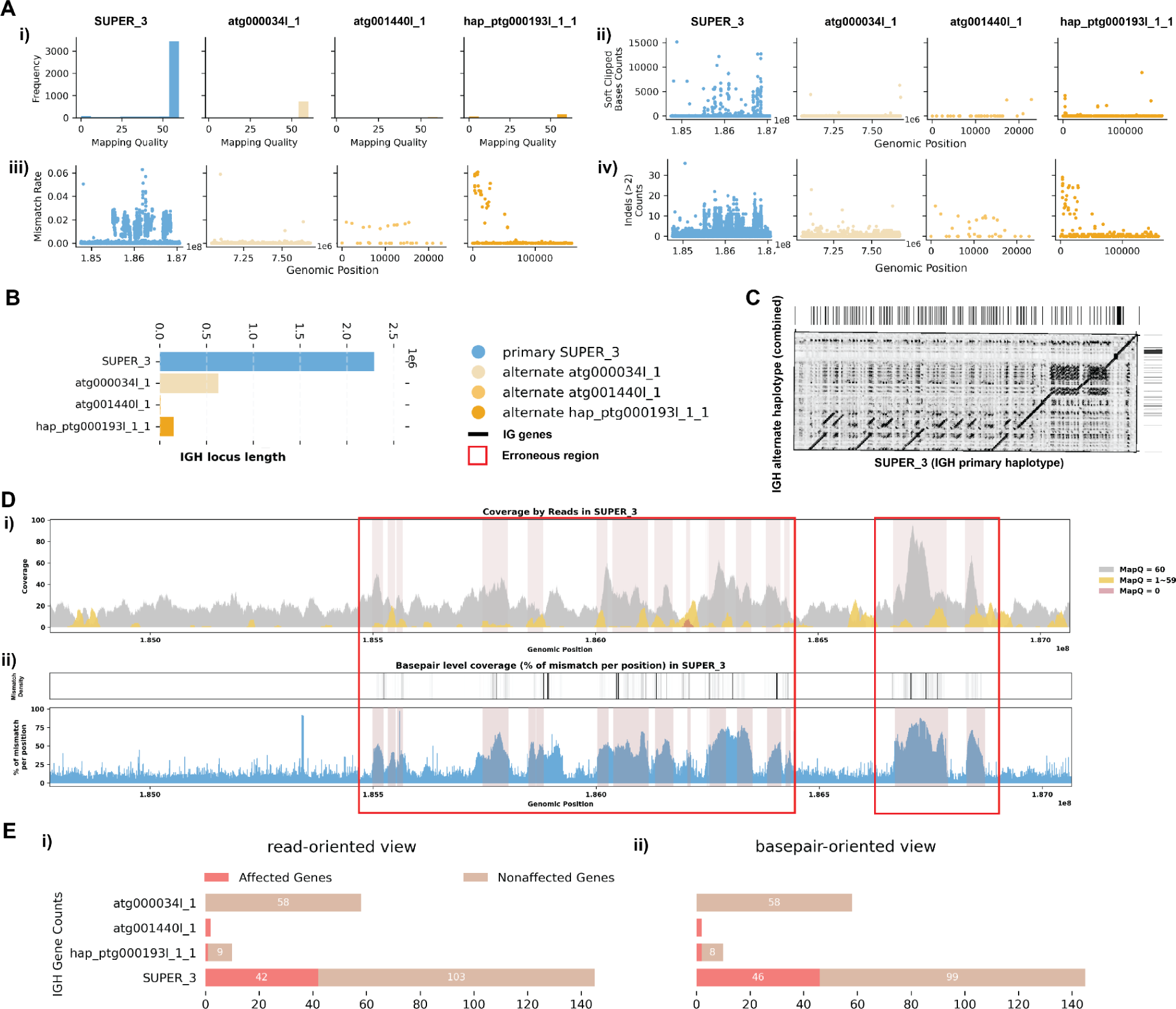
Detailed Analysis of IGH Locus Assembly Errors in Philippine Flying Lemur (*C. volans*). A. Summary statistics of the read alignment are depicted, showing mapping quality across IGH loci for both haplotypes, with blue representing the primary assembly and yellow the alternate. i) The plots illustrate the read mapping quality distribution, ii) count of soft clipped bases, iii) mismatch rates of reads, and iv) number of indels (consecutive length > 2bps) for both haplotypes across IGH loci. B. IGH locus length in both the primary and alternate assemblies are compared using bar charts. C. Dotplots comparing gene locations and alignments are shown for primary vs alternate haplotypes. We generated dotplots using Gepard(v2.1). D. A detailed analysis of alignment mismatch in the primary IGH haplotype includes i) read coverage across the entire IGH loci, color-coded by mapping quality, and ii) *basepair-oriented* mismatch rate, a heatmap above indicating the frequency of high mismatch rate base pairs, with darker colors denoting more frequent occurrences. Light red highlights positions covered by ≥5 reads with an error rate >1%, and purple bars indicate coverage breaks (coverage ≤2). E. IG gene quality from i) *read-oriented* view, affected genes are IGH genes covered by > 5 poorly aligned reads; and ii) *basepair-oriented* view, affected genes are IGH genes without perfect support (>80% confidence at every position)

For the *C. lupus* individual 1 exhibits robust coverage of approximately 30x with high mapping quality reads (**Figure 2D i, 2E i**). Similarly, the *C. volans*’ IGH loci show great coverage, around 20-40x, with most reads having a mapping quality of 60 (**Figure 3D i**). In the *C. lupus* individual 1, there are no observed breaks in coverage, but both haplotypes show sudden increases in coverage, suggesting possible issues due to duplications or repetitive regions (**Figure 2D i**). In the *C. volans*, there is also a sudden increase in coverage in the primary haplotype (SUPER_3), suggesting similar issues (**Figure 3D i**). Detailed analysis from both *read-oriented* and *basepair-oriented* perspectives reveals many regions in both species, which aligned with reads containing mismatches and substantial numbers of base pairs not well supported (2.8% and 2.2% respectively), overlap with these high-coverage regions (**Figure 2D, 2E, 3D; Supplementary Figure 4B**). These mismatches, along with the indels, soft-clipped bases and increased coverage, highlight potential errors in the assemblies. **Supplementary figure 3 and 4** also shows additional evaluation statistics. Furthermore, looking more closely at the IGH gene sequences, where having accurate representations are the most critical, 13.5%/20.9% of the genes are not well supported from the *read-oriented* perspective, and 23.8%/23.3% of the genes lack perfect base support in *C. lupus* individual 1 and *C. volans* respectively (**Figure 2F, 3E**).

To address these issues, we further analyzed the IGH loci for both the *C. lupus* individual 1 and *C. volans* using de Bruijn graphs.

#### Case 1: de Bruijn Graph Analysis and Reassembly

Our observations indicated the existence of missing contigs in both the *C. lupus* individual 1 and the *C. volans* assemblies. To reconstruct missing sequences we had to make additional manual analysis based on de Bruijn graphs. First we generated draft assemblies for these species using a different genome assembly tool. We chose to use La Jolla Assembler (LJA) as it was shown to generate the most reliable assemblies from HiFi reads (70,71). We discovered that in both species contigs reported by LJA contained regions of IG loci which were not present in the VGP assemblies. Nearly all of the reads (99.8%) with low quality alignments to the VGP assembly had nearly perfect alignments to these new regions.

Existence of an assembly supported by reads is significant additional evidence supporting the presence of the error in VGP assembly. To provide additional confirmation of errors in VGP assemblies and confirm LJA assembly results, we analyzed de Bruijn graphs of IG loci. As a part of its output, LJA provides the error-corrected de Bruijn graph, where most erroneous nodes originating from errors in reads are eliminated. The resulting graph provides valuable information about the repeat structure of the genome. **Figures 4A and 5A** show subgraphs and contigs representing IGH loci in *C. lupus* and *C. volans,* correspondingly. Below we discuss each of these two cases in more detail.

**Figure 4.**
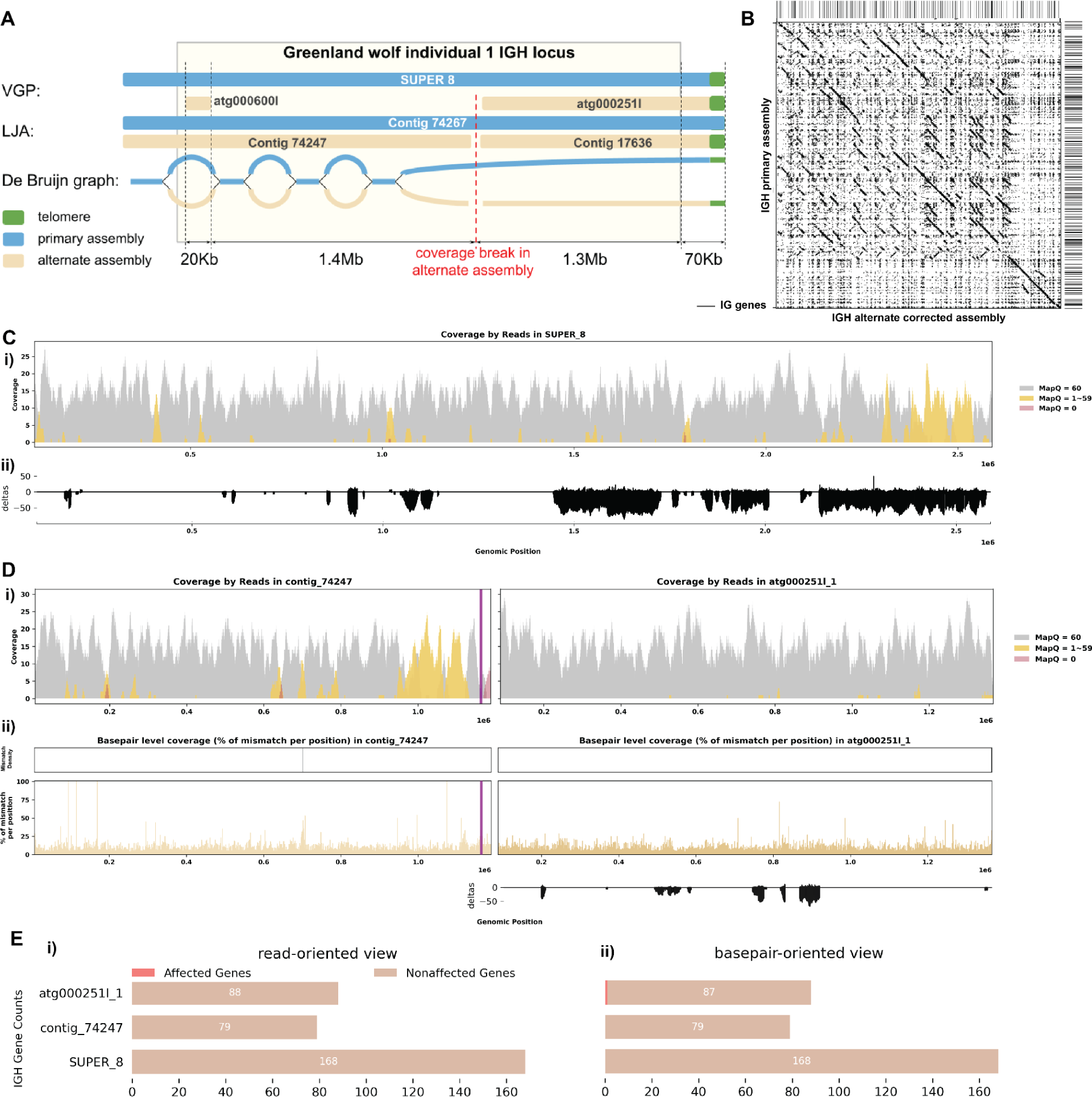
Reassembly Analysis of IGH Locus Assembly Errors in *C. lupus* individual 1. A. Primary and alternate assemblies of *C. lupus* IGH locus produced by VGP and LJA are represented by blue (primary) and tan (alternate) rectangles. Telomere sequences (tandem repeats with repeating sequence CCCTAA) are shown in green. B. Dotplots comparing gene locations and alignments are shown for the corrected alternate IGH assembly vs primary IGH assembly. C. Re-analysis of alignment mismatch in the primary IGH haplotype, i) read coverage across the entire IGH loci, color-coded by mapping quality, and ii) *basepair-oriented* mismatch rate, a heatmap above indicating the frequency of high mismatch rate base pairs, with darker colors denoting more frequent occurrences. Light red highlights positions covered by ≥5 reads with an error rate >1%, and purple bars indicate coverage breaks (coverage ≤2). D. Similar analyses are performed for the corrected alternate haplotype. E. IG gene quality from i) *read-oriented* view, affected genes are IGH genes covered by > 5 poorly aligned reads; and ii) *basepair-oriented* view, affected genes are IGH genes without perfect support (>80% confidence at every position)

**Figure 5.**
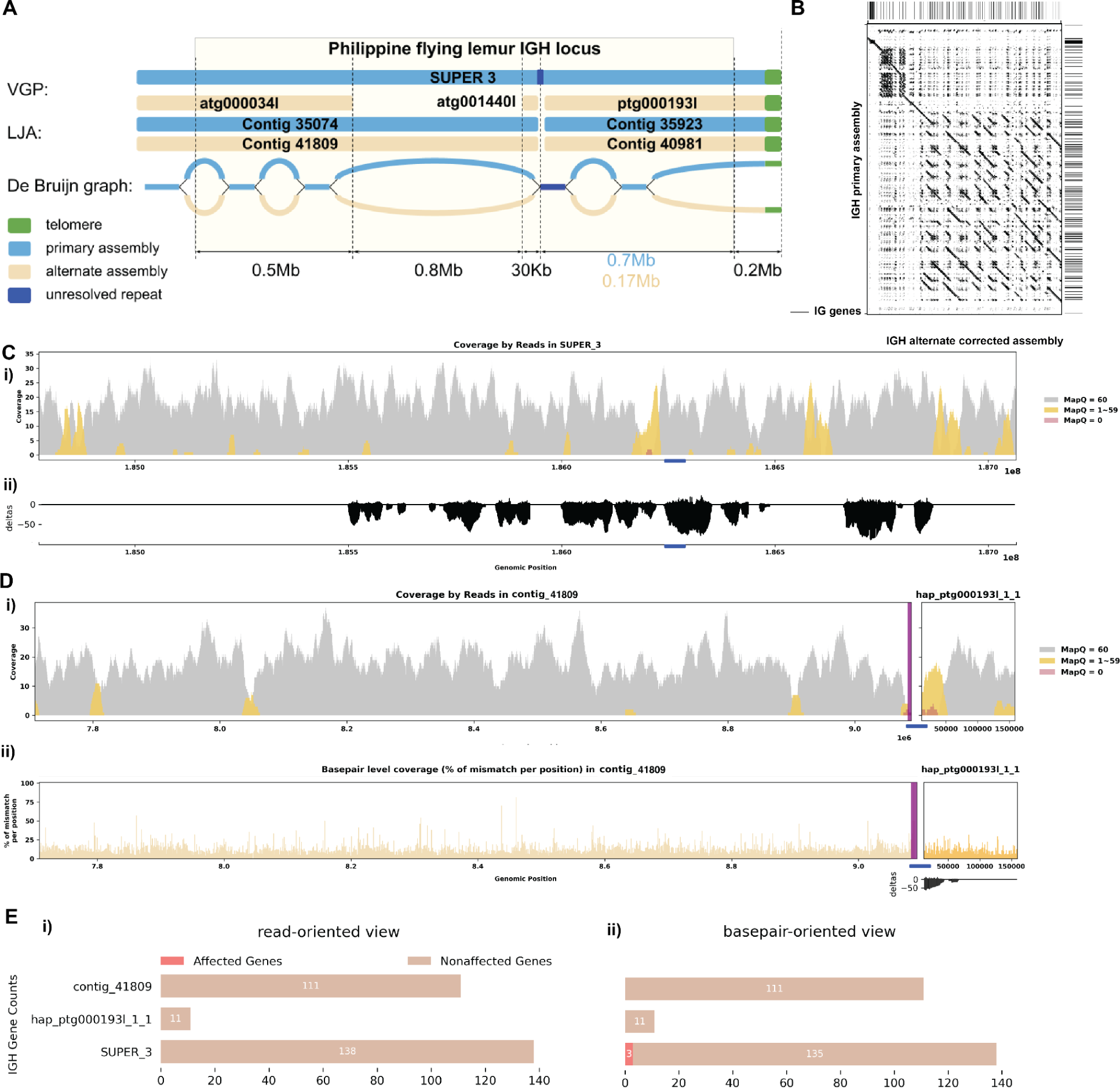
Reassembly Analysis of IGH Locus Assembly Errors in *C. volans*. A. Primary and alternate sequences of IGH locus in VGP and LJA assemblies of *C. volans* are represented by blue (primary) and tan (alternate) rectangles. Telomere sequences (tandem repeats with repeating sequence CCCTAA) are shown in green. Dark blue edge and contig segment represent a 12Kb long conserved fragment. Part of the IGH locus after the blue repeat is very diverged and its length varies from 0.17Mbp for alternate haplotype to 0.7Mbp for primary haplotype. B. Dotplots comparing gene locations and alignments are shown for the corrected alternate IGH assembly vs primary IGH assembly. C. Re-analysis of alignment mismatch in the primary IGH haplotype, i) read coverage across the entire IGH loci, color-coded by mapping quality, and ii) *basepair-oriented* mismatch rate. Light red highlights positions covered by ≥5 reads with an error rate >1%, and purple bars indicate coverage breaks (coverage ≤2). D. Similar analyses are performed for the corrected alternate haplotype. E. IG gene quality from i) *read-oriented* view, affected genes are IGH genes covered by > 5 poorly aligned reads; and ii) *basepair-oriented* view, affected genes are IGH genes without perfect support (>80% confidence at every position)

**Figure 4A** shows that LJA and VGP produced very similar primary assemblies of the *C. lupus* individual 1 IGH locus both consisting of a single contig covering the whole IGH locus and ending with a telomere. Both alternate assemblies consist of two contigs each but VGP assembly is missing a 1.4 Mbp fragment of contig “74247” generated by LJA (**Figure 4A, 4B**). The de Bruijn graph generated by LJA shows a chain of “bulges’’, representing alternating conserved and diverged homologous regions. This chain breaks approximately 1.4 Mbp before the end of the chromosome when the divergence between haplotypes becomes too high to be detected by the de Bruijn graph as it only collapses divergence-free sequences of length at least k (the value of k in the graph generated by LJA is 5001). Reads with low quality alignment to VGP assembly chained together by perfectly overlapping each other and forming a series of edges in the de Bruijn graph, which eventually formed contig “74247”. Incorporating the missing contig from the alternate assembly allowed us to bring both haplotype to length around 2.5 Mbp, and reveal 69 more IG genes than before (**Figure 4B**). Furthermore, re-performing post-correction read alignment resolved the previously found erroneous reads, despite one region with low coverage in contig “74247” near the end (**Figure 4C, 4D; Supplementary Figure 5**). The break in the alternate LJA assembly is explained by a gap in alternate haplotype coverage. However, it is unclear how to explain 1.4 Mbp of missing sequence in the VGP alternate assembly. Looking more closely at the IGH gene sequences, now all IGH genes found are well supported from the *read-oriented* perspective, and only 1 genes lack perfect base support from *basepair-oriented* perspective, with 76.9% base support at 1 position (**Figure 4E**).

**Figure 5A** shows that the primary LJA assembly of the *C. volans* IGH locus is split into two contigs while VGP assembly has a single contig covering the whole IGH locus. The de Bruijn graph shows that the two contigs generated by LJA are separated by a 12 kbp long region that is perfectly conserved between chromosomes (shown in red on **Figure 5A**). It can be assumed that LJA could not resolve this repeat and phase the haplotypes, thus making a decision to report multiple contigs rather than potentially report a longer chimeric contig that combines sequences from different haplotypes. The alternate VGP assembly is also broken by this repeat. It is unclear why this repeat is resolved in the primary assembly, but not in the alternate assembly. Even though VGP reported a less fragmented primary IGH assembly, the alternative assembly again is missing a 0.7 Mbp region from LJA contig “41809”. The De Bruijn graph indicates that the missing sequence is a part of a long fragment of the alternative haplotype that is significantly diverged from the primary haplotype. By incorporating this missing contig, it brings the two haplotype to a more comparable lengths (2.3 Mbp and 1.5 Mbp) and identifies 77 more IG genes than before (**Figure 5B**). Further analysis by re-aligning the reads resolved previously identified erroneous reads (**Figure 5C, 5D; Supplementary Figure 6**). Specifically examining the IGH gene sequences, all identified IGH genes are well supported from a read-oriented perspective. Notably, only 3 genes lack perfect base support from a *basepair-oriented* perspective: one gene has a deletion, and the other two have 66.7% and 76.9% base support at one position each (**Figure 5E**).

#### Case 2: Break in Coverage Revealing False Inversions

The assembly of the Greenland wolf (*C. lupus*) individual 2 was constructed as part of the Vertebrate Genome Project (VGP). Different from the previous cases, this individual has a haplotype-resolved assembly. Due to homology between haplotypes, not all reads were aligned with mapping quality 60 (**Figure 6A i**). We do not observe many reads aligned with mismatches, indels, soft-clipped or hard-clipped bases in either haplotype, suggesting good confidence in base pair correctness overall (**Figure 6A ii-iv**). In addition, the two haplotypes are of similar length, suggesting little likelihood of a missing sequence (**Figure 6B, 6C**). Looking at this assembly from a *read-oriented* view further confirmed that despite both haplotypes exhibit great coverage around 30x, over half of the positions are covered by reads with a mapping quality less than 60, indicating that some reads aligned have lower confidence (**Figure 6D i**). From a *basepair-oriented* view, there are few poorly supported positions (**Figure 6D ii**). **Supplementary figure 7** also shows additional evaluation statistics.

**Figure 6.**
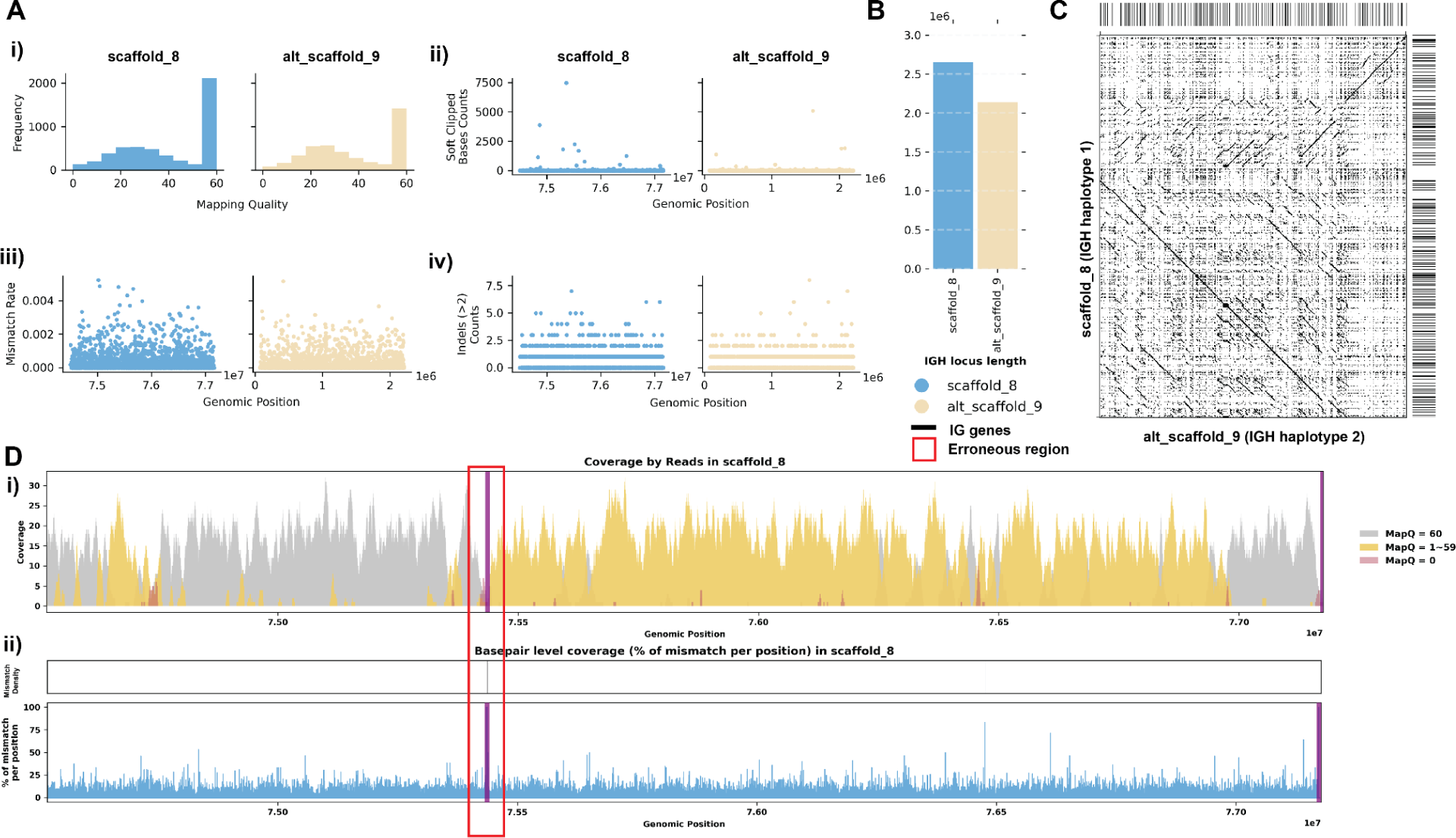
Detailed Analysis of IGH Locus Assembly Errors in Greenland Wolf (*C. lupus*) individual 2. A. Summary statistics of the read alignment are depicted, showing mapping quality across IGH loci for both haplotypes, with blue representing the primary assembly and yellow the alternate. i) The plots illustrate the read mapping quality distribution, ii) count of soft clipped bases, iii) mismatch rates of reads, and iv) number of indels (consecutive length > 2bps) for both haplotypes across IGH loci. B. IGH locus length in both the primary and alternate assemblies are compared using bar charts. C. Dotplots comparing gene locations and alignments are shown for primary vs alternate haplotypes. We generated dotplots using Gepard(v2.1). D. A detailed analysis of alignment mismatch in the primary IGH haplotype includes i) read coverage across the entire IGH loci, color-coded by mapping quality, and ii) *basepair-oriented* mismatch rate, a heatmap above indicating the frequency of high mismatch rate base pairs, with darker colors denoting more frequent occurrences. Light red highlights positions covered by ≥5 reads with an error rate >1%, and purple bars indicate coverage breaks (coverage ≤2).

However, we observe a 8 Kb long break in coverage in the middle of the primary haplotype (scaffold_8:75432695-75440919) (**Figure 6D**). Closer examination reveals that the reference assembly contains consecutive sequences of Ns at this break, indicating missing sequence information, and there are 0 reads spanning this break, confirming a lack of coverage (**Supplementary Figure 7C**). Although it is possible that this break cannot be spanned by HiFi reads and that the Ns were introduced during the scaffolding step, the short length of the break in our case makes this less likely. By comparing the sequences, we observe a significant inversion starting after this break in coverage in the *C. lupus* individual 2’s haplotype 1 IGH assembly when compared to the corresponding haplotype 2 assembly (**Figure 6C**). Considering that from case 1 we have an assembly for the same species but different individuals, it is reasonable to compare the two assemblies as they should have similar germline IG sequences. When comparing the assemblies, we confirmed again there is a 1.5Mbp inversion located at the end of the haplotype 1 IGH contig (**Supplementary** Figure 7E). While this inversion may be real, it is concerning that it coincides with the break in coverage we identified earlier. Therefore, we performed a reassembly of the IGH loci.

#### Case 2: de Bruijn Graph Analysis and Reassembly

**Figure 7A** shows that both LJA and VGP produced primary and alternative assemblies fully covering the IGH locus of Grey wolf. However the alternative assembly produced by LJA does not have the 1.5 Mbp inversion observed in VGP assembly. This inversion is bounded by the low covered region detected by read alignment analysis on one side and by telomeric sequence on the other side. Analysis of the homologous sequence in the LJA primary assembly revealed a telomere-like sequence (tandem repeat with units similar to CCCTAA) (72). We speculate that a spurious overlap between reads from this region and the real telomeric region was detected, resulting in an erroneous link and introducing an erroneous inversion in the VGP assembly (highlighted in red on **Figure 7A**). After correcting for this inversion, the break in coverage was resolved, confirming again that the inversion was indeed the source of the coverage issue (**Figure 7B-C**). This is clearly demonstrated in the post-correction read alignment, where continuous coverage is observed (**Figure 7C**). **Supplementary Figure 8** shows additional re-assembly evaluation statistics.

**Figure 7.**
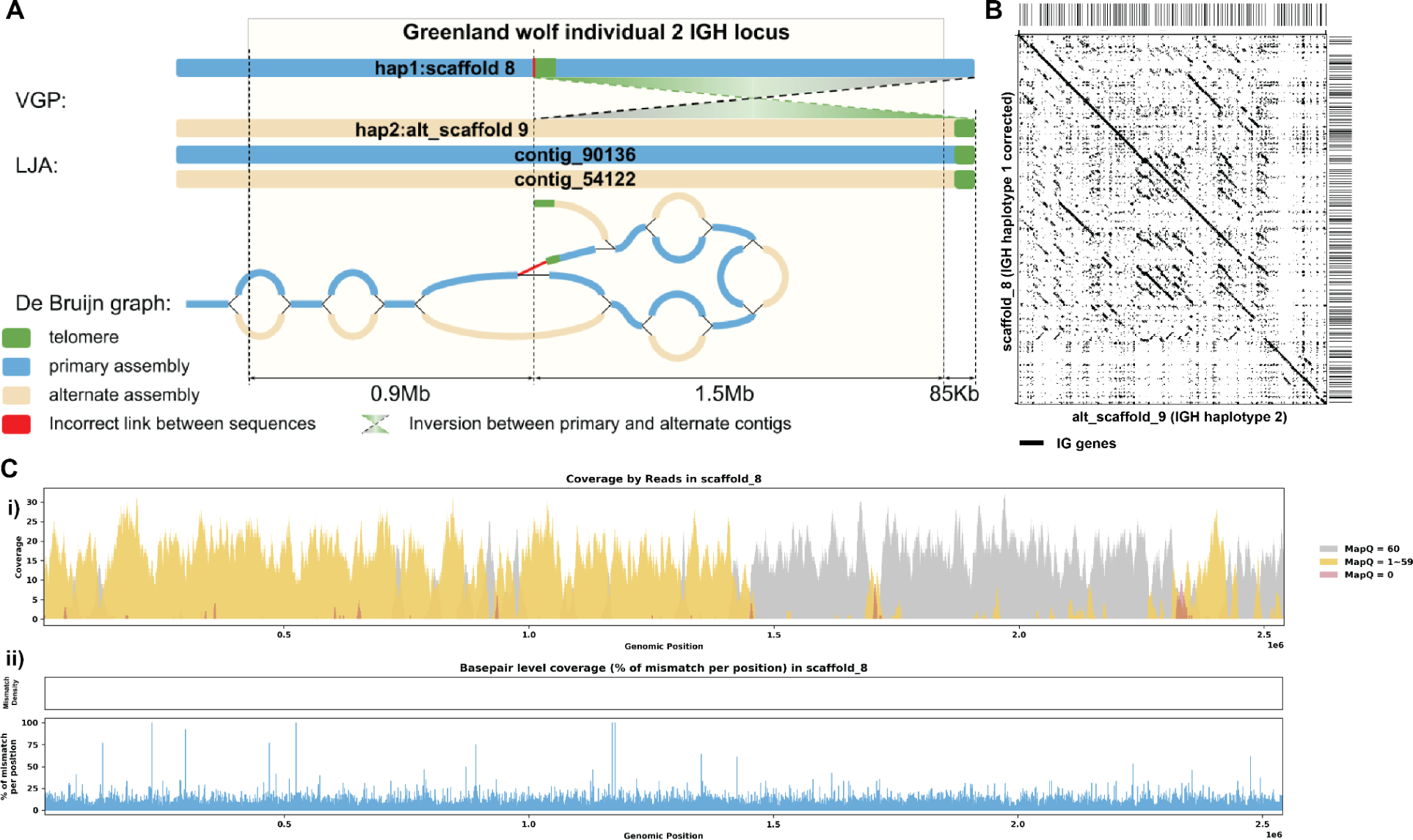
Reassembly Analysis of IGH Locus Assembly Errors in *C. volans.* Individual 2. A. Primary and alternate sequences of IGH locus in VGP and LJA assemblies of *C. lupus* individual 2 are represented by blue (primary) and tan (alternate) rectangles. Telomere sequences (tandem repeats with repeating sequence CCCTAA) are shown in green. Contigs produced by VGP for this region differ by a 1.5Mbp inversion. This inversion puts the telomeric sequence highlighted in green into the middle of contig SUPER_8. Red link corresponds to the inversion breakpoint. It does not have significant support by reading. B. Dotplots comparing gene locations and alignments are shown for the corrected haplotype 1 IGH assembly vs haplotype 2 IGH assembly. C. Re-analysis of alignment mismatch in the corrected haplotype 1 IGH assembly, i) read coverage across the entire IGH loci, color-coded by mapping quality, and ii) *basepair-oriented* mismatch rate, a heatmap above indicating the frequency of high mismatch rate base pairs, with darker colors denoting more frequent occurrences. Light red highlights positions covered by ≥5 reads with an error rate >1%, and purple bars indicate coverage breaks (coverage ≤2).

## Conclusion

In our study, we focus on publicly available genome assemblies which are often utilized by the research community without additional validation. Here we are particularly interested in assessing how well these assemblies are reconstructed for the IG loci. These loci both serve as a test case for the broader problem of reconstructing complex, repetitive regions and are of great interest in their own right, given their great biomedical importance. Accurate assemblies are critical for annotating and naming the genes in the IG loci, characterizing diversity among and within species, as well as understanding the evolutionary causes and immunological consequences of this diversity – all of which may be important for developing new vaccines and therapeutics (25).

Existing single-nucleotide resolution assembly evaluation tools like CRAQ and Inspector have limitations in detecting errors specific to the IG loci. These tools fail to identify and correctly classify errors in complex, repetitive regions like our case studies (**Supplementary Note 1**). To address these shortcomings, we developed CloseRead. This pipeline is specifically designed for complex, highly heterozygous regions like IG. CloseRead offers intuitive visualizations and metrics to highlight assembly quality, enabling easy identification and rectification of errors. Again, we emphasize that our primary motivation for developing CloseRead was to address our own empirical question; we have not rigorously benchmarked this against alternative measures across a range of scenarios – this is beyond the scope of the current paper – although we suspect that it may have wide applicability and assessing this is an obvious point for future research.

Detecting errors is only the first step; further development of scalable methods for assembly process analysis is necessary to reconstruct correct sequences or at least investigate possible alternatives. This requires transforming genome assembly from a one-button black-box solution into a transparent process with clearly outlined alternatives and decisions. Such approaches are essential for identifying and correcting errors, so researchers can make reliable inferences about genomic variation and its consequences for phenotypes. Future work should prioritize automating these approaches and integrating them into existing assembly pipelines to enhance scalability and efficiency.

In conclusion, we conducted a systematic evaluation of 74 species (most of which were mammals) revealing significant deficiencies in the representation of these loci in current assemblies. This finding is crucial as it shows that despite quality checks prior to their release, these assemblies still contain substantial inaccuracies, particularly in complex genomic regions like IG loci. Our results strongly suggest that, at least given current technologies and methodologies, that standard measures of genome-wide assembly support are not reliable indicators of the quality of complex genomic regions like the IG loci. In addition to hopefully spurring the development of more sophisticated tools for assessing and correcting, our results also serve as a useful reminder as to how poorly understood these loci are. Given that it is becoming apparent that germline variation in IG is a major determinant of variation in the adaptive immune response among individuals, it is critical that this knowledge gap is closed (25).

## STAR Methods

### Data Description

The dataset comprises genomic assemblies from three prominent projects: the Vertebrate Genome Project (VGP), the T2T Primates project, and the California Conservation Genomics Project (CCGP).

The Vertebrate Genome Project aims to generate high-quality, near-complete reference genomes for all vertebrate species. Here we utilize the assembled genomes of 59 species from VGP, with an average coverage of approximately 30x. The sequencing methods employed in VGP include high-fidelity (HiFi) reads, Illumina, and HiC. Out of the 59 assemblies, 17 are haplotype-resolved, which means they provide more detailed genetic information by distinguishing between the two sets of chromosomes inherited from each parent.

The T2T Primates project focuses on creating complete, telomere-to-telomere (T2T) assemblies for primate species. Here we utilize the assembled genomes of 4 species, with an average coverage of around 160x, using HiFi, Illumina, HiC, and Oxford Nanopore Technologies (ONT) sequencing methods. All four assemblies produced by the T2T Primates project are haplotype-resolved.

The California Conservation Genomics Project (CCGP) is dedicated to applying genomic technologies to the conservation of California’s biodiversity. Here we utilize the assembled genomes of 11 species using HiFi and HiC sequencing methods, with average coverage around 21x. Among these assemblies, 5 are haplotype-resolved.

In total, the dataset includes genomic assemblies for 74 species, contributed by these three projects. Together, these efforts have resulted in 26 haplotype-resolved assemblies. Majority of these assemblies are for mammalian species, detailed descriptions are available in **supplementary figure 9**.

### Alignment

For aligning the HiFi raw reads to the genome assemblies, we used Minimap2 (version 2.26-r1175) (60). The specific arguments used for the alignment were: minimap2 -ax map-pb. This command ensures optimal alignment settings for PacBio HiFi reads, producing .bam necessary for subsequent analyses.

### IG Gene Annotation

IG gene annotation was performed using IgDetective (version 1.1.0) (59). IgDetective is a specialized tool designed to identify and annotate IG loci in genomic assemblies. The algorithm begins by using minimap2 to align input genomic sequences against a database of known immunoglobulin gene sequences, identifying contigs likely containing IG genes. It then searches these contigs for recombination signal sequences (RSS), which mark V(D)J gene segments.

Once potential V, D, and J segments are identified, the tool refines these candidate genes through an iterative process, involving re-alignment and re-evaluation to enhance accuracy. The final output provides detailed annotations, specifying the locations and sequences of these genes within the genome.

### Error Quantification

We conducted a detailed analysis of genomic assemblies by evaluating aligned read data, with a special focus on spotting and quantifying errors at the IG loci. Initially, we processed the aligned raw data from the .bam and computed the .mpileup files using SAMtools (73). Then we focus on the IG loci using the coordinates found by IgDetective.

To ensure the genomic assemblies were of high quality, we undertook a rigorous statistical examination to catalog the frequency and types of errors. Our analysis involved counting the mismatches between the reads and the reference genome to pinpoint potential assembly errors, and then calculating mismatch rates from both a *read-oriented* and a *basepair-oriented* view. Additionally, we focused on detecting breaks in coverage, specifically areas with fewer than two reads, which could indicate potential issues with the assembly process.

We then summarized this data into comprehensive statistics and insights, which are detailed in the Results section of our paper. Our summary included a detailed breakdown analysis of errors by gene type within the IG loci, with a specific focus on the IGH, IGK, and IGL loci. We also assessed how these errors impact the reliability of genomic assemblies across various species.

This thorough approach has allowed us to assess the quality of the assembly, providing valuable insights into the accuracy and completeness of the IG loci assemblies for the species we studied.

### De Bruijn graph analysis

De Bruijn graph is a powerful instrument for visualization and analysis of sequencing reads. Observing low quality alignments of reads to assembly may just indicate an elevated error rate in reads. But if the de Bruijn graph shows that these reads perfectly overlap each other and form a path, we can conclude that these reads originate from a genome fragment that is missing from the assembly.

We analyzed the corrected de Bruijn graph printed by LJA as a part of its output. In addition we used jumboDBG tool (70) that allowed us to specify the value of k and visualize paths of contigs through the graph. Below we describe basic principles that we used for analysis of the de Bruijn graph.

De Bruijn graph collapses together perfectly matching fragments of the genome of length at least k. In particular homologous heterozygosity-free regions of length at least k would be collapsed, while divergent homologous regions would be represented as pairs of paths that share start and end vertex. Thus homology between haplotypes can be seen in de Bruijn graphs as a “bulge path” representing alternating diverged and non-diverged regions of the genome. In our case studies we identified edges in the graph, corresponding to contig regions with poor read alignments and identified bulge paths containing these edges. Further analyses of edges in these bulge paths and their alignments to primary and alternate assemblies revealed the abnormalities described in case studies.

## Supporting information

Assembly Evaluation Summary

IGH Loci Evaluations

IGK Loci Evaluations

IGL Loci Evaluations

## Declarations

### Availability of data and materials

All datasets used in this manuscript are publicly available, as described under Methods. CloseRead: https://github.com/phylo-lab-usc/ig-assembly-eval. Curated assembly sequences are on github within the “curated_IGH” folder, and is also available on zenodo at https://doi.org/10.5281/zenodo.13125505.

### Ethics approval and consent to participate

Ethical approval is not applicable for this article.

### Consent for publication

Not applicable.

### Authors’ contributions

All authors contributed to the design of the study and to writing the paper. YZ developed the bioinformatics tool, with help from other authors. YZ, YS, and AB analyzed the data.

## Acknowledgements

We thank Katalin Voss, Eric Engelbrecht, and the rest of the Pennell, Edge, and Mooney lab groups for helpful discussion of this work.

## Declaration of generative AI and AI-assisted technologies

During the preparation of this work the authors used ChatGPT in order to improve the manuscript’s language. After using this tool, the authors reviewed and edited the content as needed and take full responsibility for the content of the published article.

## Supplementary Information

Document S1. Figures S1–S9 and Note S1

Table S2. Excel file containing additional data too large to fit in a PDF, related to summary of the 74 species’ IG assembly evaluation

Document S2. Results of the 74 species’ IGH loci assembly

Document S3. Results of the 57 species’ IGK loci assembly

Document S4. Results of the 61 species’ IGL loci assembly

## Supplementary Information

### Simulation

**Supplementary Figure 1.**
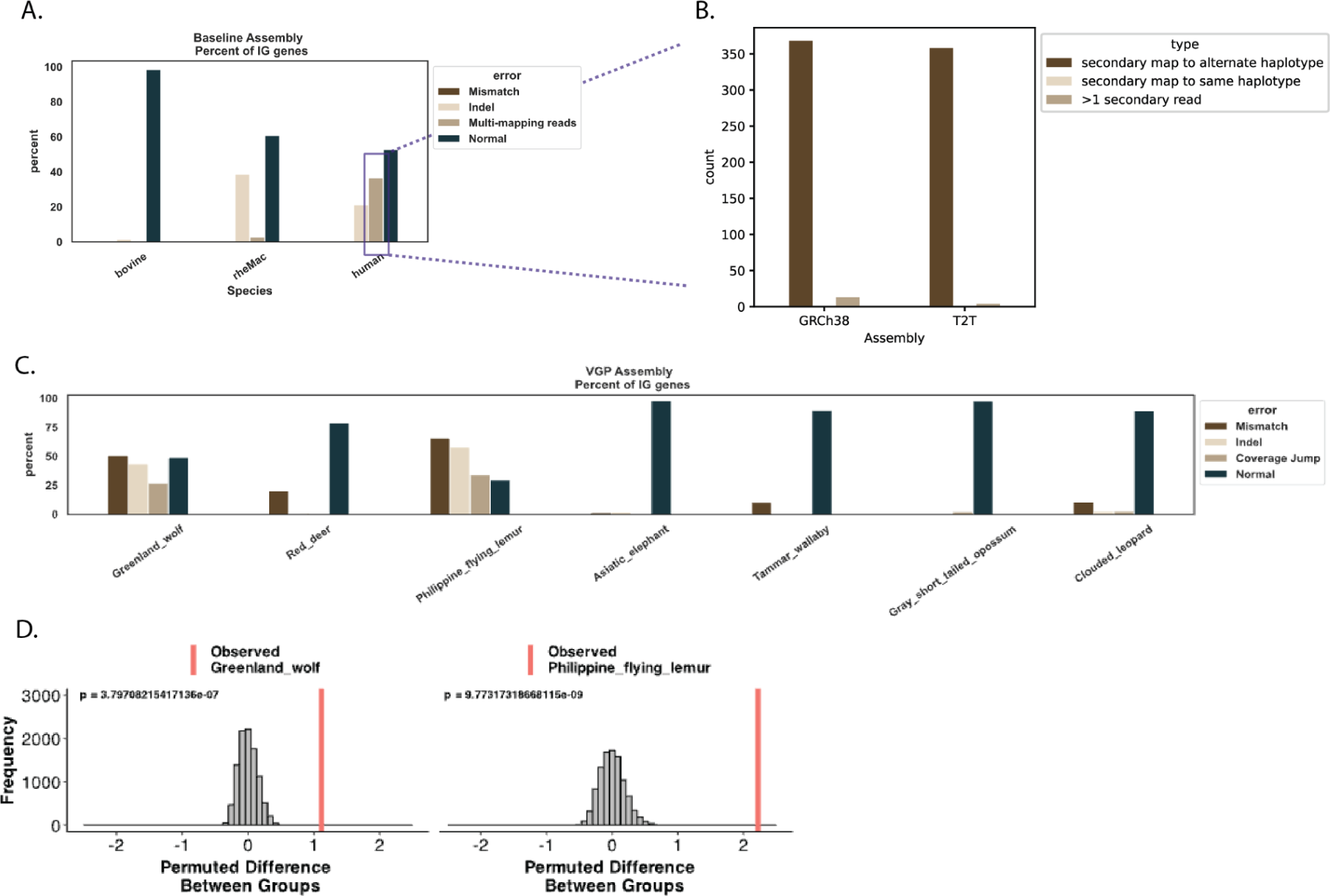
Benchmarking Assembly Errors Using Simulated HiFi reads from Human, Bovine, and Rhesus assembly. All genomes are diploid, with the human genome created as a combination of the GRCh38 and T2T assemblies. A. Percent of IG genes displaying each type of alignment error within human, bovine, and rhesus assemblies using simulated reads. B. Multi-mapped reads in the human assembly were found due to low heterozygosity between the two haplotypes. C. Percent of IG genes displaying each type of alignment error within eight VGP assemblies using real sequencing reads. D. A permutation test was used to evaluate whether the observed errors in the VGP genomes were consistent with those of the reference species with simulated reads.

### Breaks in Coverage

For breaks with consecutive ’N’ reference sequences, which represent unknown bases, these are often linked to complex genomic structures or repetitive elements. This includes cases with one or two reads mapped, where reads may still align if the surrounding sequences provide sufficient context for alignment algorithms to operate despite uncertainties. It also covers breaks with zero reads aligned, referred as gaps in this paper, indicating completely unsequenced or unassembled regions.

In contrast, for non-’N’ reference sequences, breaks with fewer than two (non zero) reads mapped suggest regions with sparse sequencing data, potentially highlighting unsupported sequences that might reflect errors in the assembly. Breaks with no reads aligned point to more significant data gaps, where the reference sequence is known but completely lacks sequencing support, providing stronger evidence of possible assembly inaccuracies.

**Supplementary Figure 2.**
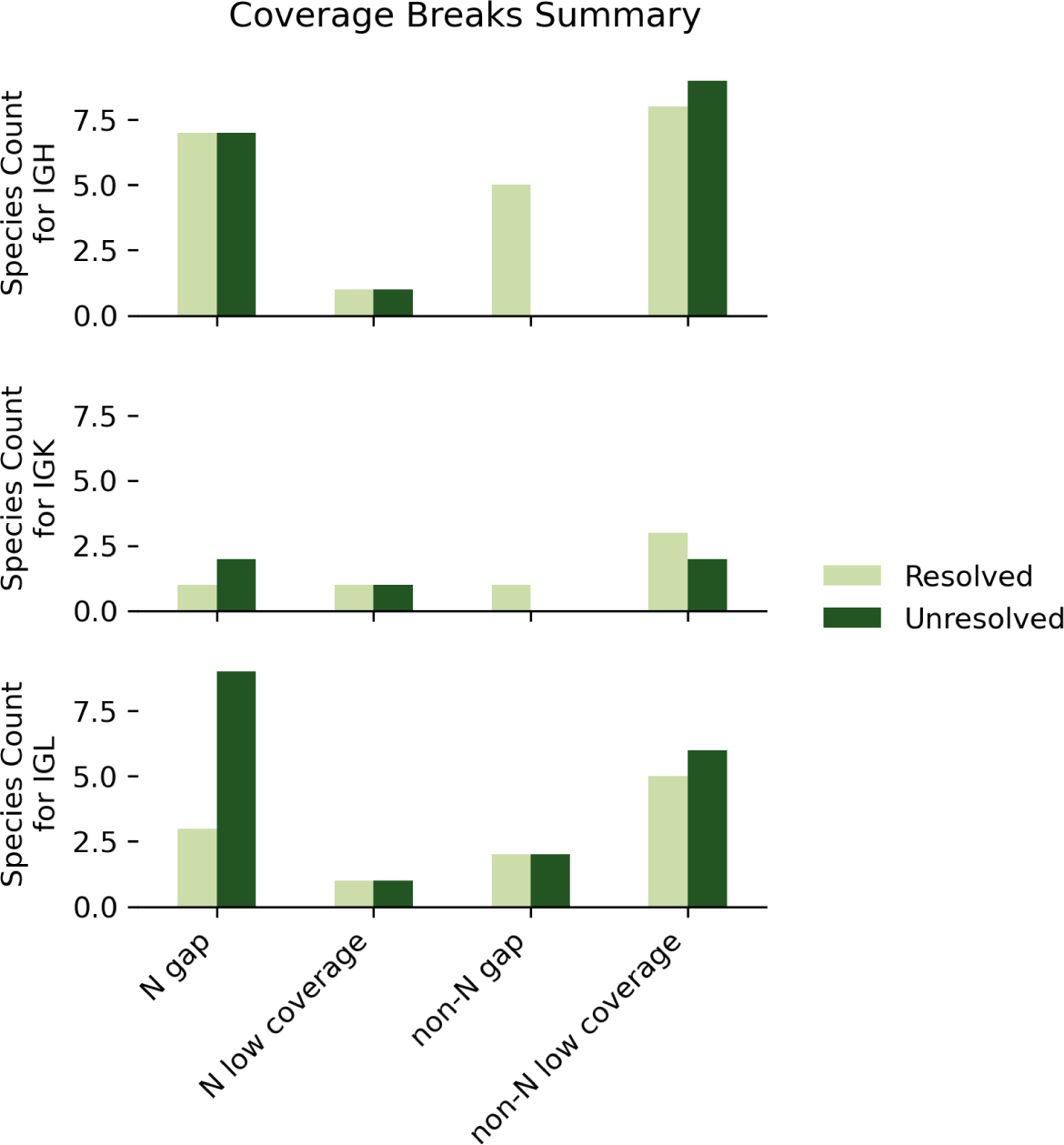
Breakdown of Assembly Break Types Across Species in IGH, IGK and IGL loci. This figure displays the frequency of four distinct types of assembly breaks across various species within the IGH, IGK, and IGL loci. Break types are categorized as follows: N low coverage (fewer than two reads mapped in regions with consecutive ’N’ reference sequences); N gap (zero reads mapped in regions with consecutive ’N’ reference sequences); non-’N’ low coverage (fewer than two reads mapped in regions with non-’N’ reference sequence); and non-N gap (no reads mapped in regions with non-’N’ reference sequence). Each bar is color-coded to indicate whether the species’ assemblies are haplotype-resolved.

### Case Studies Supplementary

**Supplementary Figure 3.**
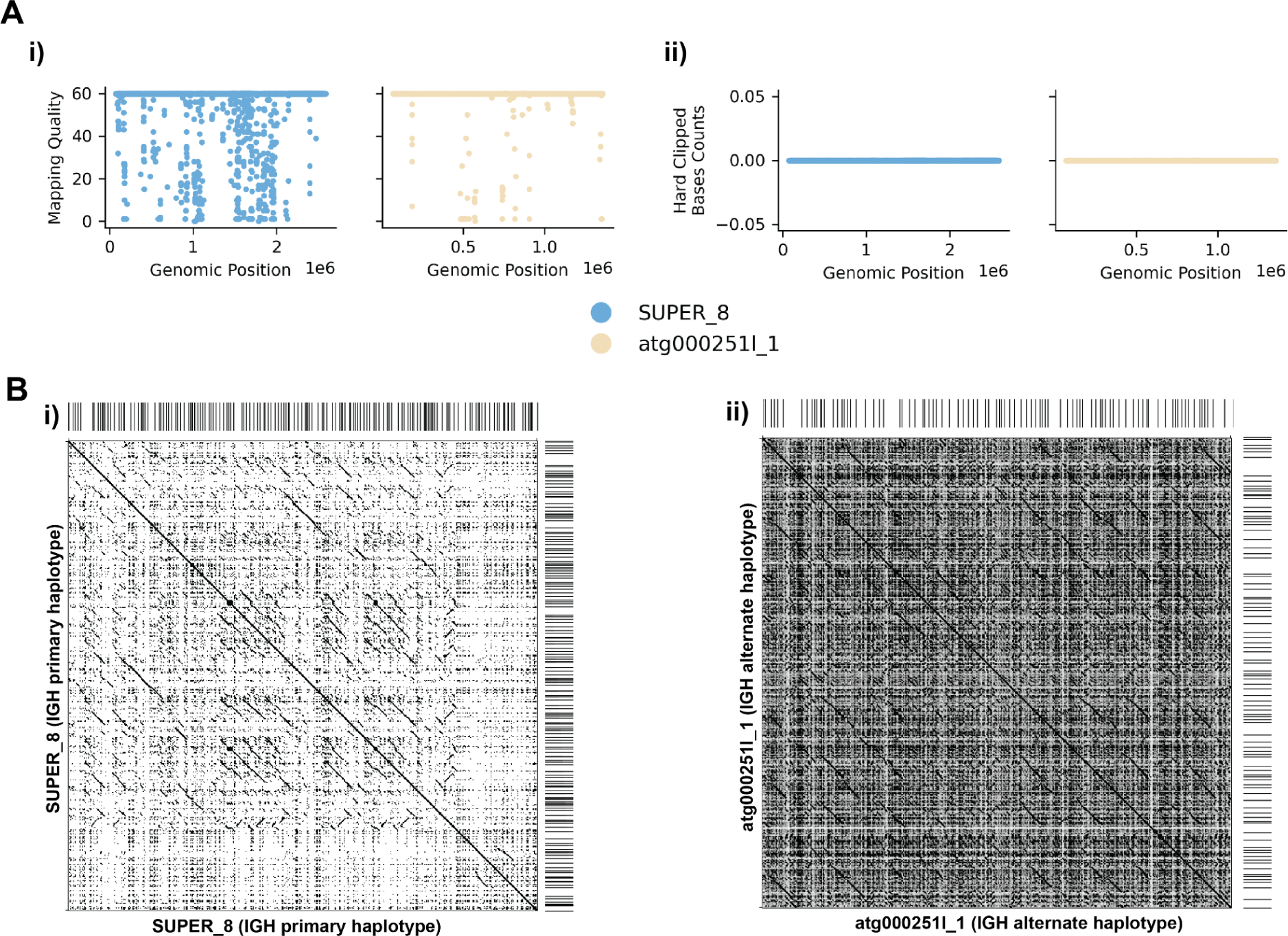
Additional Analysis of IGH Locus Assembly Errors in Greenland wolf (*C. lupus*) individual 1. A. Summary statistics of the read alignment situation are depicted, showing i) mapping quality across IGH loci for both haplotypes, with blue representing the primary assembly and yellow the alternate. ii) count of hard clipped bases. B. Dotplots comparing gene locations and alignments are shown for i) primary vs primary and ii) alternate vs alternate haplotypes.

**Supplementary Figure 4.**
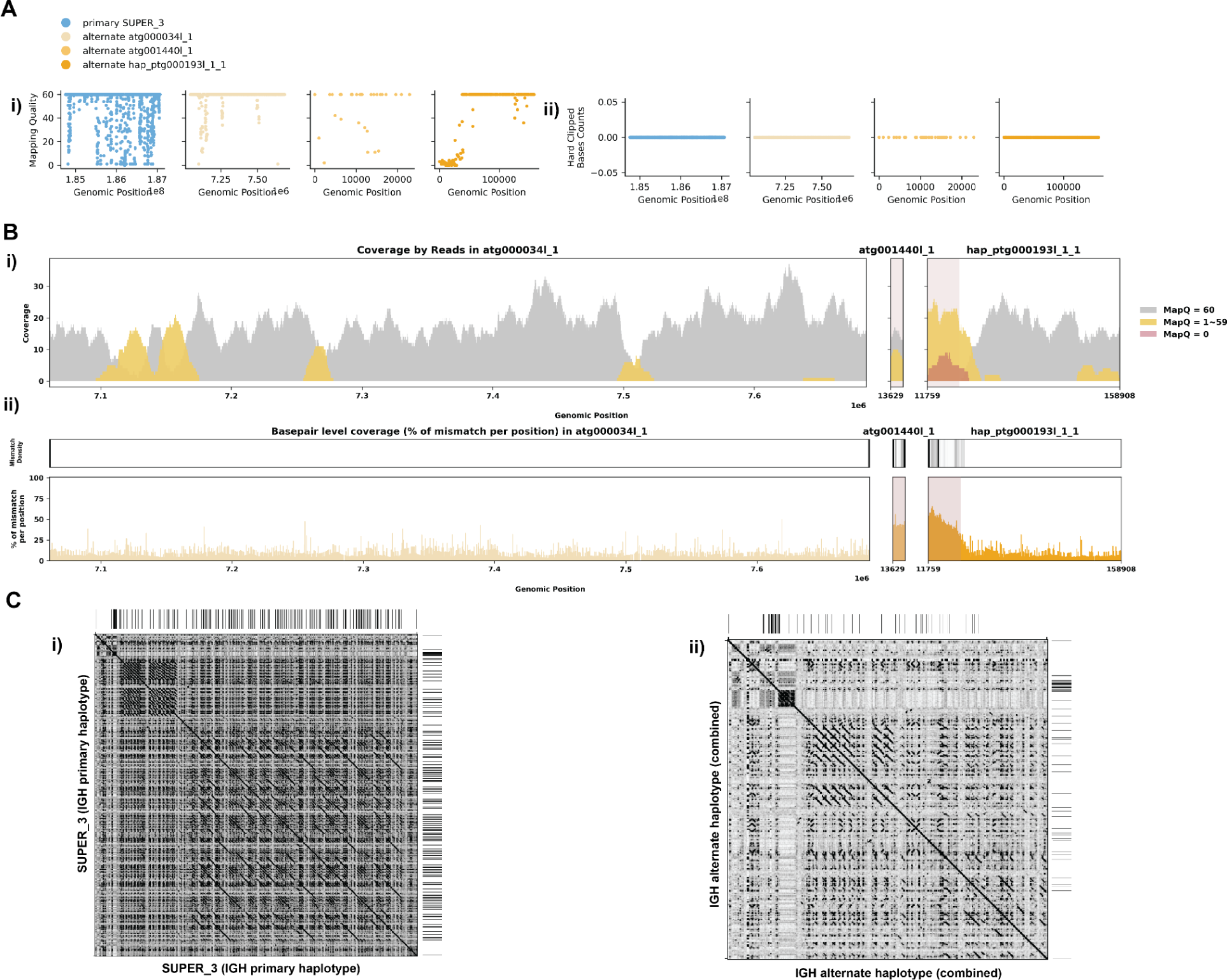
Additional Analysis of IGH Locus Assembly Errors in Philippine Flying Lemur (*C. volans).* A. Summary statistics of the read alignment situation are depicted, showing i) mapping quality across IGH loci for both haplotypes, with blue representing the primary assembly and yellow the alternate. ii) count of hard clipped bases. B. A detailed analysis of alignment mismatch in the alternate IGH haplotype includes i) read coverage across the entire IGH loci, color-coded by mapping quality, and ii) *basepair-oriented* mismatch rate, a heatmap above indicating the frequency of high mismatch rate base pairs, with darker colors denoting more frequent occurrences. Light red highlights positions covered by ≥5 reads with an error rate >1%, and purple bars indicate coverage breaks (coverage ≤2). C. Dotplots comparing gene locations and alignments are shown for i) primary vs primary and ii) alternate vs alternate haplotypes.

**Supplementary Figure 5.**
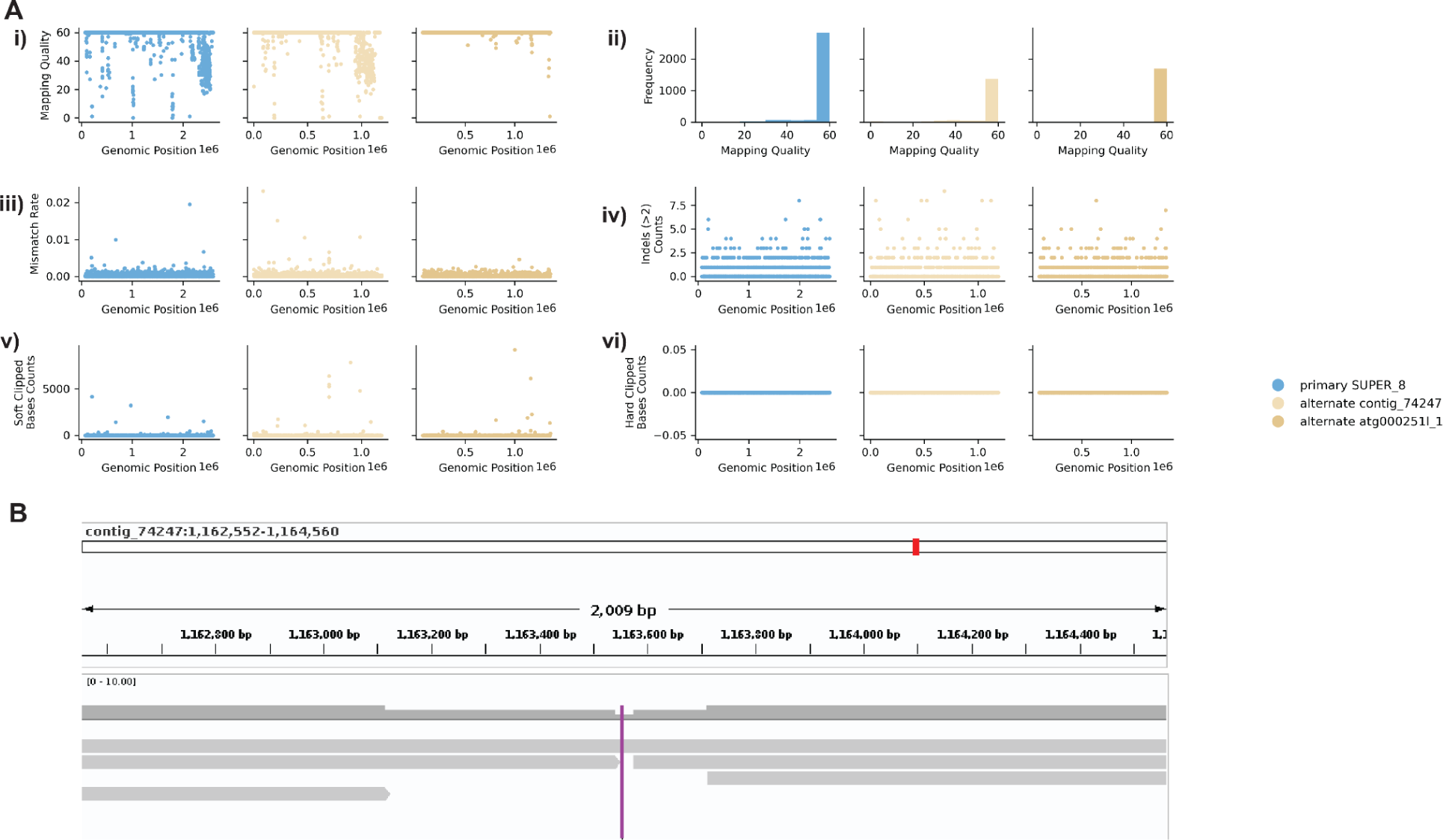
Additional Reassembly Analysis of IGH Locus Assembly Errors in *C. lupus* individual 1. A. Summary statistics of the read alignment are significantly improved. Blue represents the primary assembly and yellow the alternate. i) The plots illustrate the read mapping quality across the IGH loci, ii) read mapping quality distribution, iii) mismatch rates of reads, and iv) number of indels (consecutive length > 2bps), v) count of soft clipped bases, vi) count of hard clipped bases, for both haplotypes across IGH loci. B. IGV screenshot of the low coverage region in contig “74247”, purple indicate where the break in coverage is.

**Supplementary Figure 6.**
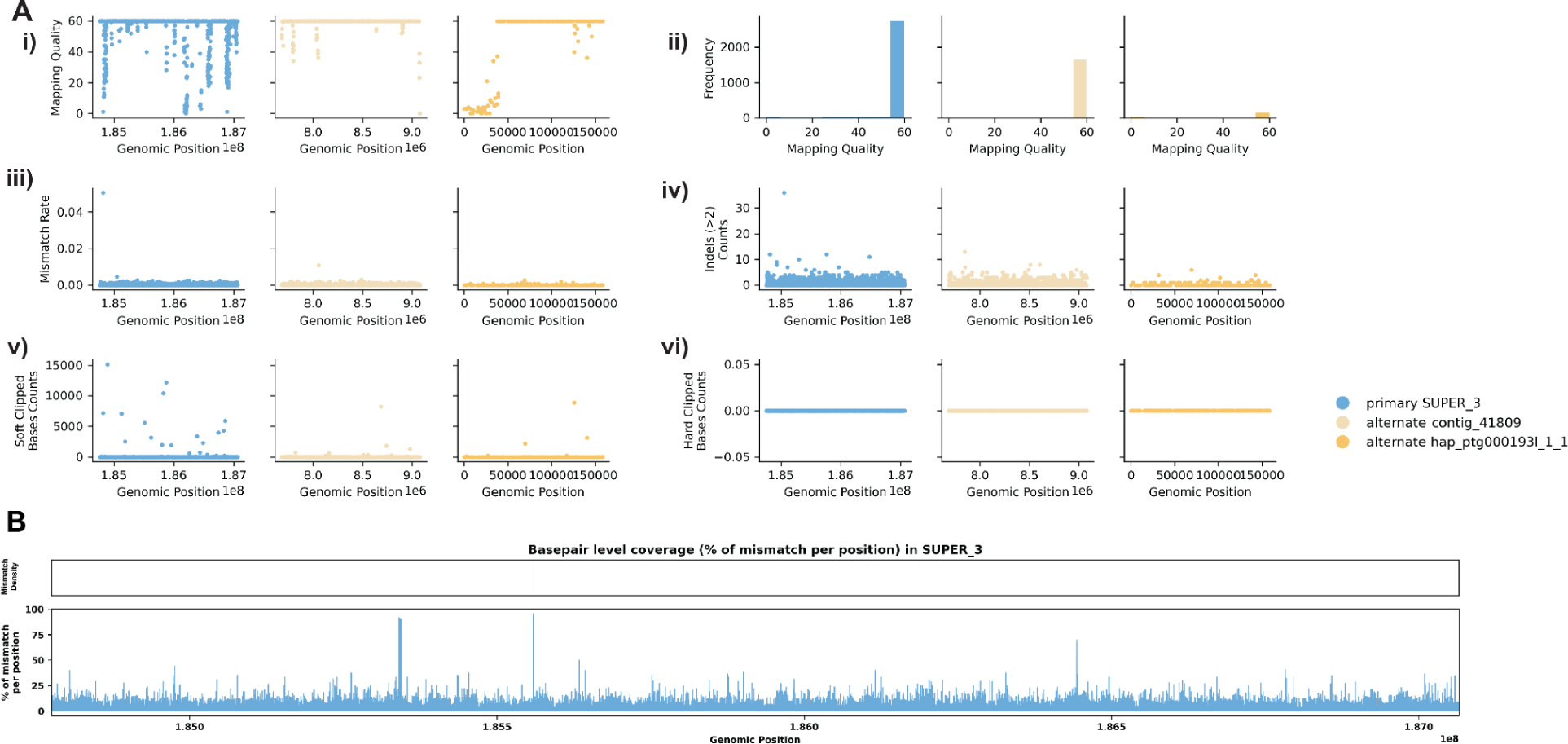
Additional Reassembly Analysis of IGH Locus Assembly Errors in *C. volans.* A. Summary statistics of the read alignment are significantly improved. Blue represents the primary assembly and yellow the alternate. i) The plots illustrate the read mapping quality across the IGH loci, ii) read mapping quality distribution, iii) mismatch rates of reads, and iv) number of indels (consecutive length > 2bps), v) count of soft clipped bases, vi) count of hard clipped bases, for both haplotypes across IGH loci. B. A detailed analysis of alignment mismatch in the primary IGH haplotype from *basepair-oriented* view, a heatmap above indicating the frequency of high mismatch rate base pairs, with darker colors denoting more frequent occurrences. Purple bars indicate coverage breaks (coverage ≤2).

**Supplementary Figure 7.**
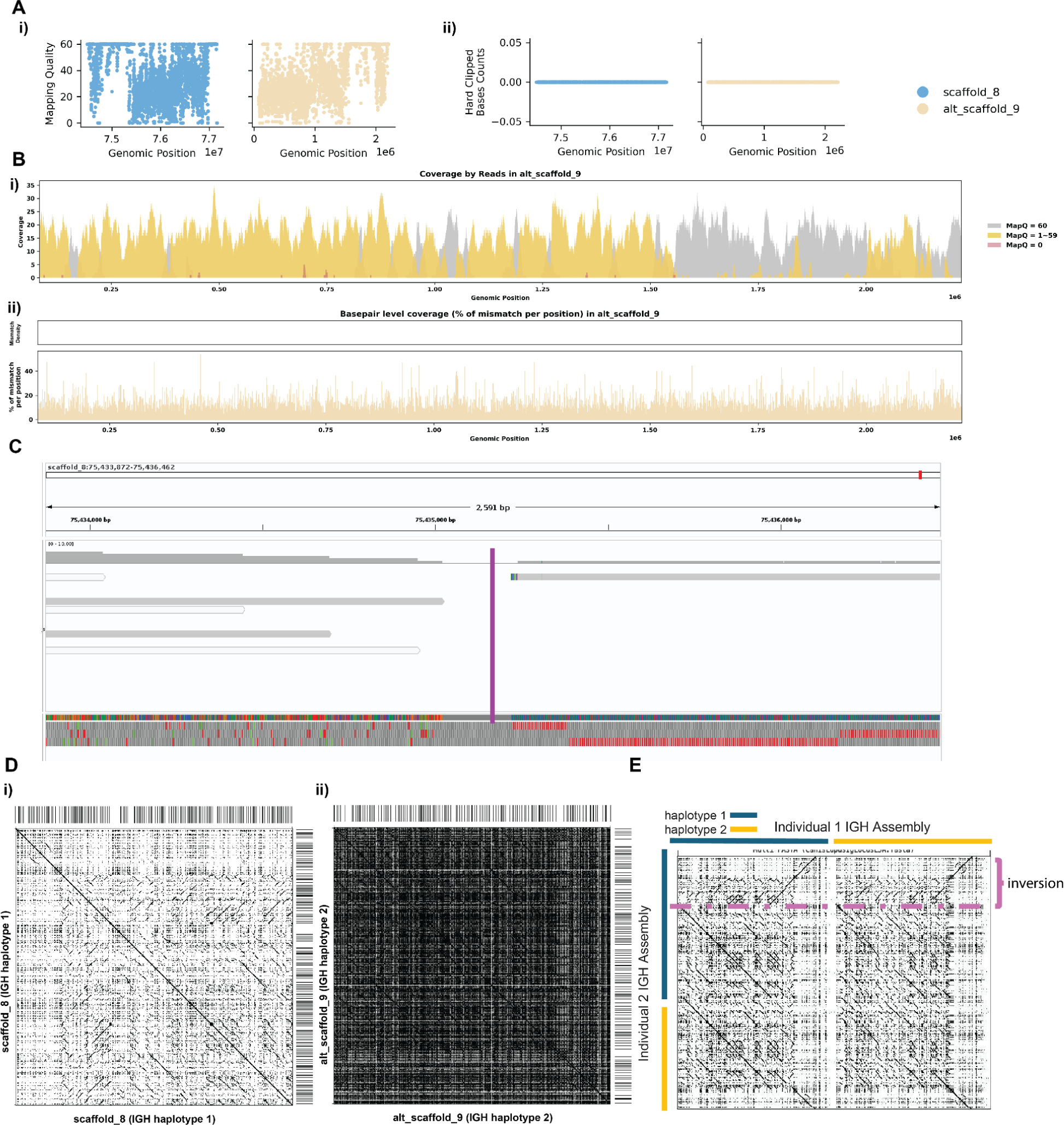
Additional Detailed Analysis of IGH Locus Assembly Errors in Greenland Wolf (*C. lupus*) individual 2. A. Summary statistics of the read alignment situation are depicted, showing i) mapping quality across IGH loci for both haplotypes, with blue representing the primary assembly and yellow the alternate. ii) count of hard clipped bases. B. A detailed analysis of alignment mismatch in the alternate IGH haplotype includes i) read coverage across the entire IGH loci, color-coded by mapping quality, and ii) *basepair-oriented* mismatch rate, a heatmap above indicating the frequency of high mismatch rate base pairs, with darker colors denoting more frequent occurrences. Light red highlights positions covered by ≥5 reads with an error rate >1%, and purple bars indicate coverage breaks (coverage ≤2). C. IGV screenshot of the break in coverage D. Dotplots comparing gene locations and alignments are shown for i) primary vs primary and ii) alternate vs alternate haplotypes. E. Dotplots comparing *C. lupus* individual 1 assembly vs individual 2 assembly. Purple dashed line indicate the inversion observed.

**Supplementary Figure 8.**
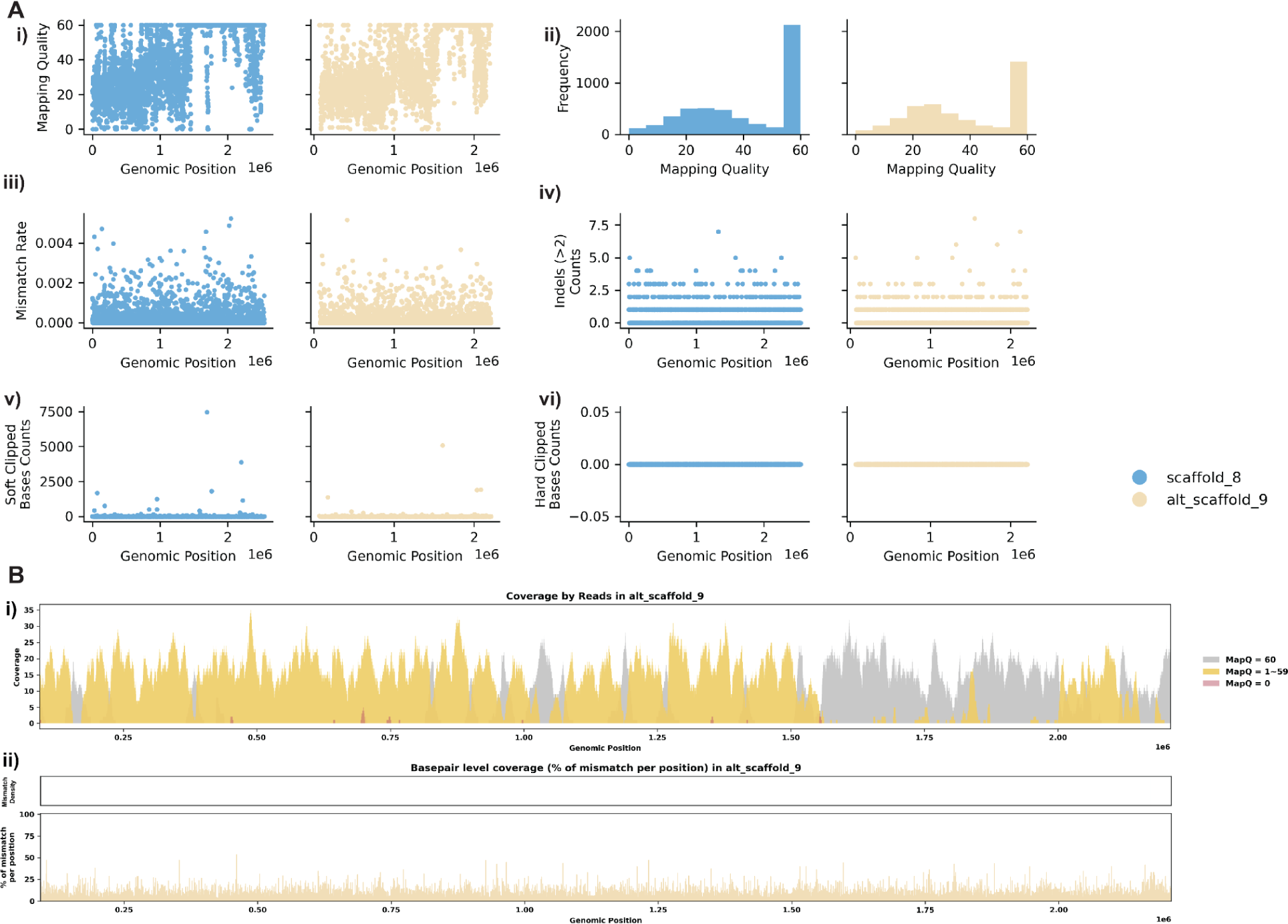
Additional Reassembly Analysis of IGH Locus Assembly Errors in *C. volans.* Individual 2. A. Summary statistics of the read alignment are significantly improved. Blue represents the primary assembly and yellow the alternate. i) The plots illustrate the read mapping quality across the IGH loci, ii) read mapping quality distribution, iii) mismatch rates of reads, and iv) number of indels (consecutive length > 2bps), v) count of soft clipped bases, vi) count of hard clipped bases, for both haplotypes across IGH loci. B. A detailed analysis of alignment mismatch in the alternate IGH haplotype includes i) read coverage across the entire IGH loci, color-coded by mapping quality, and ii) *basepair-oriented* mismatch rate, a heatmap above indicating the frequency of high mismatch rate base pairs, with darker colors denoting more frequent occurrences. Light red highlights positions covered by ≥5 reads with an error rate >1%, and purple bars indicate coverage breaks (coverage ≤2).

### Species Overview

**Supplementary Figure 9.**
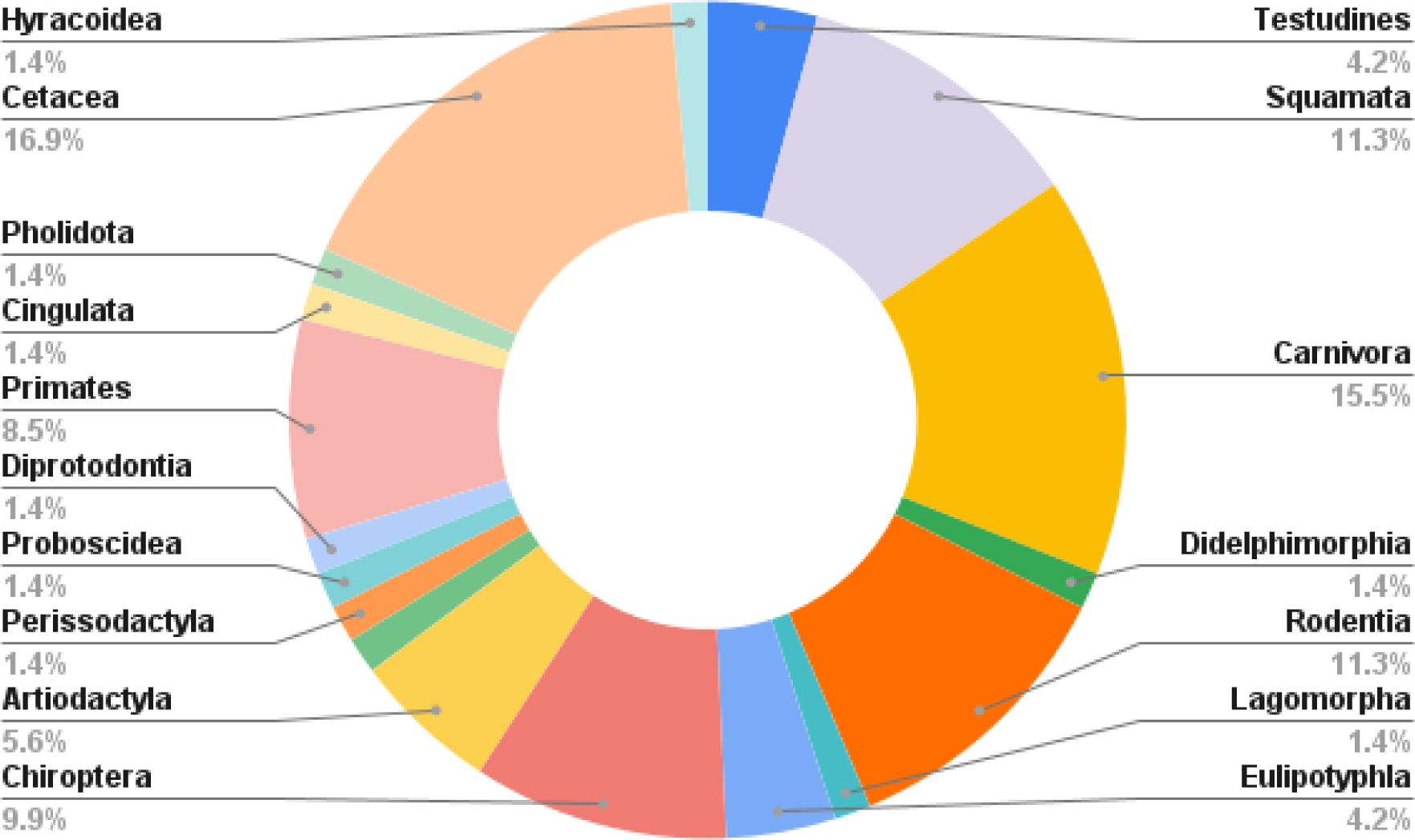
Pie chart summarizing the distribution of species by order.

**Supplementary Note 1**: Comparison to CRAQ and Inspector

We ran CRAQ and Inspector on *C. lupus* individual 1 and *C. lupus* individual 2 to evaluate their effectiveness. In the structural error output BED file provided by Inspector, the IGH loci for both individuals were missing, indicating that Inspector failed to detect structural errors in these regions. Similarly, CRAQ’s results showed that the IGH loci for both individuals did not appear in the CSE (Clip-based Structural Errors) and CSH (Clip-based Structural Heterozygosity) outputs. Although CRAQ did identify these errors in its regional error output file, it classified them incorrectly as small-scale errors. This misclassification creates confusion and demonstrates the limitations of both tools in accurately detecting and categorizing errors in the IG loci. This underscores the necessity of developing CloseRead, as relying solely on existing tools like CRAQ and Inspector would have left us unaware of these critical inaccuracies. CloseRead provides a targeted approach to ensure precise detection and classification of errors, addressing the shortcomings of current tools.

