## Supplementary material for "Assessing Assembly Errors in Immunoglobulin Loci: A Comprehensive Evaluation of Long-read Genome Assemblies Across Vertebrates": Assembly Evaluation Summary

### IGH

| Class | IndividualID | LatinName | CommonName | Source | SourceLink | Haplotype Resolved | Evaluation Result | Special Condition | Curation Status |
|---|---|---|---|---|---|---|---|---|---|
| Mammalia | mApoSyl1 | Apodemus sylvaticus | wood mouse | VGP | o/genomeark-all/A | No | Mismatch; Break in Primary assembly; Abnormal high coverage | alternate locus too short/not found | |
| Mammalia | mBalAcu1 | Balaenoptera_acutorostrata | minke whale | VGP | rg/vgp-all/Balaen | No | Good | Slight mismatch | |
| Mammalia | mCamDro1 | Camelus dromedarius | dromedary | VGP | g/genomeark-all/C | Yes | Good | Slight Mismatch | |
| Mammalia | mCanLor1 | Canis_lupus | Greenland Wolf | VGP | e/s3/genomeark/s | No | Mismatch | Short Alternate Assembly | Missing Contig in Alternate assembly |
| Mammalia | mCanLor2 | Canis_lupus | Greenland Wolf | VGP | e/s3/genomeark/s | Yes | Break in Primary assembly | | false inversion |
| Mammalia | mCerEla1 | Cervus_elaphus | Red Deer | VGP | s3/genomeark/spe | No | Good | Slight Mismatch; Short Alternate Assembly | |
| Mammalia | mChiNiv1 | Chionomys nivalis | European snow vole | VGP | .io/genomeark-all | No | Good | Slight Mismatch; alternate locus too short/not found | |
| Mammalia | mCorTow1.0 | Corynorhinus townsendii | Townsend's Big-eared Bat | CCGP | s/corynorhinus-tov | Yes | Break in Primary + Alternate assembly | 2 IGH; Slight Mismatch | |
| Mammalia | mCynVol1 | Cynocephalus volans | Philippine flying lemur | VGP | genomeark/specie | No | Mismatch | Short Alternate Assembly | Missing Contig in Alternate assembly |
| Mammalia | mDasNov1 | Dasypus novemcinctus | nine-banded armadillo | VGP | /genomeark-all/D | Yes | Break in Primary + Alternate Assembly | | |
| Mammalia | mDelDel1 | Delphinus delphis | saddleback dolphin | VGP | ark.org/vgp-all/De | No | Mismatch | Short Alternate Assembly | |
| Mammalia | mDicBic1 | Diceros bicornis | black rhinoceros | VGP | b.io/genomeark-a | Yes | Mismatch; Break in Primary assembly | | |
| Mammalia | mDipMer1 | Dipodomys merriami | Merriam's Kangaroo Rat | CCGP | cies/dipodomys-m | No | Good | | |
| Mammalia | mEleMax1 | Elephas maximus | Asiatic Elephant | VGP | 3/genomeark/spec | No | Good | Short Alternate Assembly | |
| Mammalia | mEptNil1 | Eptesicus nilssonii | northern bat | VGP | ark.org/vgp-all/Ept | No | Mismatch; Break in Alternate Assembly | 2 IGH | |
| Mammalia | mEriEur2 | Erinaceus europaeus | western European hedgehog | VGP | k.org/vgp-all/Erina | No | Mismatch; Break in Primary assembly | alternate locus too short/not found | |
| Mammalia | mEscRob2 | Eschrichtius robustus | grey whale | VGP | k.org/vgp-all/Esch | No | Good | | |
| Mammalia | mEubGla1 | Eubalaena glacialis | North Atlantic right whale | VGP | rk.org/vgp-all/Eub | Yes | Good | | |
| Mammalia | mGloMel1 | Globicephala melas | long-finned pilot whale | VGP | rk.org/vgp-all/Glol | No | Good | Short Alternate Assembly | |
| Mammalia | mGorGor1 | Gorilla_gorilla | Gorilla | T2T Primate | /s3/genomeark/sp | Yes | Good | | |
| Mammalia | mHetBru1 | Heterohyrax brucei | Yellow-spotted hyrax | VGP | .io/genomeark-all/ | No | Good | Short Alternate Assembly | |
| Mammalia | mHipAmp2 | Hippopotamus amphibius kiboko | hippopotamus | VGP | ark-curated-asser | Yes | Good | Slight Mismatch; Most MapQ 0 reads; Short Alternate Assembly | |
| Mammalia | mHypAmp2 | Hyperoodon ampullatus | northern bottlenose whale | VGP | .org/vgp-all/Hyper | No | Good | Short Alternate Assembly | |
| Mammalia | mLagAlb1 | Lagenorhynchus albirostris | white-beaked dolphin | VGP | rg/vgp-all/Lageno | No | Mismatch | alternate locus too short/not found | |
| Mammalia | mLemCat1 | Lemur catta | Ring-tailed lemur | VGP | hub.io/genomeark | No | Break in Primary assembly (but also is the beginning of chrom) | | |
| Mammalia | mLynRuf1 | Lynx rufus | Bobcat | CCGP | oject.org/species/ | No | Good | alternate locus too short/not found | |
| Mammalia | mMacEug1 | Macropus eugenii | tammar wallaby | VGP | .io/genomeark-all | No | Mismatch; Break in Primary assembly | alternate locus too short/not found | |
| Mammalia | mManPen7 | Manis pentadactyla | Chinese pangolin | VGP | io/genomeark-all/ | Yes | Good | Slight Mismatch | |
| Mammalia | mMarMar1 | Martes martes | European pine marten | VGP | eark.org/vgp-all/M | No | Mismatch | alternate locus too short/not found | |
| Mammalia | mMelMel3 | Meles meles | European badger | VGP | hub.io/genomeark- | Yes | Good | Slight Mismatch | |
| Mammalia | mMesDen1 | Mesoplodon densirostris | Blainville's beaked whale | VGP | org/vgp-all/Mesop | No | Good | | |
| Mammalia | mMicCal1.0 | Microtus californicus | California Vole | CCGP | /species/microtus- | Yes | Break in Primary + Alternate Assembly | | |
| Mammalia | mMicMin1 | Micromys minutus | European harvest mouse | VGP | ark.org/vgp-all/Mic | No | Mismatch | alternate locus too short/not found | |
| Mammalia | mMirAng1 | Mirounga angustirostris | Northern Elephant Seal | CCGP | es/mirounga-angu | Yes | Break in Alternate assembly | | |
| Mammalia | mMonDom1 | Monodelphis_domestica | gray short-tailed opossum | VGP | nomeark/species/ | No | Mismatch | | |
| Mammalia | mMunRee1 | Muntiacus reevesi | Reeves' muntjac | VGP | ark.org/vgp-all/Mu | No | Good | | |
| Mammalia | mMusAve1 | Muscardinus avellanarius | hazel dormouse | VGP | org/vgp-all/Musca | No | Good | alternate locus too short/not found | |
| Mammalia | mMusLut2 | Mustela lutreola | European mink | VGP | .org/genomeark-a | No | Good | | |
| Mammalia | mMusNiv1 | Mustela nivalis | Least weasel | VGP | .org/genomeark-a | Yes | Break in Alternate Assembly | | |
| Mammalia | mMyoDau2 | Myotis daubentonii | Daubenton's bat | VGP | ark.org/vgp-all/My | No | Mismatch | 2 IGH | |
| Mammalia | mMyoYum1.0 | Myotis yumanensis | Yuma myotis | CCGP | g/species/myotis-y | Yes | Break in Primary + Alternate Assembly | 2 IGH | |
| Mammalia | mNeoNeb1 | Neofelis_nebulosa | Clouded Leopard | VGP | 3/genomeark/spec | No | Mismatch | Abnormal 2x coverage region; Short Alternate Assembly | |
| Mammalia | mNycCou1 | Nycticebus coucang | slow loris | VGP | io/genomeark-all/ | No | Break in Alternate assembly | Short Primary Assembly | |
| Mammalia | mOrcOrc1 | Orcinus orca | killer whale | VGP | hub.io/genomeark- | No | Good | Short Alternate Assembly | |
| Mammalia | mOryCun1 | Oryctolagus cuniculus | rabbit | VGP | k.org/vgp-all/Oryc | No | Mismatch | alternate locus too short/not found | |
| Mammalia | mPanPan1 | Pan paniscus | Bonobo | T2T Primate | /s3/genomeark/sp | Yes | Good | | |
| Mammalia | mPerMan1 | Peromyscus maniculatus | deer mouse | CCGP | species/peromysc | No | Break in Primary Assembly | Slight Mismatch | |
| Mammalia | mPhoPho1 | Phocoena phocoena | harbor porpoise | VGP | k.org/vgp-all/Phoc | No | Mismatch | | |
| Mammalia | mPipPyg2 | Pipistrellus pygmaeus | soprano pipistrelle | VGP | k.org/vgp-all/Pipis | No | Good | Slight Mismatch | |
| Mammalia | mPleAur1 | Plecotus auritus | brown big-eared bat | VGP | eark.org/vgp-all/Pl | No | Mismatch; Break in Primary Assembly | 2 IGH; alternate locus too short/not found | |
| Mammalia | mPonAbe1 | Pongo_abelii | Sumatran orangutan | T2T Primate | /s3/genomeark/sp | Yes | Good | | |
| Mammalia | mPonPyg2 | Pongo_pygmaeus | Bornean orangutan | T2T Primate | 3/genomeark/spec | Yes | Good | | |
| Mammalia | mPseCra1 | Pseudorca crassidens | false killer whale | VGP | k.org/vgp-all/Pseu | Yes | Good | | |
| Mammalia | mPumCon1.1 | Puma concolor | Mountain Lion | CCGP | org/species/puma- | Yes | Good | | |
| Mammalia | mSorAra2/1 | Sorex araneus | Common shrew | VGP | ub.io/genomeark-a | No | Mismatch | Abnormal hig coverage; Short Primary Assembly | |
| Mammalia | mSteCoe1 | Stenella coeruleoalba | striped dolphin | VGP | k.org/vgp-all/Sten | No | Mismatch | | |
| Mammalia | mTalEur1 | Talpa europaea | European mole | VGP | eark.org/vgp-all/Ta | No | Mismatch; Break in Primary assembly | alternate locus too short/not found | |
| Mammalia | mThoBot1 | Thomomys bottae | Botta's pocket gopher | CCGP | g/species/thomom | No | Good | | |
| Mammalia | mUrsAme1 | Ursus americanus | American black bear | CCGP | gov/datasets/gen | No | Good | Slight Mismatch; alternate locus too short/not found | |
| Mammalia | mUrsArc2 | Ursus arctos | brown bear | NCBI | gov/datasets/gen | No | Good | 2 IGH; Slight Mismatch | |
| Mammalia | mVesMur1 | Vespertilio murinus | particolored bat | VGP | rk.org/vgp-all/Ves | No | Mismatch | 2 IGH | |
| Crocodilia | rAllMis2 | Alligator mississippiensis | American alligator | VGP | genomeark/specie | Yes | Break in Primary Assembly | | |
| Testudines | rCarCar2 | Caretta caretta | Loggerhead turtle | VGP | eark.org/vgp-all/C | Yes | Break in Alternate Assembly; Double Coverage in Primary Assembly | Abnormal 2x coverage region | |
| Testudines | rEmyOrb1 | Emys orbicularis | European pond turtle | VGP | ark.org/vgp-all/En | Yes | Break in Primary + Alternate Assembly | | |
| Testudines | rEryReg1 | Erythrolamprus reginae | royal ground snake | VGP | .org/vgp-all/Erythr | No | Mismatch; Break in Primary Assembly | | |
| Squamata | rLiaOli1 | Liasis olivaceus | olive python | VGP | eark.org/vgp-all/Li | Yes | Break in Primary assembly | Slight Mismatch | |
| Squamata | rLiaOli2 | Liasis olivaceus | olive python | VGP | eark.org/vgp-all/Li | Yes | Break in Primary assembly | Slight Mismatch | |
| Testudines | rMalTer1 | Malaclemys terrapin | diamondback terrapin | VGP | rk.org/vgp-all/Mala | Yes | Break in Primary assembly | Slight Mismatch | |
| Squamata | rPodCre2 | Podarcis cretensis | cretan wall lizard | VGP | ark.org/vgp-all/Poc | No | Break in Primary assembly | Slight Mismatch; alternate locus too short/not found | |

| Class | IndividualID | LatinName | CommonName | Source | SourceLink | Haplotype Resolved | Evaluation Result | Special Condition | Curation Status |
|---|---|---|---|---|---|---|---|---|---|
| Squamata | rPodRaf1 | Podarcis raffonei | Aeolian wall lizard | VGP | ark.org/vgp-all/Po | No | Good | alternate locus too short/not found | |
| Squamata | rRhiFlo1 | Rhineura floridana | Florida worm lizard | VGP | ark.org/vgp-all/Rhi | Yes | Break in Primary + Alternate Assembly | Slight Mismatch; Short Alternate Assembly | |
| Squamata | rVipLat1 | Vipera latastei | snub-nosed viper | VGP | eark.org/vgp-all/V | No | Mismatch; Break in Primary Assembly | alternate locus too short/not found | |
| Squamata | rVipUrs1 | Vipera ursinii | Hungarian meadow viper | VGP | neark.org/vgp-all/\ | No | Mismatch; Break in Primary Assembly | alternate locus too short/not found | |
| Squamata | rZooViv1 | Zootoca vivipara | common lizard | VGP | ark.org/vgp-all/Zo | No | Mismatch | alternate locus too short/not found | |

### IGK

| Class | IndividualID | LatinName | CommonName | Source | SourceLink | Haplotype Resolved | Evaluation Result | Special Condition |
|---|---|---|---|---|---|---|---|---|
| Mammalia | mApoSyl1 | Apodemus sylvaticus | wood mouse | VGP | o/genomeark-all/A | No | Mismatch; Break in Primary + Alternate Assembly | Short Alternate Assembly |
| Mammalia | mBalAcu1 | Balaenoptera_acutorostrata | minke whale | VGP | rg/vgp-all/Balaen | No | Good | alternate locus too short/not found |
| Mammalia | mCamDro1 | Camelus dromedarius | dromedary | VGP | g/genomeark-all/C | Yes | Good | |
| Mammalia | mCanLor1 | Canis_lupus | Greenland Wolf | VGP | e/s3/genomeark/s | No | Good | |
| Mammalia | mCanLor2 | Canis_lupus | Greenland Wolf | VGP | e/s3/genomeark/s | Yes | Good | All MapQ 0 reads |
| Mammalia | mCerEla1 | Cervus_elaphus | Red Deer | VGP | s3/genomeark/spe | No | Good | |
| Mammalia | mChiNiv1 | Chionomys nivalis | European snow vole | VGP | .io/genomeark-all | No | Mismatch | Short Alternate Assembly |
| Mammalia | mDasNov1 | Dasypus novemcinctus | nine-banded armadillo | VGP | /genomeark-all/D | Yes | Mismatch; Break in Primary + Alternate Assembly | |
| Mammalia | mDelDel1 | Delphinus delphis | saddleback dolphin | VGP | ark.org/vgp-all/De | No | Good | |
| Mammalia | mDicBic1 | Diceros bicornis | black rhinoceros | VGP | b.io/genomeark-a | Yes | Good | |
| Mammalia | mDipMer1 | Dipodomys merriami | Merriam's Kangaroo Rat | CCGP | cies/dipodomys-m | No | Good | 3 IGK |
| Mammalia | mEleMax1 | Elephas maximus | Asiatic Elephant | VGP | 3/genomeark/spec | No | Good | |
| Mammalia | mEriEur2 | Erinaceus europaeus | western European hedgehog | VGP | k.org/vgp-all/Erina | No | Good | alternate locus too short/not found |
| Mammalia | mEscRob2 | Eschrichtius robustus | grey whale | VGP | k.org/vgp-all/Esch | No | Good | |
| Mammalia | mEubGla1 | Eubalaena glacialis | North Atlantic right whale | VGP | rk.org/vgp-all/Eub | Yes | Good | |
| Mammalia | mGloMel1 | Globicephala melas | long-finned pilot whale | VGP | rk.org/vgp-all/Glol | No | Good | alternate locus too short/not found |
| Mammalia | mGorGor1 | Gorilla_gorilla | Gorilla | T2T Primate | /s3/genomeark/sp | Yes | Good | |
| Mammalia | mHetBru1 | Heterohyrax brucei | Yellow-spotted hyrax | VGP | .io/genomeark-all/ | No | Good | alternate locus too short/not found |
| Mammalia | mHipAmp2 | Hippopotamus amphibius kiboko | hippopotamus | VGP | ark-curated-asser | Yes | Good | Slight Mismatch |
| Mammalia | mHypAmp2 | Hyperoodon ampullatus | northern bottlenose whale | VGP | .org/vgp-all/Hyper | No | Mismatch | |
| Mammalia | mLagAlb1 | Lagenorhynchus albirostris | white-beaked dolphin | VGP | rg/vgp-all/Lageno | No | Good | |
| Mammalia | mLemCat1 | Lemur catta | Ring-tailed lemur | VGP | hub.io/genomeark | No | Good | |
| Mammalia | mLynRuf1 | Lynx rufus | Bobcat | CCGP | oject.org/species/ | No | Good | Short Alternate Assembly |
| Mammalia | mMacEug1 | Macropus eugenii | tammar wallaby | VGP | .io/genomeark-all | No | Mismatch | alternate locus too short/not found |
| Mammalia | mManPen7 | Manis pentadactyla | Chinese pangolin | VGP | io/genomeark-all/ | Yes | Break in Alternate Assembly | |
| Mammalia | mMarMar1 | Martes martes | European pine marten | VGP | eark.org/vgp-all/M | No | Mismatch | alternate locus too short/not found |
| Mammalia | mMelMel3 | Meles meles | European badger | VGP | hub.io/genomeark | Yes | Good | All MapQ 0 reads |
| Mammalia | mMesDen1 | Mesoplodon densirostris | Blainville's beaked whale | VGP | org/vgp-all/Mesop | No | Good | |
| Mammalia | mMicCal1.0 | Microtus californicus | California Vole | CCGP | /species/microtus- | Yes | Good | |
| Mammalia | mMicMin1 | Micromys minutus | European harvest mouse | VGP | ark.org/vgp-all/Mic | No | Mismatch | alternate locus too short/not found |
| Mammalia | mMirAng1 | Mirounga angustirostris | Northern Elephant Seal | CCGP | es/mirounga-angu | Yes | Good | All MapQ 0 reads |
| Mammalia | mMonDom1 | Monodelphis_domestica | gray short-tailed opossum | VGP | enomeark/species/ | No | Mismatch; Break in Primary Assembly | alternate locus too short/not found |
| Mammalia | mMunRee1 | Muntiacus reevesi | Reeves' muntjac | VGP | ark.org/vgp-all/Mu | No | Good | |
| Mammalia | mMusAve1 | Muscardinus avellanarius | hazel dormouse | VGP | org/vgp-all/Musca | No | Good | Short Alternate Assembly |
| Mammalia | mMusLut2 | Mustela lutreola | European mink | VGP | .org/genomeark-a | No | Good | Short Alternate Assembly |
| Mammalia | mMusNiv1 | Mustela nivalis | Least weasel | VGP | .org/genomeark-a | Yes | Break in Alternate Assembly | Most MapQ 0 reads |
| Mammalia | mNeoNeb1 | Neofelis_nebulosa | Clouded Leopard | VGP | 3/genomeark/spec | No | Good | |
| Mammalia | mNycCou1 | Nycticebus coucang | slow loris | VGP | io/genomeark-all/ | No | Good | |
| Mammalia | mOrcOrc1 | Orcinus orca | killer whale | VGP | hub.io/genomeark- | No | Good | alternate locus too short/not found |
| Mammalia | mOryCun1 | Oryctolagus cuniculus | rabbit | VGP | .org/vgp-all/Oryc | No | Mismatch | alternate locus too short/not found; abnormal 2x coverage |
| Mammalia | mPanPan1 | Pan paniscus | Bonobo | T2T Primate | /s3/genomeark/sp | Yes | Good | |
| Mammalia | mPerMan1 | Peromyscus maniculatus | deer mouse | CCGP | species/peromyscu | No | Break in Alternate Assembly | Short Primary Assembly |
| Mammalia | mPhoPho1 | Phocoena phocoena | harbor porpoise | VGP | rk.org/vgp-all/Phoc | No | Good | |
| Mammalia | mPonAbe1 | Pongo_abelii | Sumatran orangutan | T2T Primate | e/s3/genomeark/sp | Yes | Good | |
| Mammalia | mPonPyg2 | Pongo_pygmaeus | Bornean orangutan | T2T Primate | 3/genomeark/spec | Yes | Good | |
| Mammalia | mPseCra1 | Pseudorca crassidens | false killer whale | VGP | .org/vgp-all/Pseu | Yes | Good | |
| Mammalia | mPumCon1.1 | Puma concolor | Mountain Lion | CCGP | org/species/puma- | Yes | Good | |
| Mammalia | mSorAra2/1 | Sorex araneus | Common shrew | VGP | ub.io/genomeark-a | No | Mismatch; Break in Primary Assembly | HUGE BREAK , Short Alternate Assembly |
| Mammalia | mSteCoe1 | Stenella coeruleoalba | striped dolphin | VGP | k.org/vgp-all/Sten | No | Good | |
| Mammalia | mTalEur1 | Talpa europaea | European mole | VGP | eark.org/vgp-all/Ta | No | Good | |
| Mammalia | mThoBot1 | Thomomys bottae | Botta's pocket gopher | CCGP | g/species/thomom | No | Good | Short Primary Assembly |
| Mammalia | mUrsAme1 | Ursus americanus | American black bear | CCGP | gov/datasets/gen | No | Good | alternate locus too short/not found |
| Mammalia | mUrsArc2 | Ursus arctos | brown bear | NCBI | gov/datasets/gen | No | Good | |
| Crocodilia | rAllMis2 | Alligator mississippiensis | American alligator | VGP | genomeark/specie | Yes | Good | Most MapQ 0 reads |
| Testudines | rCarCar2 | Caretta caretta | Loggerhead turtle | VGP | eark.org/vgp-all/C | Yes | Good | |
| Testudines | rEmyOrb1 | Emys orbicularis | European pond turtle | VGP | ark.org/vgp-all/Er | Yes | Good | Slight Mismatch; Short Primary Assembly |
| Testudines | rMalTer1 | Malaclemys terrapin | diamondback terrapin | VGP | rk.org/vgp-all/Mala | Yes | Good | Short Primary Assembly |

### IGL

| Class | IndividualID | LatinName | CommonName | Source | SourceLink | Haplotype Resolved | Evaluation Result | Special Condition |
|---|---|---|---|---|---|---|---|---|
| Mammalia | mApoSyl1 | Apodemus sylvaticus | wood mouse | VGP | o/genomeark-all/A | No | Mismatch; Break in Primary Assembly | alternate locus too short/not found |
| Mammalia | mBalAcu1 | Balaenoptera_acutorostrata | minke whale | VGP | rg/vgp-all/Balaen | No | Good | |
| Mammalia | mCamDro1 | Camelus dromedarius | dromedary | VGP | g/genomeark-all/C | Yes | Good | |
| Mammalia | mCanLor1 | Canis_lupus | Greenland Wolf | VGP | e/s3/genomeark/s | No | Mismatch; Break in Primary Assembly | alternate locus too short/not found |
| Mammalia | mCanLor2 | Canis_lupus | Greenland Wolf | VGP | e/s3/genomeark/s | Yes | Break in Primary + Alternate Assembly | |
| Mammalia | mCerEla1 | Cervus_elaphus | Red Deer | VGP | s3/genomeark/spe | No | Good | Short Alternate Assembly |
| Mammalia | mCorTow1.0 | Corynorhinus townsendii | Townsend's Big-eared Bat | CCGP | s/corynorhinus-tov | Yes | Good | Slight Mismatch |
| Mammalia | mCynVol1 | Cynocephalus volans | Philiphine flying lemur | VGP | genomeark/specie | No | Mismatch | alternate locus too short/not found |
| Mammalia | mDasNov1 | Dasypus novemcinctus | nine-banded armadillo | VGP | /genomeark-all/D | Yes | Mismatch; Break in Primary + Alternate Assembly | |
| Mammalia | mDelDel1 | Delphinus delphis | saddleback dolphin | VGP | ark.org/vgp-all/De | No | Mismatch | |
| Mammalia | mDicBic1 | Diceros bicornis | black rhinoceros | VGP | b.io/genomeark-a | Yes | Break in Alternate Assembly | Slight Mismatch |
| Mammalia | mEleMax1 | Elephas maximus | Asiatic Elephant | VGP | 3/genomeark/spec | No | Good | |
| Mammalia | mEptNil1 | Eptesicus nilssonii | northern bat | VGP | ark.org/vgp-all/Ept | No | Good | Slight Mismatch |
| Mammalia | mEriEur2 | Erinaceus europaeus | western European hedgehog | VGP | k.org/vgp-all/Erina | No | Mismatch; Break in Primary Assembly | alternate locus too short/not found |
| Mammalia | mEscRob2 | Eschrichtius robustus | grey whale | VGP | k.org/vgp-all/Esch | No | Good | |
| Mammalia | mEubGla1 | Eubalaena glacialis | North Atlantic right whale | VGP | rk.org/vgp-all/Eub | Yes | Good | All MapQ 0 reads |
| Mammalia | mGloMel1 | Globicephala melas | long-finned pilot whale | VGP | rk.org/vgp-all/Glol | No | Mismatch; Break in Primary Assembly | alternate locus too short/not found |
| Mammalia | mGorGor1 | Gorilla_gorilla | Gorilla | T2T Primate | /s3/genomeark/sp | Yes | Good | |
| Mammalia | mHetBru1 | Heterohyrax brucei | Yellow-spotted hyrax | VGP | .io/genomeark-all/ | No | Mismatch | |
| Mammalia | mHipAmp2 | Hippopotamus amphibius kiboko | hippopotamus | VGP | ark-curated-asser | Yes | Break in Alternate Assembly | All MapQ 0 reads |
| Mammalia | mHypAmp2 | Hyperoodon ampullatus | northern bottlenose whale | VGP | .org/vgp-all/Hyper | No | Good | Short Alternate Assembly |
| Mammalia | mLagAlb1 | Lagenorhynchus albirostris | white-beaked dolphin | VGP | rg/vgp-all/Lageno | No | Mismatch | alternate locus too short/not found |
| Mammalia | mLemCat1 | Lemur catta | Ring-tailed lemur | VGP | hub.io/genomeark | No | Break in Primary + Alternate Assembly | Slight Mismatch |
| Mammalia | mLynRuf1 | Lynx rufus | Bobcat | CCGP | oject.org/species/ | No | Good | Slight Mismatch; alternate locus too short/not found |
| Mammalia | mMacEug1 | Macropus eugenii | tammar wallaby | VGP | .io/genomeark-all | No | Break in Primary Assembly | alternate locus too short/not found |
| Mammalia | mManPen7 | Manis pentadactyla | Chinese pangolin | VGP | io/genomeark-all/ | Yes | Good | All MapQ 0 reads |
| Mammalia | mMarMar1 | Martes martes | European pine marten | VGP | eark.org/vgp-all/M | No | Mismatch | alternate locus too short/not found |
| Mammalia | mMelMel3 | Meles meles | European badger | VGP | hub.io/genomeark- | Yes | Good | Most MapQ 0 reads |
| Mammalia | mMesDen1 | Mesoplodon densirostris | Blainville's beaked whale | VGP | org/vgp-all/Mesop | No | Good | |
| Mammalia | mMicCal1.0 | Microtus californicus | California Vole | CCGP | /species/microtus- | Yes | Good | Short Primary Assembly |
| Mammalia | mMicMin1 | Micromys minutus | European harvest mouse | VGP | ark.org/vgp-all/Mic | No | Mismatch; Break in Primary Assembly | alternate locus too short/not found |
| Mammalia | mMirAng1 | Mirounga angustirostris | Northern Elephant Seal | CCGP | es/mirounga-angu | Yes | Break in Primary + Alternate Assembly | 2 IGL |
| Mammalia | mMonDom1 | Monodelphis_domestica | gray short-tailed opossum | VGP | nomeark/species/ | No | Break in Primary Assembly | alternate locus too short/not found |
| Mammalia | mMunRee1 | Muntiacus reevesi | Reeves' muntjac | VGP | ark.org/vgp-all/Mu | No | Good | Slight Mismatch; alternate locus too short/not found |
| Mammalia | mMusAve1 | Muscardinus avellanarius | hazel dormouse | VGP | org/vgp-all/Musca | No | Mismatch | alternate locus too short/not found |
| Mammalia | mMusLut2 | Mustela lutreola | European mink | VGP | .org/genomeark-a | No | Good | Slight Mismatch; Short Alternate Assembly |
| Mammalia | mMusNiv1 | Mustela nivalis | Least weasel | VGP | .org/genomeark-a | Yes | Mismatch; Break in Primary + Alternate Assembly | |
| Mammalia | mMyoDau2 | Myotis daubentonii | Daubenton's bat | VGP | ark.org/vgp-all/My | No | Mismatch | alternate locus too short/not found |
| Mammalia | mMyoYum1.0 | Myotis yumanensis | Yuma myotis | CCGP | g/species/myotis- | Yes | Good | Short Primary Assembly |
| Mammalia | mNeoNeb1 | Neofelis_nebulosa | Clouded Leopard | VGP | 3/genomeark/spec | No | Good | alternate locus too short/not found |
| Mammalia | mNycCou1 | Nycticebus coucang | slow loris | VGP | io/genomeark-all/ | No | Mismatch; Break in Primary Assembly | extremly abnormal coverage in Alternate Assembly |
| Mammalia | mOrcOrc1 | Orcinus orca | killer whale | VGP | hub.io/genomeark- | No | Good | alternate locus too short/not found |
| Mammalia | mOryCun1 | Oryctolagus cuniculus | rabbit | VGP | k.org/vgp-all/Oryc | No | Good | alternate locus too short/not found |
| Mammalia | mPanPan1 | Pan paniscus | Bonobo | T2T Primate | /s3/genomeark/sp | Yes | Good | |
| Mammalia | mPerMan1 | Peromyscus maniculatus | deer mouse | CCGP | pecies/peromysc | No | Good | Slight Mismatch; Short Alternate Assembly |
| Mammalia | mPhoPho1 | Phocoena phocoena | harbor porpoise | VGP | k.org/vgp-all/Pho | No | Mismatch | Short Alternate Assembly |
| Mammalia | mPipPyg2 | Pipistrellus pygmaeus | soprano pipistrelle | VGP | k.org/vgp-all/Pipis | No | Good | |
| Mammalia | mPonAbe1 | Pongo_abelii | Sumatran orangutan | T2T Primate | e/s3/genomeark/sp | Yes | Good | |
| Mammalia | mPonPyg2 | Pongo_pygmaeus | Bornean orangutan | T2T Primate | 3/genomeark/spec | Yes | Good | |
| Mammalia | mPseCra1 | Pseudorca crassidens | false killer whale | VGP | k.org/vgp-all/Pseu | Yes | Good | |
| Mammalia | mPumCon1.1 | Puma concolor | Mountain Lion | CCGP | org/species/puma | Yes | Good | |
| Mammalia | mSorAra2/1 | Sorex araneus | Common shrew | VGP | ub.io/genomeark-a | No | Mismatch; Break in Primary Assembly | Short Alternate Assembly |
| Mammalia | mSteCoe1 | Stenella coeruleoalba | striped dolphin | VGP | k.org/vgp-all/Sten | No | Mismatch | Short Alternate Assembly |
| Mammalia | mTalEur1 | Talpa europaea | European mole | VGP | eark.org/vgp-all/Ta | No | Break in Primary Assembly | alternate locus too short/not found |
| Mammalia | mUrsAme1 | Ursus americanus | American black bear | CCGP | gov/datasets/gen | No | Break in Primary Assembly | 2 IGL but on same chrom |
| Mammalia | mUrsArc2 | Ursus arctos | brown bear | NCBI | gov/datasets/gen | No | Mismatch | |
| Mammalia | mVesMur1 | Vespertilio murinus | particolored bat | VGP | rk.org/vgp-all/Ves | No | Good | |
| Crocodilia | rAllMis2 | Alligator mississippiensis | American alligator | VGP | genomeark/specie | Yes | Good | |
| Testudines | rCarCar2 | Caretta caretta | Loggerhead turtle | VGP | eark.org/vgp-all/C | Yes | Break in Primary Assembly | Slight Mismatch |
| Testudines | rEmyOrb1 | Emys orbicularis | European pond turtle | VGP | ark.org/vgp-all/Er | Yes | Good | |
| Testudines | rMalTer1 | Malaclemys terrapin | diamondback terrapin | VGP | rk.org/vgp-all/Mala | Yes | Good | |
