## Supplementary material for "Assessing Assembly Errors in Immunoglobulin Loci: A Comprehensive Evaluation of Long-read Genome Assemblies Across Vertebrates": IGH Loci Evaluations

Table of Contents

1. mApoSyl1 - Apodemus sylvaticus - Wood mouse
2. mBalAcu1 - Balaenoptera acutorostrata - Minke whale
3. mCamDro1 - Camelus dromedarius - Dromedary
4. mCerEla1 - Cervus elaphus - Red Deer
5. mChiNiv1 - Chionomys nivalis - European snow vole
6. mCorTow1.0 - Corynorhinus townsendii - Townsend's Big-eared Bat
7. mDasNov1 - Dasypus novemcinctus - Nine-banded armadillo
8. mDelDel1 - Delphinus delphis - Saddleback dolphin
9. mDicBic1 - Diceros bicornis - Black rhinoceros
10. mDipMer1 - Dipodomys merriami - Merriam's Kangaroo Rat
11. mEleMax1 - Elephas maximus - Asiatic Elephant
12. mEptNil1 - Eptesicus nilssonii - Northern bat
13. mEriEur2 - Erinaceus europaeus - Western European hedgehog
14. mEscRob2 - Eschrichtius robustus - Grey whale
15. mEubGla1 - Eubalaena glacialis - North Atlantic right whale
16. mGloMel1 - Globicephala melas - Long-finned pilot whale
17. mGorGor1 - Gorilla gorilla - Gorilla
18. mHetBru1 - Heterohyrax brucei - Yellow-spotted hyrax
19. mHipAmp2 - Hippopotamus amphibius kiboko - Hippopotamus
20. mHypAmp2 - Hyperoodon ampullatus - Northern bottlenose whale
21. mLagAlb1 - Lagenorhynchus albirostris - White-beaked dolphin
22. mLemCat1 - Lemur catta - Ring-tailed lemur
23. mLynRuf1 - Lynx rufus - Bobcat
24. mMacEug1 - Macropus eugenii - Tammar wallaby
25. mManPen7 - Manis pentadactyla - Chinese pangolin
26. mMarMar1 - Martes martes - European pine marten
27. mMelMel3 - Meles meles - European badger
28. mMesDen1 - Mesoplodon densirostris - Blainville's beaked whale
29. mMicCal1.0 - Microtus californicus - California Vole
30. mMicMin1 - Micromys minutus - European harvest mouse
31. mMirAng1 - Mirounga angustirostris - Northern Elephant Seal
32. mMonDom1 - Monodelphis domestica - Gray short-tailed opossum
33. mMunRee1 - Muntiacus reevesi - Reeves' muntjac
34. mMusAve1 - Muscardinus avellanarius - Hazel dormouse
35. mMusLut2 - Mustela lutreola - European mink
36. mMusNiv1 - Mustela nivalis - Least weasel
37. mMyoDau2 - Myotis daubentonii - Daubenton's bat
38. mMyoYum1.0 - Myotis yumanensis - Yuma myotis
39. mNeoNeb1 - Neofelis nebulosa - Clouded Leopard
40. mNycCou1 - Nycticebus coucang - Slow loris
41. mOrcOrc1 - Orcinus orca - Killer whale
42. mOryCun1 - Oryctolagus cuniculus - Rabbit
43. mPanPan1 - Pan paniscus - Bonobo
44. mPerMan1 - Peromyscus maniculatus - Deer mouse
45. mPhoPho1 - Phocoena phocoena - Harbor porpoise
46. mPipPyg2 - Pipistrellus pygmaeus - Soprano pipistrelle
47. mPleAur1 - Plecotus auritus - Brown big-eared bat
48. mPonAbe1 - Pongo abelii - Sumatran orangutan
49. mPonPyg2 - Pongo pygmaeus - Bornean orangutan
50. mPseCra1 - Pseudorca crassidens - False killer whale
51. mPumCon1.1 - Puma concolor - Mountain Lion
52. mSorAra2/1 - Sorex araneus - Common shrew
53. mSteCoe1 - Stenella coeruleoalba - Striped dolphin
54. mTalEur1 - Talpa europaea - European mole
55. mThoBot1 - Thomomys bottae - Botta's pocket gopher
56. mUrsAme1 - Ursus americanus - American black bear
57. mUrsArc2 - Ursus arctos - Brown bear
58. mVesMur1 - Vespertilio murinus - Particolored bat
59. rAllMis2 - Alligator mississippiensis - American alligator
60. rCarCar2 - Caretta caretta - Loggerhead turtle
61. rEmyOrb1 - Emys orbicularis - European pond turtle
62. rEryReg1 - Erythrolamprus reginae - Royal ground snake
63. rLiaOli1 - Liasis olivaceus - Olive python
64. rLiaOli2 - Liasis olivaceus - Olive python
65. rMalTer1 - Malaclemys terrapin - Diamondback terrapin
66. rPodCre2 - Podarcis cretensis - Cretan wall lizard
67. rPodRaf1 - Podarcis raffonei - Aeolian wall lizard
68. rRhiFlo1 - Rhineura floridana - Florida worm lizard
69. rVipLat1 - Vipera latastei - Snub-nosed viper
70. rVipUrs1 - Vipera ursinii - Hungarian meadow viper
71. rZooViv1 - Zootoca vivipara - Common lizard

Note: This supplementary material only displays alternate IG loci if they are longer than one-quarter of the corresponding primary IG locus length. Loci shorter than this threshold are typically too fragmented to be shown. 18 species had alternate IGH loci shorter than this threshold and are not shown.

Species ID: mApoSyl1

Common Name: wood mouse

Scientific Name: Apodemus sylvaticus

Assembly Type: Not Haplotype Resolved

Data Source: VGP

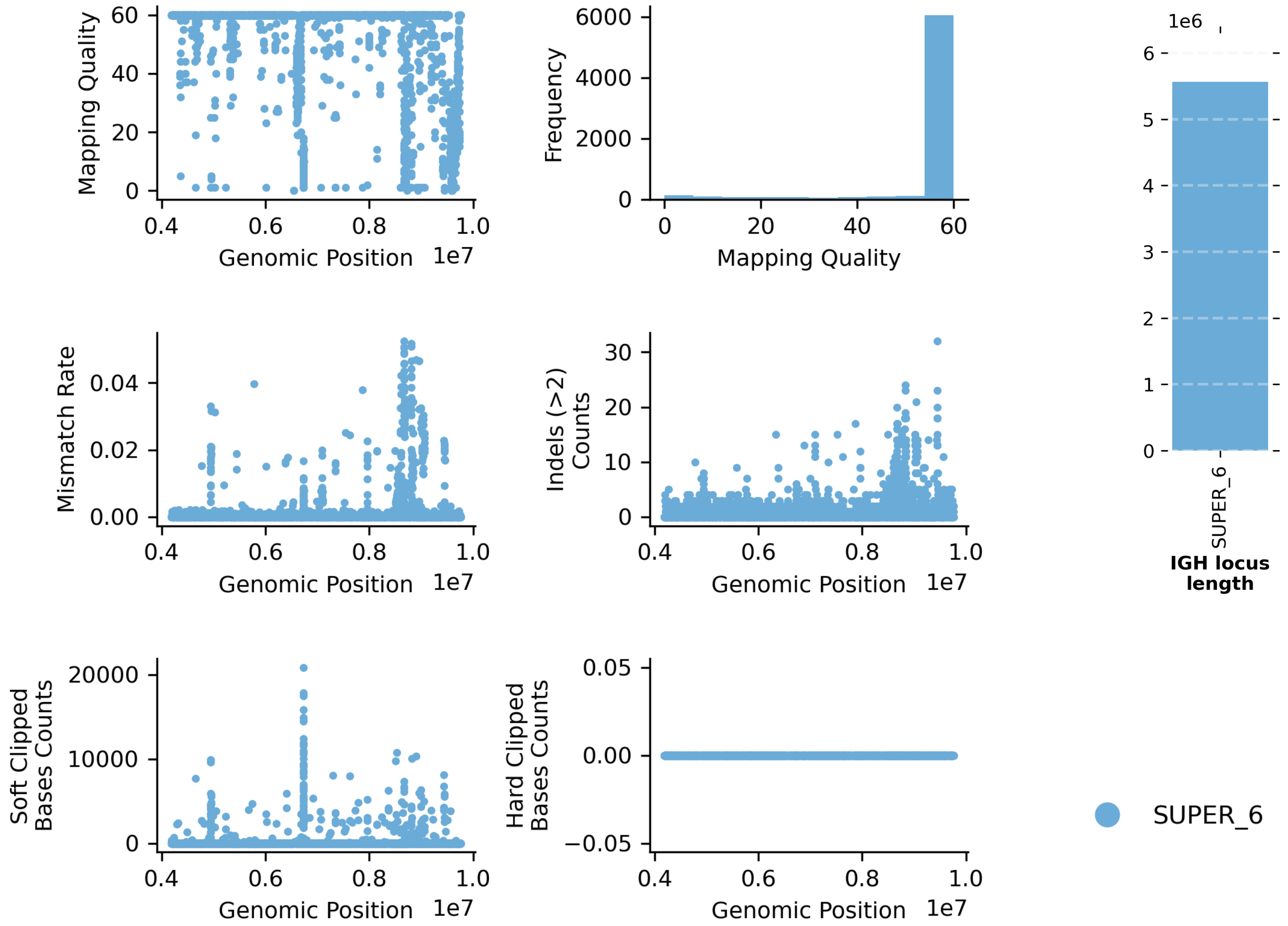


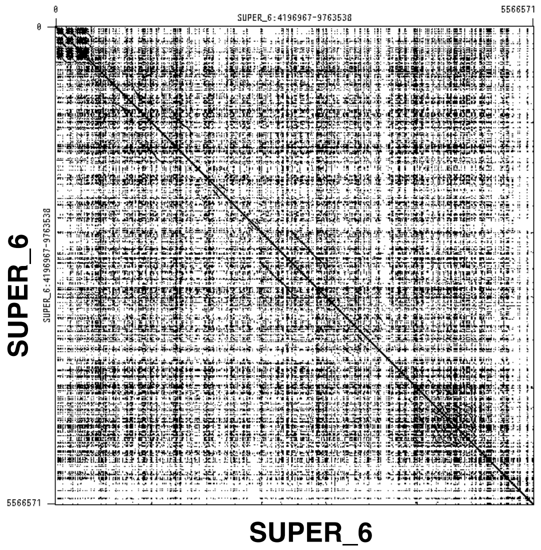


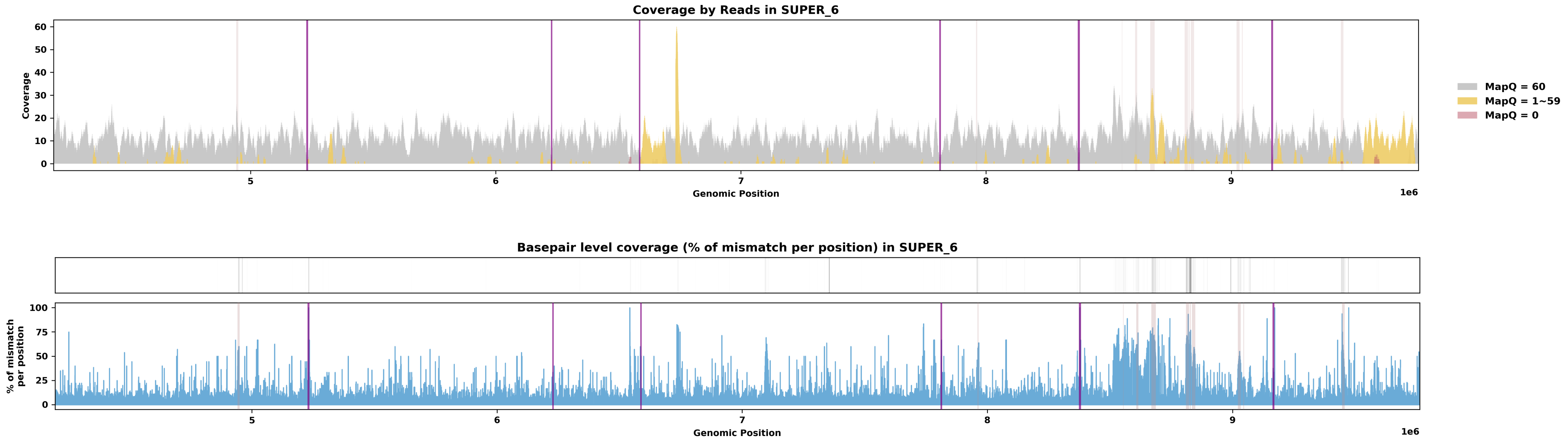

Species ID: mBalAcu1

Common Name: minke whale

Scientific Name: Balaenoptera_acutorostrata

Assembly Type: Not Haplotype Resolved

Data Source: VGP

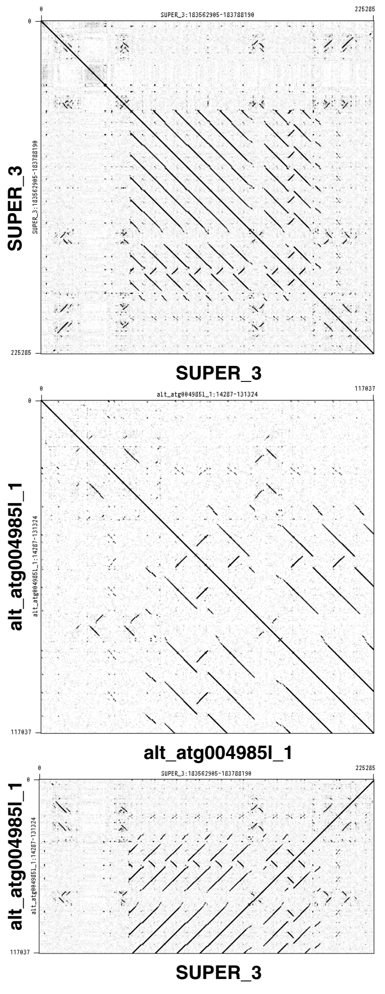


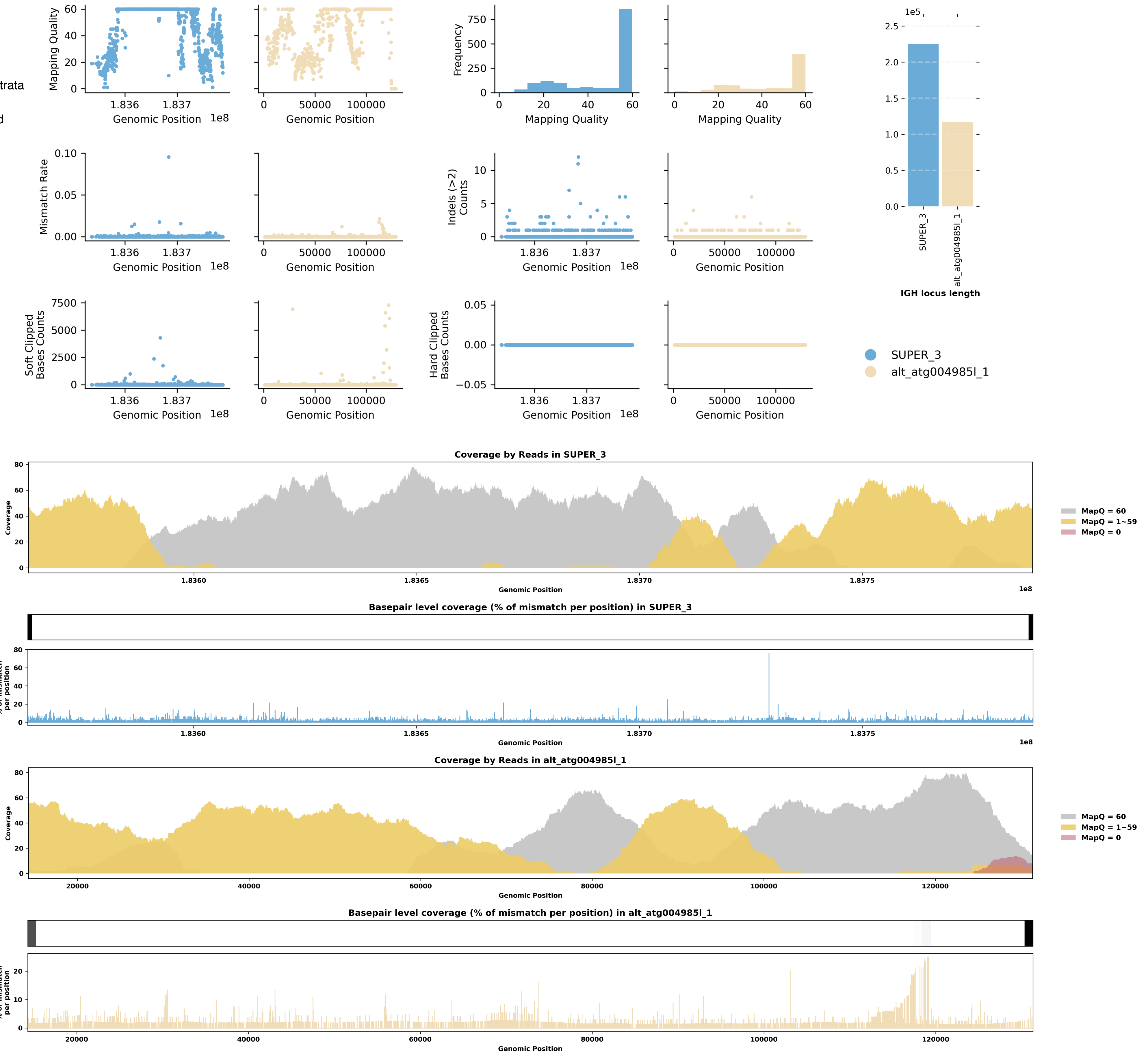

Species ID: mCamDro1

Common Name: dromedary

Scientific Name: Camelus dromedarius

Assembly Type: Haplotype Resolved

Data Source: VGP

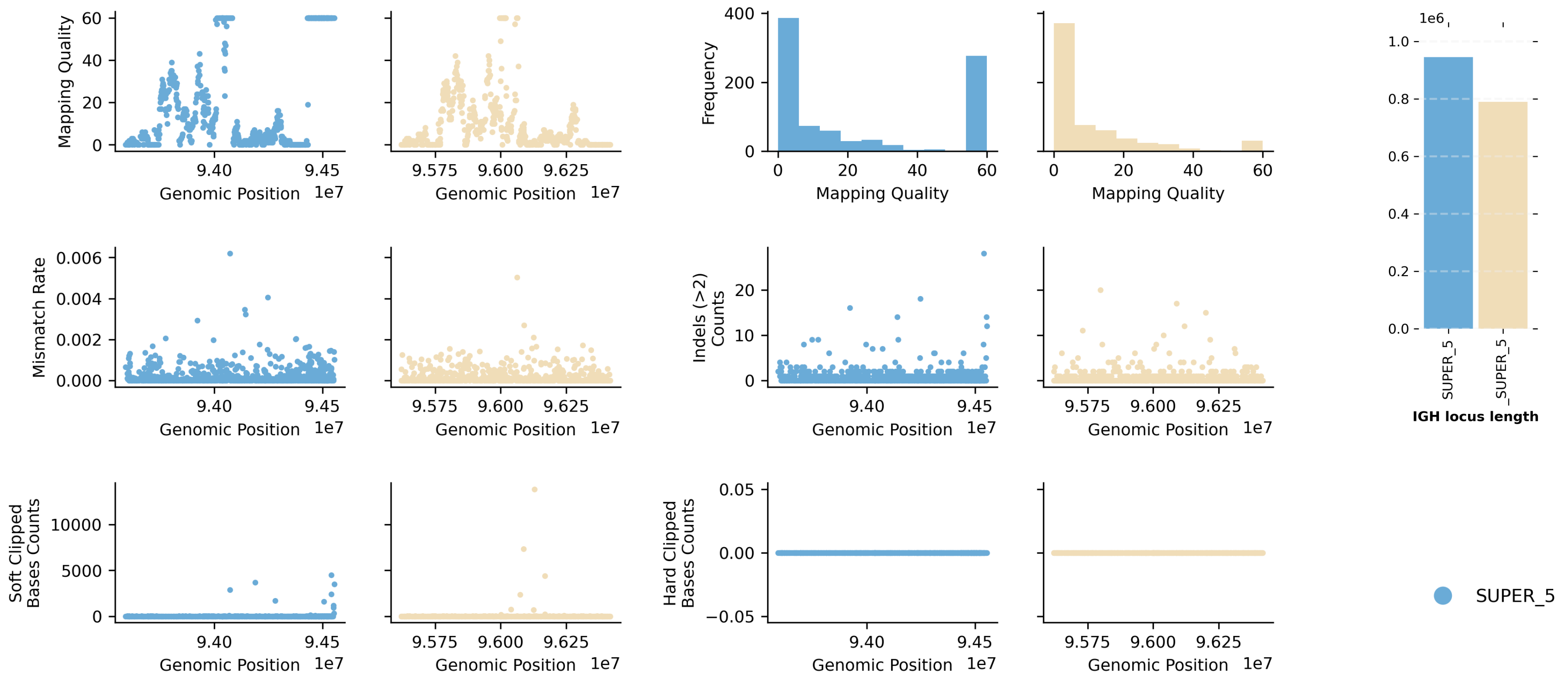


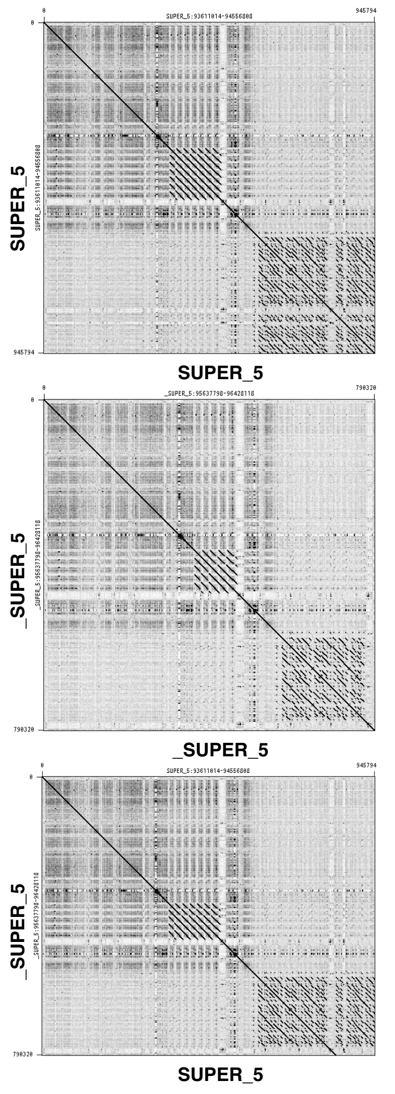


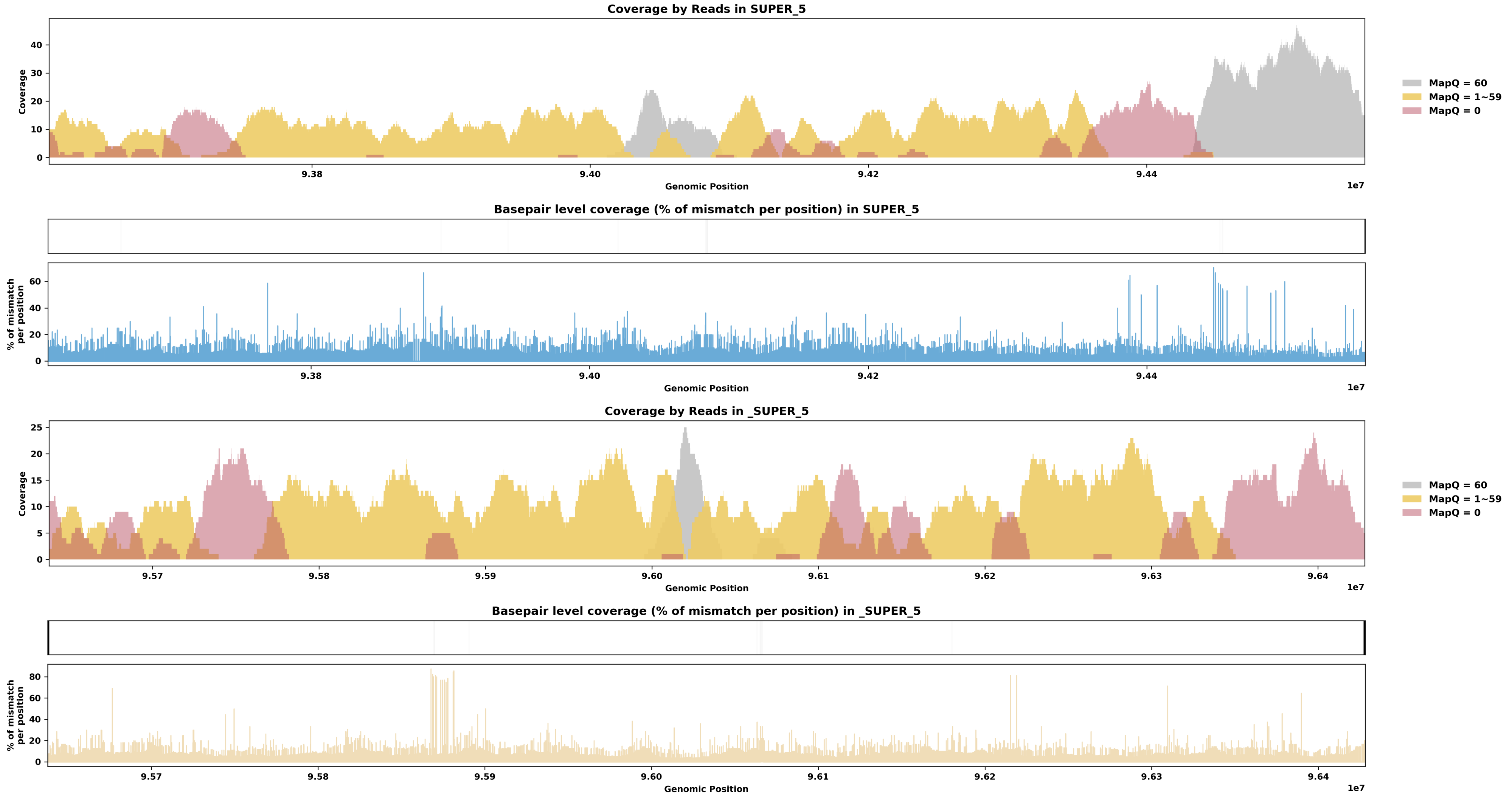

Species ID: mCerEla1

Common Name: Red Deer

Scientific Name: Cervus_elaphus

Assembly Type: Not Haplotype Resolved

Data Source: VGP

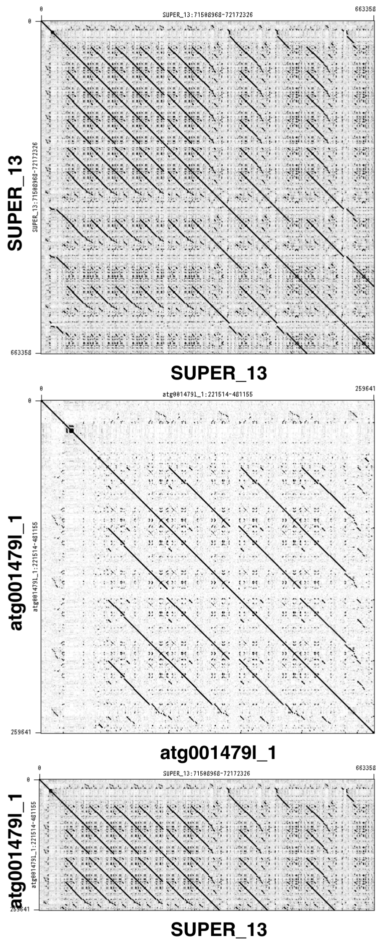


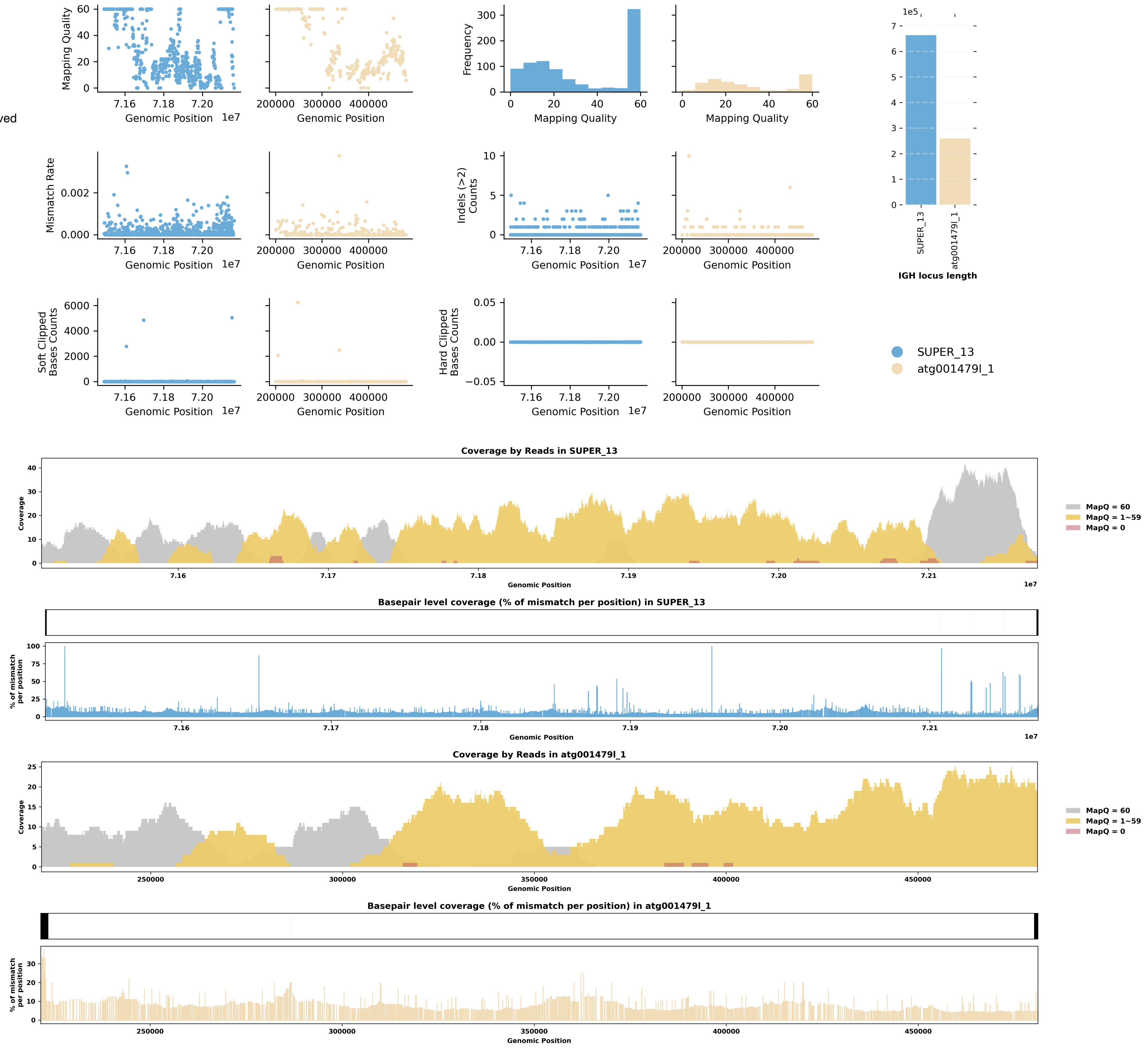

Species ID: mChiNiv1

Common Name: European snow vole

Scientific Name: Chionomys nivalis

Assembly Type: Not Haplotype Resolved

Data Source: VGP

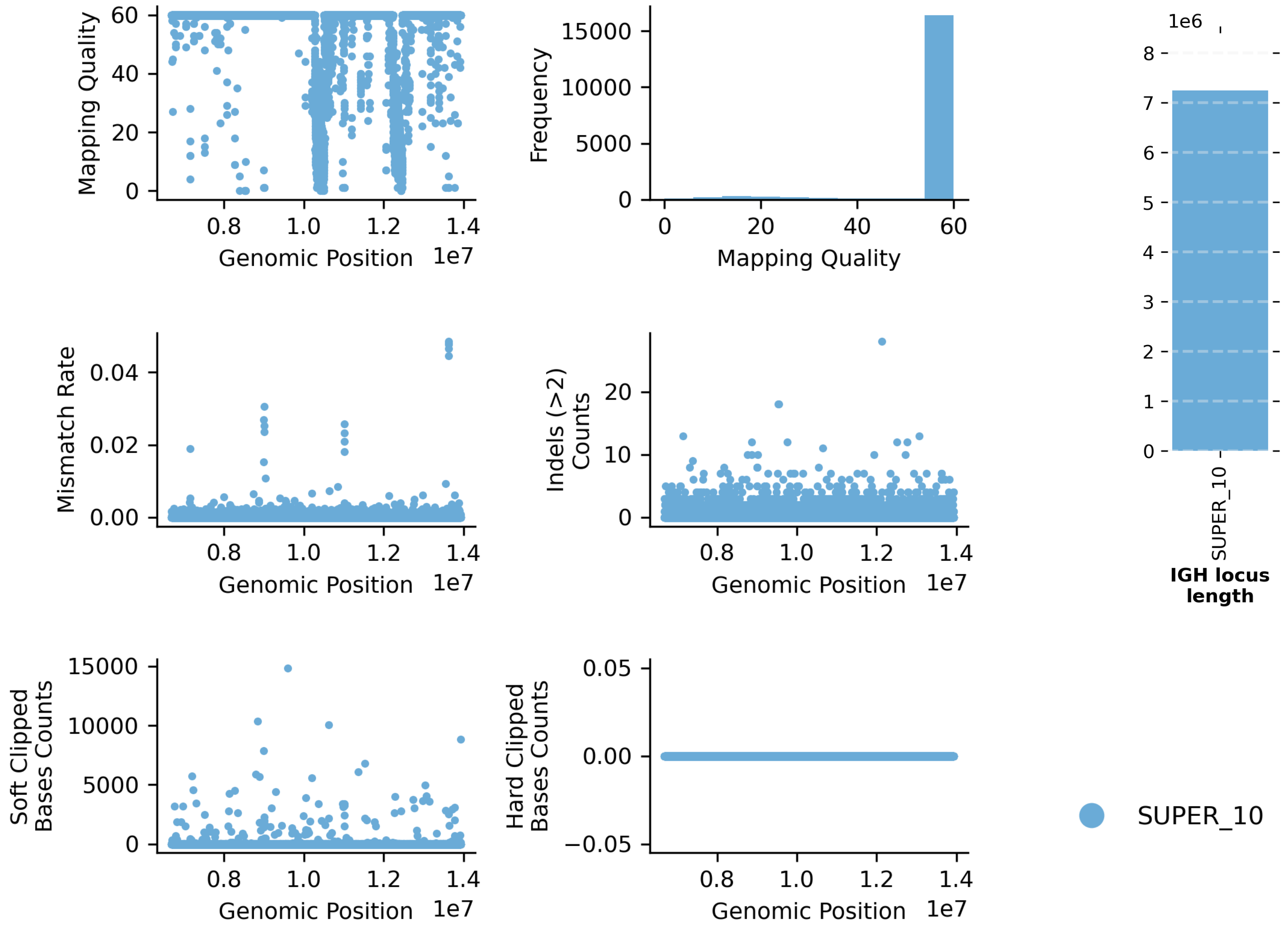


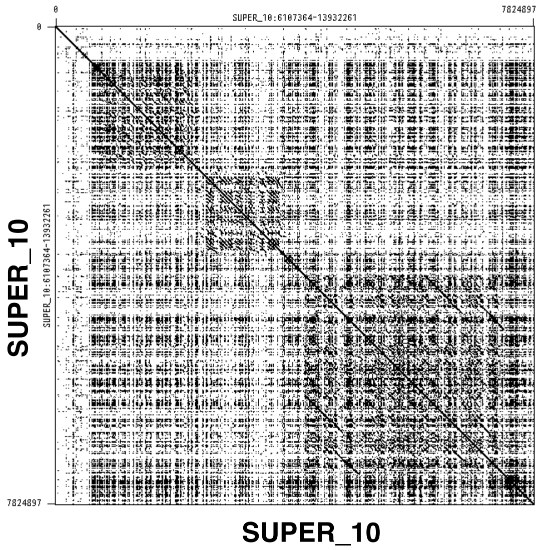


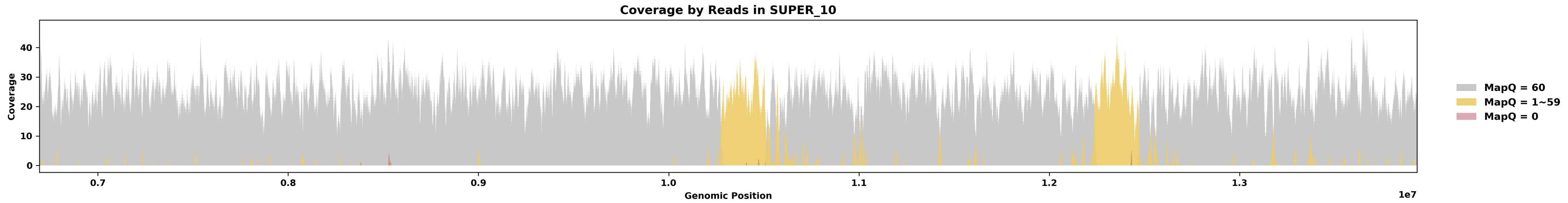


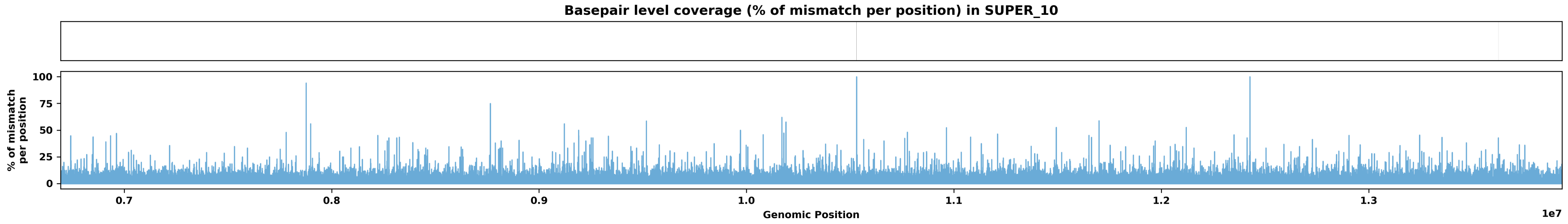

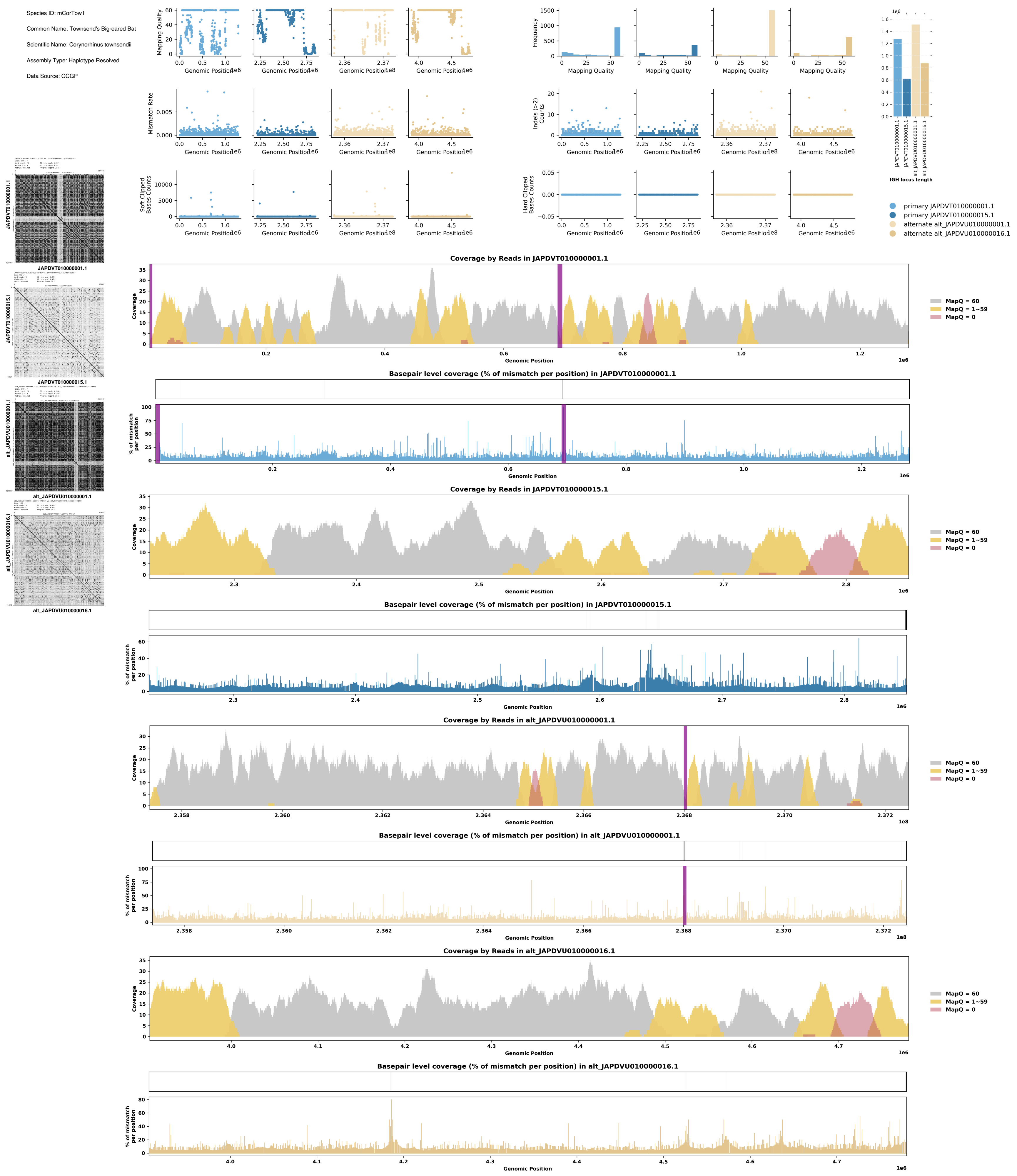
Species ID: mCorTow1
Common Name: Townsend's Big-eared Bat
Scientific Name: Corynorhinus townsendii
Assembly Type: Haplotype Resolved
Data Source: CCGP
Mapping Quality
Genomic Position
Frequency
Mismatch Rate
Indels (>2) Counts
Soft Clipped Bases Counts
Hard Clipped Bases Counts
IGH locus length
primary JAPDVT010000001.1
primary JAPDVT010000015.1
alternate alt_JAPDVU010000001.1
alternate alt_JAPDVU010000016.1
JAPDVT010000001.1
JAPDVT010000015.1
alt_JAPDVU010000001.1
alt_JAPDVU010000016.1
Coverage by Reads in JAPDVT010000001.1
Basepair level coverage (% of mismatch per position) in JAPDVT010000001.1
Coverage by Reads in JAPDVT010000015.1
Basepair level coverage (% of mismatch per position) in JAPDVT010000015.1
Coverage by Reads in alt_JAPDVU010000001.1
Basepair level coverage (% of mismatch per position) in alt_JAPDVU010000001.1
Coverage by Reads in alt_JAPDVU010000016.1
Basepair level coverage (% of mismatch per position) in alt_JAPDVU010000016.1
Coverage
% of mismatch per position
MapQ = 60
MapQ = 1~59
MapQ = 0

Species ID: mDasNov1

Common Name: nine-banded armadillo

Scientific Name: Dasypus novemcinctus

Assembly Type: Haplotype Resolved

Data Source: VGP

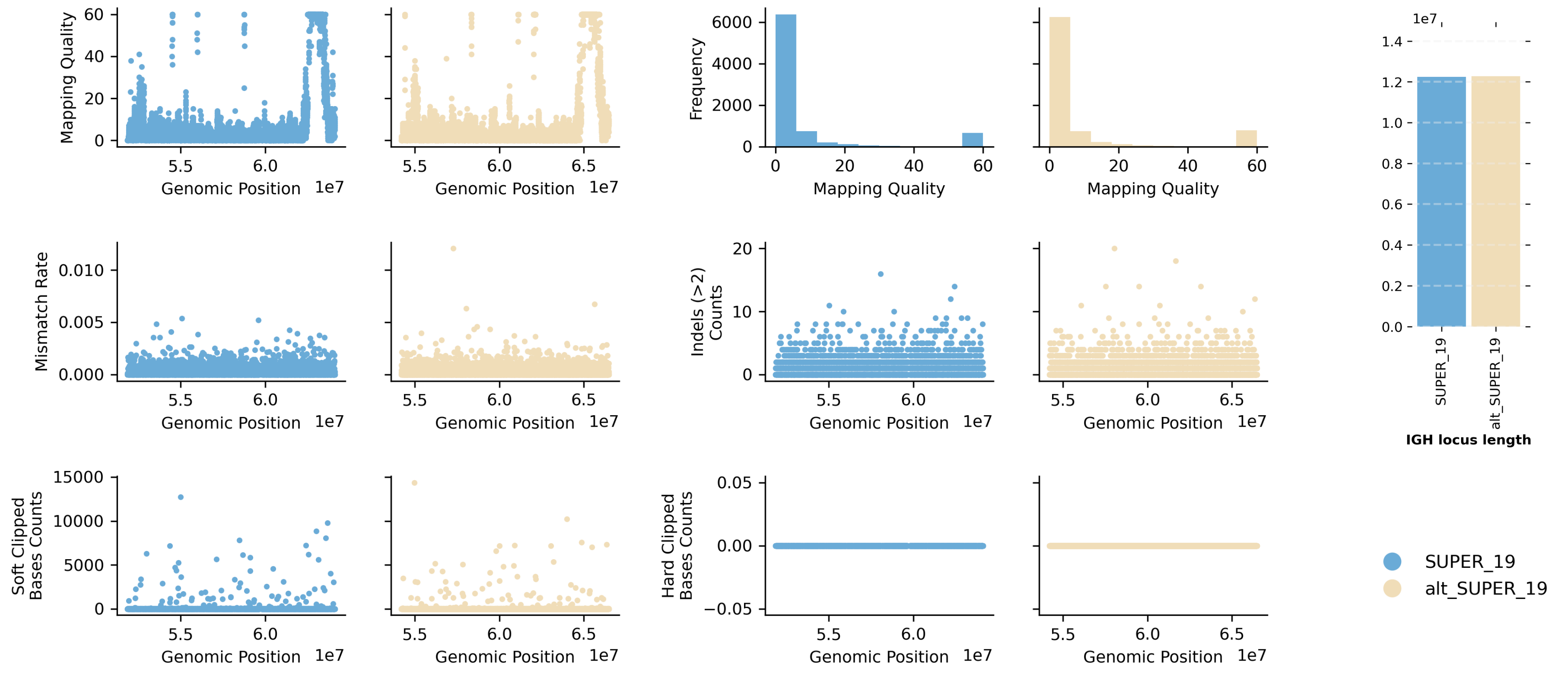


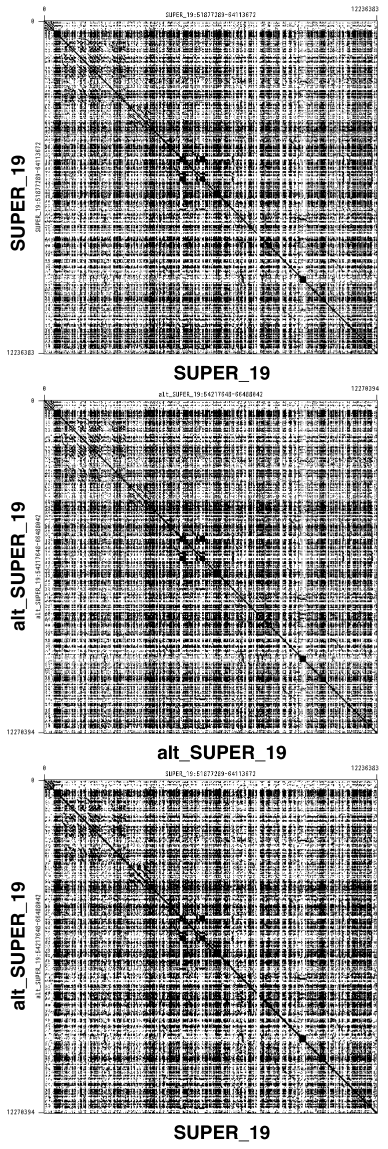


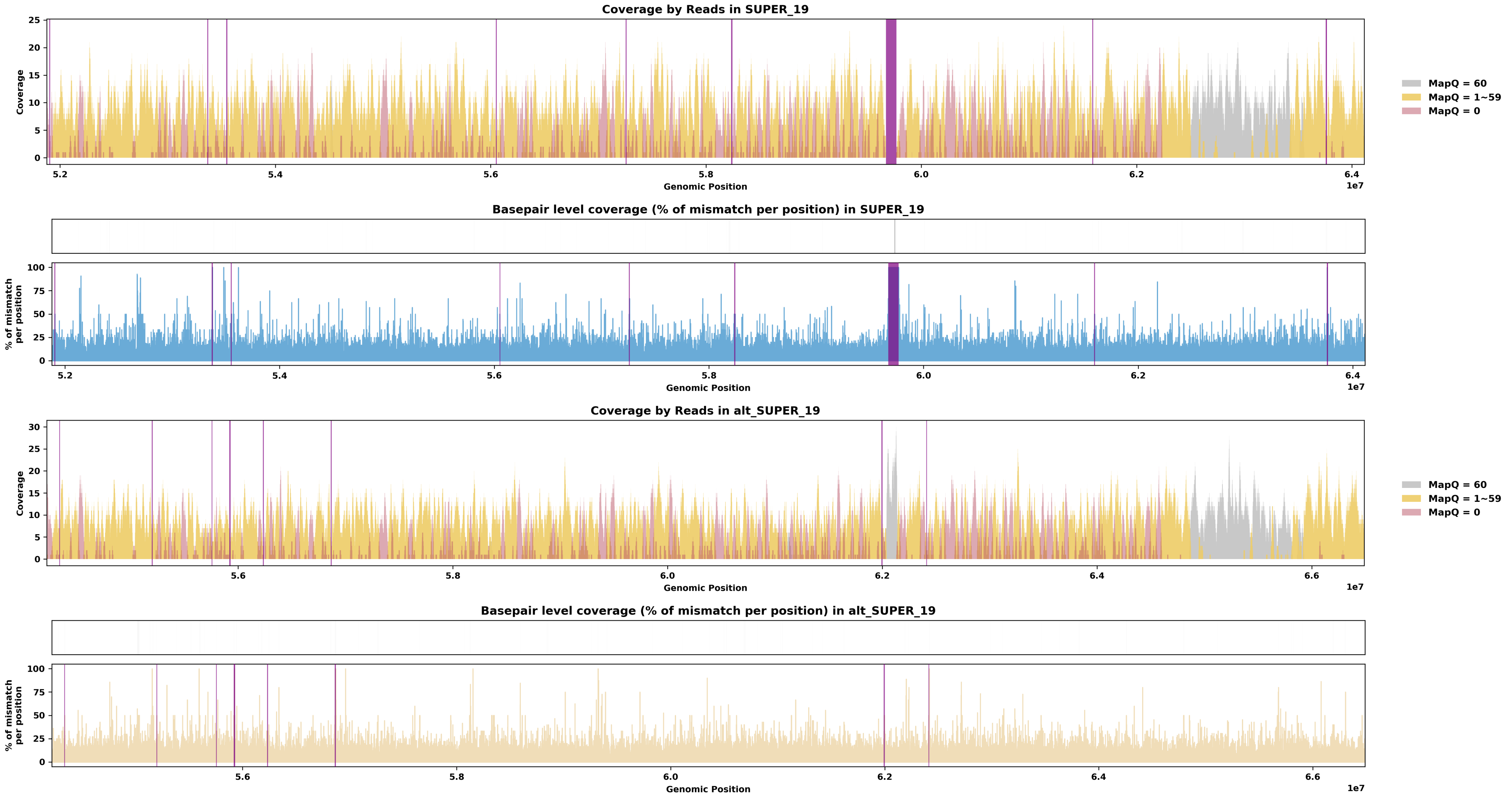

Species ID: mDelDel1

Common Name: saddleback dolphin

Scientific Name: Delphinus delphis

Assembly Type: Not Haplotype Resolved

Data Source: VGP

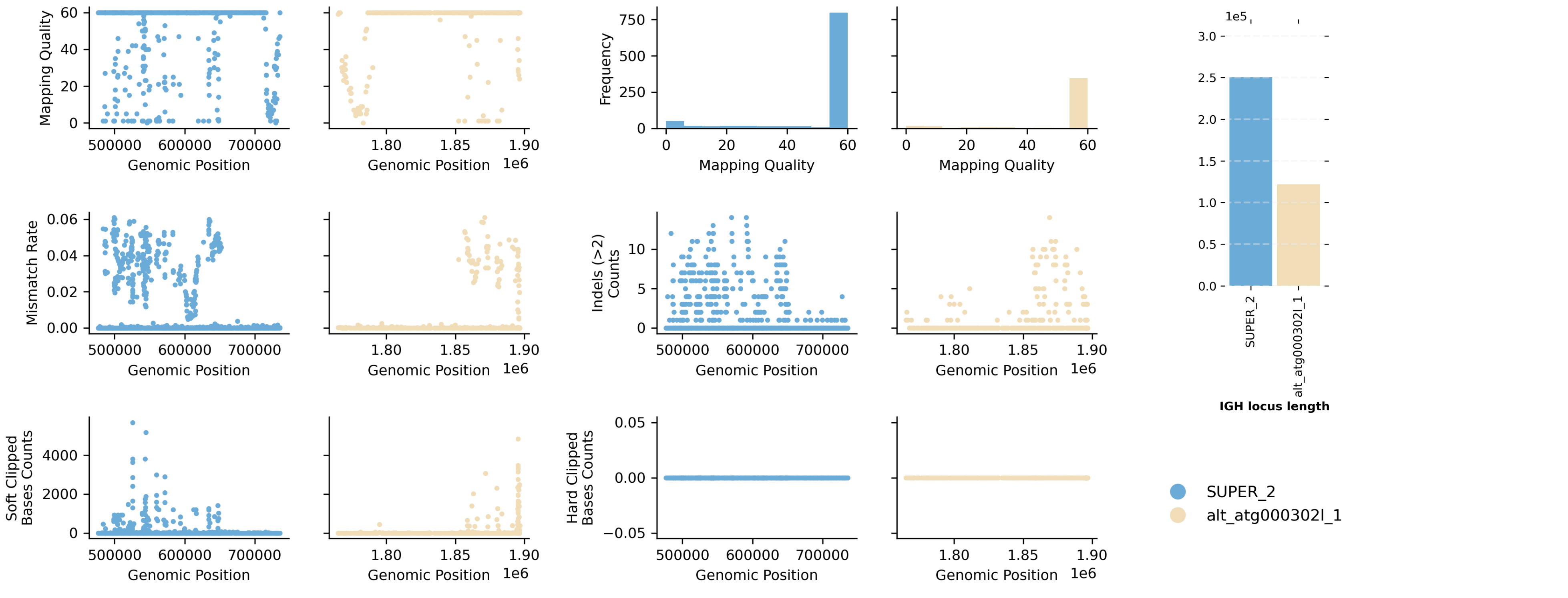


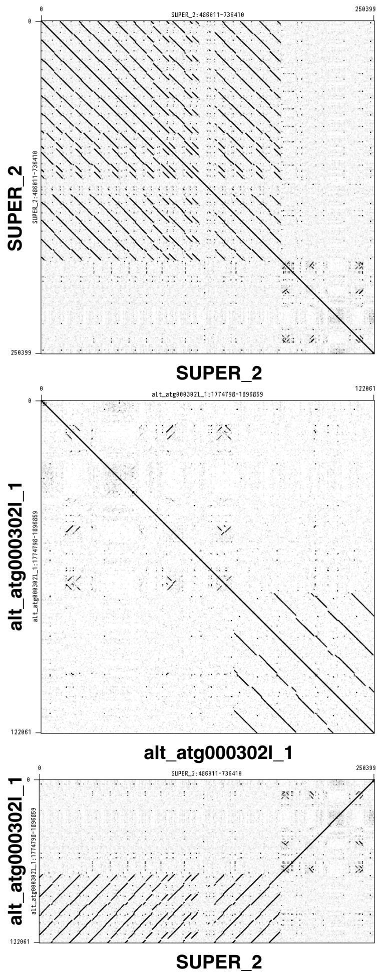


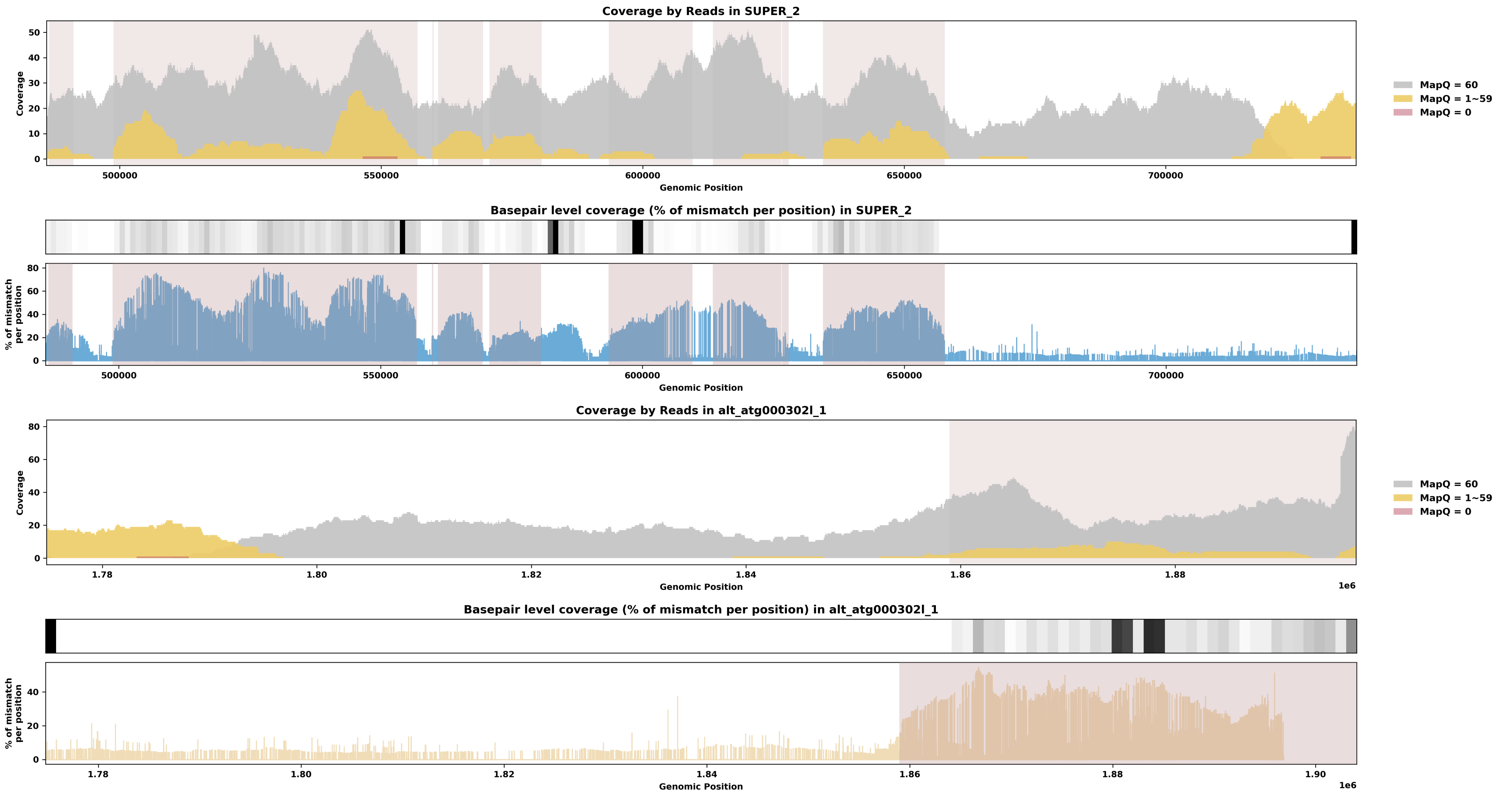

Species ID: mDicBic1

Common Name: black rhinoceros

Scientific Name: Diceros bicornis

Assembly Type: Haplotype Resolved

Data Source: VGP

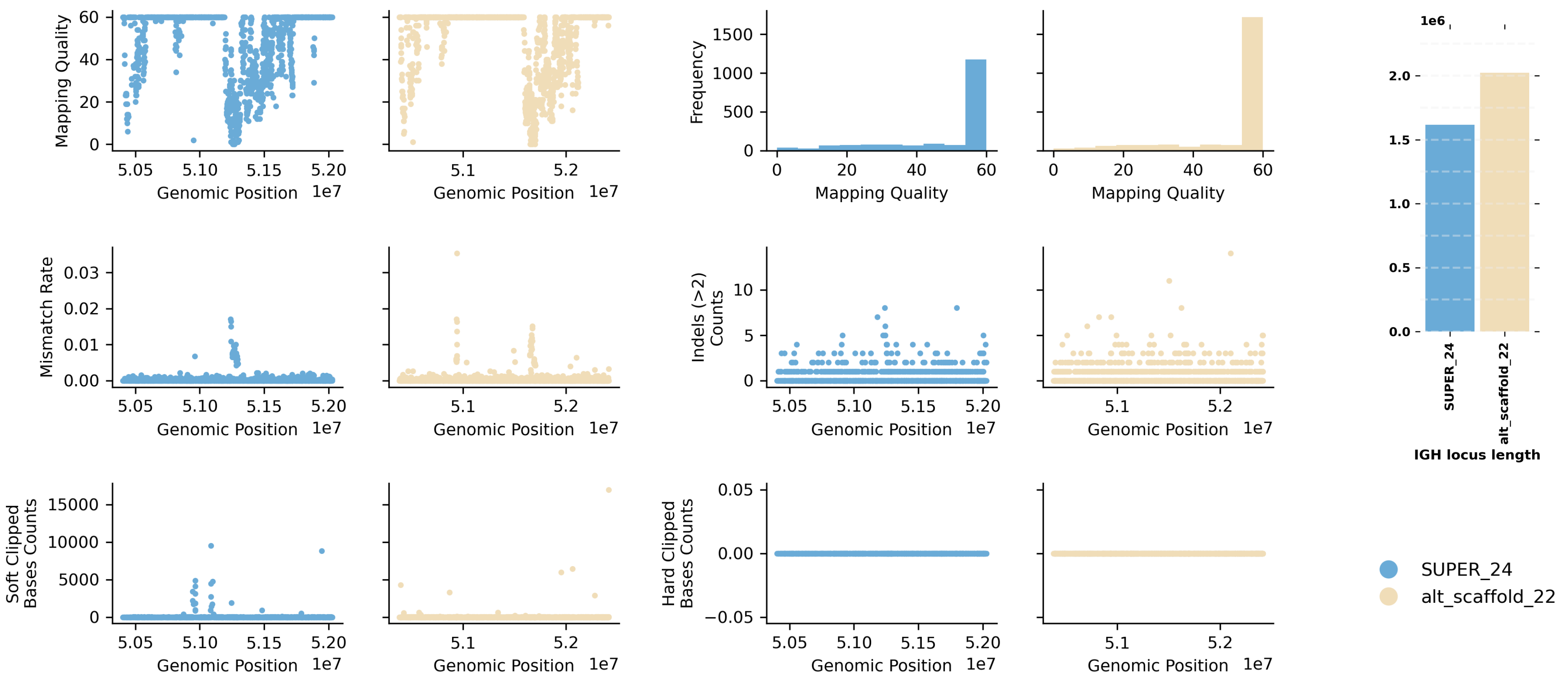


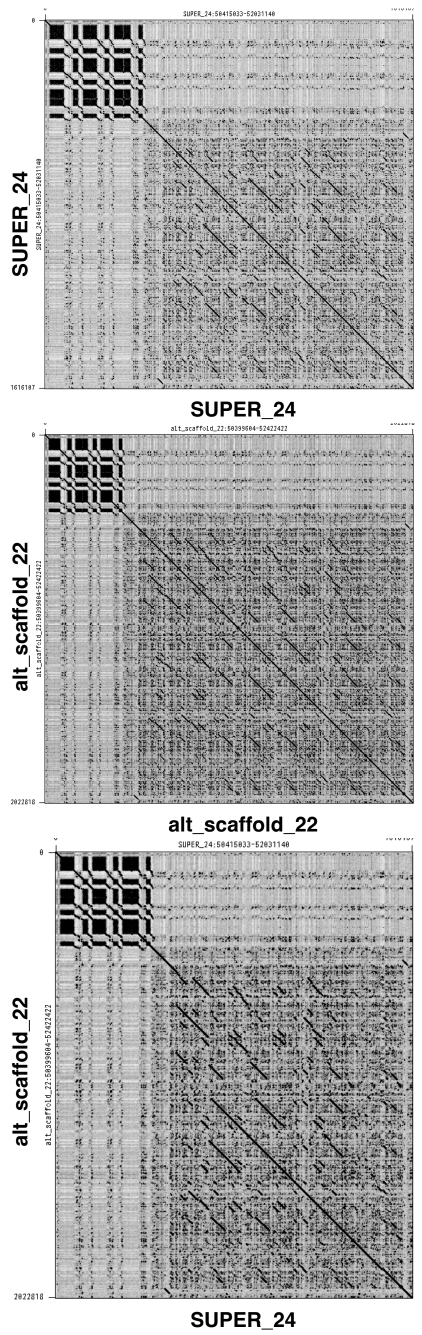


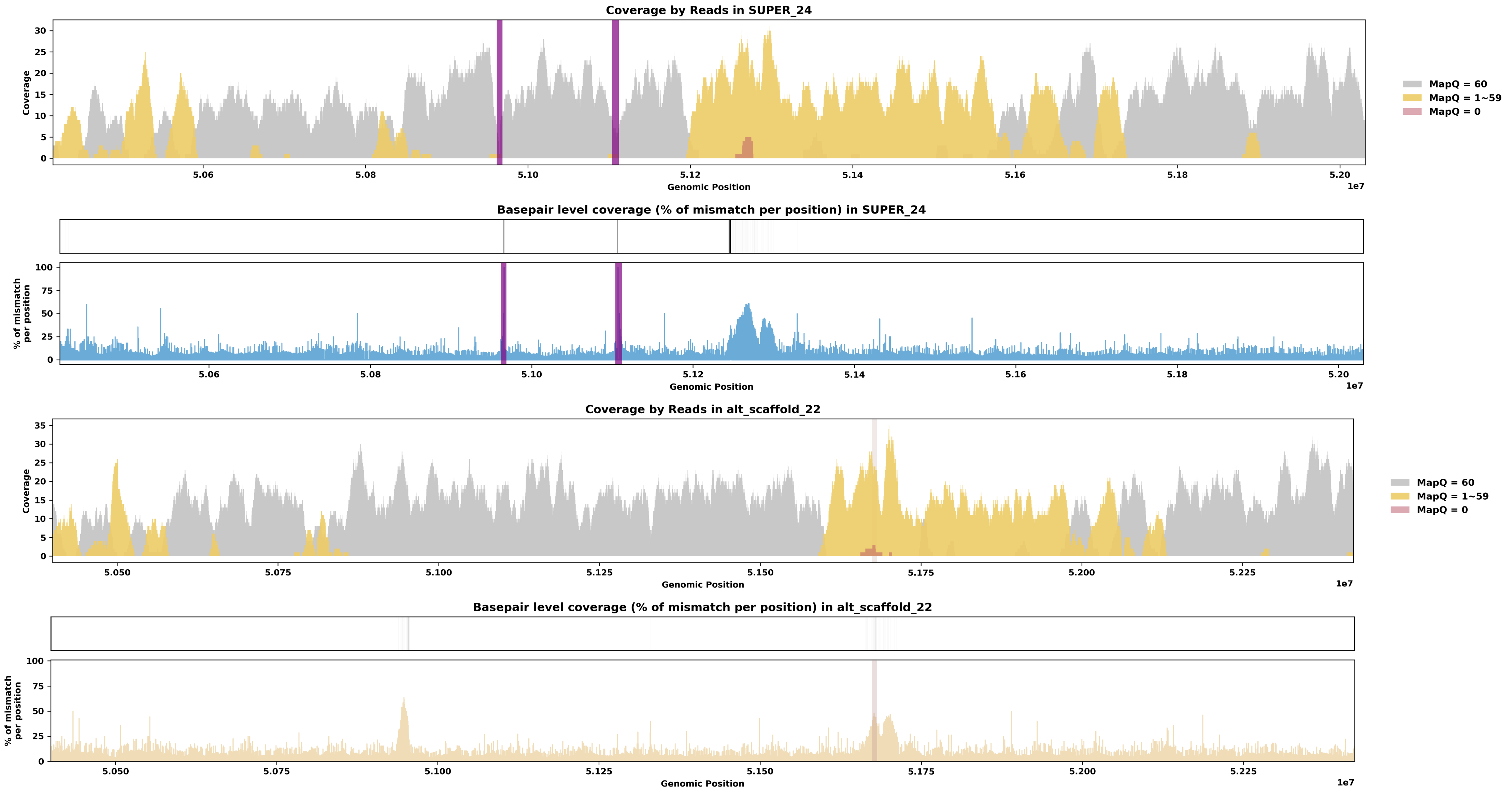

Species ID: mDipMer1

Common Name: Merriam Kangaroo Rat

Scientific Name: Dipodomys merriami

Assembly Type: Not Haplotype Resolved

Data Source: CCGP

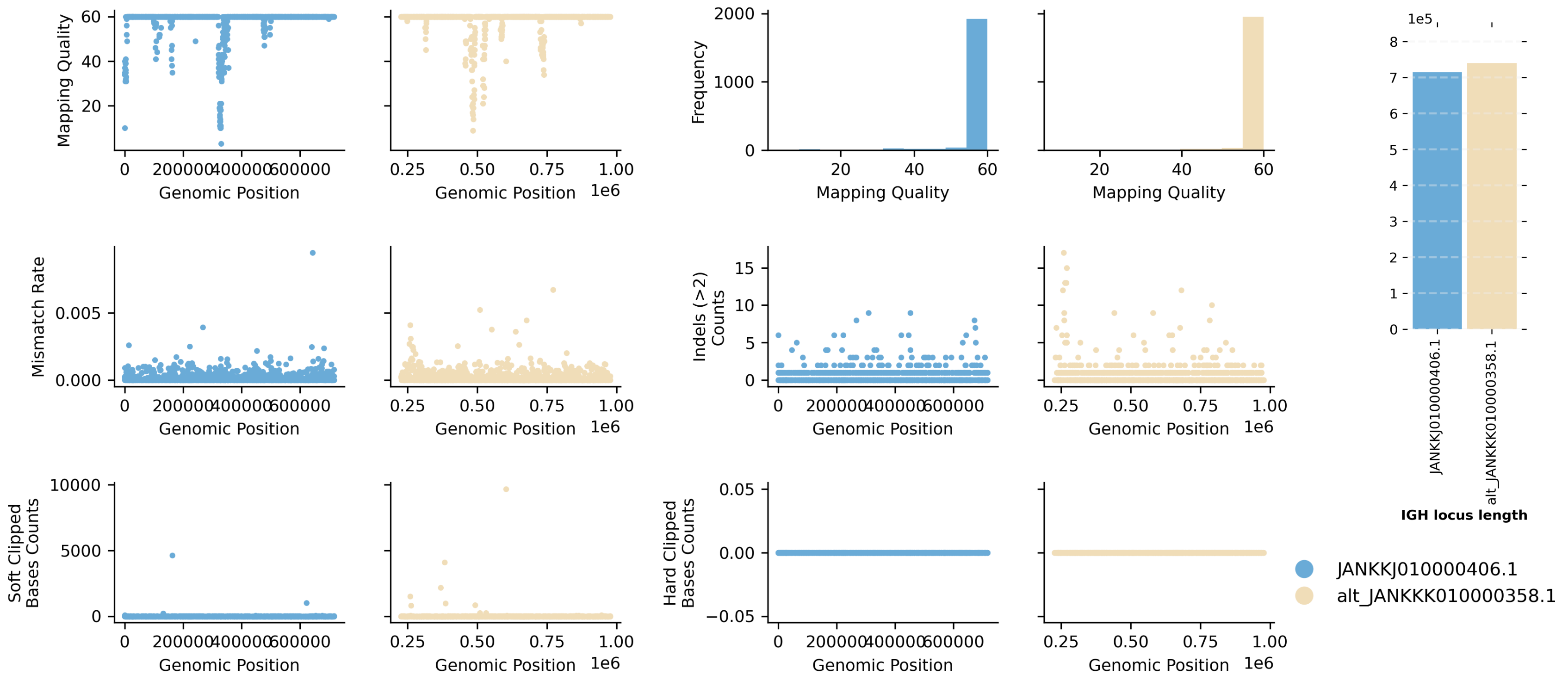


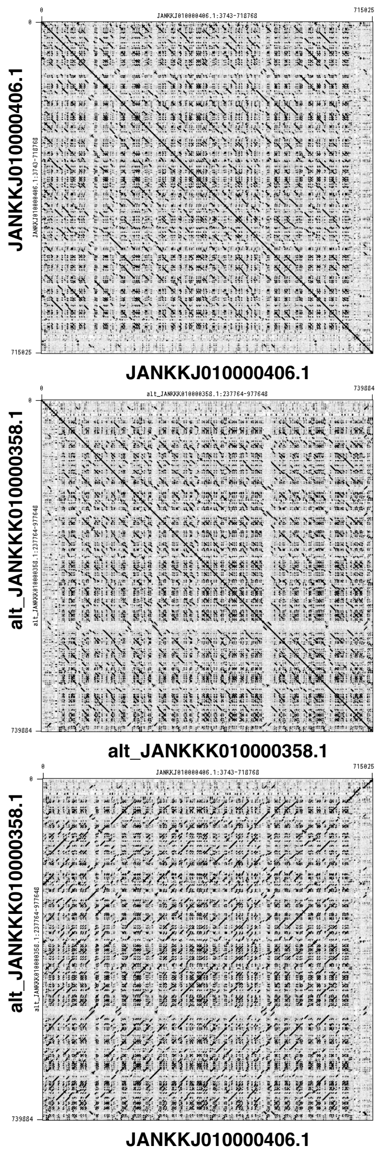


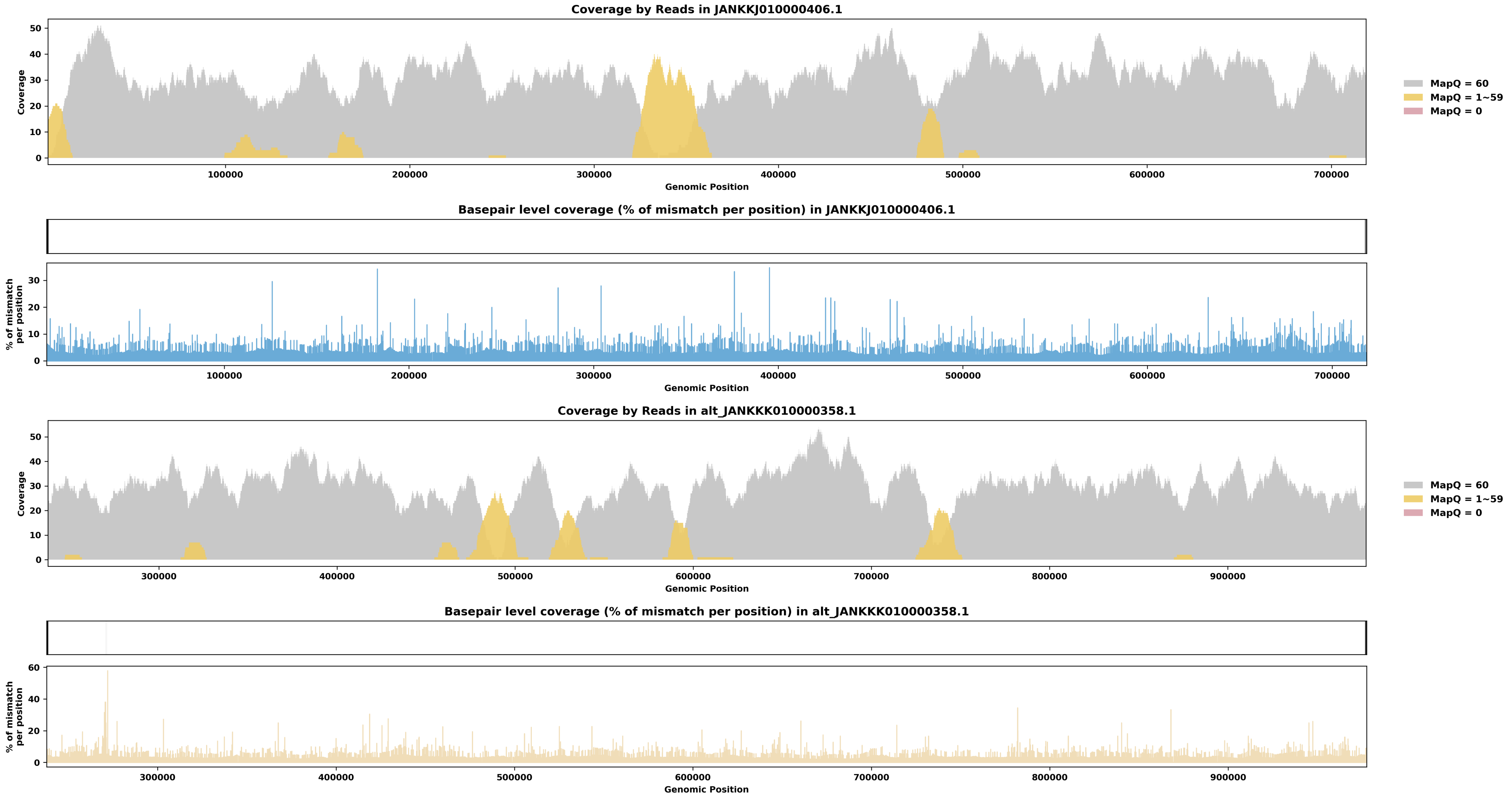

Species ID: mEleMax1

Common Name: Asiatic Elephant

Scientific Name: Elephas maximus

Assembly Type: Not Haplotype Resolved

Data Source: VGP

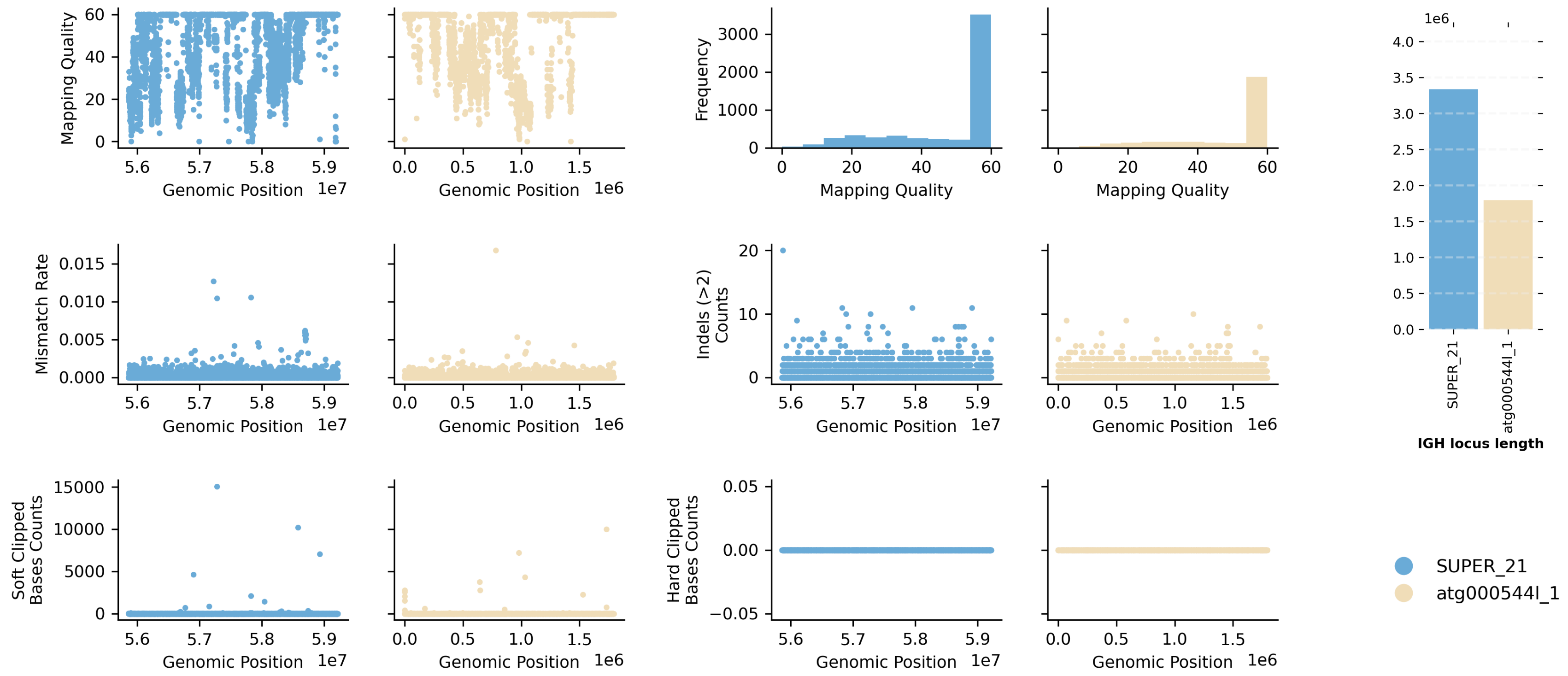


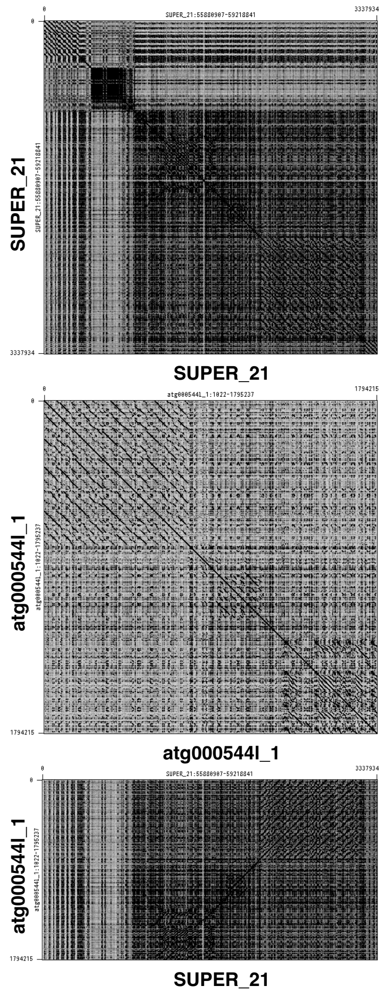


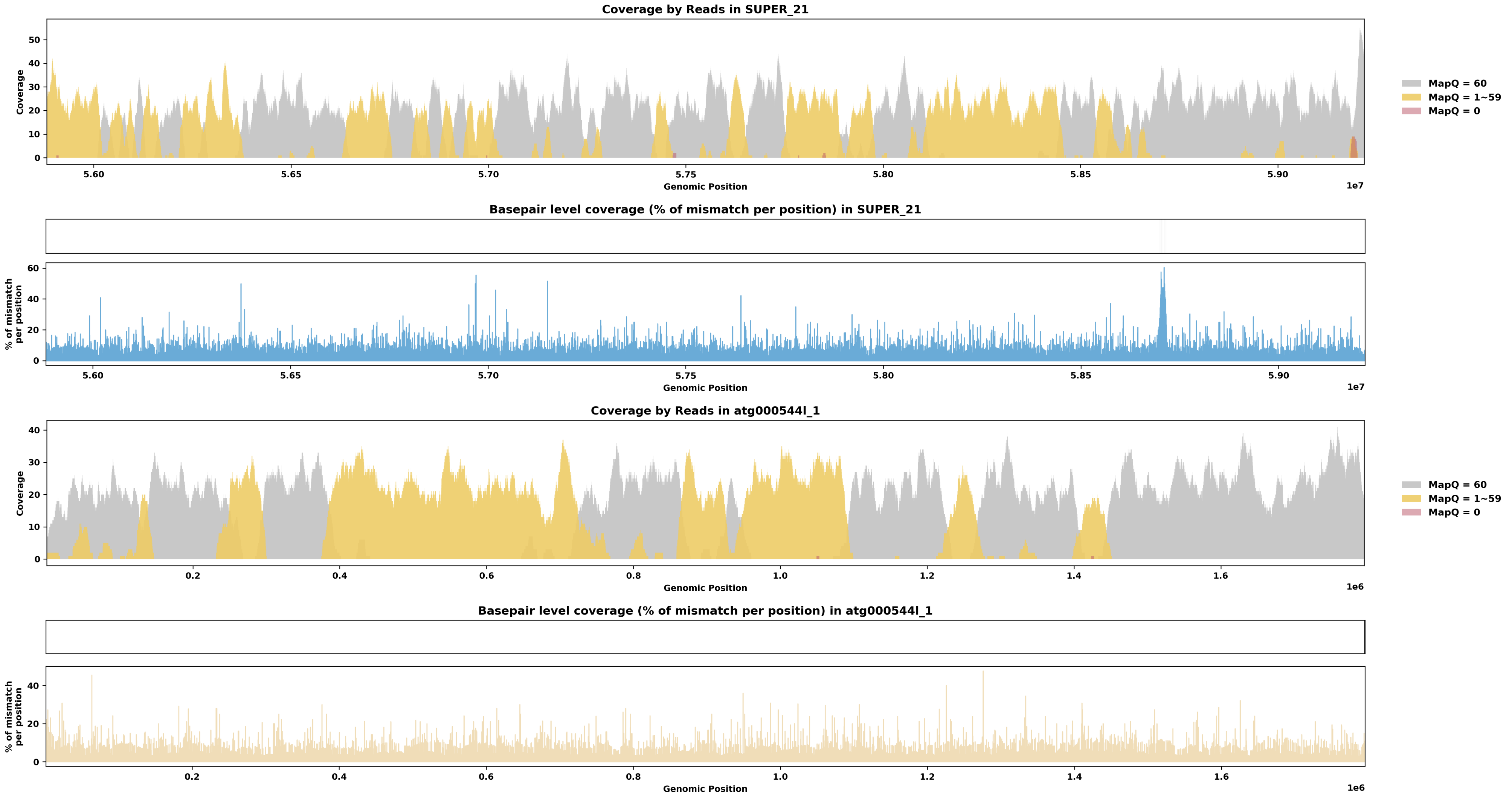

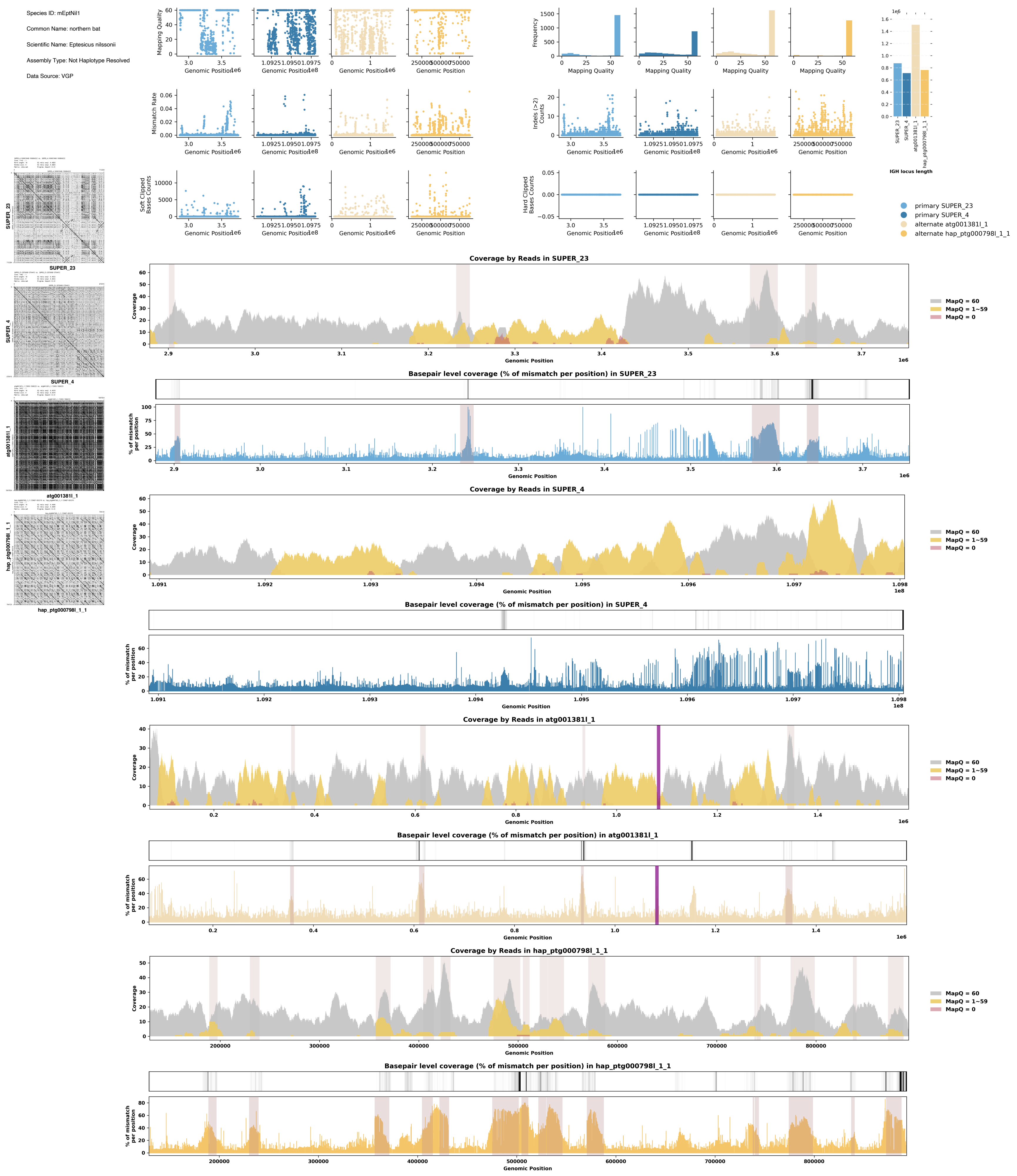
Species ID: mEptNil1
Common Name: northern bat
Scientific Name: Eptesicus nilssonii
Assembly Type: Not Haplotype Resolved
Data Source: VGP
Mapping Quality
Genomic Position
Frequency
Mismatch Rate
Indels (>2) Counts
Soft Clipped Bases Counts
Hard Clipped Bases Counts
IGH locus length
SUPER_23
SUPER_4
atg001381l_1
hap_ptg000798l_1_1
primary SUPER_23
primary SUPER_4
alternate atg001381l_1
alternate hap_ptg000798l_1_1
Coverage by Reads in SUPER_23
Coverage
MapQ = 60
MapQ = 1~59
MapQ = 0
Basepair level coverage (% of mismatch per position) in SUPER_23
% of mismatch per position
Coverage by Reads in SUPER_4
Basepair level coverage (% of mismatch per position) in SUPER_4
Coverage by Reads in atg001381l_1
Basepair level coverage (% of mismatch per position) in atg001381l_1
Coverage by Reads in hap_ptg000798l_1_1
Basepair level coverage (% of mismatch per position) in hap_ptg000798l_1_1

Species ID: mEriEur2

Common Name: western European hedgehog

Scientific Name: Erinaceus europaeus

Assembly Type: Not Haplotype Resolved

Data Source: VGP

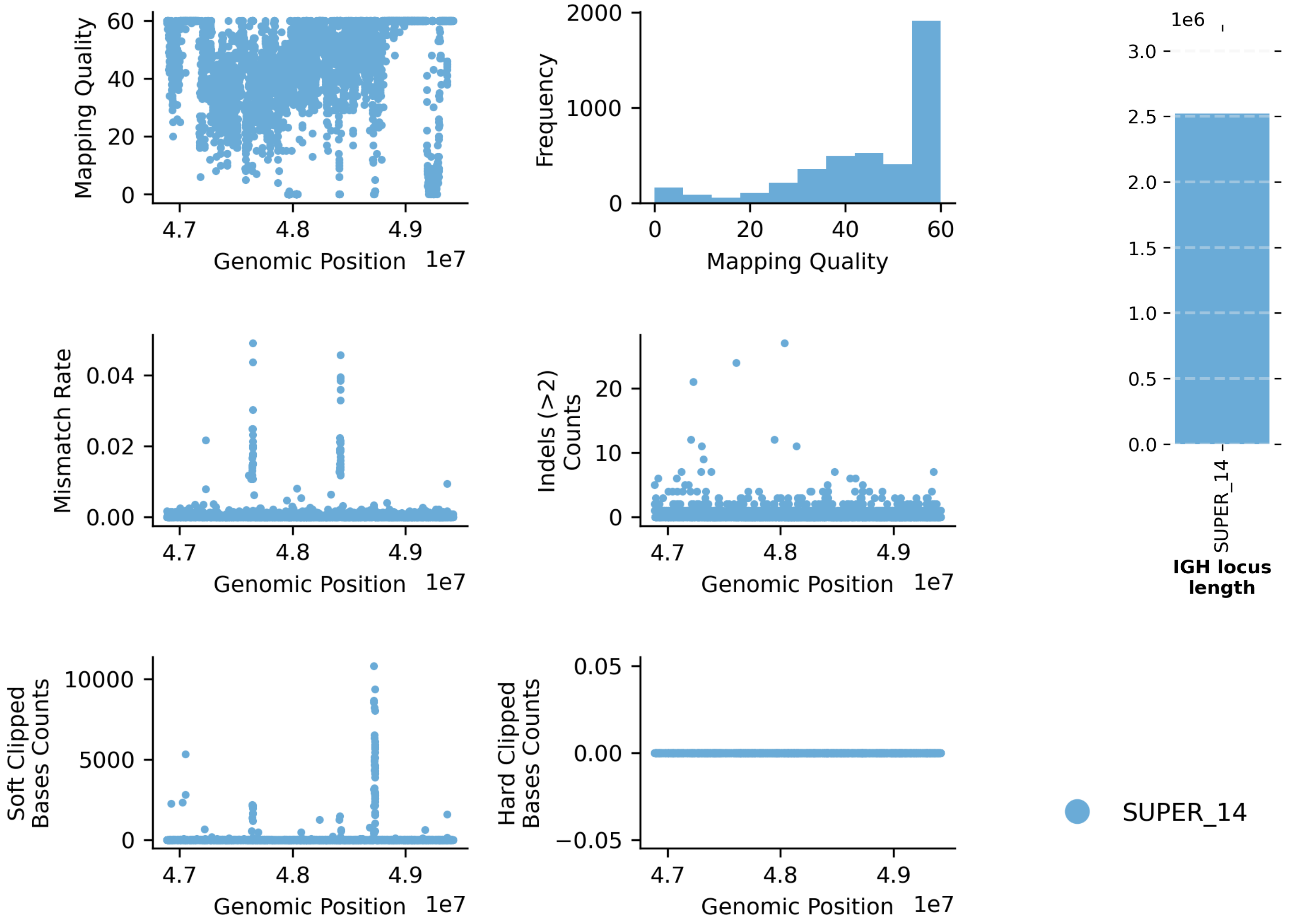


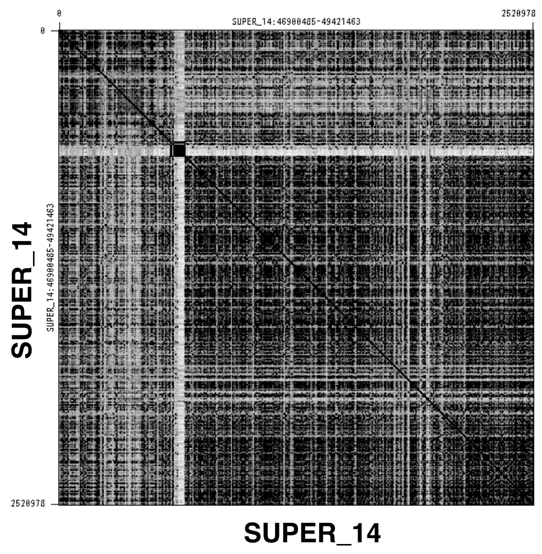


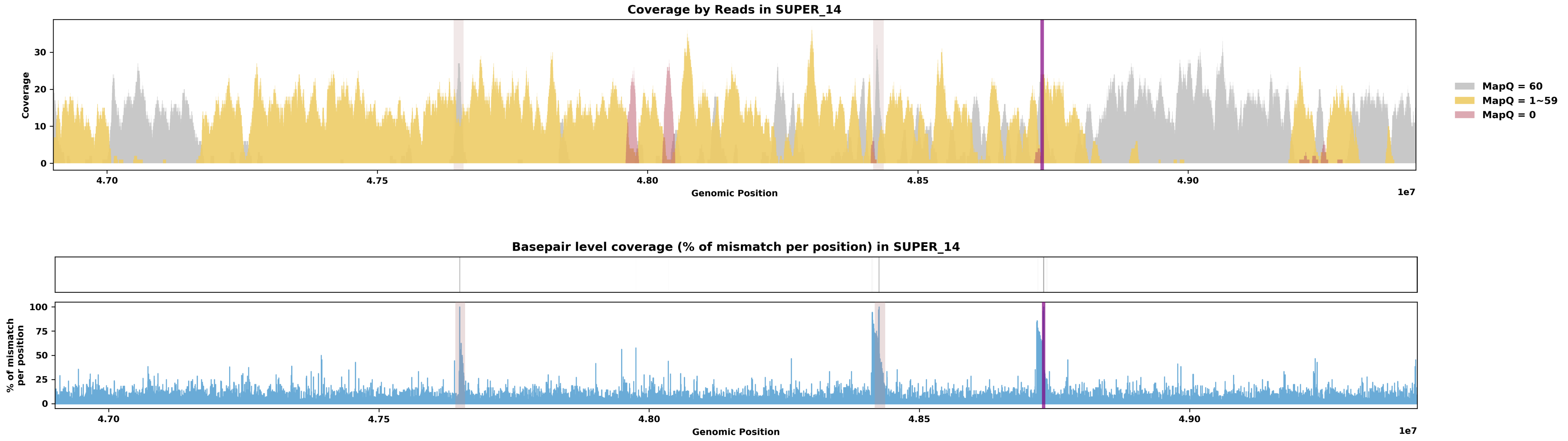

Species ID: mEscRob2

Common Name: grey whale

Scientific Name: Eschrichtius robustus

Assembly Type: Not Haplotype Resolved

Data Source: VGP

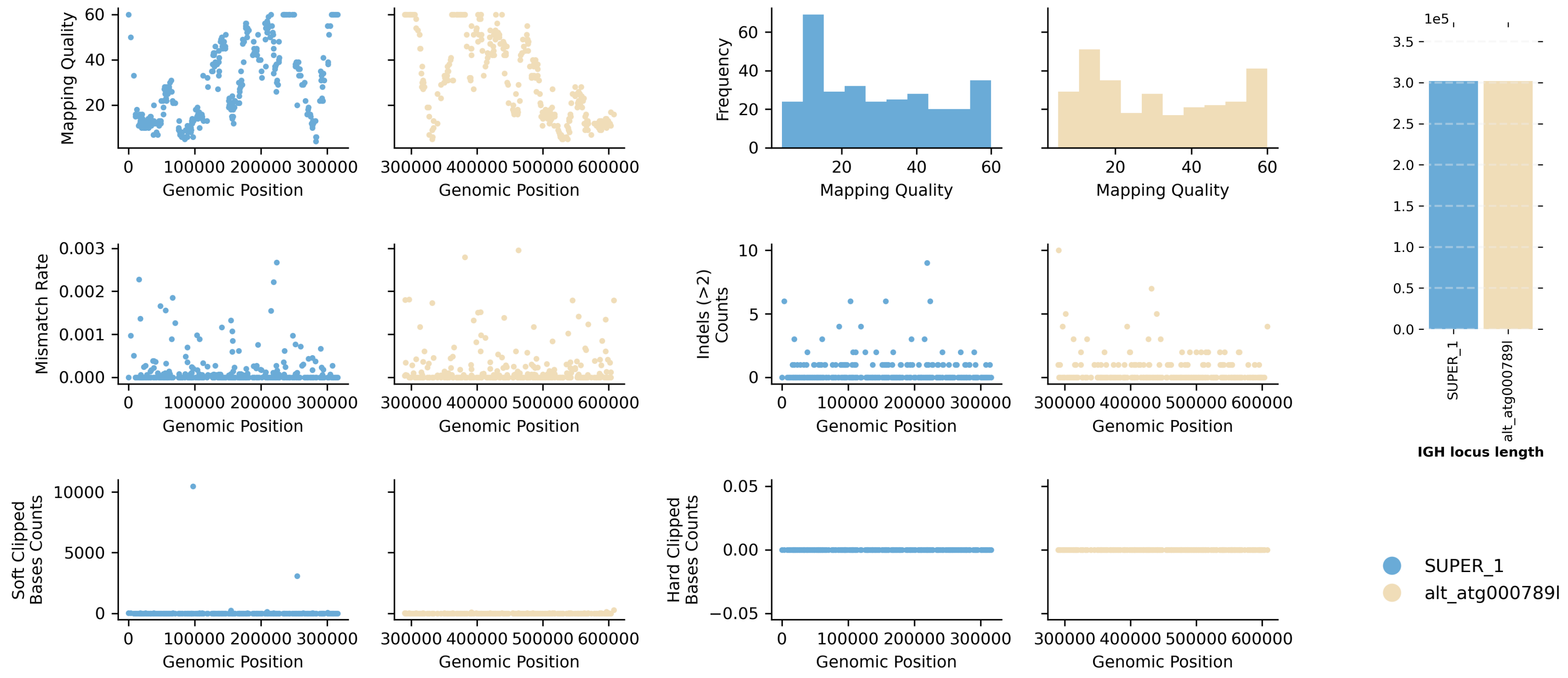


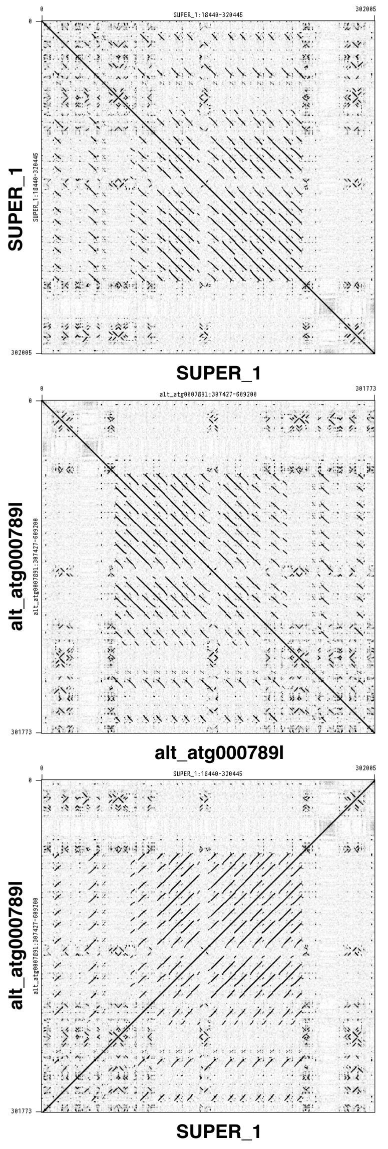


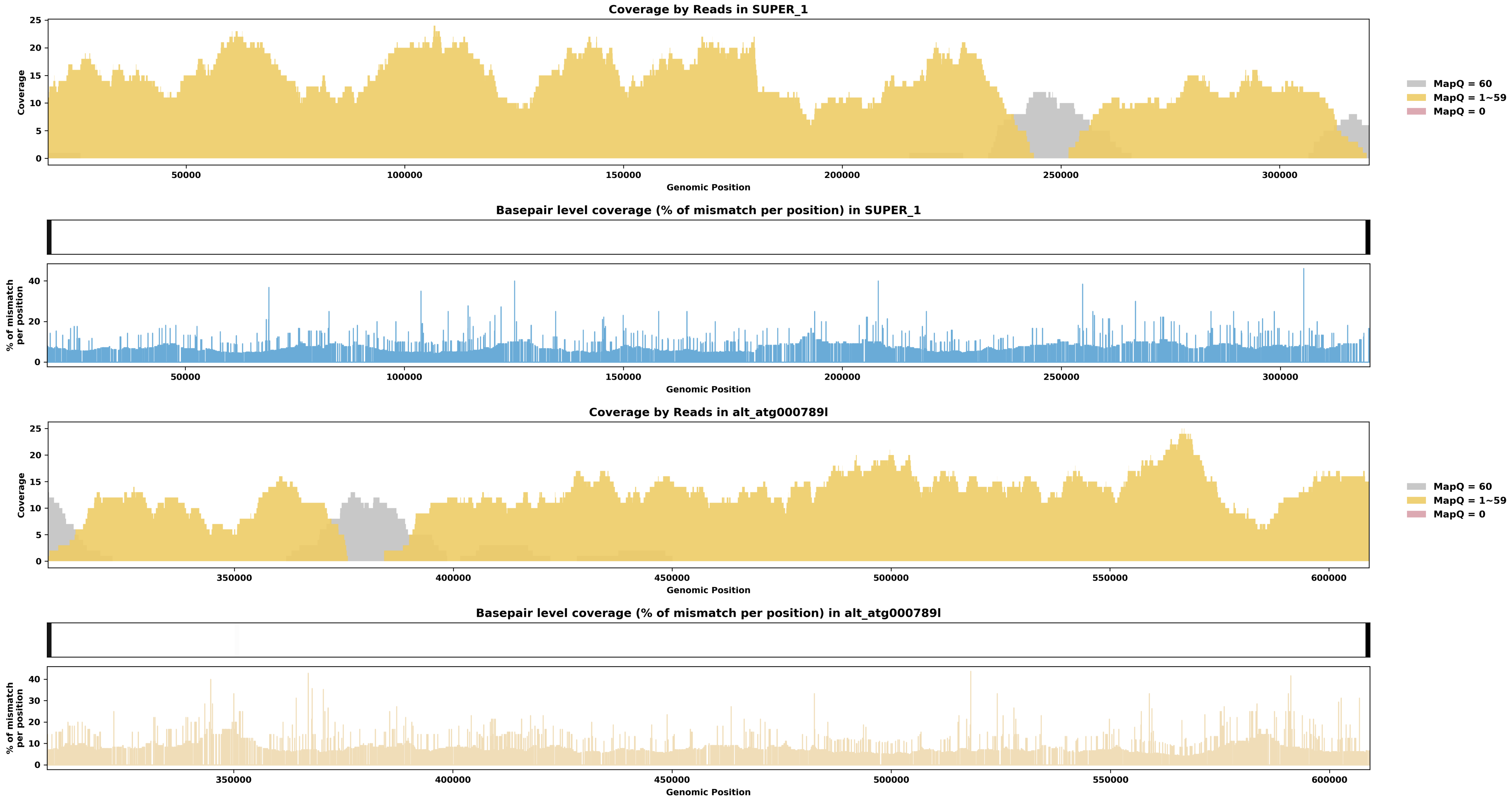

Species ID: mEubGla1

Common Name: North Atlantic right whale

Scientific Name: Eubalaena glacialis

Assembly Type: Haplotype Resolved

Data Source: VGP

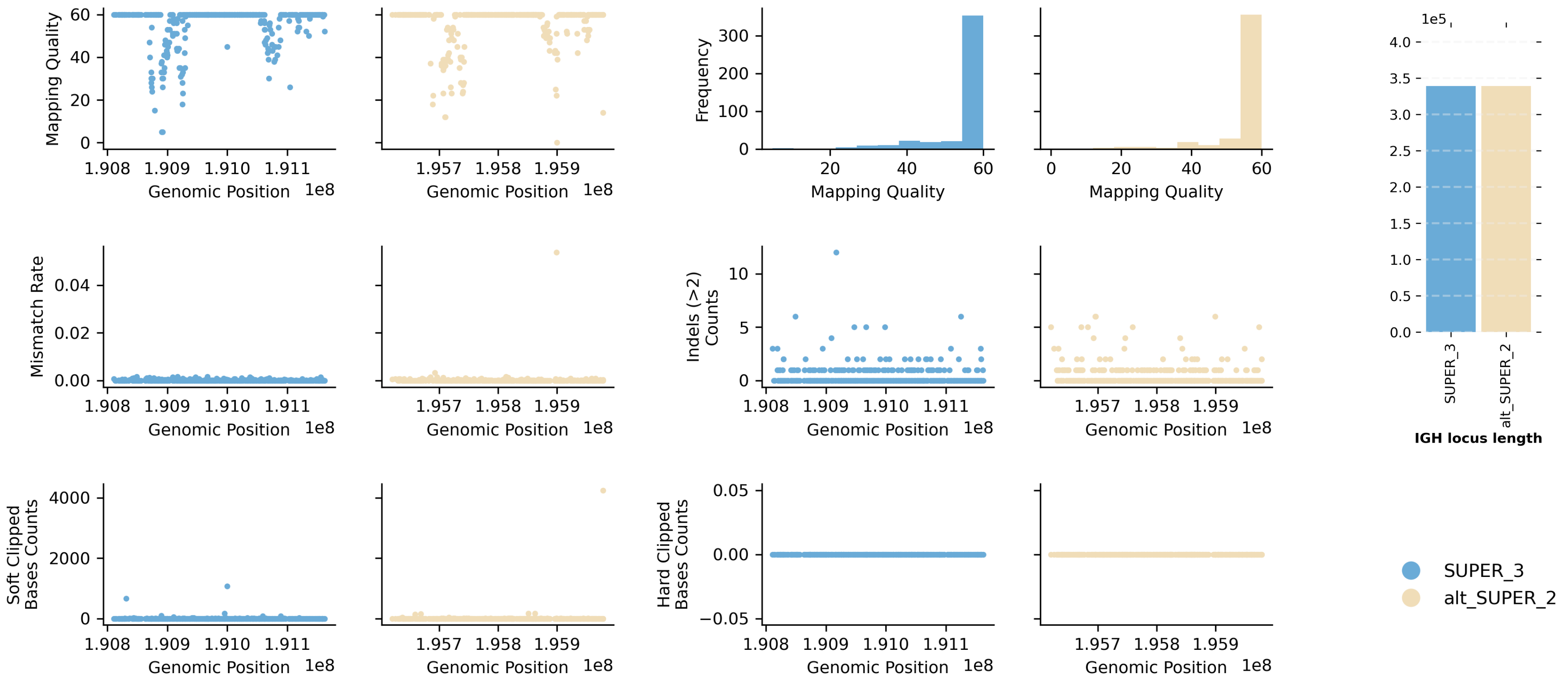


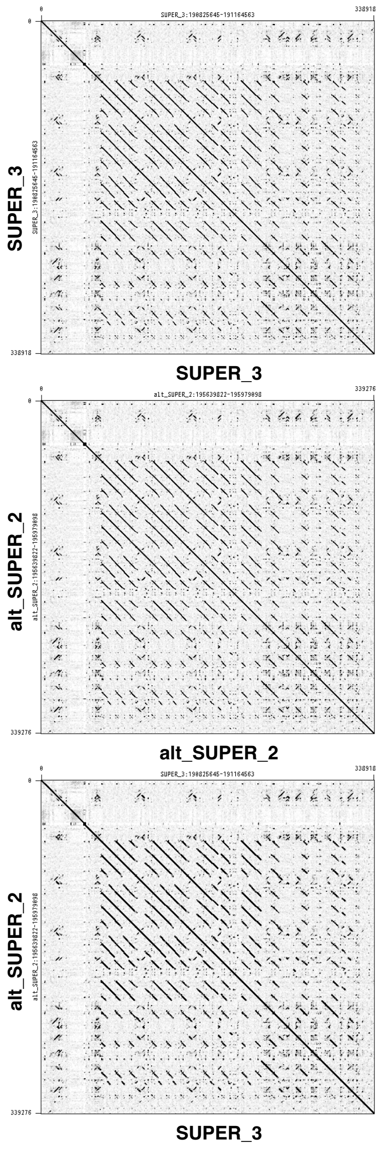


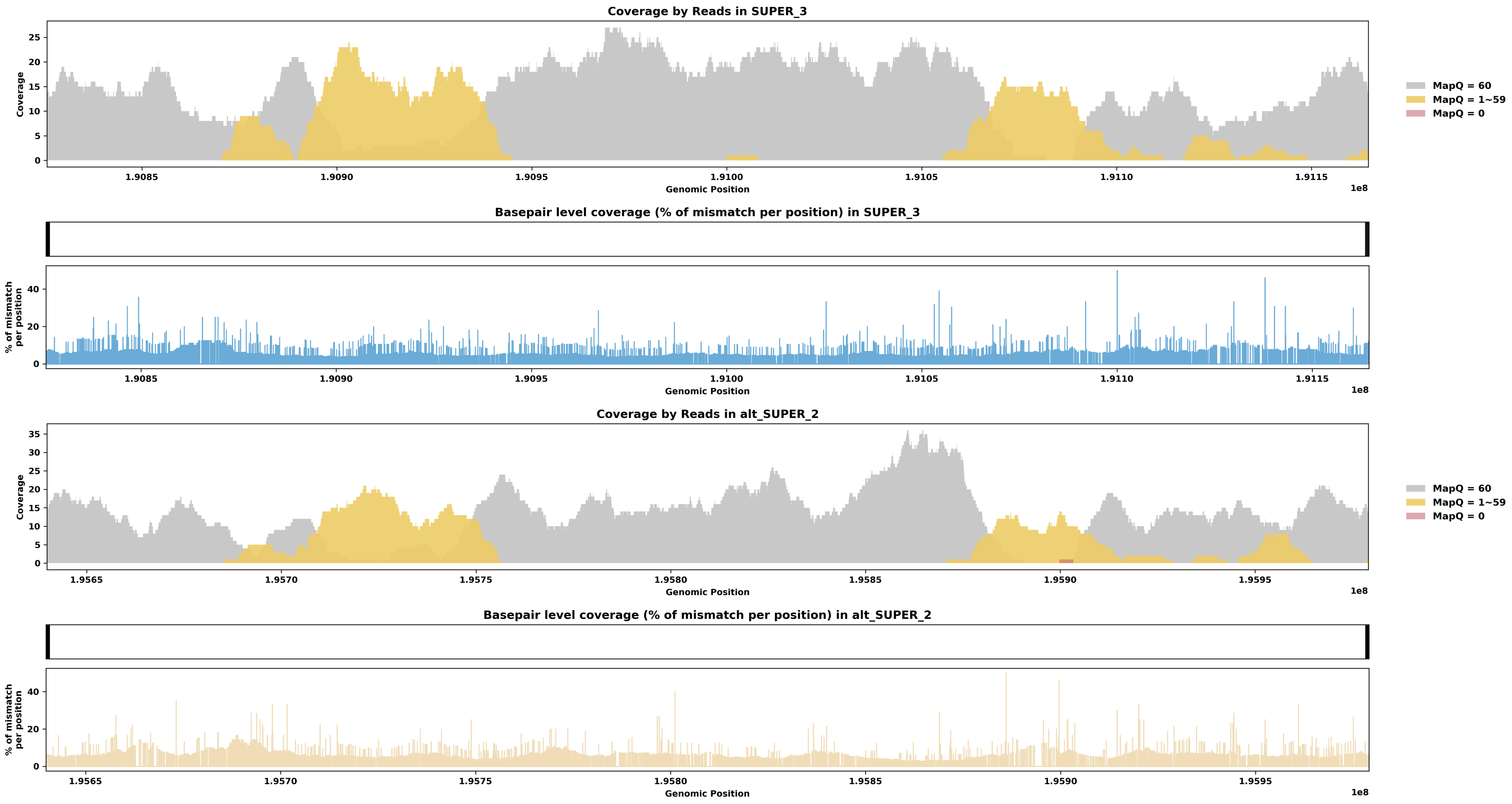

Species ID: mGloMel1

Common Name: long-finned pilot whale

Scientific Name: Globicephala melas

Assembly Type: Not Haplotype Resolved

Data Source: VGP

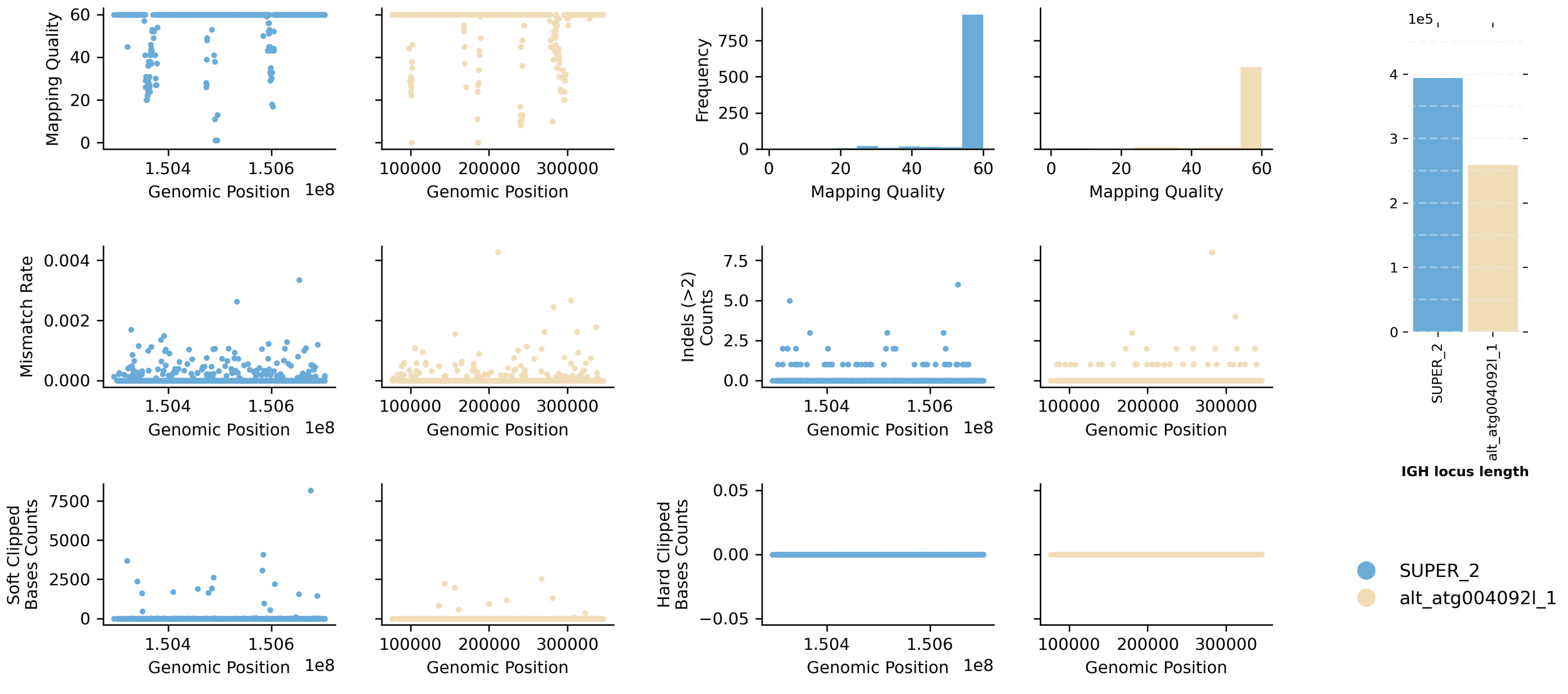


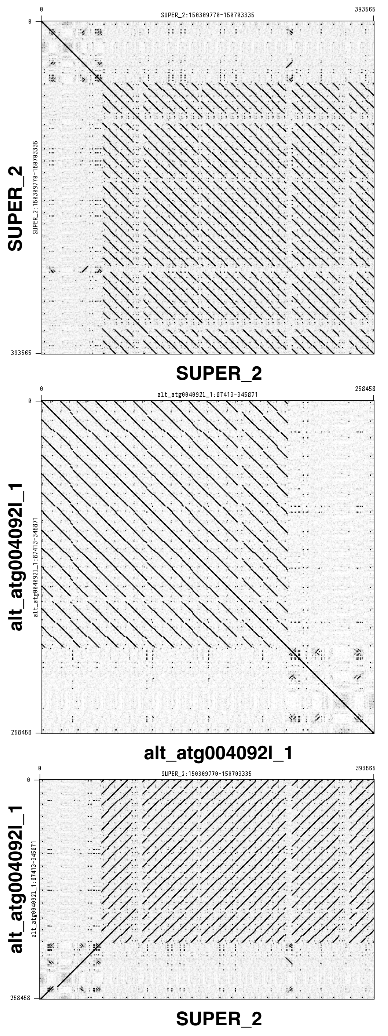


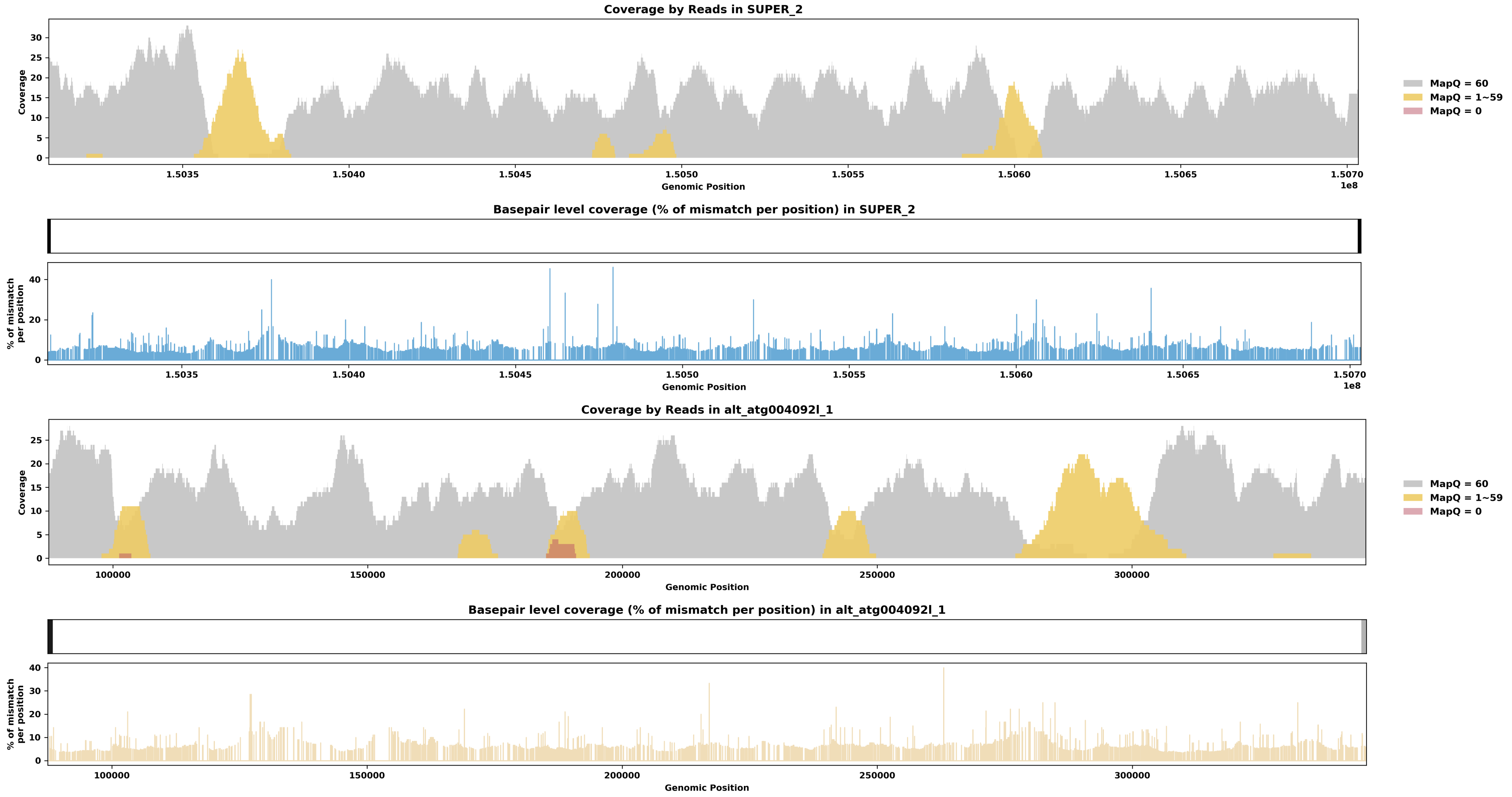

Species ID: mGorGor1

Common Name: Gorilla

Scientific Name: Gorilla_gorilla

Assembly Type: Haplotype Resolved

Data Source: T2T Primate

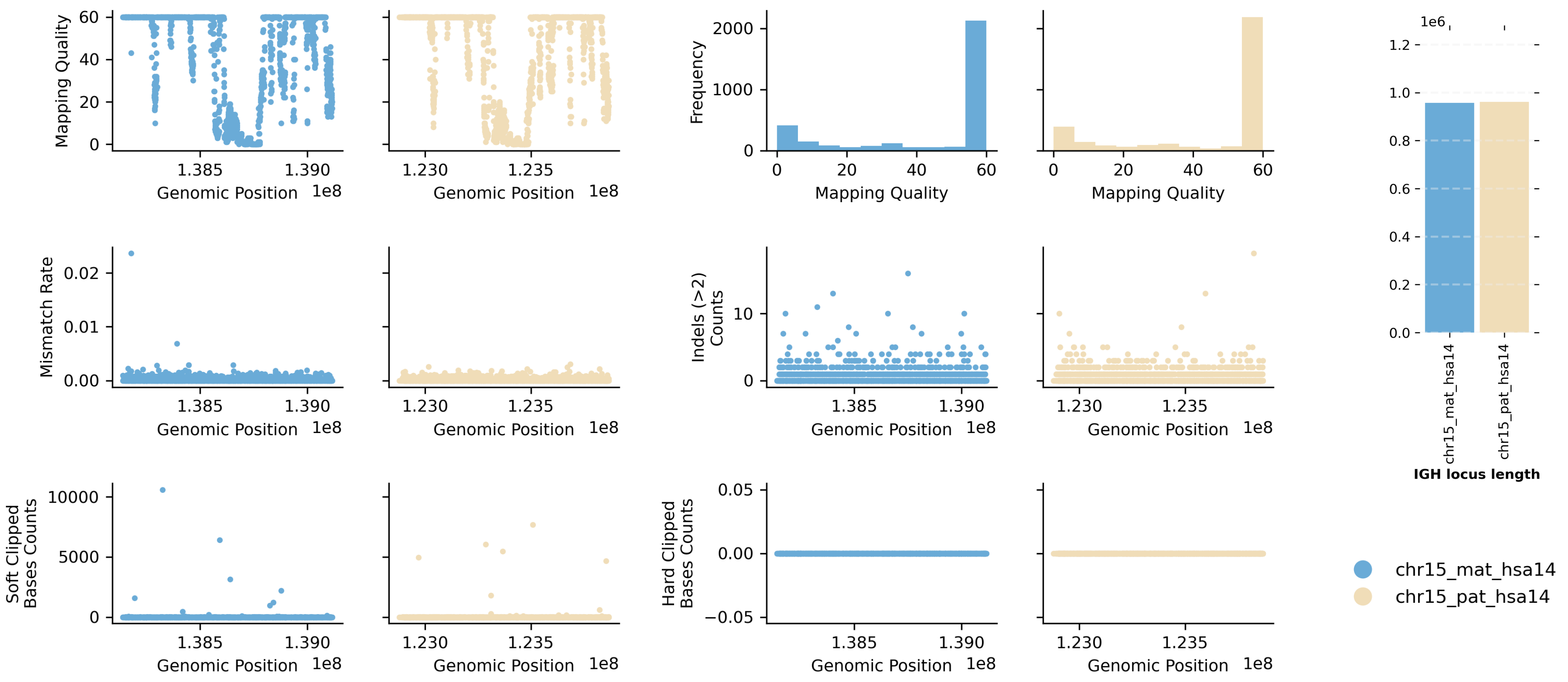


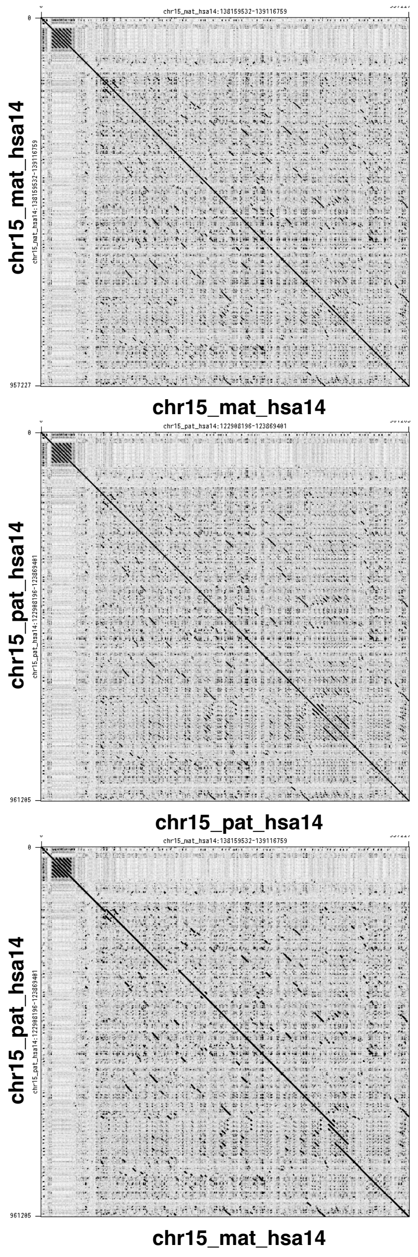


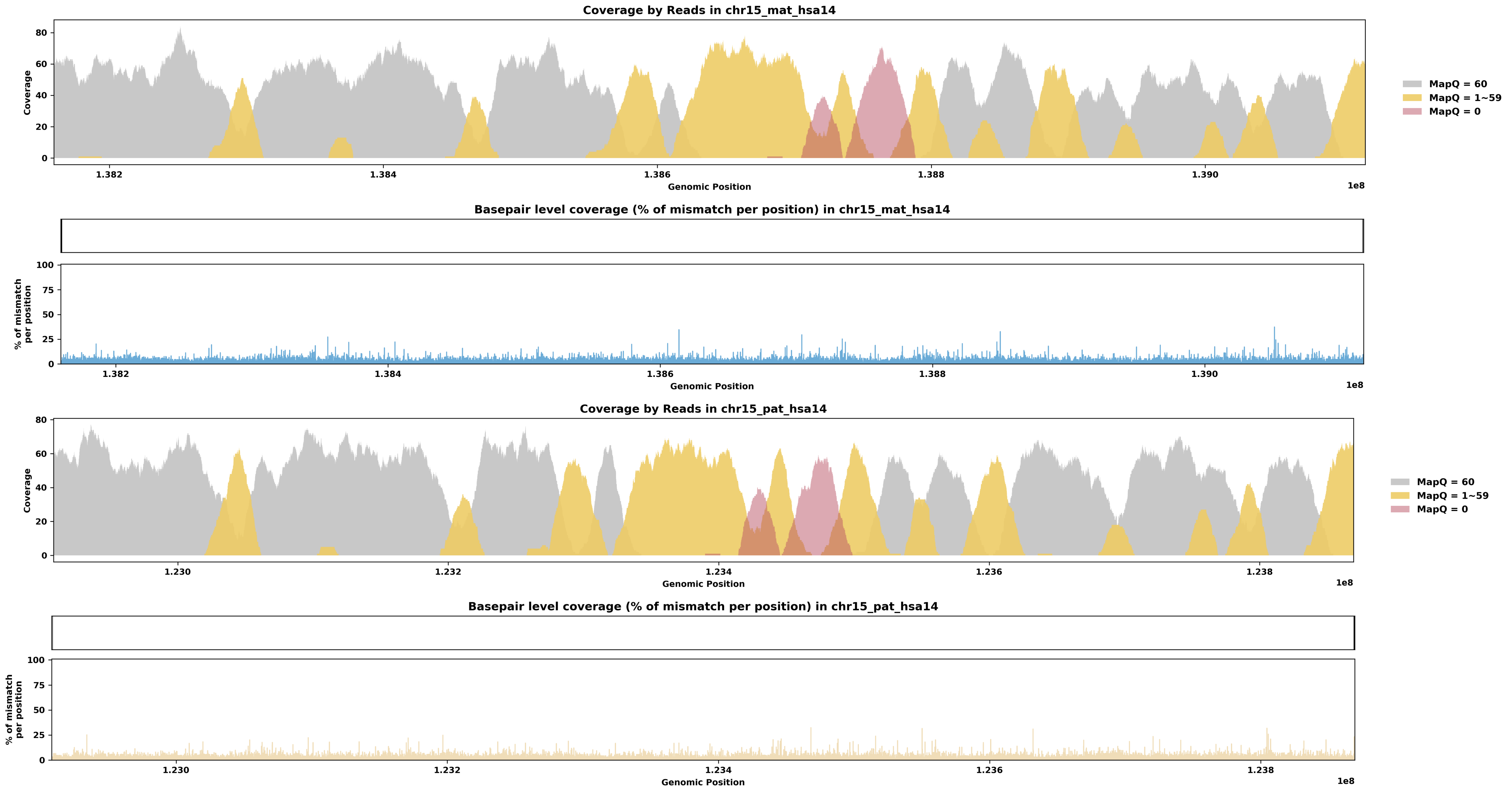

Species ID: mHetBru1

Common Name: Yellow-spotted hyrax

Scientific Name: Heterohyrax brucei

Assembly Type: Not Haplotype Resolved

Data Source: VGP

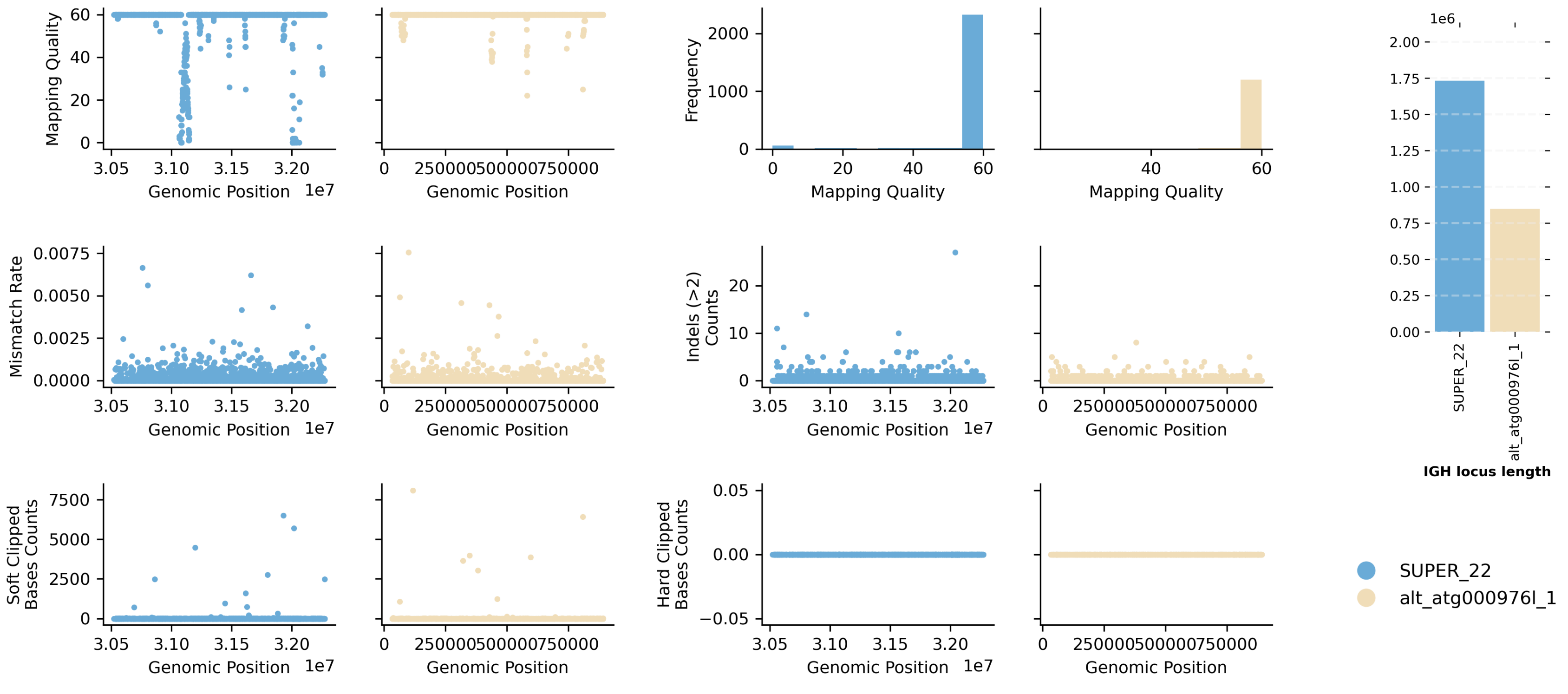


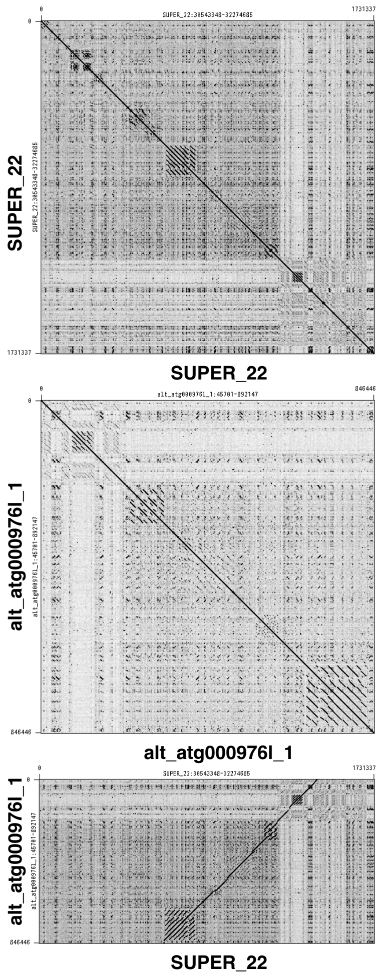


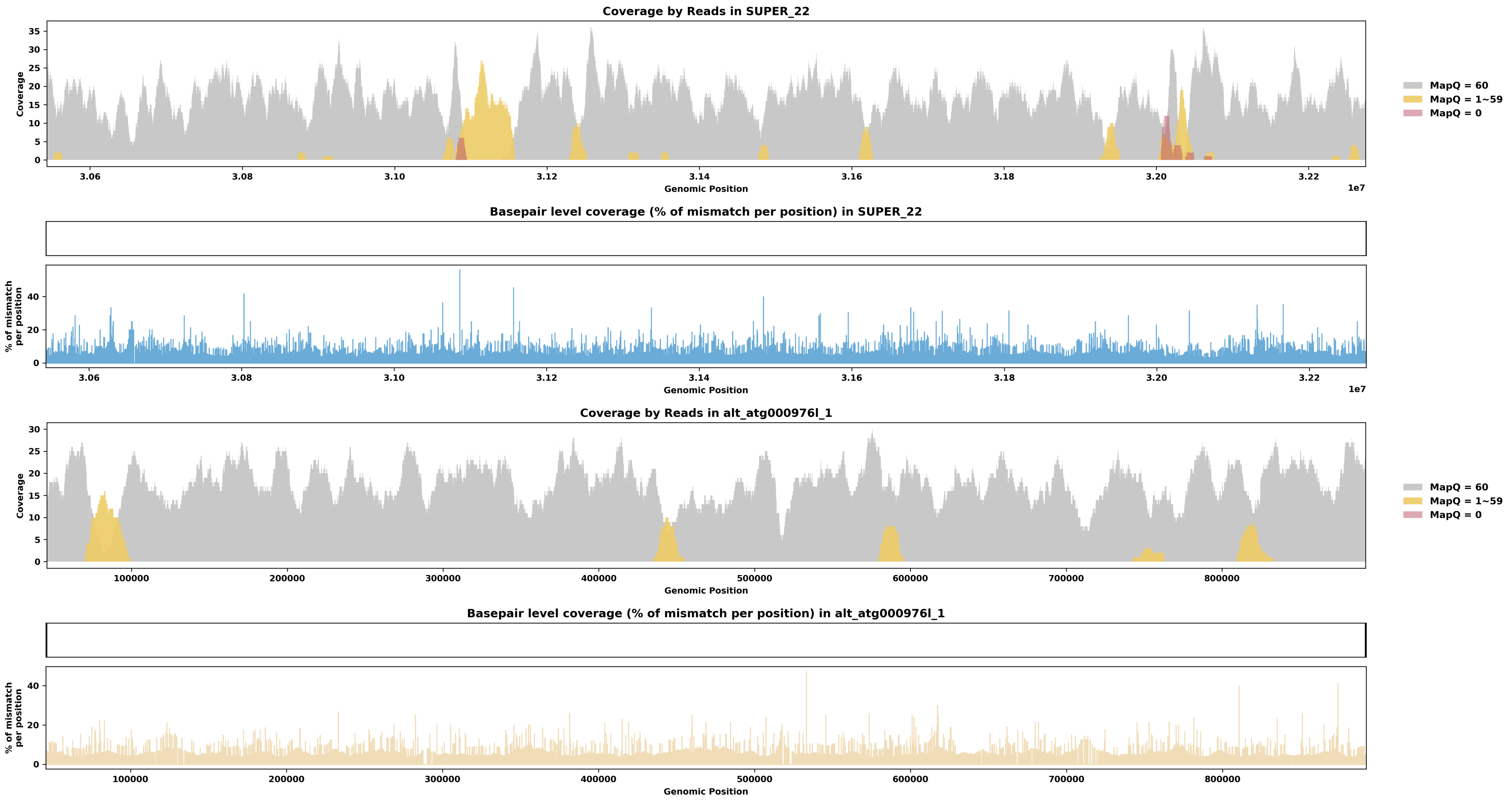

Species ID: mHipAmp2

Common Name: hippopotamus

Scientific Name: Hippopotamus amphibius kiboko

Assembly Type: Haplotype Resolved

Data Source: VGP

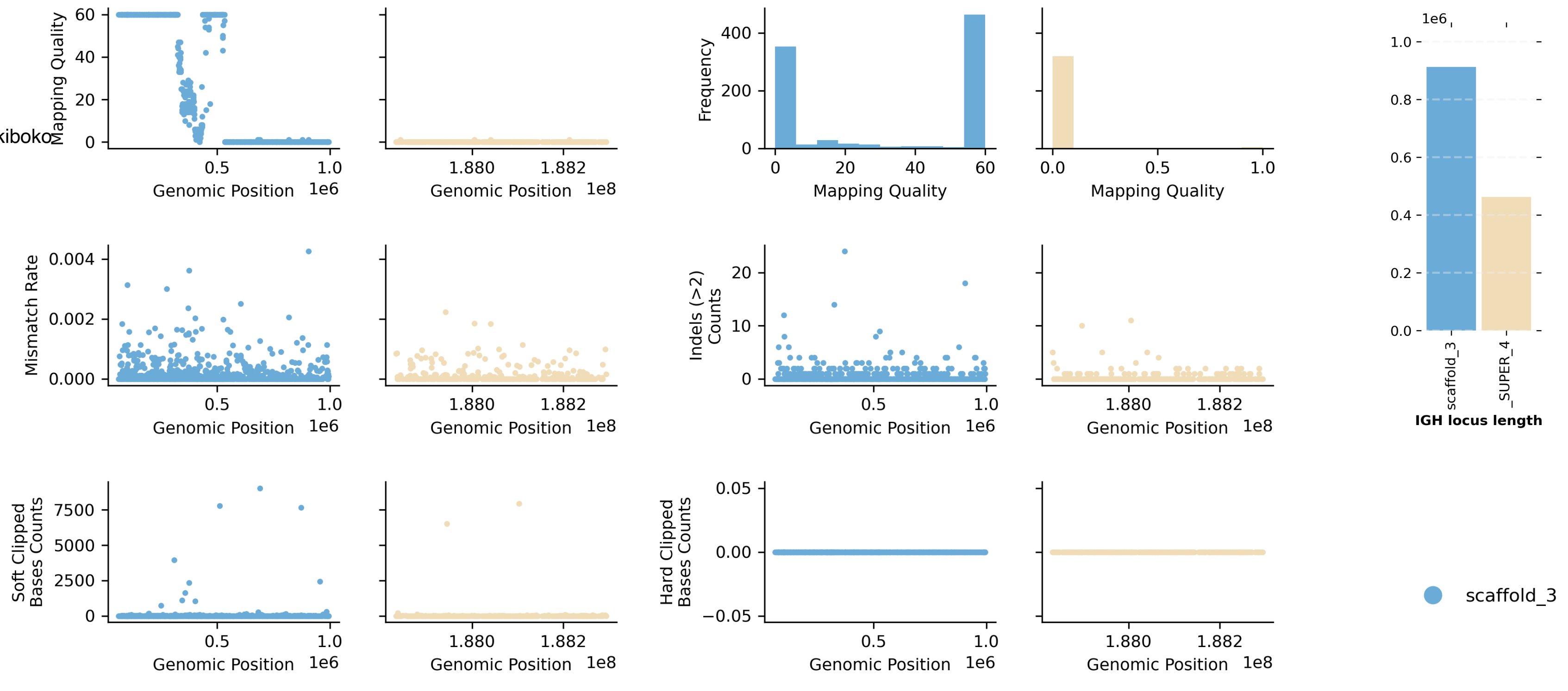


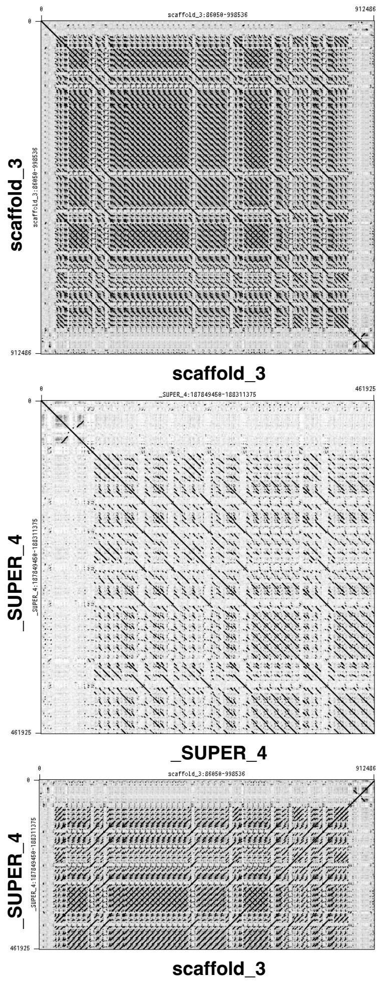


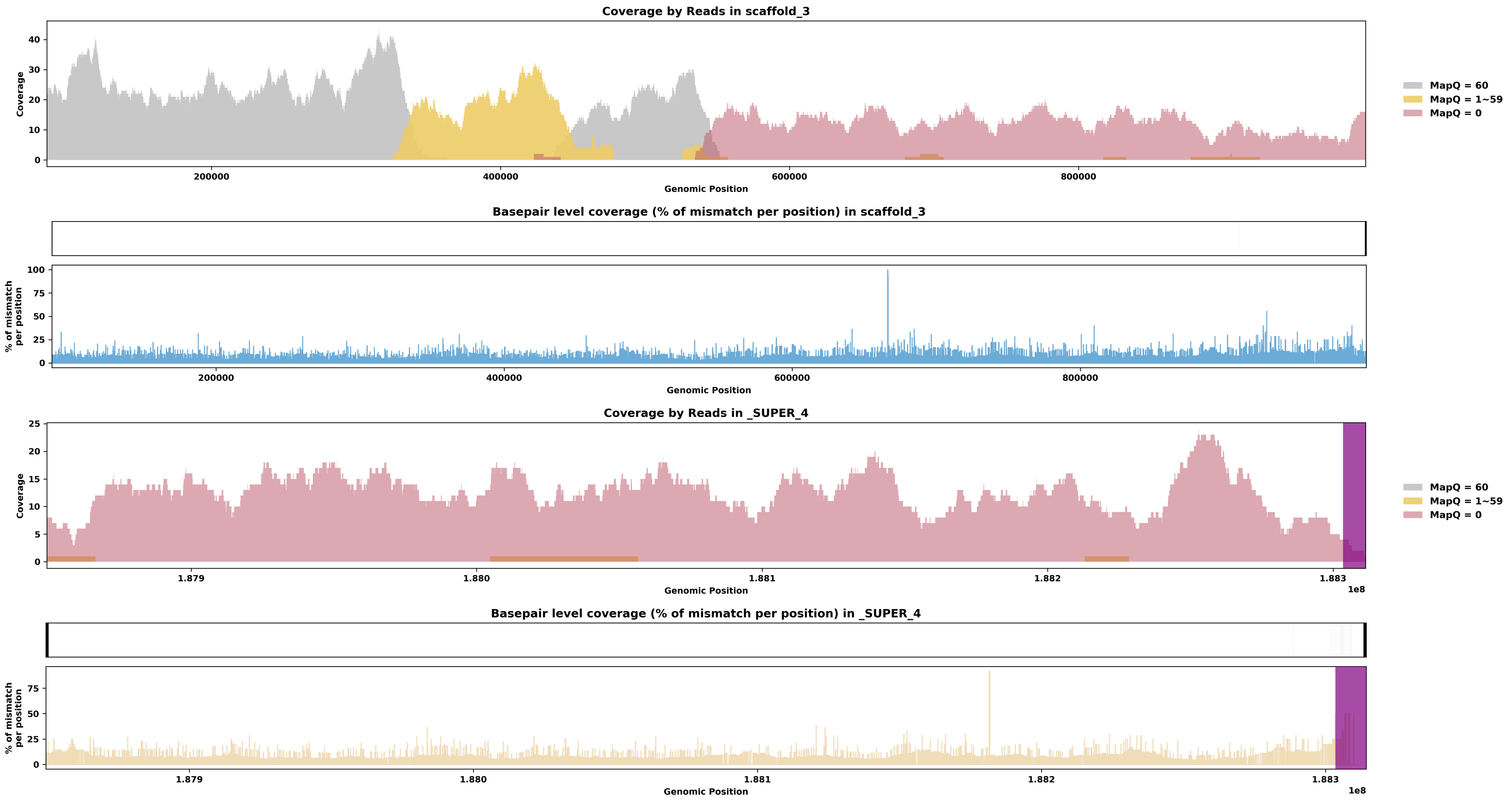

Species ID: mHypAmp2

Common Name: northern bottlenose whale

Scientific Name: Hyperoodon ampullatus

Assembly Type: Not Haplotype Resolved

Data Source: VGP

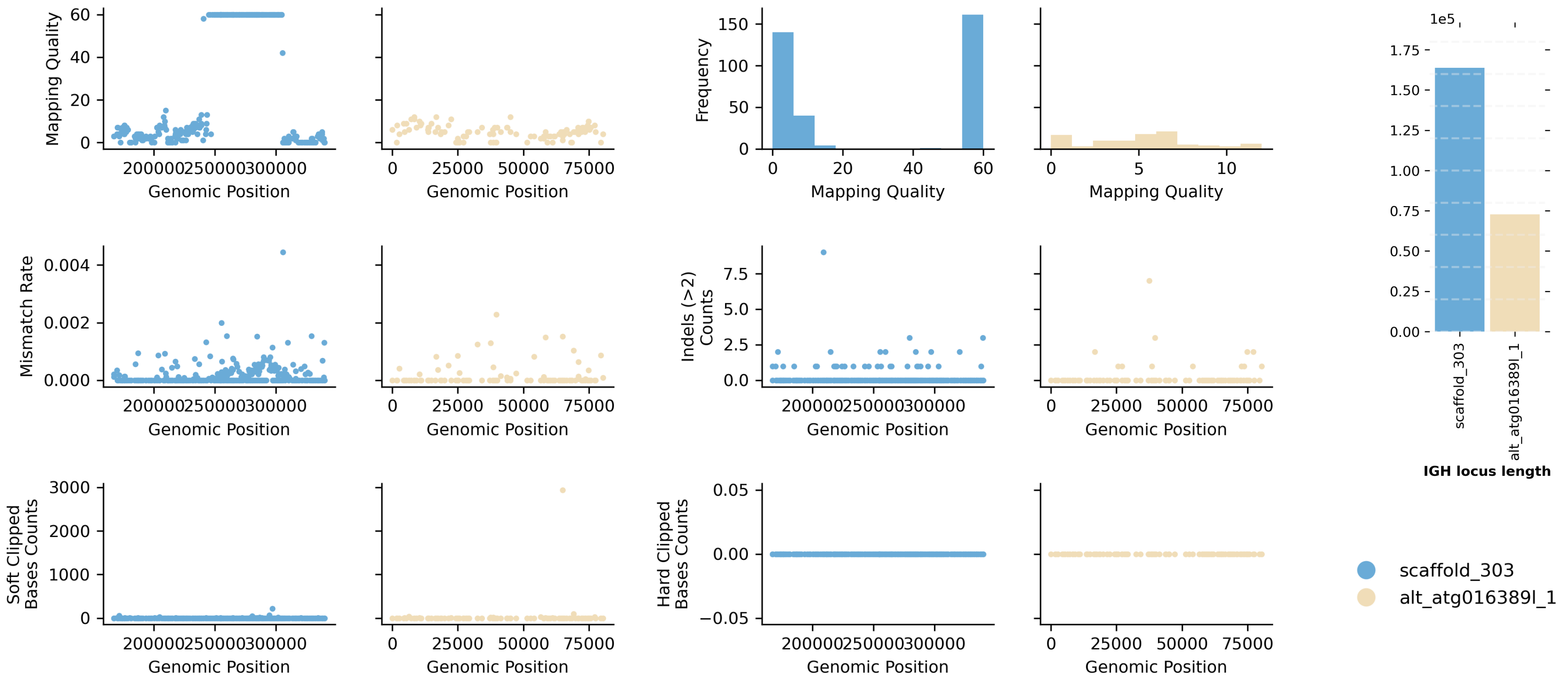


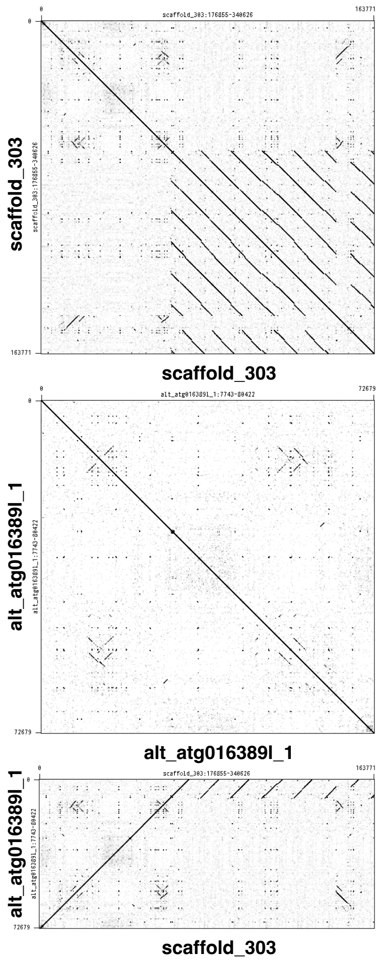


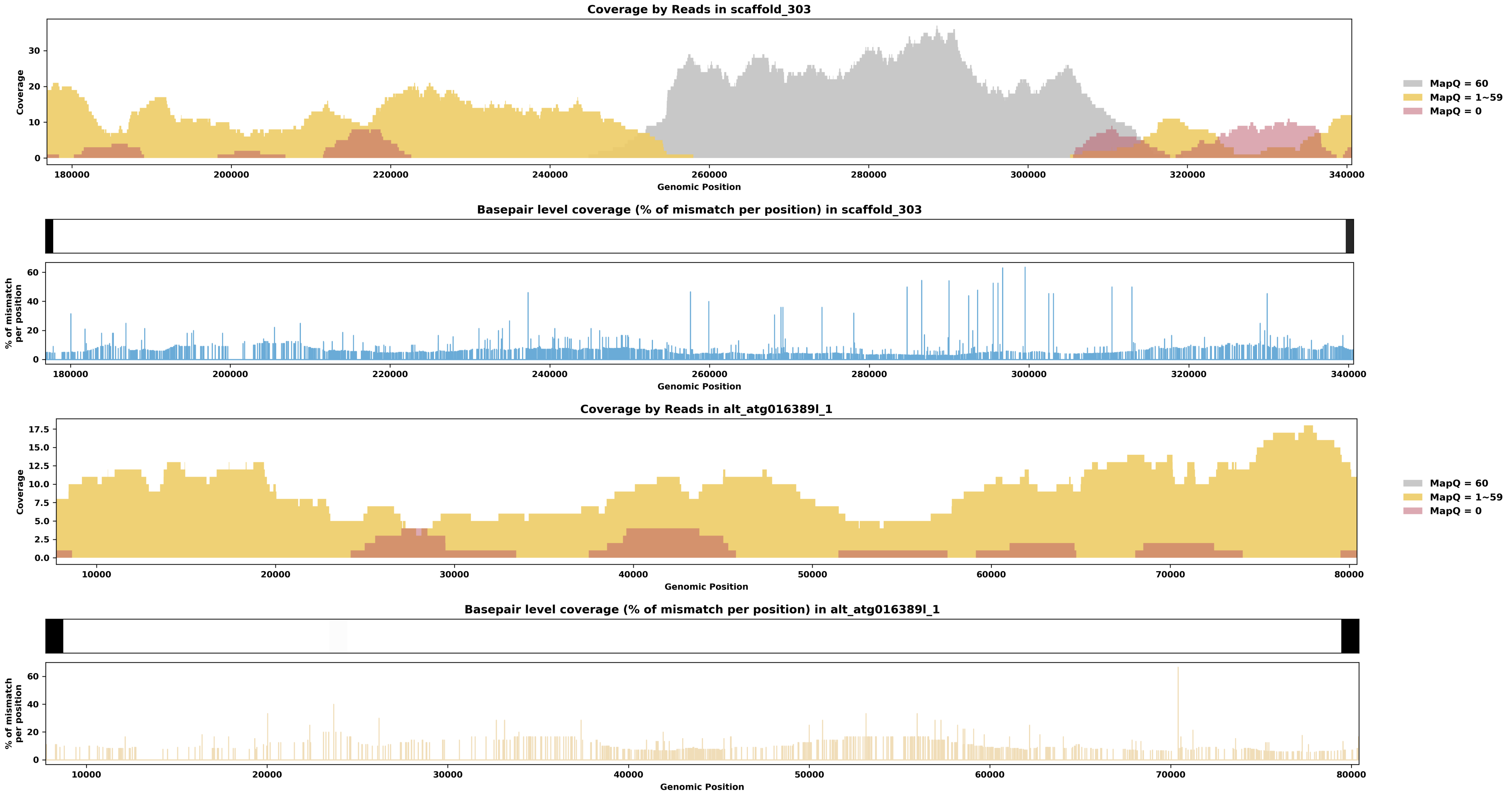

Species ID: mLagAlb1

Common Name: white-beaked dolphin

Scientific Name: Lagenorhynchus albirostris

Assembly Type: Not Haplotype Resolved

Data Source: VGP

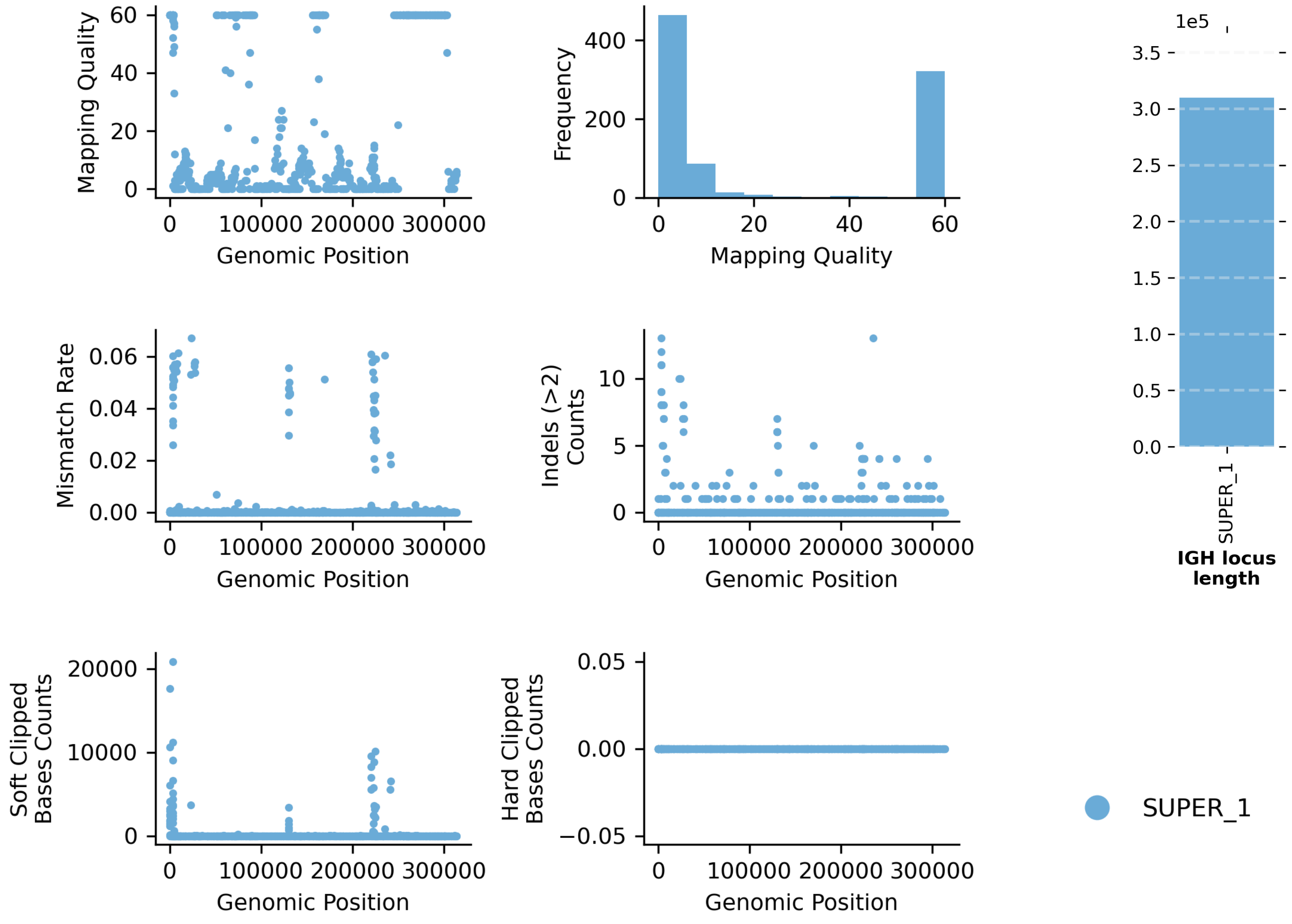


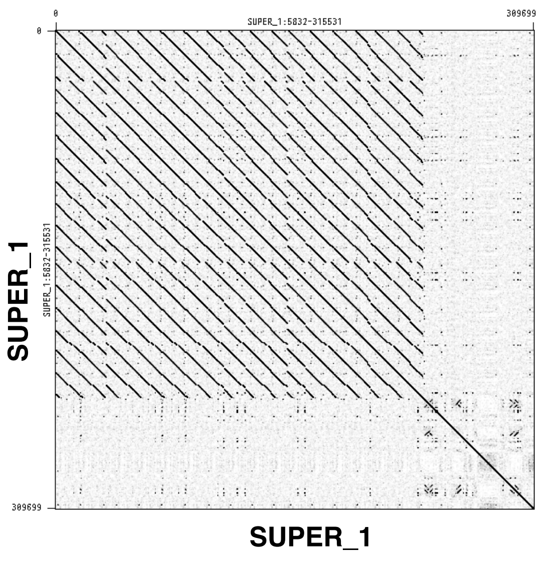


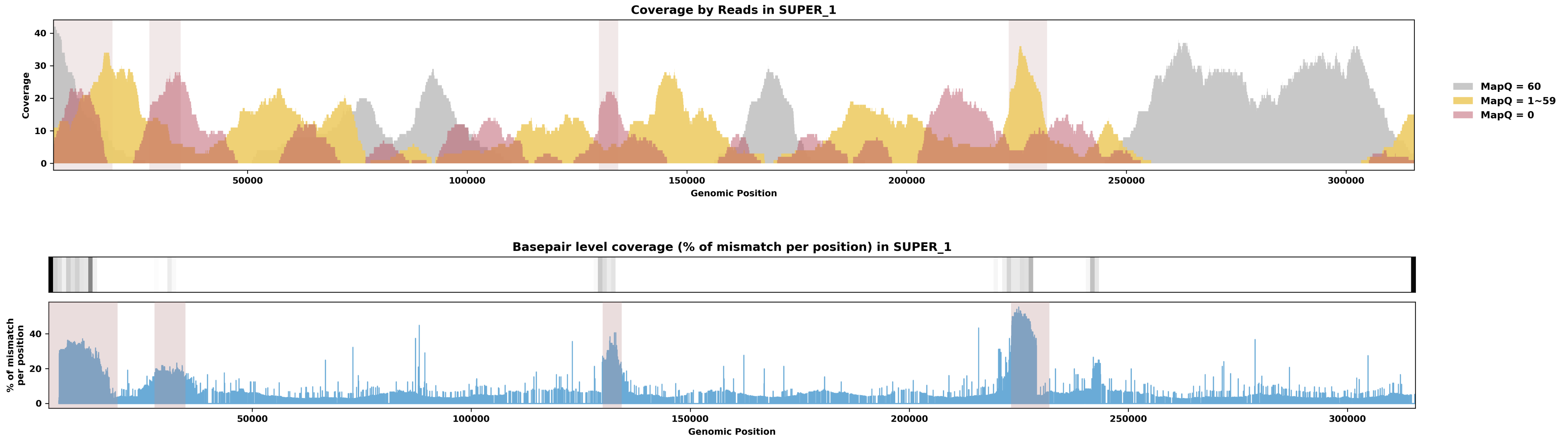

Species ID: mLemCat1

Common Name: Ring-tailed lemur

Scientific Name: Lemur catta

Assembly Type: Not Haplotype Resolved

Data Source: VGP

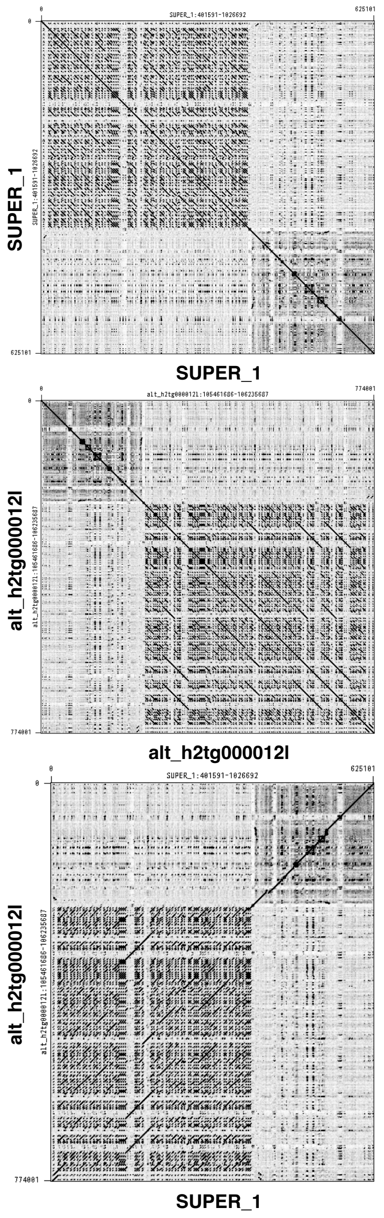


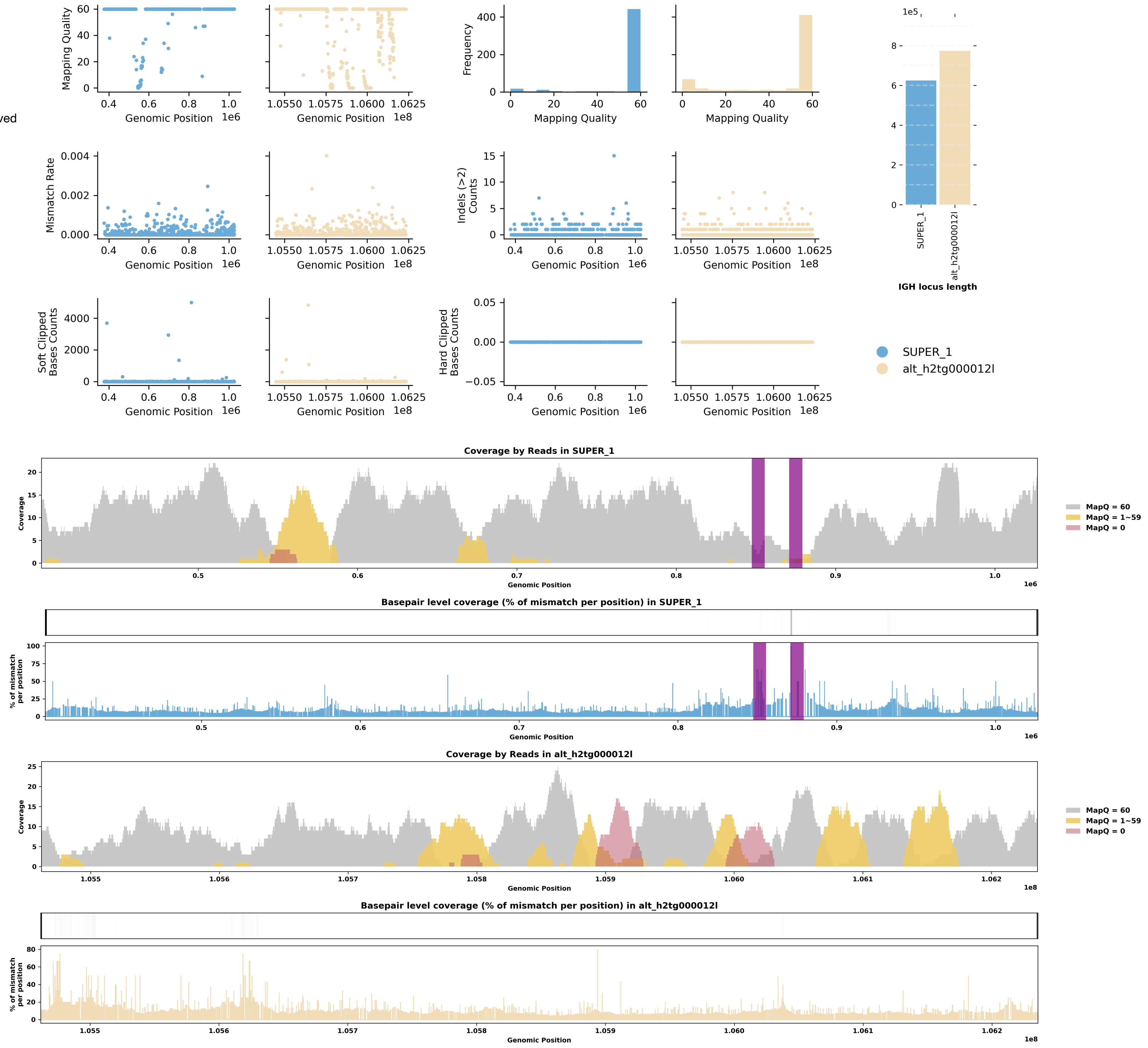

Species ID: mLynRuf1

Common Name: Bobcat

Scientific Name: Lynx rufus

Assembly Type: Not Haplotype Resolved

Data Source: CCGP

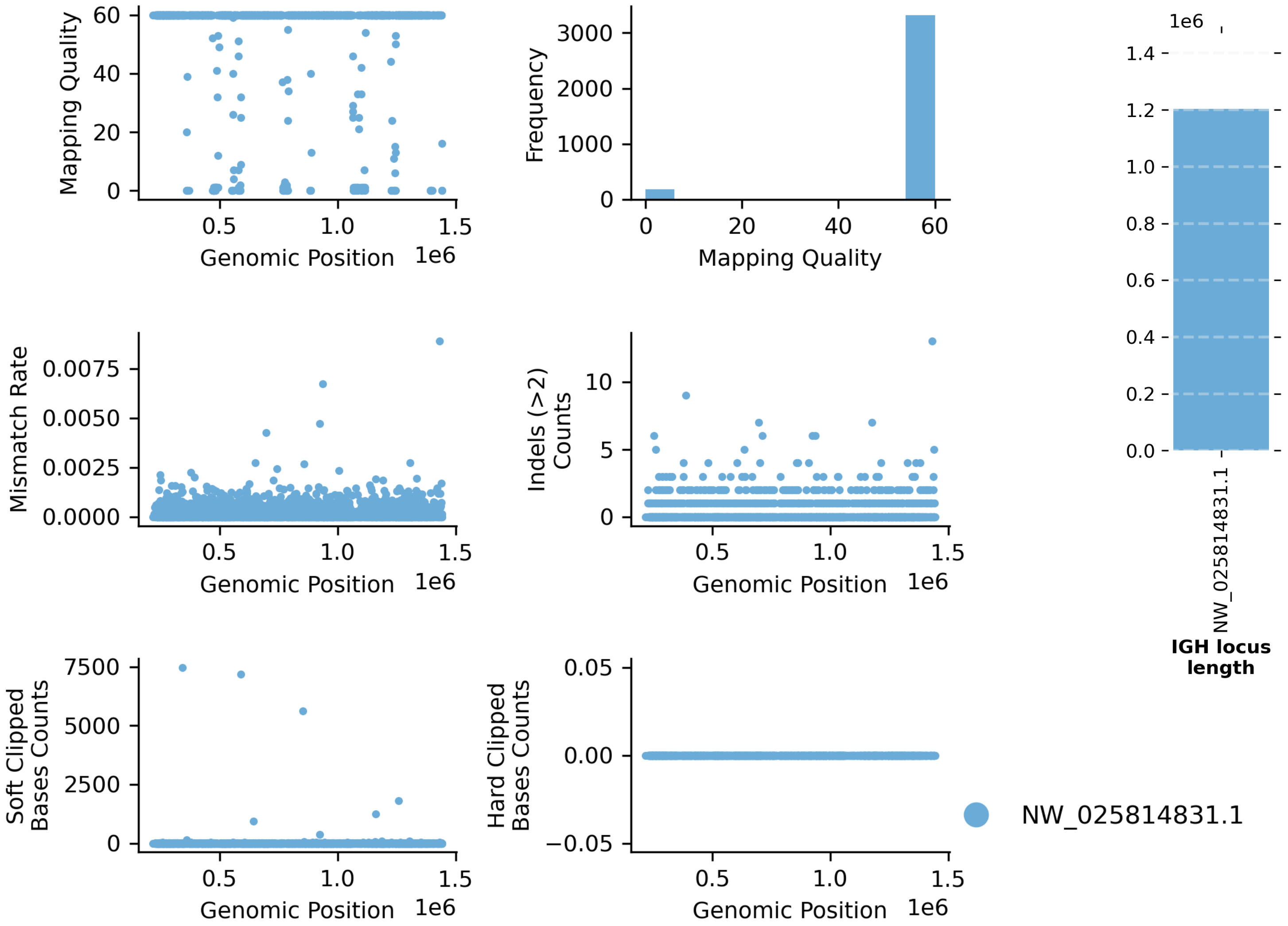


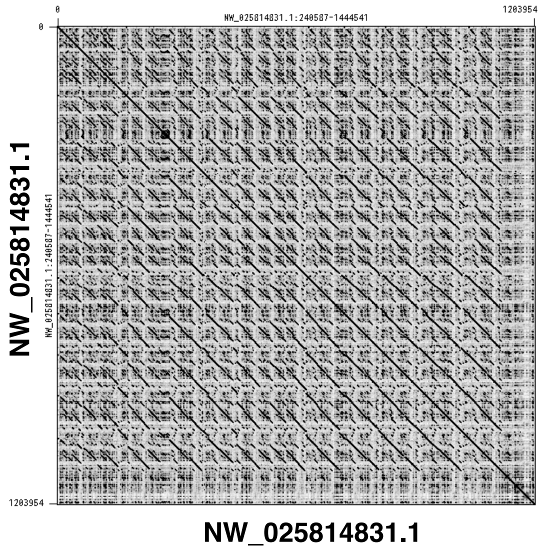


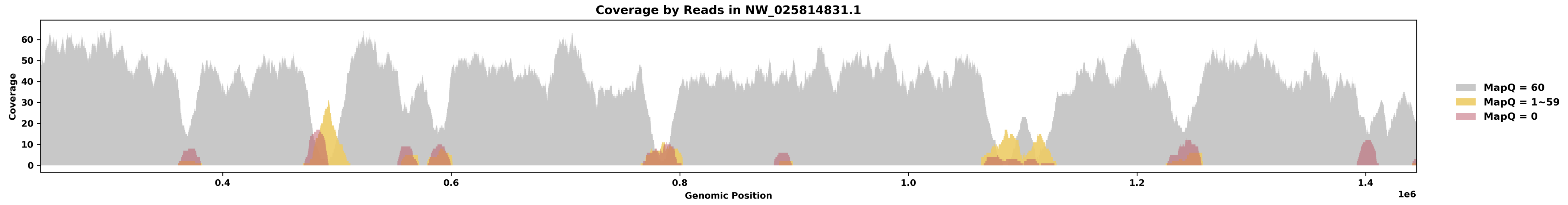


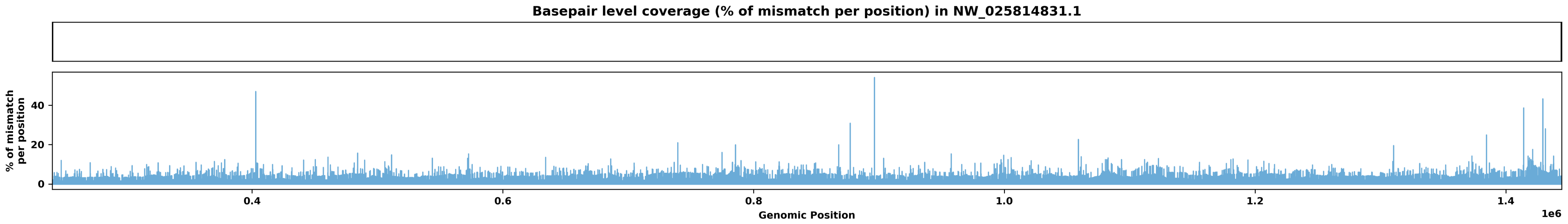

Species ID: mMacEug1

Common Name: tammar wallaby

Scientific Name: Macropus eugenii

Assembly Type: Not Haplotype Resolved

Data Source: VGP

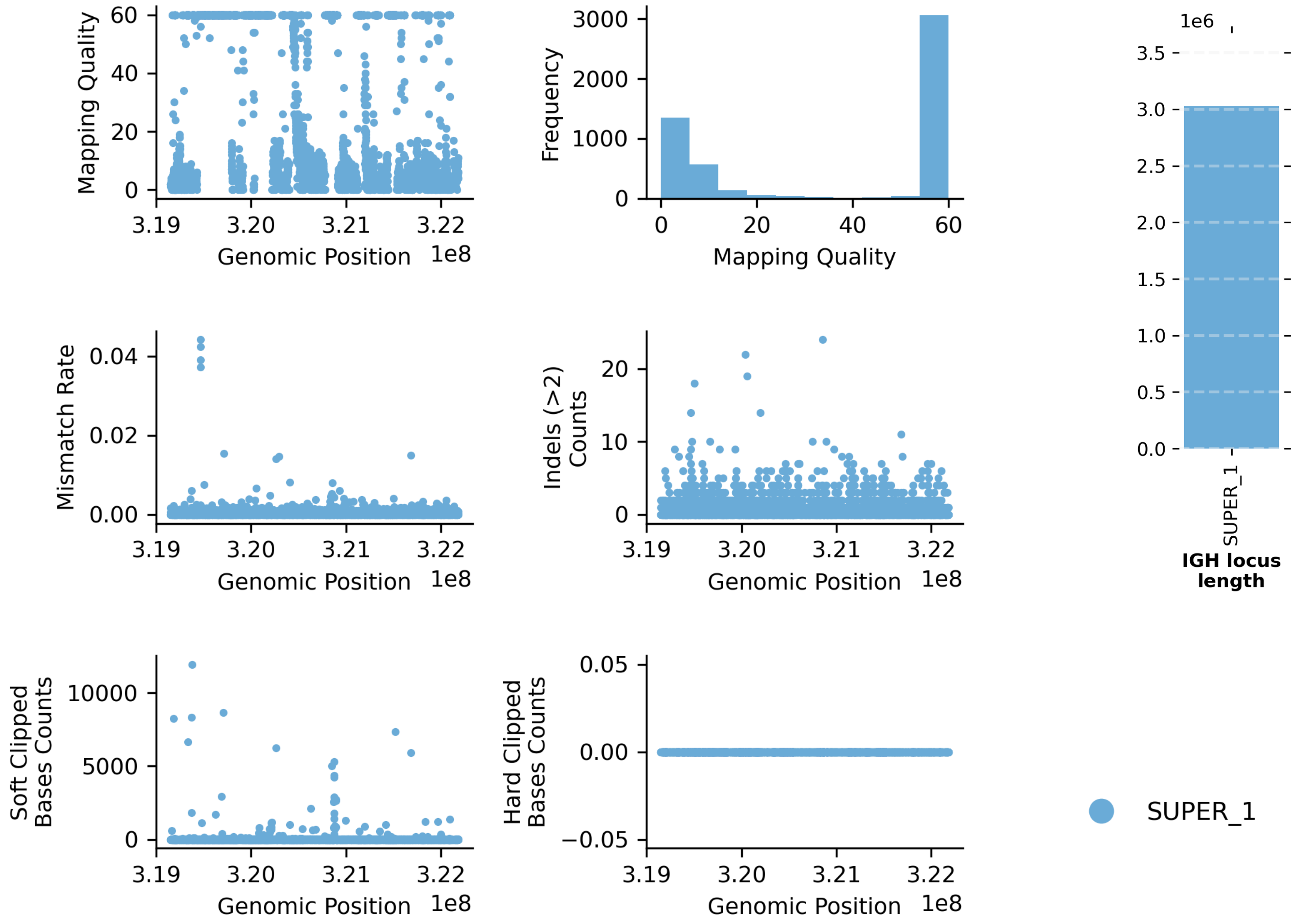


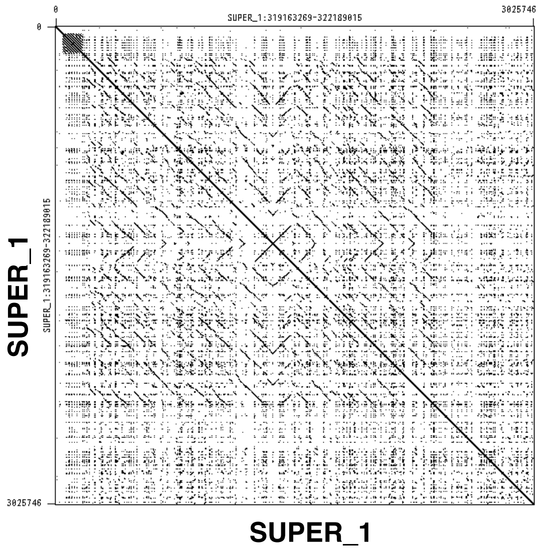


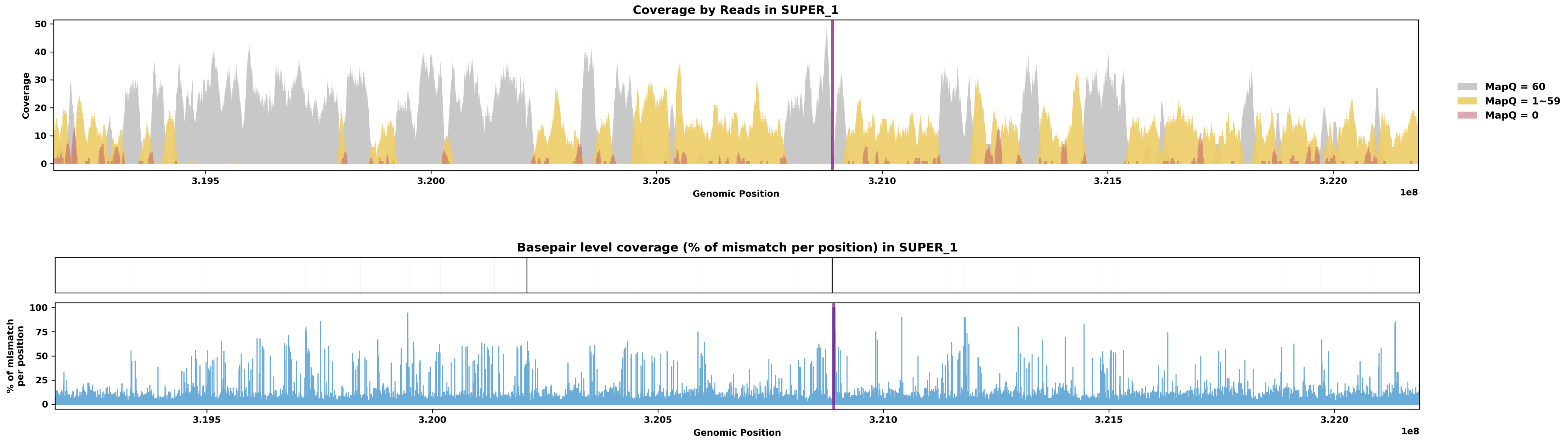

Species ID: mManPen7

Common Name: Chinese pangolin

Scientific Name: Manis pentadactyla

Assembly Type: Haplotype Resolved

Data Source: VGP

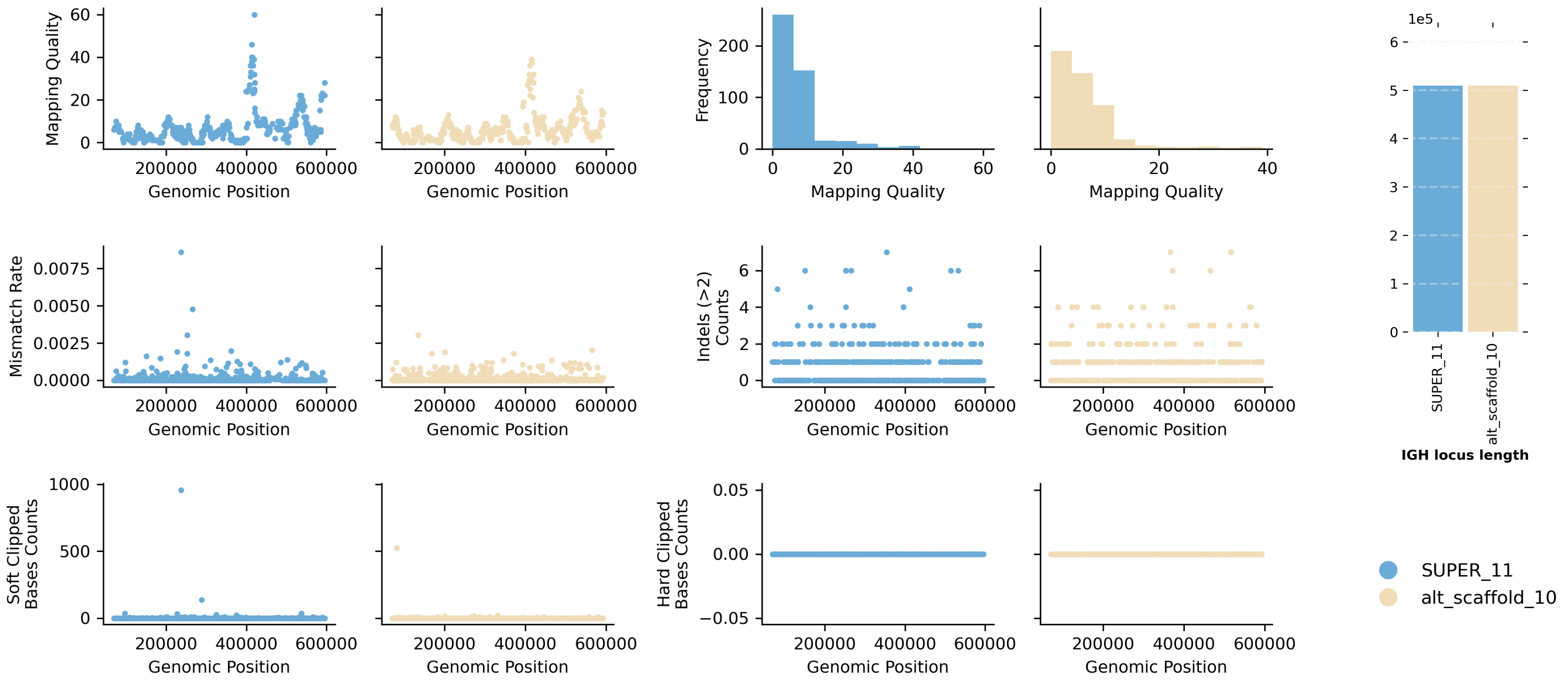


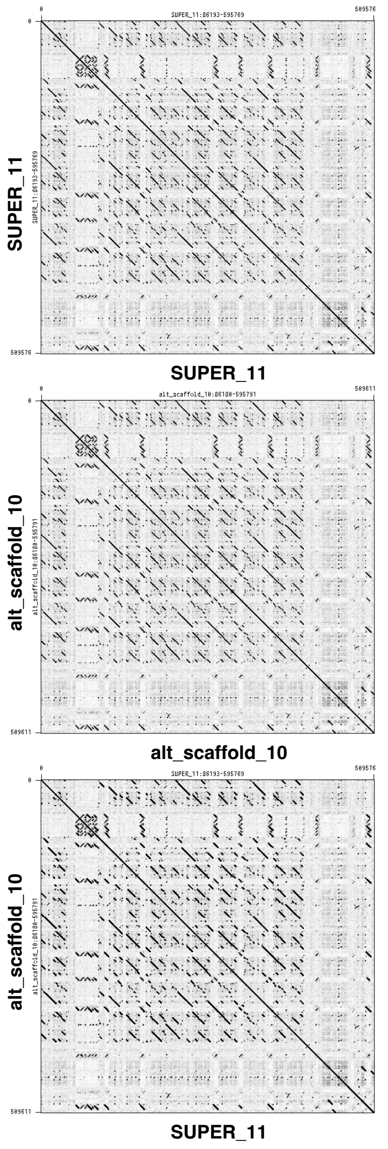


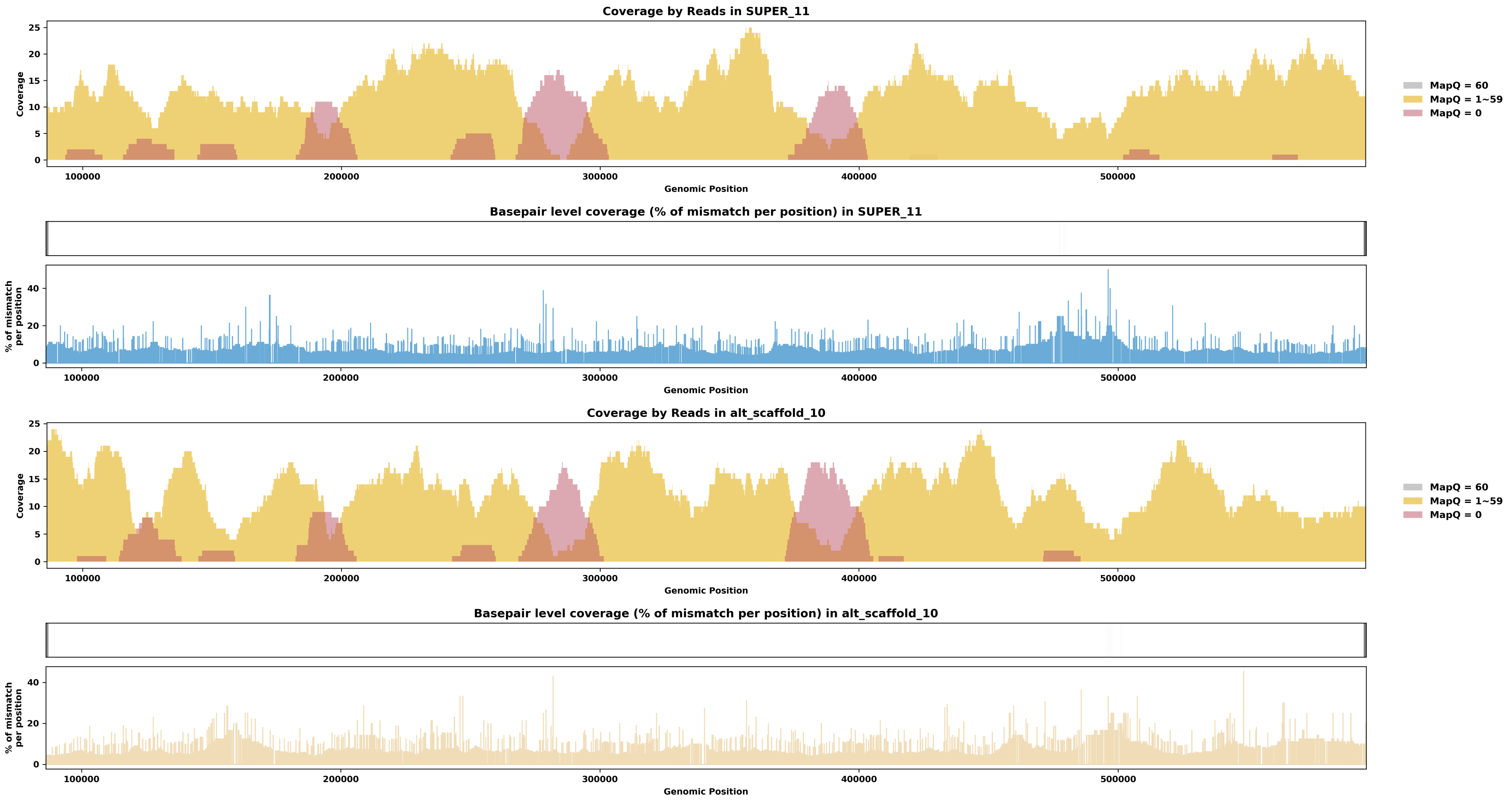

Species ID: mMarMar1

Common Name: European pine marten

Scientific Name: Martes martes

Assembly Type: Not Haplotype Resolved

Data Source: VGP

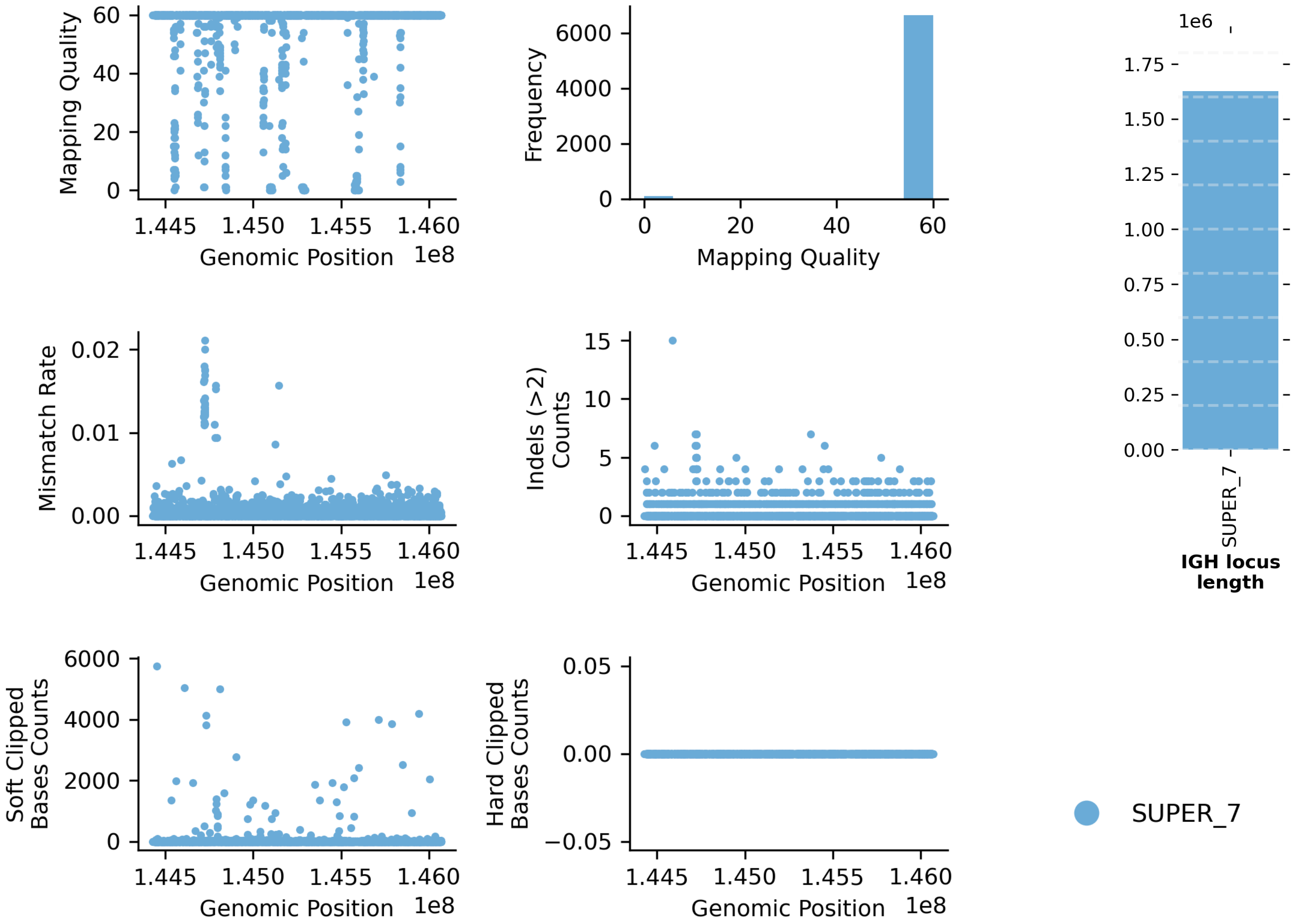


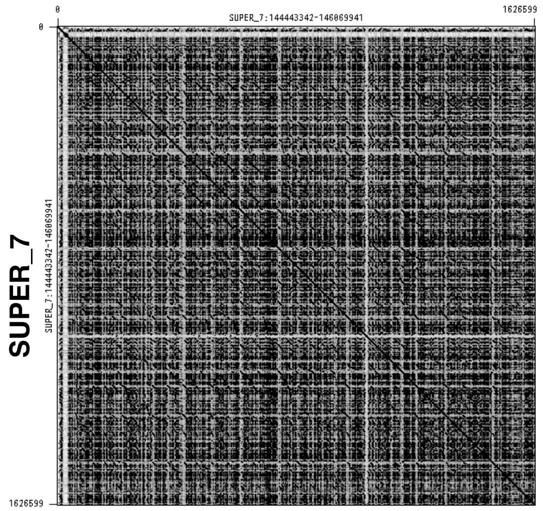


SUPER_7

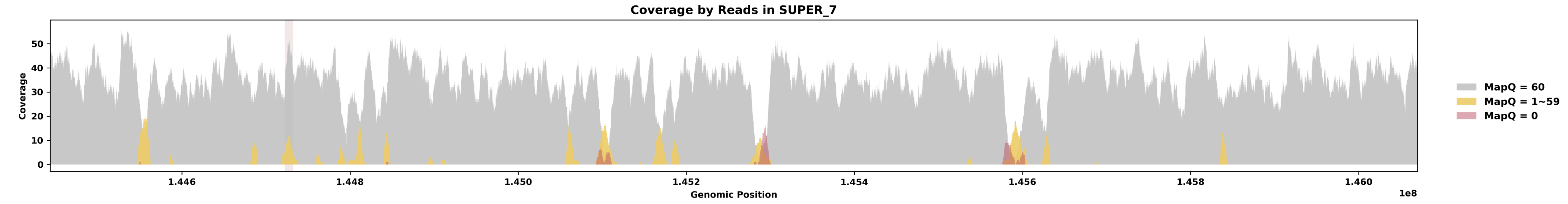


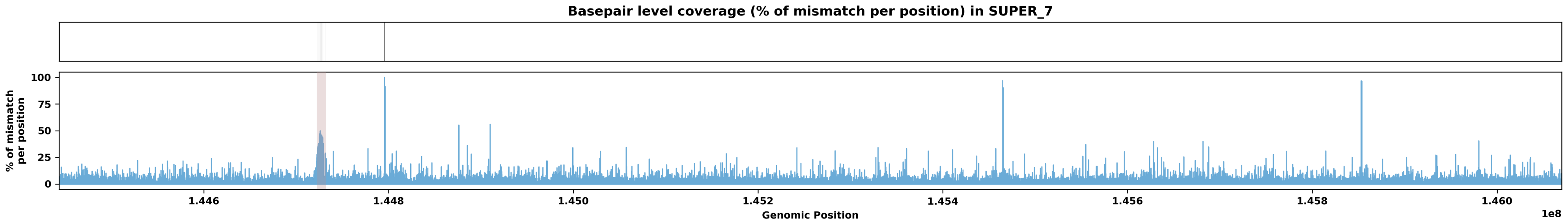

Species ID: mMelMel3

Common Name: European badger

Scientific Name: Meles meles

Assembly Type: Haplotype Resolved

Data Source: VGP

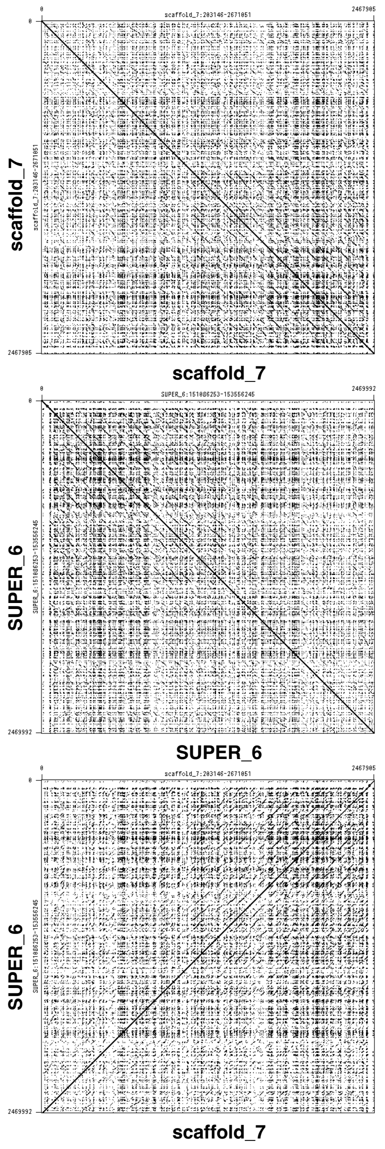


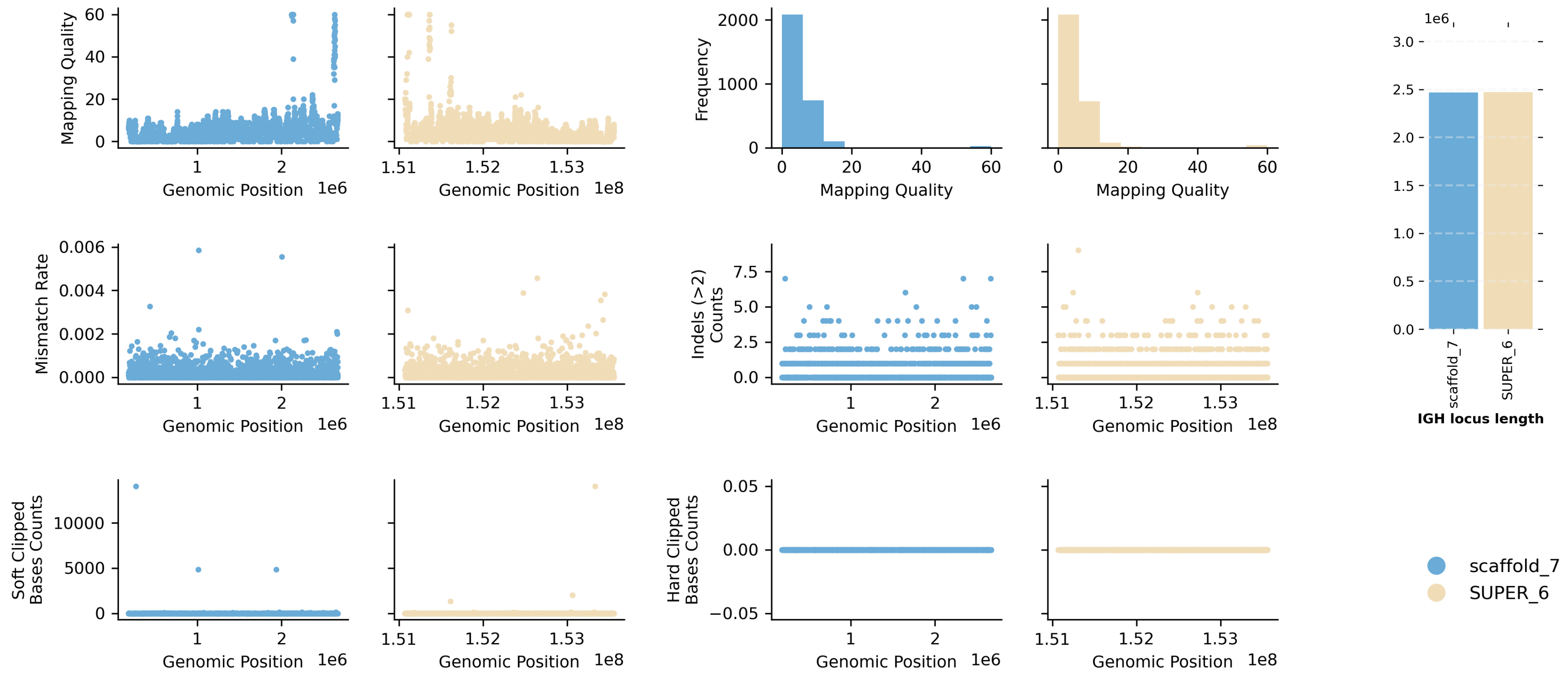


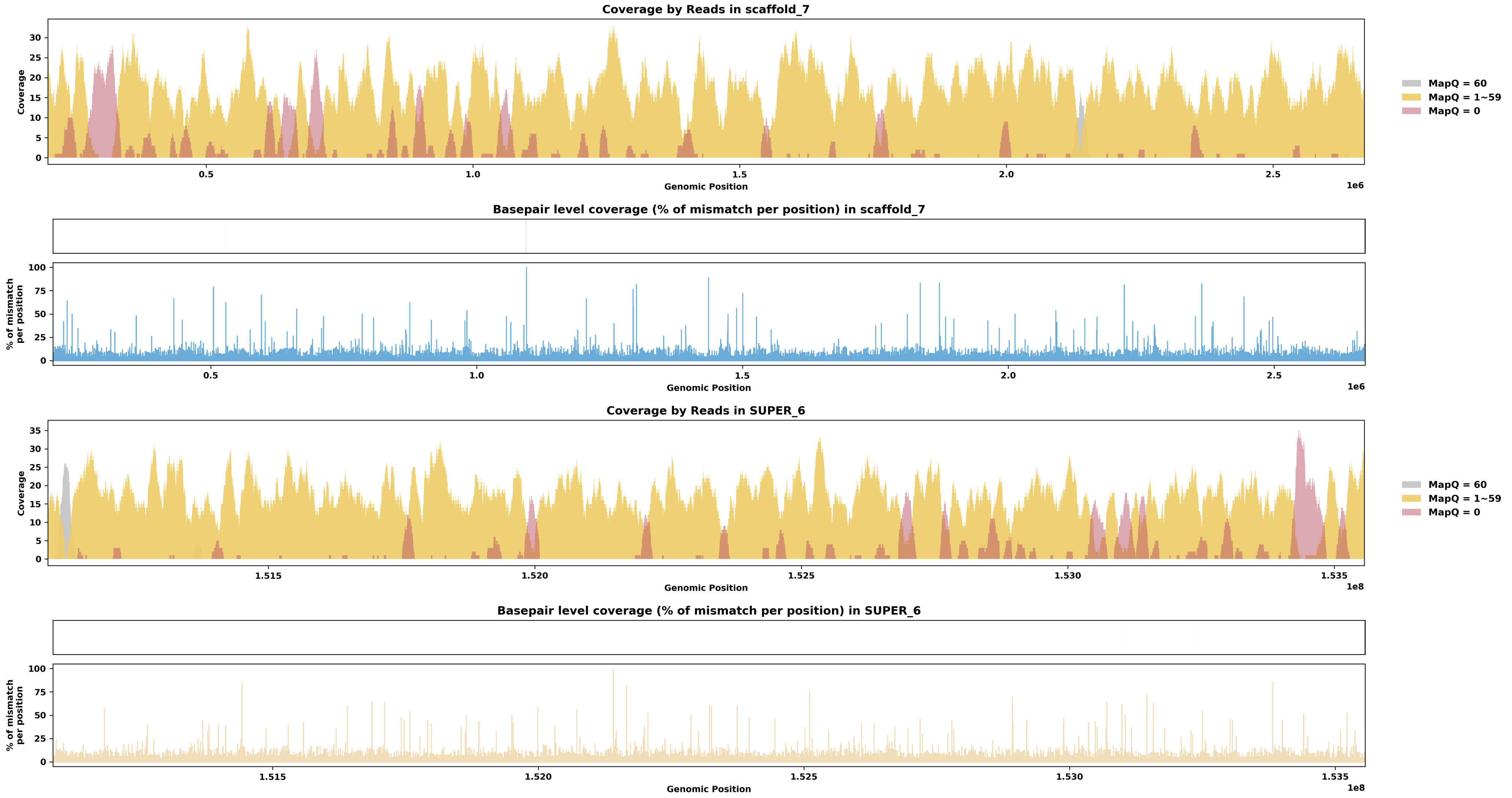

Species ID: mMesDen1

Common Name: Blainville's beaked whale

Scientific Name: Mesoplodon densirostris

Assembly Type: Not Haplotype Resolved

Data Source: VGP

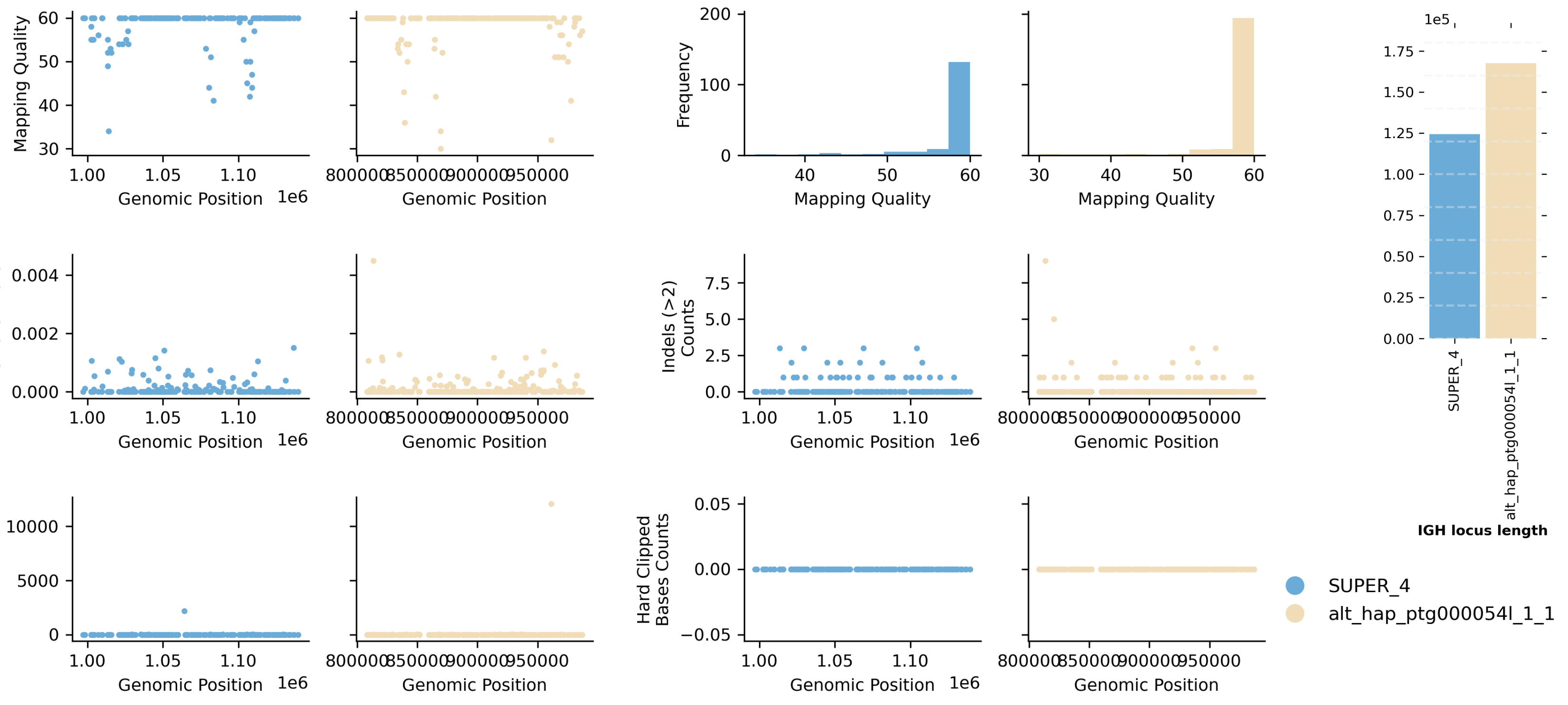


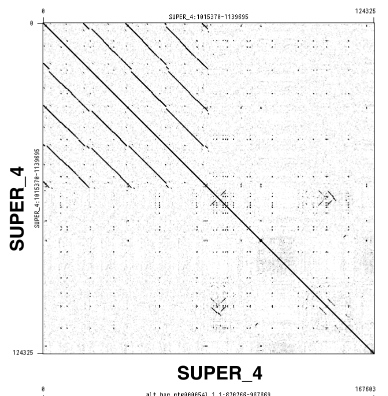


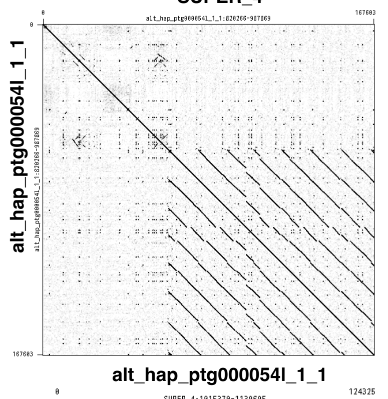


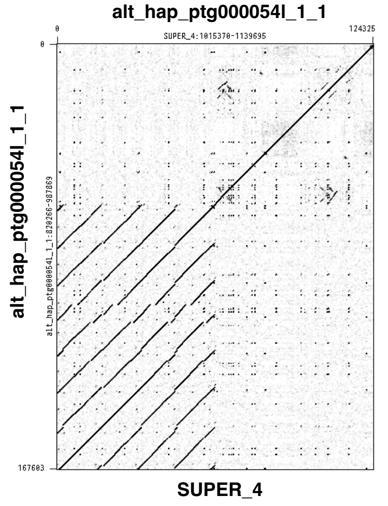


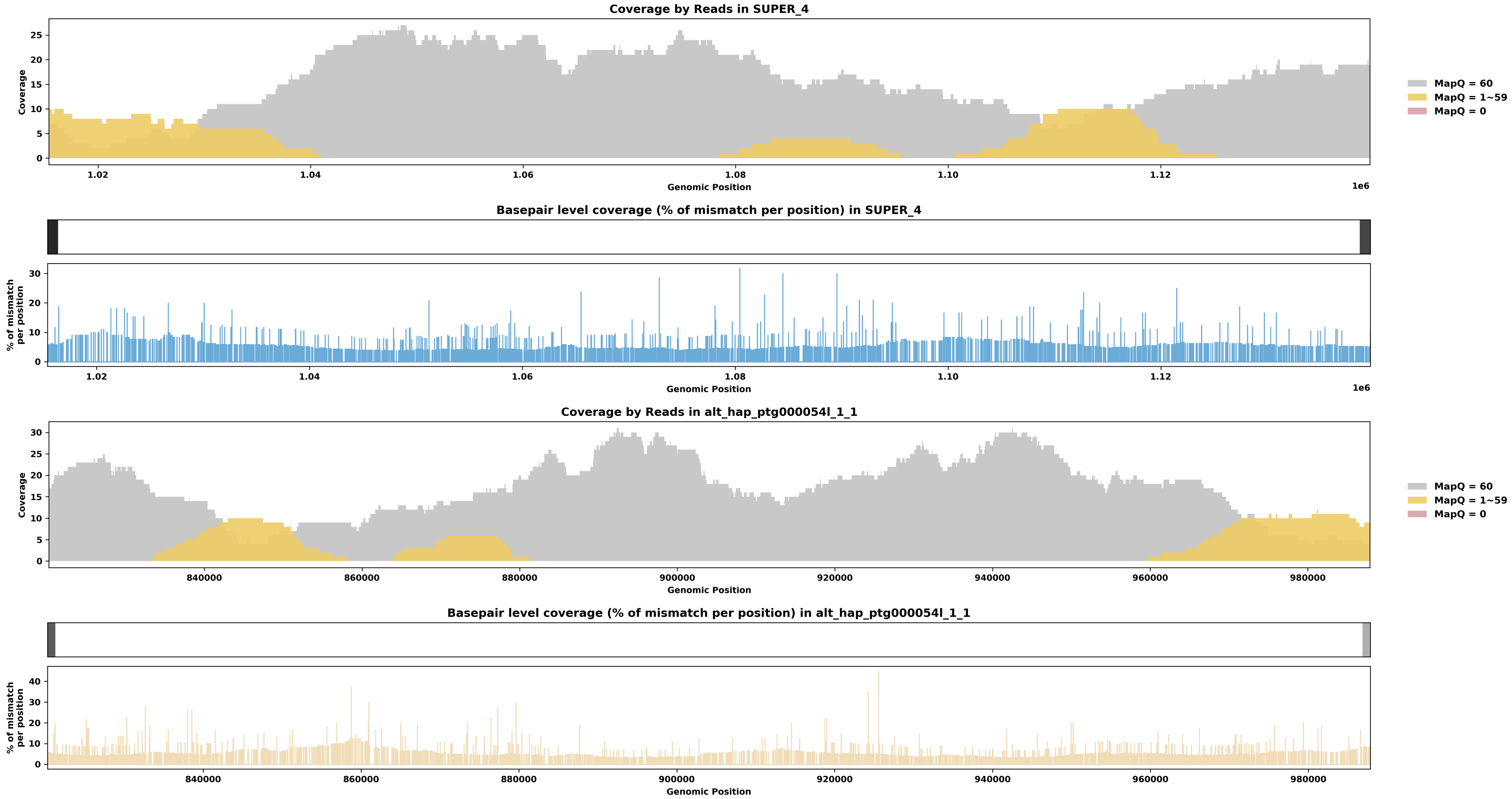

Species ID: mMicCal1

Common Name: California Vole

Scientific Name: Microtus californicus

Assembly Type: Haplotype Resolved

Data Source: CCGP

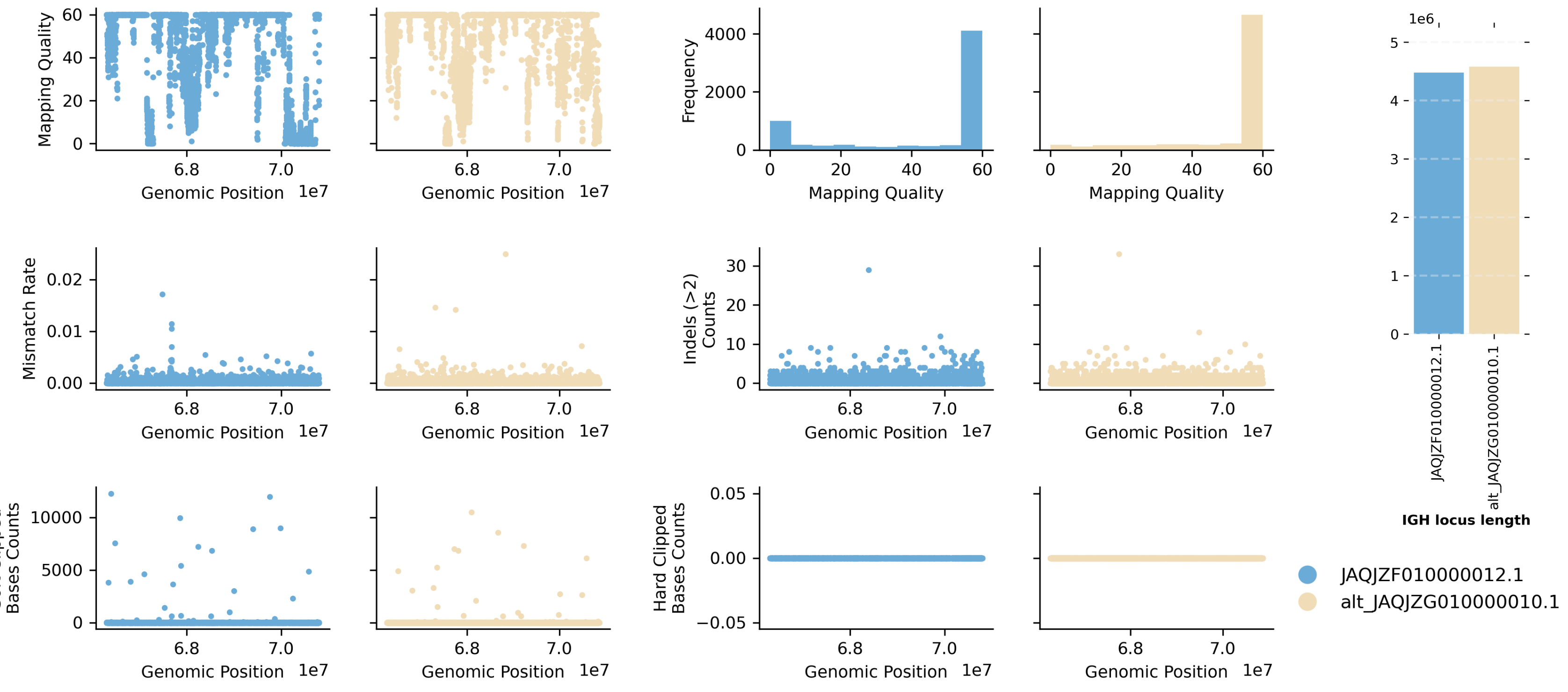


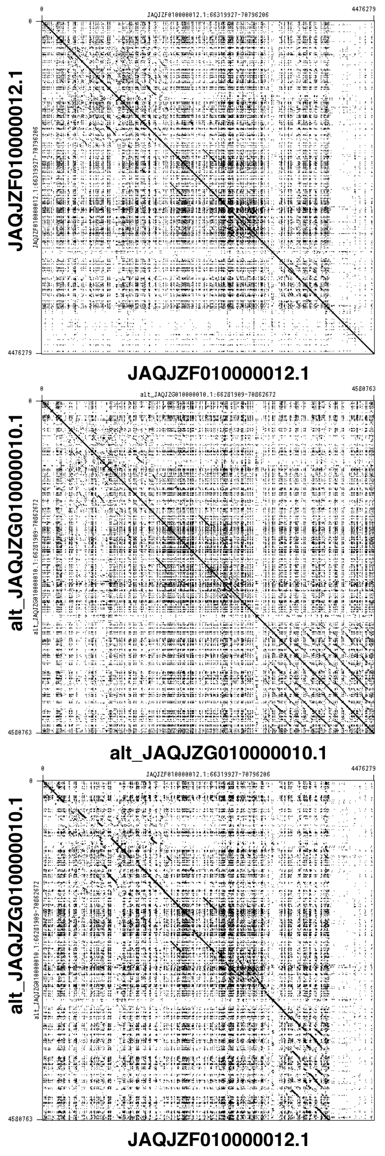


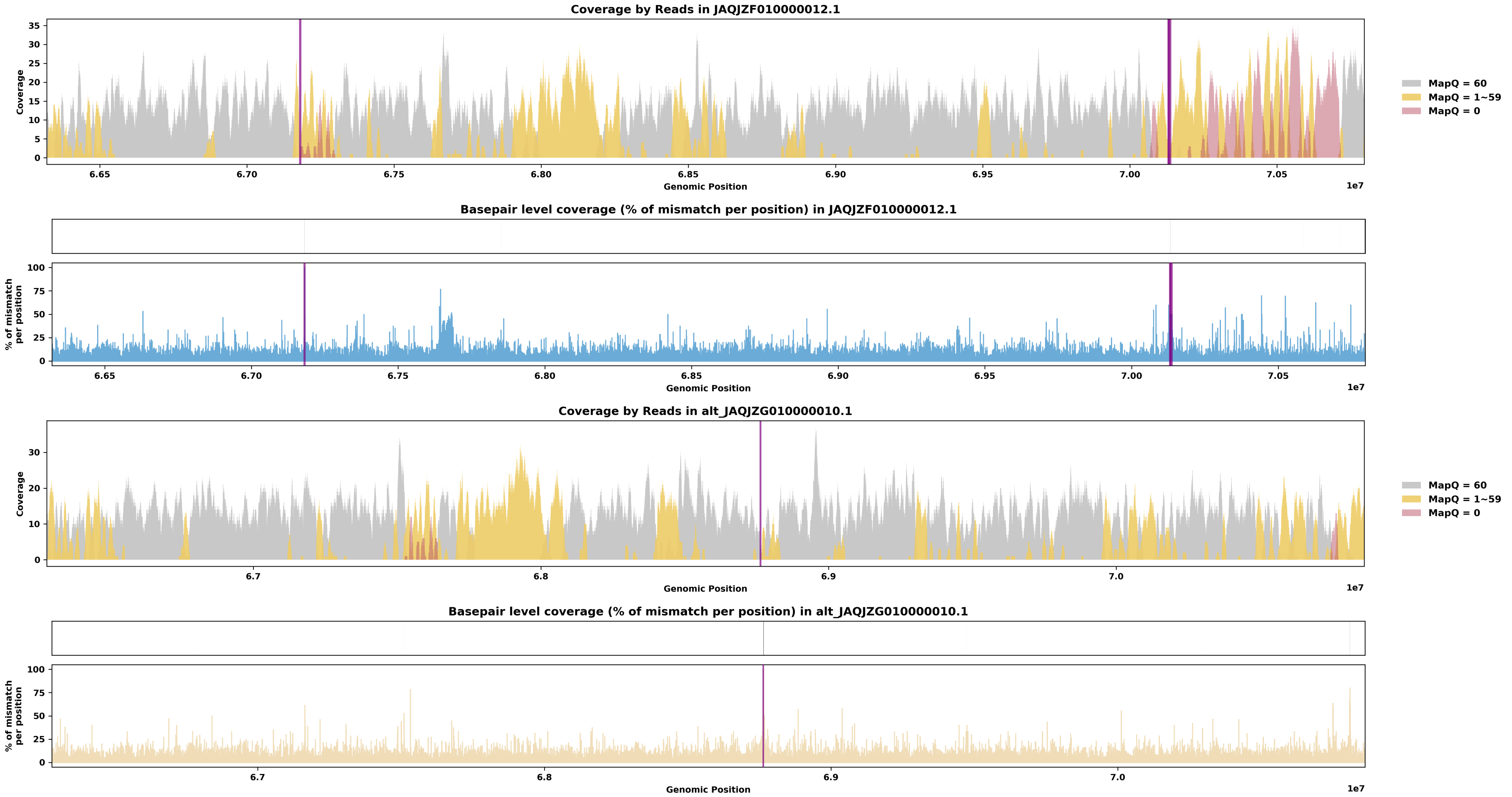

Species ID: mMicMin1

Common Name: European harvest mouse

Scientific Name: Micromys minutus

Assembly Type: Not Haplotype Resolved

Data Source: VGP

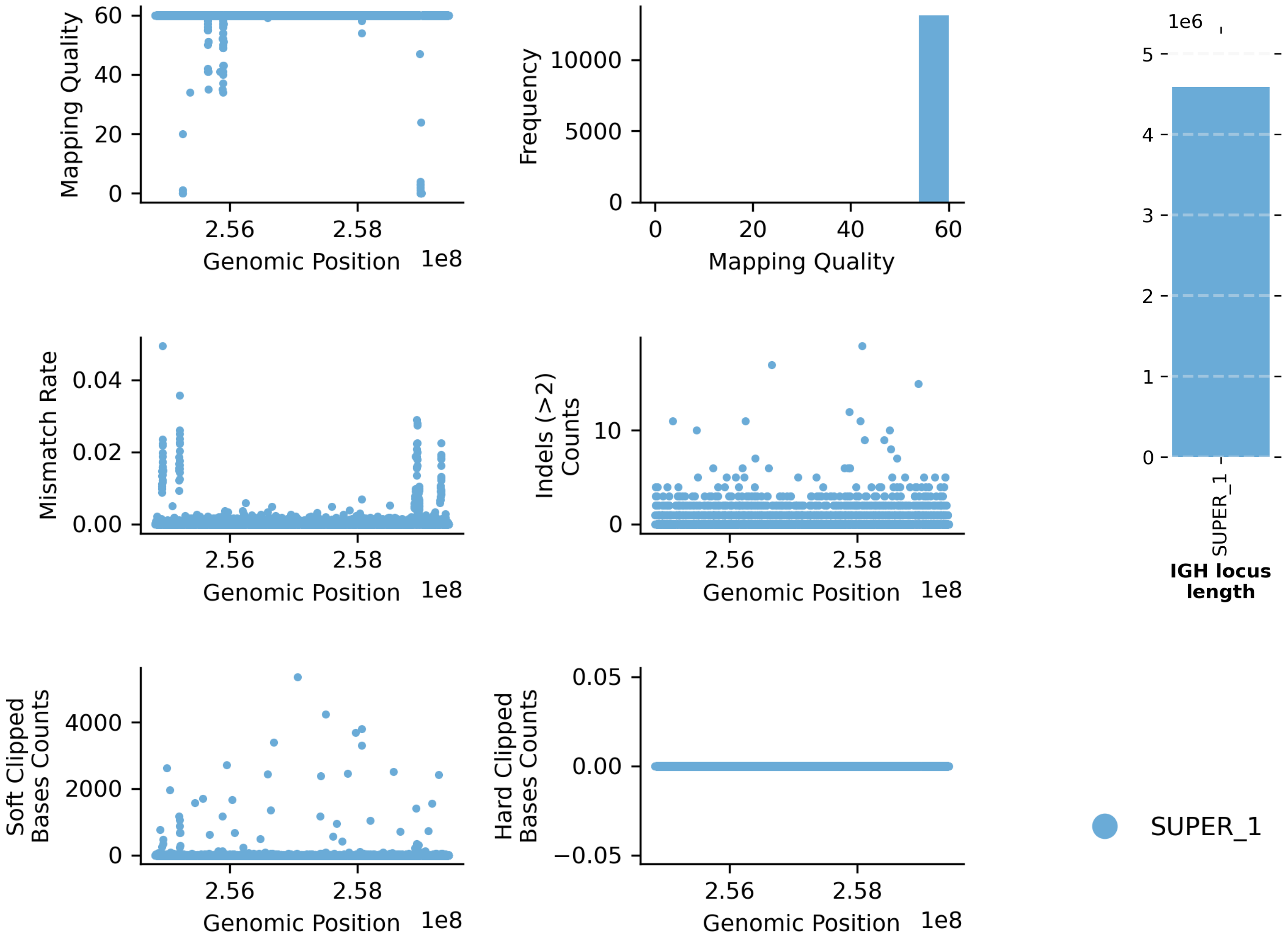


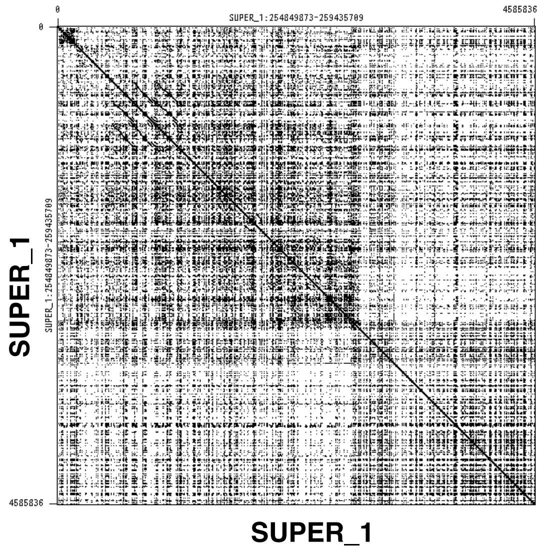


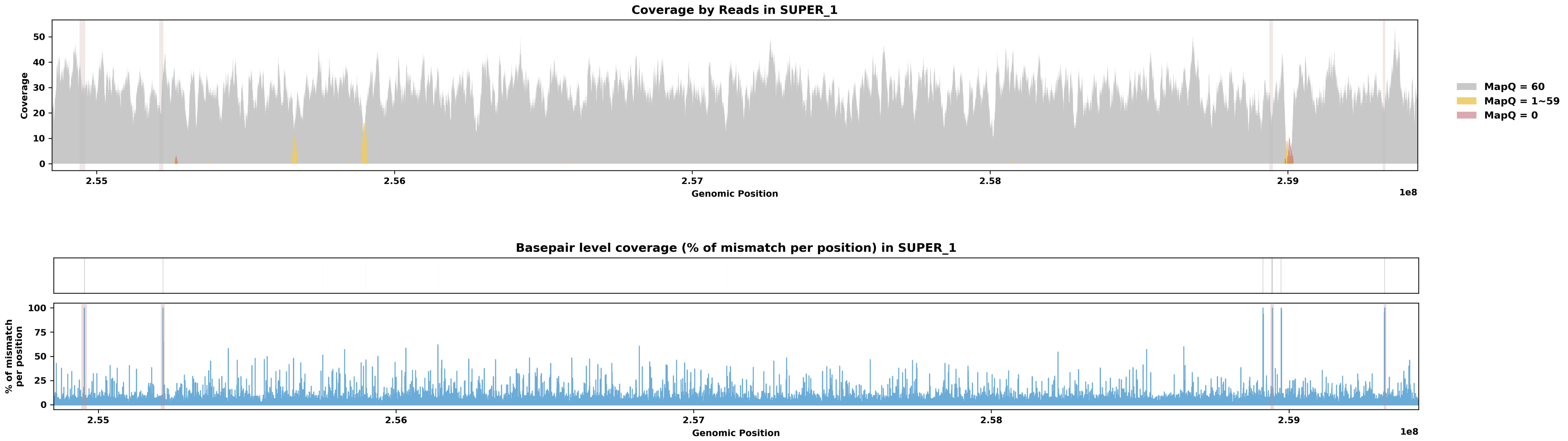

Species ID: mMirAng1

Common Name: Northern Elephant Seal

Scientific Name: Mirounga angustirostris

Assembly Type: Haplotype Resolved

Data Source: CCGP

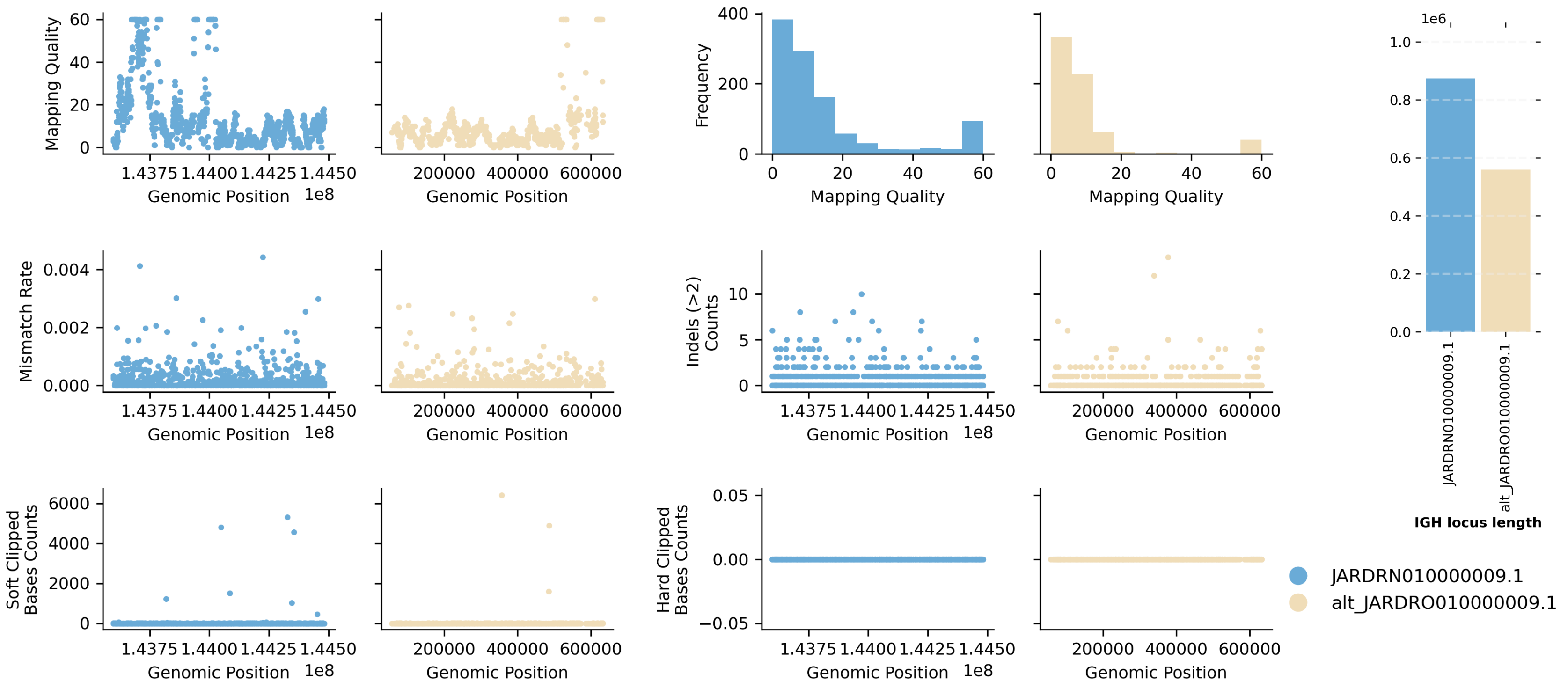


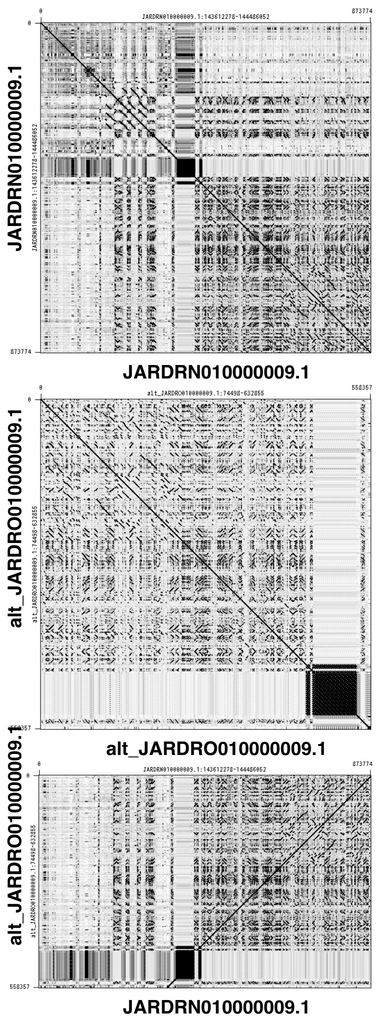


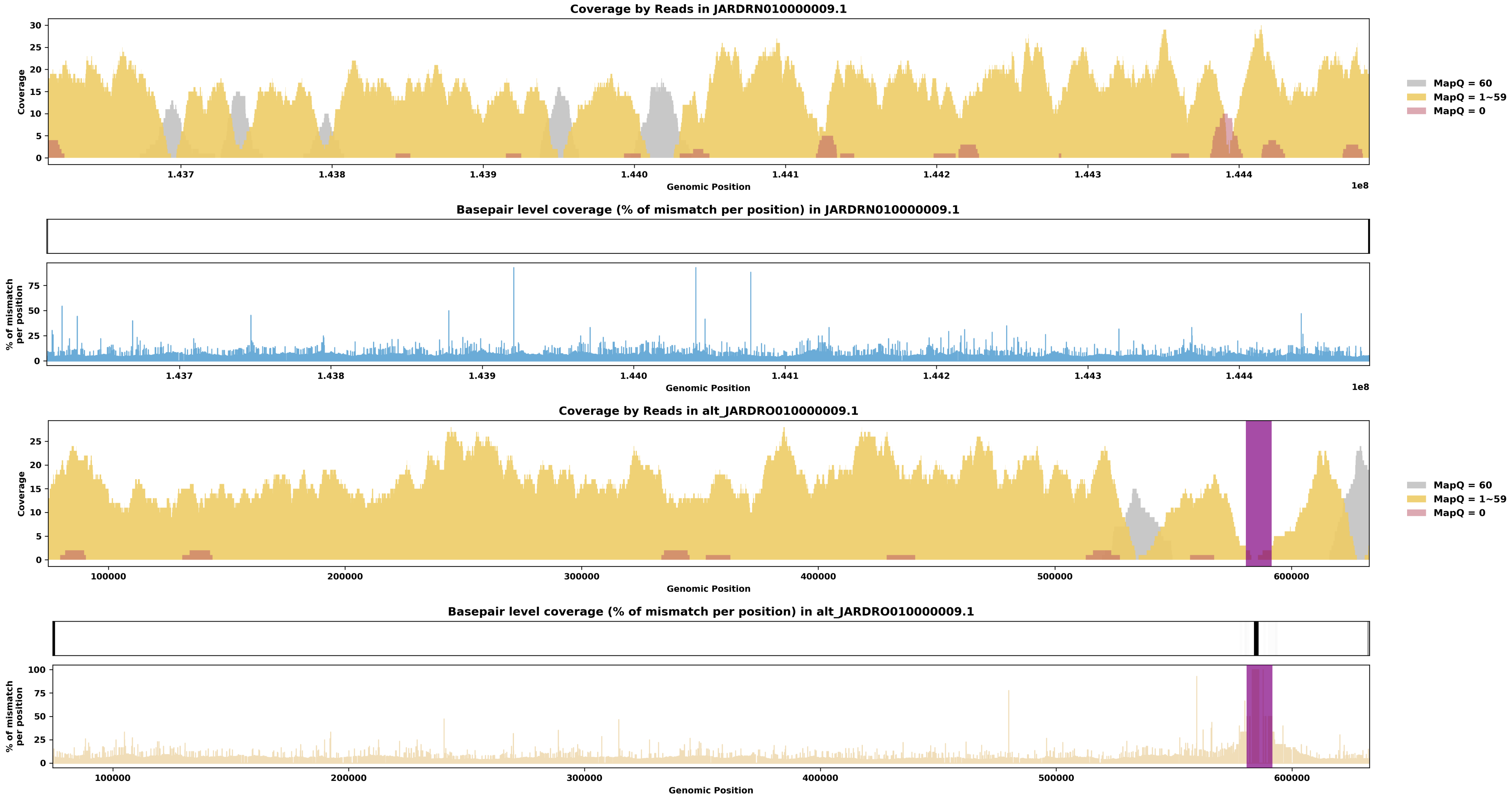

Species ID: mMonDom1

Common Name: gray short-tailed opossum

Scientific Name: Monodelphis_domestica

Assembly Type: Not Haplotype Resolved

Data Source: VGP

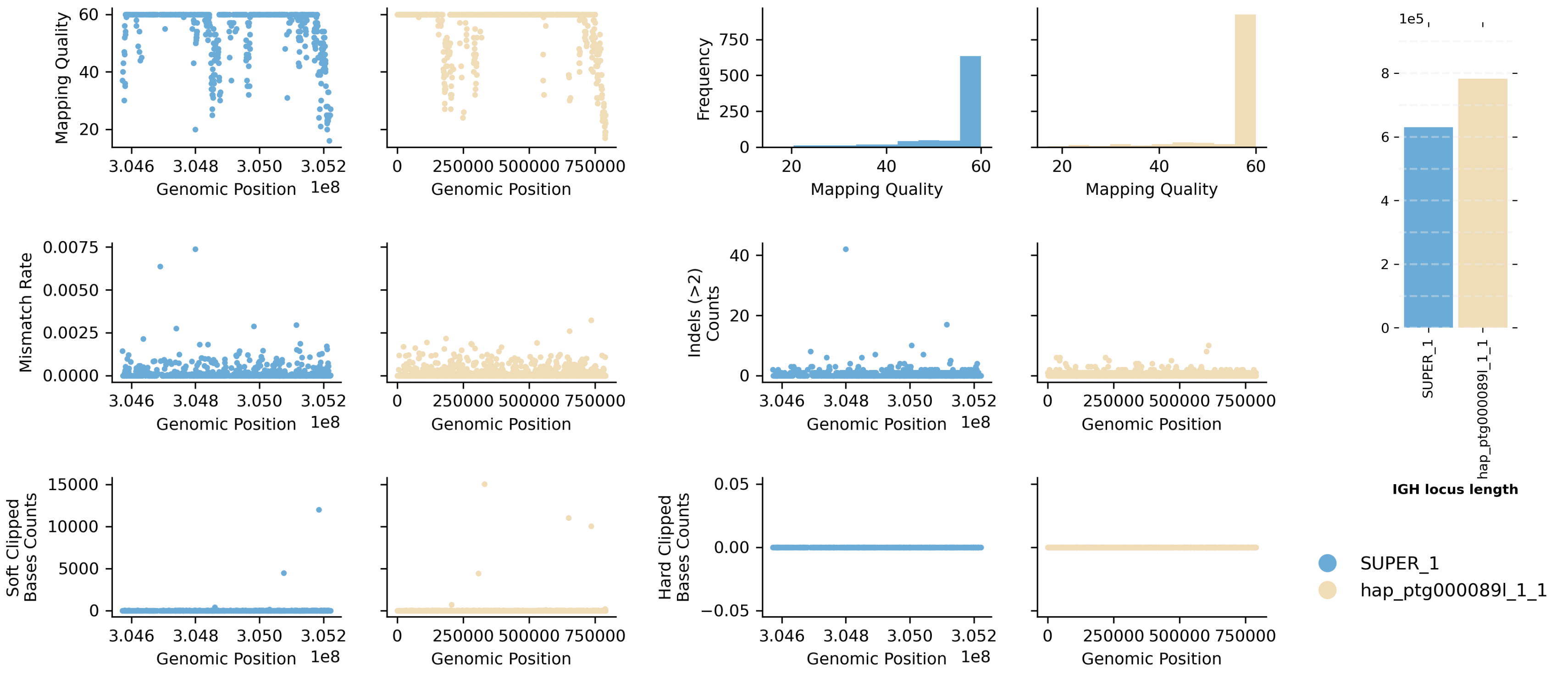


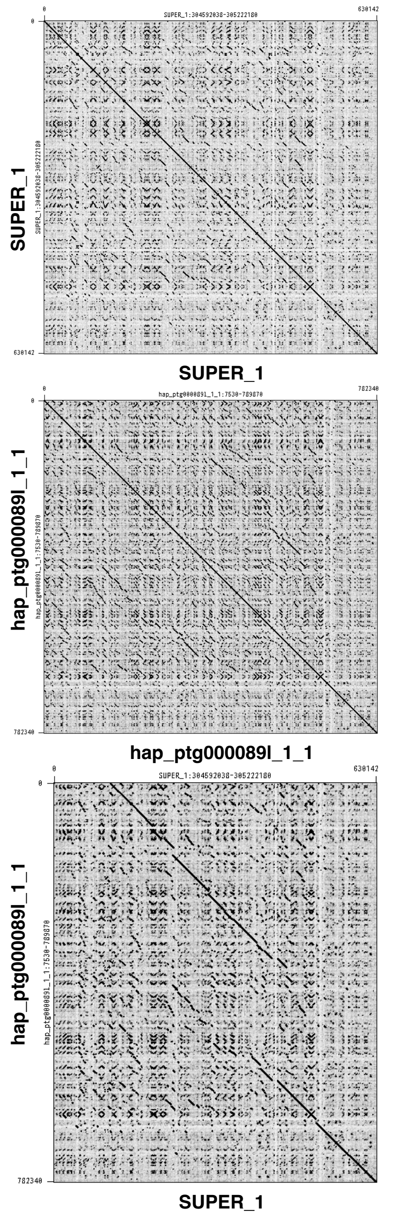


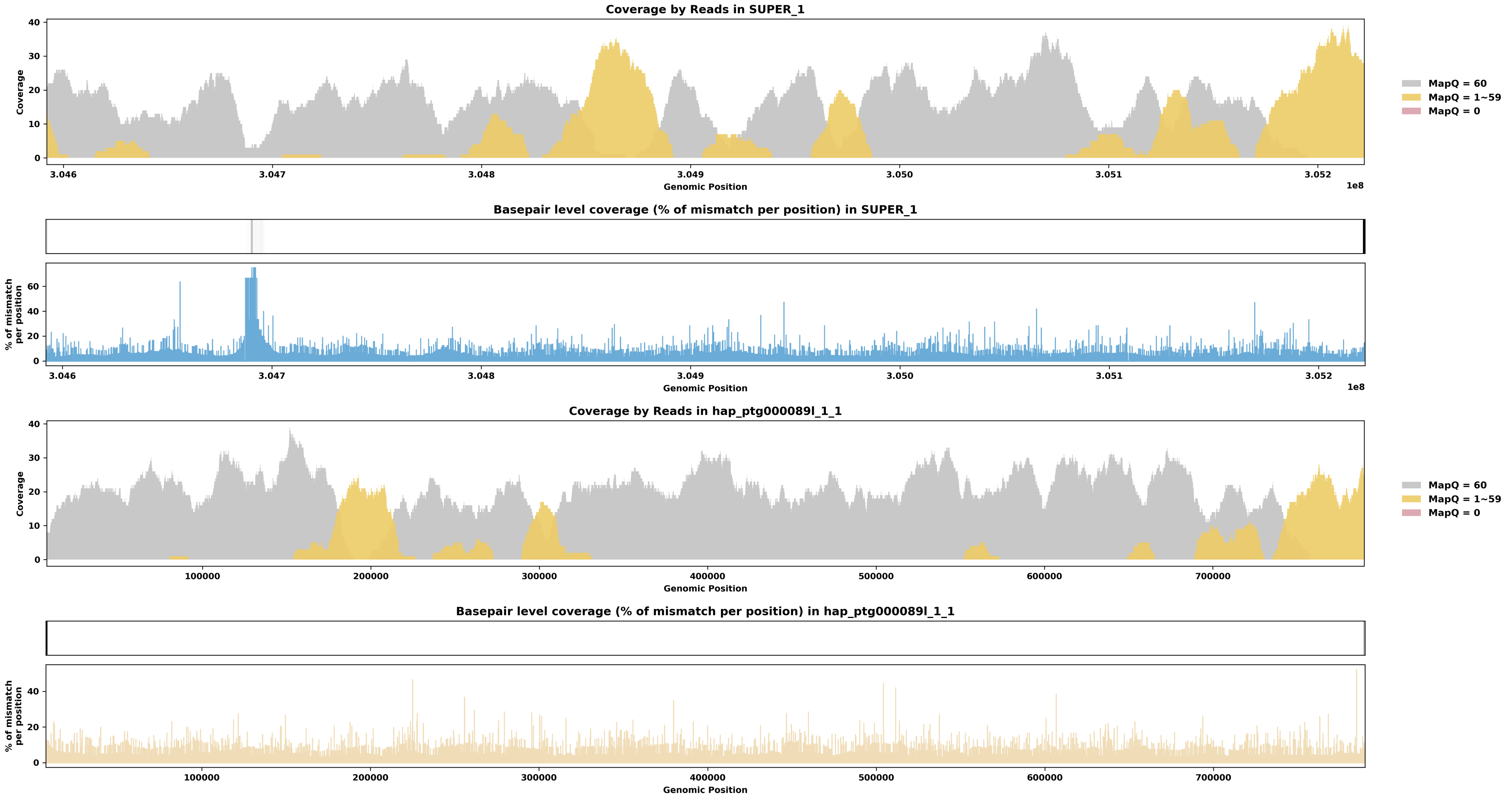

Species ID: mMunRee1

Common Name: Reeves' muntjac

Scientific Name: Muntiacus reevesi

Assembly Type: Not Haplotype Resolved

Data Source: VGP

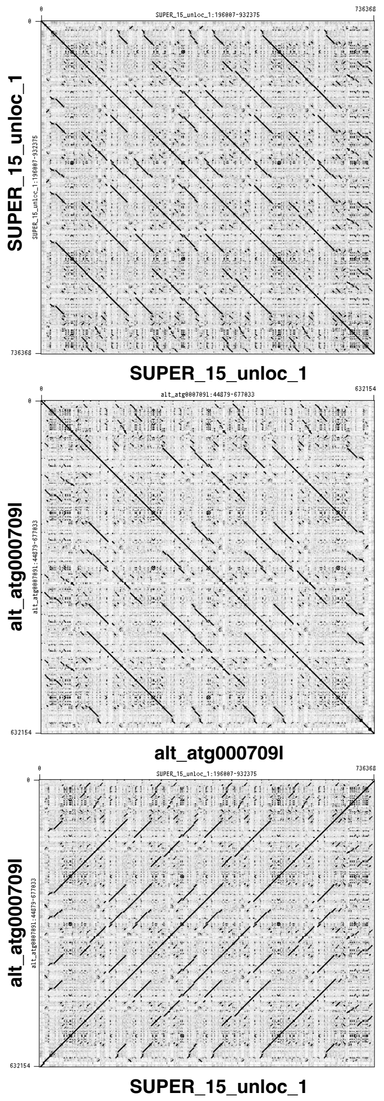


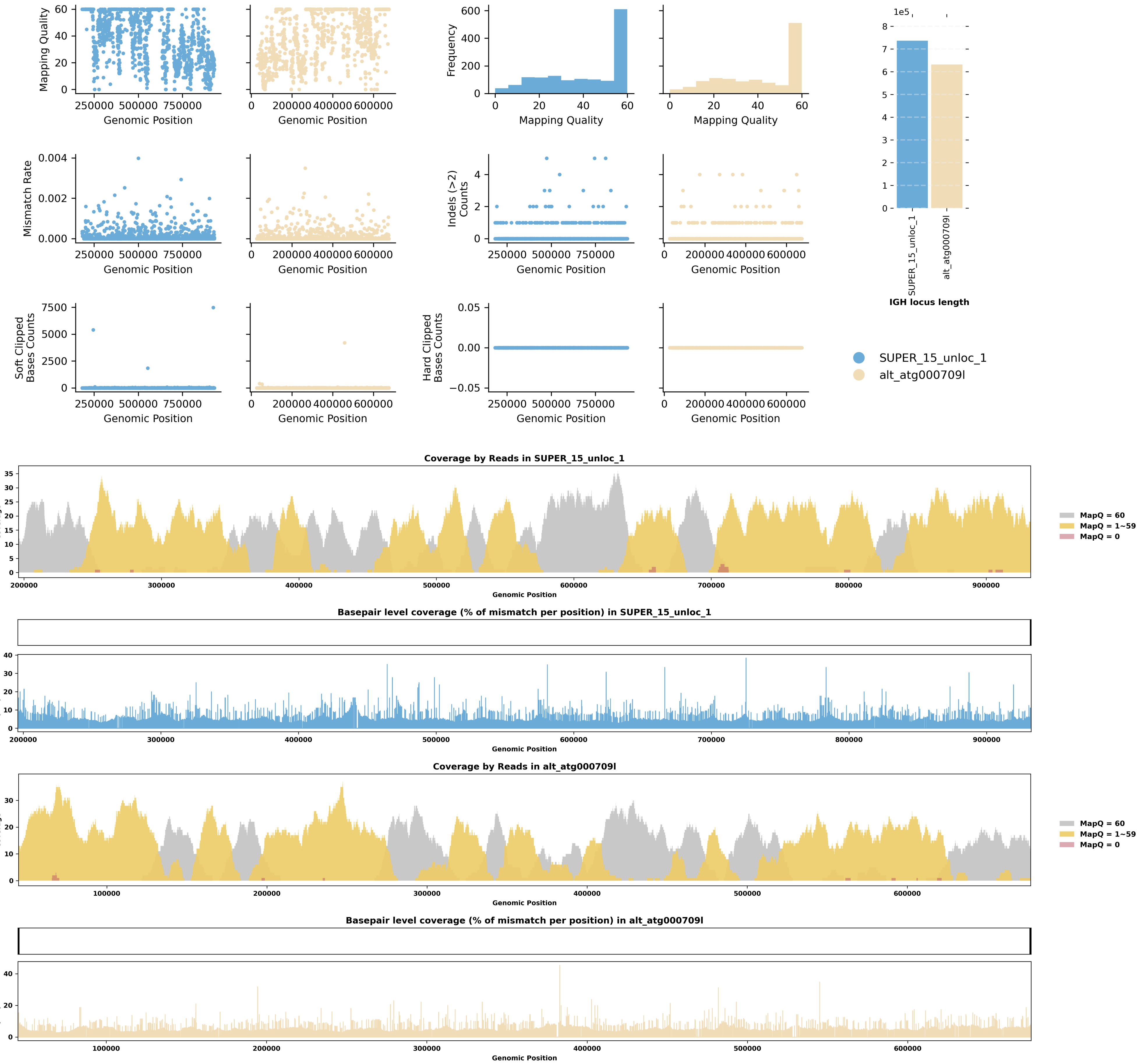

Species ID: mMusAve1

Common Name: hazel dormouse

Scientific Name: Muscardinus avellanarius

Assembly Type: Not Haplotype Resolved

Data Source: VGP

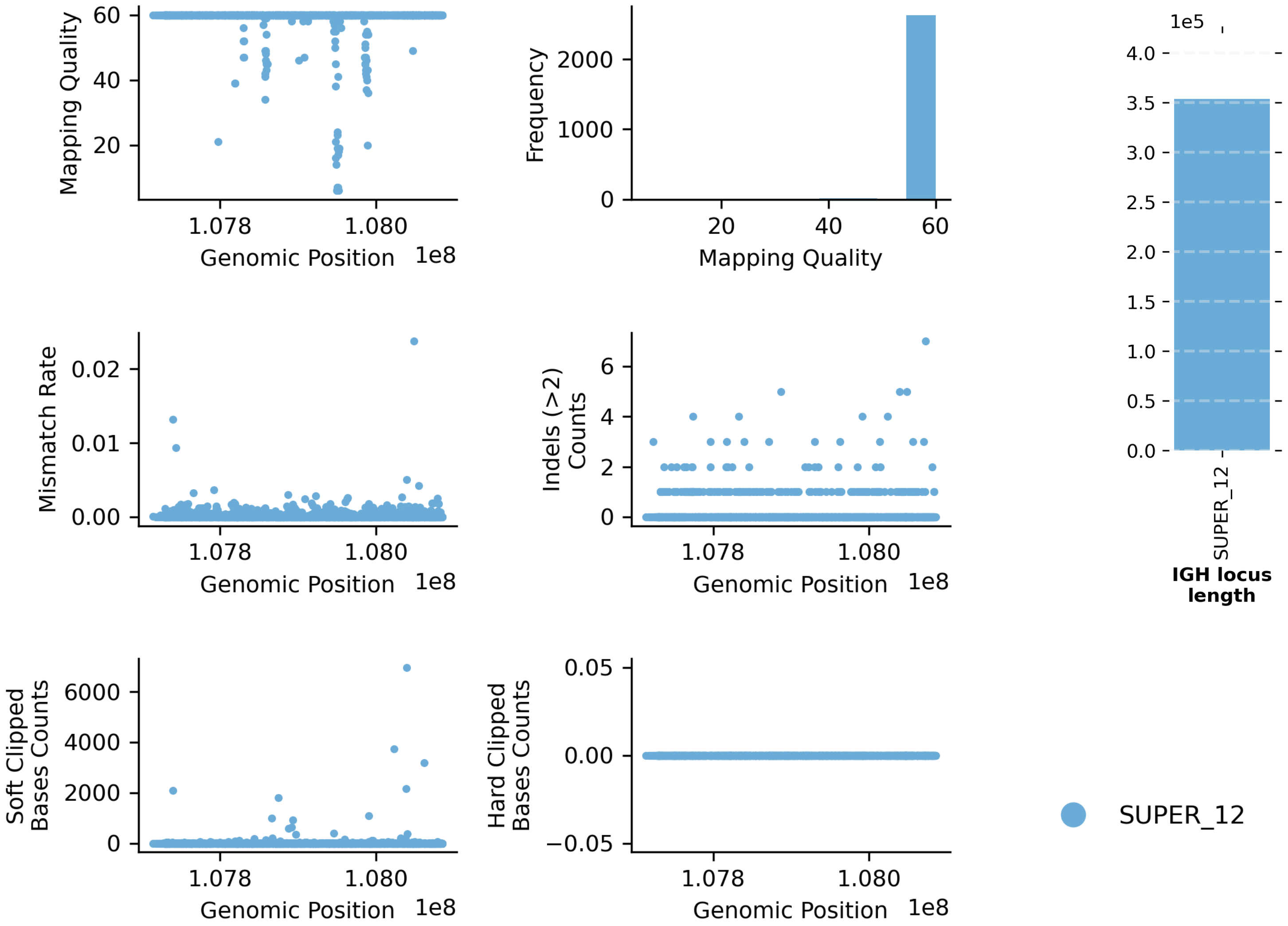


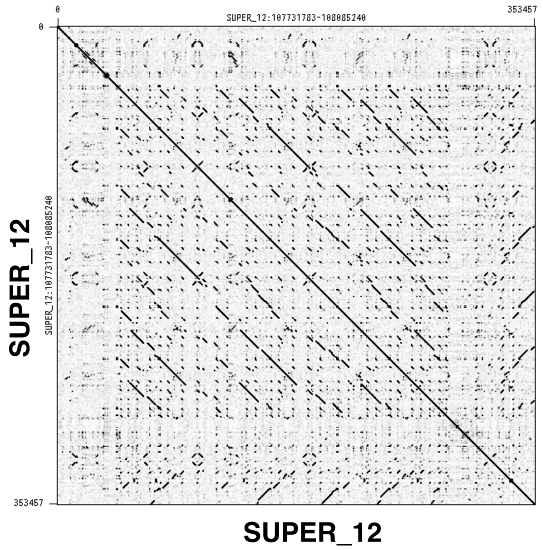


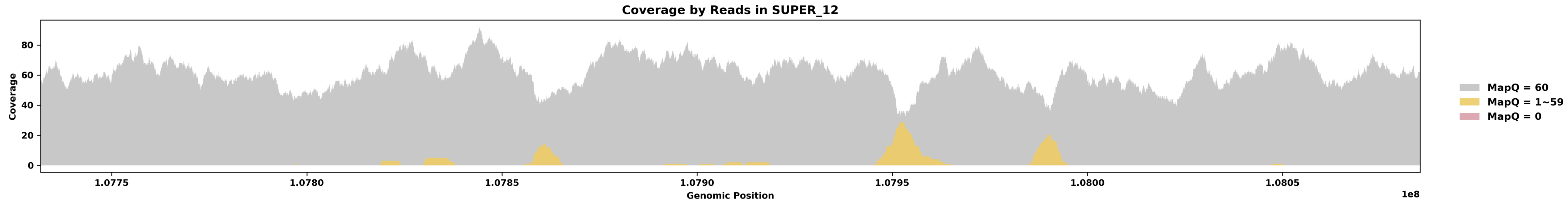


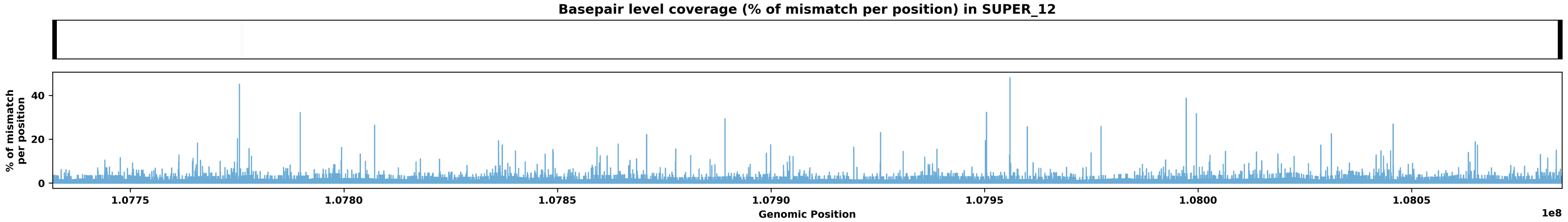

Species ID: mMusLut2

Common Name: European mink

Scientific Name: Mustela lutreola

Assembly Type: Not Haplotype Resolved

Data Source: VGP

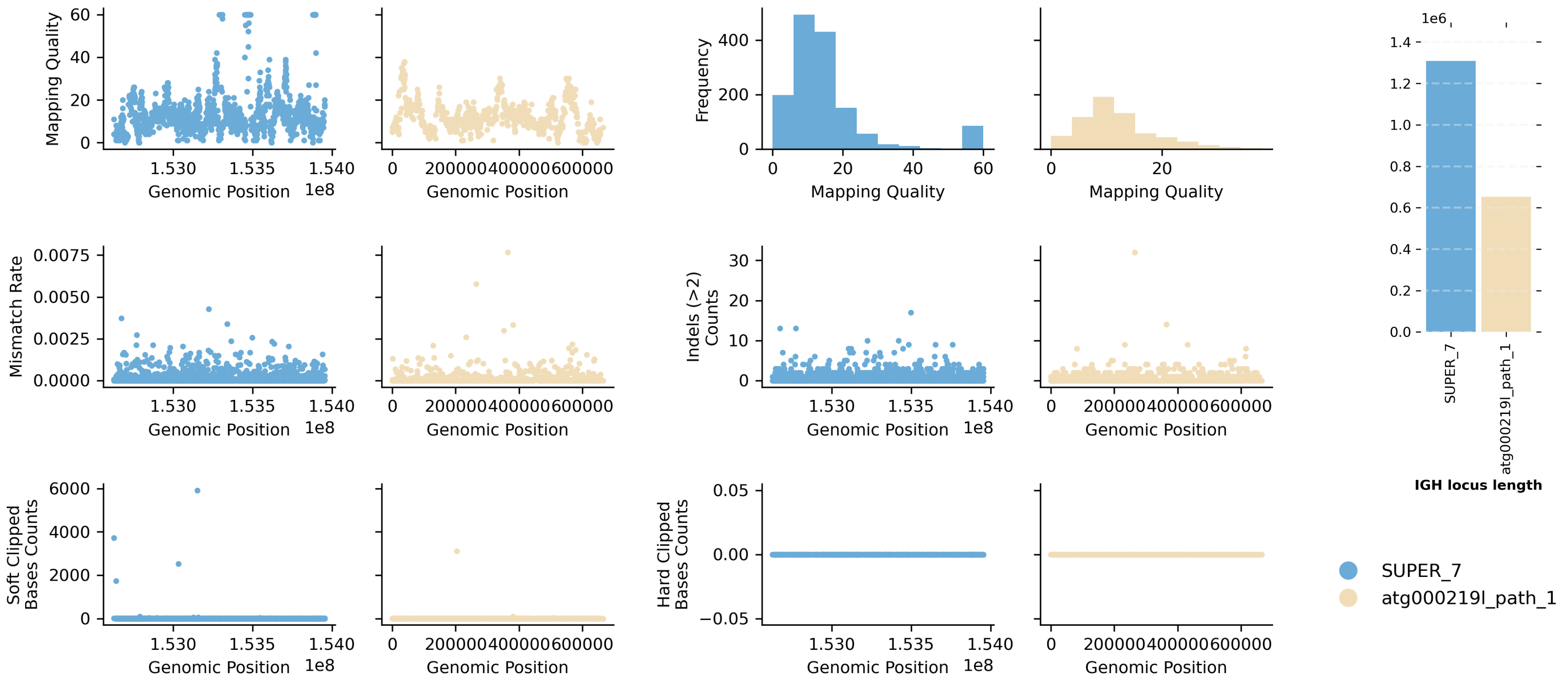


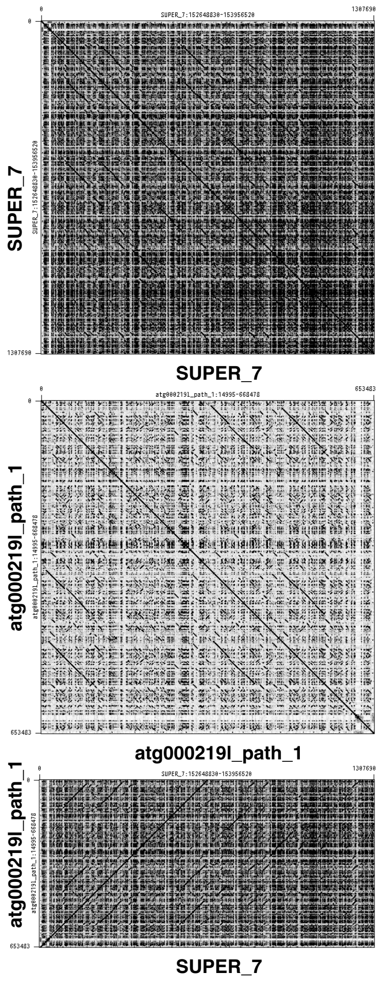


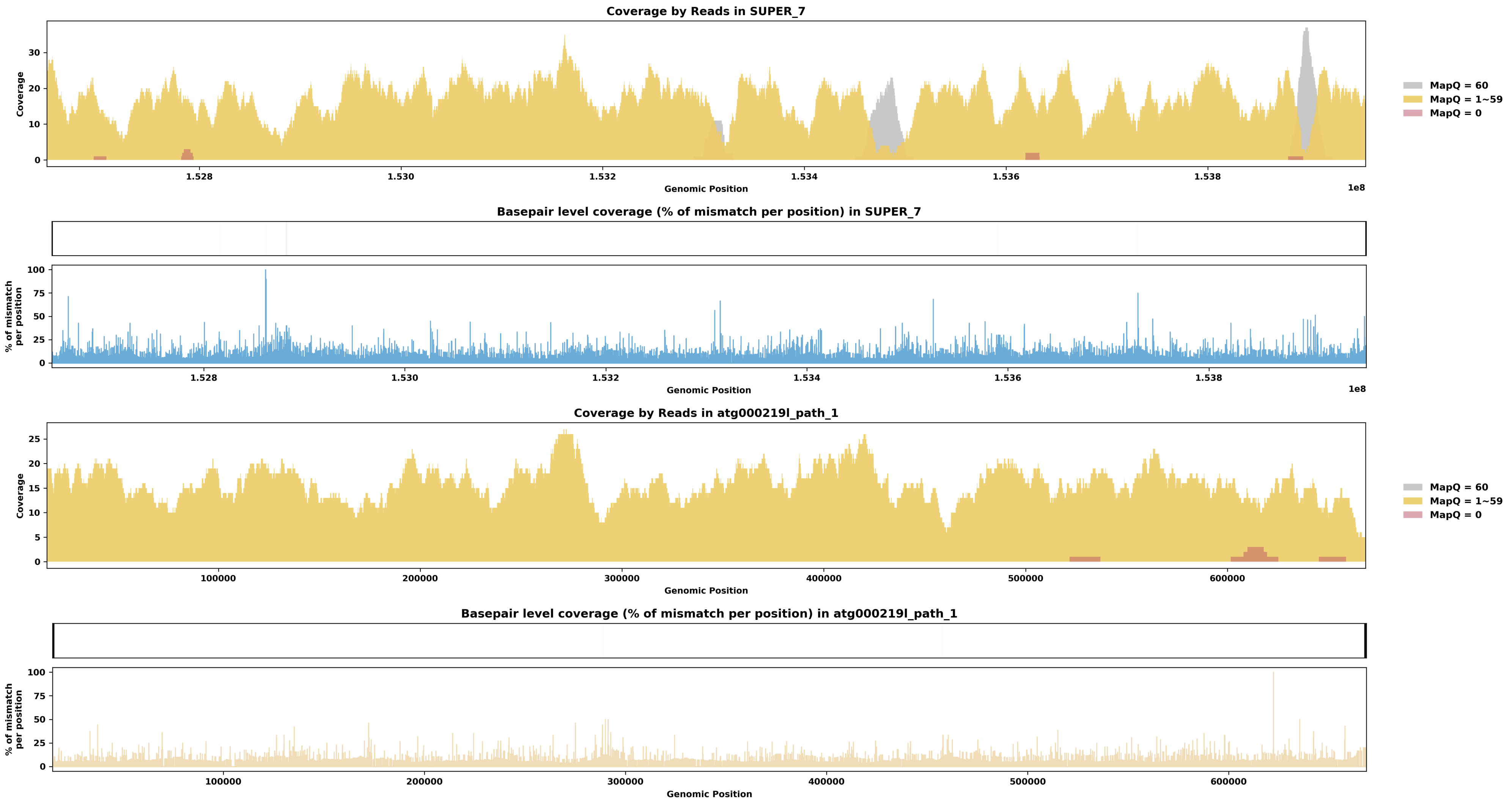

Species ID: mMusNiv1

Common Name: Least weasel

Scientific Name: Mustela nivalis

Assembly Type: Haplotype Resolved

Data Source: VGP

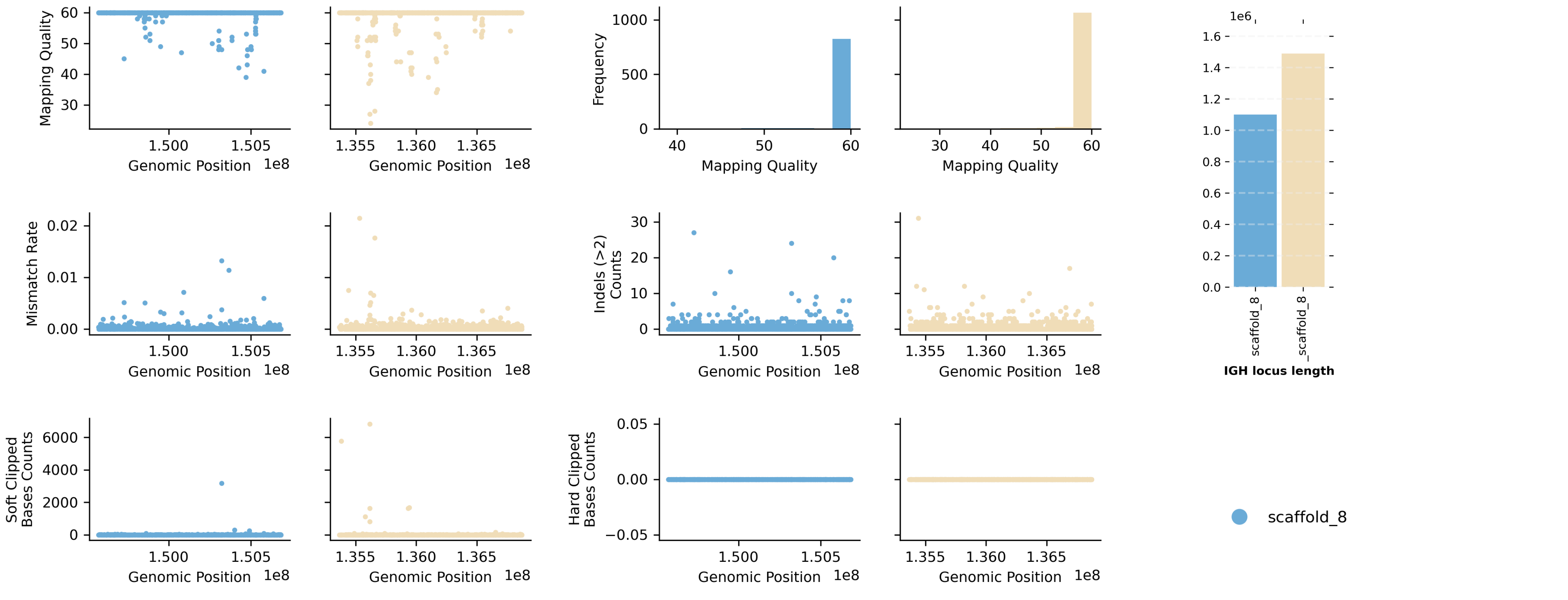


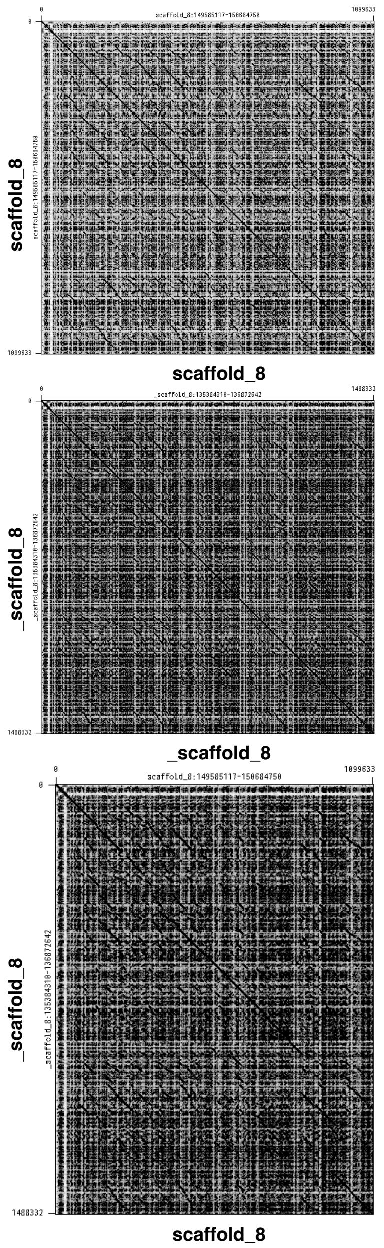


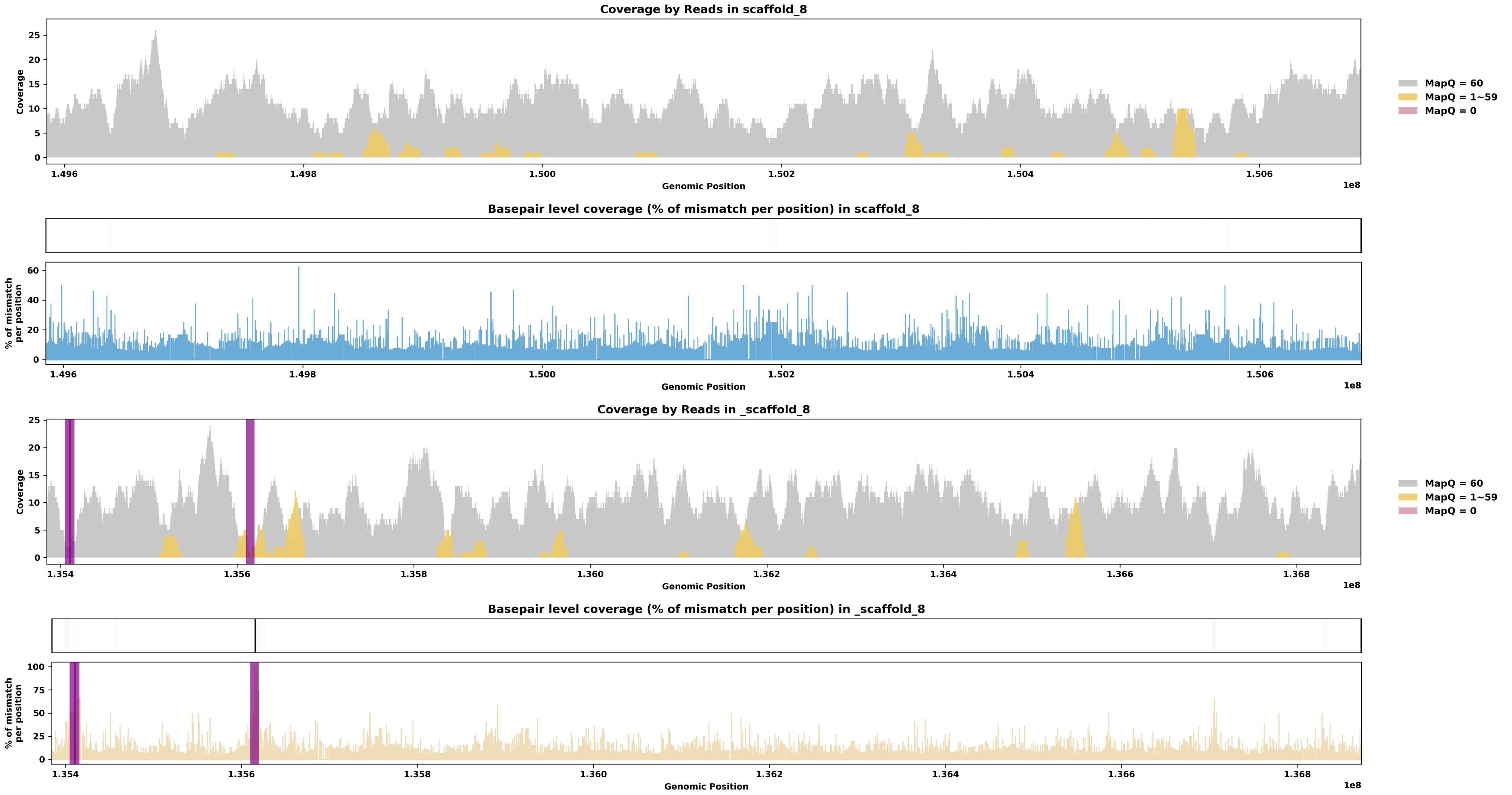

Species ID: mMyoDau2

Common Name: Daubenton's bat

Scientific Name: Myotis daubentonii

Assembly Type: Not Haplotype Resolved

Data Source: VGP

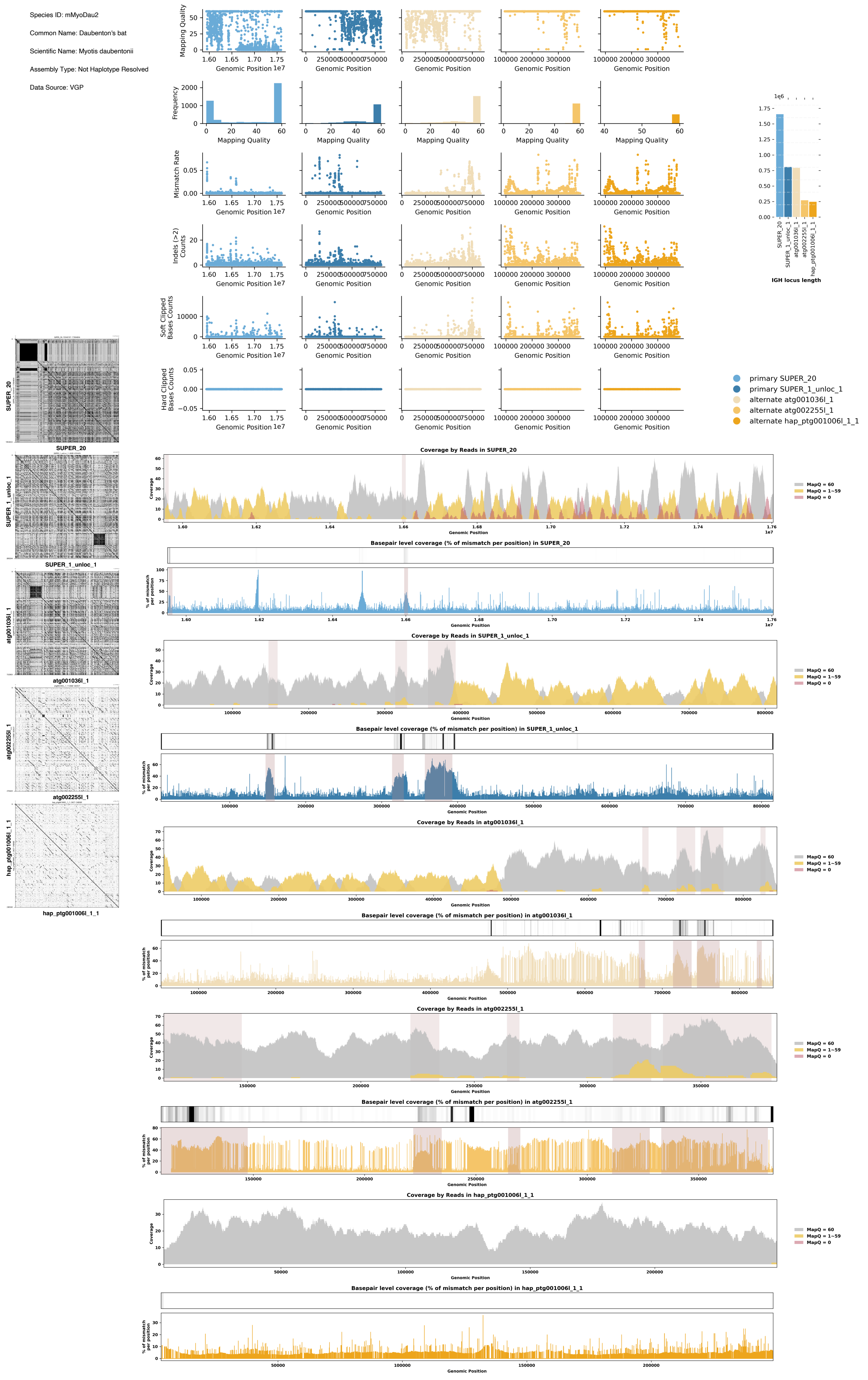

Species ID: mMyoYum1

Common Name: Yuma myotis

Scientific Name: Myotis yumanensis

Assembly Type: Haplotype Resolved

Data Source: CCGP

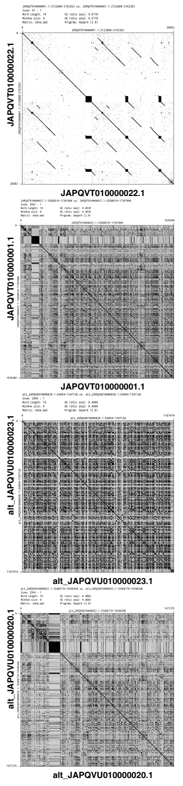


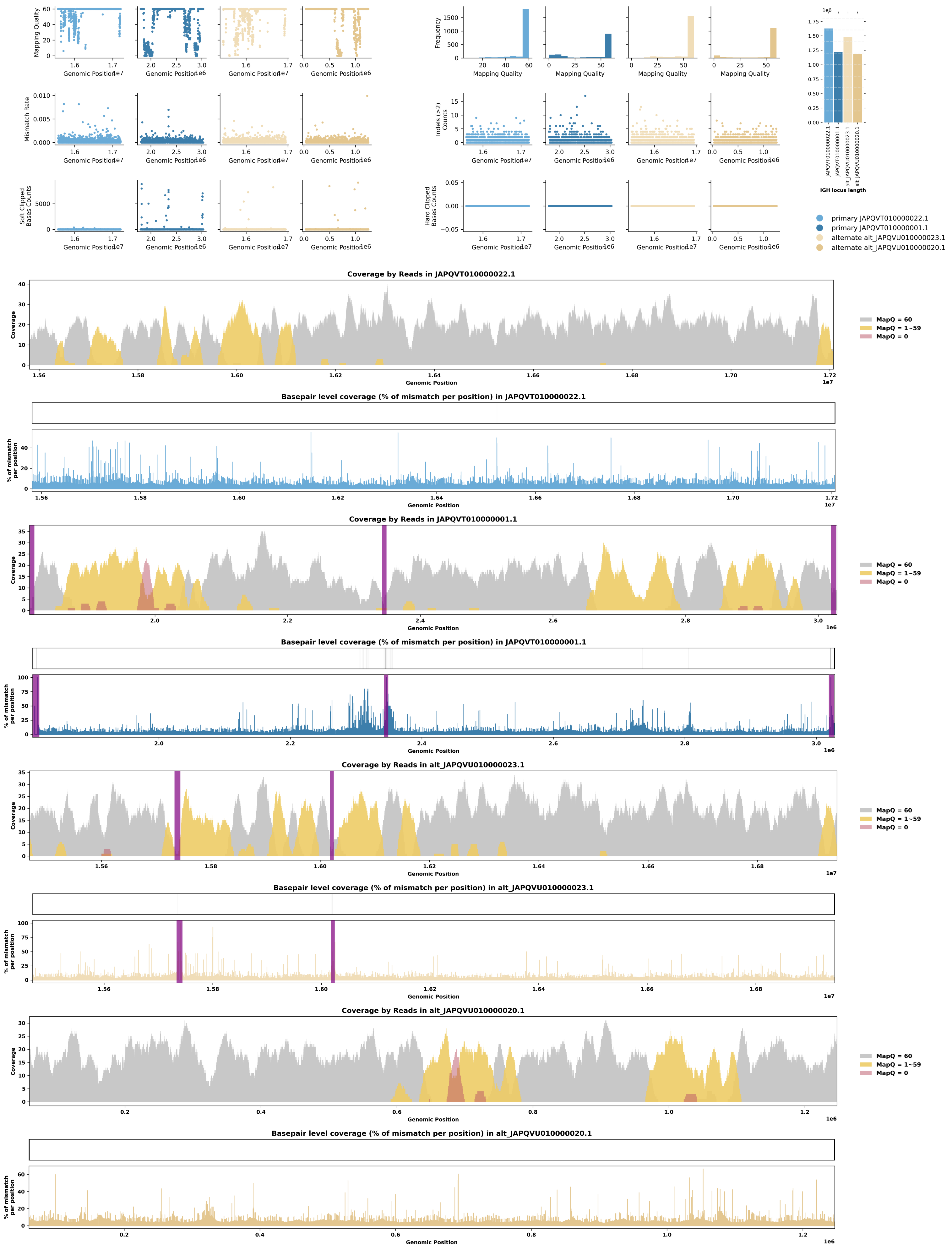

Species ID: mNeoNeb1

Common Name: Clouded Leopard

Scientific Name: Neofelis_nebulosa

Assembly Type: Not Haplotype Resolved

Data Source: VGP

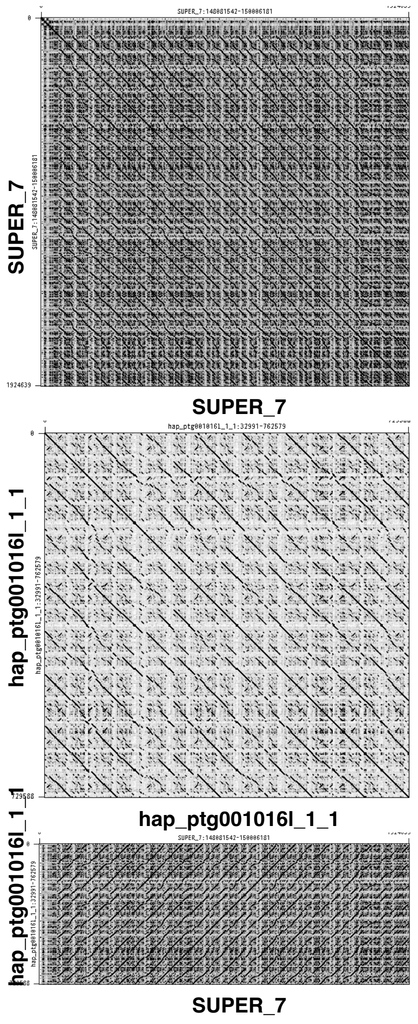


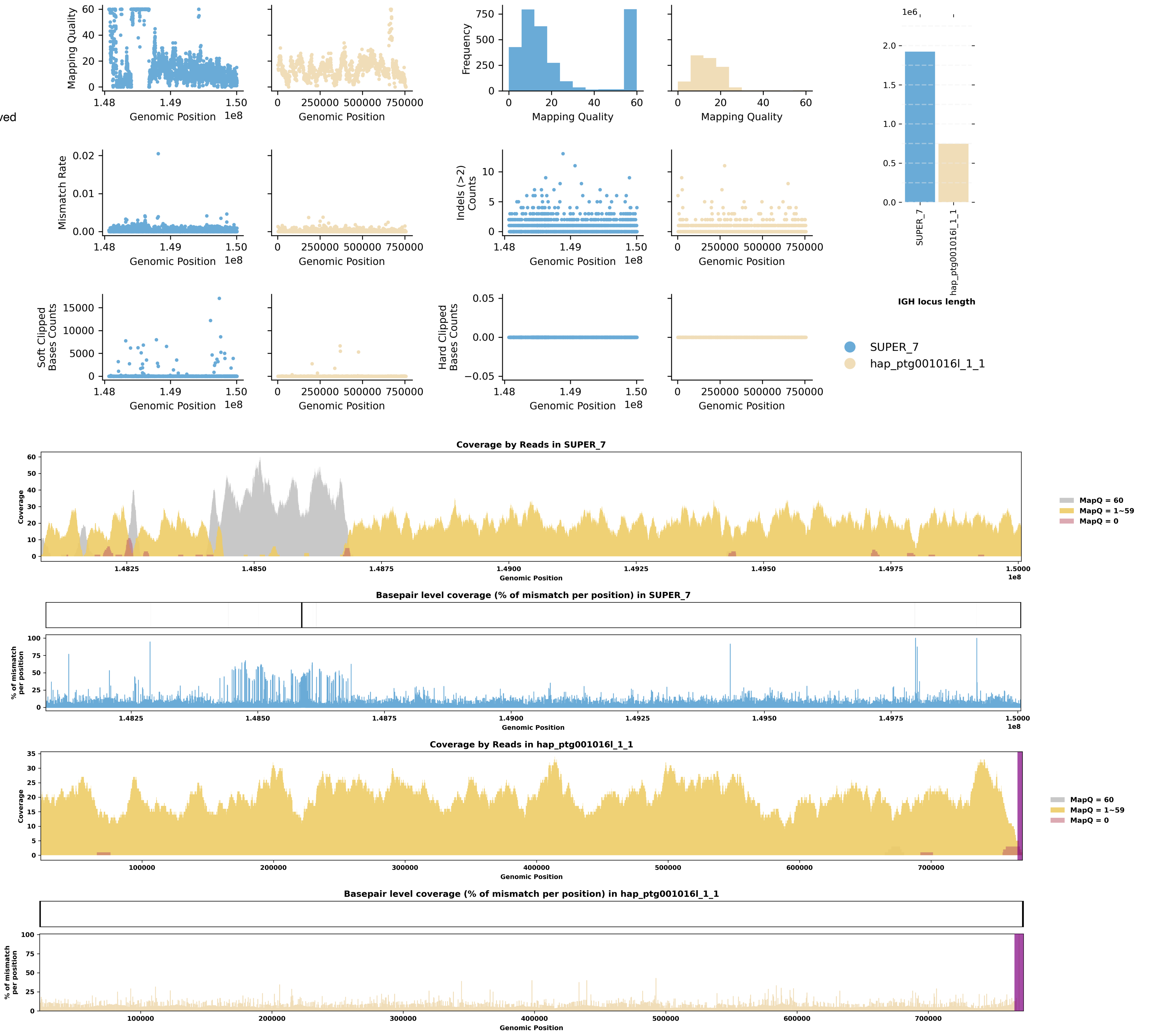

Species ID: mNycCou1

Common Name: slow loris

Scientific Name: Nycticebus coucang

Assembly Type: Not Haplotype Resolved

Data Source: VGP

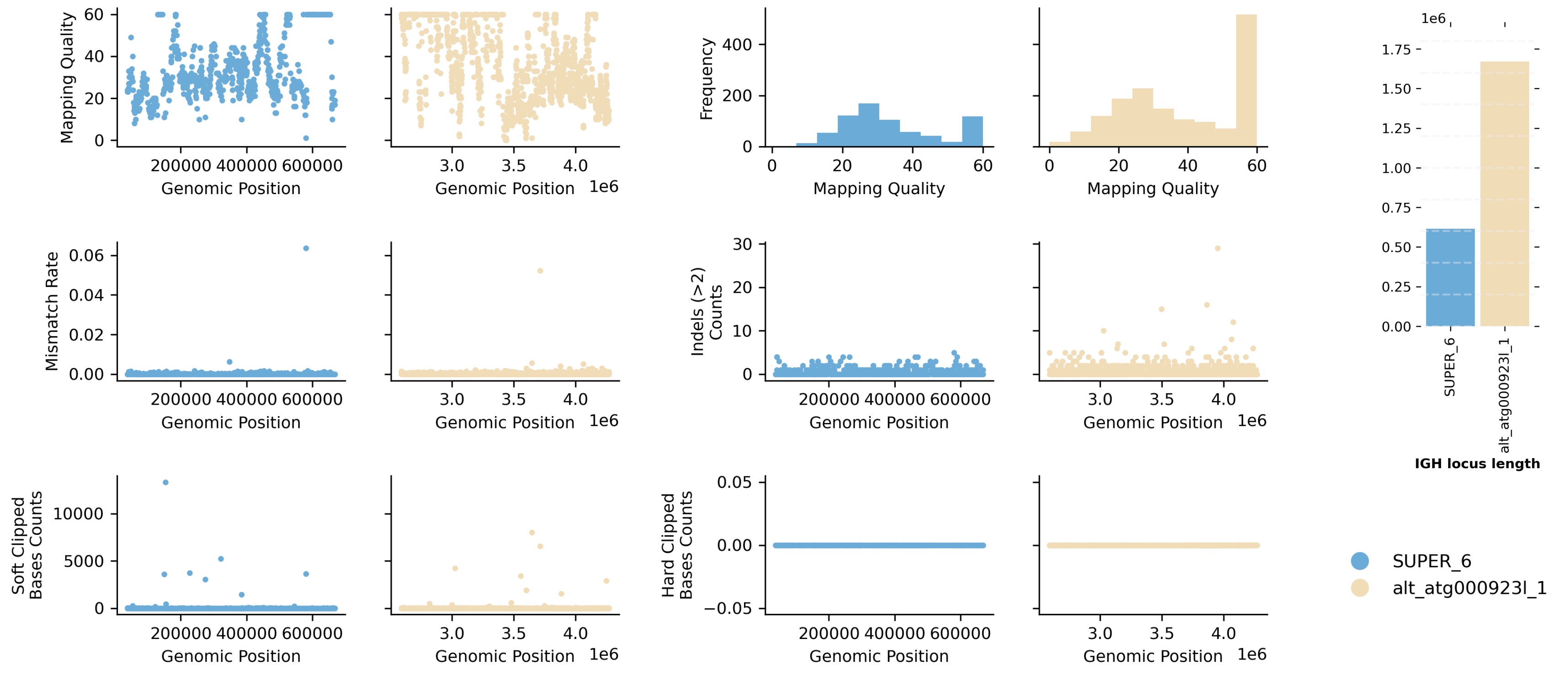


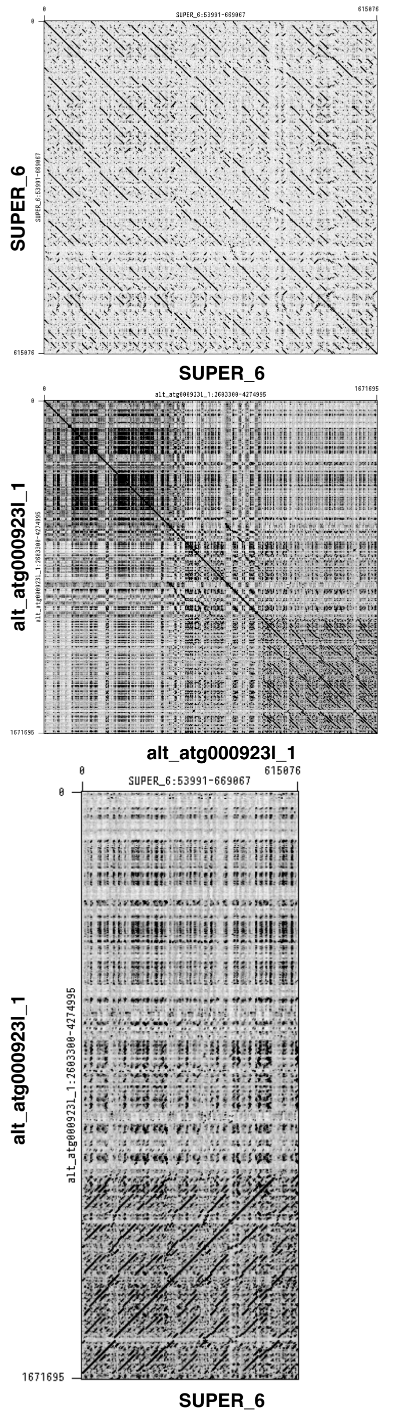


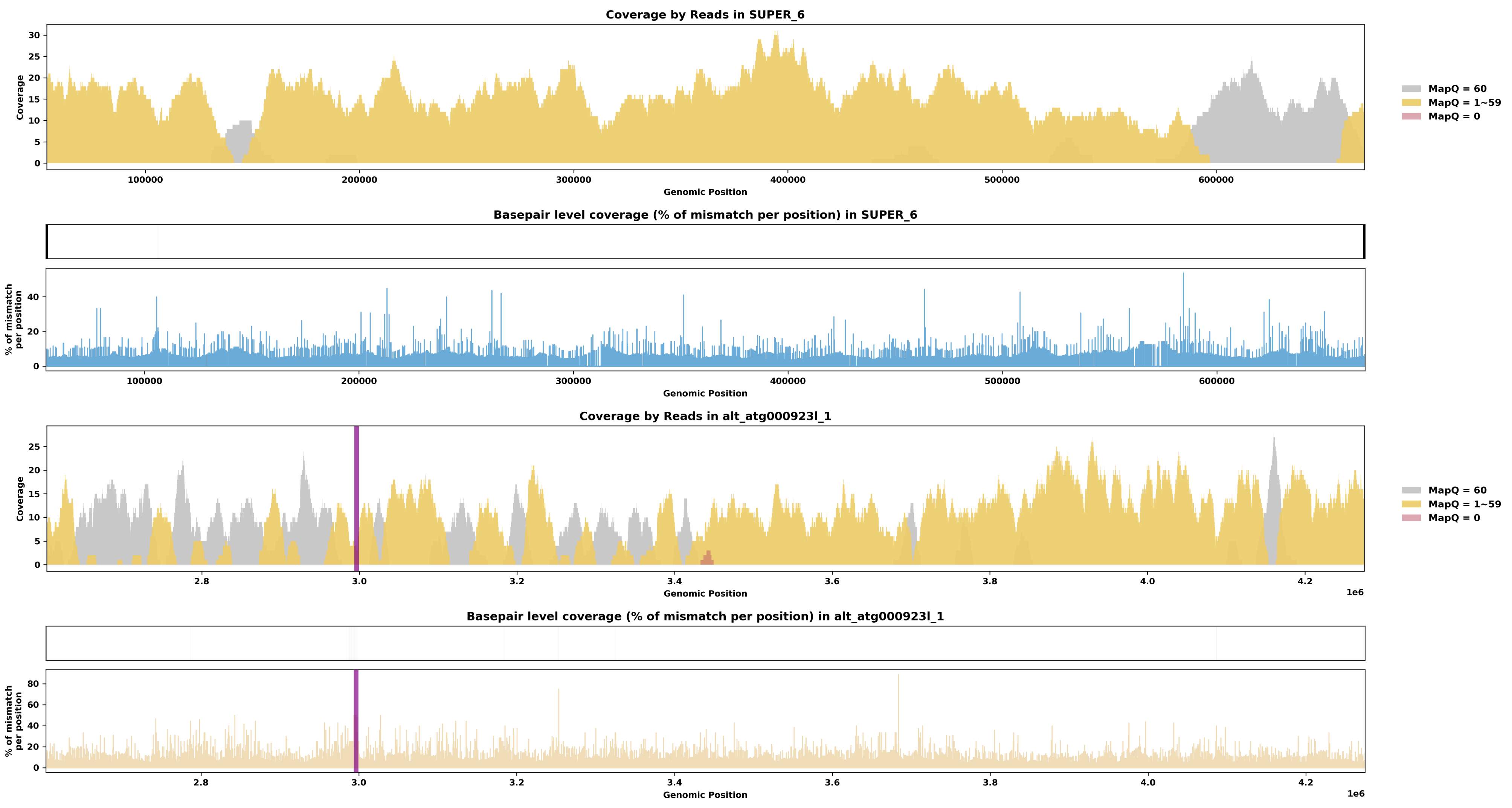

Species ID: mOrcOrc1

Common Name: killer whale

Scientific Name: Orcinus orca

Assembly Type: Not Haplotype Resolved

Data Source: VGP

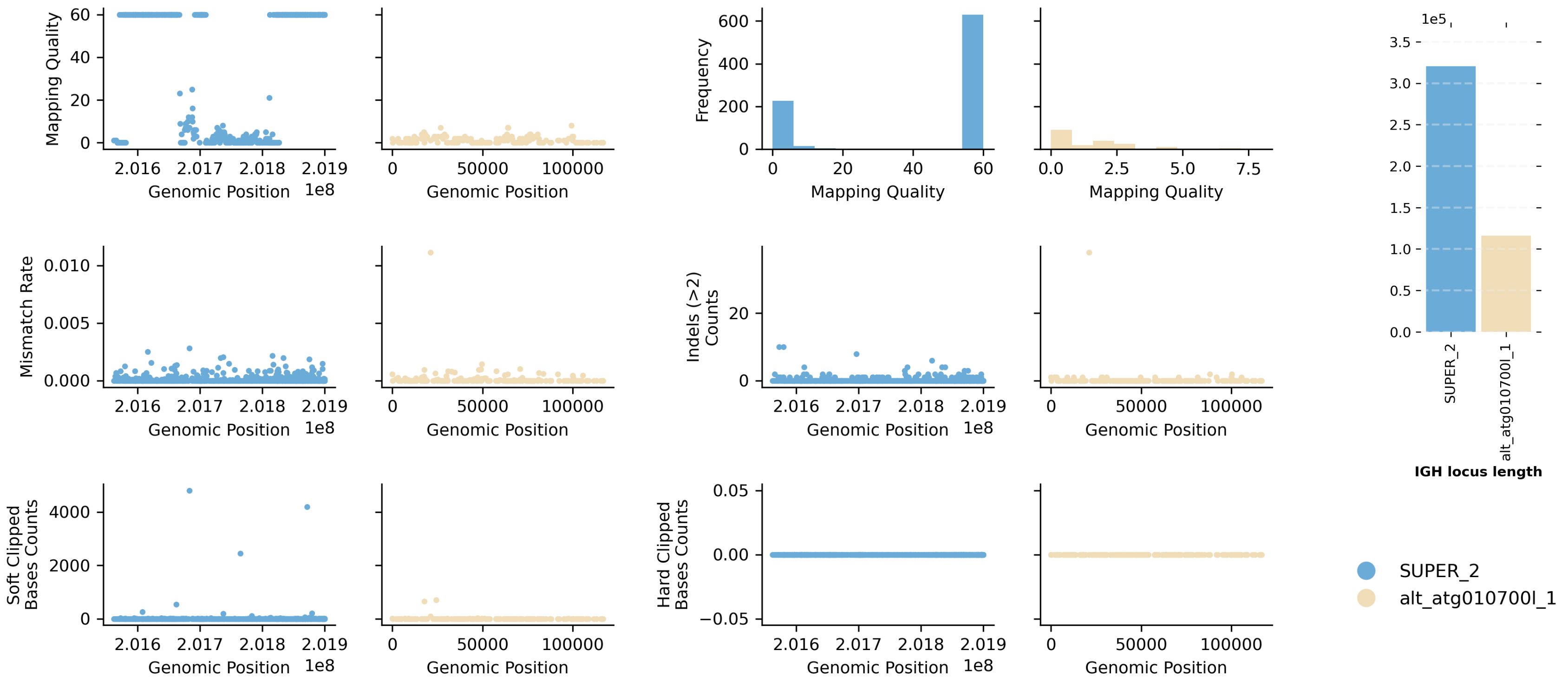


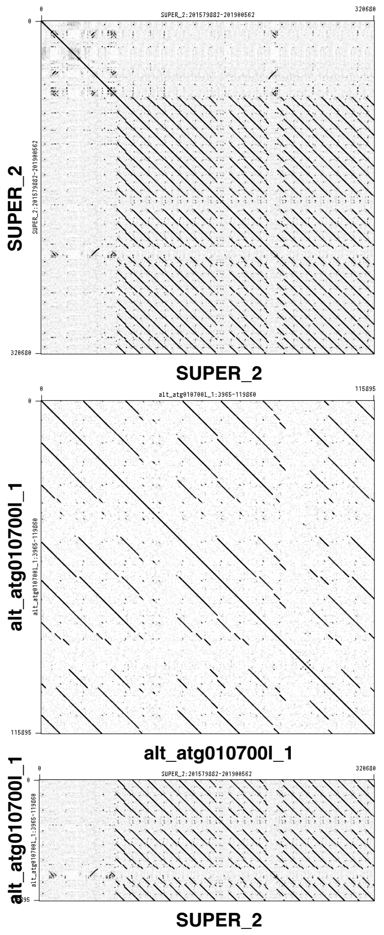


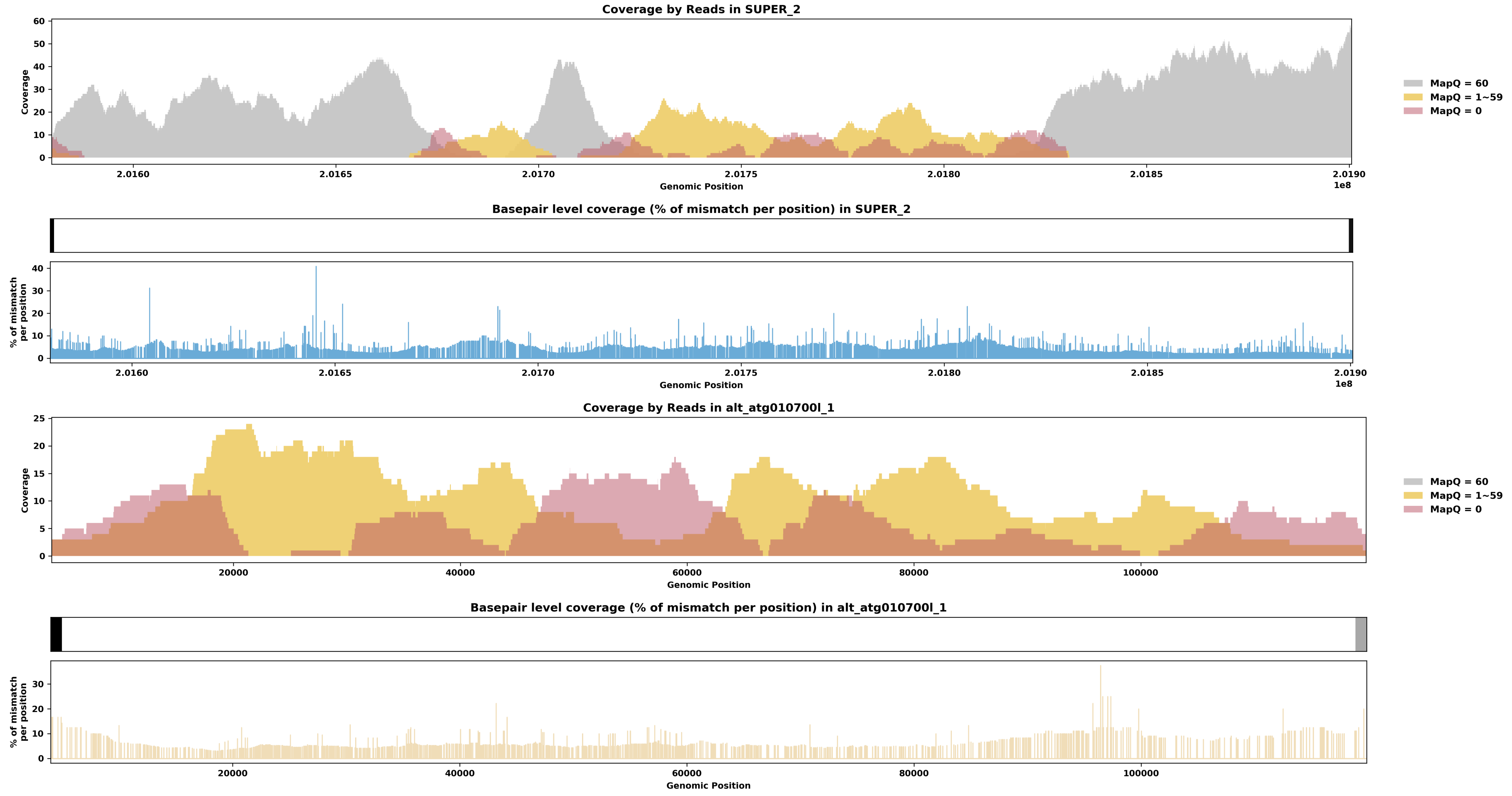

Species ID: mOryCun1

Common Name: rabbit

Scientific Name: Oryctolagus cuniculus

Assembly Type: Not Haplotype Resolved

Data Source: VGP

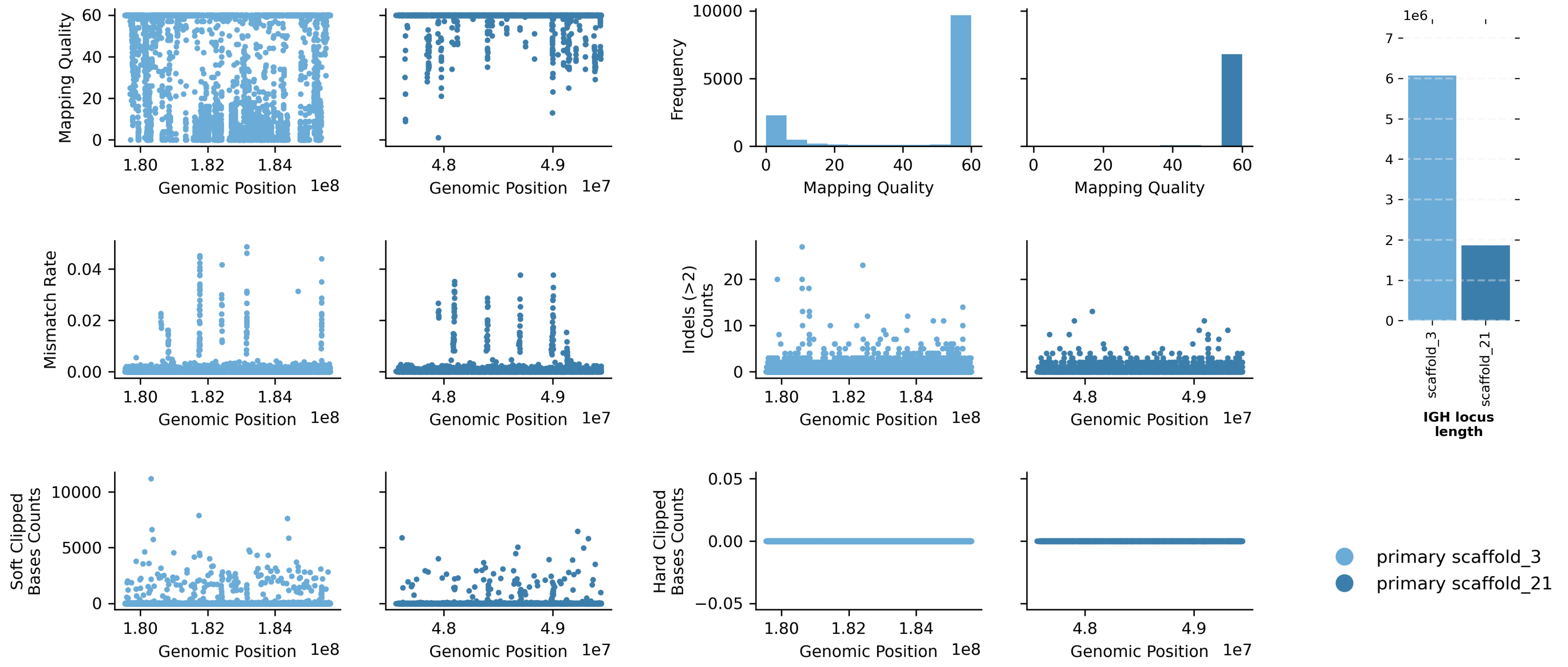


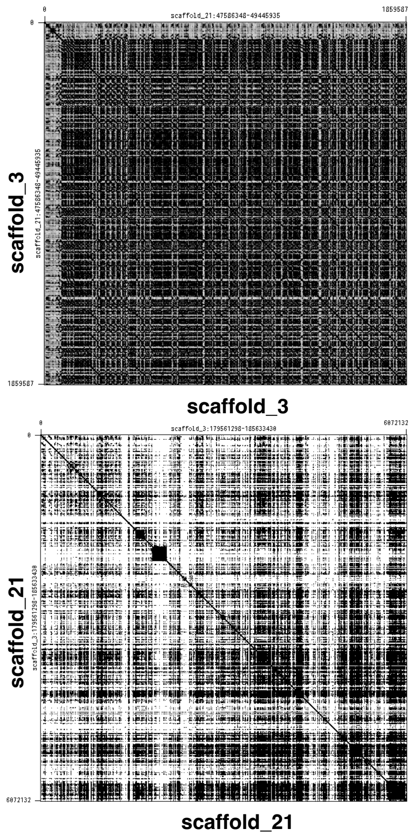


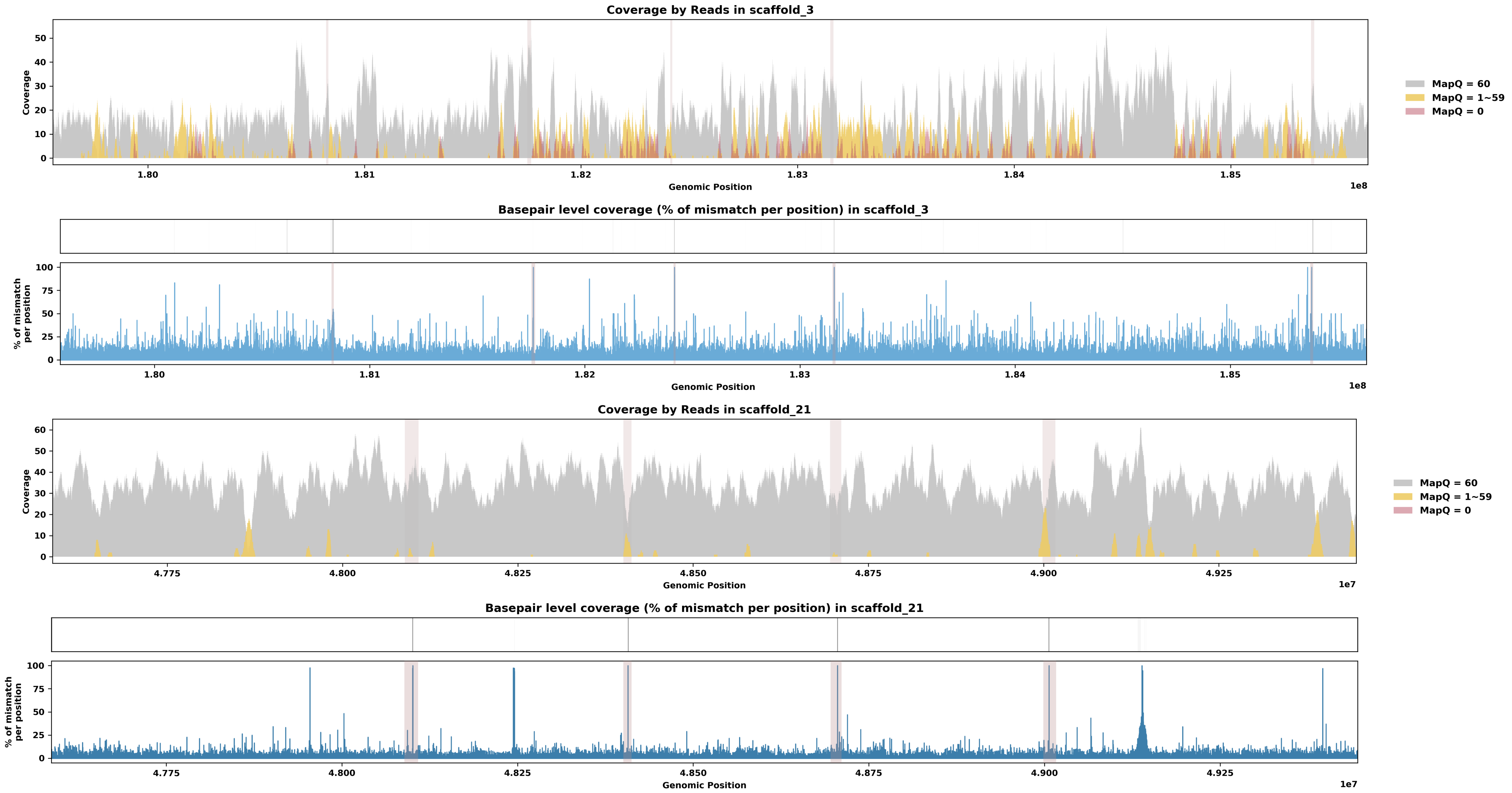

Species ID: mPanPan1

Common Name: Bonobo

Scientific Name: Pan paniscus

Assembly Type: Haplotype Resolved

Data Source: T2T Primate

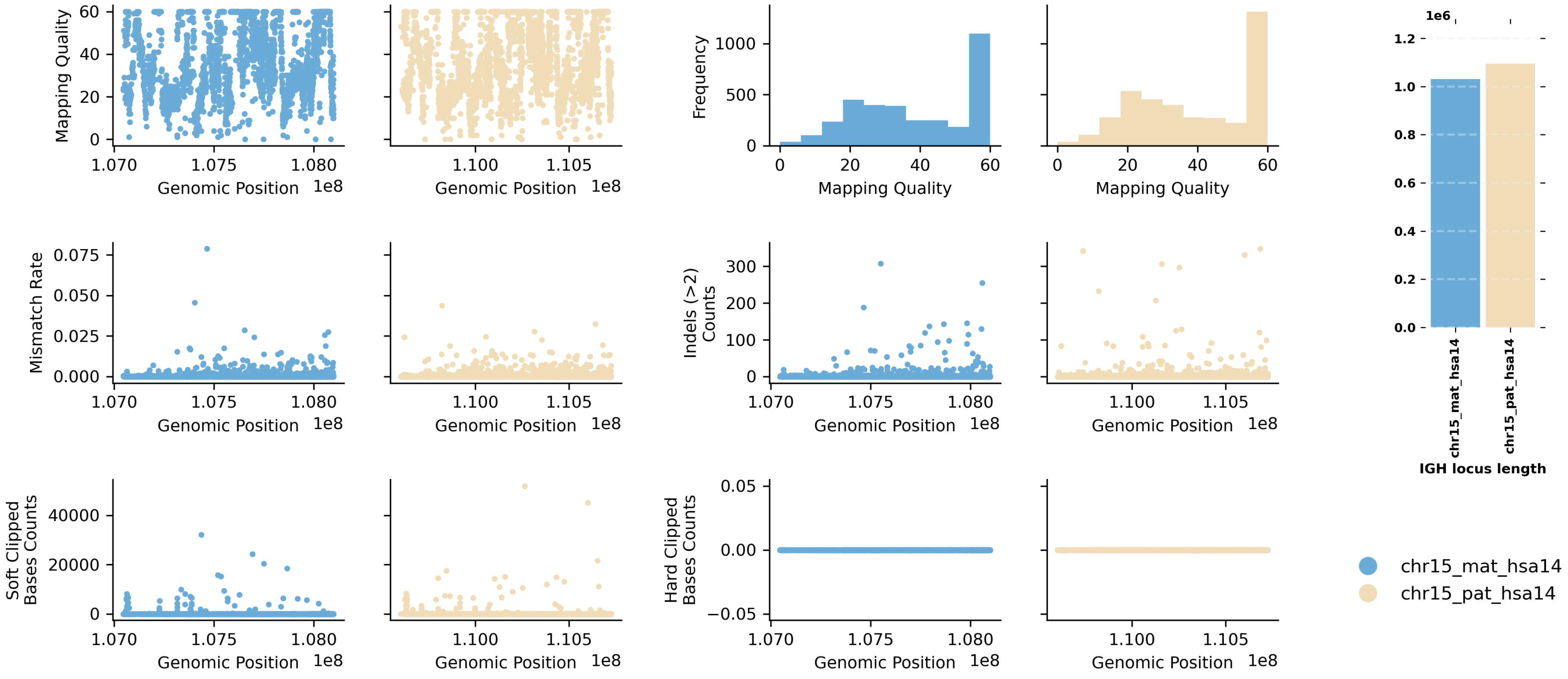


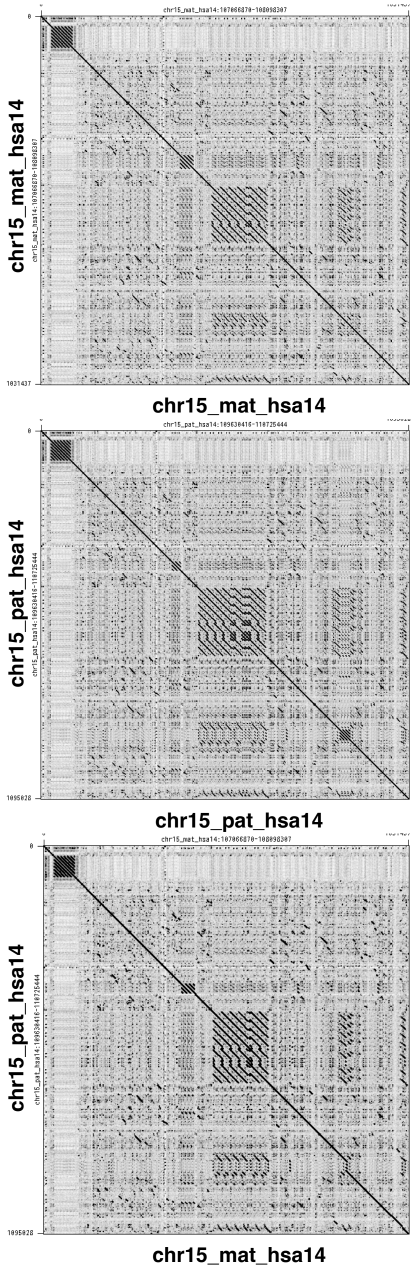


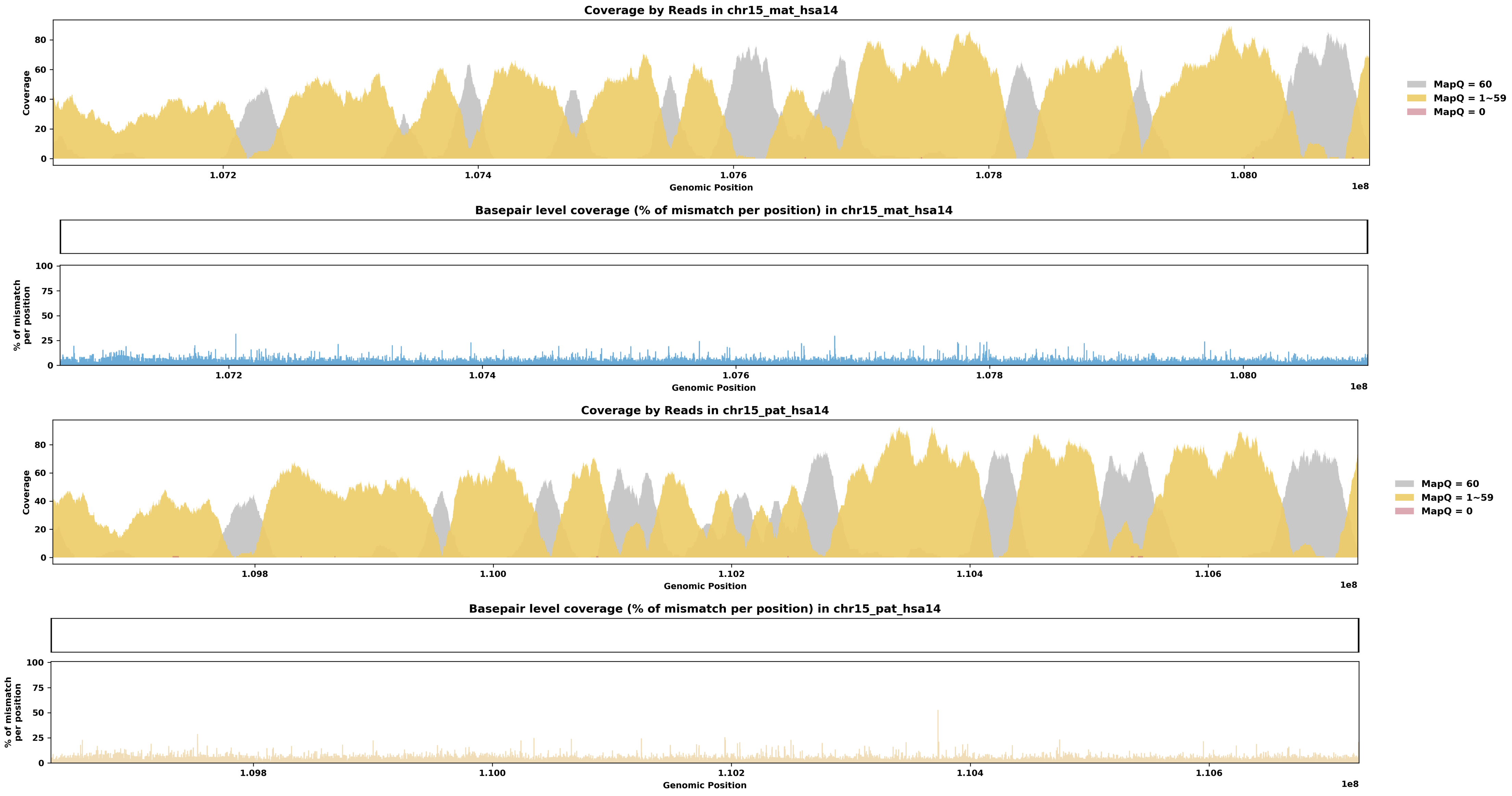

Species ID: mPerMan1

Common Name: deer mouse

Scientific Name: Peromyscus maniculatus

Assembly Type: Not Haplotype Resolved

Data Source: CCGP

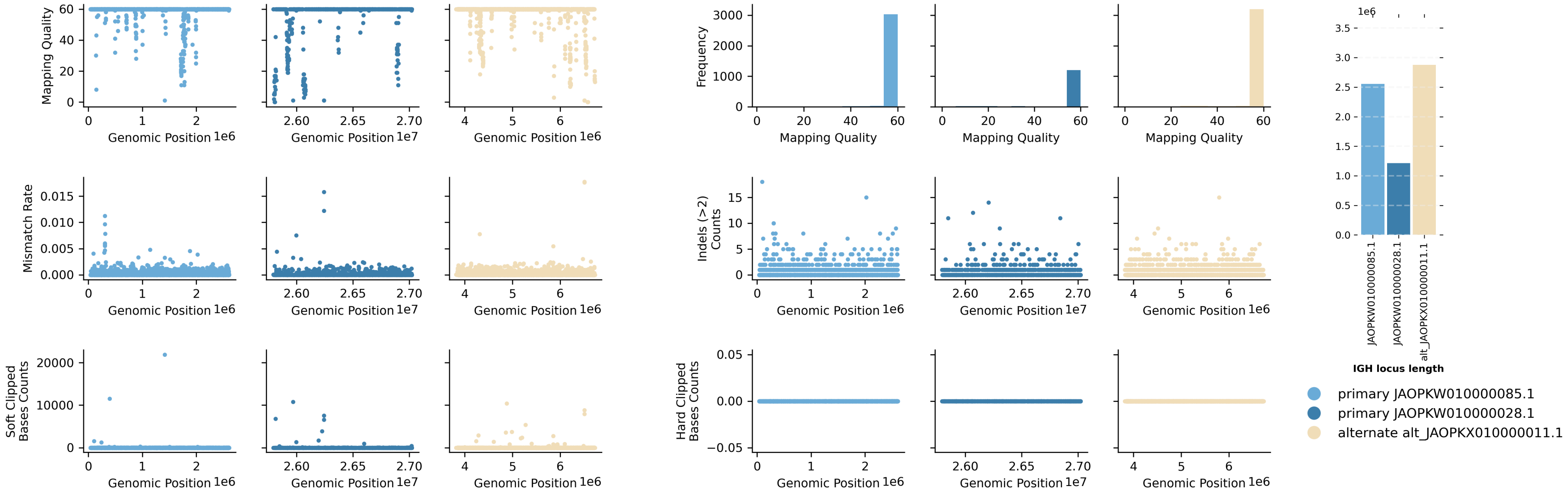


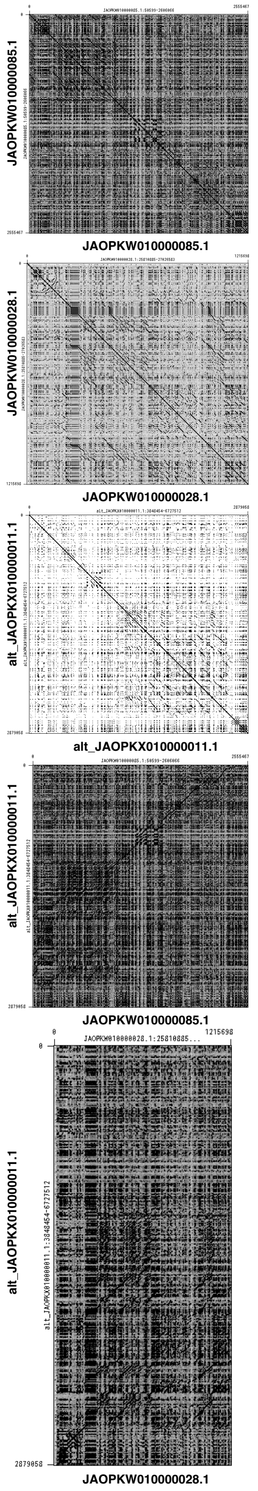


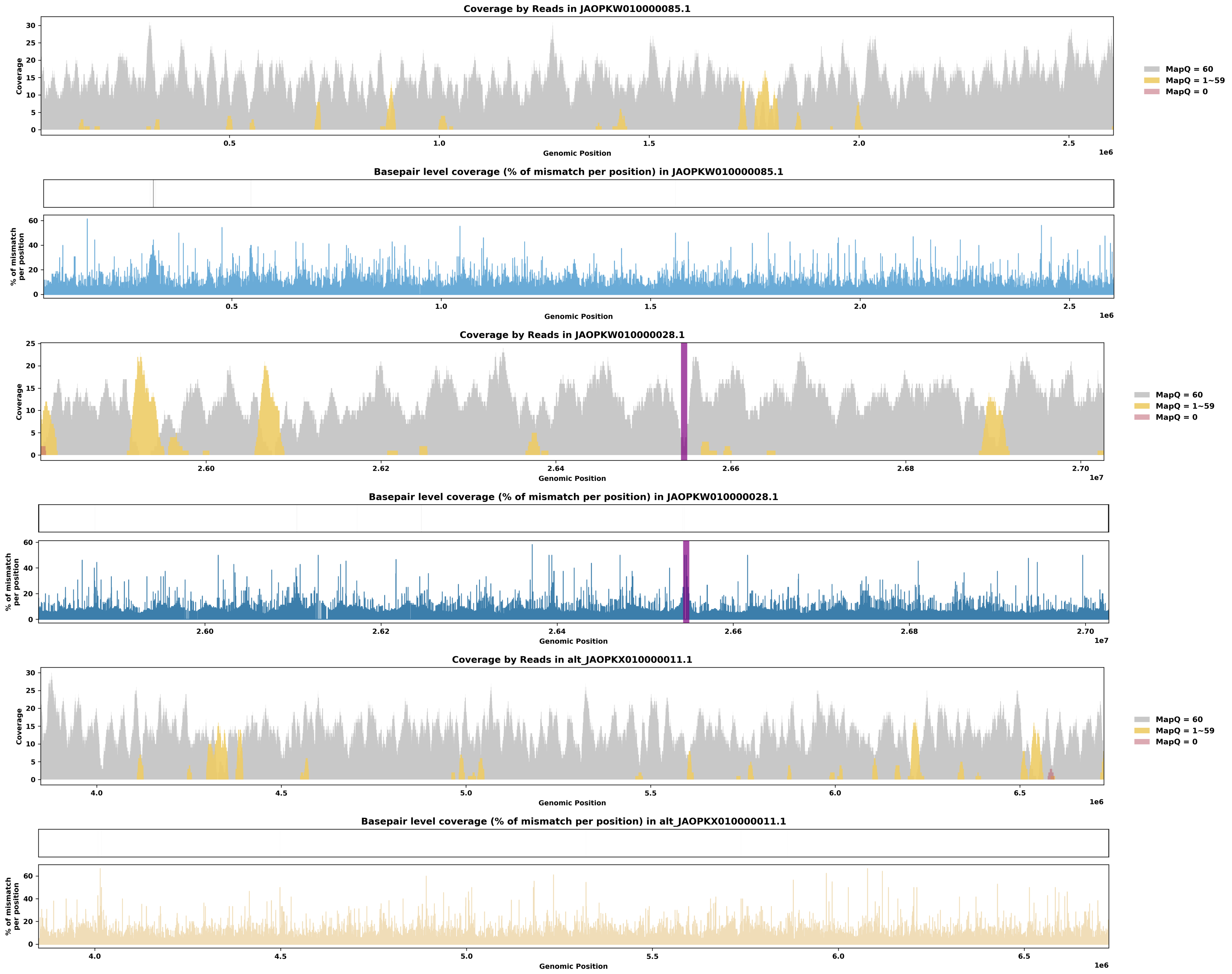

Species ID: mPhoPho1

Common Name: harbor porpoise

Scientific Name: Phocoena phocoena

Assembly Type: Not Haplotype Resolved

Data Source: VGP

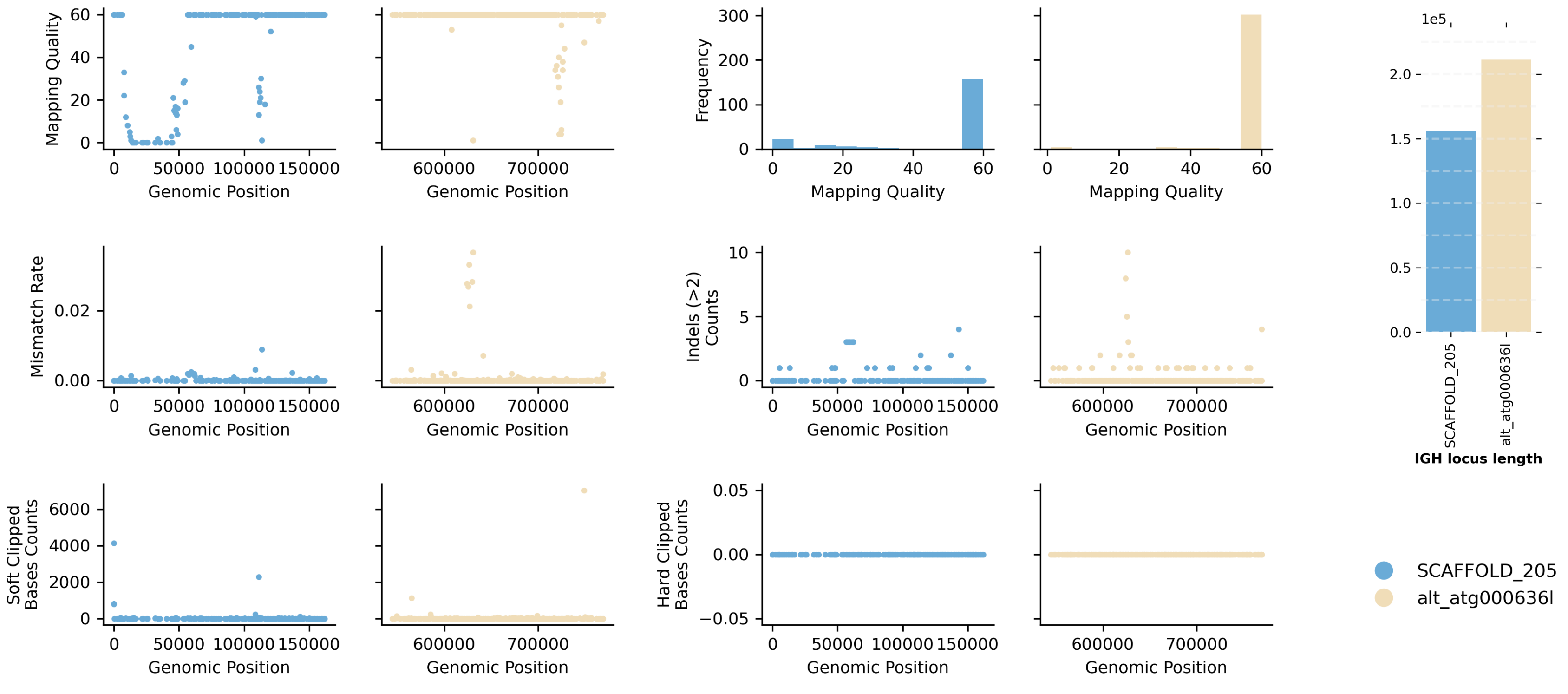


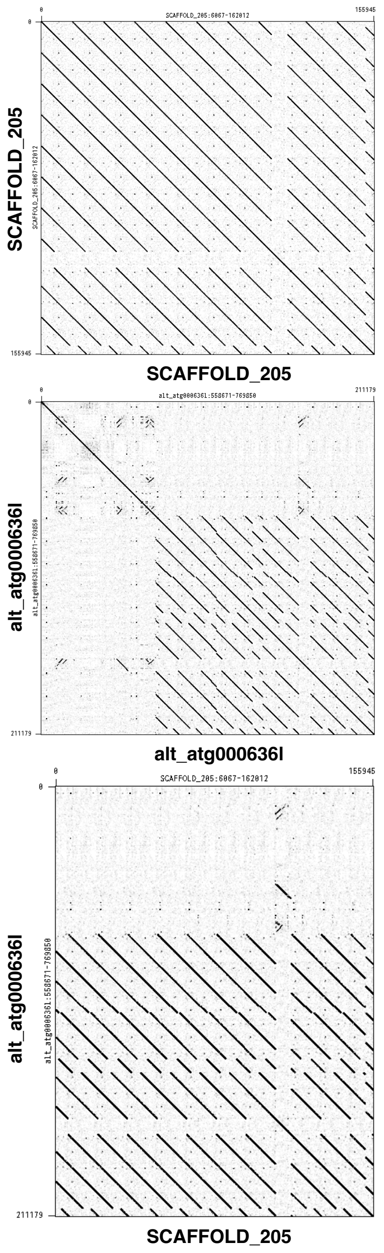


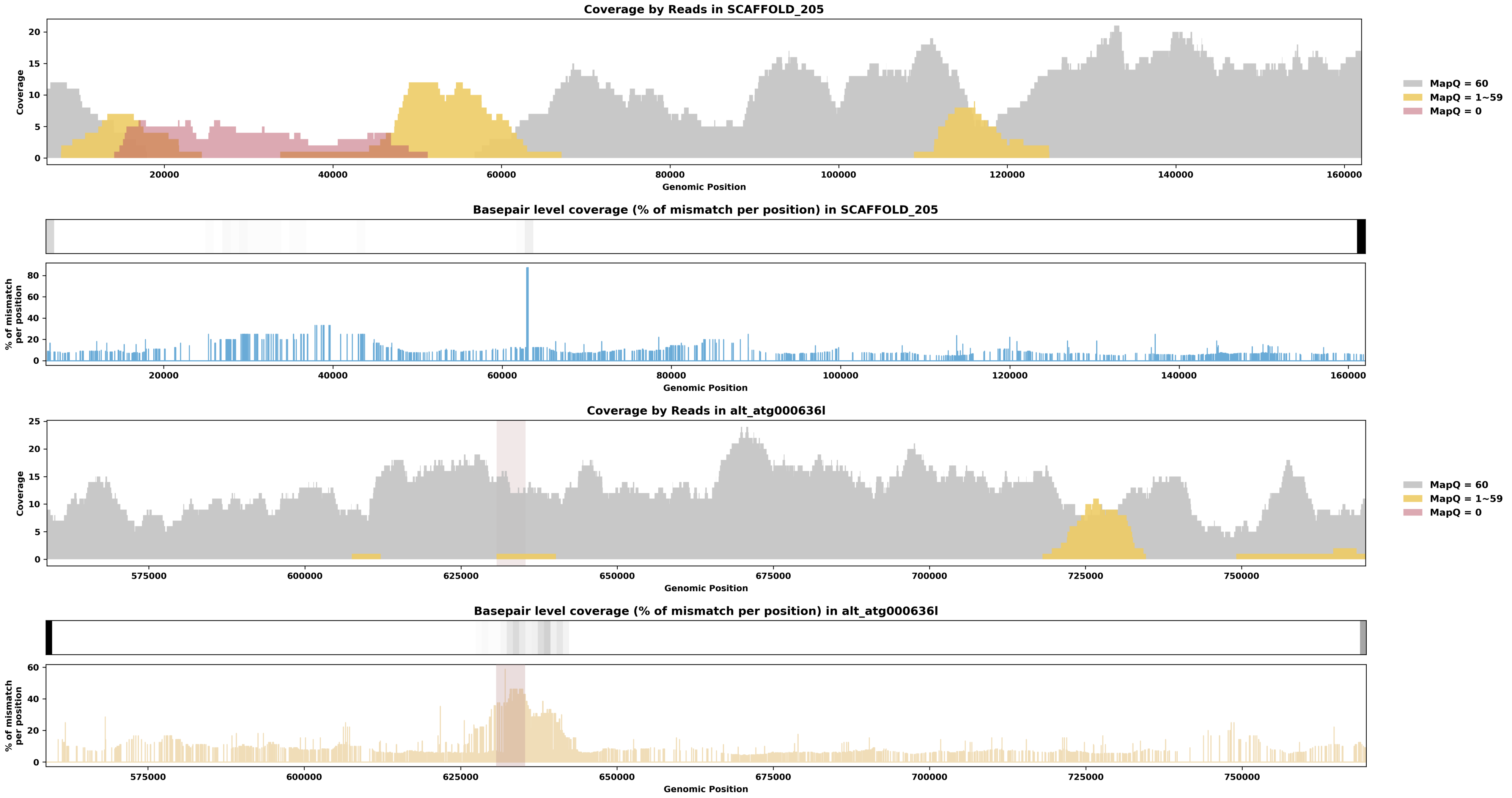

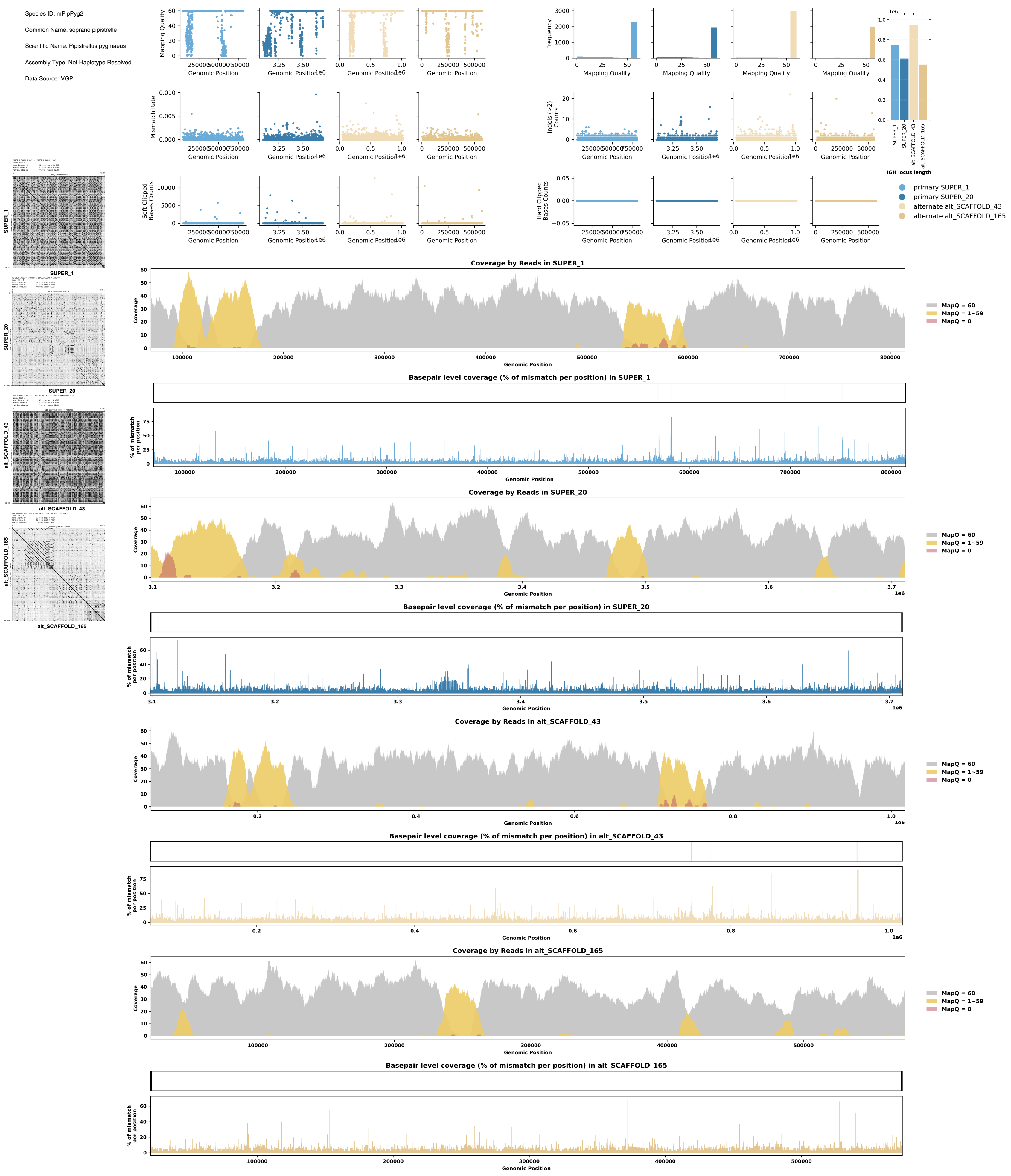
Species ID: mPipPyg2
Common Name: soprano pipistrelle
Scientific Name: Pipistrellus pygmaeus
Assembly Type: Not Haplotype Resolved
Data Source: VGP
Mapping Quality
Genomic Position
Frequency
Mismatch Rate
Indels (>2) Counts
Soft Clipped Bases Counts
Hard Clipped Bases Counts
IGH locus length
primary SUPER_1
primary SUPER_20
alternate alt_SCAFFOLD_43
alternate alt_SCAFFOLD_165
SUPER_1
SUPER_20
alt_SCAFFOLD_43
alt_SCAFFOLD_165
Coverage by Reads in SUPER_1
Coverage
MapQ = 60
MapQ = 1~59
MapQ = 0
Basepair level coverage (% of mismatch per position) in SUPER_1
% of mismatch per position
Coverage by Reads in SUPER_20
Basepair level coverage (% of mismatch per position) in SUPER_20
Coverage by Reads in alt_SCAFFOLD_43
Basepair level coverage (% of mismatch per position) in alt_SCAFFOLD_43
Coverage by Reads in alt_SCAFFOLD_165
Basepair level coverage (% of mismatch per position) in alt_SCAFFOLD_165

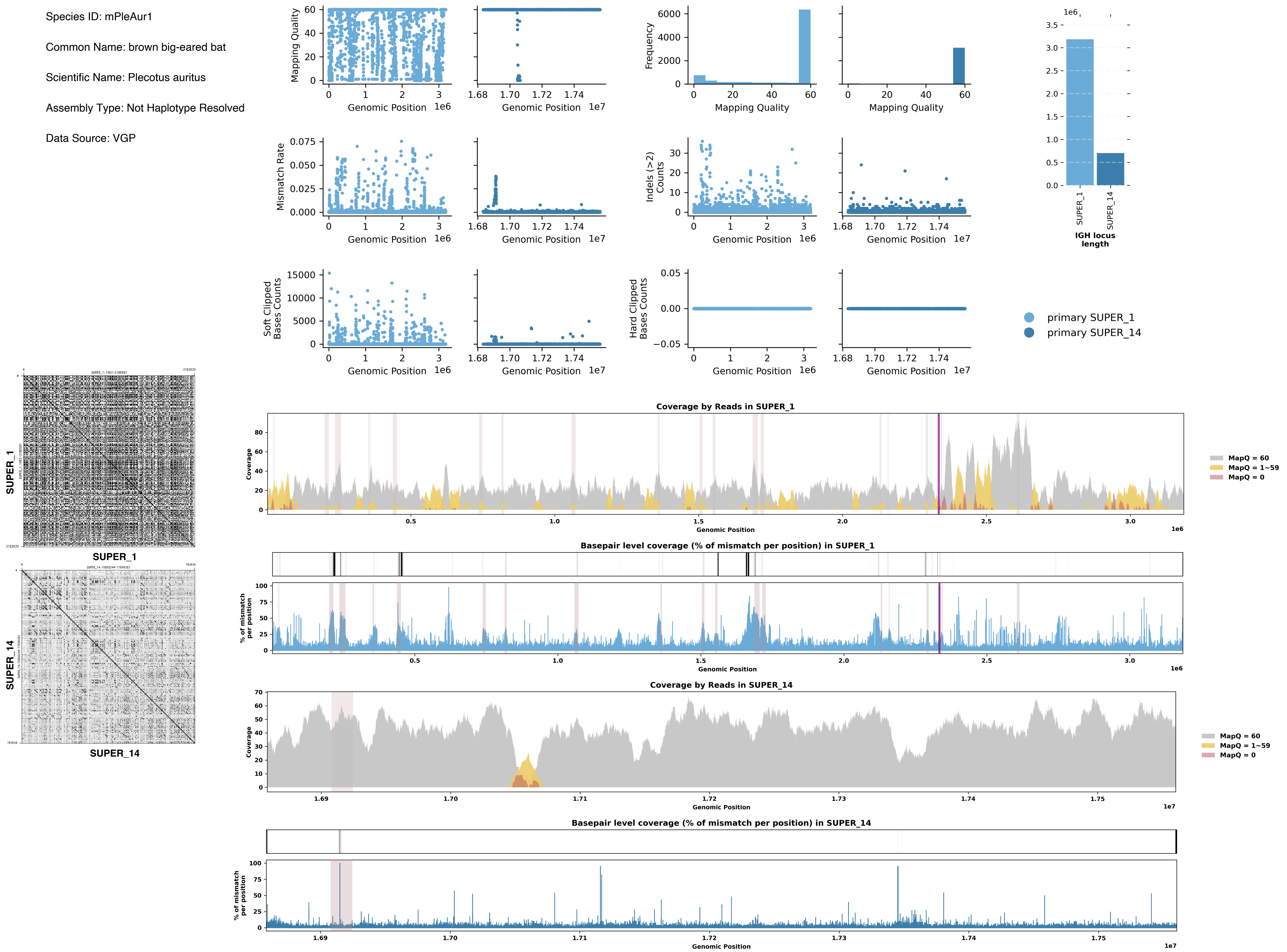
Species ID: mPleAur1
Common Name: brown big-eared bat
Scientific Name: Plecotus auritus
Assembly Type: Not Haplotype Resolved
Data Source: VGP
Mapping Quality
Genomic Position
Frequency
Mismatch Rate
Indels (>2) Counts
Soft Clipped Bases Counts
Hard Clipped Bases Counts
IGH locus length
SUPER_1
SUPER_14
primary SUPER_1
primary SUPER_14
Coverage by Reads in SUPER_1
Coverage
MapQ = 60
MapQ = 1~59
MapQ = 0
Basepair level coverage (% of mismatch per position) in SUPER_1
% of mismatch per position
Coverage by Reads in SUPER_14
Basepair level coverage (% of mismatch per position) in SUPER_14

Species ID: mPonAbe1

Common Name: Sumatran orangutan

Scientific Name: Pongo_abelii

Assembly Type: Haplotype Resolved

Data Source: T2T Primate

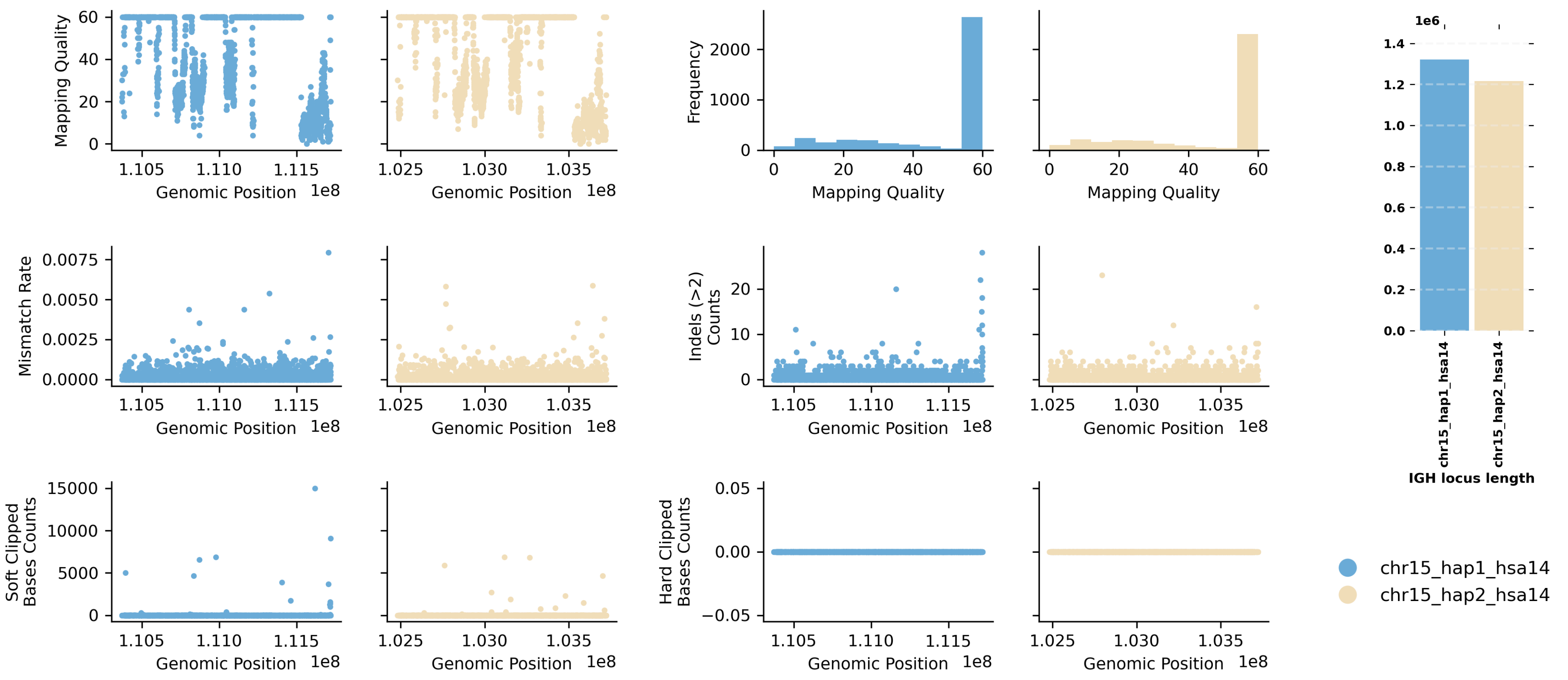


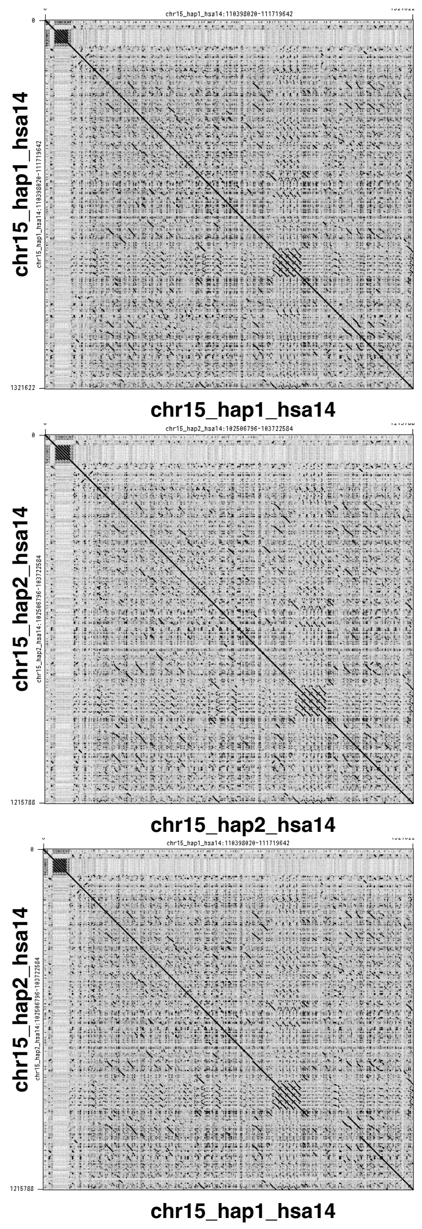


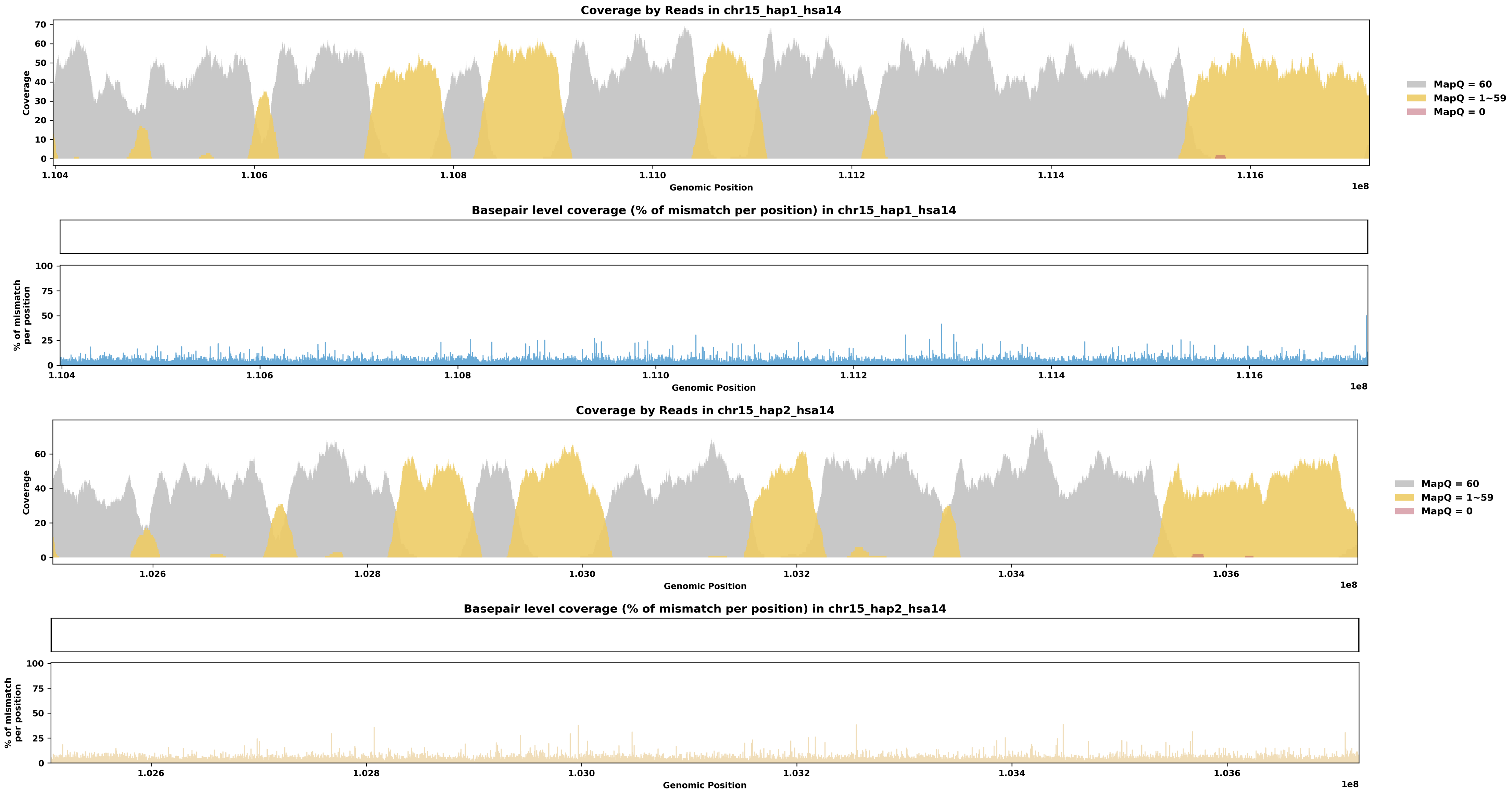

Species ID: mPonPyg2

Common Name: Bornean orangutan

Scientific Name: Pongo_pygmaeus

Assembly Type: Haplotype Resolved

Data Source: T2T Primate

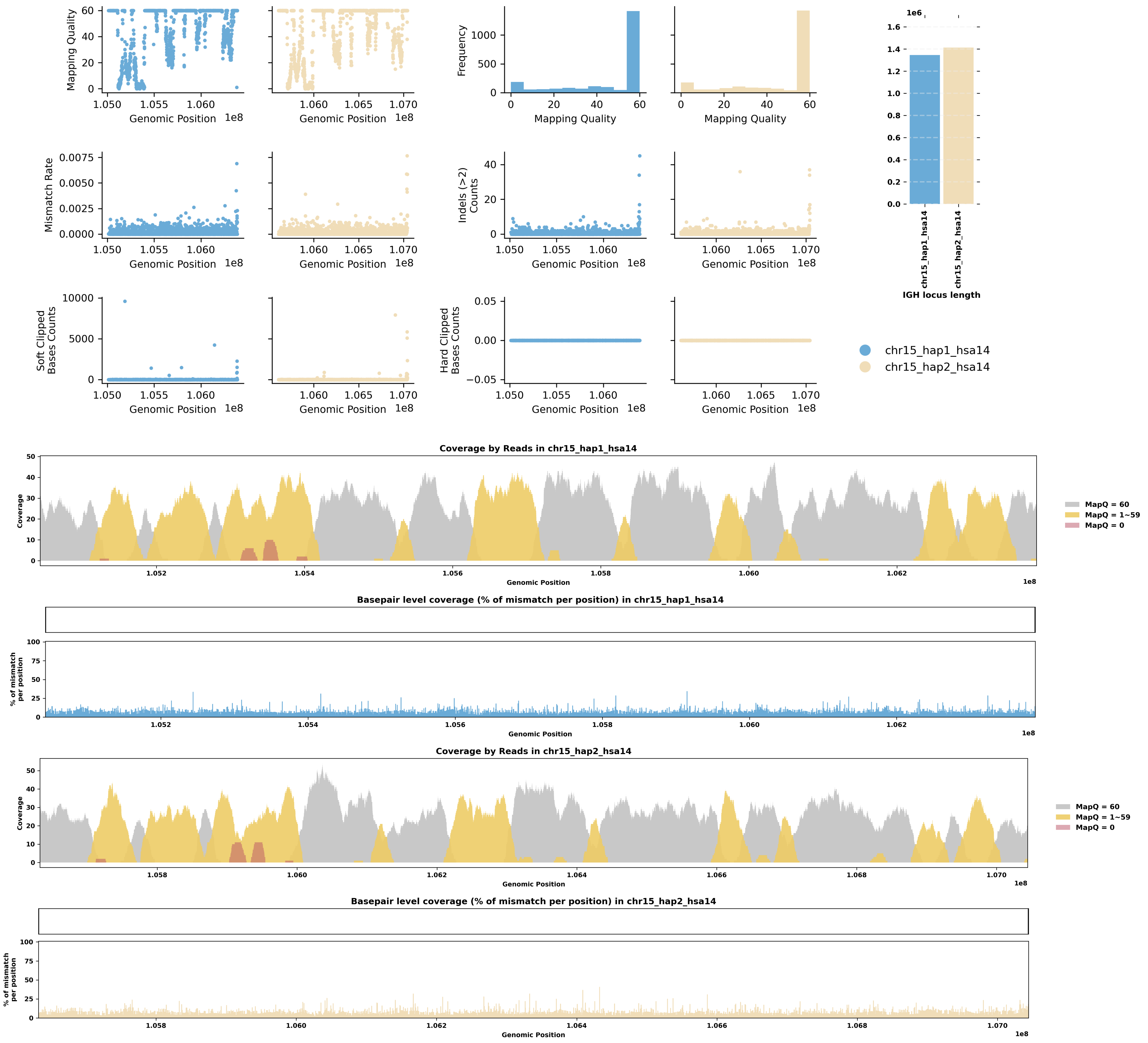


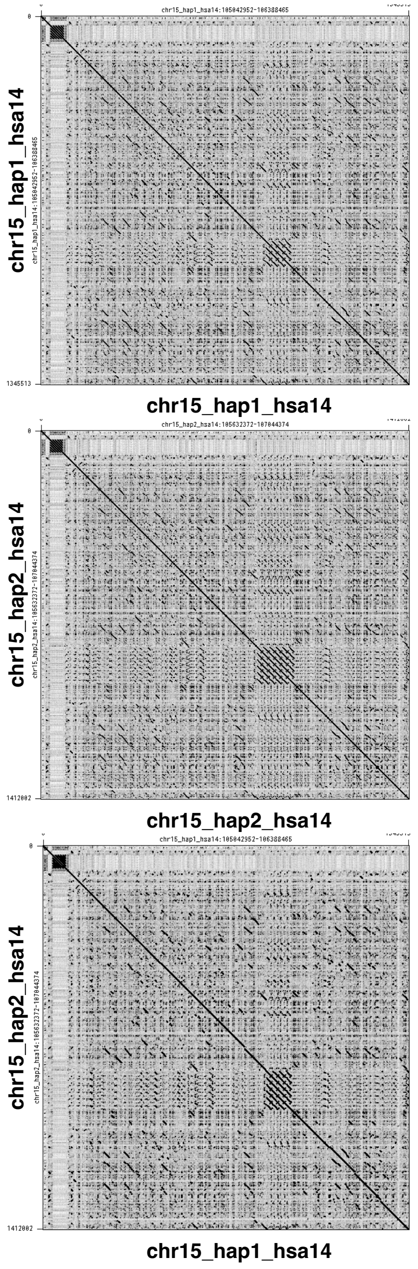

Species ID: mPseCra1

Common Name: false killer whale

Scientific Name: Pseudorca crassidens

Assembly Type: Haplotype Resolved

Data Source: VGP

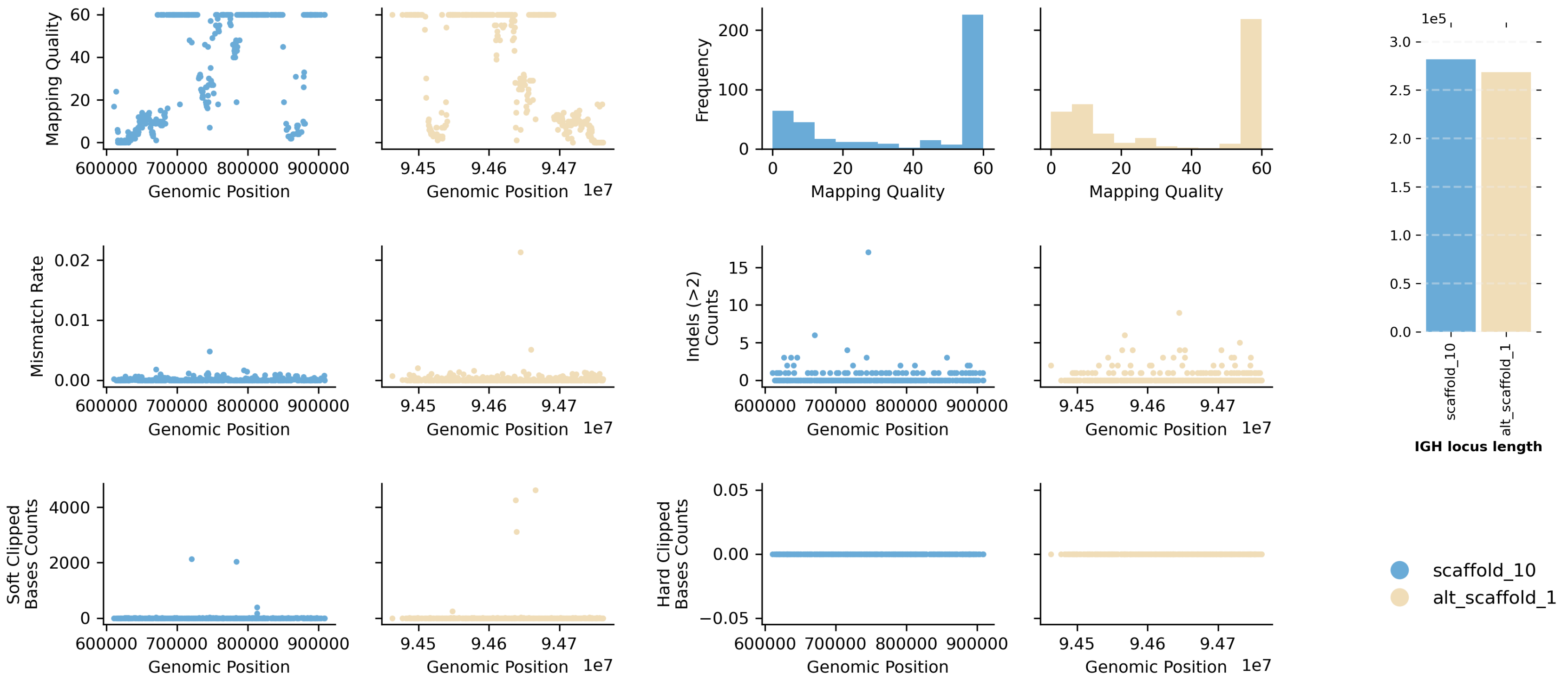


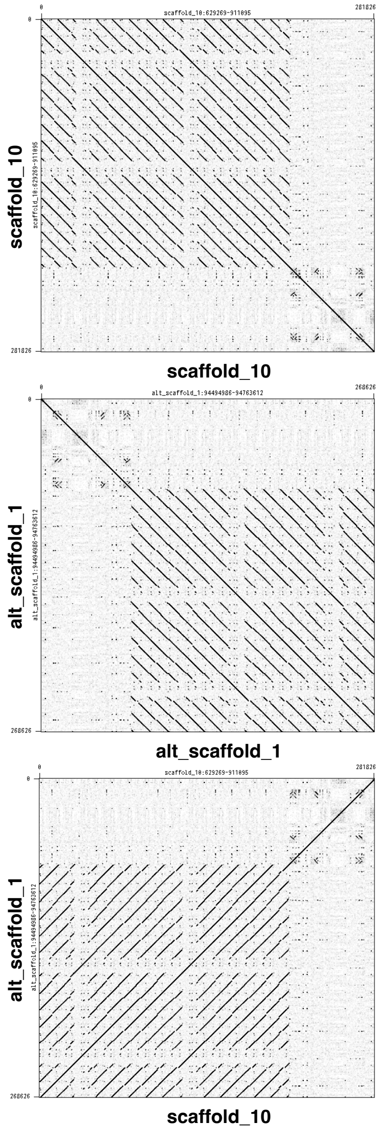


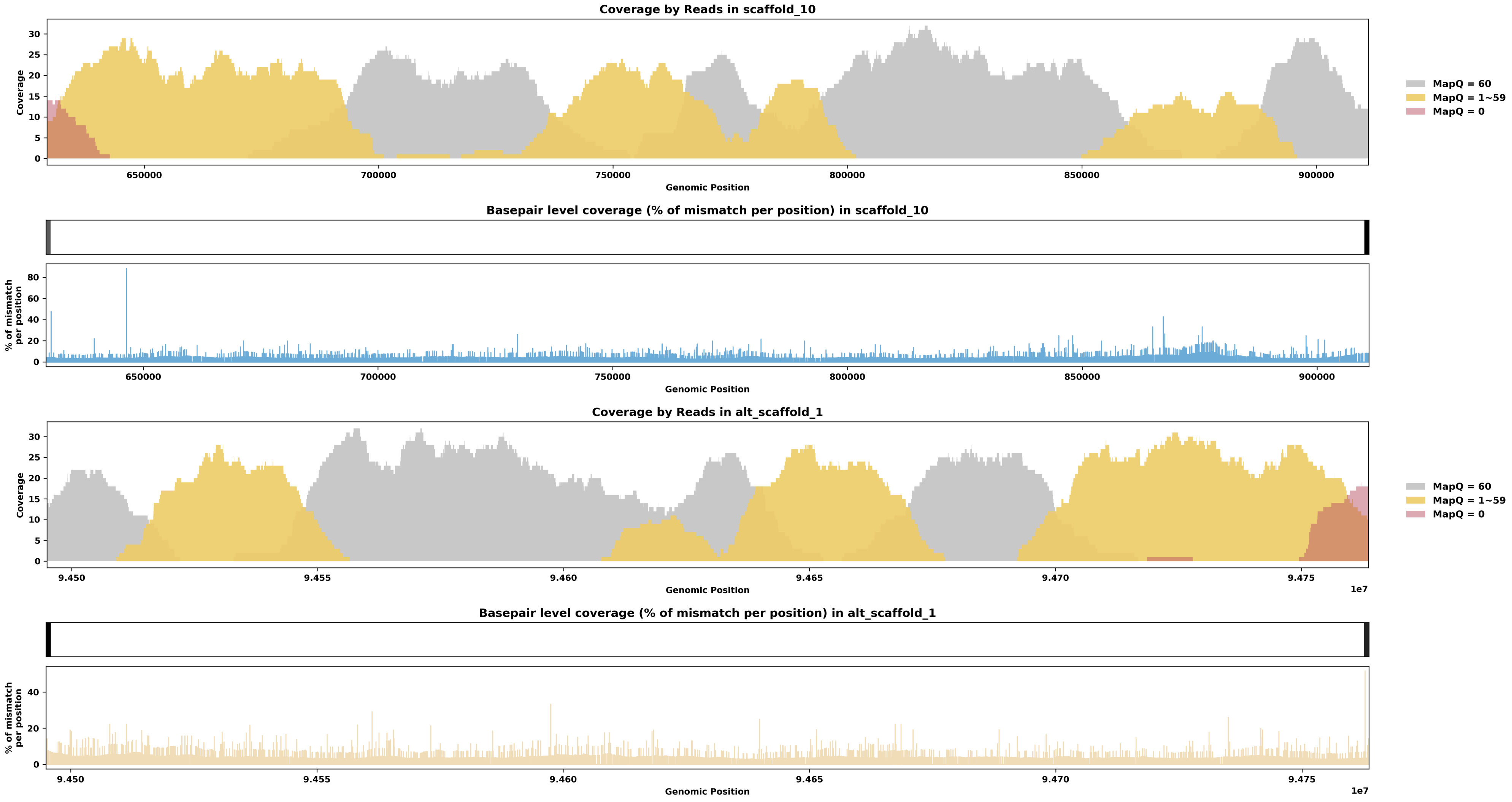

Species ID: mPumCon1

Common Name: Mountain Lion

Scientific Name: Puma concolor

Assembly Type: Haplotype Resolved

Data Source: CCGP

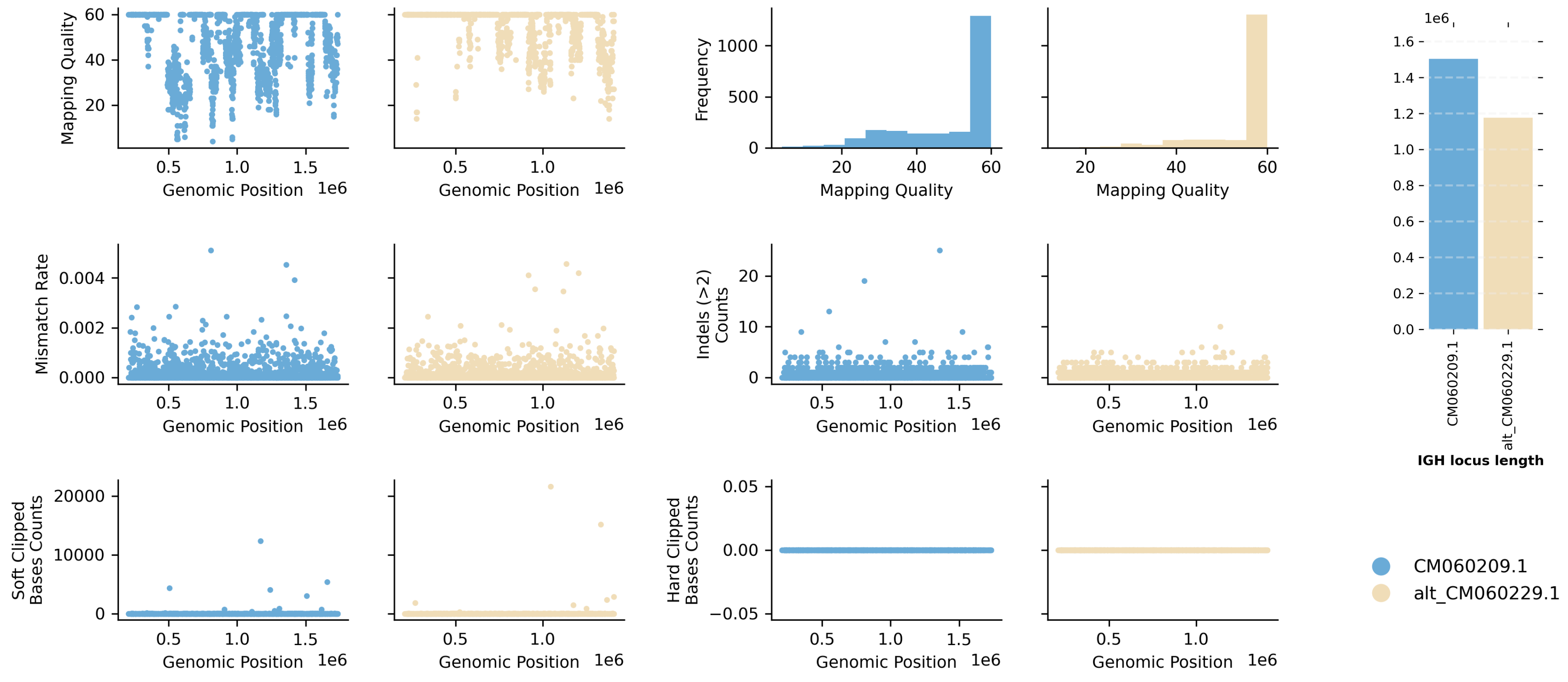


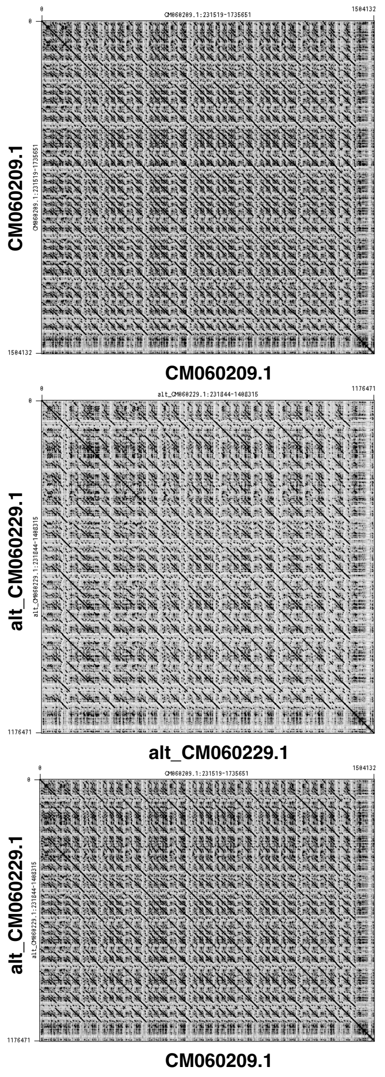


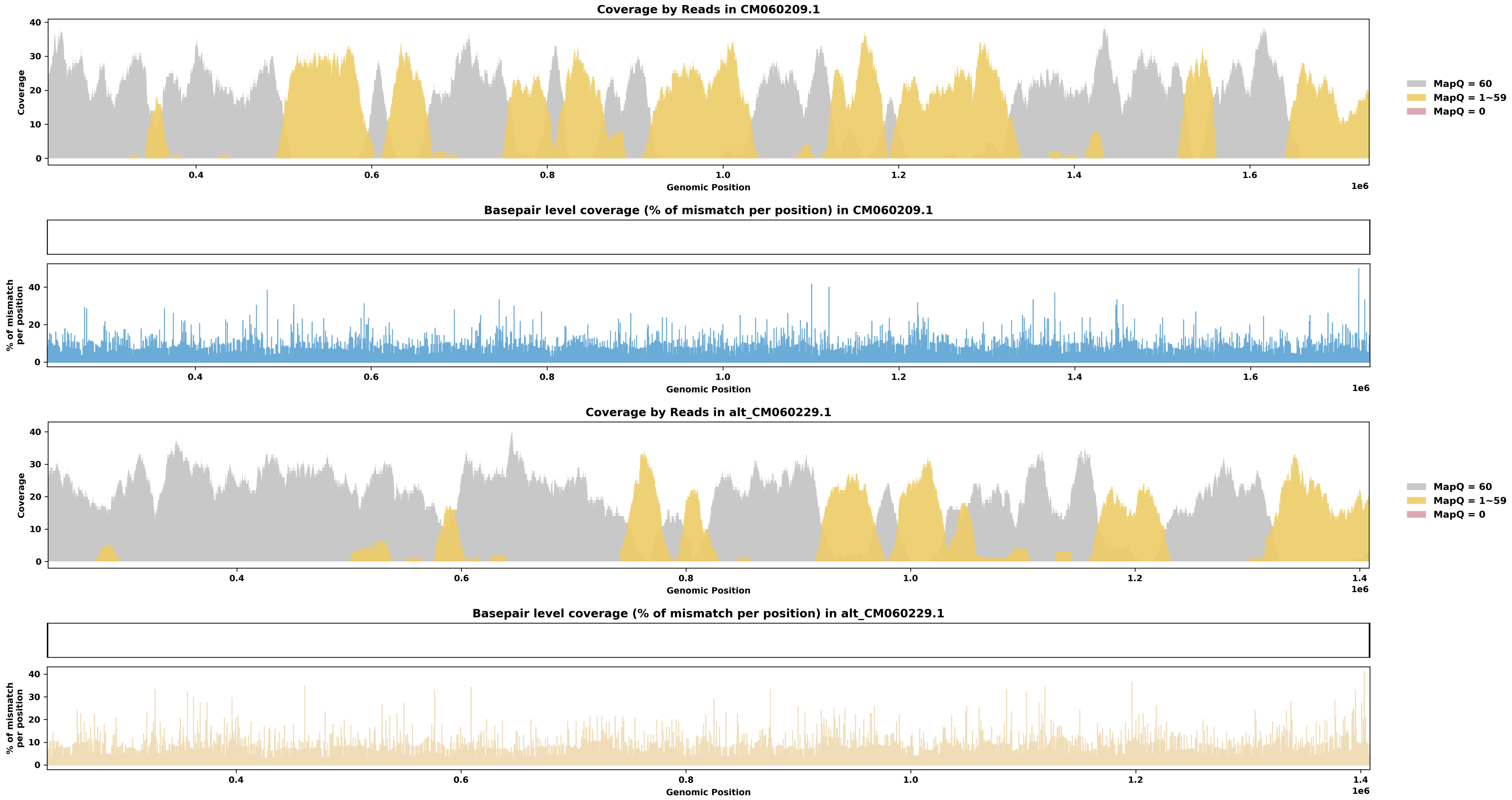

Species ID: mSorAra2

Common Name: Common shrew

Scientific Name: Sorex araneus

Assembly Type: Not Haplotype Resolved

Data Source: VGP

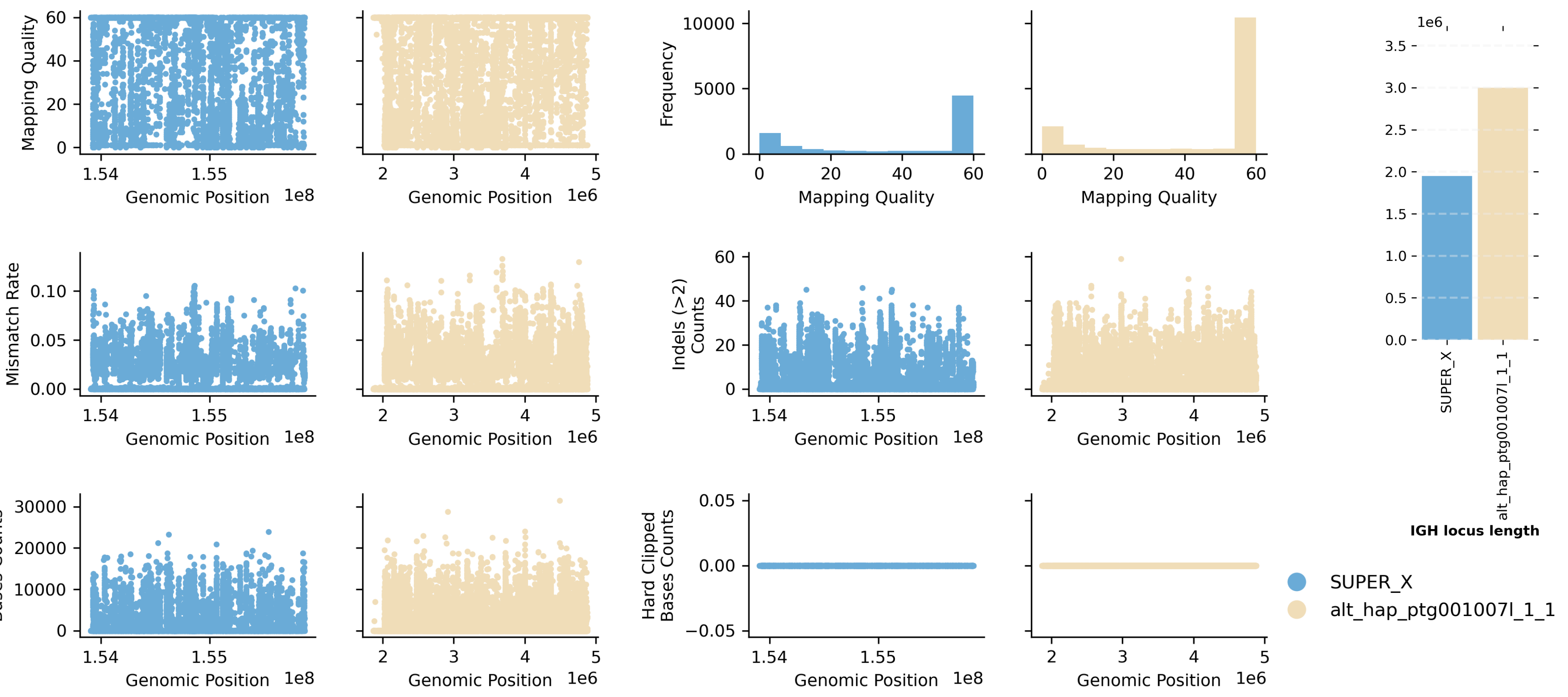


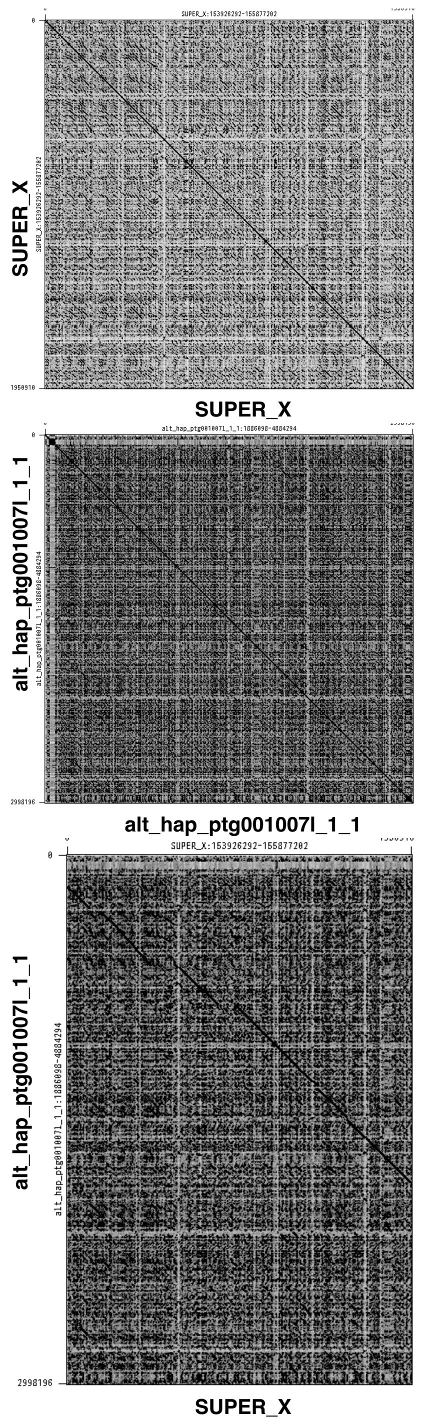


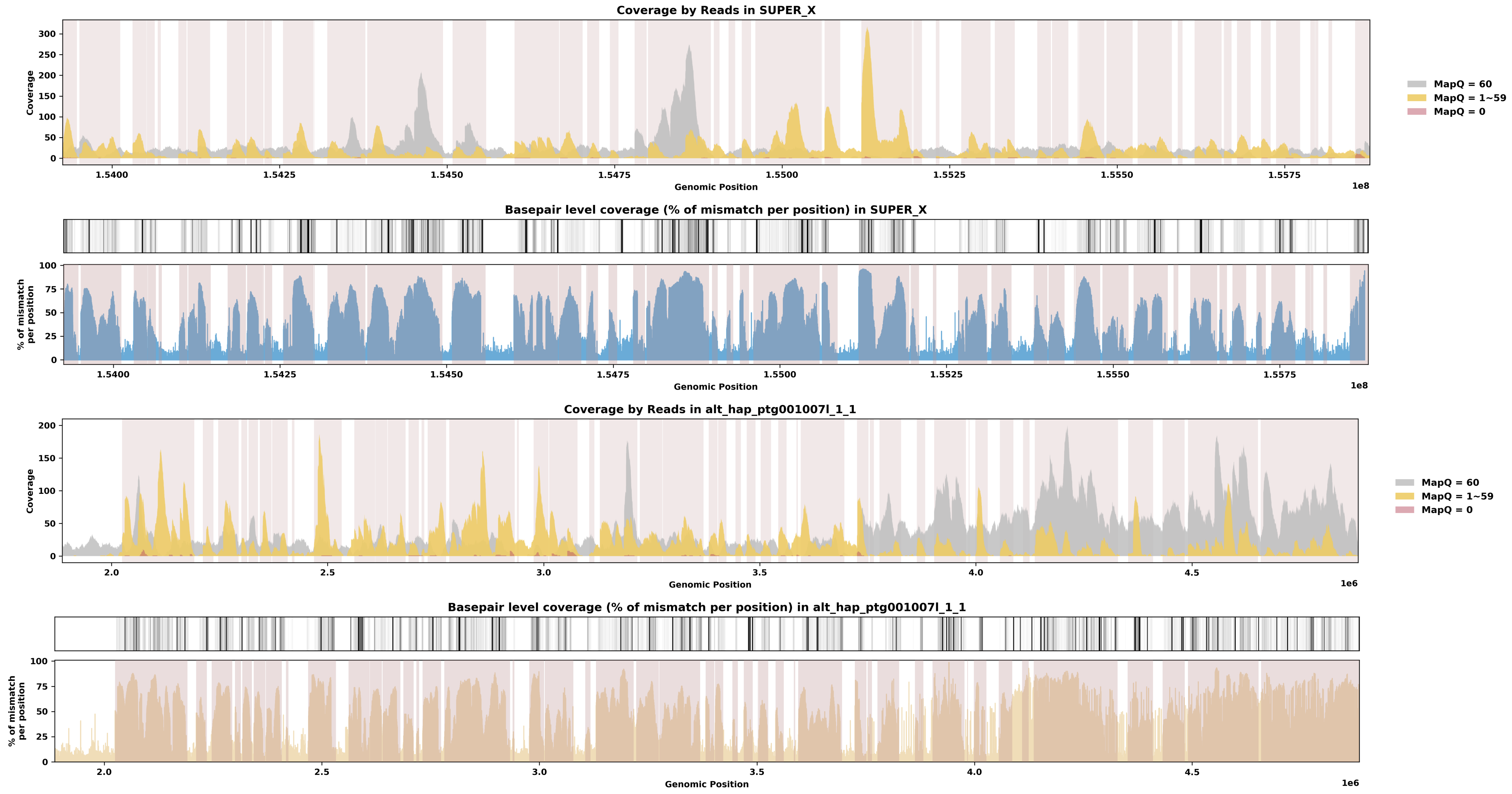

Species ID: mSteCoe1

Common Name: striped dolphin

Scientific Name: Stenella coeruleoalba

Assembly Type: Not Haplotype Resolved

Data Source: VGP

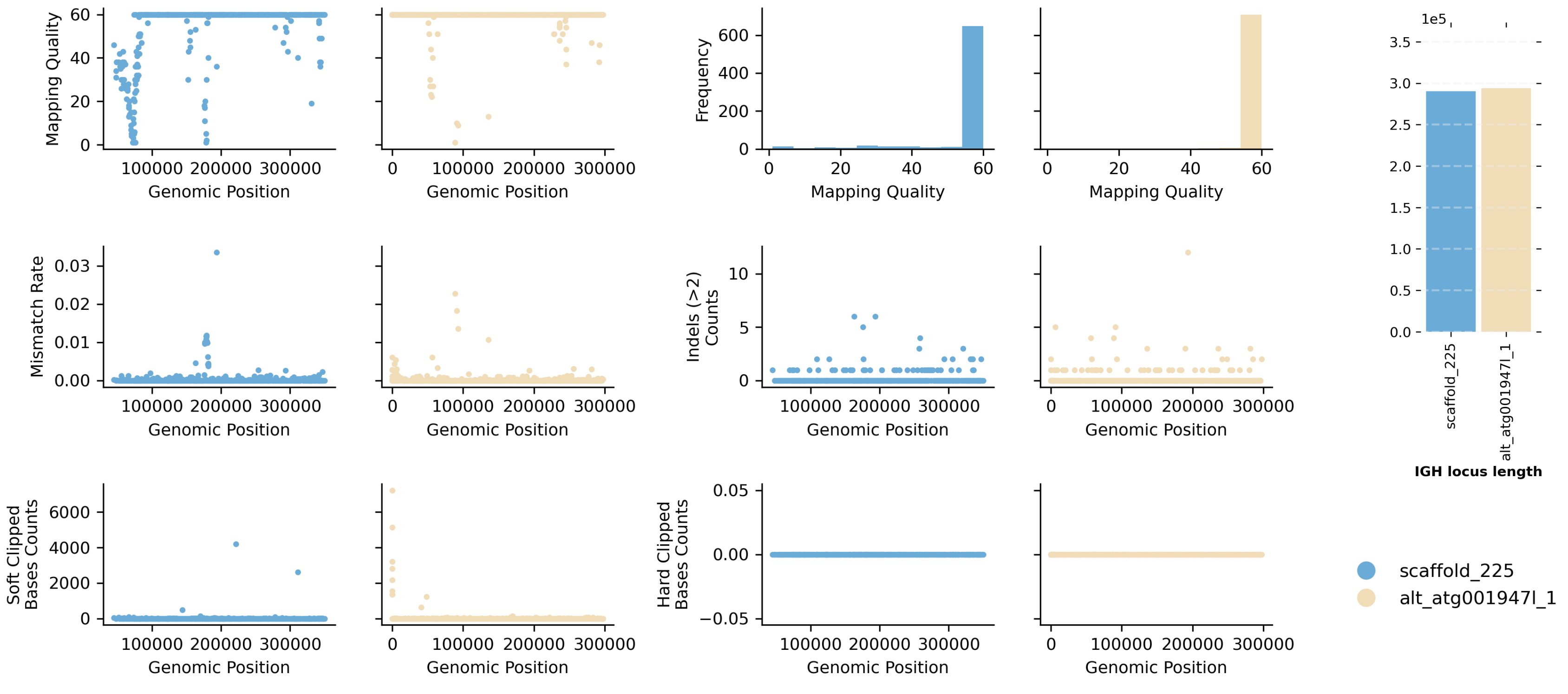


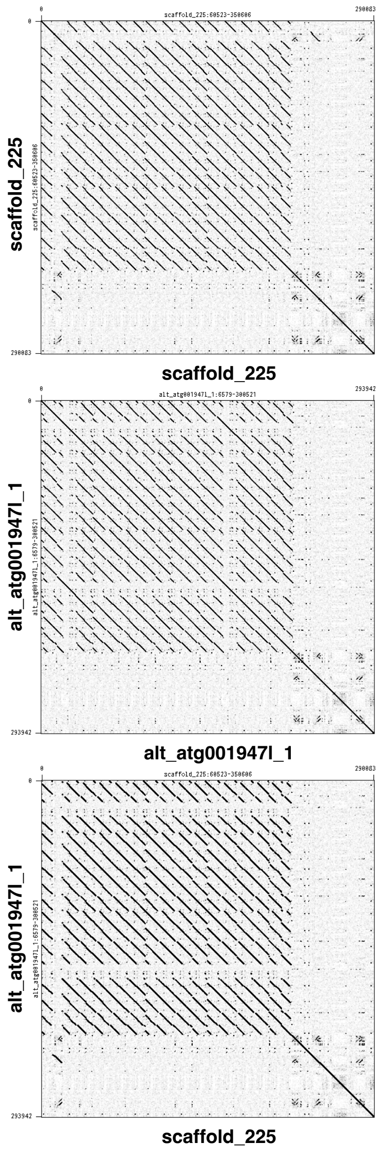


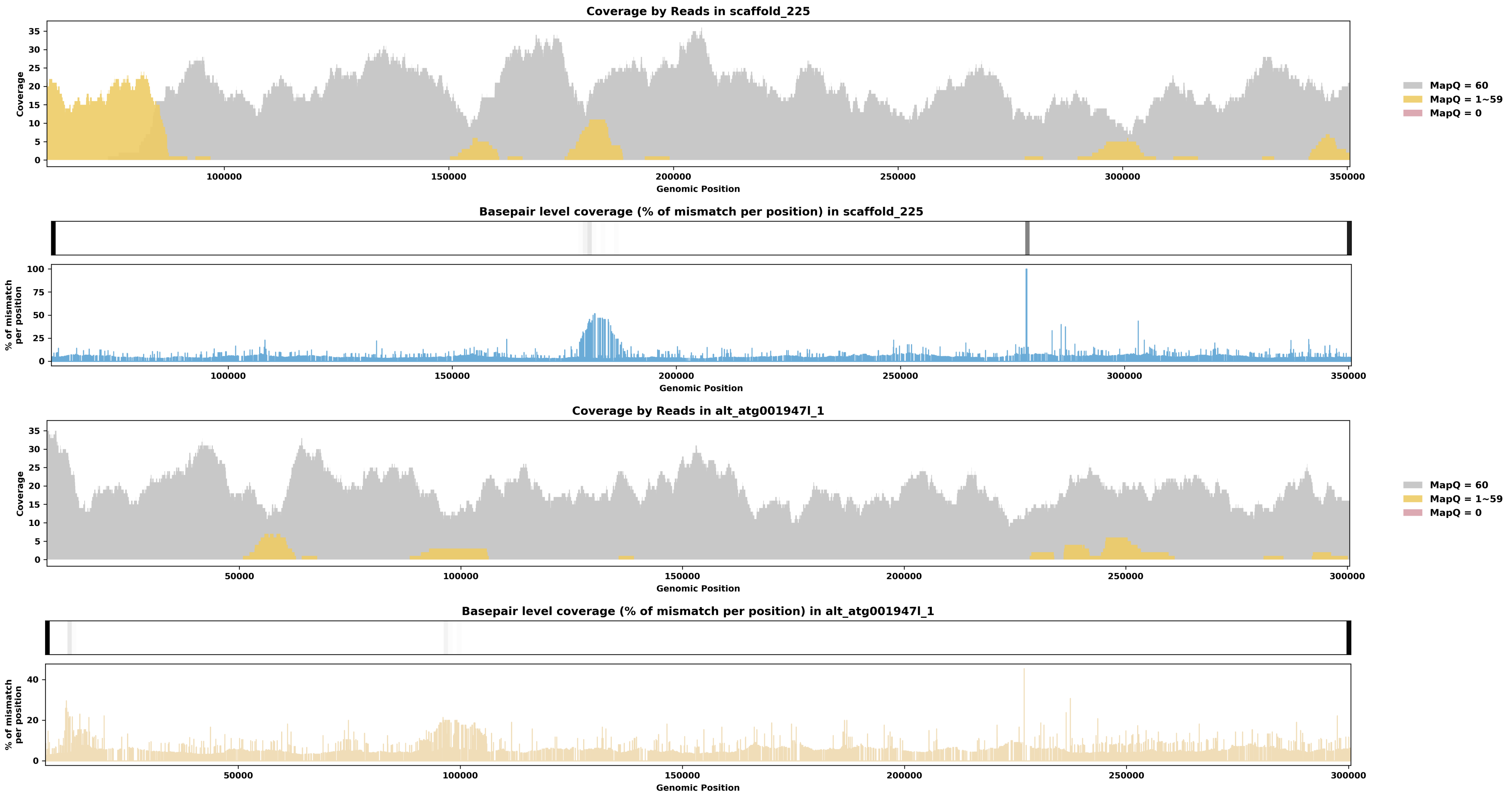

Species ID: mTalEur1

Common Name: European mole

Scientific Name: Talpa europaea

Assembly Type: Not Haplotype Resolved

Data Source: VGP

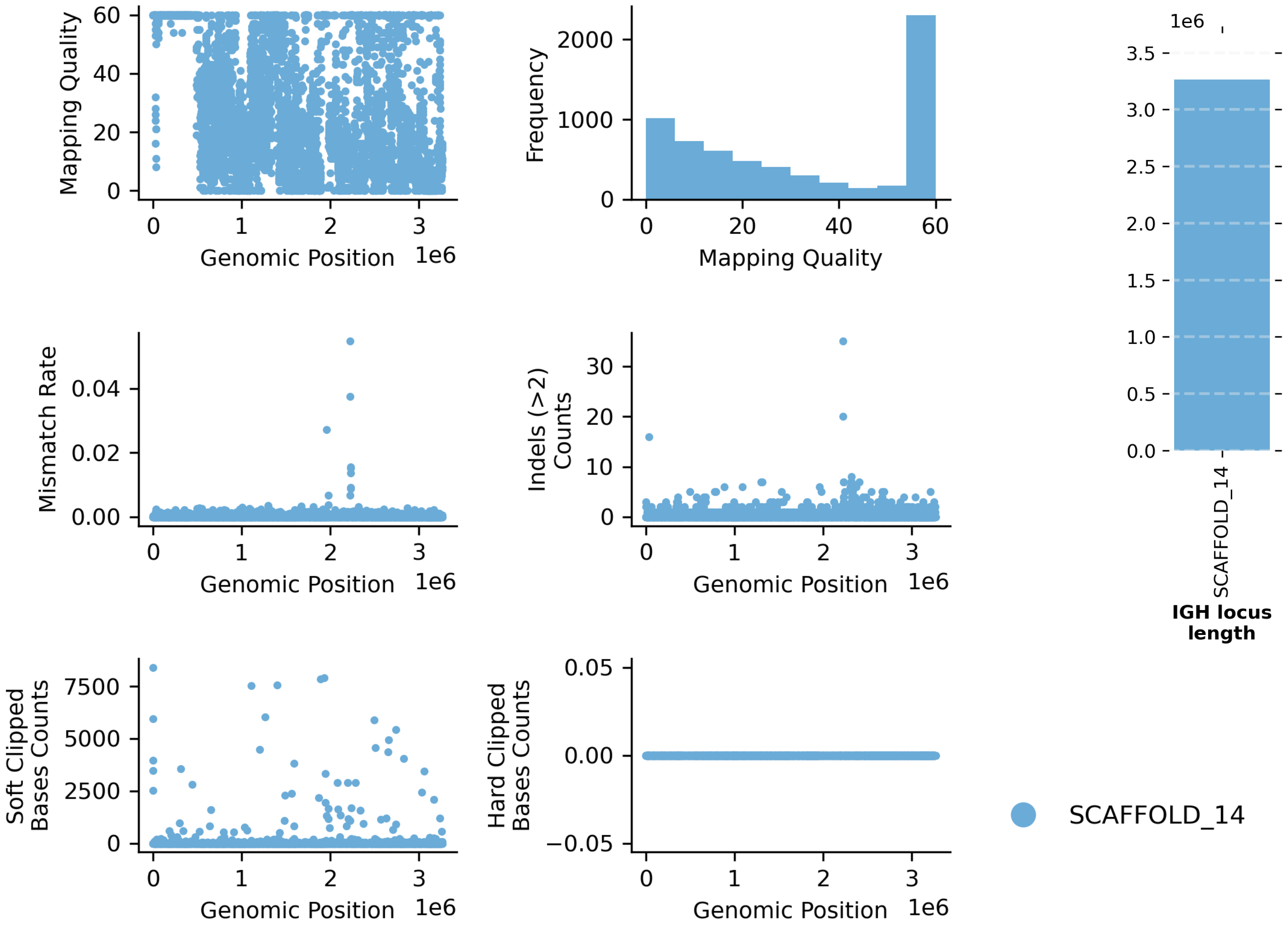


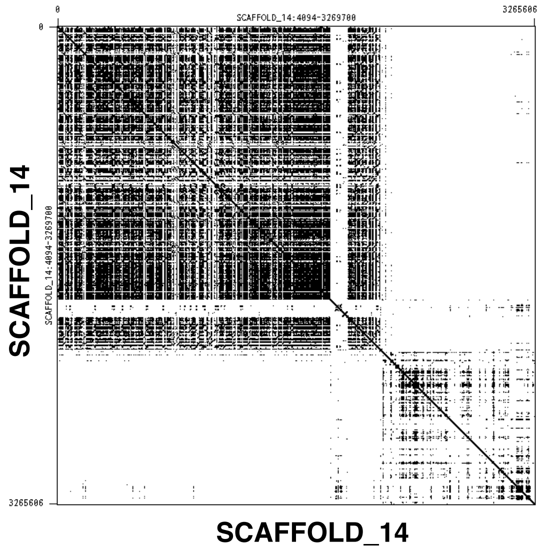


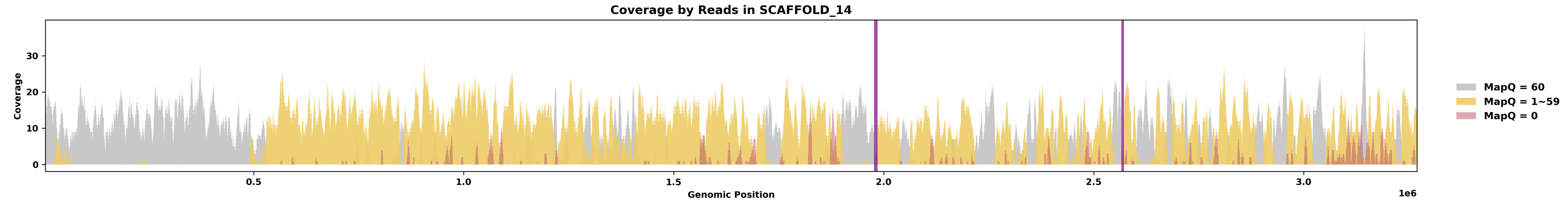


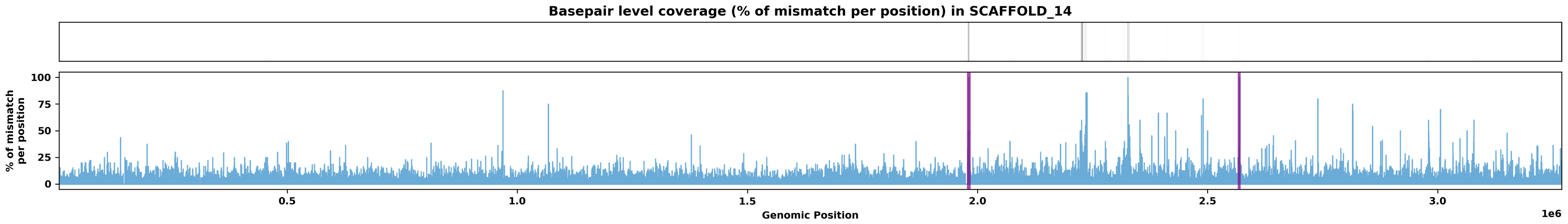

Species ID: mThoBot1

Common Name: Botta's pocket gopher

Scientific Name: Thomomys bottae

Assembly Type: Not Haplotype Resolved

Data Source: CCGP

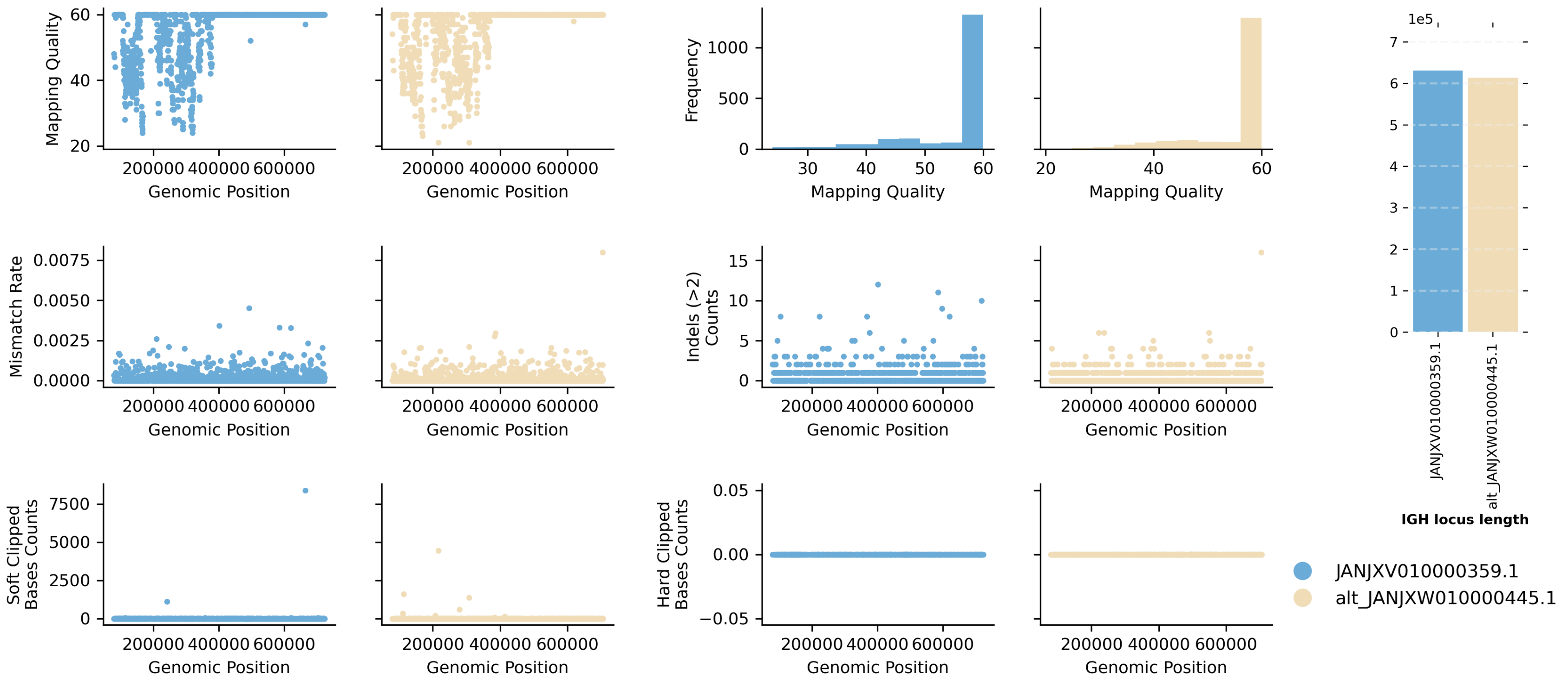


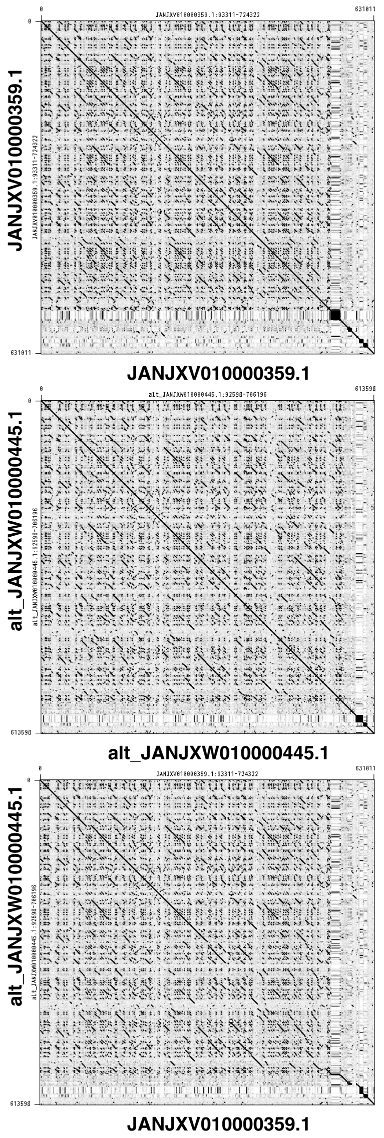


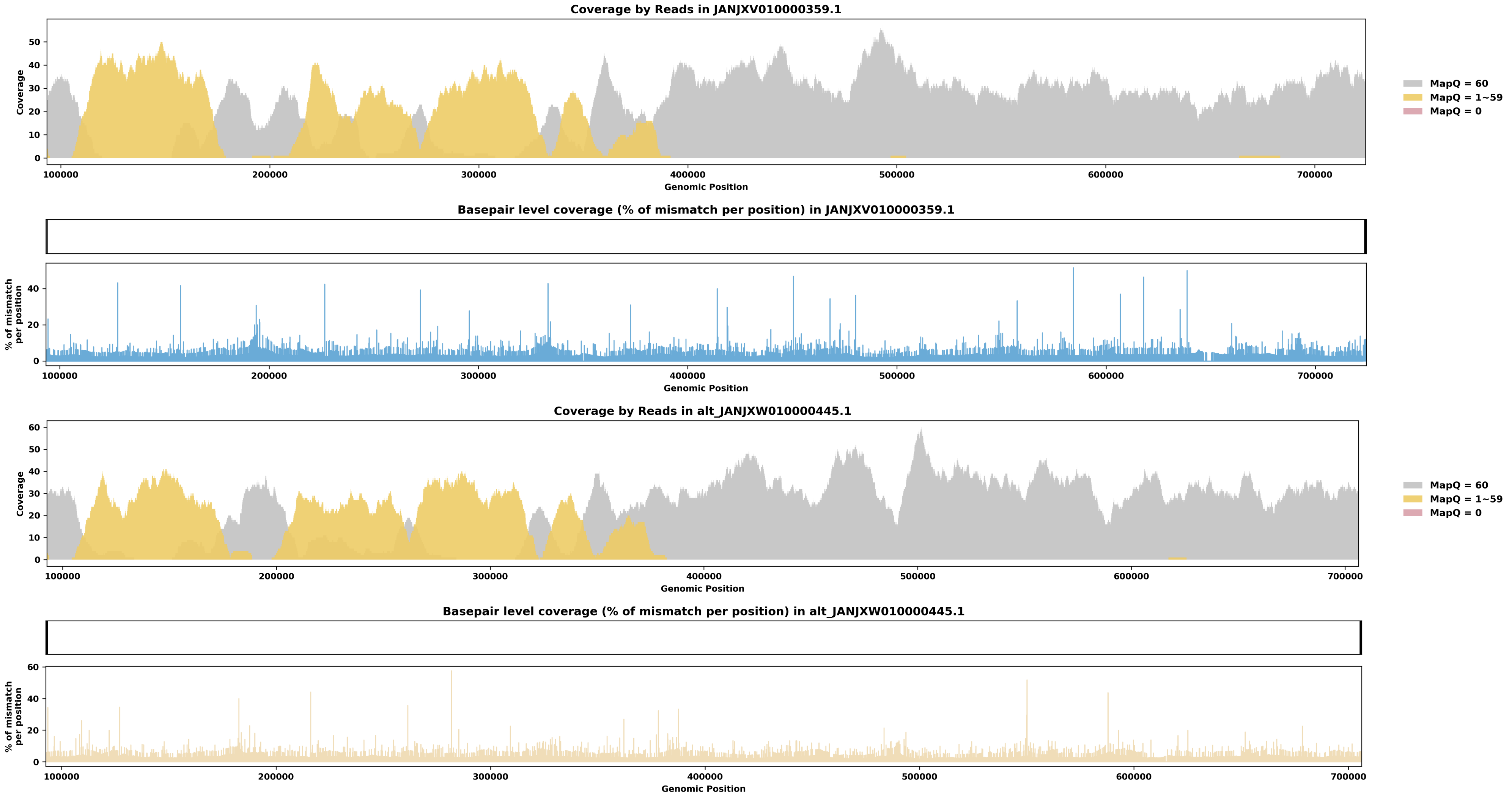

Species ID: mUrsAme1

Common Name: American black bear

Scientific Name: Ursus americanus

Assembly Type: Not Haplotype Resolved

Data Source: CCGP

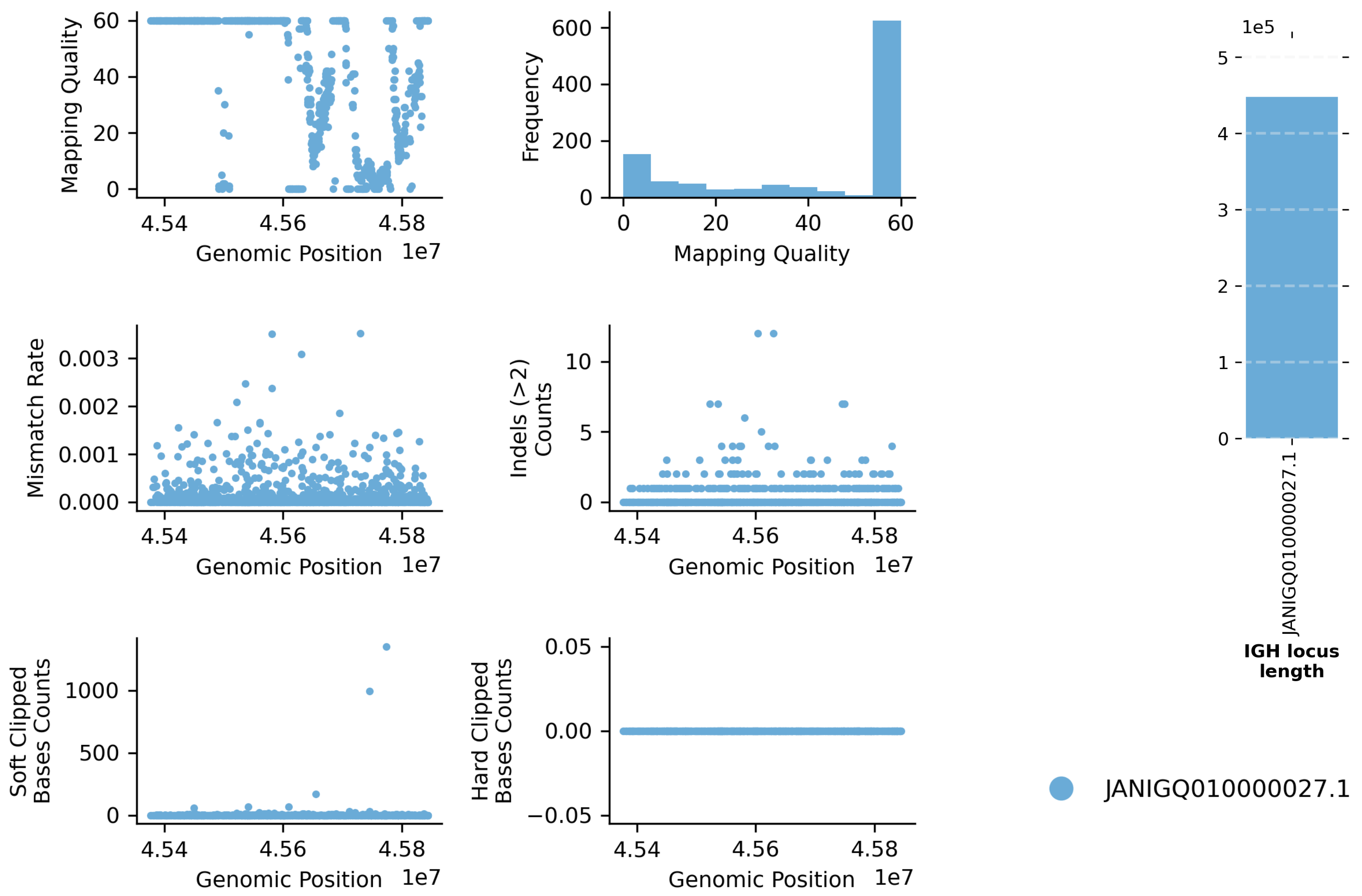


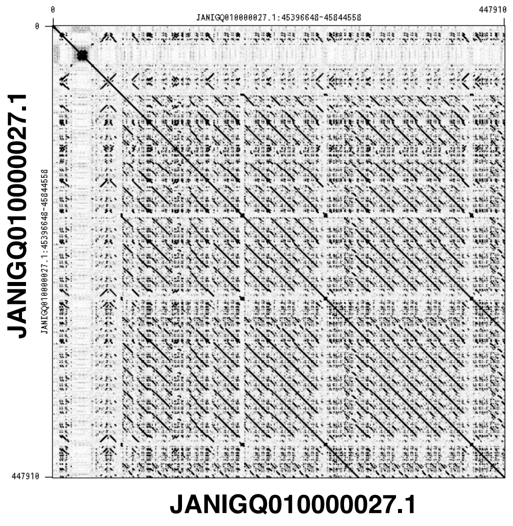


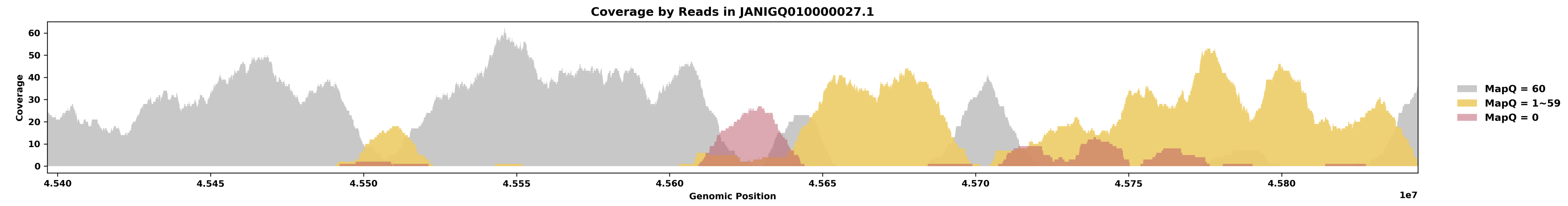


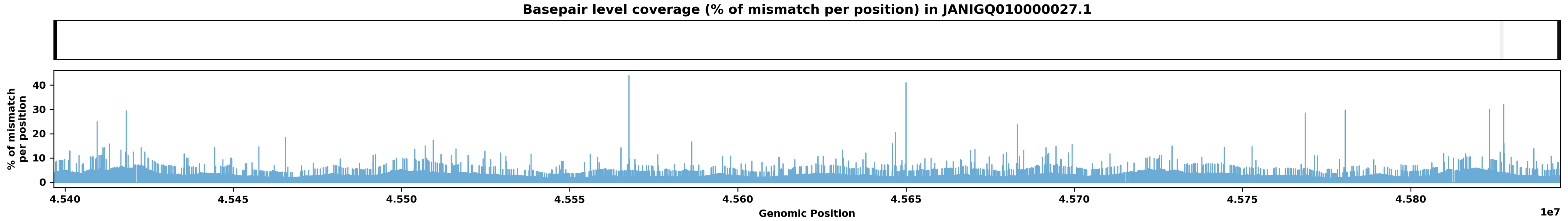

Species ID: mUrsArc2

Common Name: brown bear

Scientific Name: Ursus arctos

Assembly Type: Not Haplotype Resolved

Data Source: NCBI

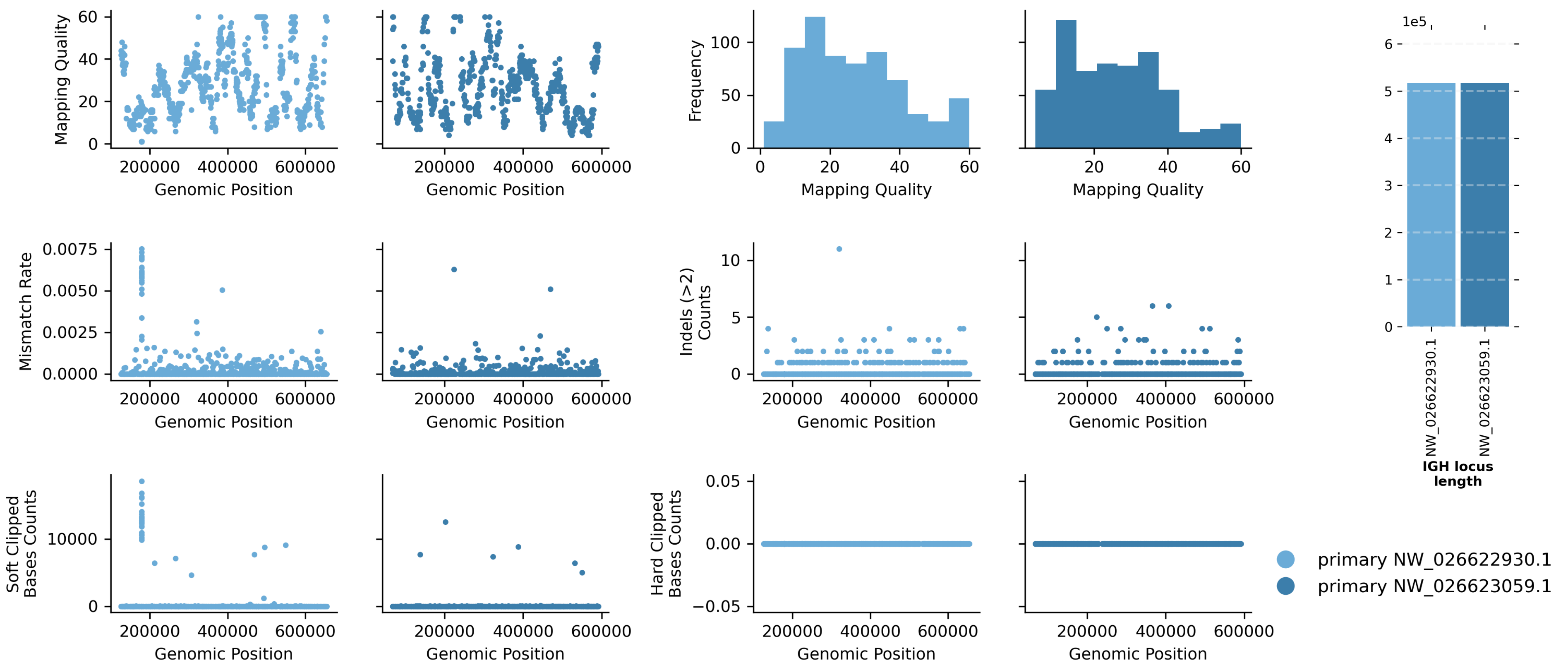


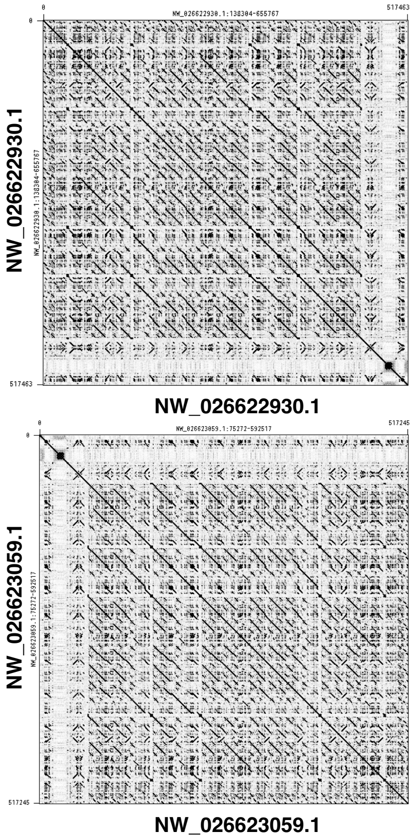


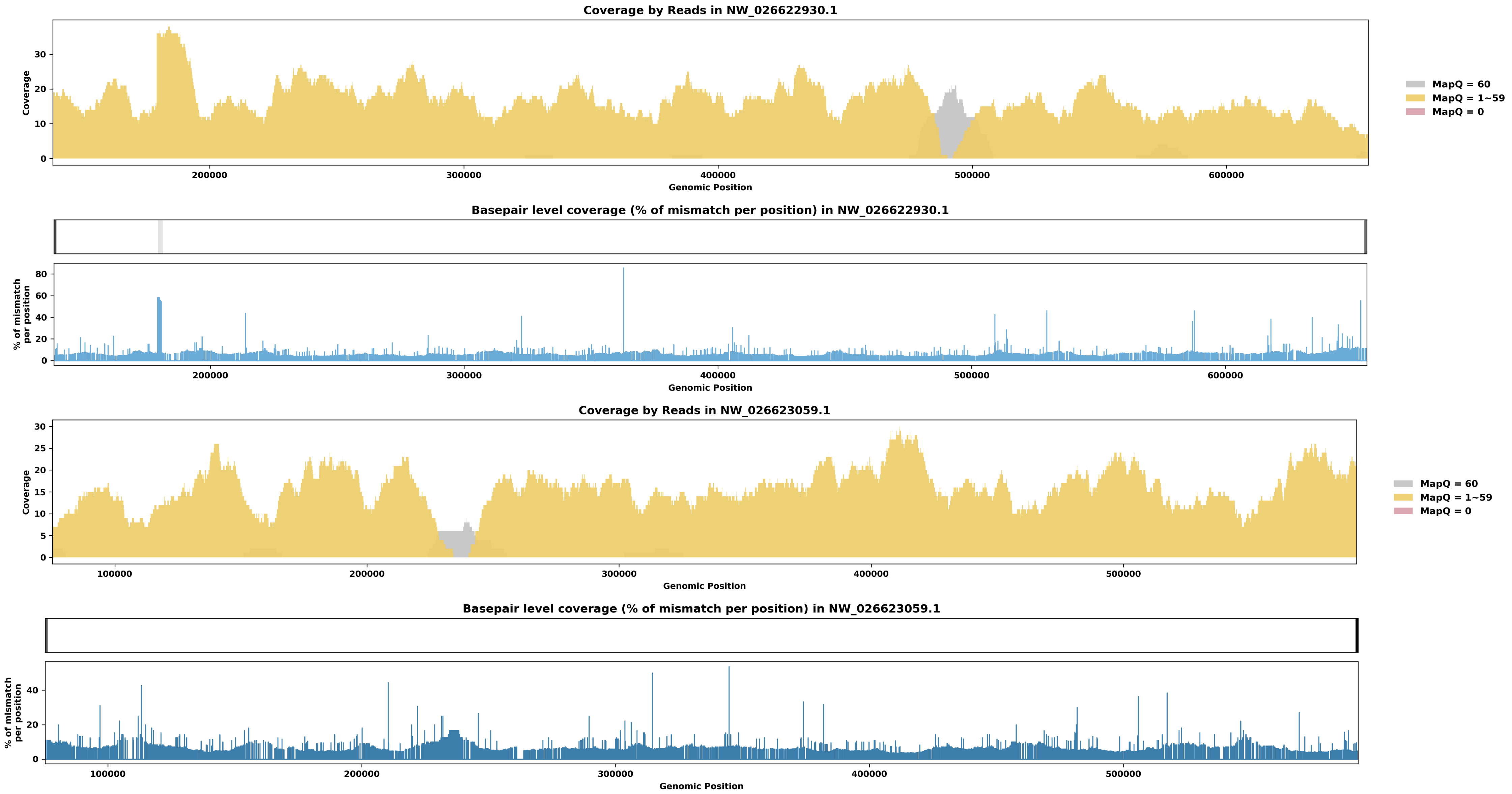

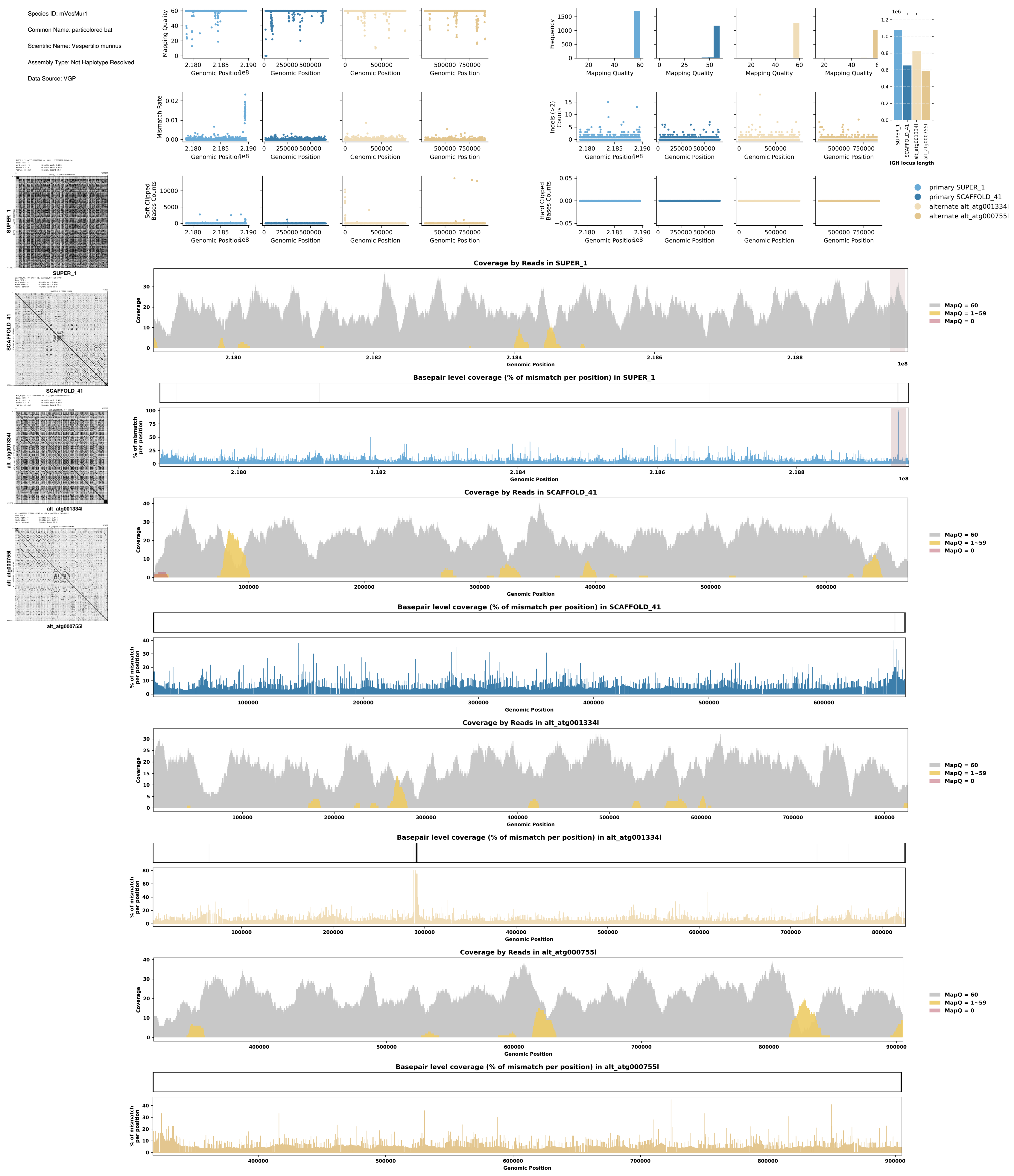
Species ID: mVesMur1
Common Name: particolored bat
Scientific Name: Vespertilio murinus
Assembly Type: Not Haplotype Resolved
Data Source: VGP
Mapping Quality
Genomic Position
Frequency
Mismatch Rate
Indels (>2) Counts
Soft Clipped Bases Counts
Hard Clipped Bases Counts
IGH locus length
primary SUPER_1
primary SCAFFOLD_41
alternate alt_atg001334l
alternate alt_atg000755l
SUPER_1
SCAFFOLD_41
alt_atg001334l
alt_atg000755l
Coverage by Reads in SUPER_1
Coverage
MapQ = 60
MapQ = 1~59
MapQ = 0
Basepair level coverage (% of mismatch per position) in SUPER_1
% of mismatch per position
Coverage by Reads in SCAFFOLD_41
Basepair level coverage (% of mismatch per position) in SCAFFOLD_41
Coverage by Reads in alt_atg001334l
Basepair level coverage (% of mismatch per position) in alt_atg001334l
Coverage by Reads in alt_atg000755l
Basepair level coverage (% of mismatch per position) in alt_atg000755l

Species ID: rAllMis2

Common Name: American alligator

Scientific Name: Alligator mississippiensis

Assembly Type: Haplotype Resolved

Data Source: VGP

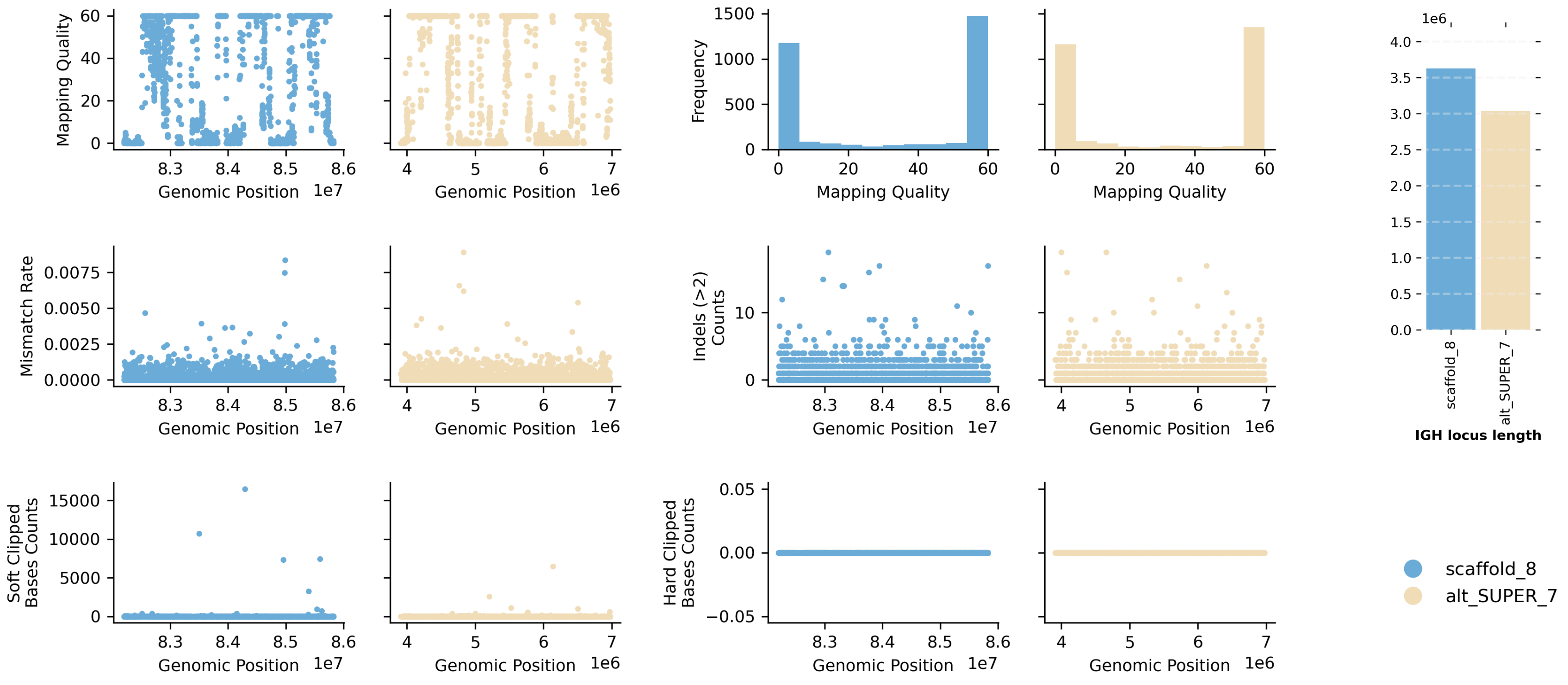


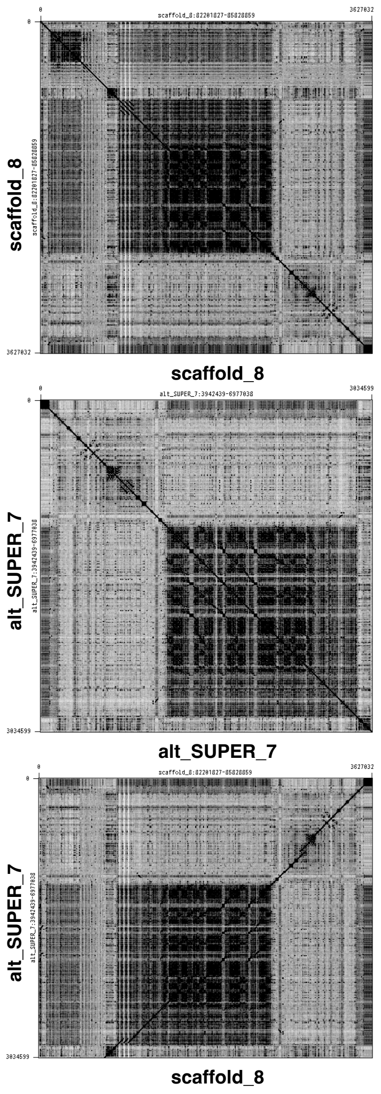


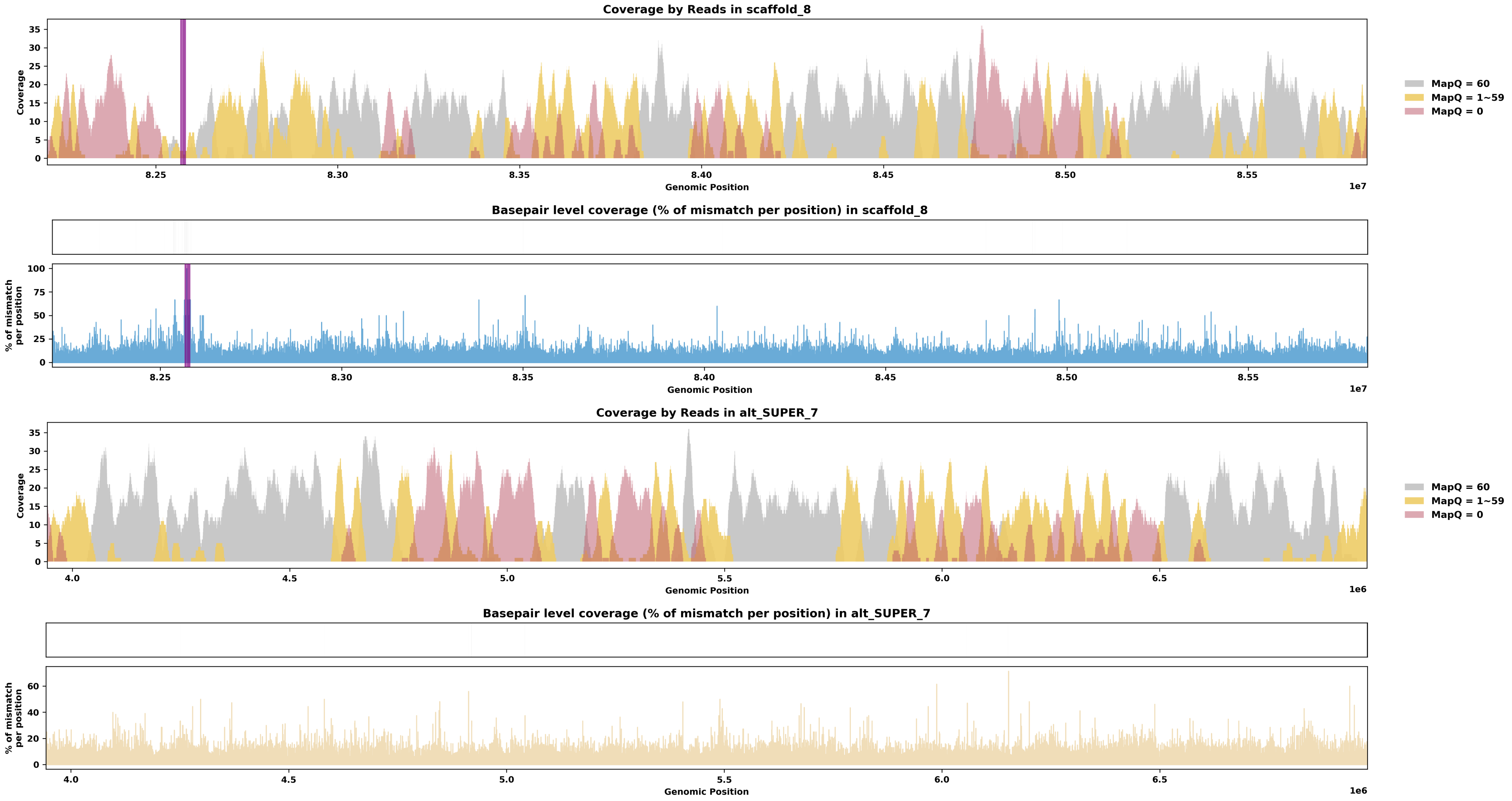

Species ID: rCarCar2

Common Name: Loggerhead turtle

Scientific Name: Caretta caretta

Assembly Type: Haplotype Resolved

Data Source: VGP

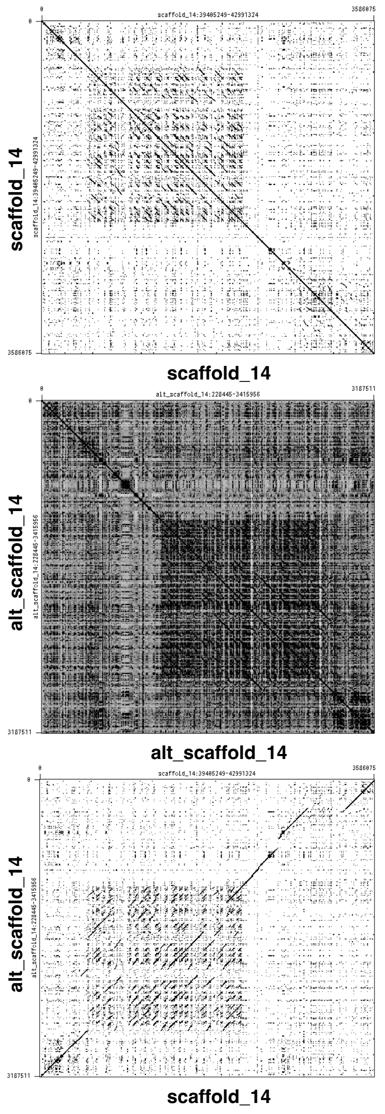


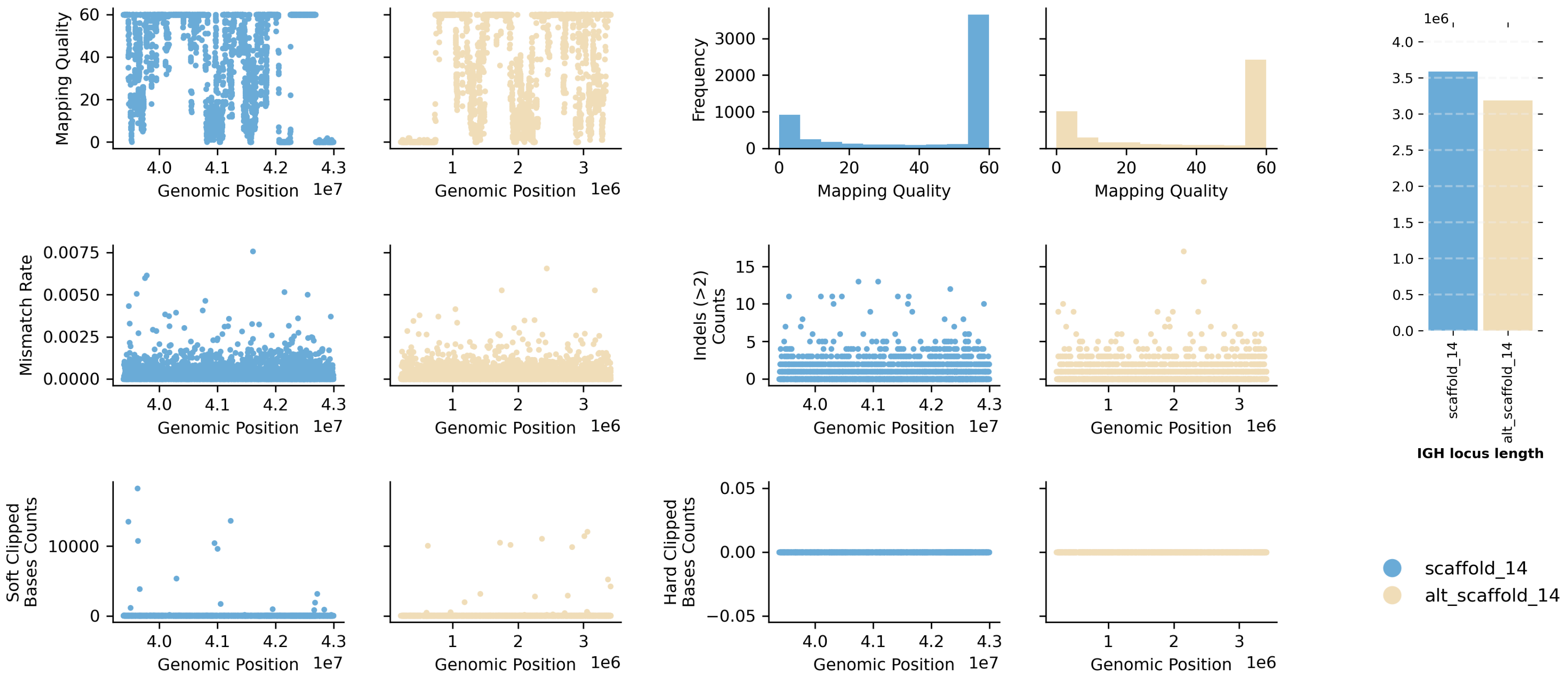


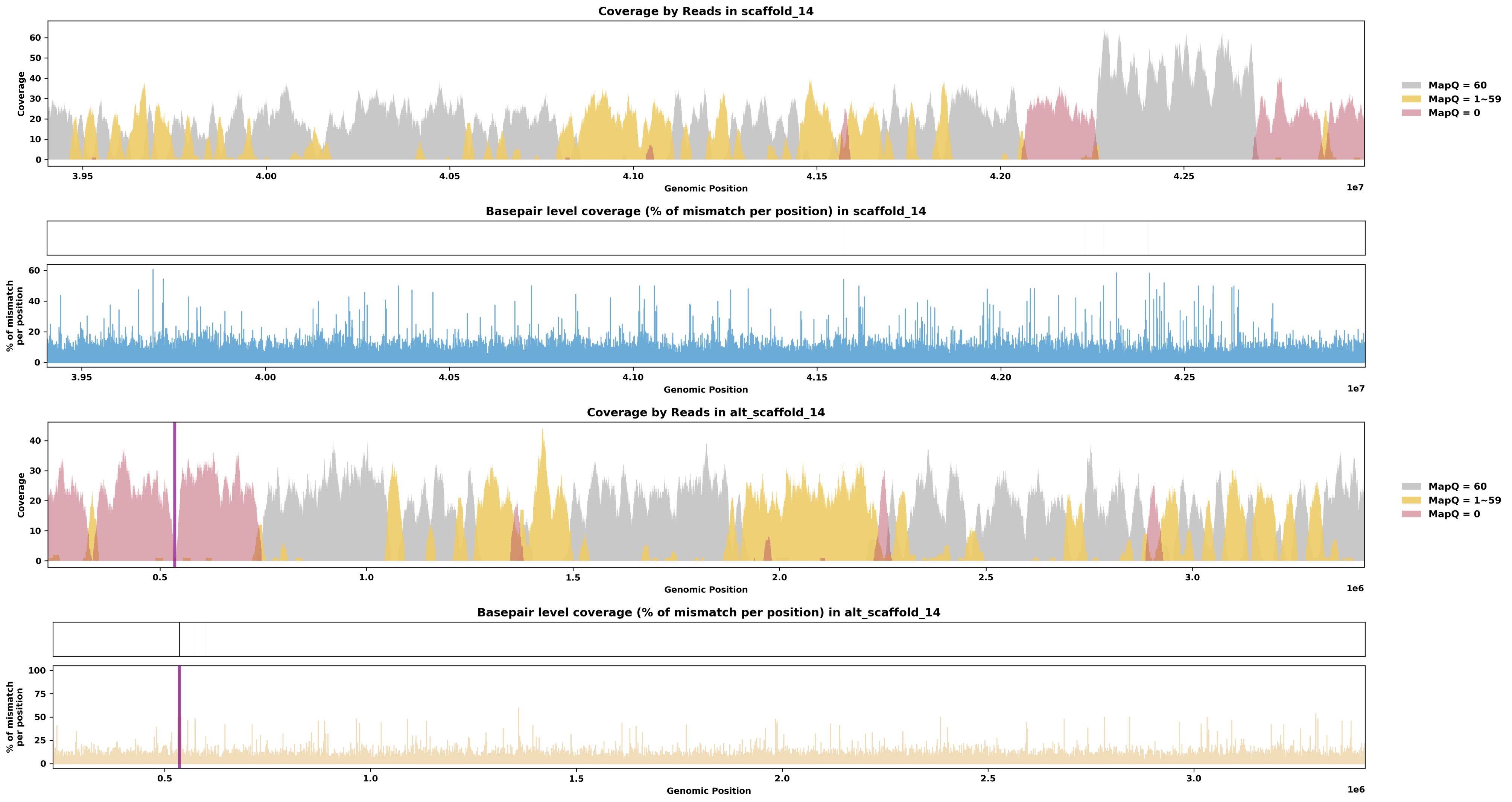

Species ID: rEmyOrb1

Common Name: European pond turtle

Scientific Name: Emys orbicularis

Assembly Type: Haplotype Resolved

Data Source: VGP

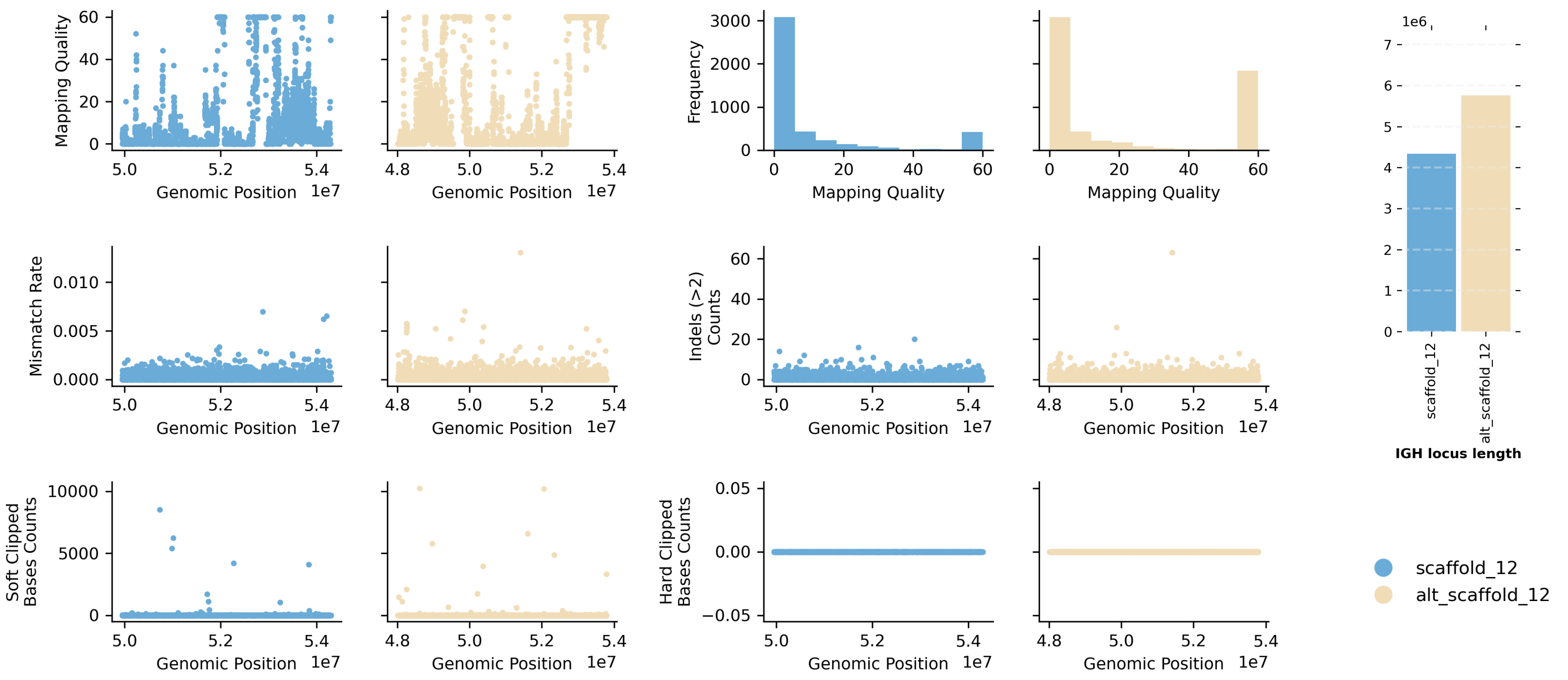


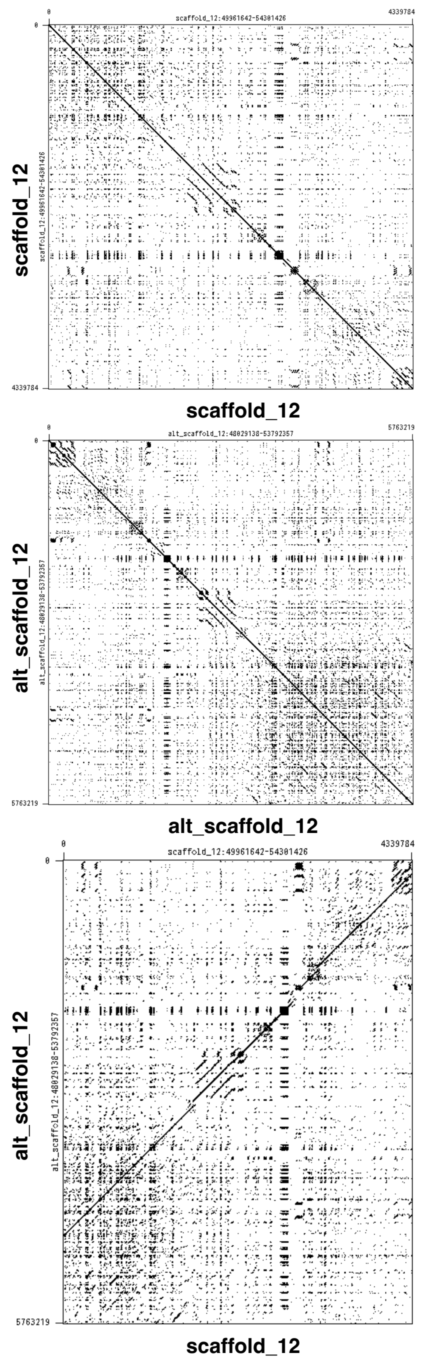


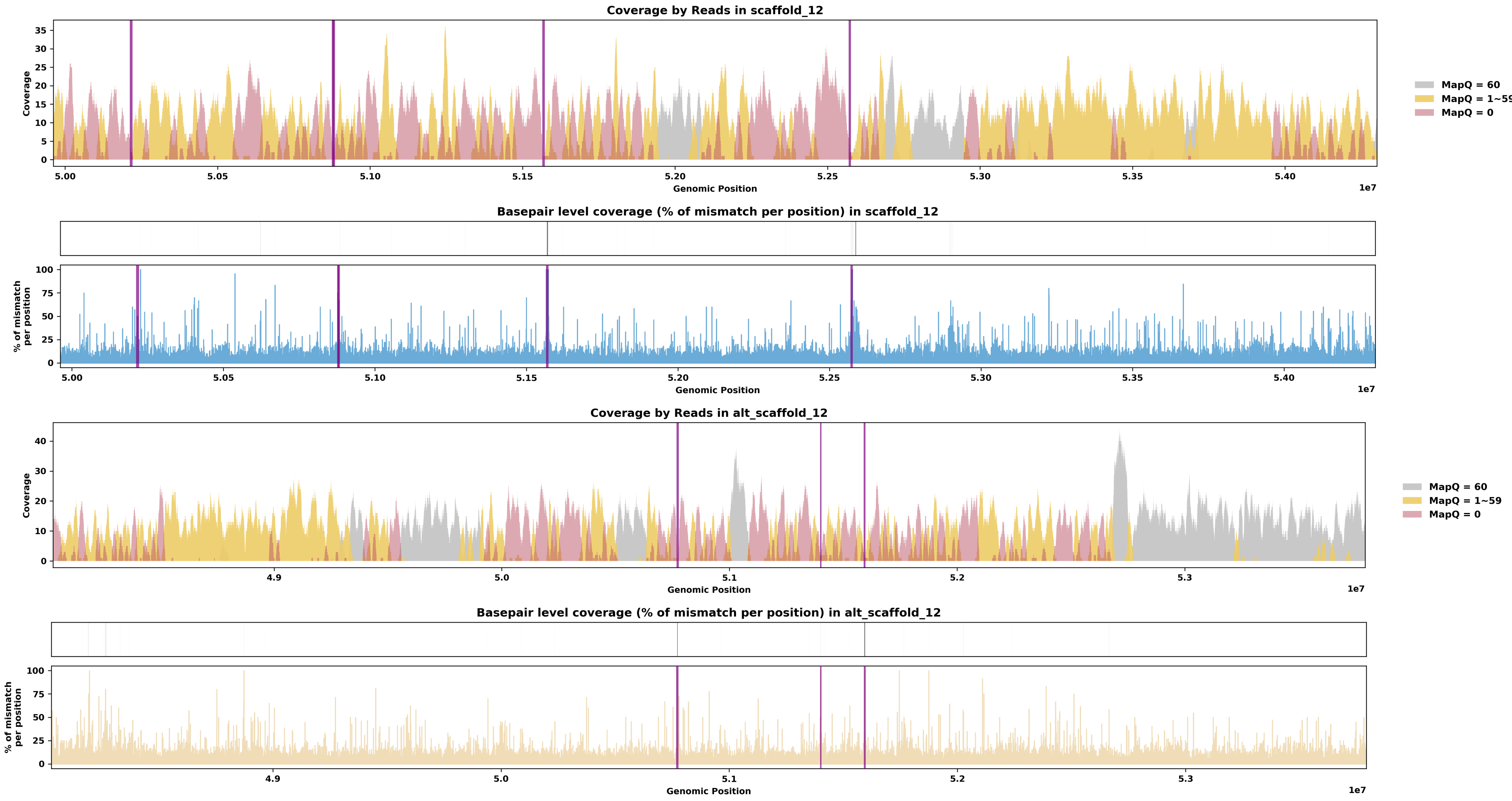

Species ID: rEryReg1

Common Name: royal ground snake

Scientific Name: Erythrolamprus reginae

Assembly Type: Not Haplotype Resolved

Data Source: VGP

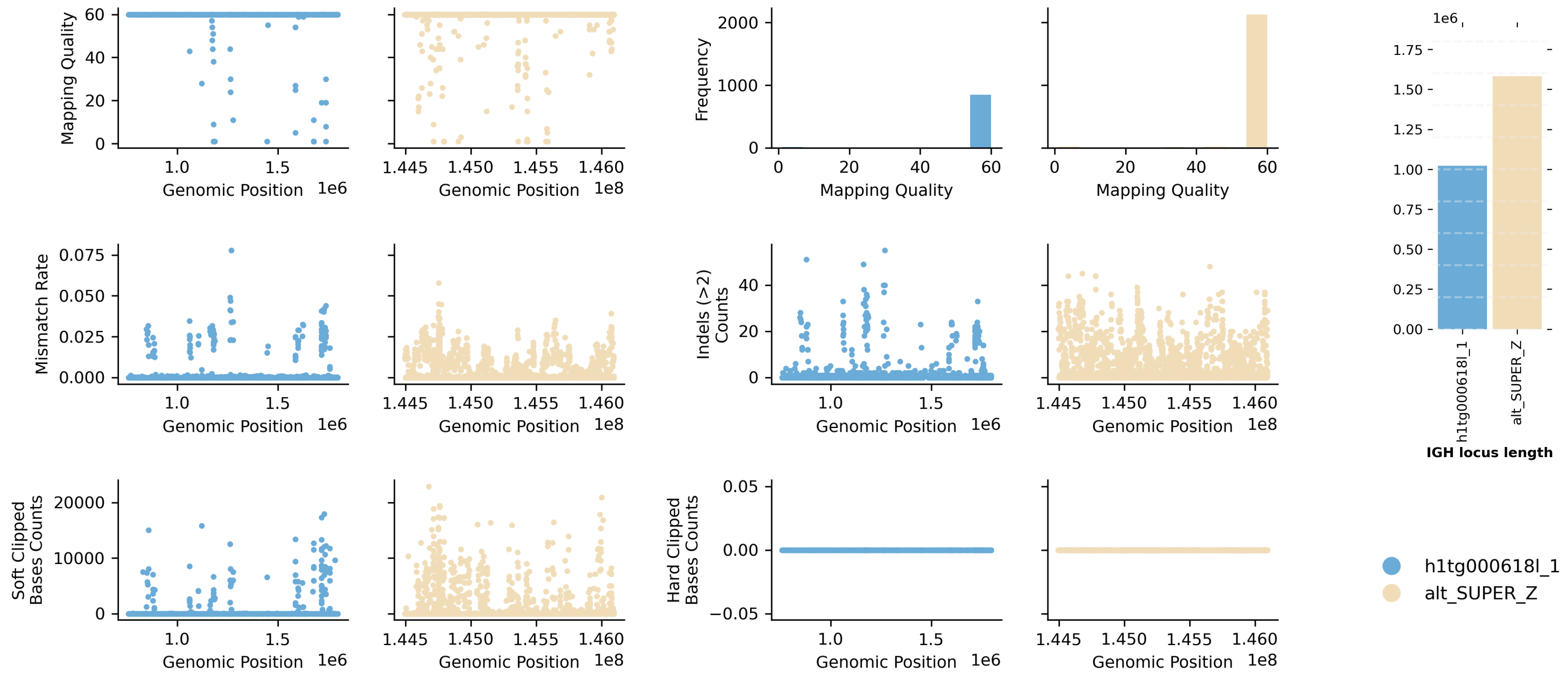


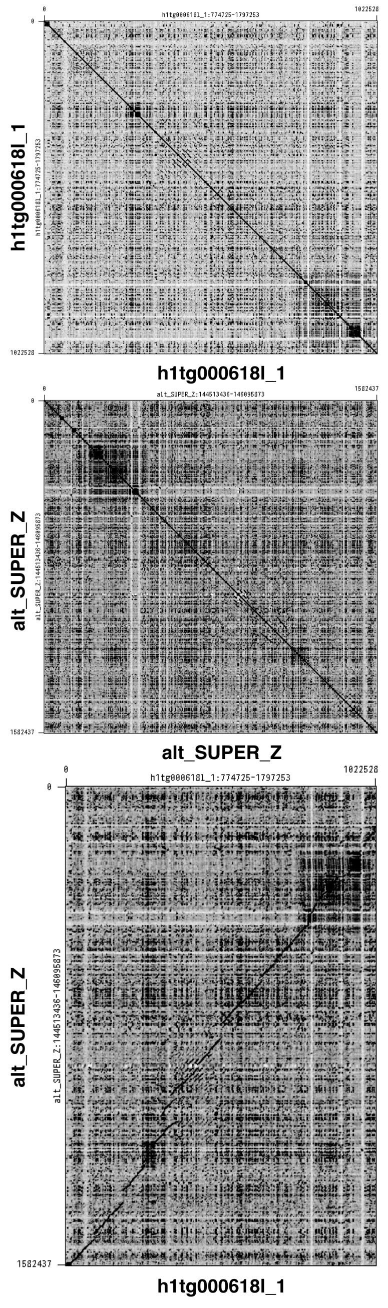


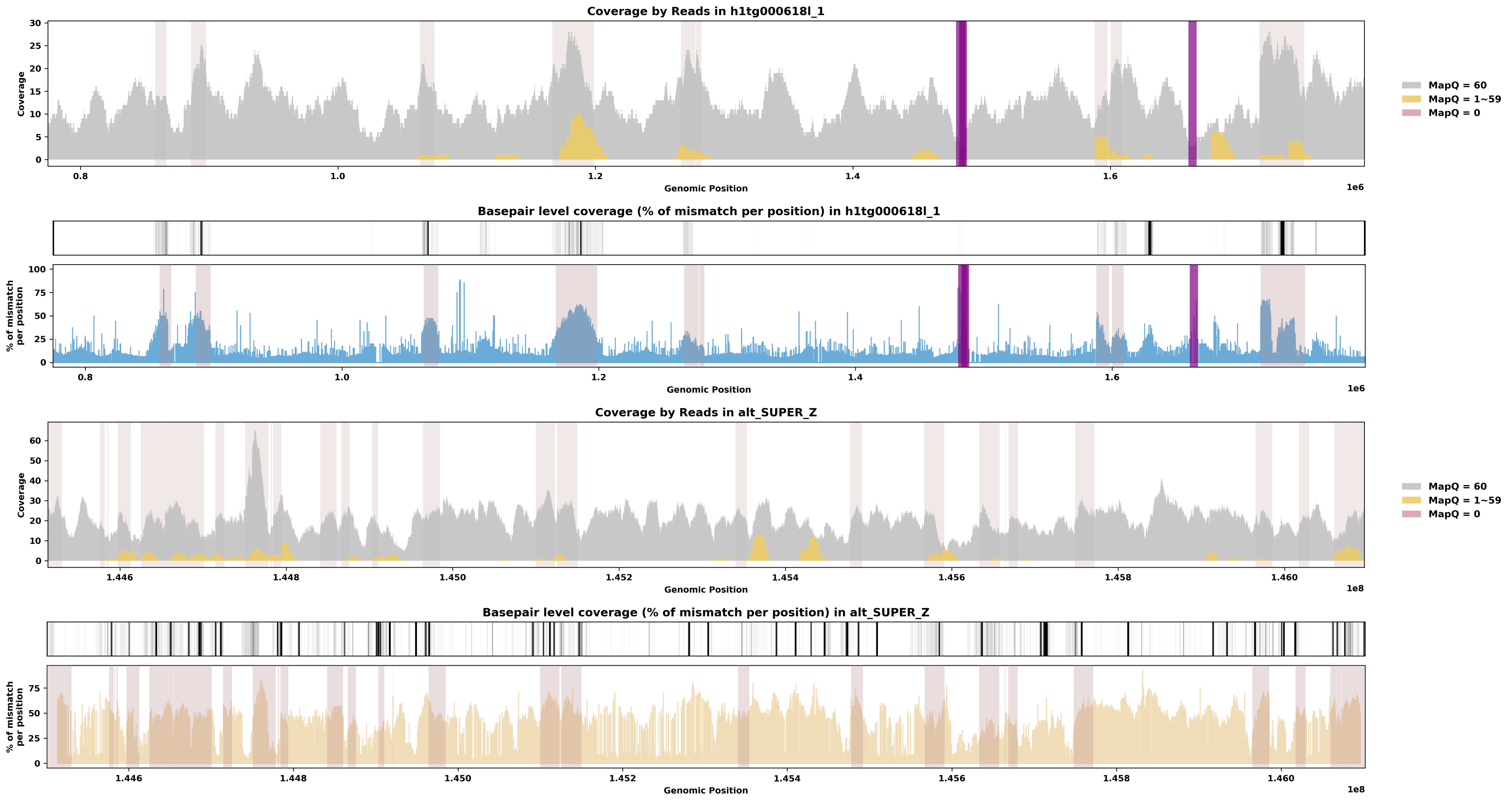

Species ID: rLiaOli1

Common Name: olive python

Scientific Name: Liasis olivaceus

Assembly Type: Haplotype Resolved

Data Source: VGP

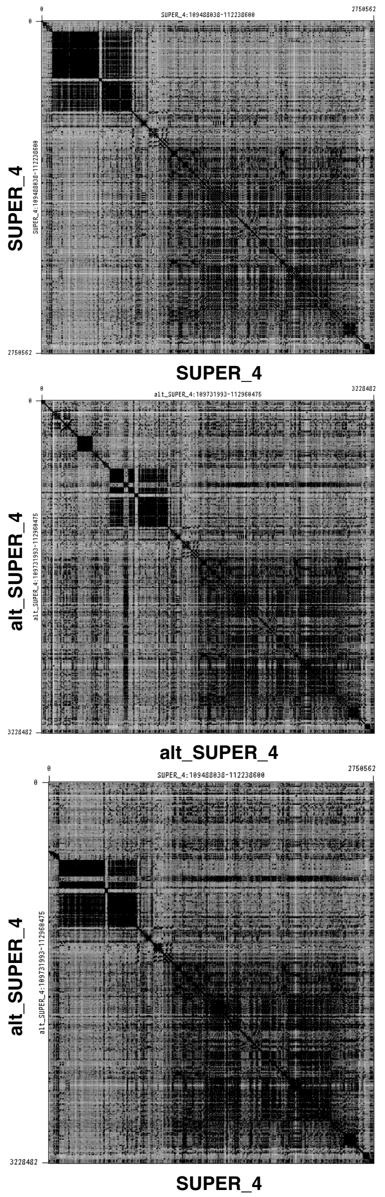


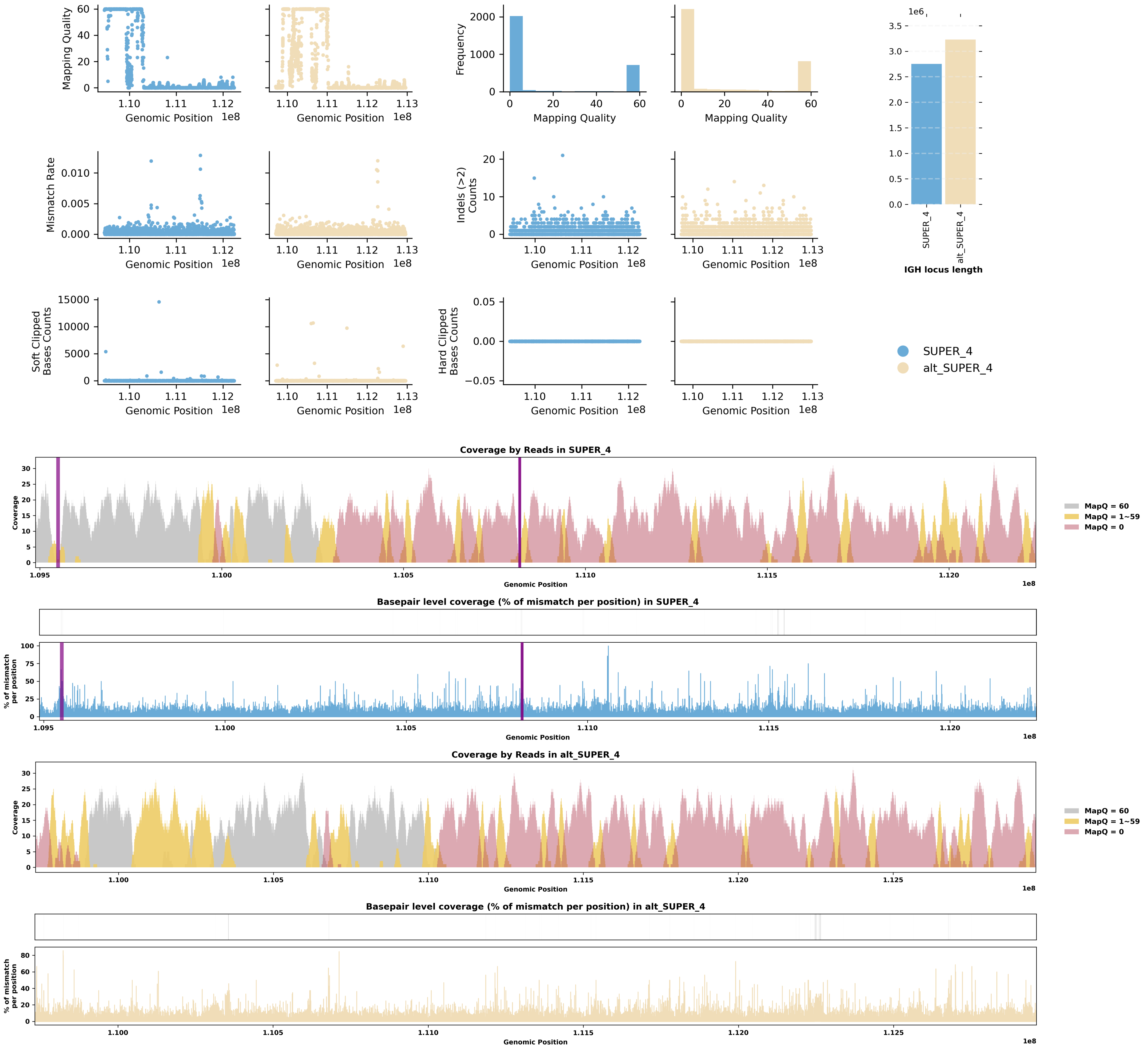

Species ID: rLiaOli2

Common Name: olive python

Scientific Name: Liasis olivaceus

Assembly Type: Haplotype Resolved

Data Source: VGP

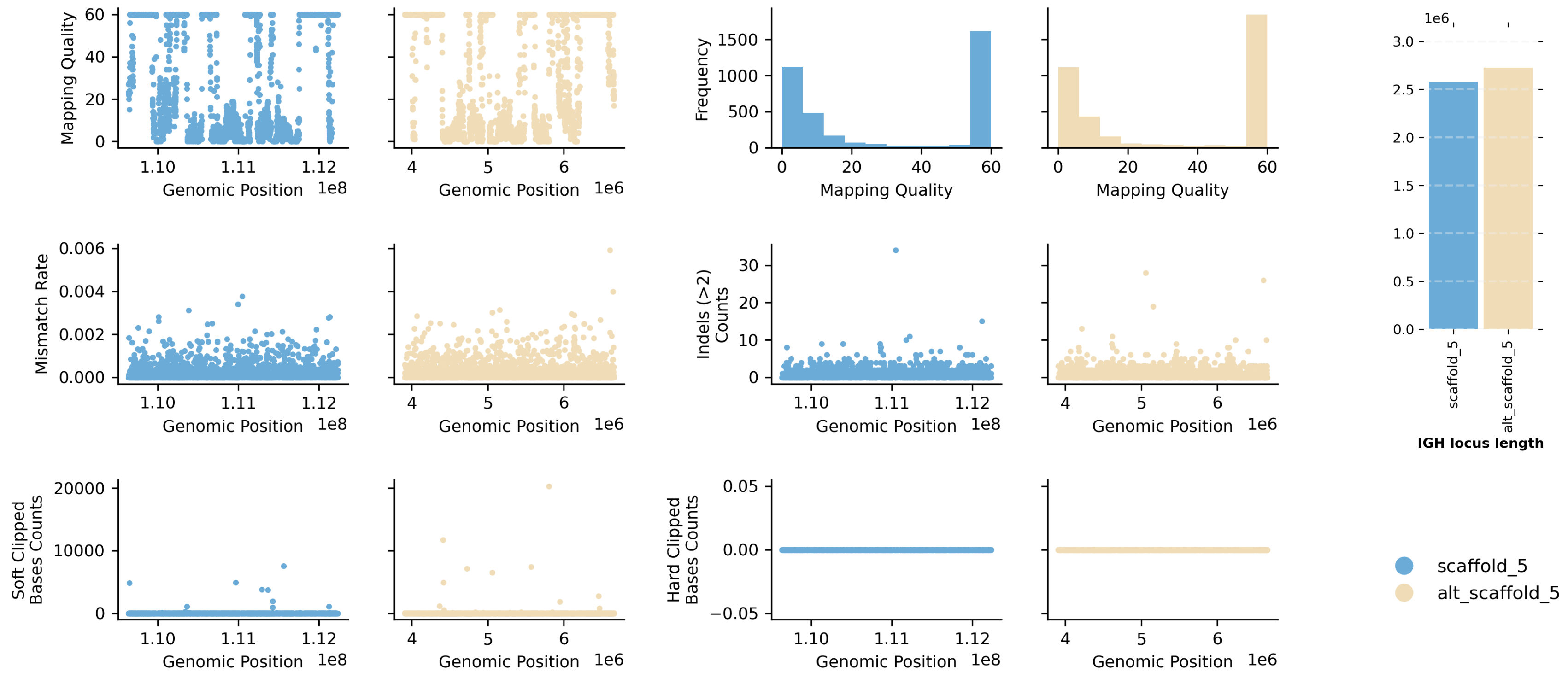


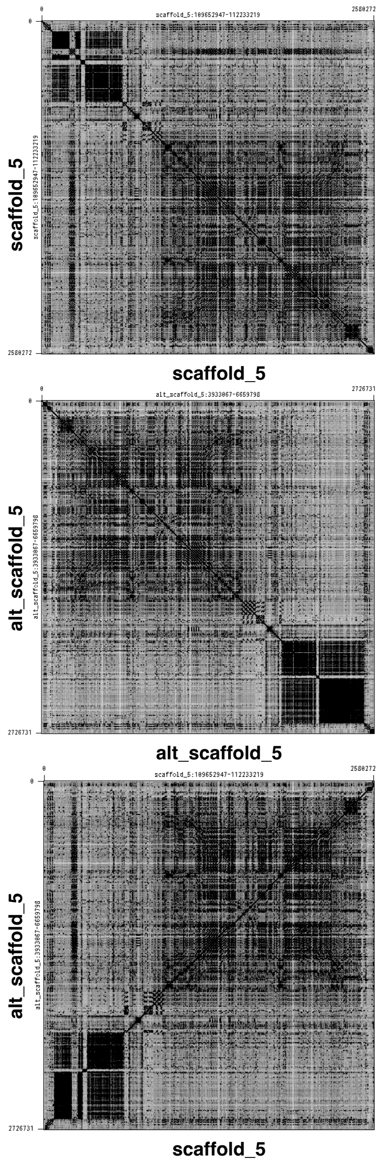


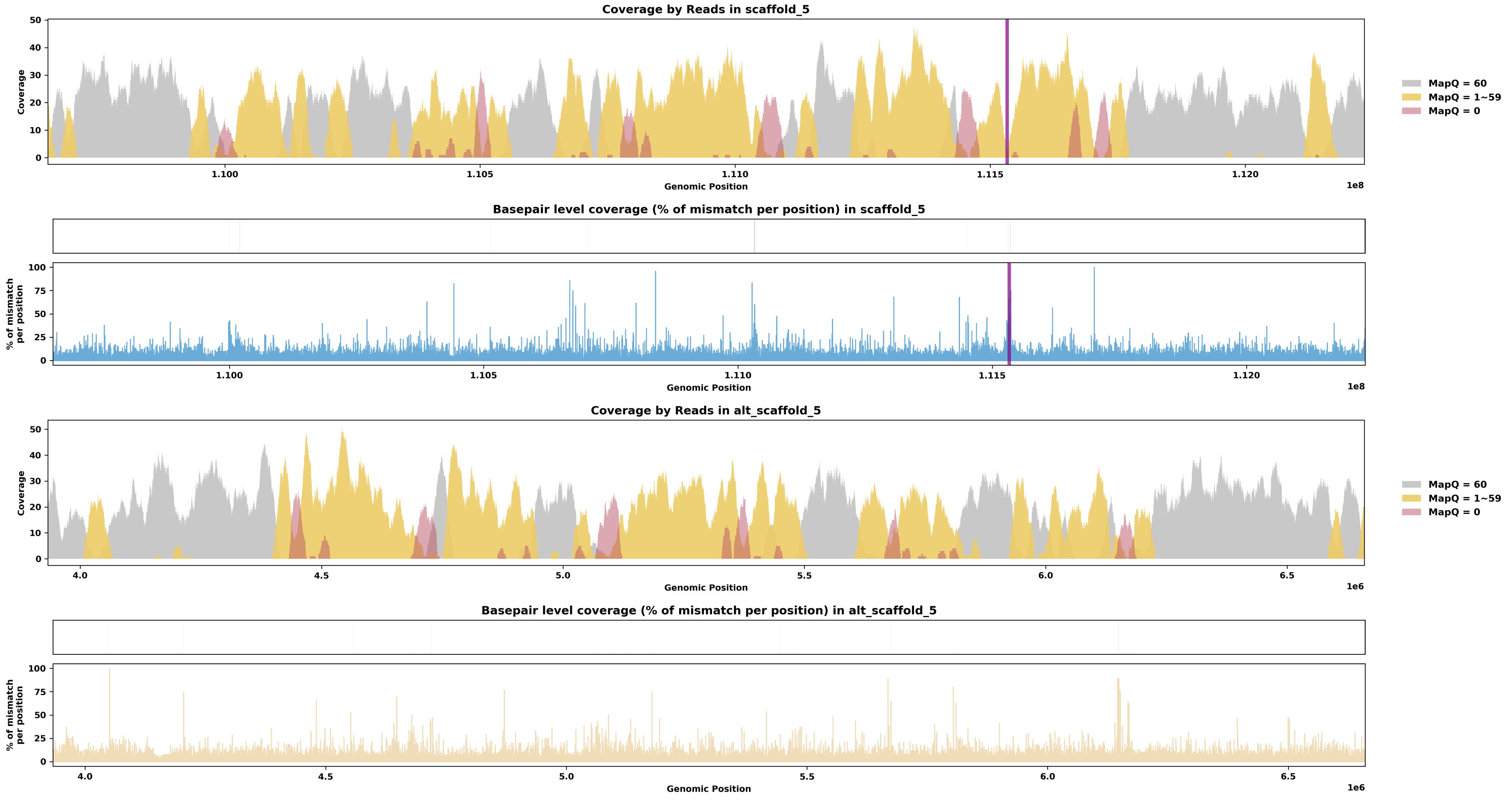

Species ID: rMalTer1

Common Name: diamondback terrapin

Scientific Name: Malaclemys terrapin

Assembly Type: Haplotype Resolved

Data Source: VGP

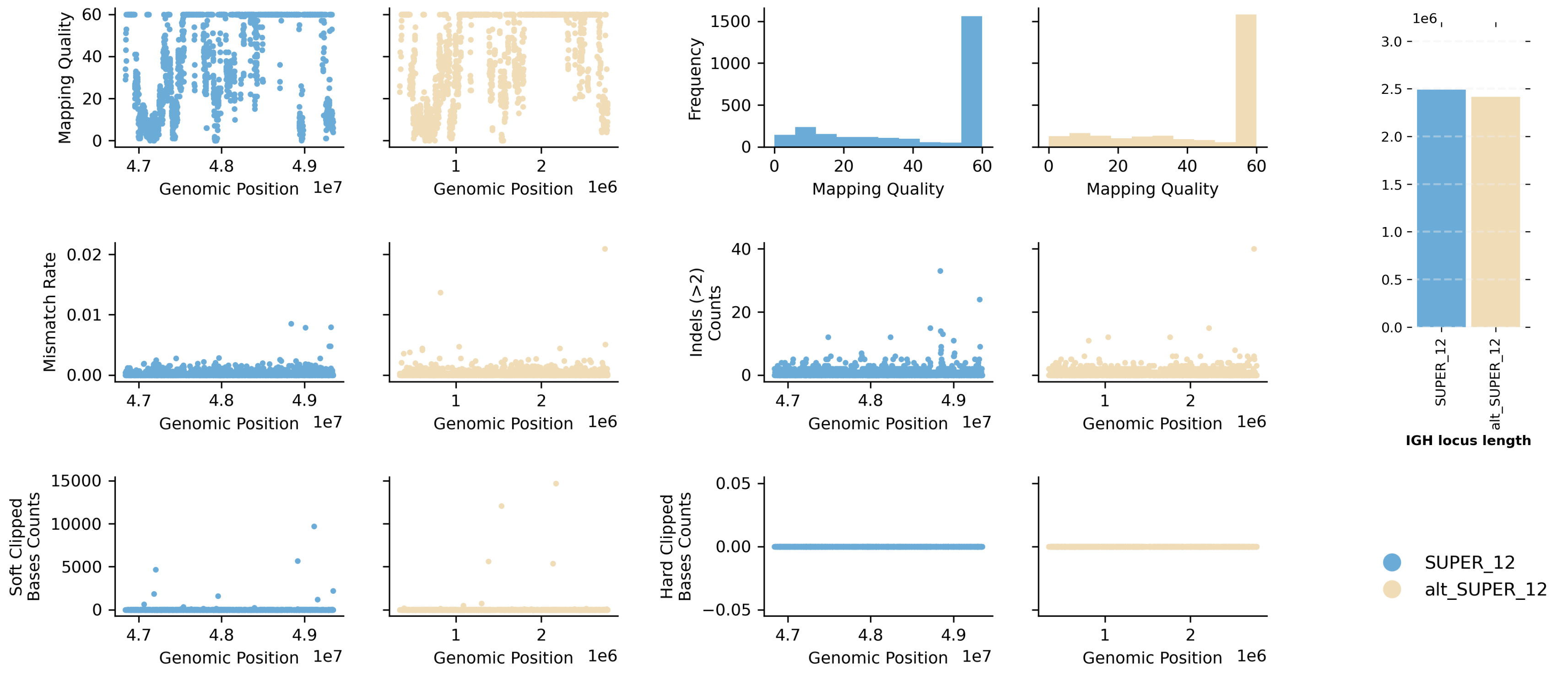


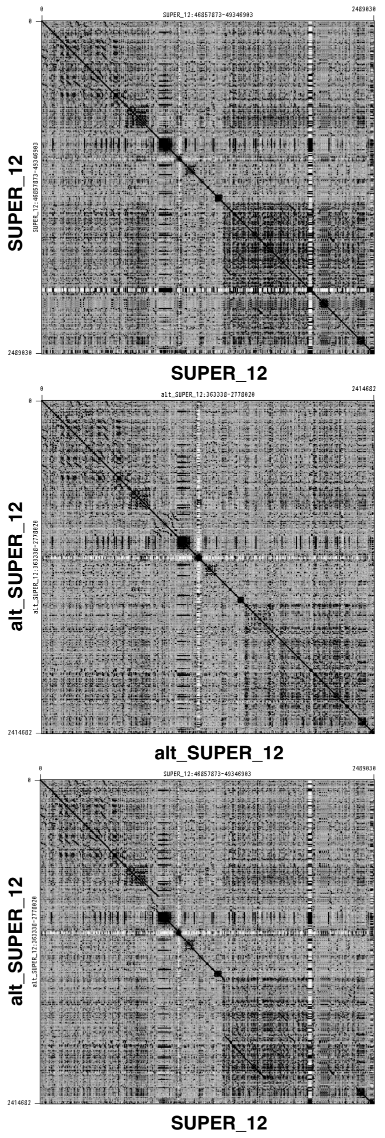


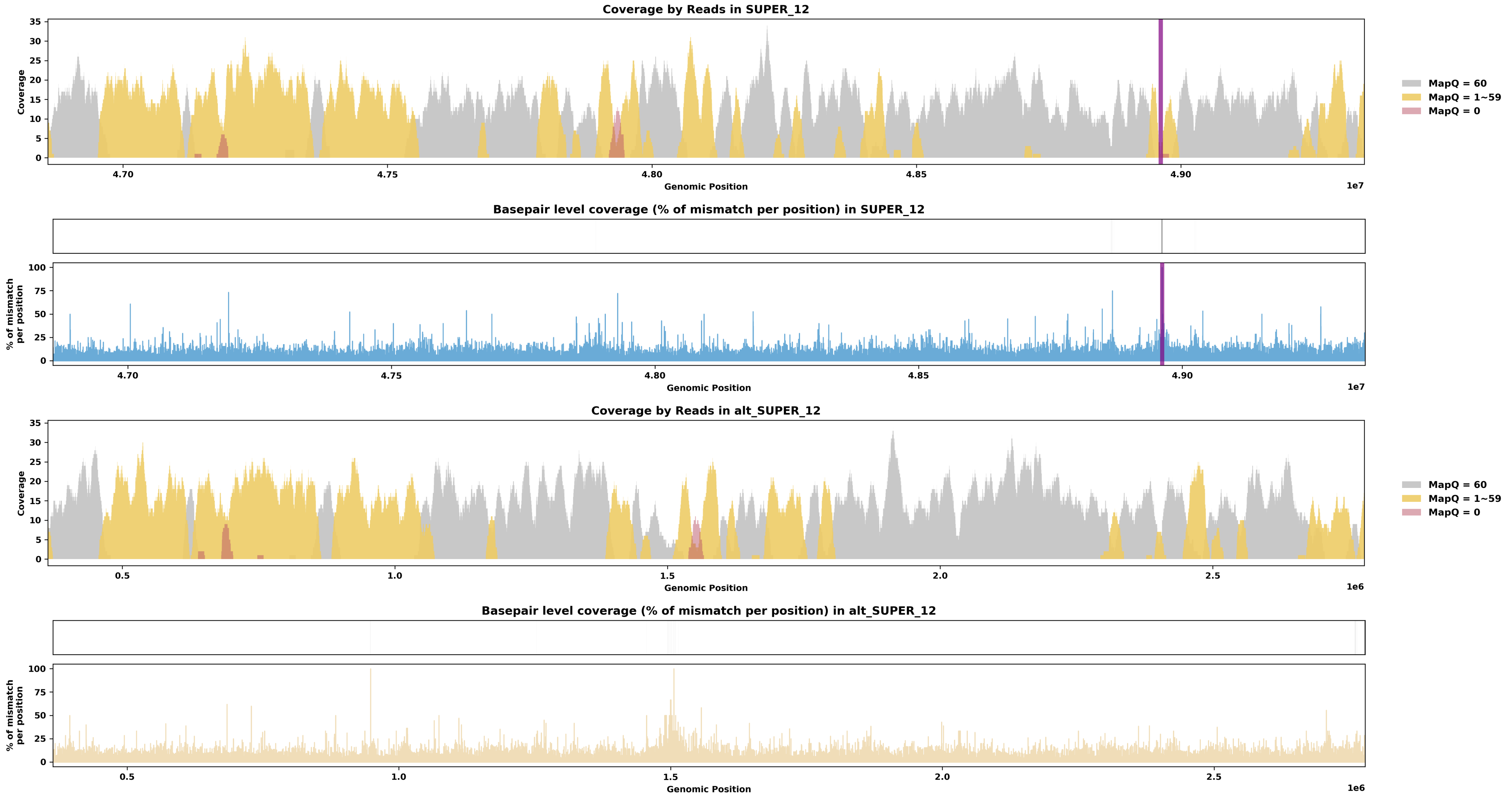

Species ID: rPodCre2

Common Name: cretan wall lizard

Scientific Name: Podarcis cretensis

Assembly Type: Not Haplotype Resolved

Data Source: VGP

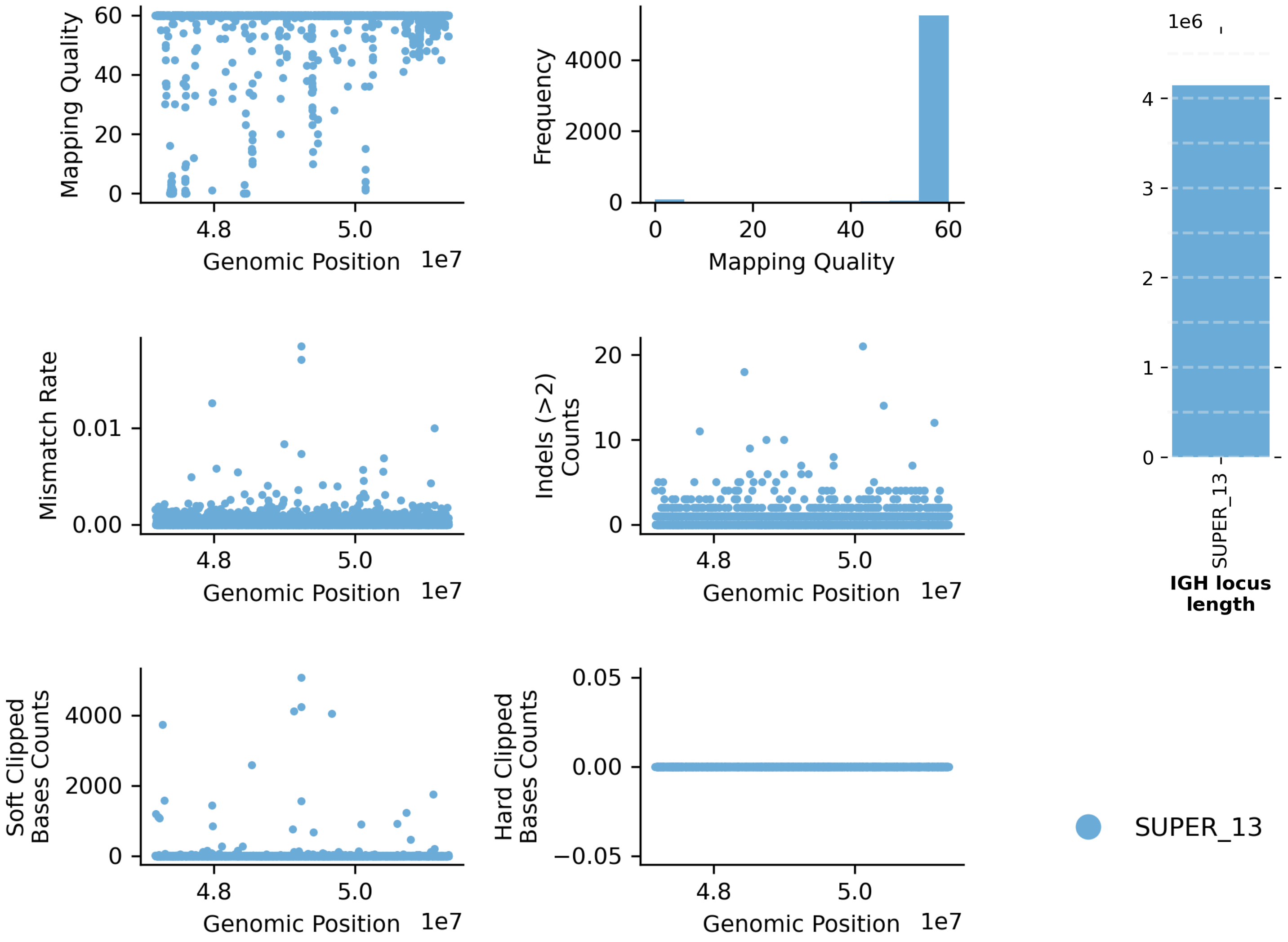


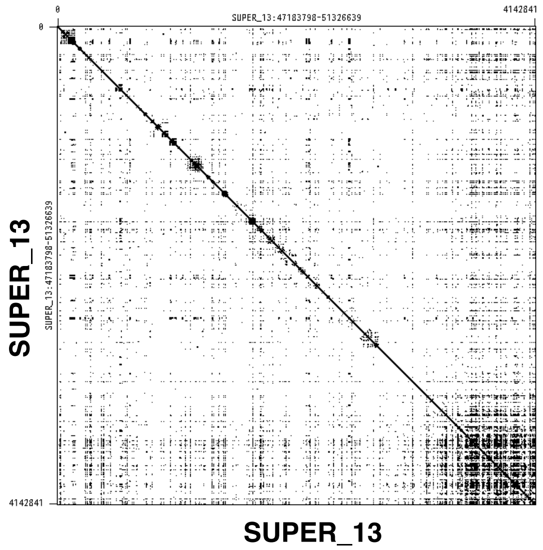


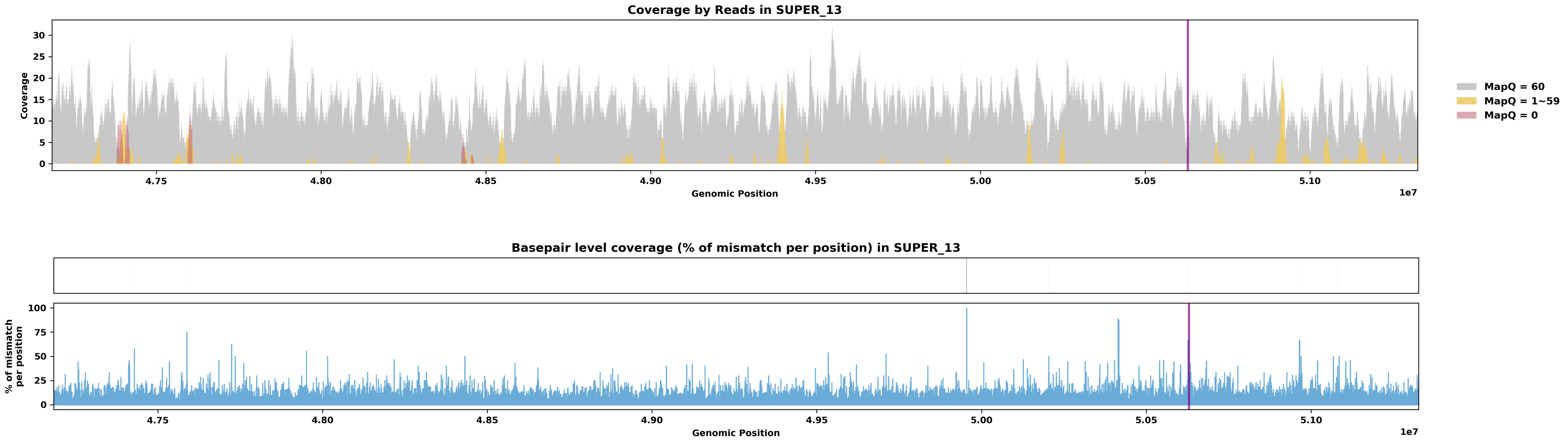

Species ID: rPodRaf1

Common Name: Aeolian wall lizard

Scientific Name: Podarcis raffonei

Assembly Type: Not Haplotype Resolved

Data Source: VGP

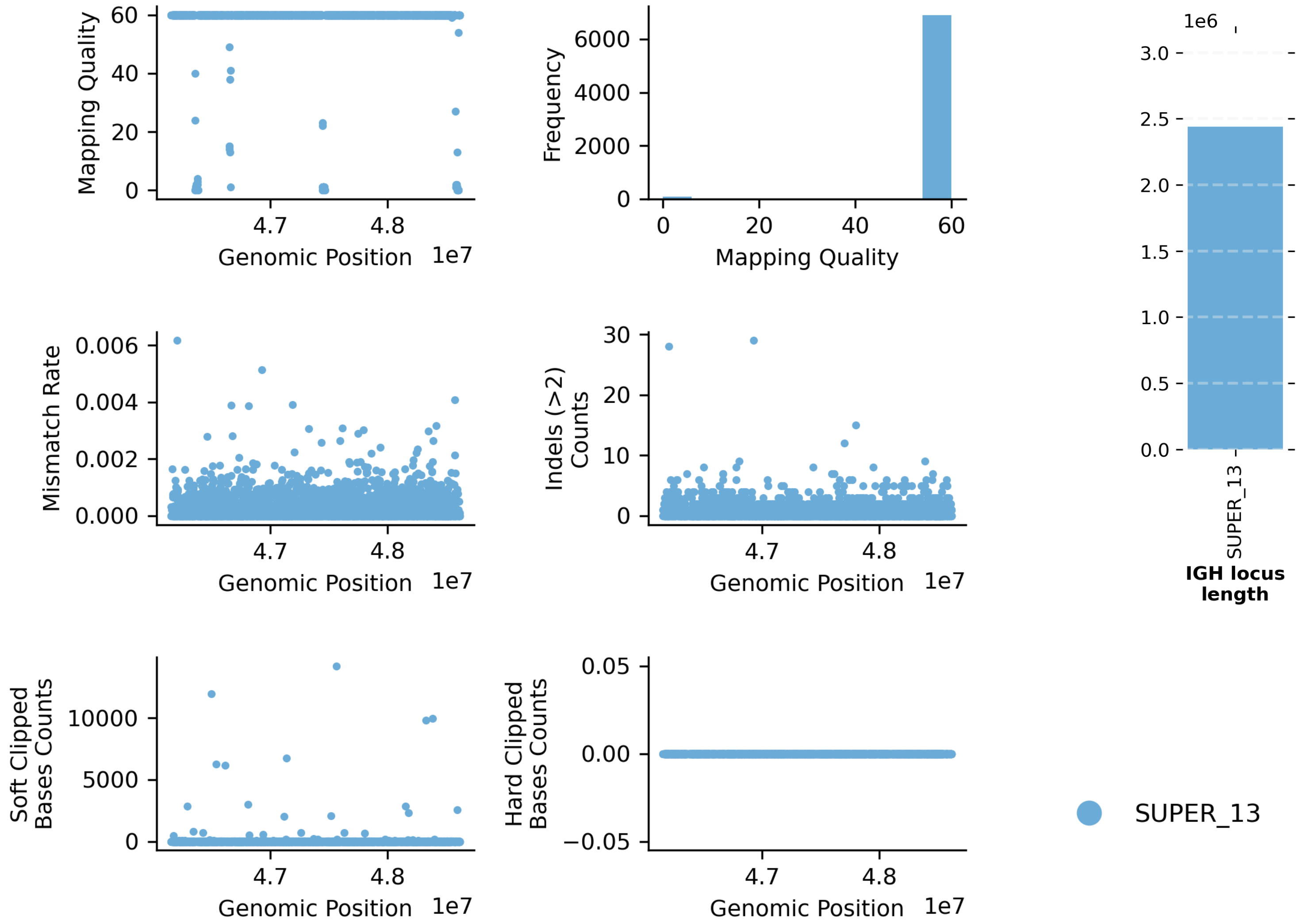


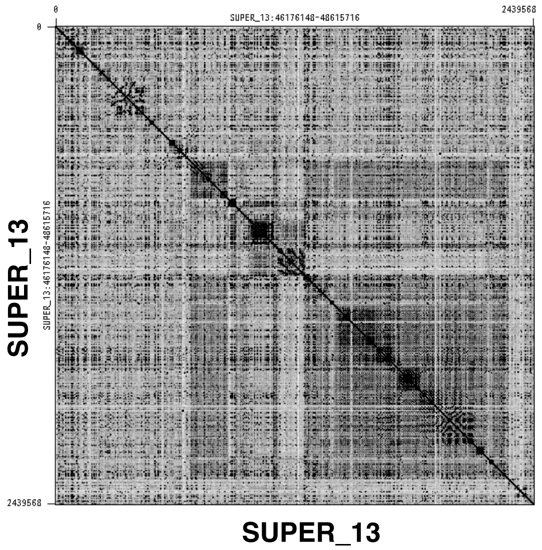


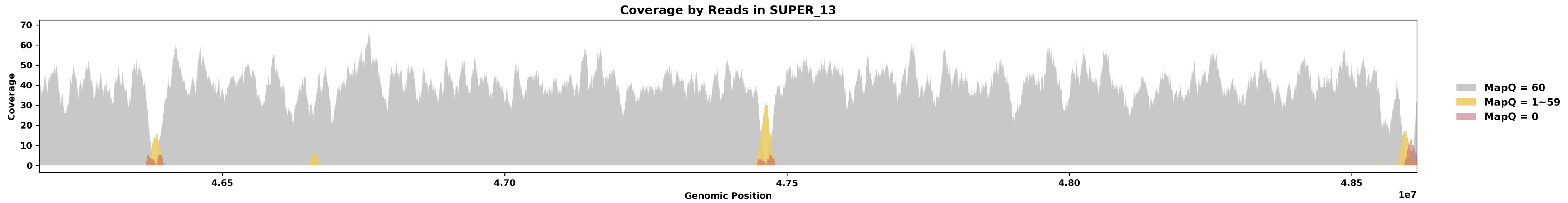


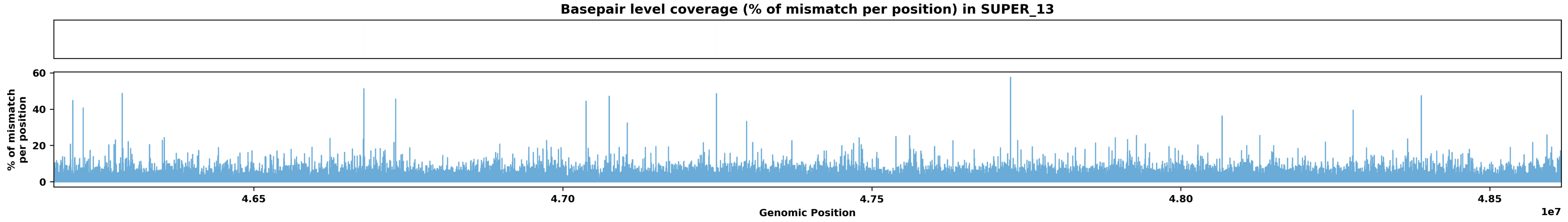

Species ID: rRhiFlo1

Common Name: Florida worm lizard

Scientific Name: Rhineura floridana

Assembly Type: Haplotype Resolved

Data Source: VGP

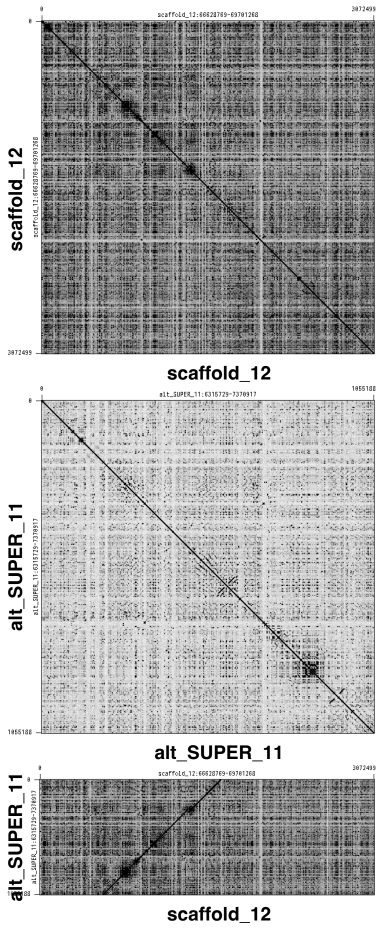


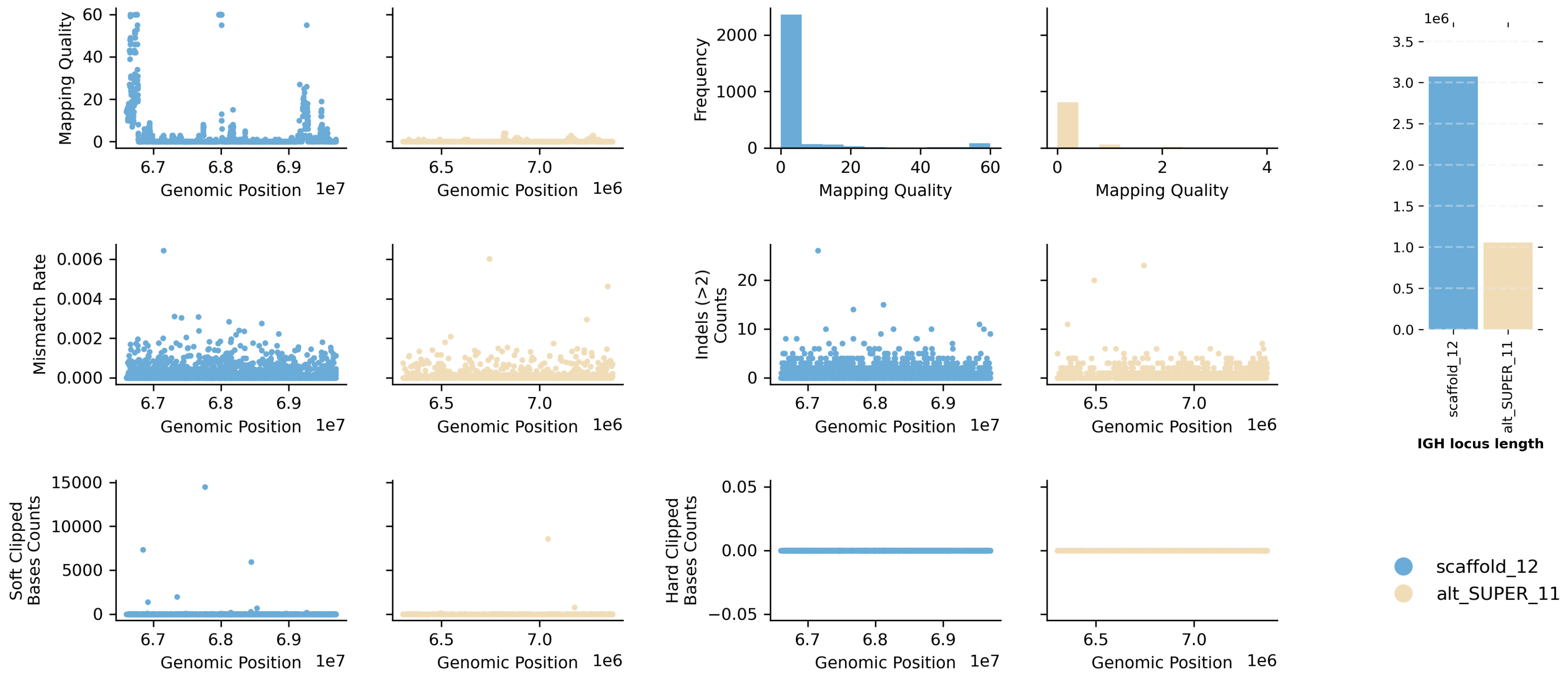


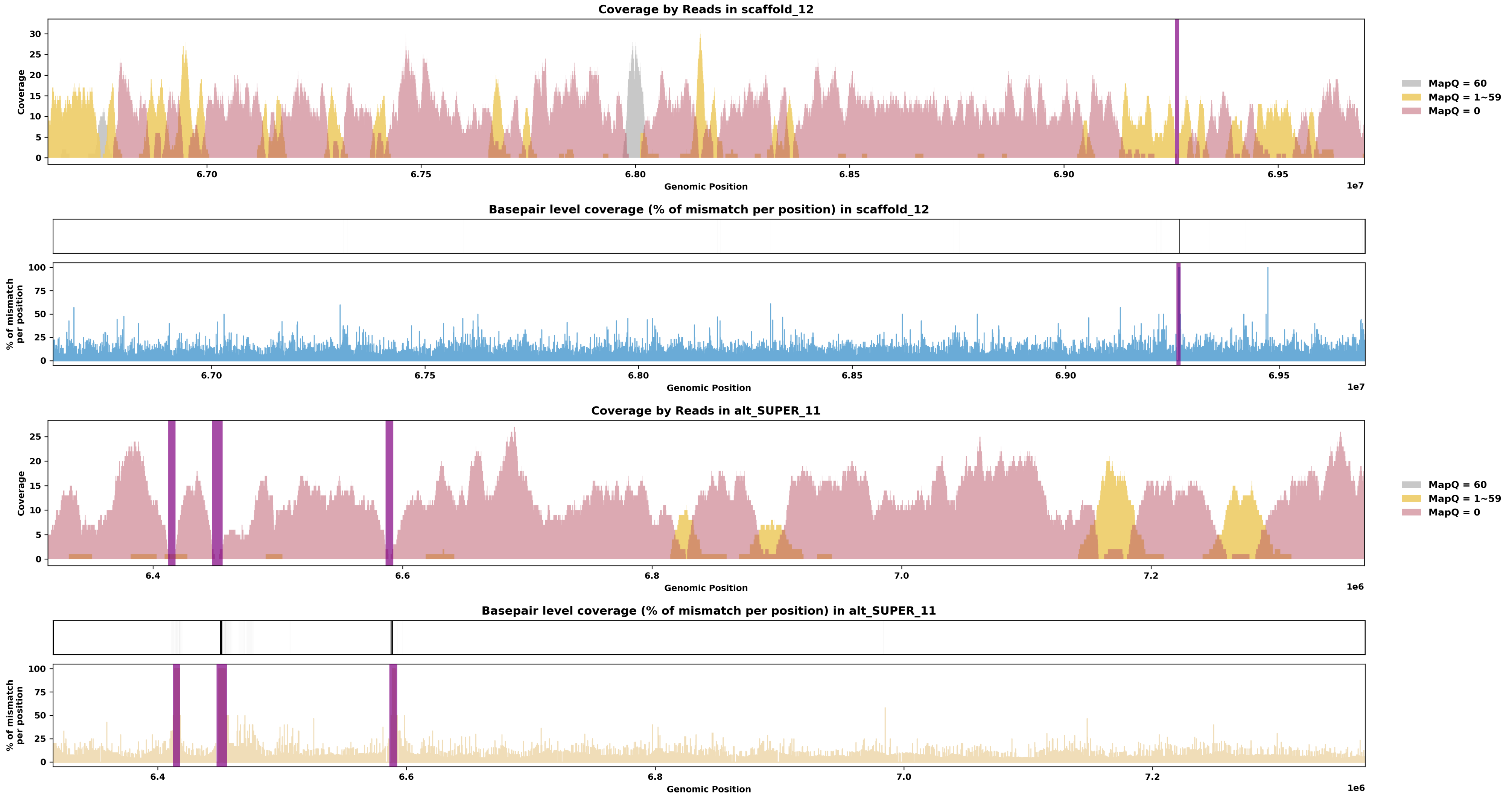

Species ID: rVipLat1

Common Name: snub-nosed viper

Scientific Name: Vipera latastei

Assembly Type: Not Haplotype Resolved

Data Source: VGP

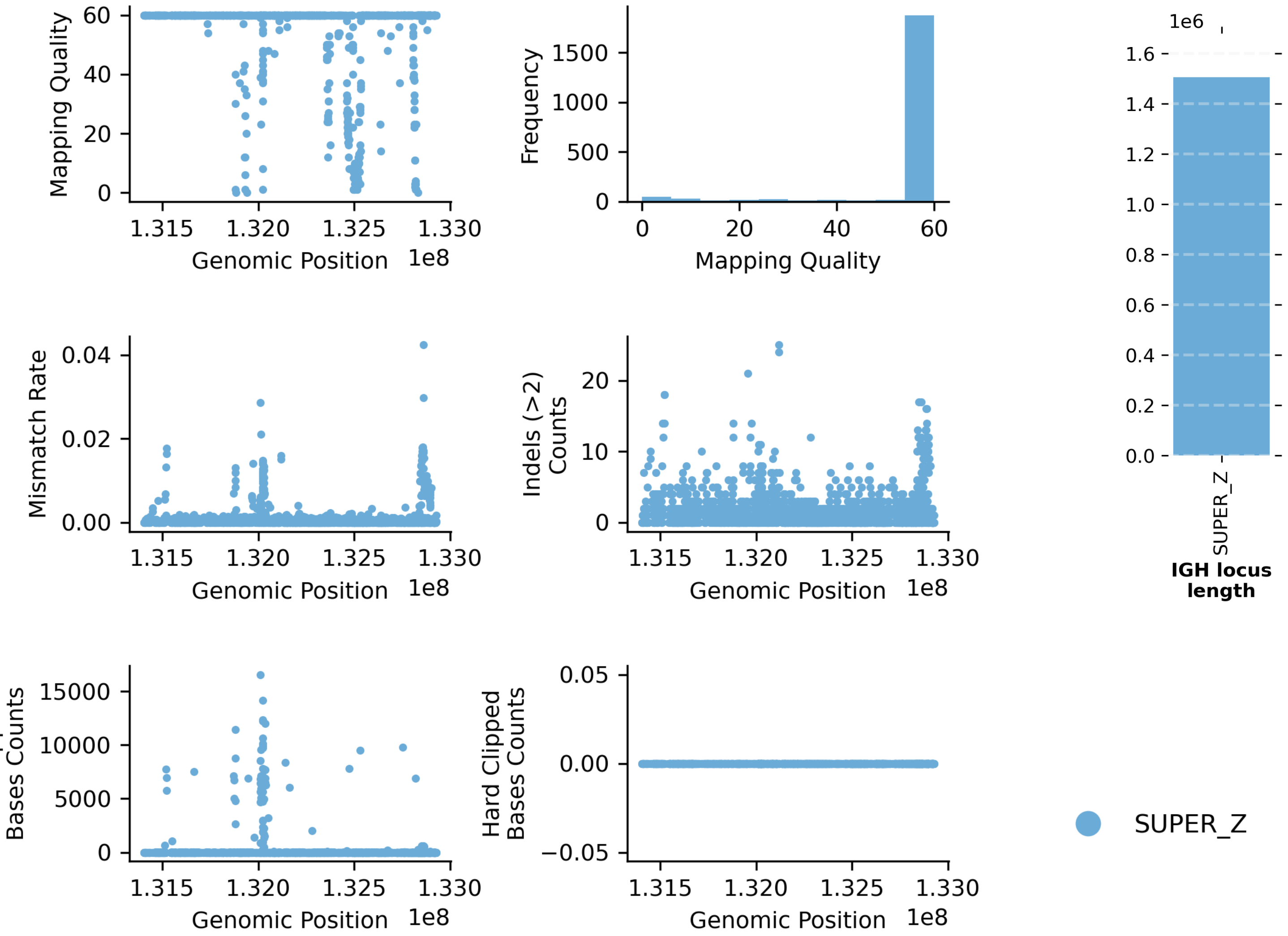


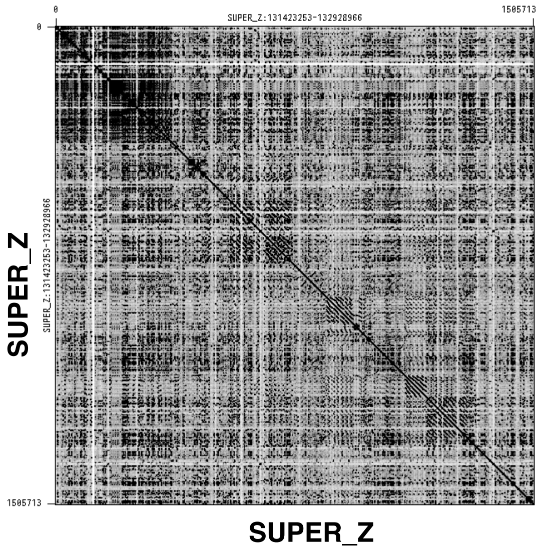


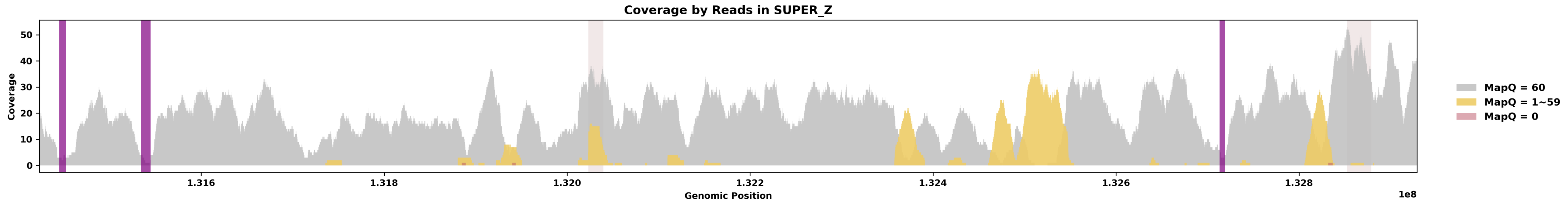


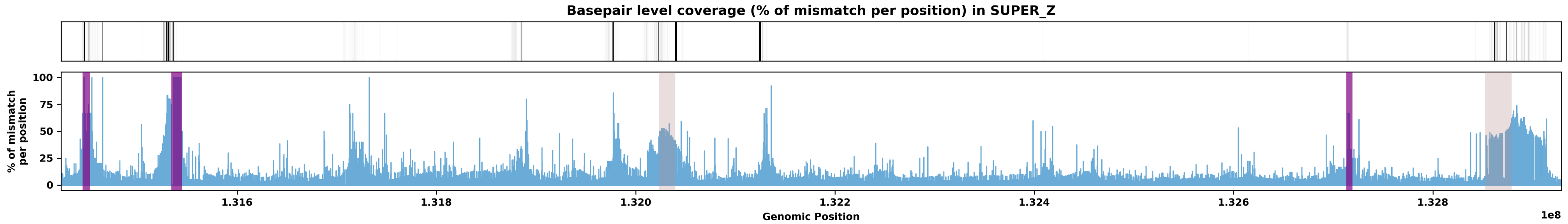

Species ID: rVipUrs1

Common Name: Hungarian meadow viper

Scientific Name: Vipera ursinii

Assembly Type: Not Haplotype Resolved

Data Source: VGP

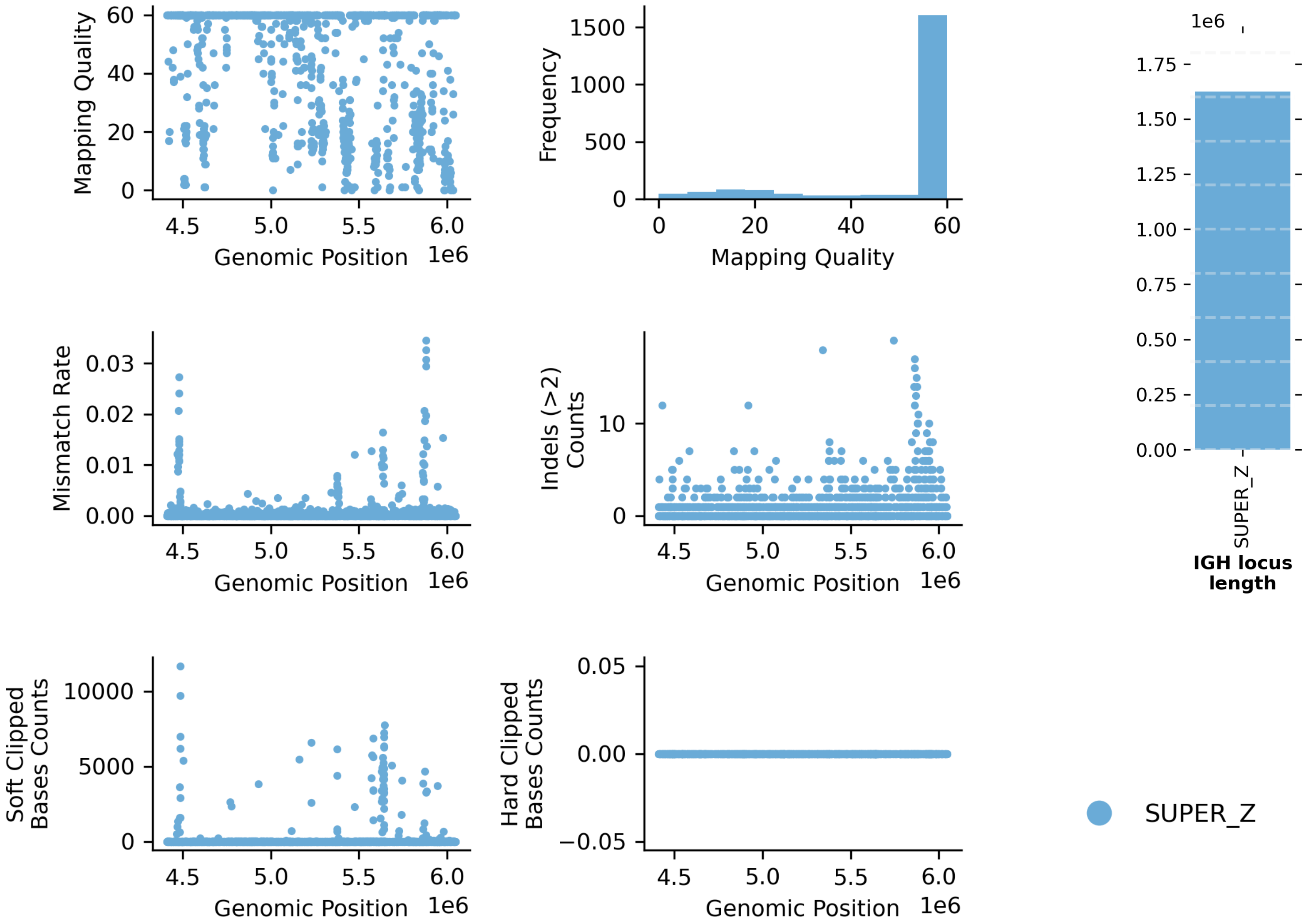


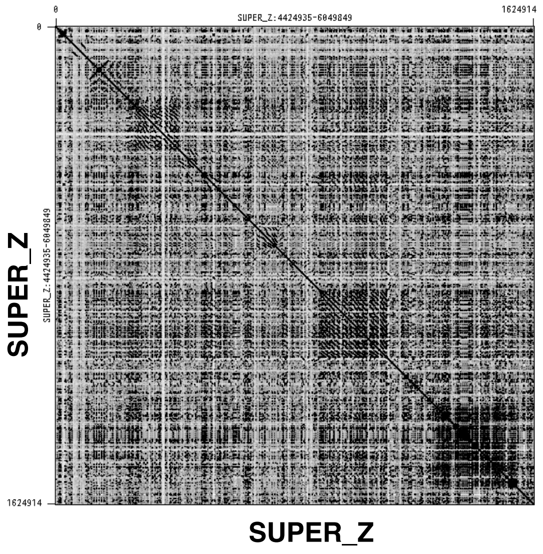


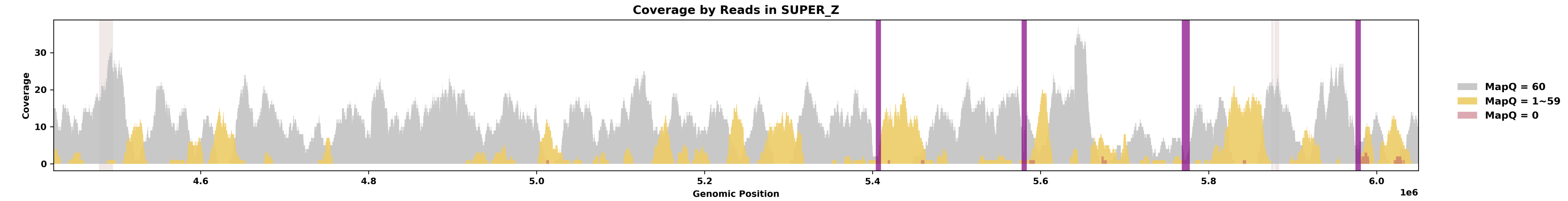


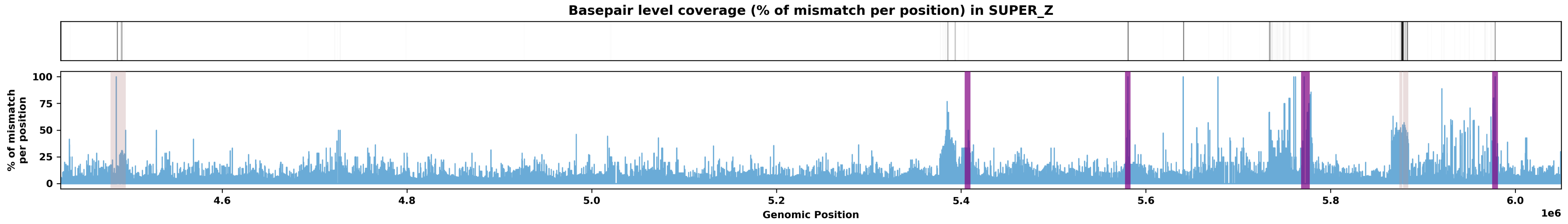

Species ID: rZooViv1

Common Name: common lizard

Scientific Name: Zootoca vivipara

Assembly Type: Not Haplotype Resolved

Data Source: VGP

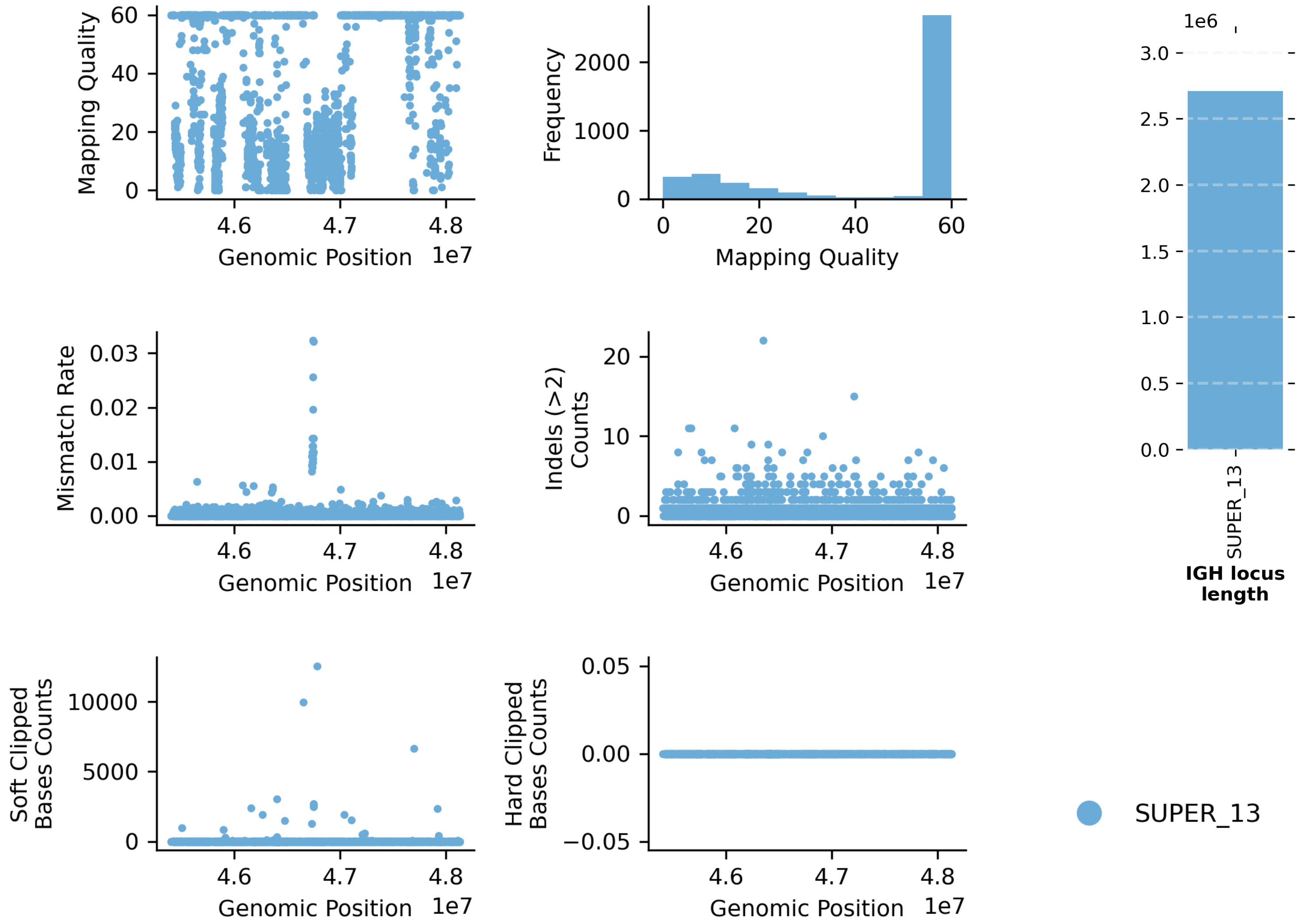


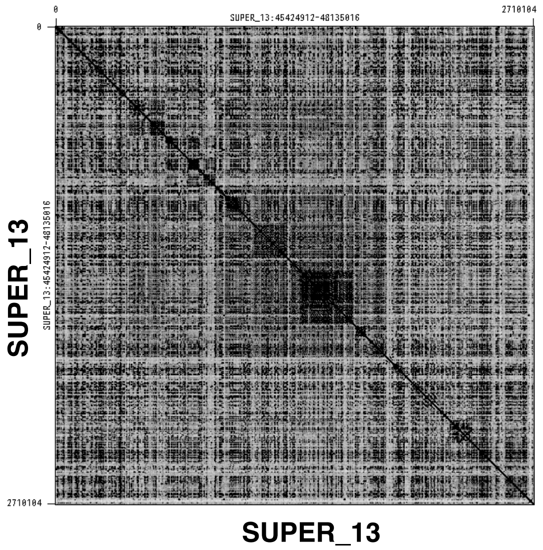


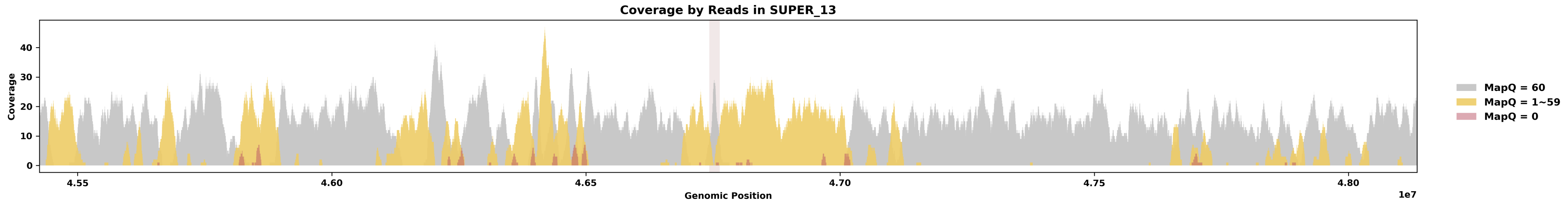


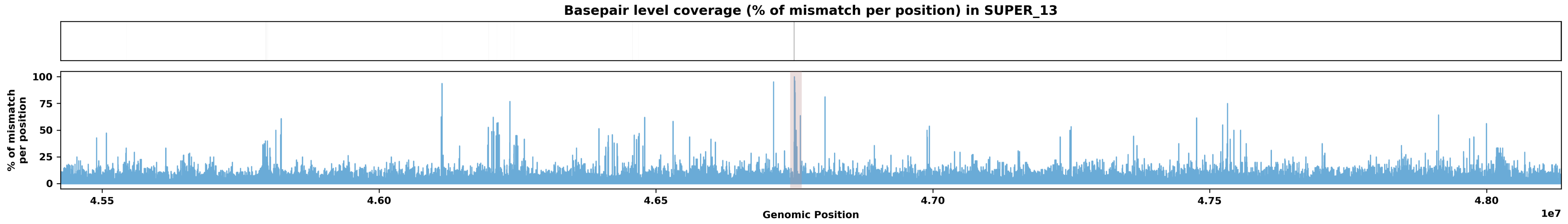
