## Supplementary material for "Assessing Assembly Errors in Immunoglobulin Loci: A Comprehensive Evaluation of Long-read Genome Assemblies Across Vertebrates": IGK Loci Evaluations

Table of Contents

1. mApoSyl1 - Apodemus sylvaticus - Wood mouse
2. mBalAcu1 - Balaenoptera acutorostrata - Minke whale
3. mCamDro1 - Camelus dromedarius - Dromedary
4. mCanLor1 - Canis lupus - Greenland Wolf
5. mCanLor2 - Canis lupus - Greenland Wolf
6. mCerEla1 - Cervus elaphus - Red Deer
7. mChiNiv1 - Chionomys nivalis - European snow vole
8. mDasNov1 - Dasypus novemcinctus - Nine-banded armadillo
9. mDelDel1 - Delphinus delphis - Saddleback dolphin
10. mDicBic1 - Diceros bicornis - Black rhinoceros
11. mDipMer1 - Dipodomys merriami - Merriam's Kangaroo Rat
12. mEleMax1 - Elephas maximus - Asiatic Elephant
13. mEriEur2 - Erinaceus europaeus - Western European hedgehog
14. mEscRob2 - Eschrichtius robustus - Grey whale
15. mEubGla1 - Eubalaena glacialis - North Atlantic right whale
16. mGloMel1 - Globicephala melas - Long-finned pilot whale
17. mGorGor1 - Gorilla gorilla - Gorilla
18. mHetBru1 - Heterohyrax brucei - Yellow-spotted hyrax
19. mHipAmp2 - Hippopotamus amphibius kiboko - Hippopotamus
20. mHypAmp2 - Hyperoodon ampullatus - Northern bottlenose whale
21. mLagAlb1 - Lagenorhynchus albirostris - White-beaked dolphin
22. mLemCat1 - Lemur catta - Ring-tailed lemur
23. mLynRuf1 - Lynx rufus - Bobcat
24. mMacEug1 - Macropus eugenii - Tammar wallaby
25. mManPen7 - Manis pentadactyla - Chinese pangolin
26. mMarMar1 - Martes martes - European pine marten
27. mMelMel3 - Meles meles - European badger
28. mMesDen1 - Mesoplodon densirostris - Blainville's beaked whale
29. mMicCal1.0 - Microtus californicus - California Vole
30. mMicMin1 - Micromys minutus - European harvest mouse
31. mMirAng1 - Mirounga angustirostris - Northern Elephant Seal
32. mMonDom1 - Monodelphis domestica - Gray short-tailed opossum
33. mMunRee1 - Muntiacus reevesi - Reeves' muntjac
34. mMusAve1 - Muscardinus avellanarius - Hazel dormouse
35. mMusLut2 - Mustela lutreola - European mink
36. mMusNiv1 - Mustela nivalis - Least weasel
37. mNeoNeb1 - Neofelis nebulosa - Clouded Leopard
38. mNycCou1 - Nycticebus coucang - Slow loris
39. mOrcOrc1 - Orcinus orca - Killer whale
40. mOryCun1 - Oryctolagus cuniculus - Rabbit
41. mPanPan1 - Pan paniscus - Bonobo
42. mPerMan1 - Peromyscus maniculatus - Deer mouse
43. mPhoPho1 - Phocoena phocoena - Harbor porpoise
44. mPonAbe1 - Pongo abelii - Sumatran orangutan
45. mPonPyg2 - Pongo pygmaeus - Bornean orangutan
46. mPseCra1 - Pseudorca crassidens - False killer whale
47. mPumCon1.1 - Puma concolor - Mountain Lion
48. mSorAra2/1 - Sorex araneus - Common shrew
49. mSteCoe1 - Stenella coeruleoalba - Striped dolphin
50. mTalEur1 - Talpa europaea - European mole
51. mThoBot1 - Thomomys bottae - Botta's pocket gopher
52. mUrsAme1 - Ursus americanus - American black bear
53. mUrsArc2 - Ursus arctos - Brown bear
54. rAllMis2 - Alligator mississippiensis - American alligator
55. rCarCar2 - Caretta caretta - Loggerhead turtle
56. rEmyOrb1 - Emys orbicularis - European pond turtle
57. rMalTer1 - Malaclemys terrapin - Diamondback terrapin

Note: This supplementary material only displays alternate IG loci if they are longer than one-quarter of the corresponding primary IG locus length. Loci shorter than this threshold are typically too fragmented to be shown. 11 species had alternate IGK loci shorter than this threshold and are not shown.

Species ID: mApoSyl1

Common Name: wood mouse

Scientific Name: Apodemus sylvaticus

Assembly Type: Not Haplotype Resolved

Data Source: VGP

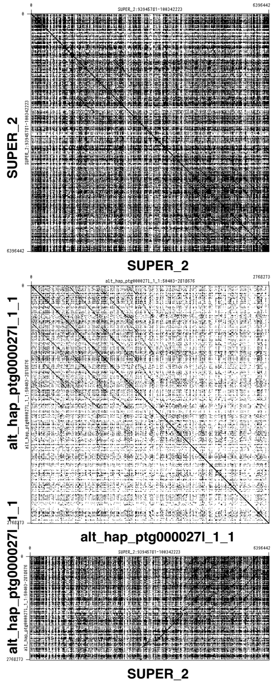


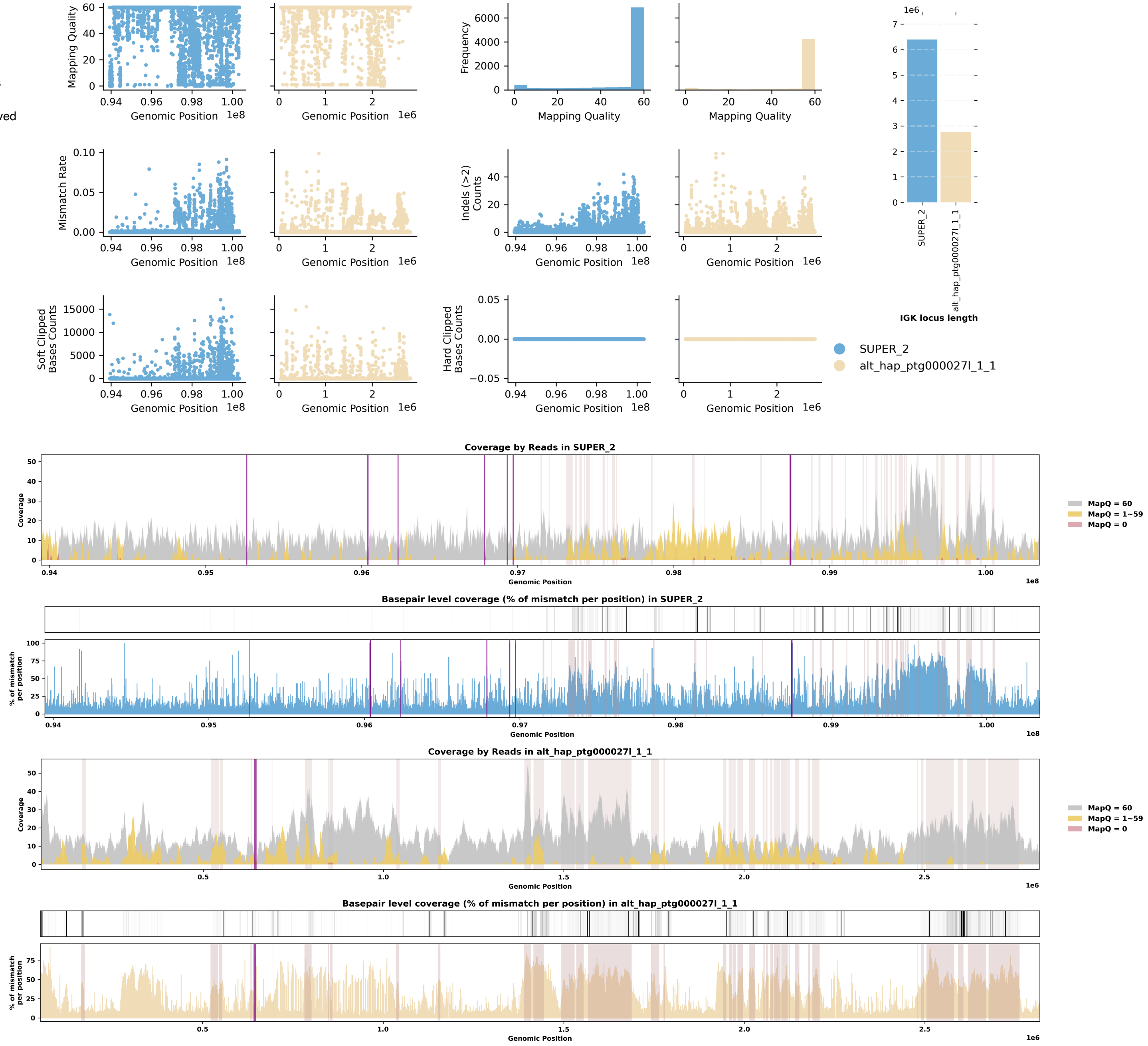

Species ID: mBalAcu1

Common Name: minke whale

Scientific Name: Balaenoptera_acutorostrata

Assembly Type: Not Haplotype Resolved

Data Source: VGP

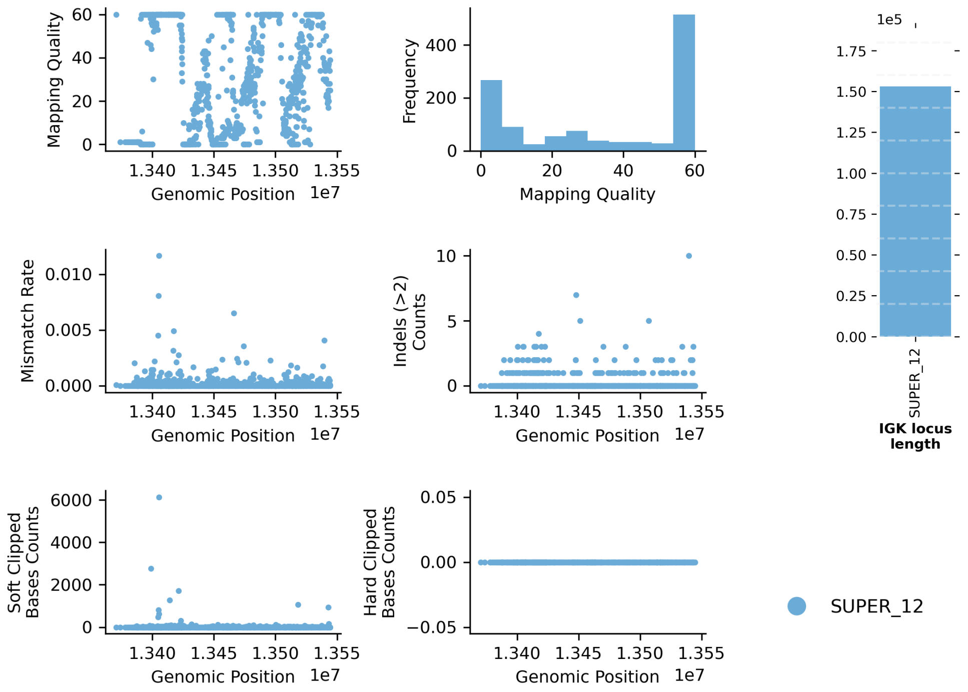


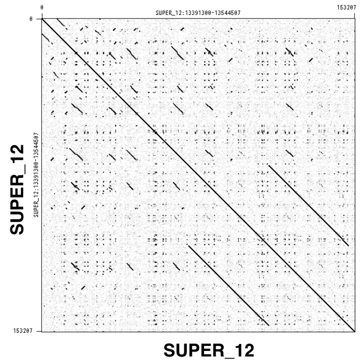


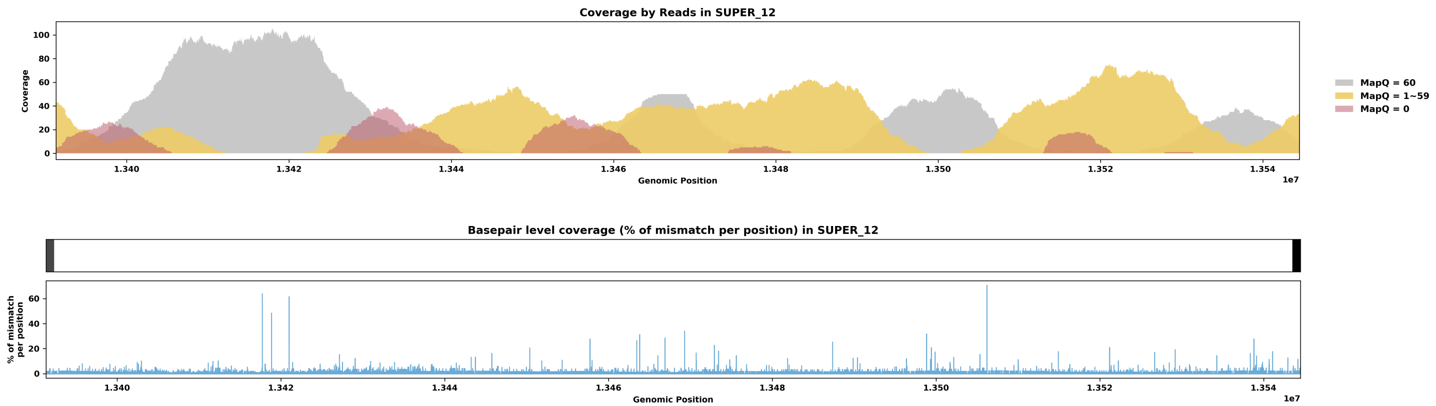

Species ID: mCamDro1

Common Name: dromedary

Scientific Name: Camelus dromedarius

Assembly Type: Haplotype Resolved

Data Source: VGP

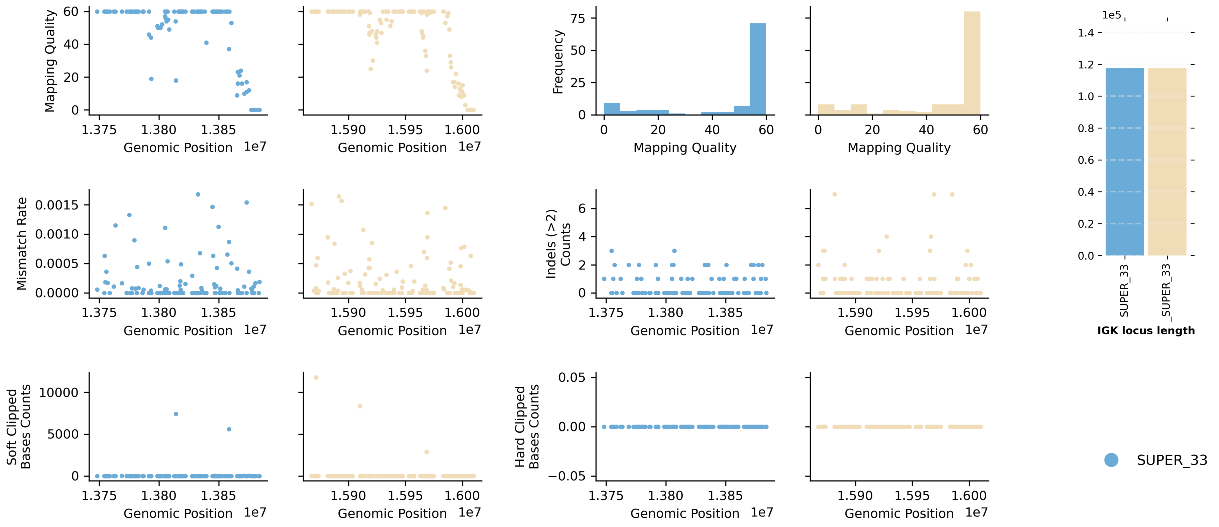


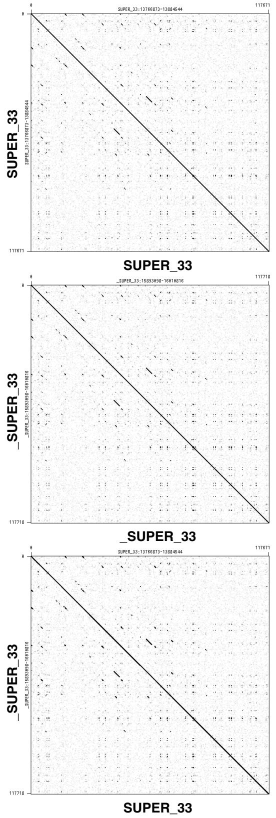


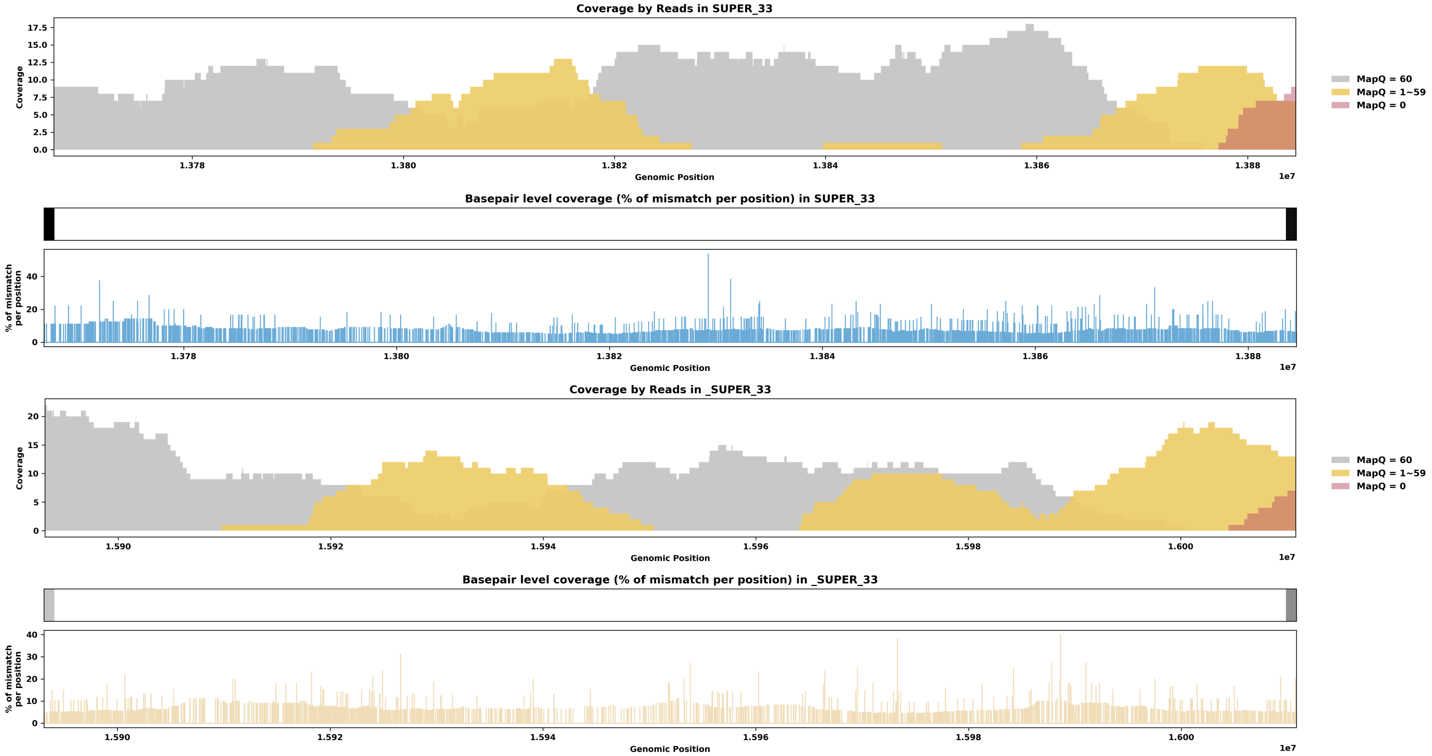

Species ID: mCanLor1

Common Name: Greenland Wolf

Scientific Name: Canis_lupus

Assembly Type: Not Haplotype Resolved

Data Source: VGP

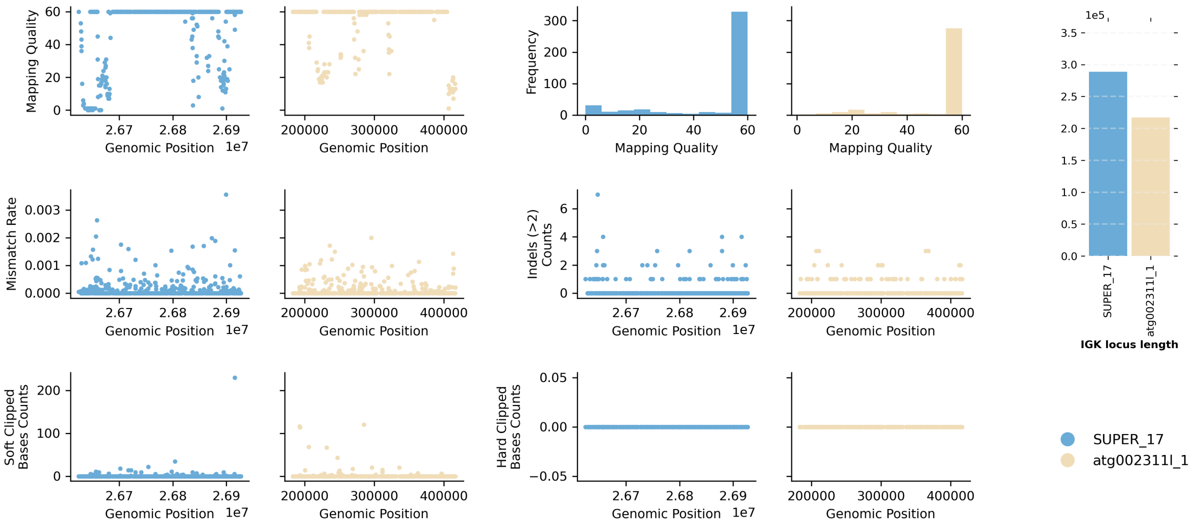


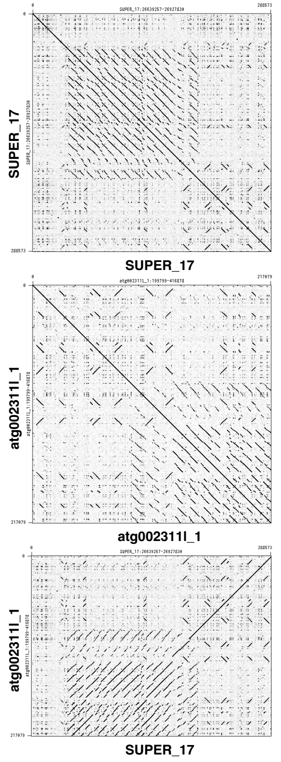


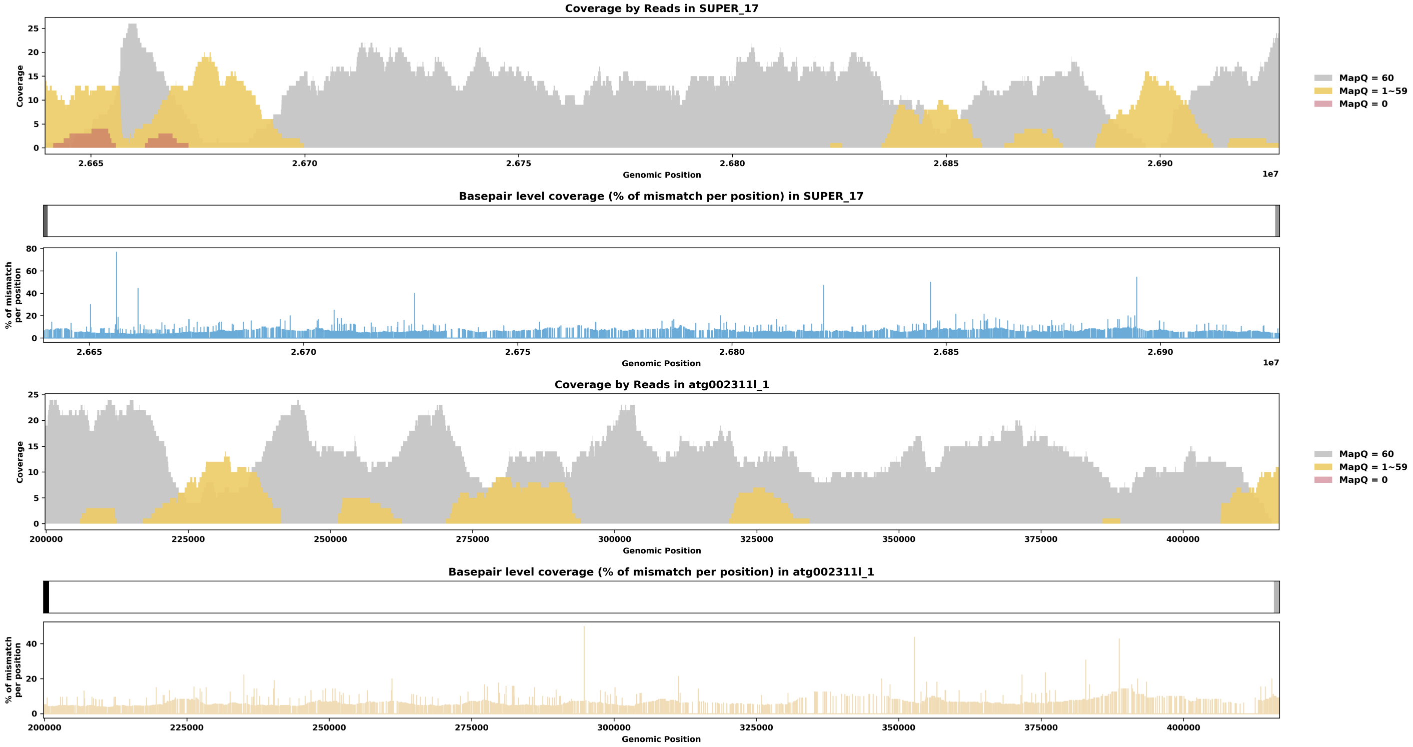

Species ID: mCanLor2

Common Name: Greenland Wolf

Scientific Name: Canis_lupus

Assembly Type: Haplotype Resolved

Data Source: VGP

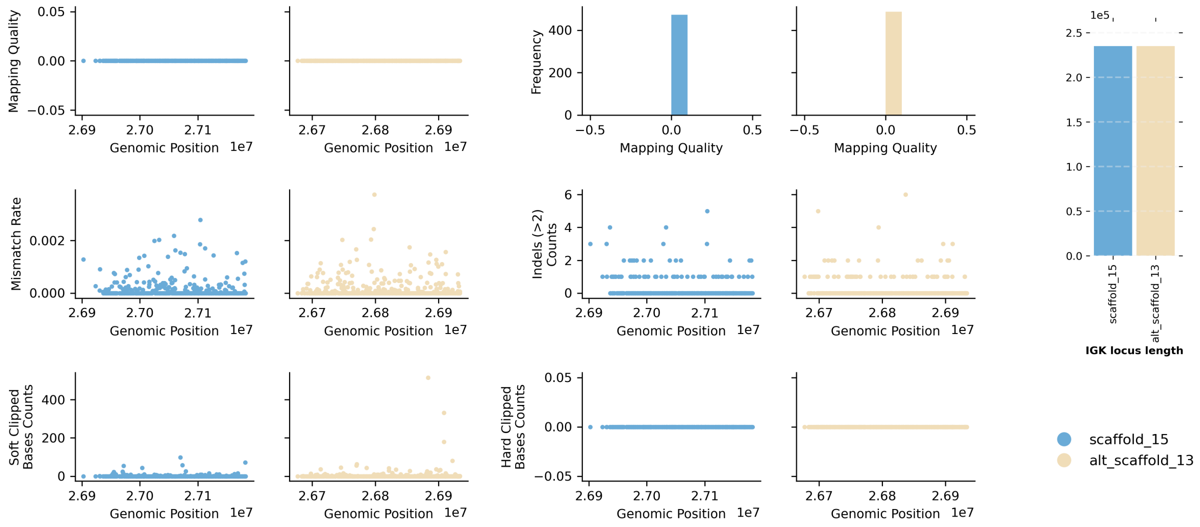


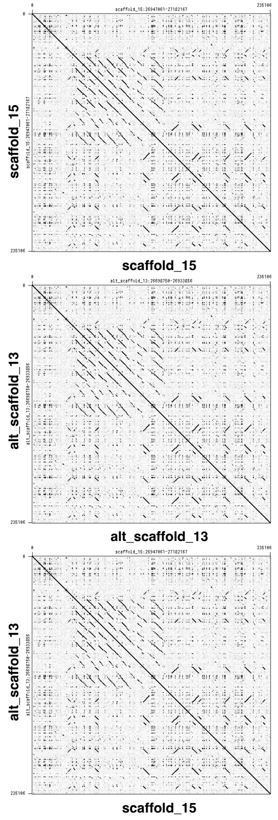


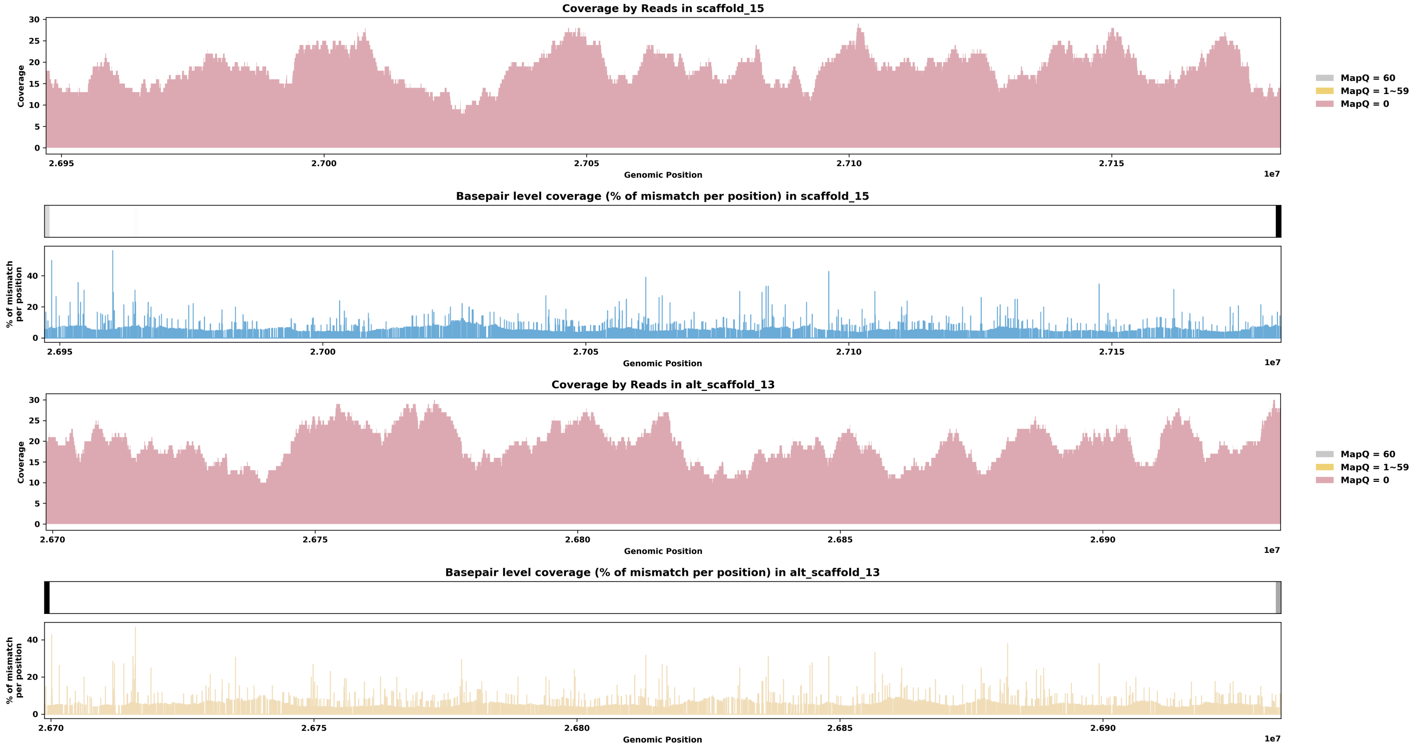

Species ID: mCerEla1

Common Name: Red Deer

Scientific Name: Cervus_elaphus

Assembly Type: Not Haplotype Resolved

Data Source: VGP

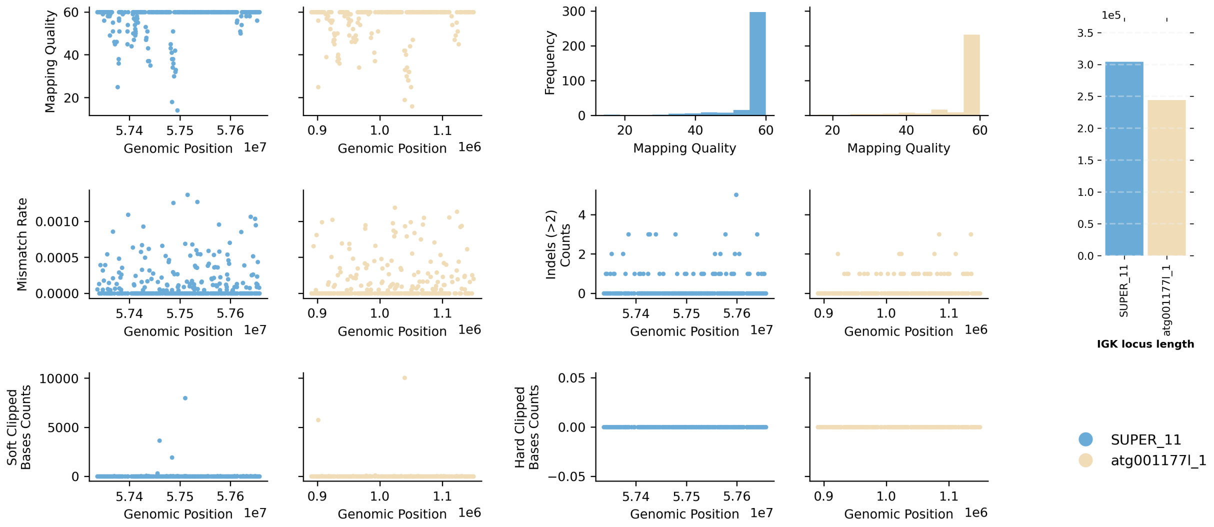


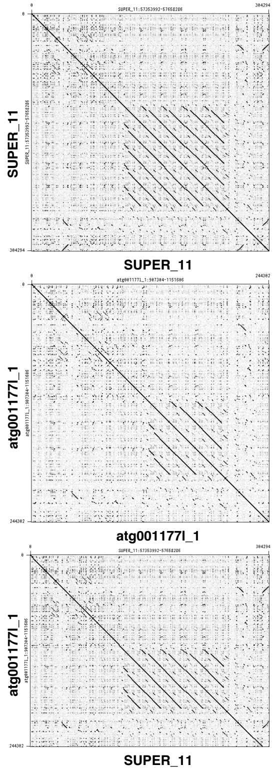


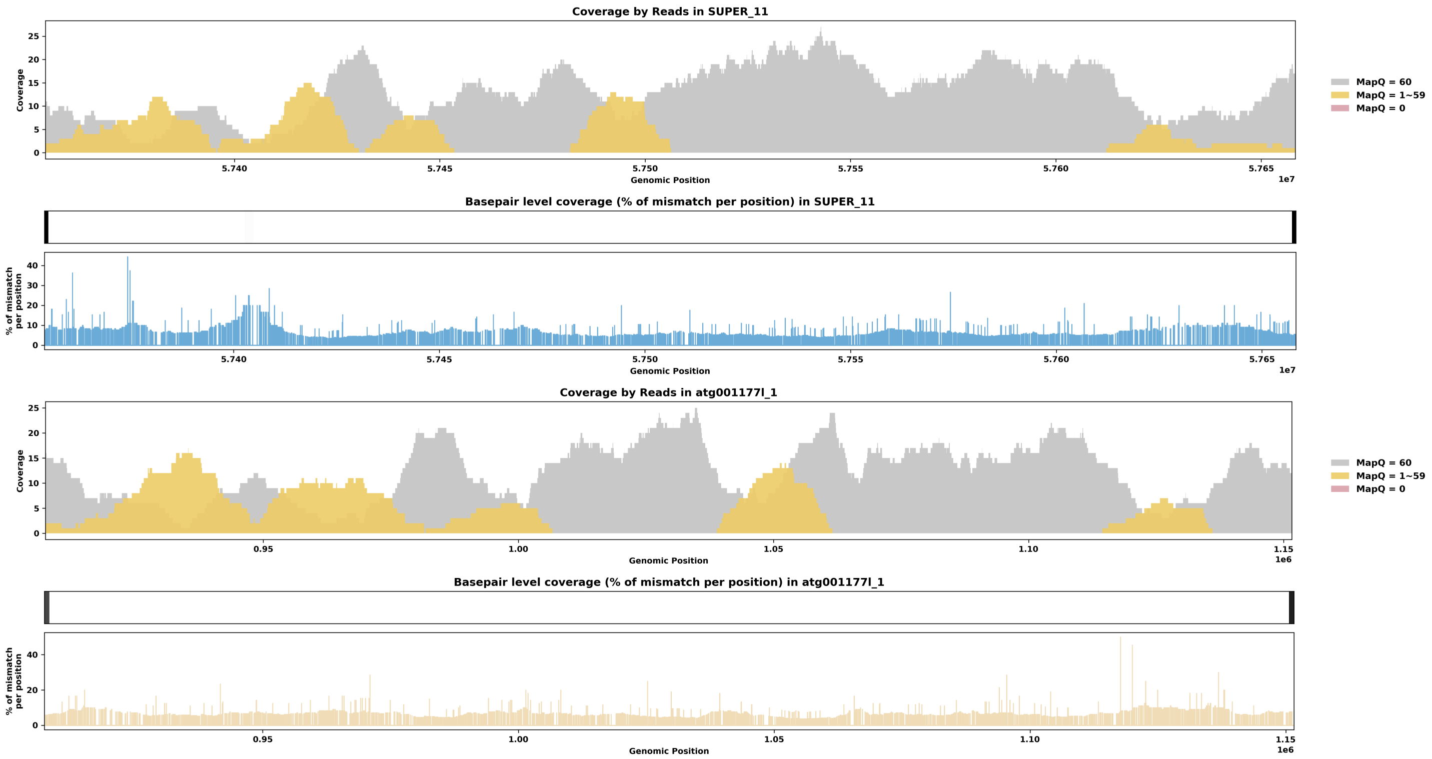

Species ID: mChiNiv1

Common Name: European snow vole

Scientific Name: Chionomys nivalis

Assembly Type: Not Haplotype Resolved

Data Source: VGP

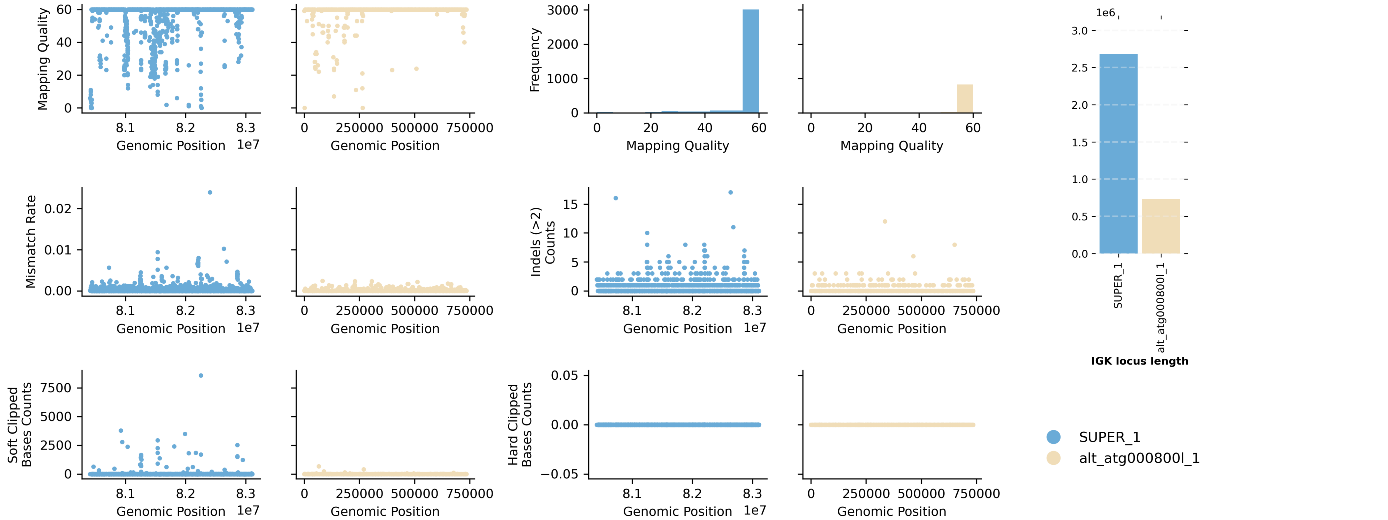


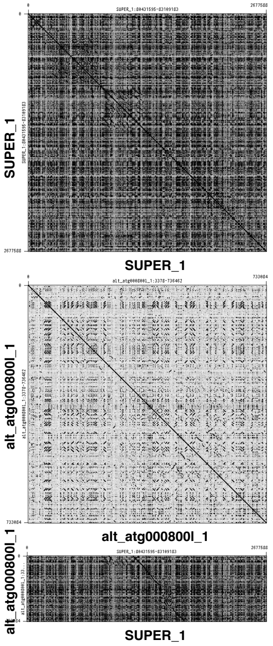


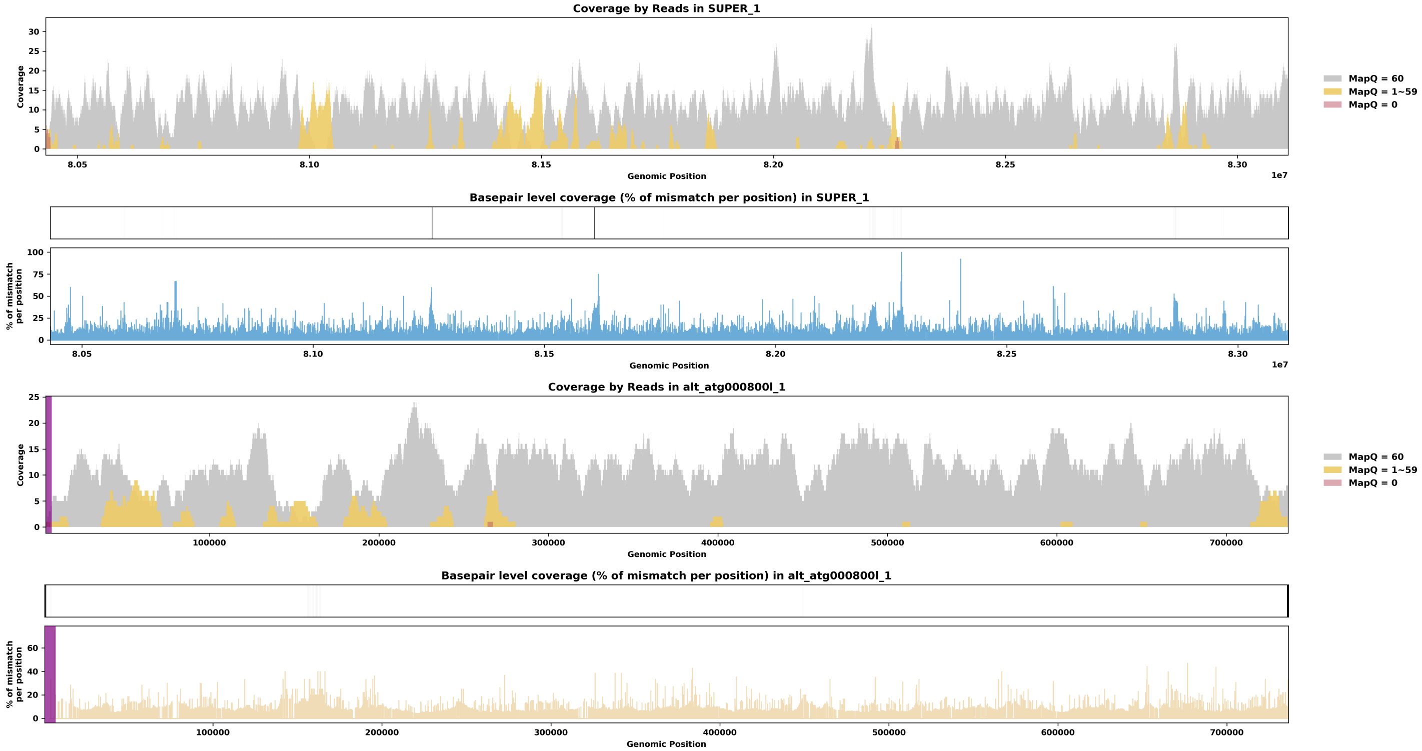

Species ID: mDasNov1

Common Name: nine-banded armadillo

Scientific Name: Dasypus novemcinctus

Assembly Type: Haplotype Resolved

Data Source: VGP

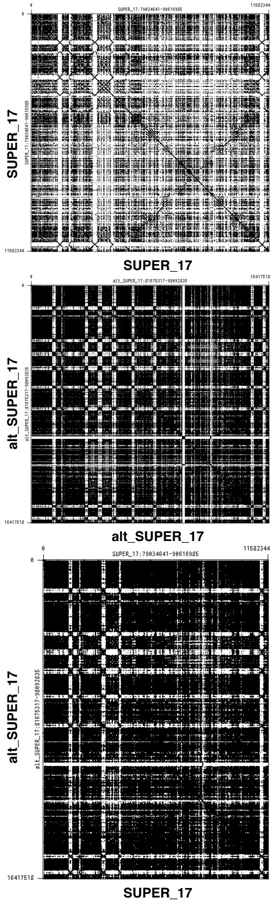


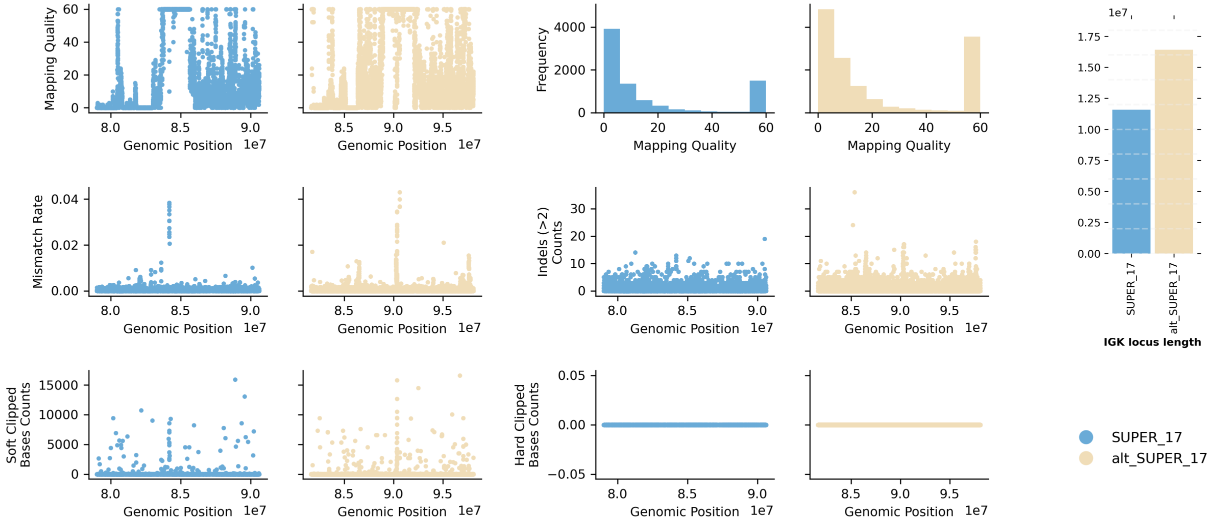


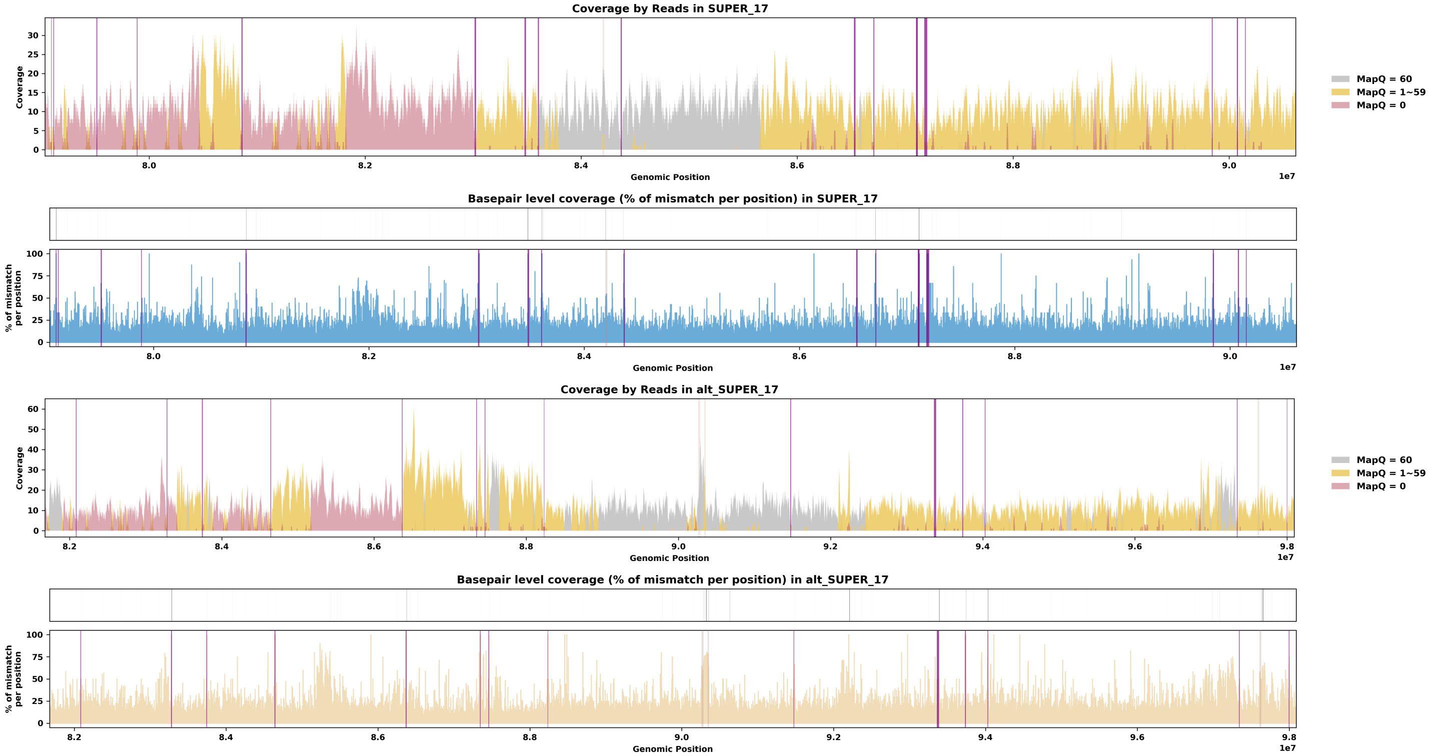

Species ID: mDelDel1

Common Name: saddleback dolphin

Scientific Name: Delphinus delphis

Assembly Type: Not Haplotype Resolved

Data Source: VGP

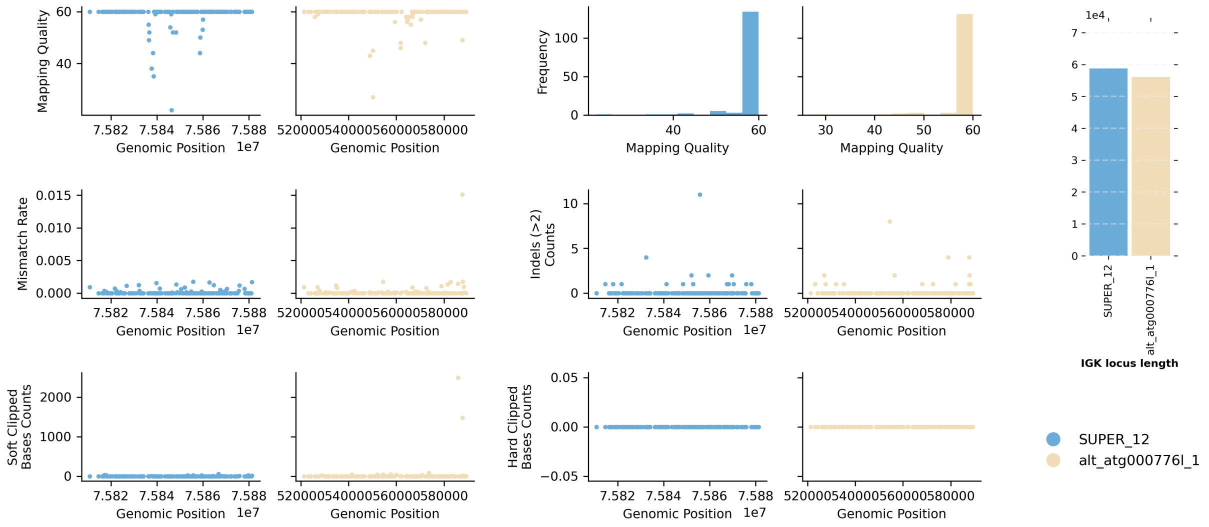


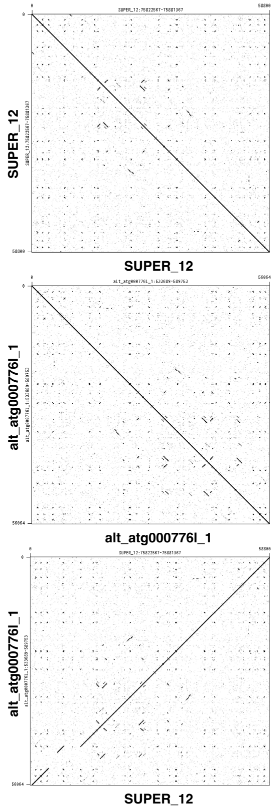


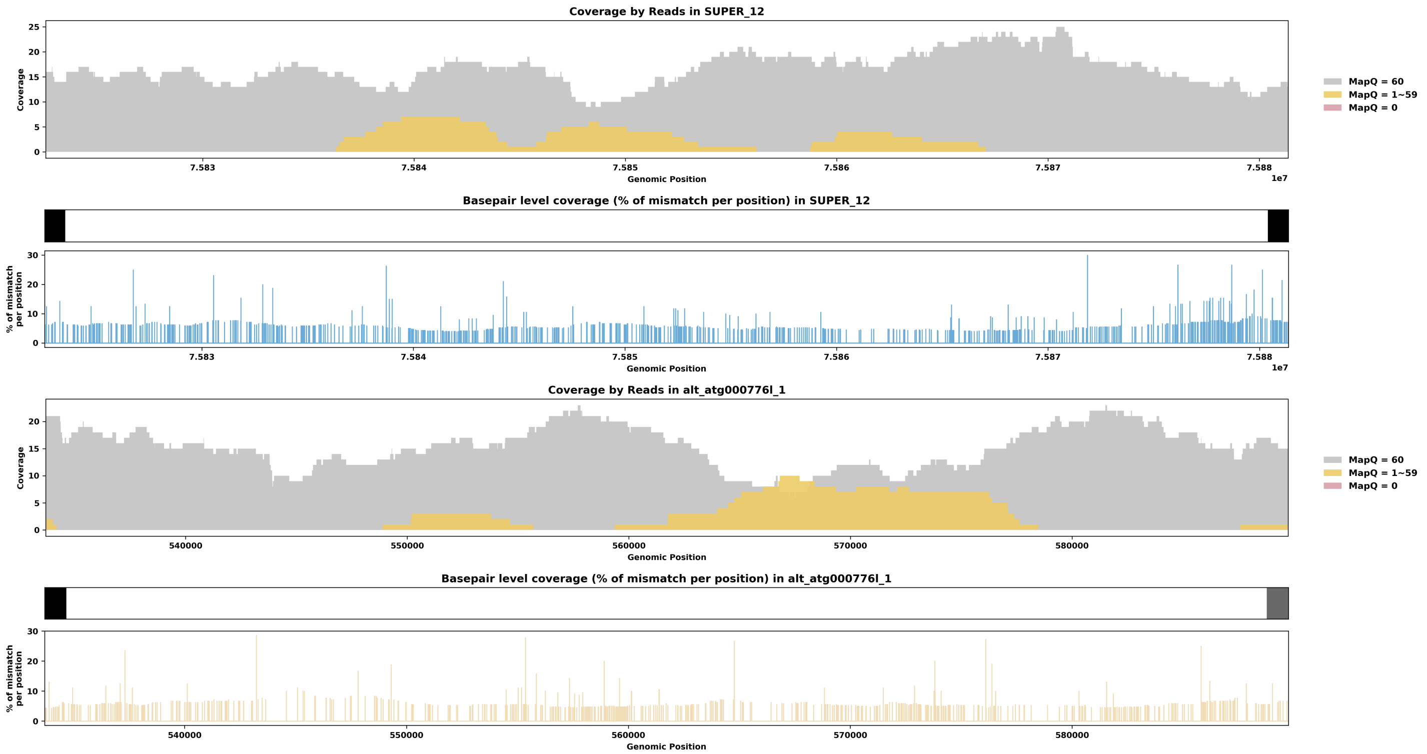

Species ID: mDicBic1

Common Name: black rhinoceros

Scientific Name: Diceros bicornis

Assembly Type: Haplotype Resolved

Data Source: VGP

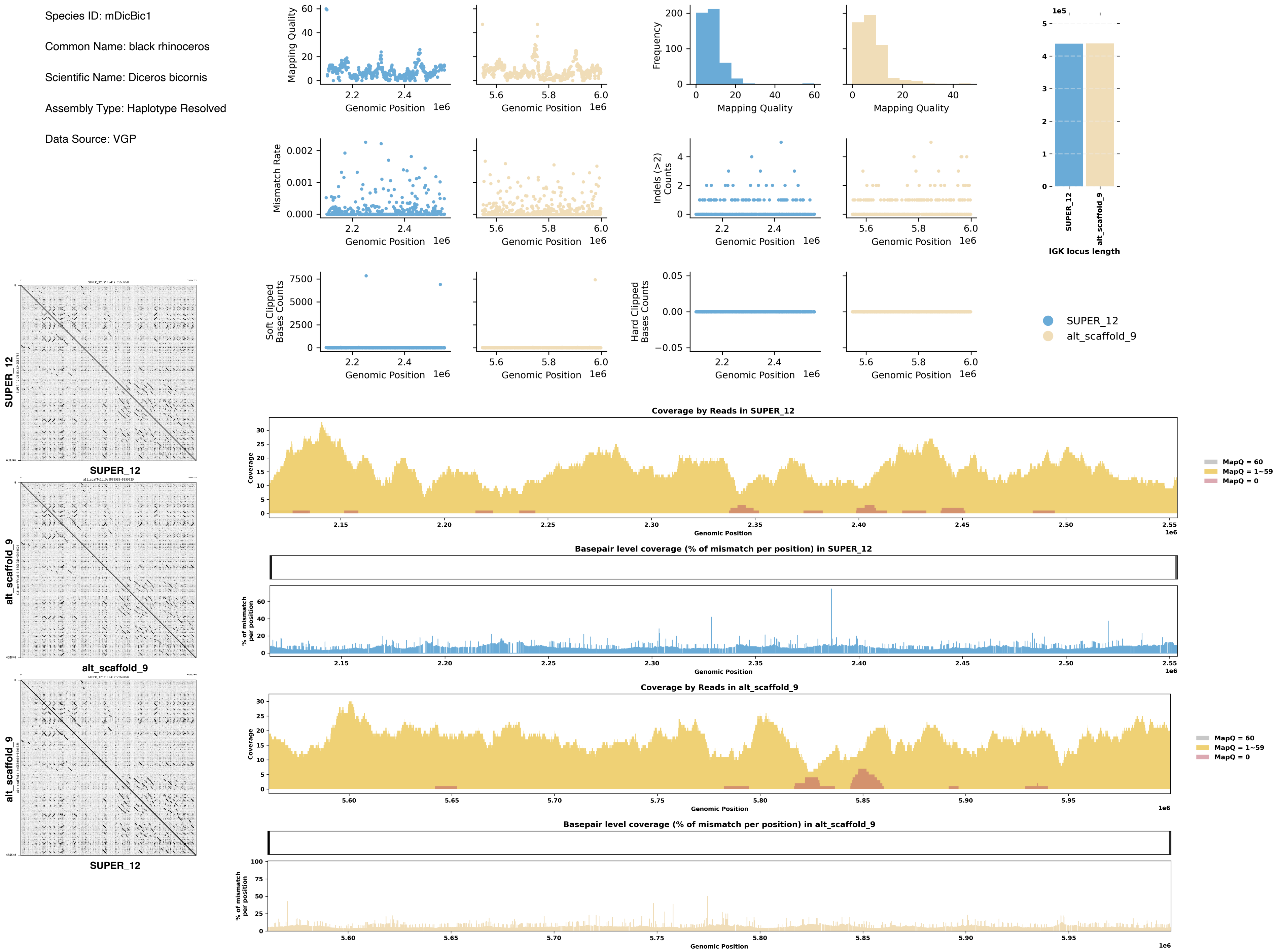

Species ID: mDipMer1

Common Name: Merriam Kangaroo Rat

Scientific Name: Dipodomys merriami

Assembly Type: Not Haplotype Resolved

Data Source: CCGP

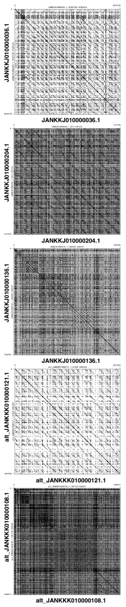


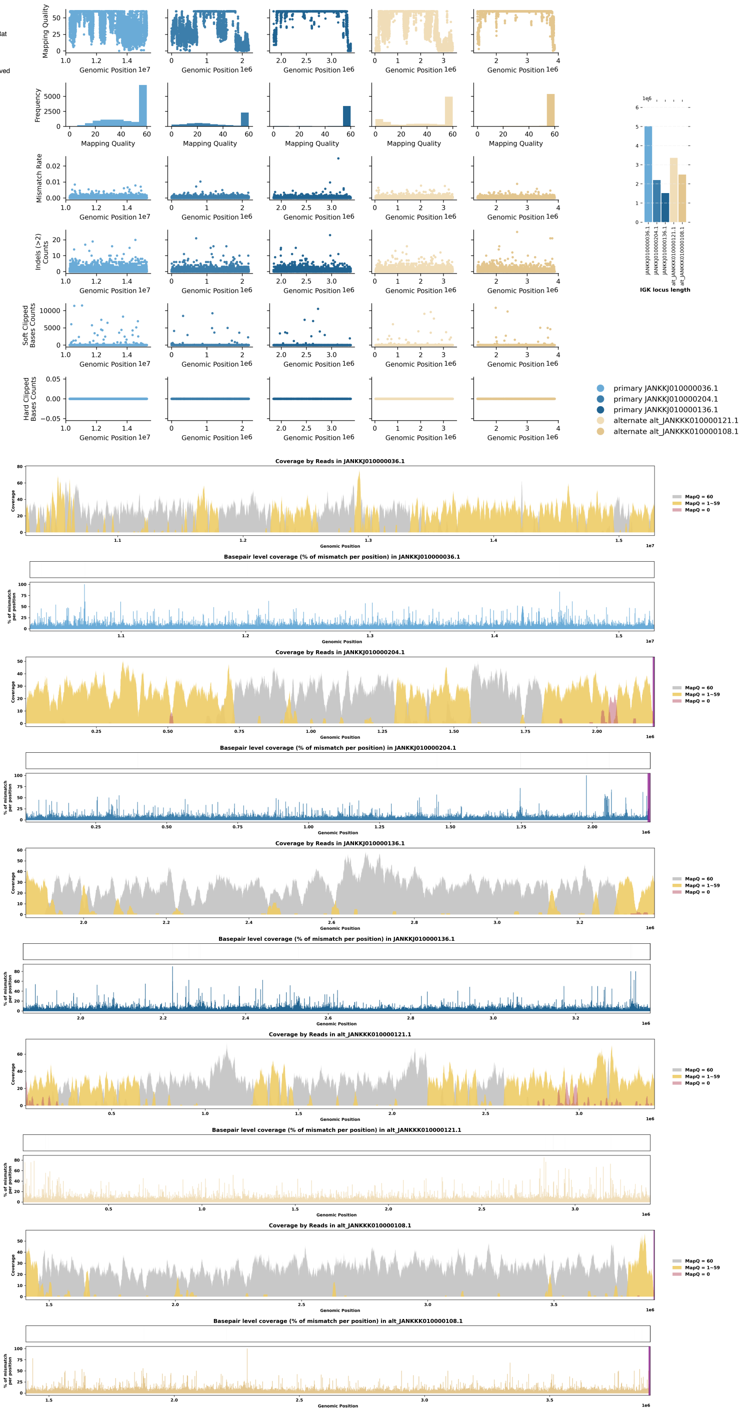

Species ID: mEleMax1

Common Name: Asiatic Elephant

Scientific Name: Elephas maximus

Assembly Type: Not Haplotype Resolved

Data Source: VGP

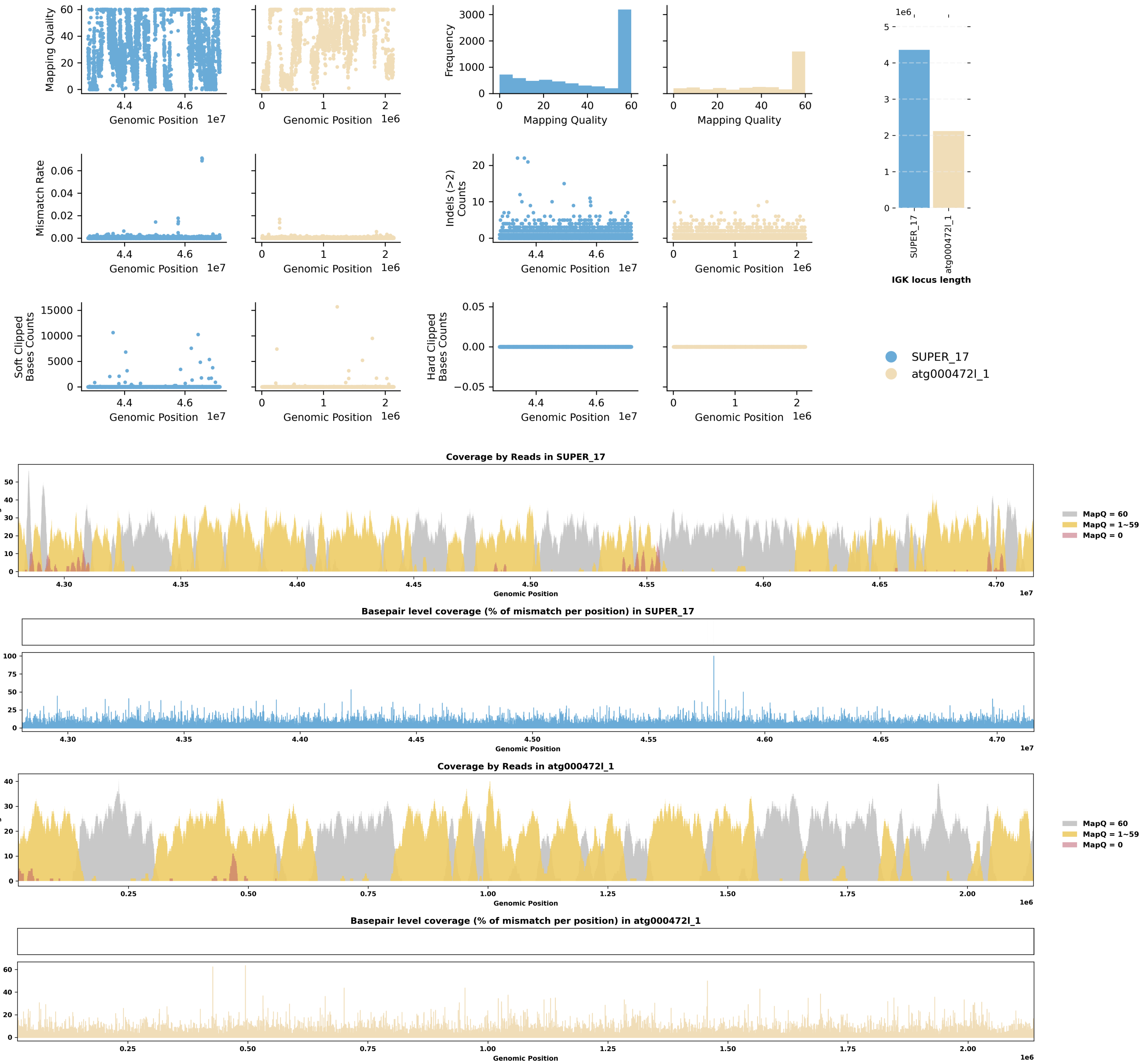


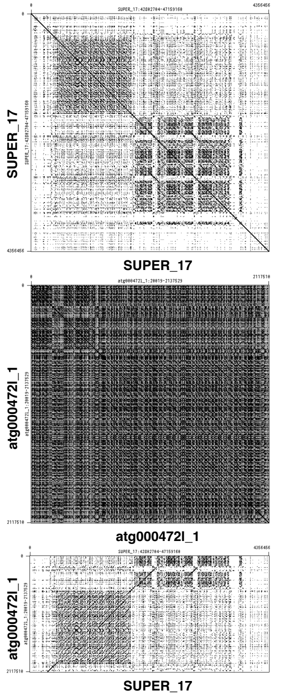

Species ID: mEriEur2

Common Name: western European hedgehog

Scientific Name: Erinaceus europaeus

Assembly Type: Not Haplotype Resolved

Data Source: VGP

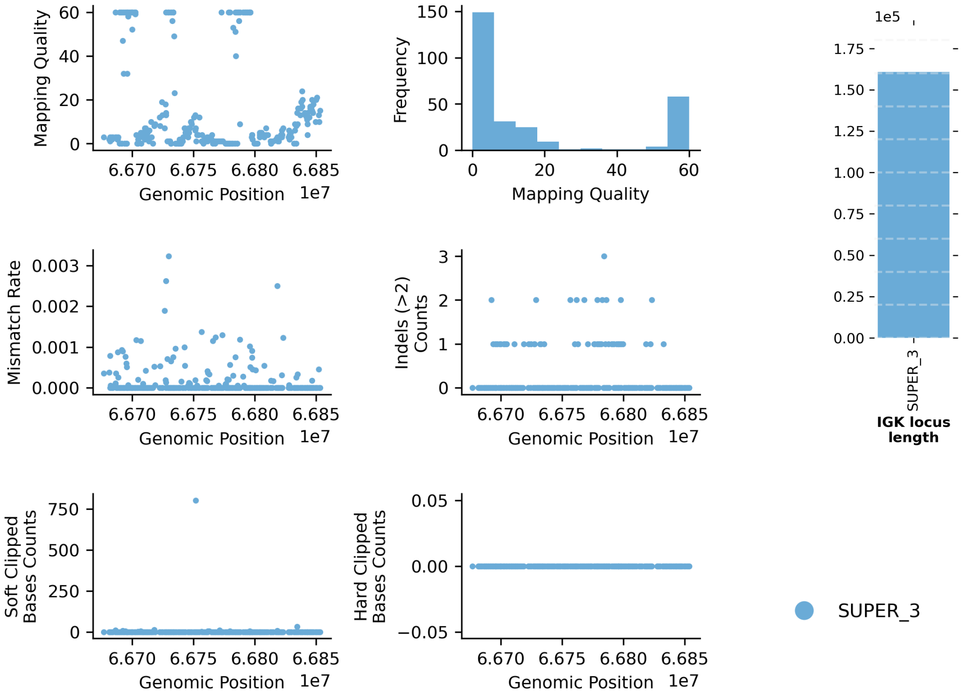


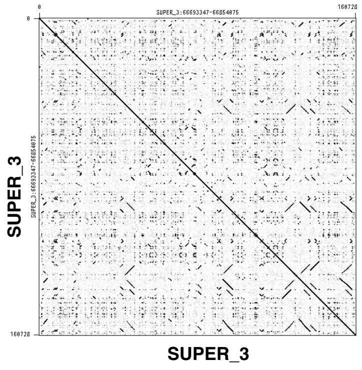


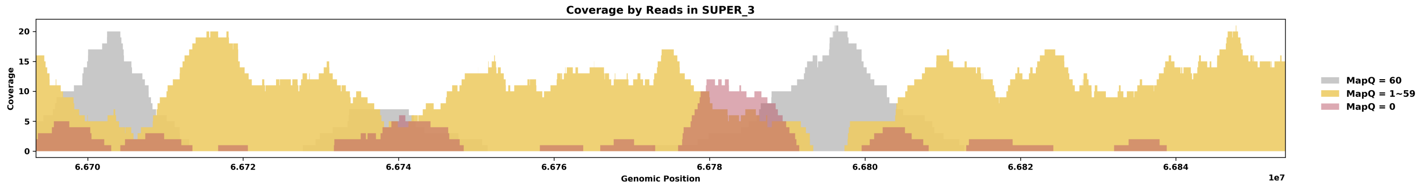


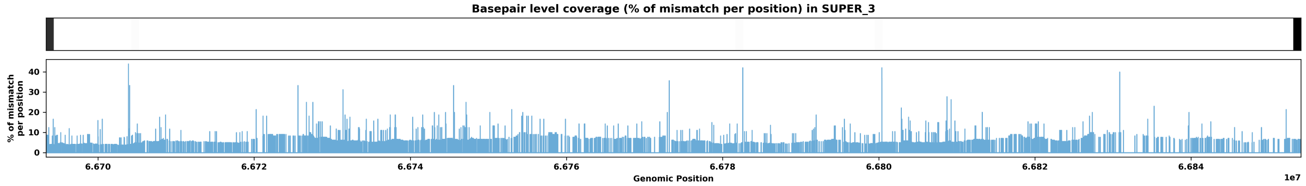

Species ID: mEscRob2

Common Name: grey whale

Scientific Name: Eschrichtius robustus

Assembly Type: Not Haplotype Resolved

Data Source: VGP

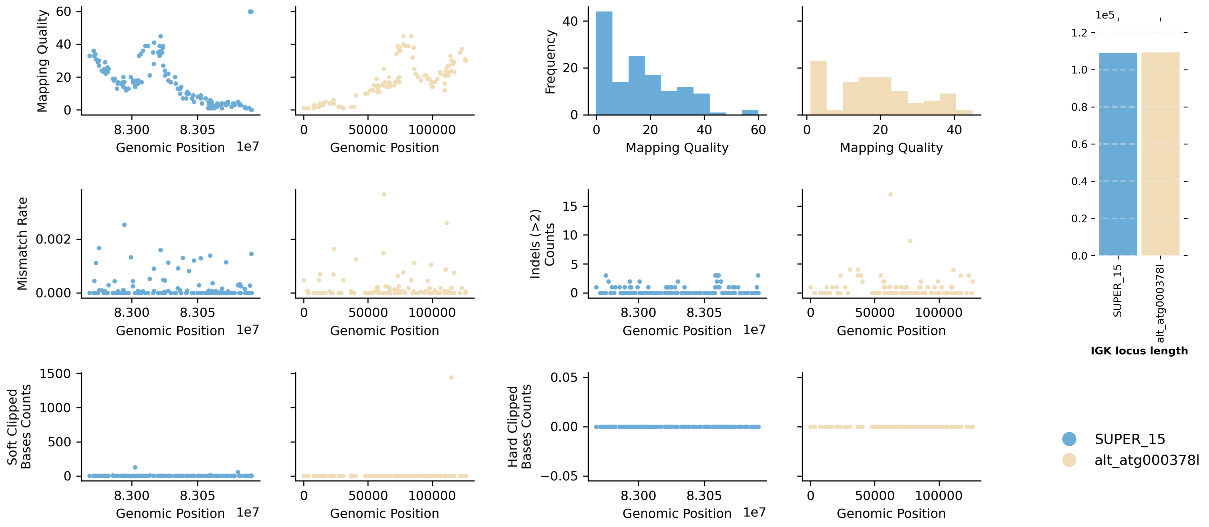


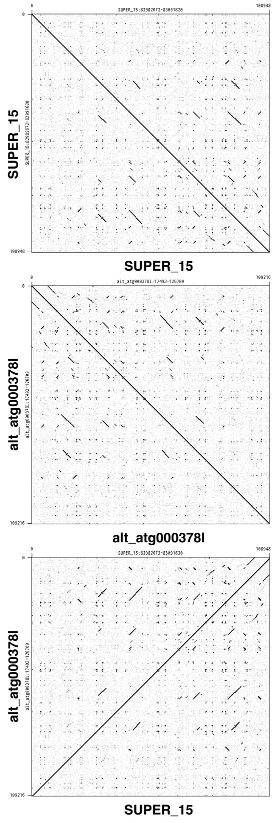


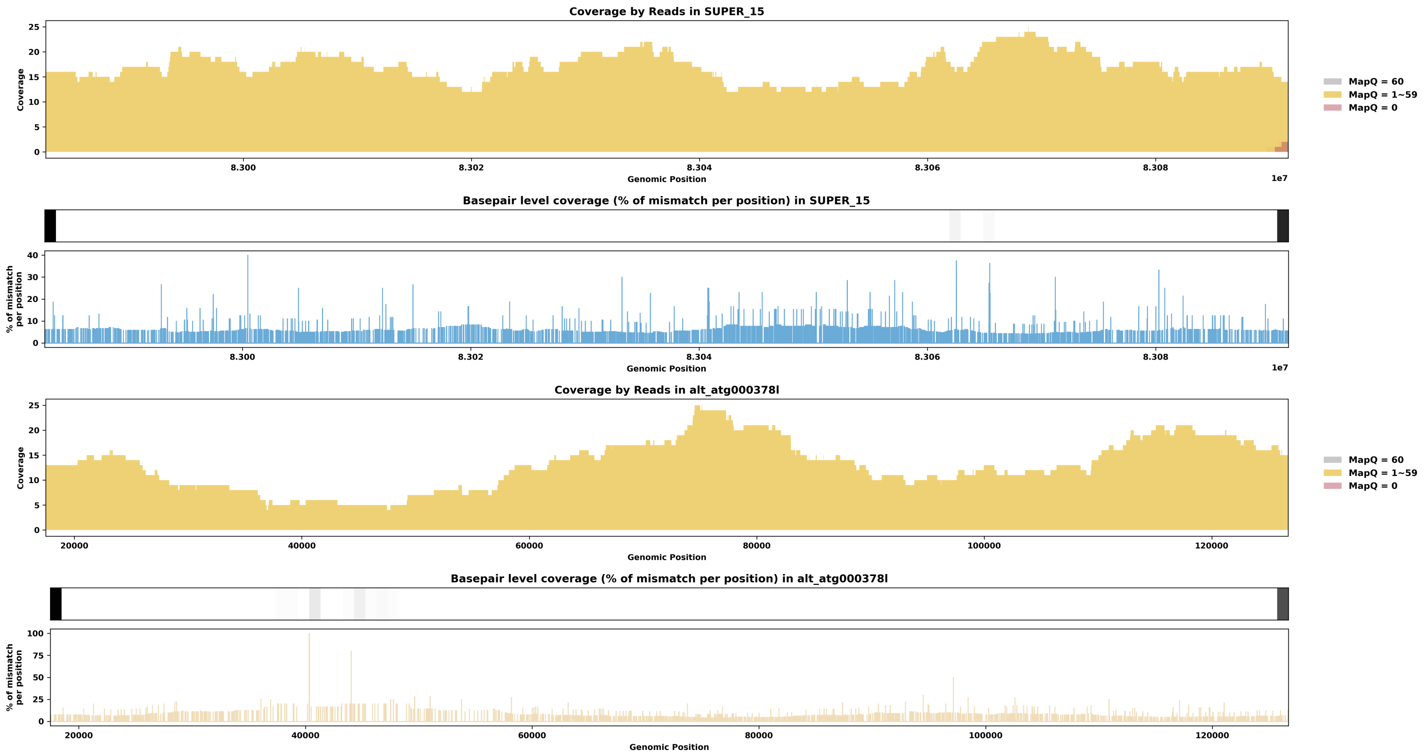

Species ID: mEubGla1

Common Name: North Atlantic right whale

Scientific Name: Eubalaena glacialis

Assembly Type: Haplotype Resolved

Data Source: VGP

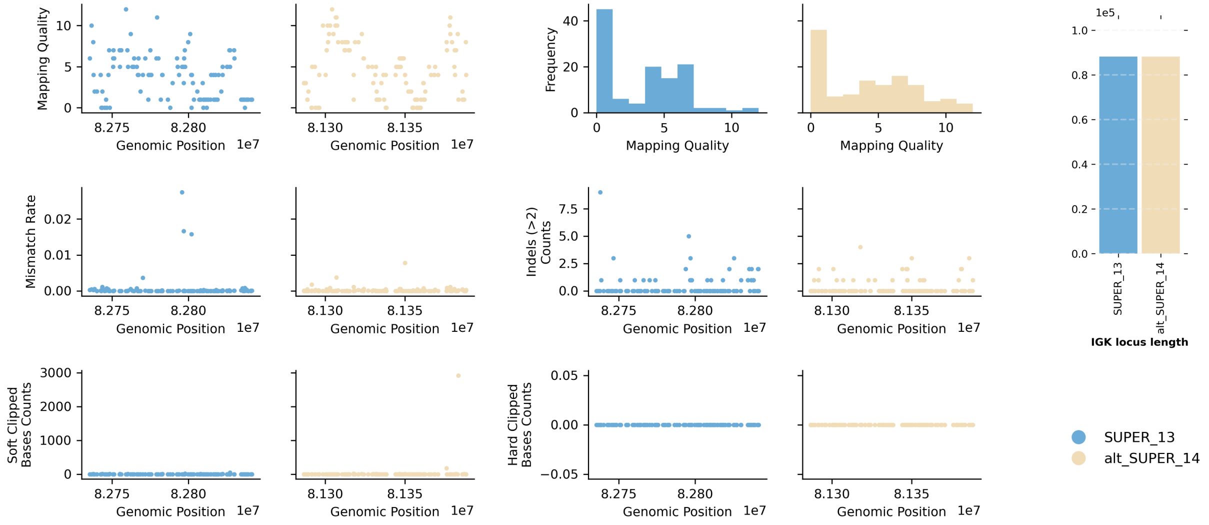


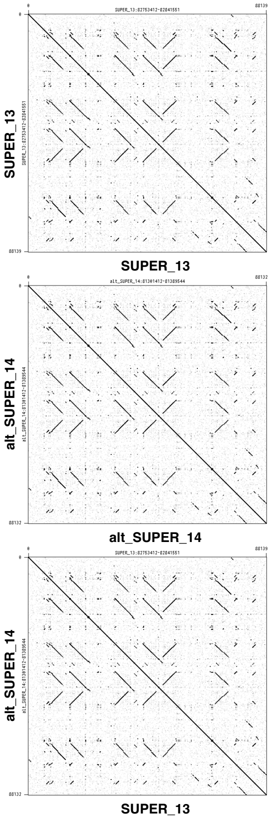


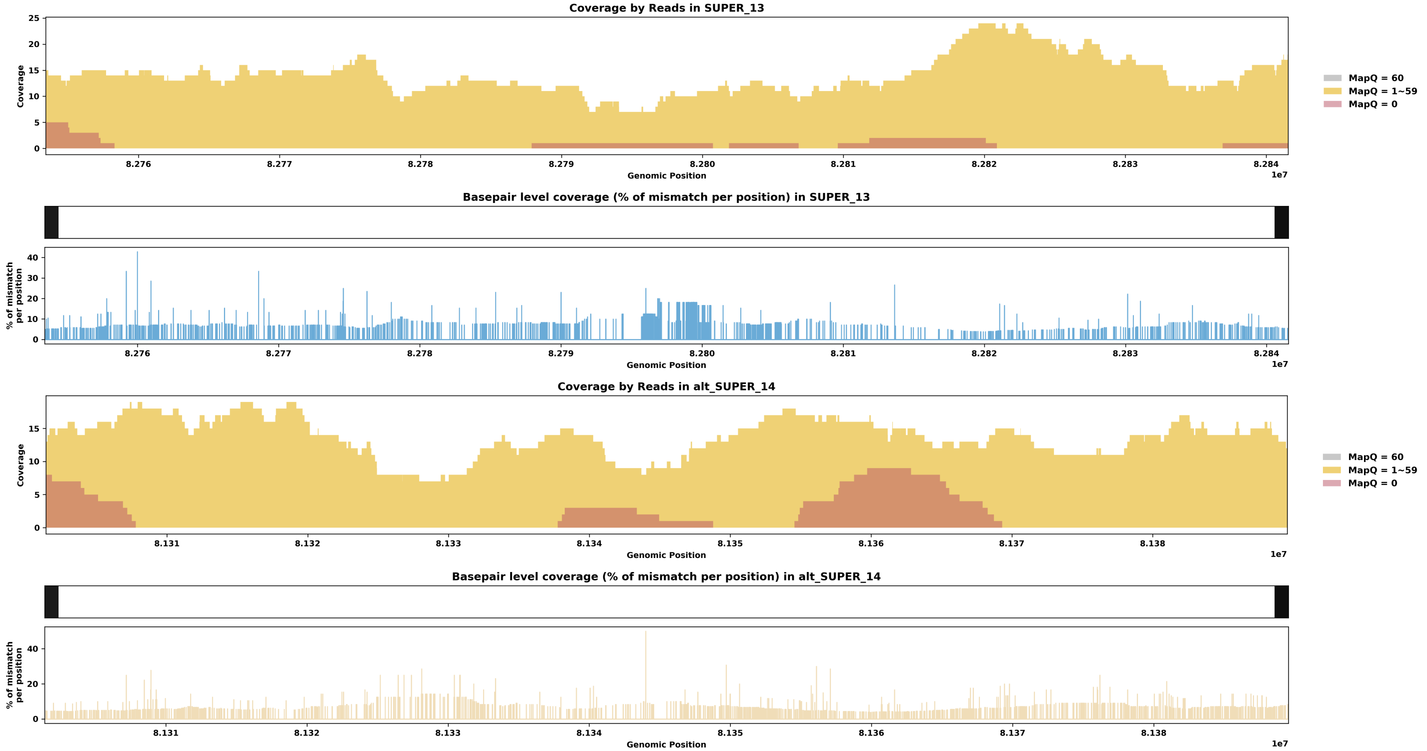

Species ID: mGloMel1

Common Name: long-finned pilot whale

Scientific Name: Globicephala melas

Assembly Type: Not Haplotype Resolved

Data Source: VGP

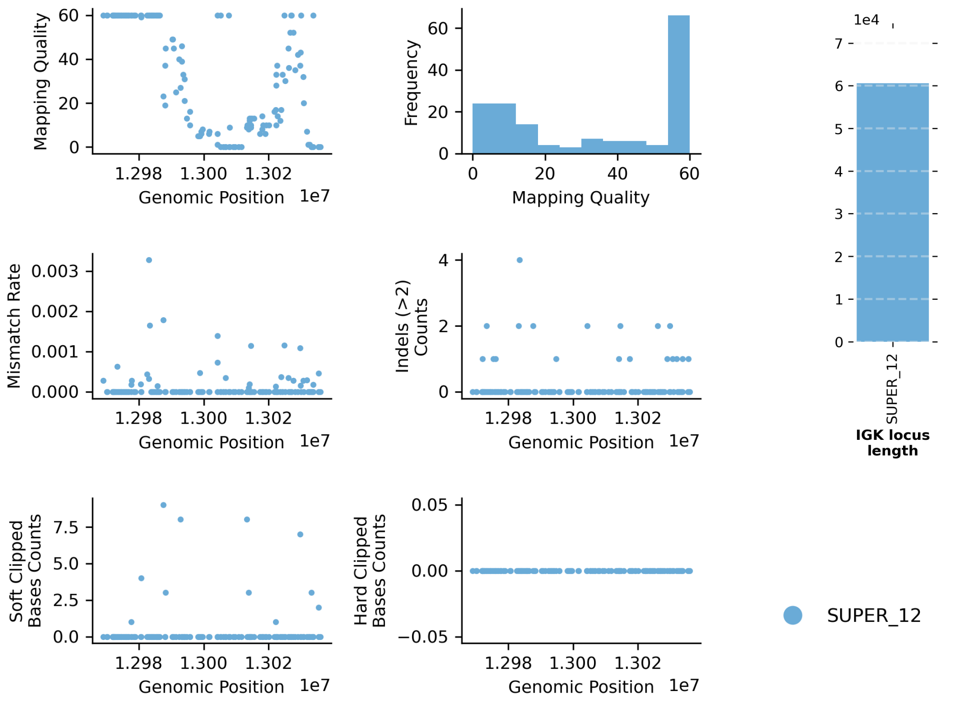


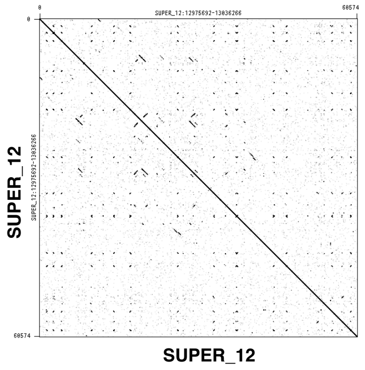


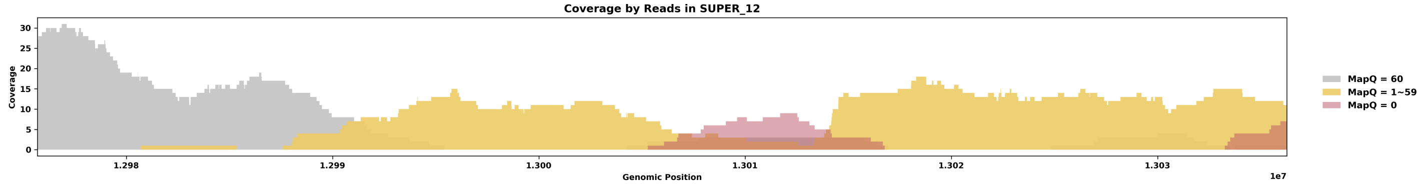


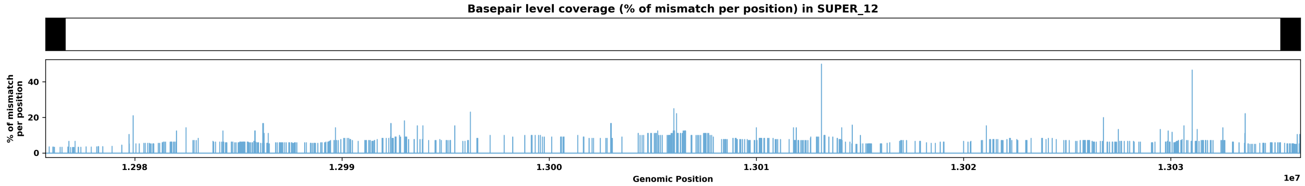

Species ID: mGorGor1

Common Name: Gorilla

Scientific Name: Gorilla_gorilla

Assembly Type: Haplotype Resolved

Data Source: T2T Primate

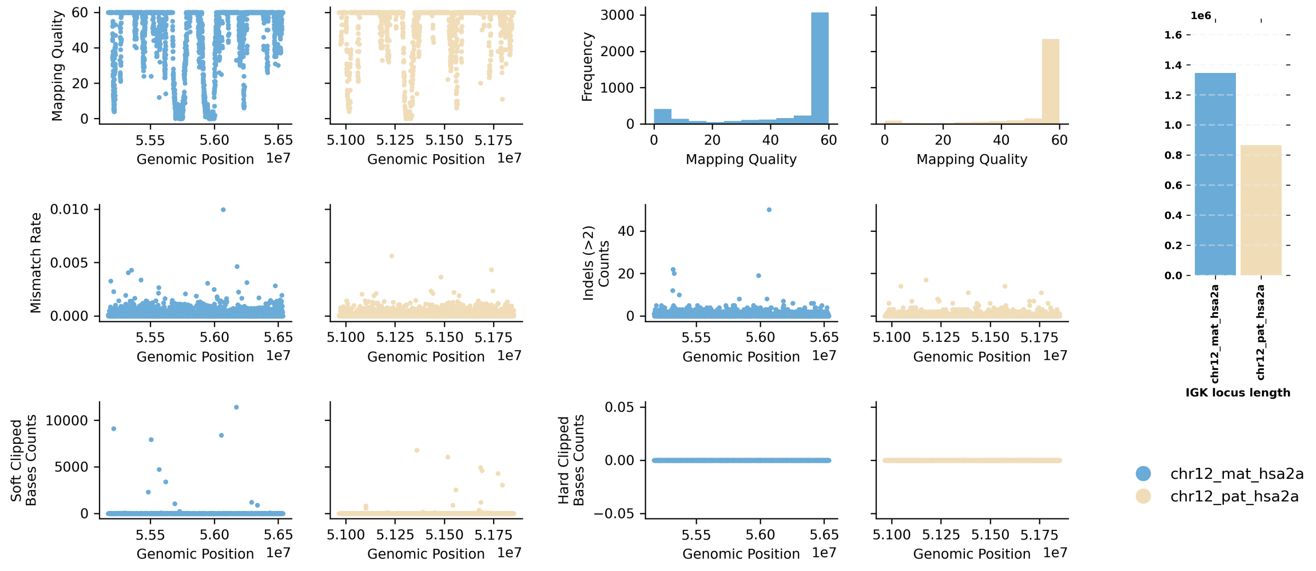


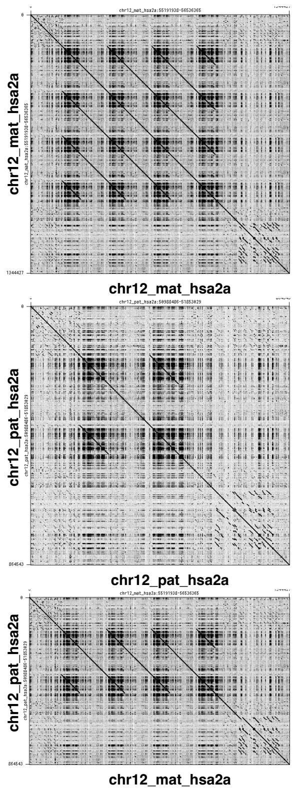


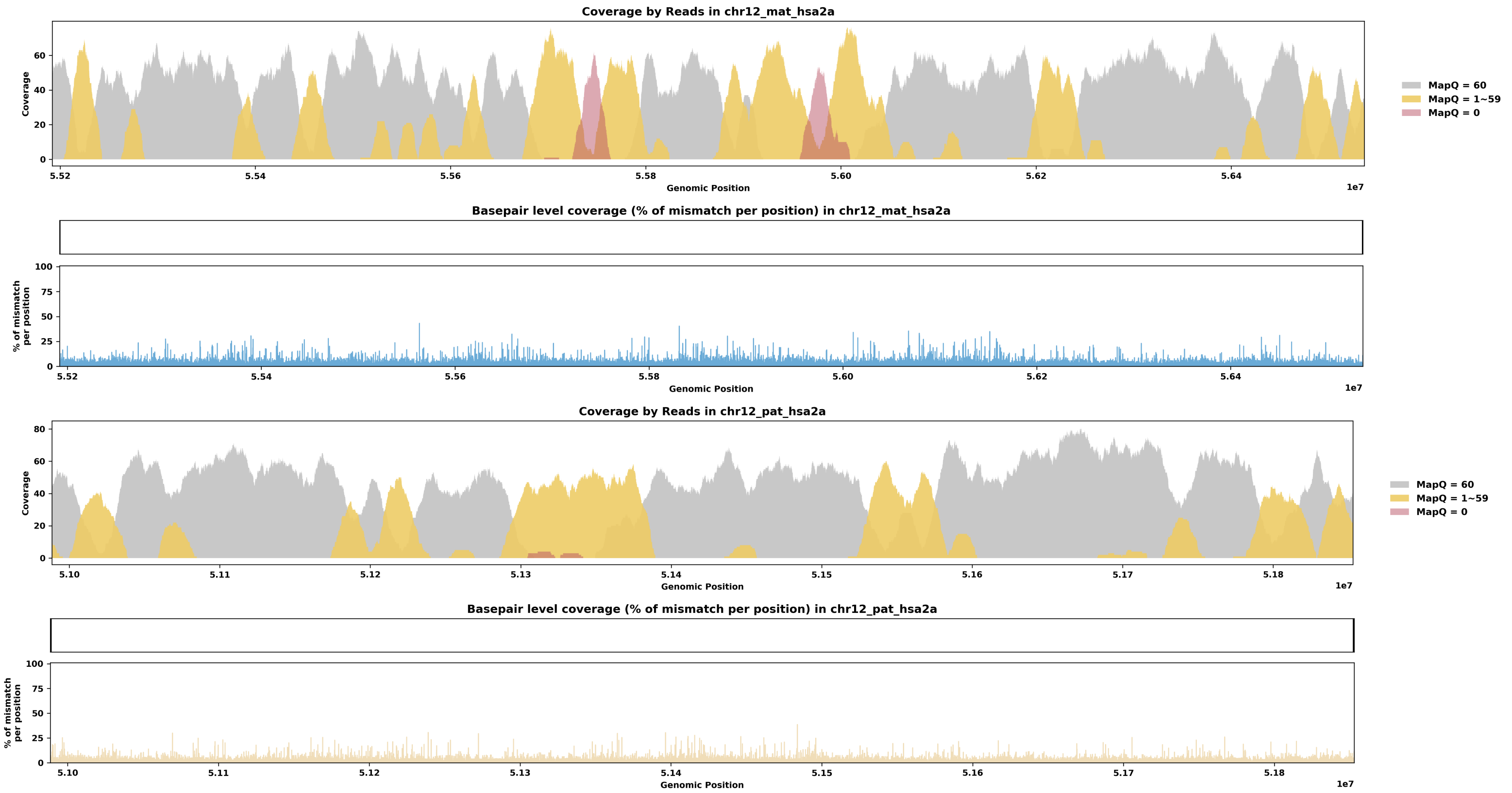

Species ID: mHetBru1

Common Name: Yellow-spotted hyrax

Scientific Name: Heterohyrax brucei

Assembly Type: Not Haplotype Resolved

Data Source: VGP

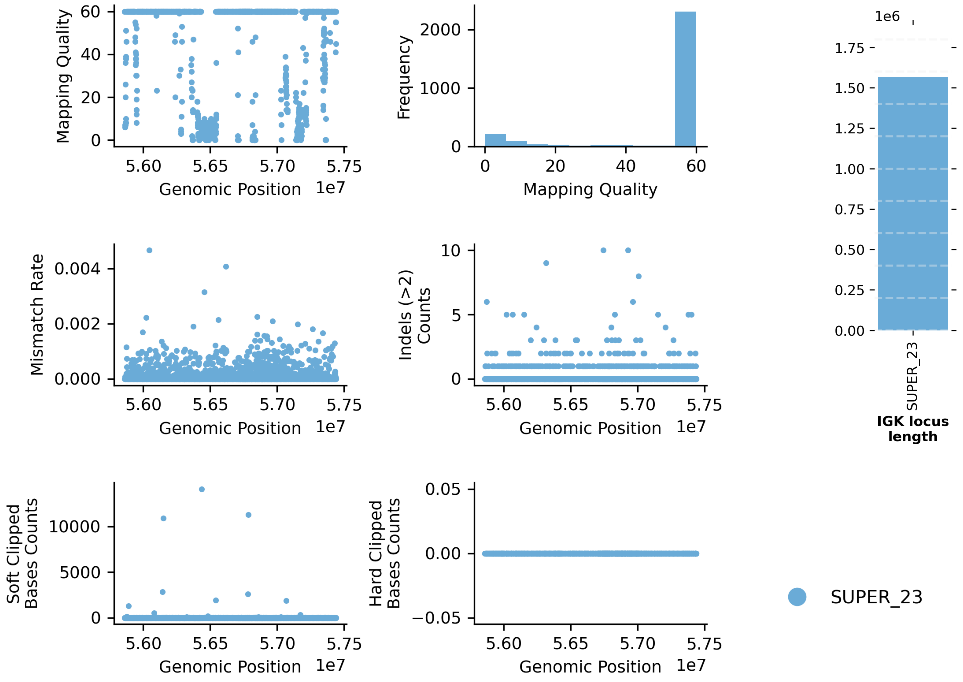


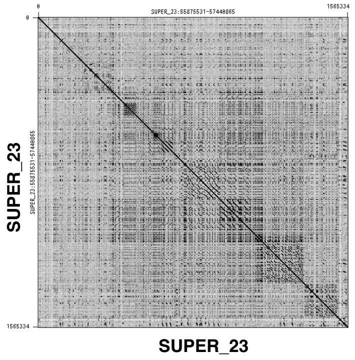


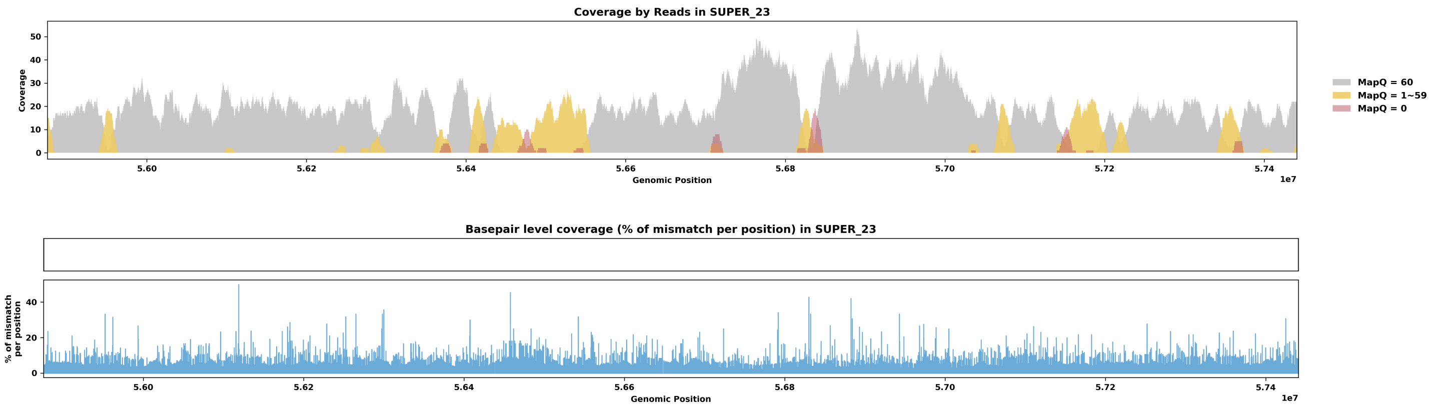

Species ID: mHipAmp2

Common Name: hippopotamus

Scientific Name: Hippopotamus amphibius kiboko

Assembly Type: Haplotype Resolved

Data Source: VGP

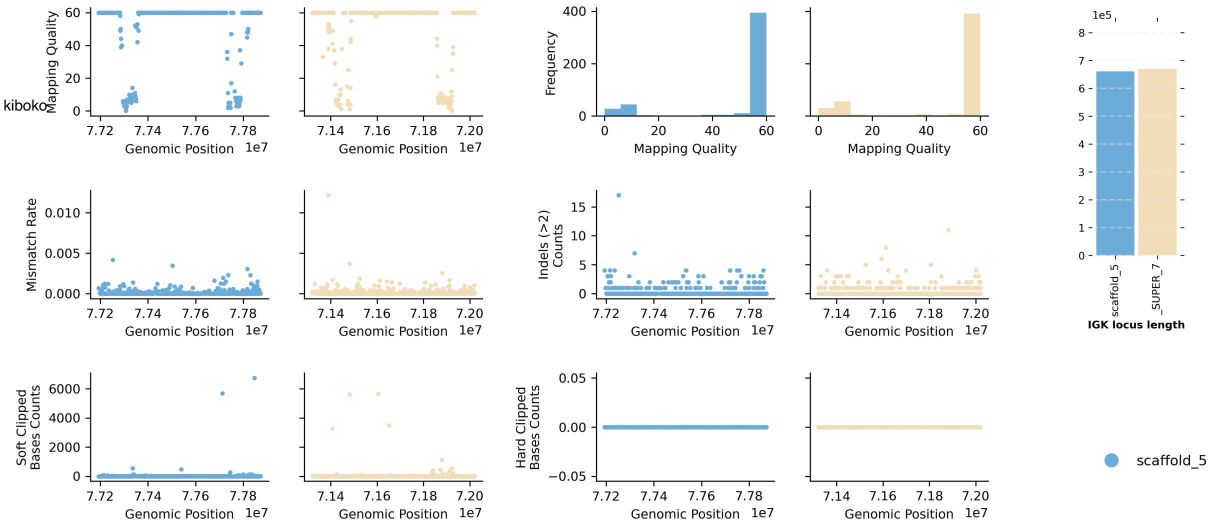


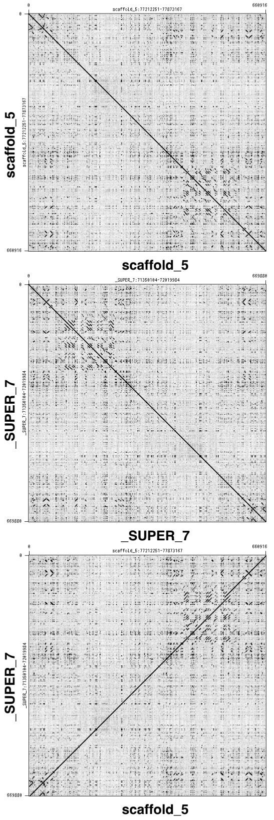


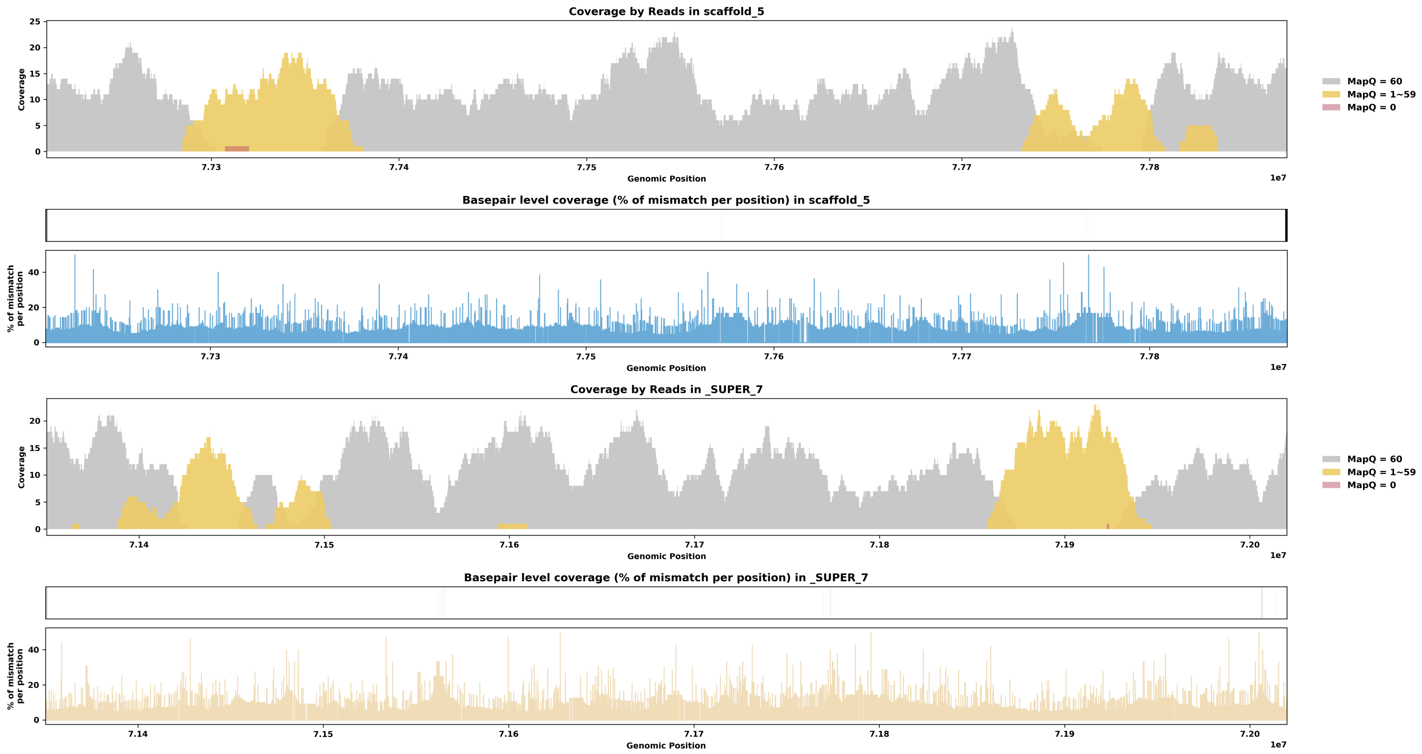

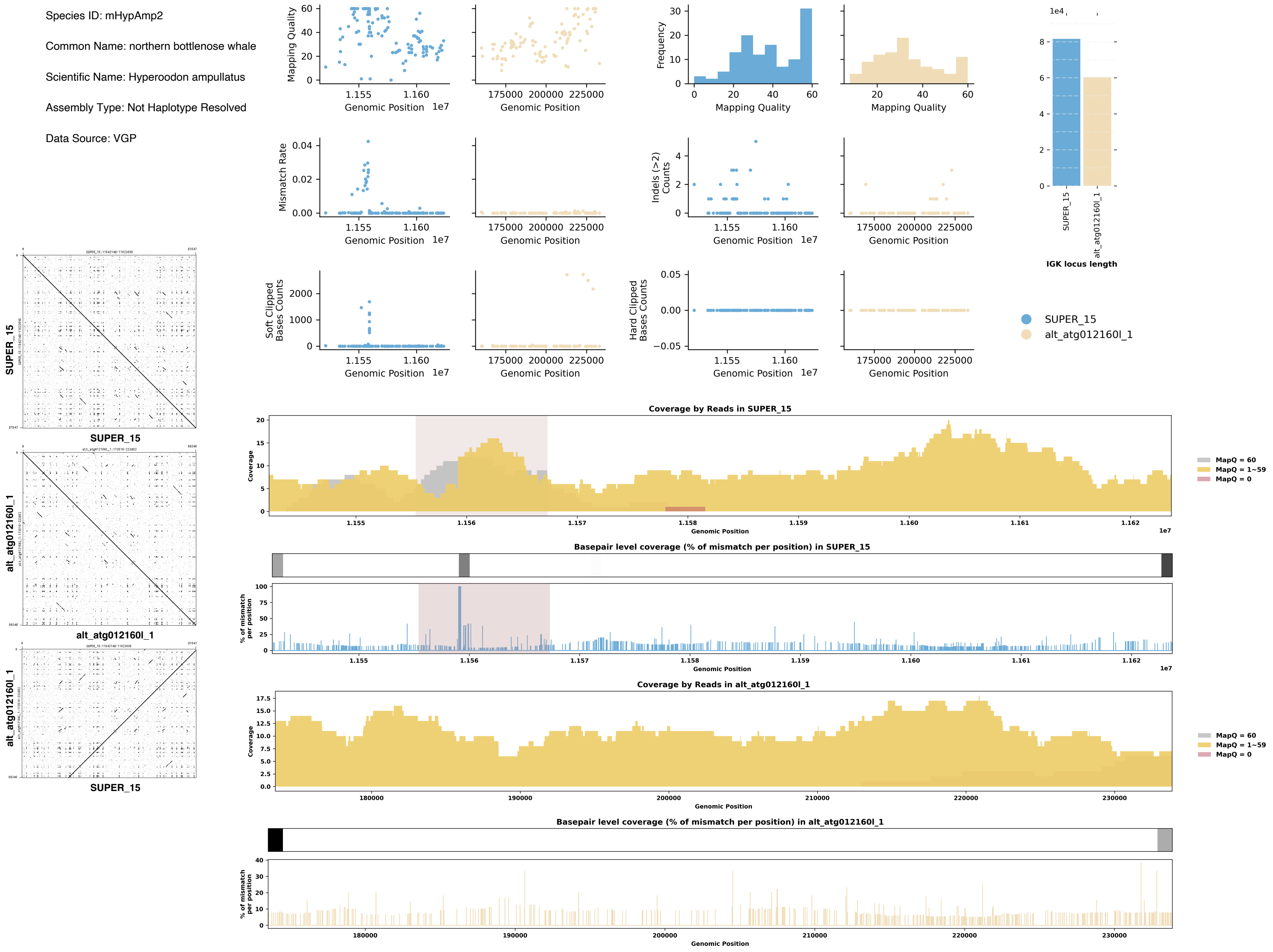
Species ID: mHypAmp2
Common Name: northern bottlenose whale
Scientific Name: Hyperoodon ampullatus
Assembly Type: Not Haplotype Resolved
Data Source: VGP
Mapping Quality
Genomic Position
Frequency
Mismatch Rate
Indels (>2) Counts
Soft Clipped Bases Counts
Hard Clipped Bases Counts
IGK locus length
SUPER_15
alt_atg012160l_1
Coverage by Reads in SUPER_15
Coverage
MapQ = 60
MapQ = 1~59
MapQ = 0
Basepair level coverage (% of mismatch per position) in SUPER_15
% of mismatch per position
Coverage by Reads in alt_atg012160l_1
Basepair level coverage (% of mismatch per position) in alt_atg012160l_1

Species ID: mLagAlb1

Common Name: white-beaked dolphin

Scientific Name: Lagenorhynchus albirostris

Assembly Type: Not Haplotype Resolved

Data Source: VGP

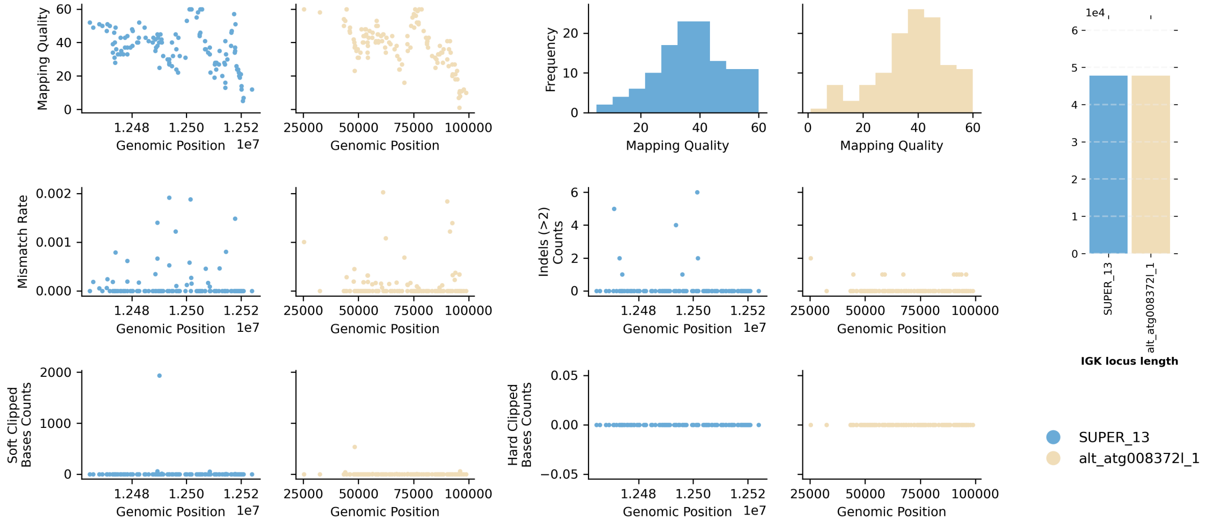


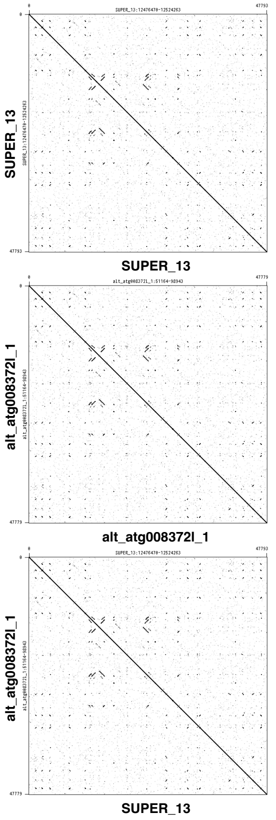


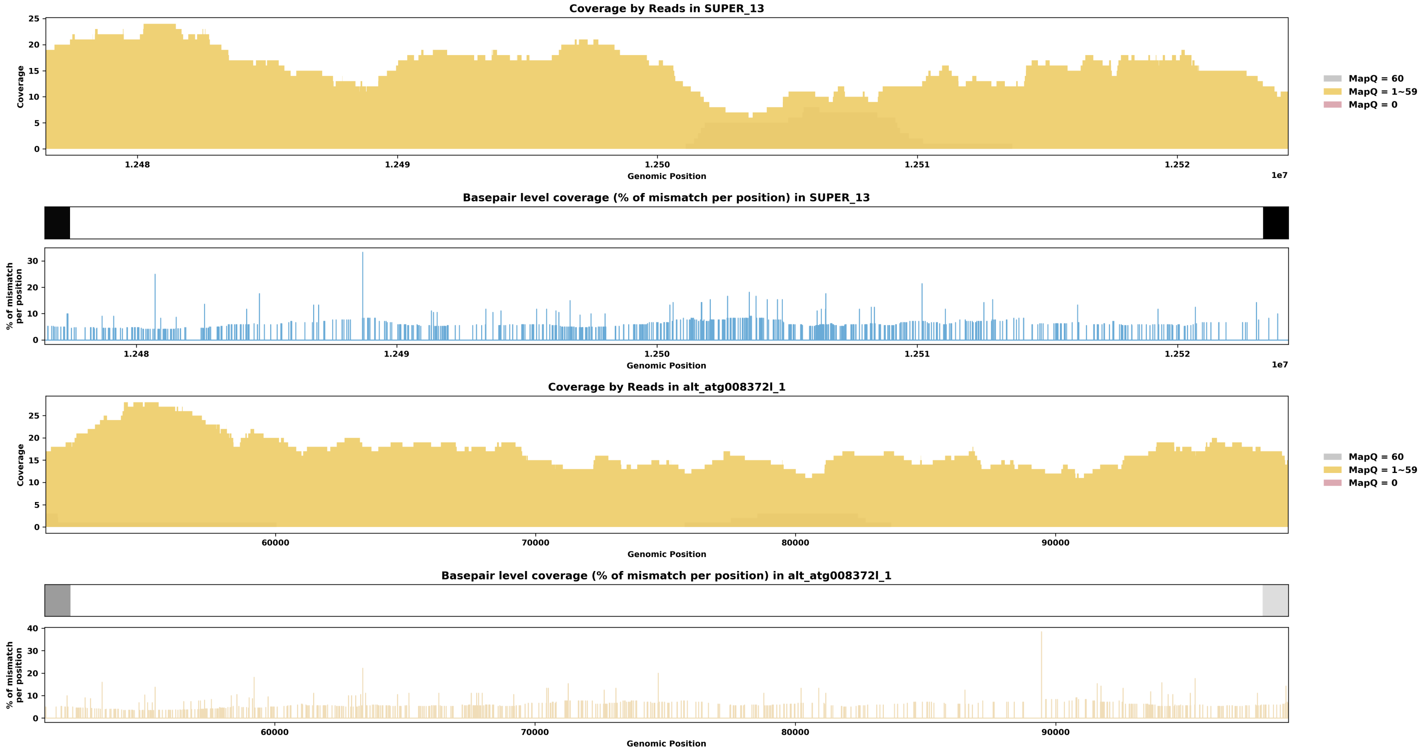

Species ID: mLemCat1

Common Name: Ring-tailed lemur

Scientific Name: Lemur catta

Assembly Type: Not Haplotype Resolved

Data Source: VGP

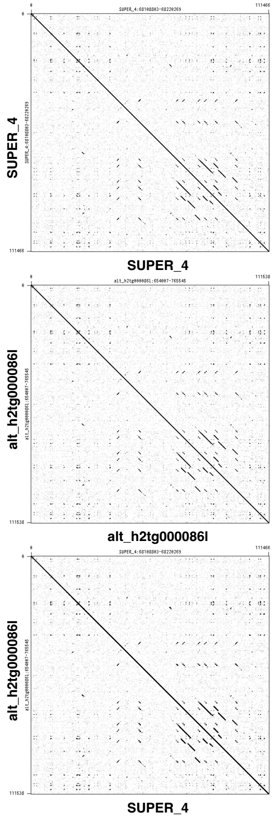


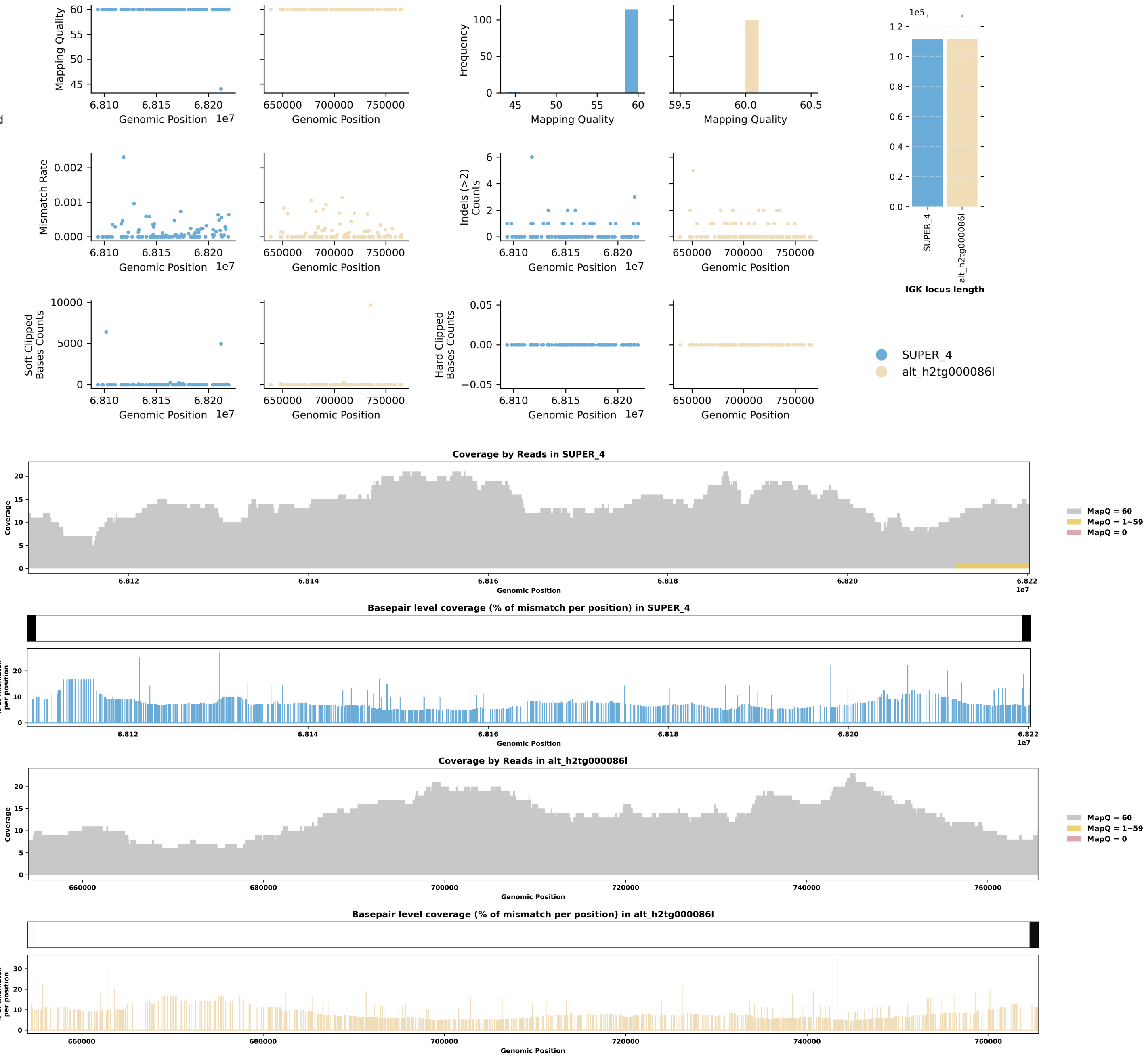

Species ID: mLynRuf1

Common Name: Bobcat

Scientific Name: Lynx rufus

Assembly Type: Not Haplotype Resolved

Data Source: CCGP

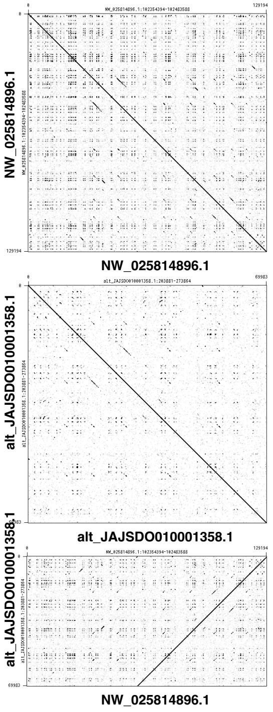


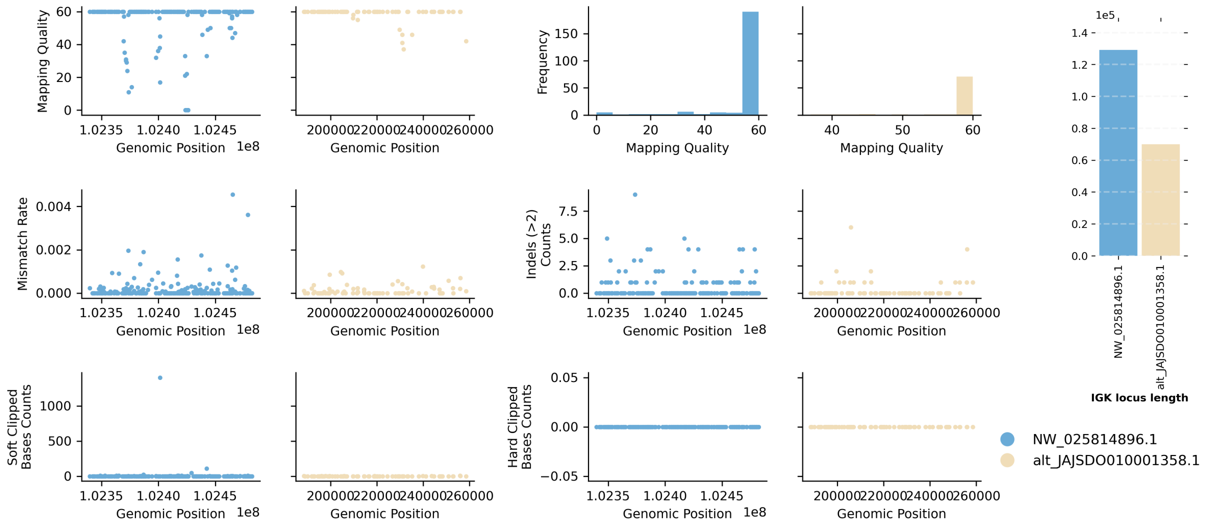


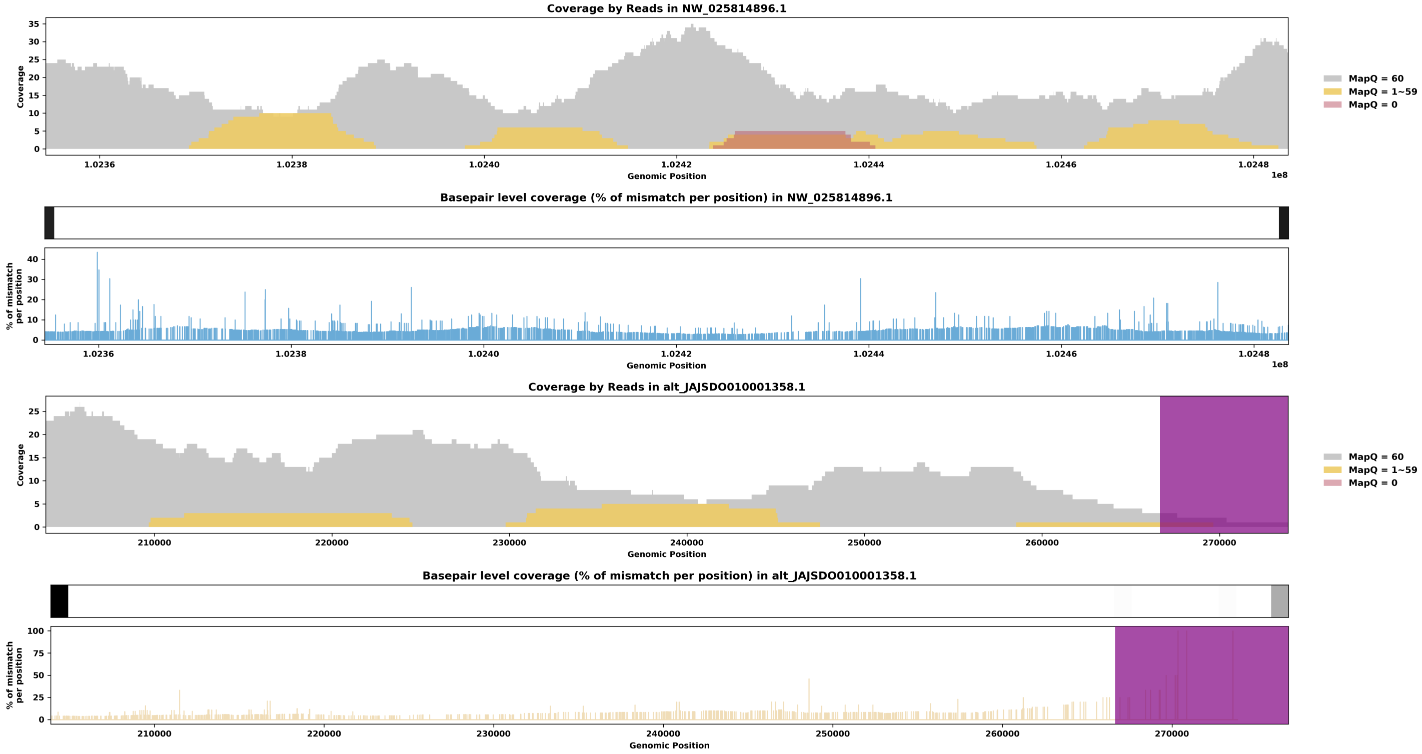

Species ID: mMacEug1

Common Name: tammar wallaby

Scientific Name: Macropus eugenii

Assembly Type: Not Haplotype Resolved

Data Source: VGP

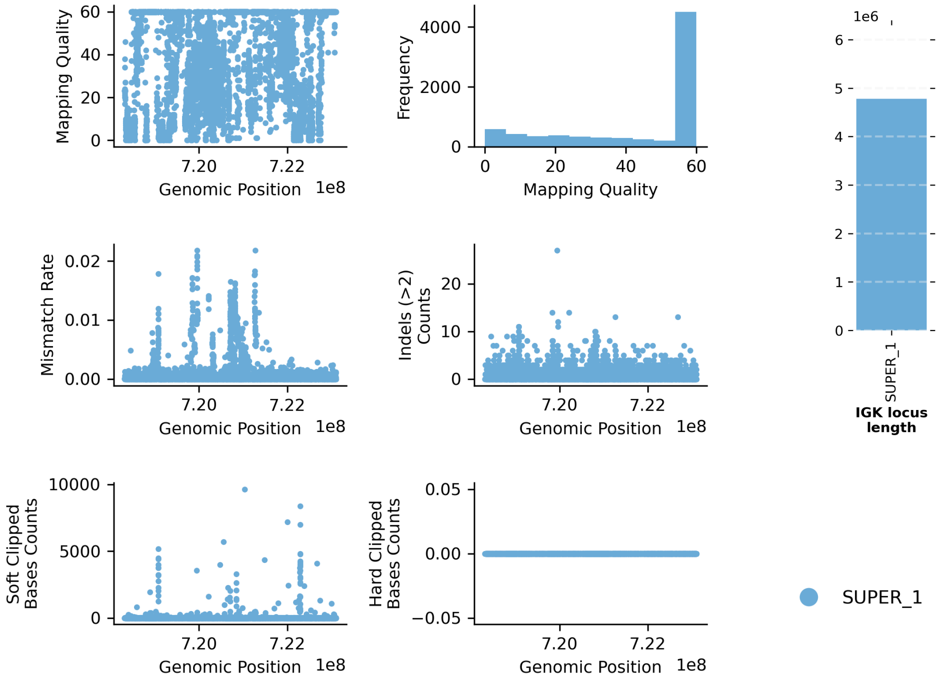


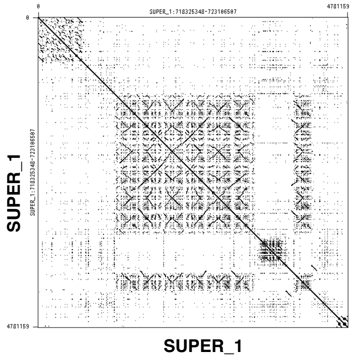


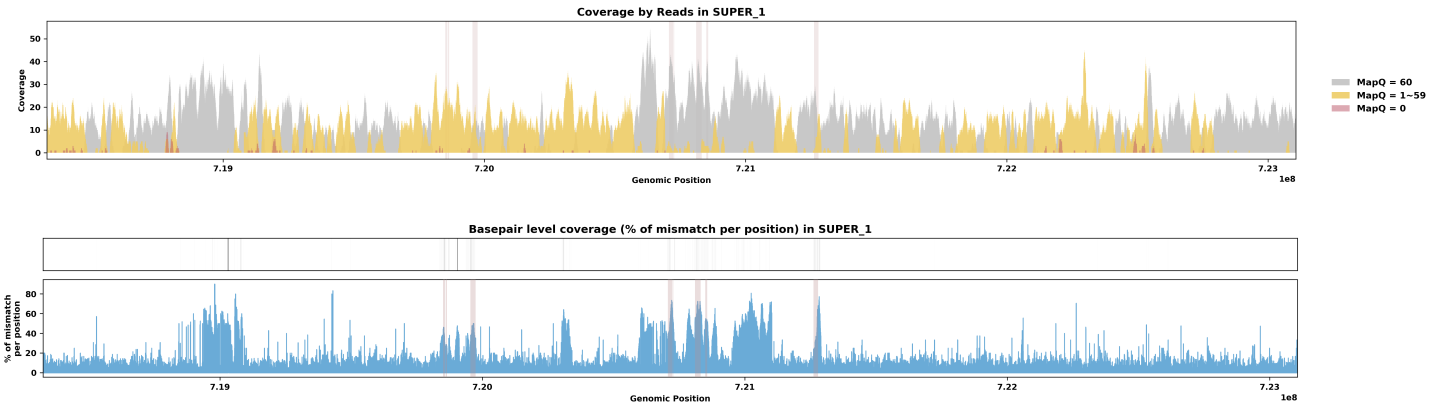

Species ID: mManPen7

Common Name: Chinese pangolin

Scientific Name: Manis pentadactyla

Assembly Type: Haplotype Resolved

Data Source: VGP

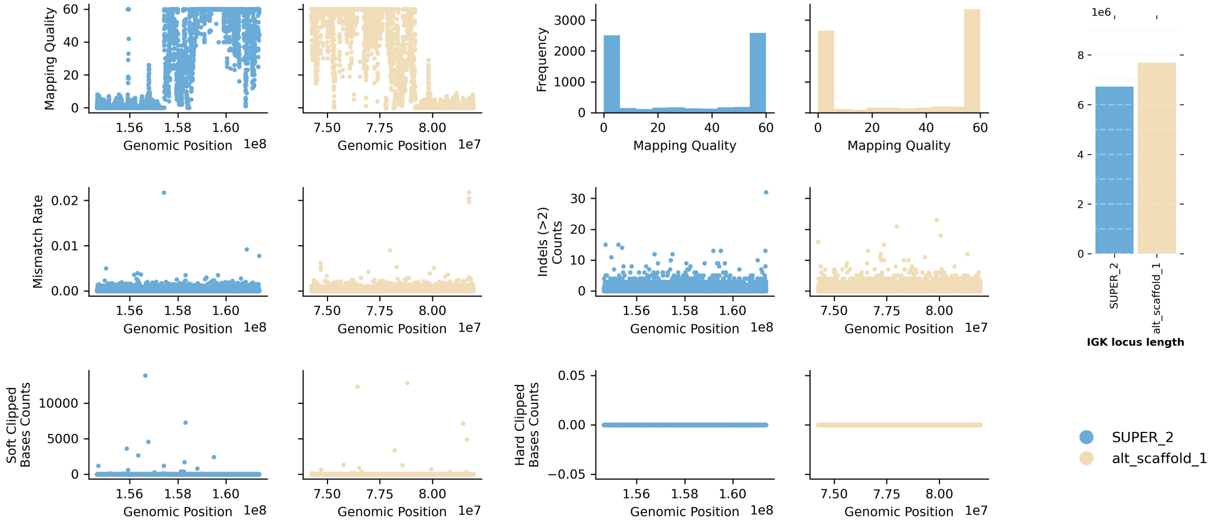


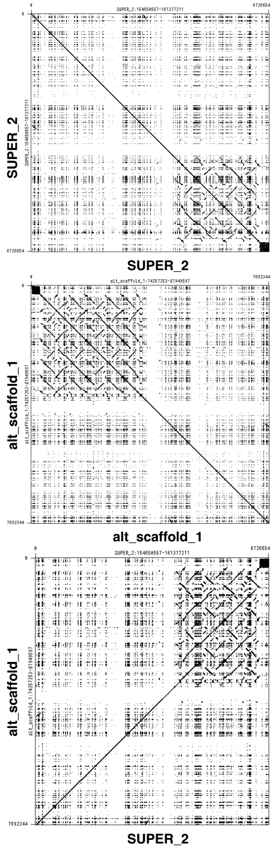


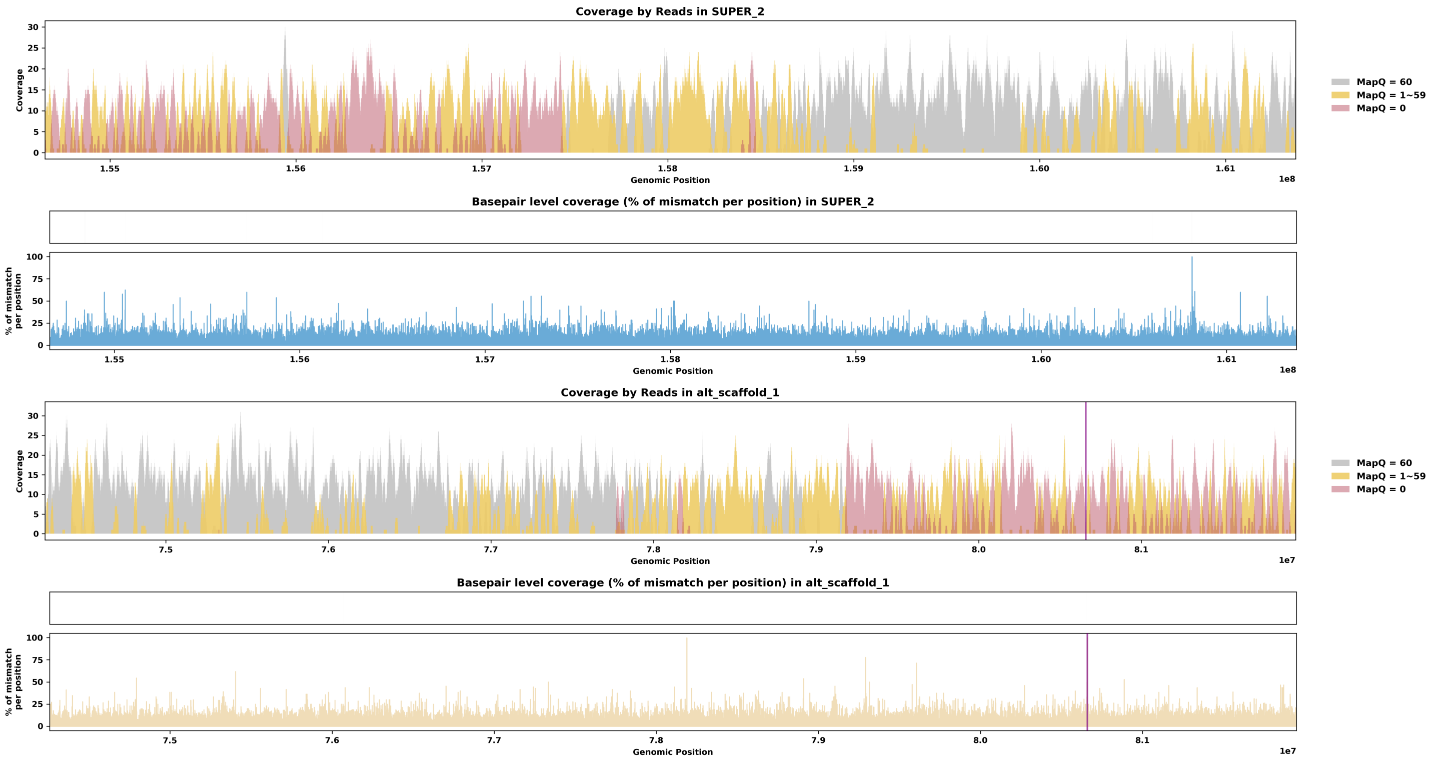

Species ID: mMarMar1

Common Name: European pine marten

Scientific Name: Martes martes

Assembly Type: Not Haplotype Resolved

Data Source: VGP

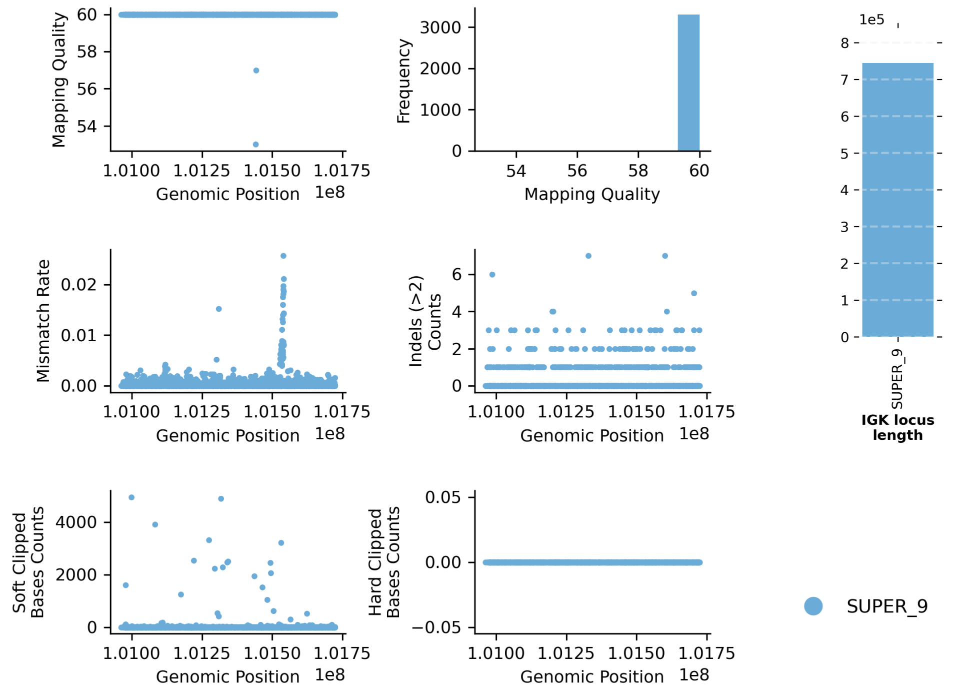


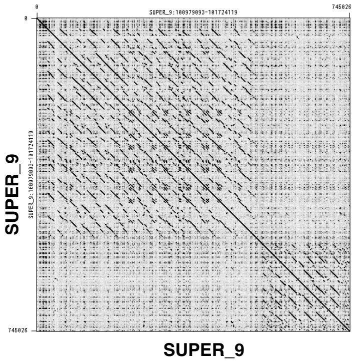


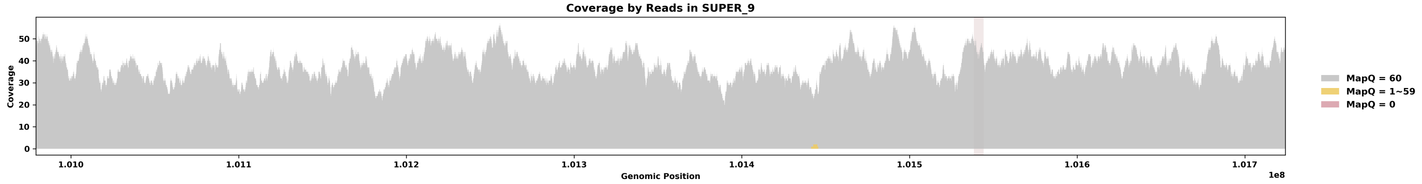


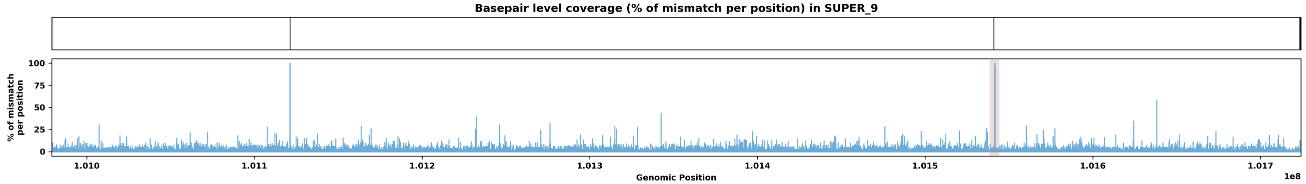

Species ID: mMelMel3

Common Name: European badger

Scientific Name: Meles meles

Assembly Type: Haplotype Resolved

Data Source: VGP

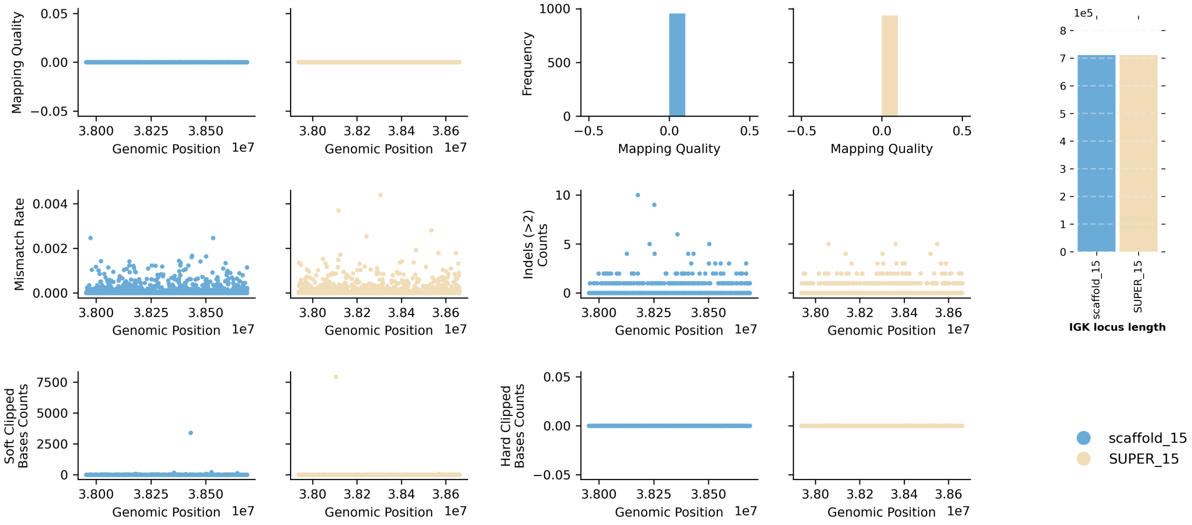


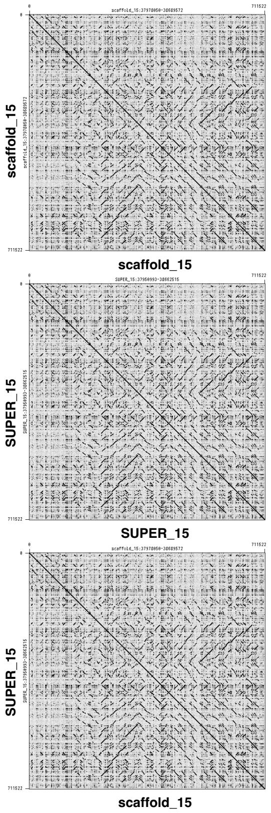


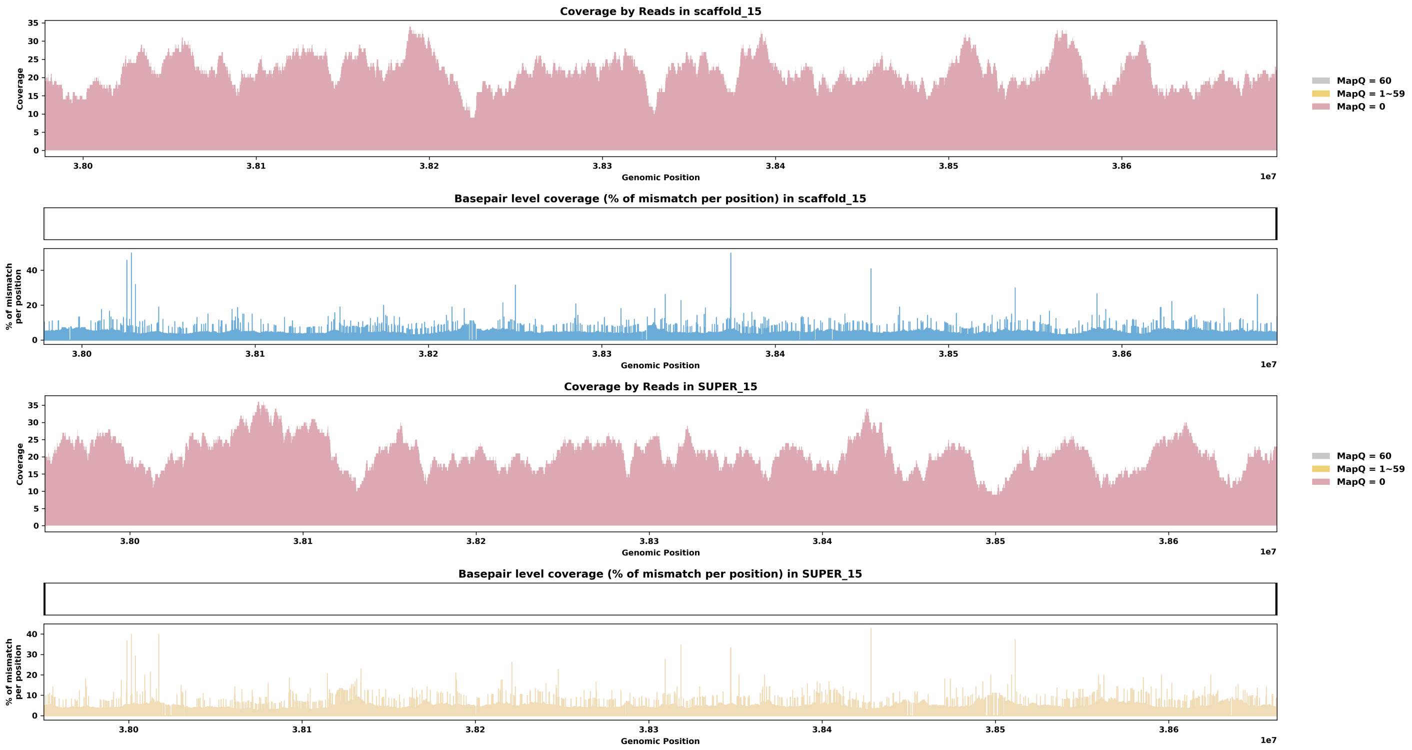

Species ID: mMesDen1

Common Name: Blainville's beaked whale

Scientific Name: Mesoplodon densirostris

Assembly Type: Not Haplotype Resolved

Data Source: VGP

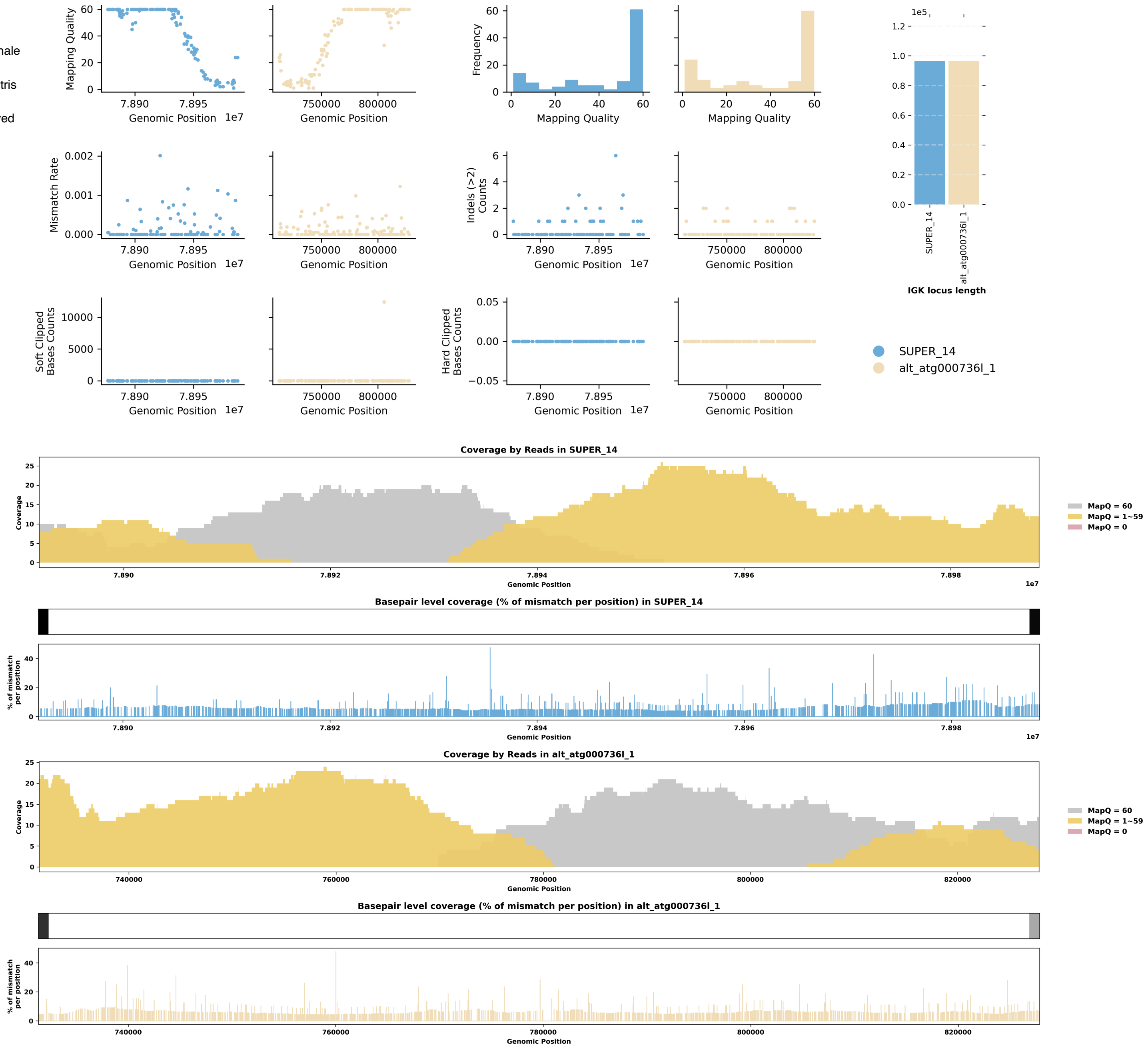


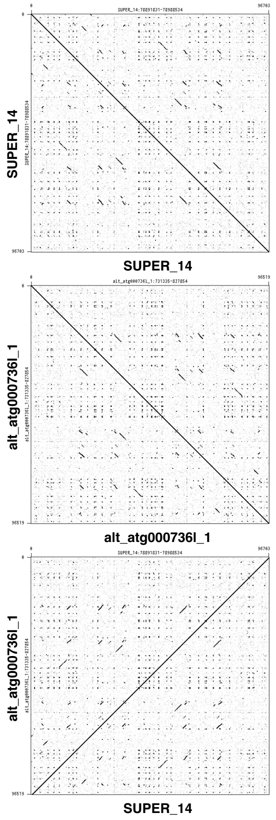

Species ID: mMicCal1

Common Name: California Vole

Scientific Name: Microtus californicus

Assembly Type: Haplotype Resolved

Data Source: CCGP

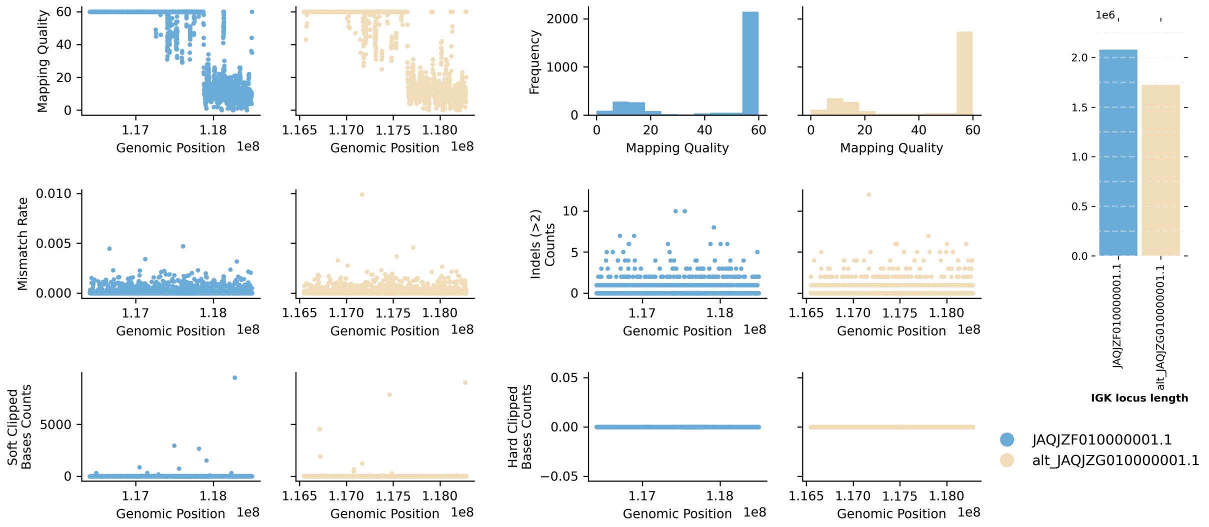


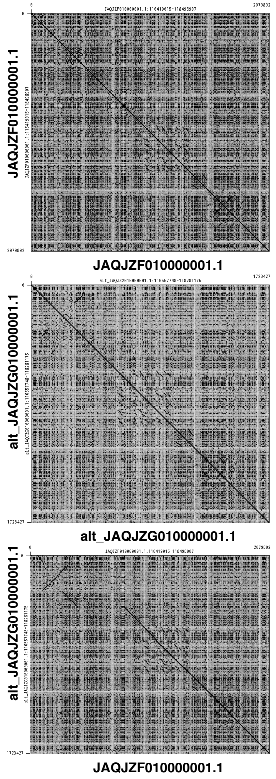


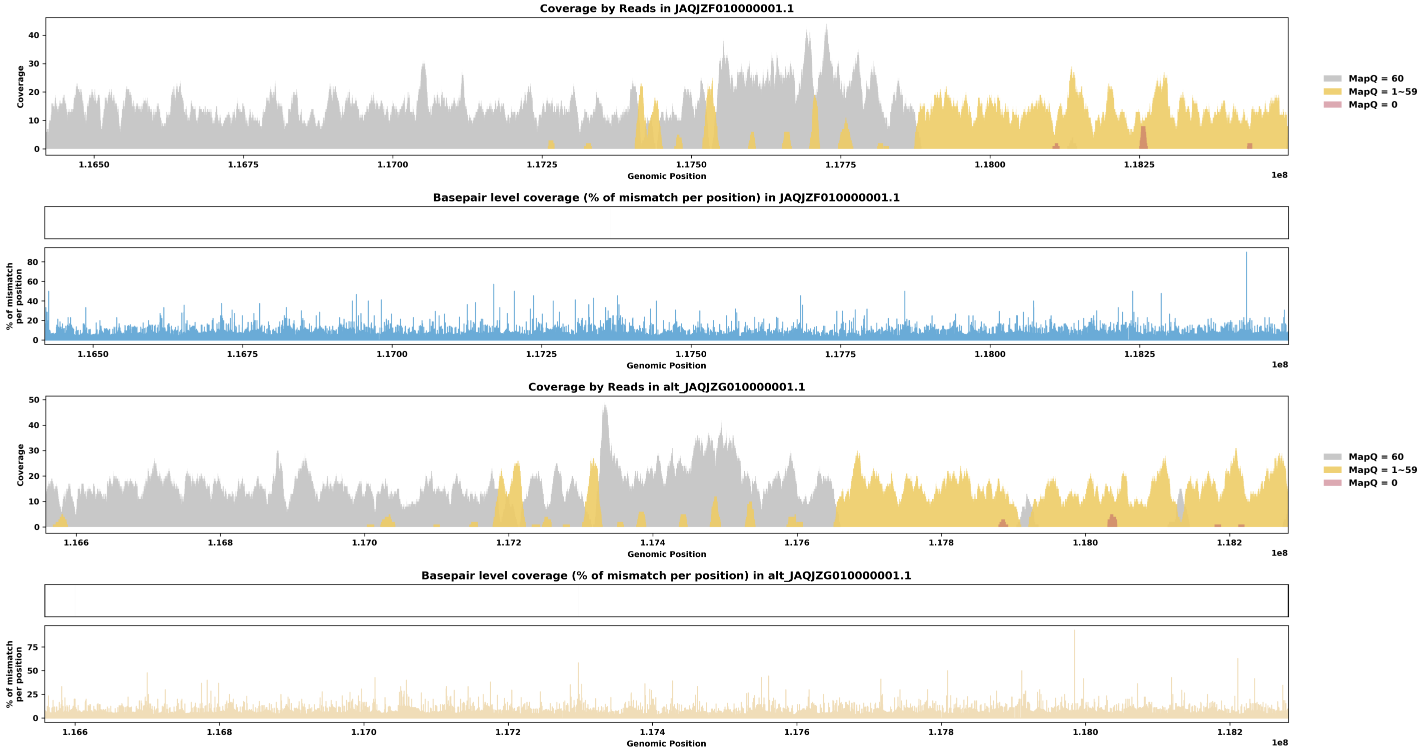

Species ID: mMicMin1

Common Name: European harvest mouse

Scientific Name: Micromys minutus

Assembly Type: Not Haplotype Resolved

Data Source: VGP

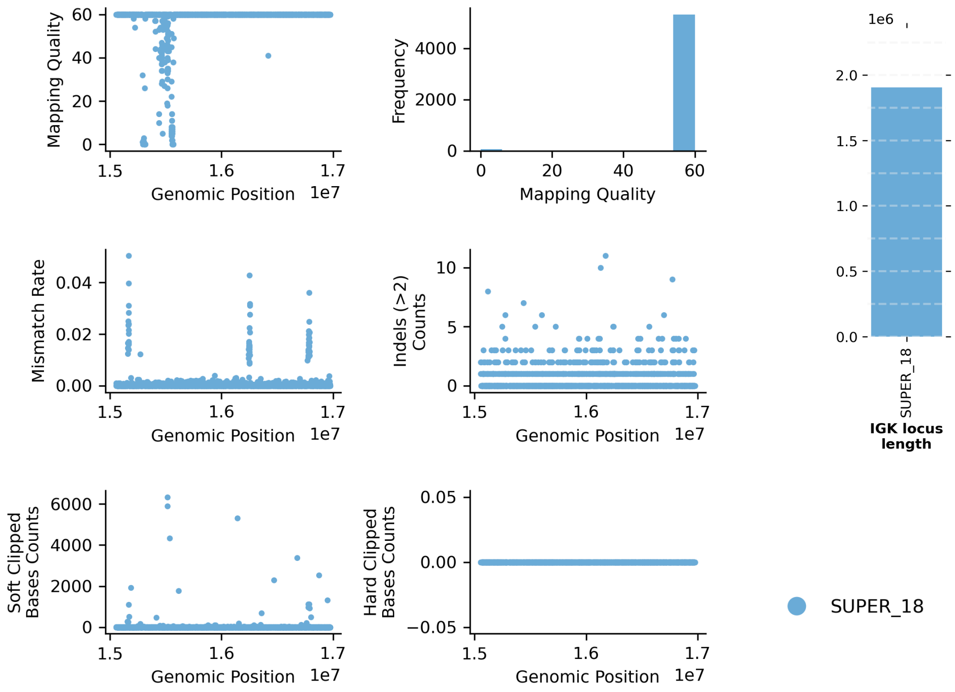


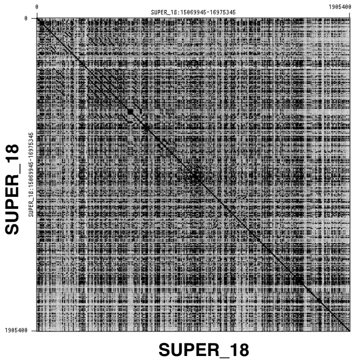


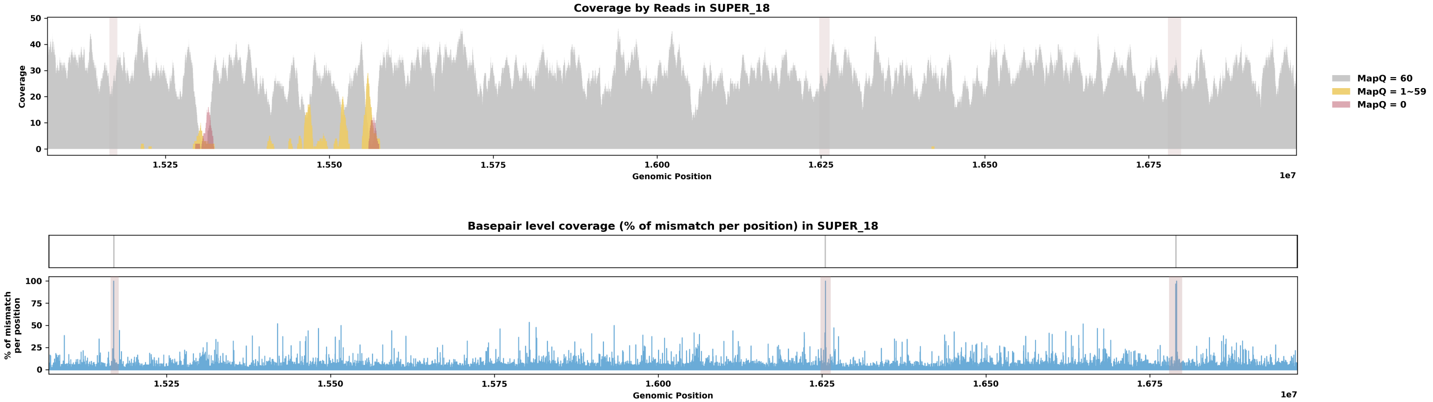

Species ID: mMirAng1

Common Name: Northern Elephant Seal

Scientific Name: Mirounga angustirostris

Assembly Type: Haplotype Resolved

Data Source: CCGP

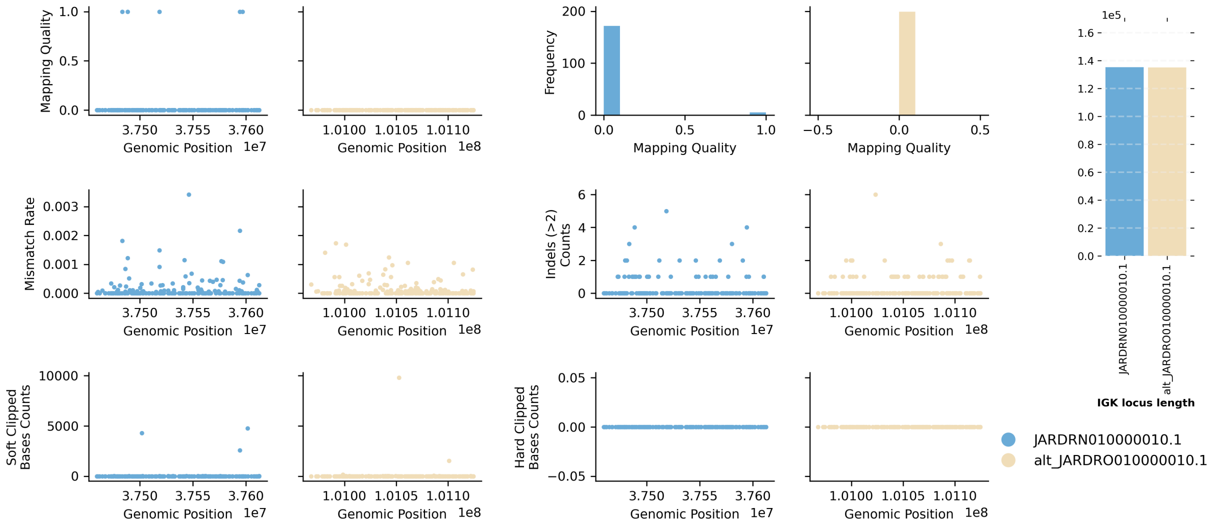


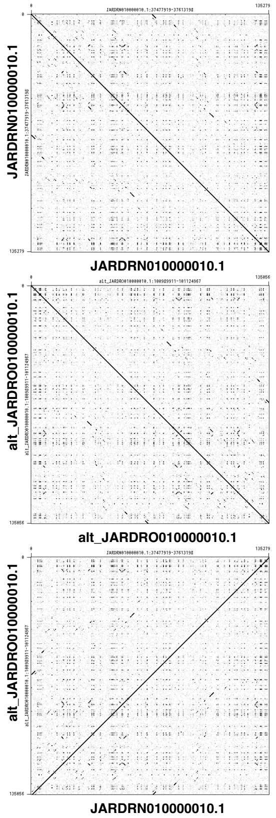


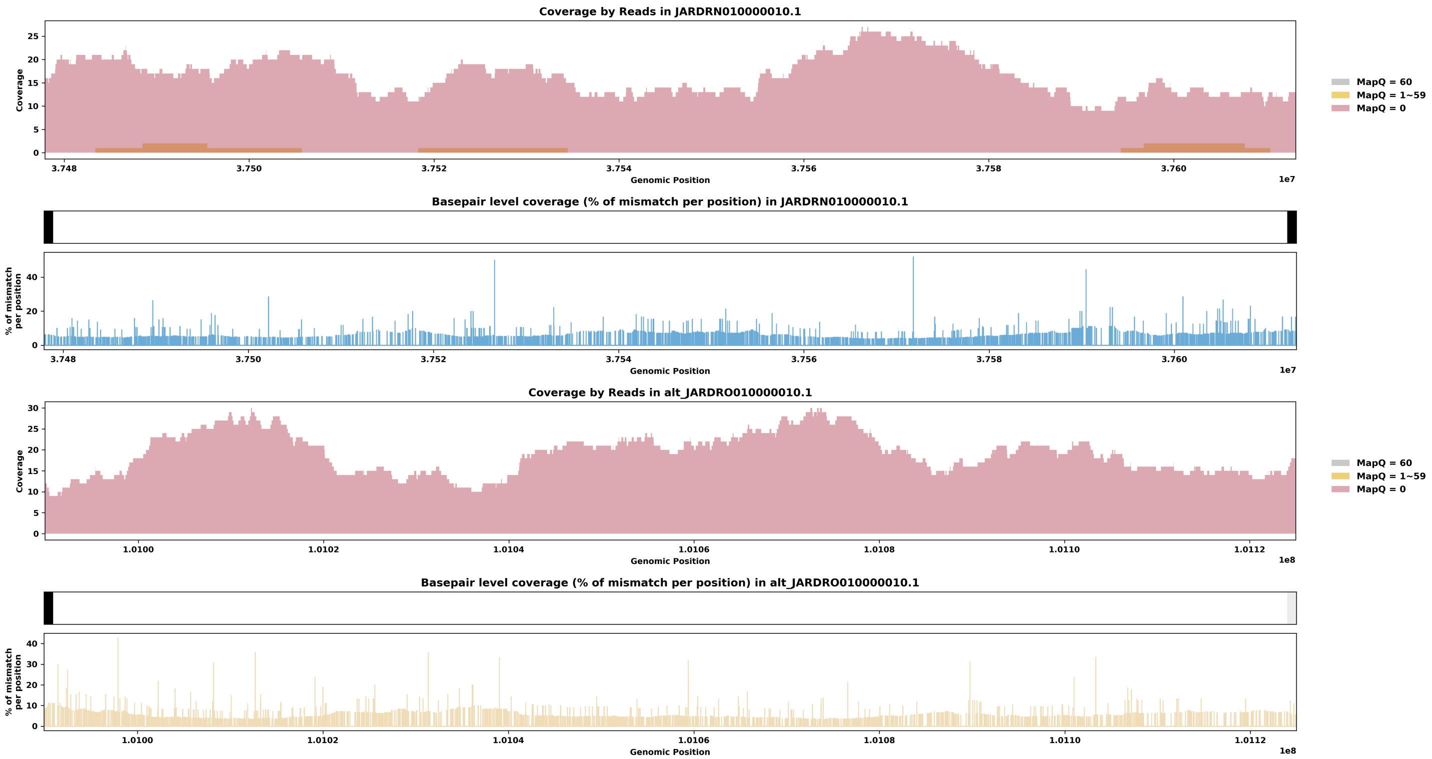

Species ID: mMonDom1

Common Name: gray short-tailed opossum

Scientific Name: Monodelphis_domestica

Assembly Type: Not Haplotype Resolved

Data Source: VGP

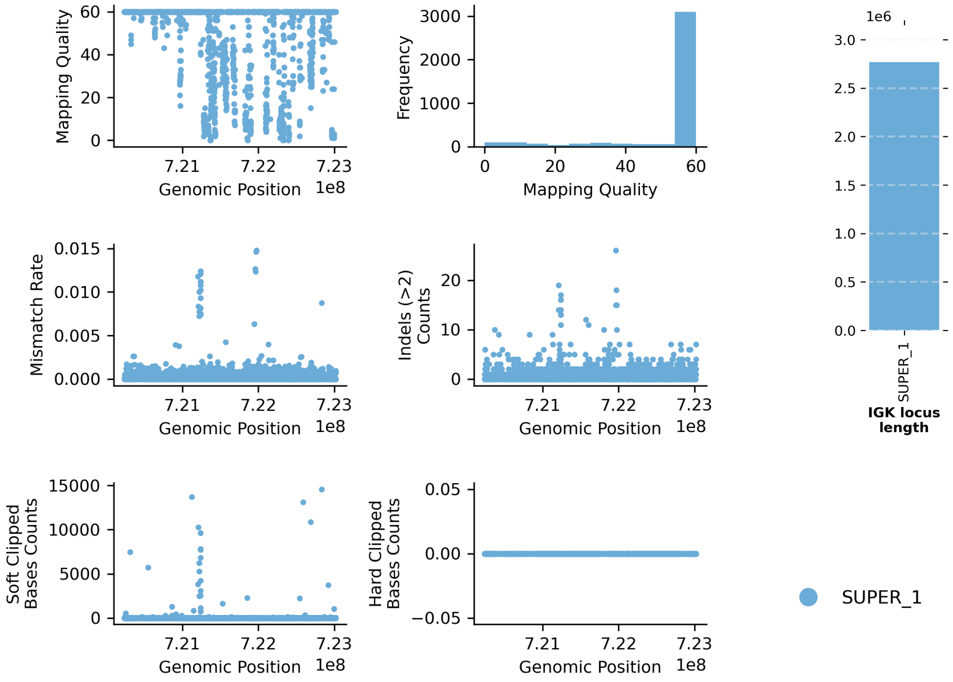


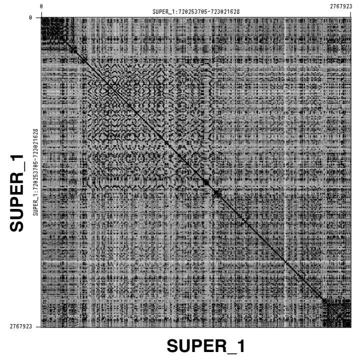


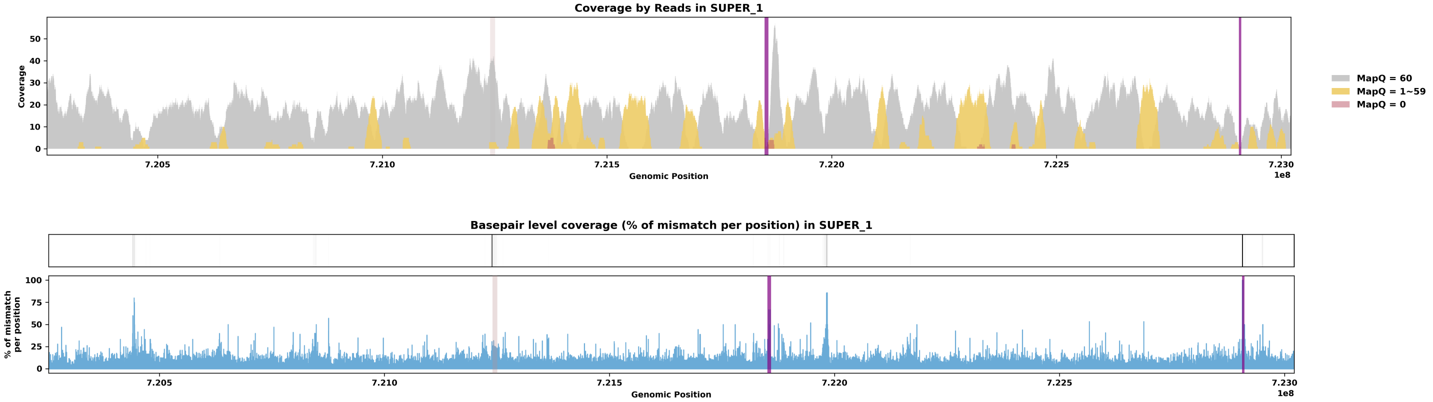

Species ID: mMunRee1

Common Name: Reeves' muntjac

Scientific Name: Muntiacus reevesi

Assembly Type: Not Haplotype Resolved

Data Source: VGP

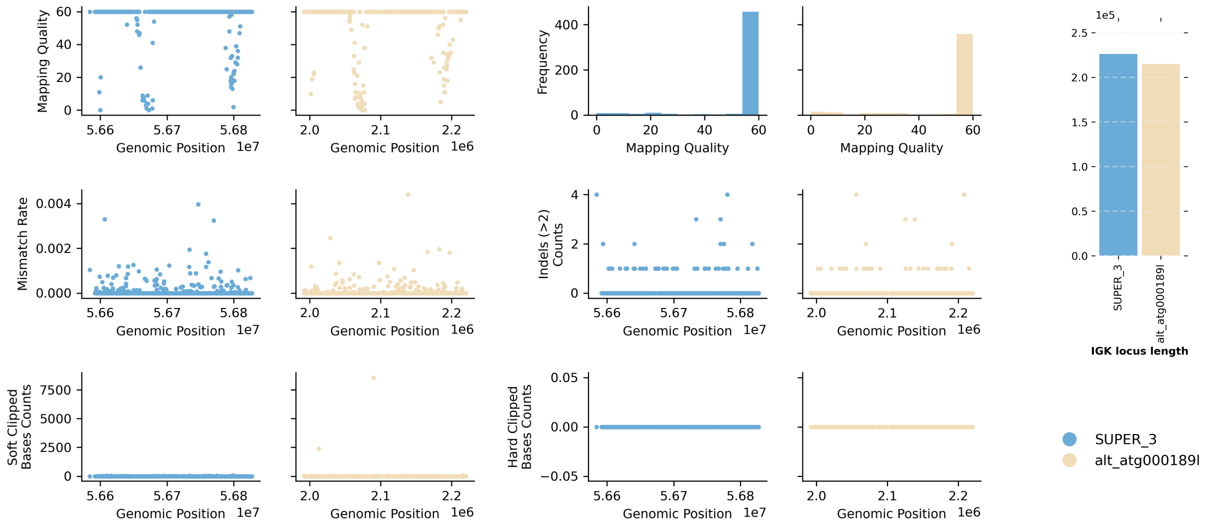


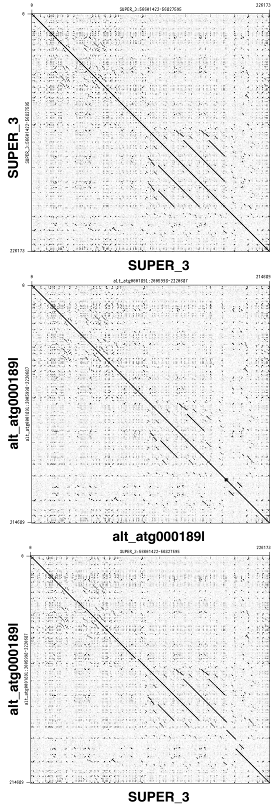


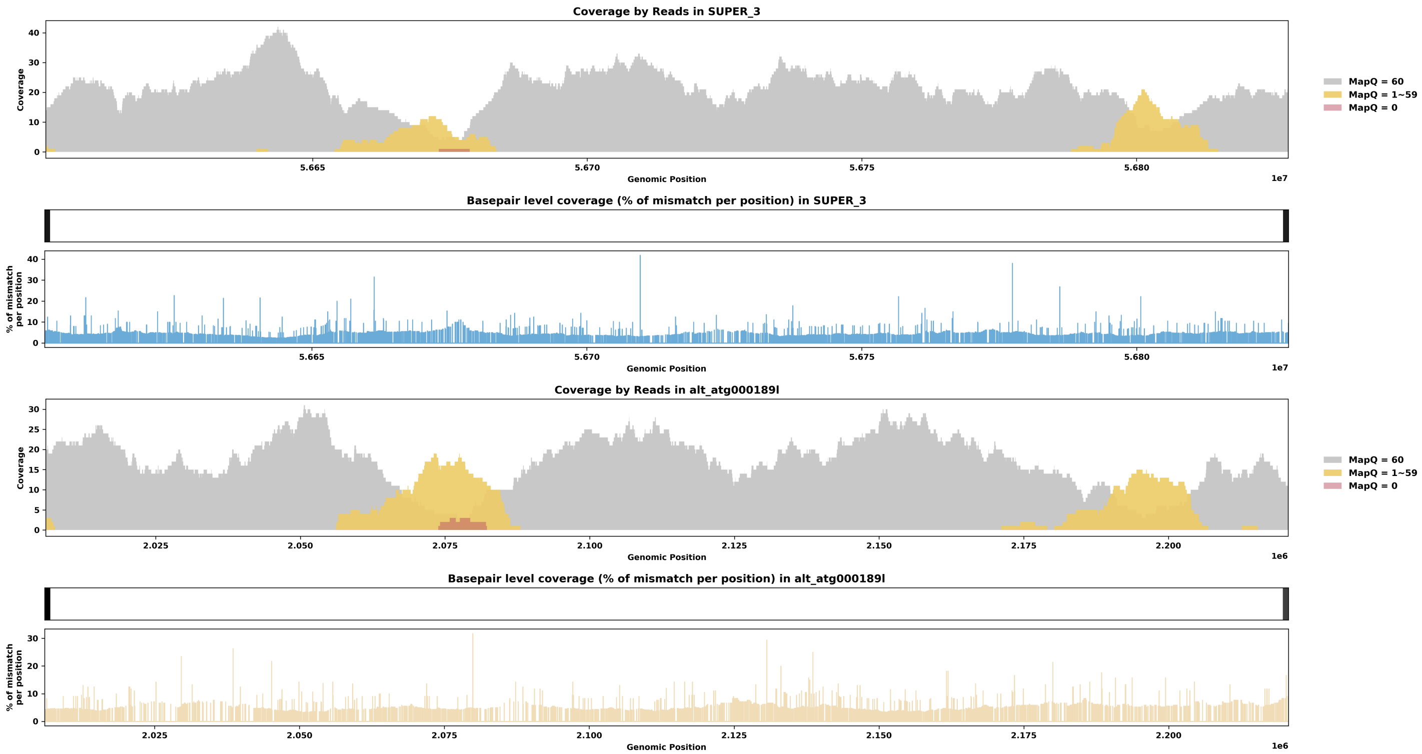

Species ID: mMusAve1

Common Name: hazel dormouse

Scientific Name: Muscardinus avellanarius

Assembly Type: Not Haplotype Resolved

Data Source: VGP

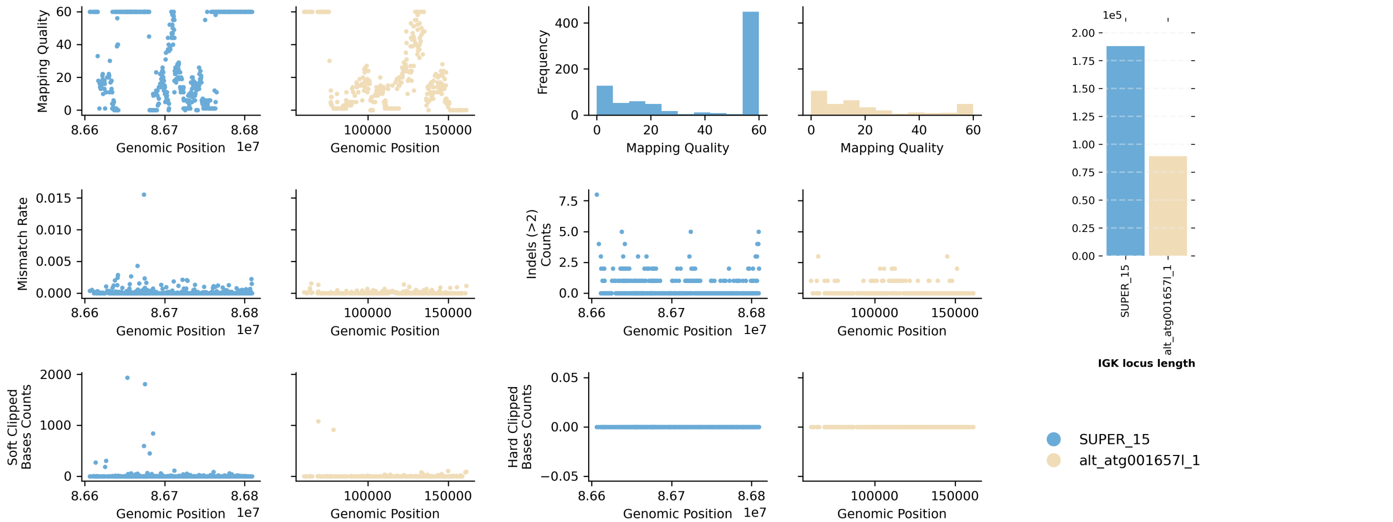


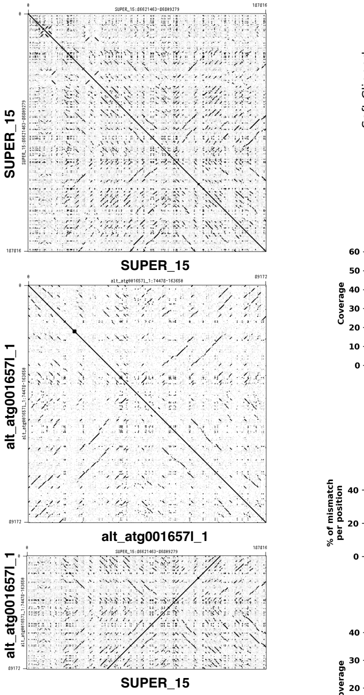


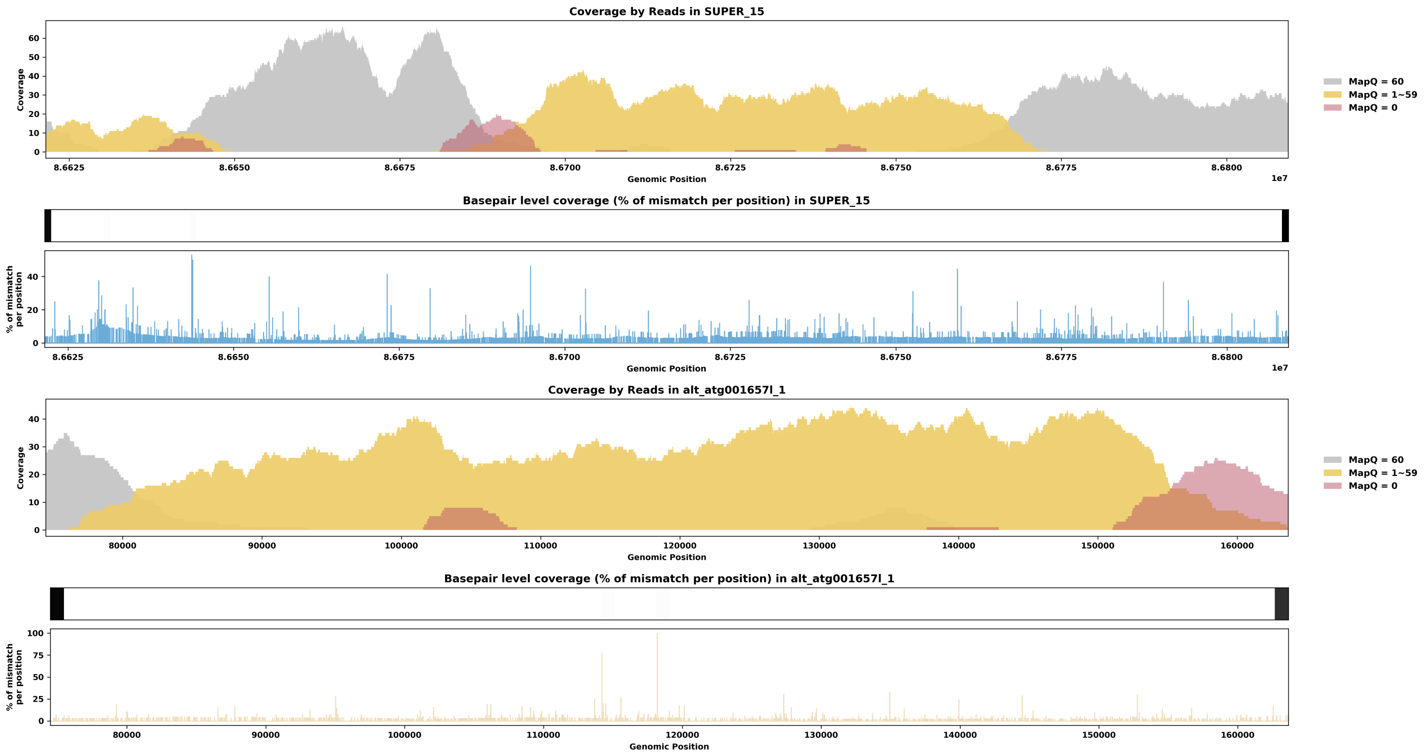

Species ID: mMusLut2

Common Name: European mink

Scientific Name: Mustela lutreola

Assembly Type: Not Haplotype Resolved

Data Source: VGP

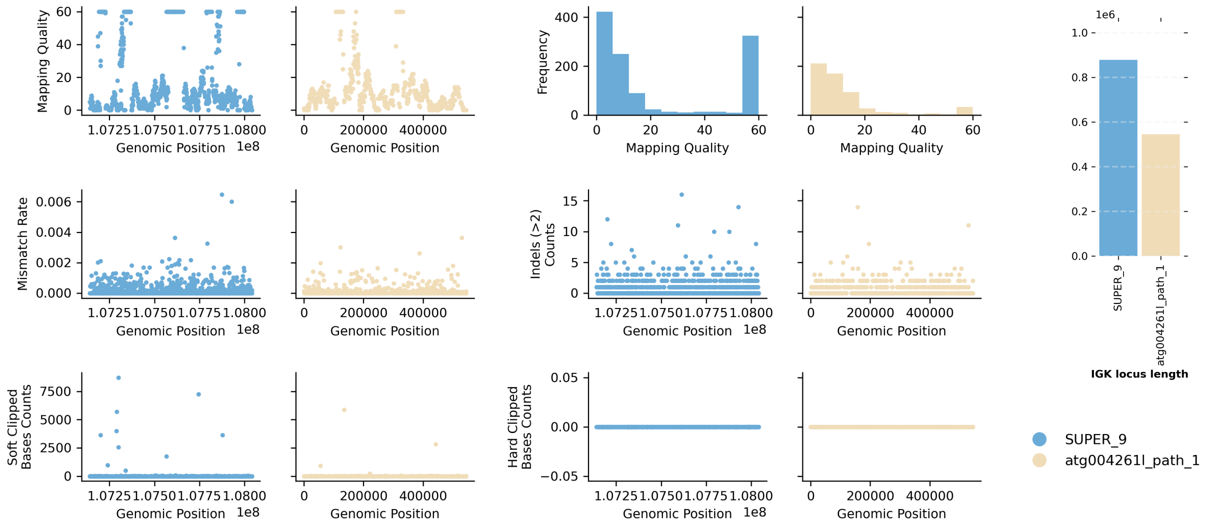


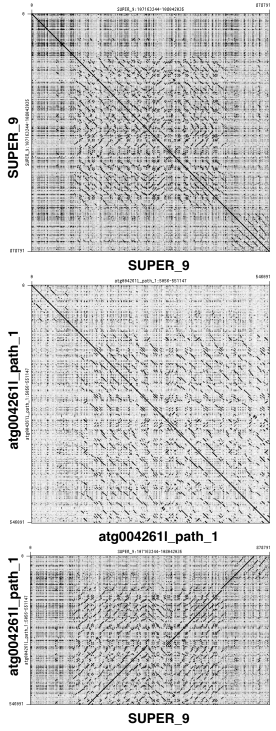


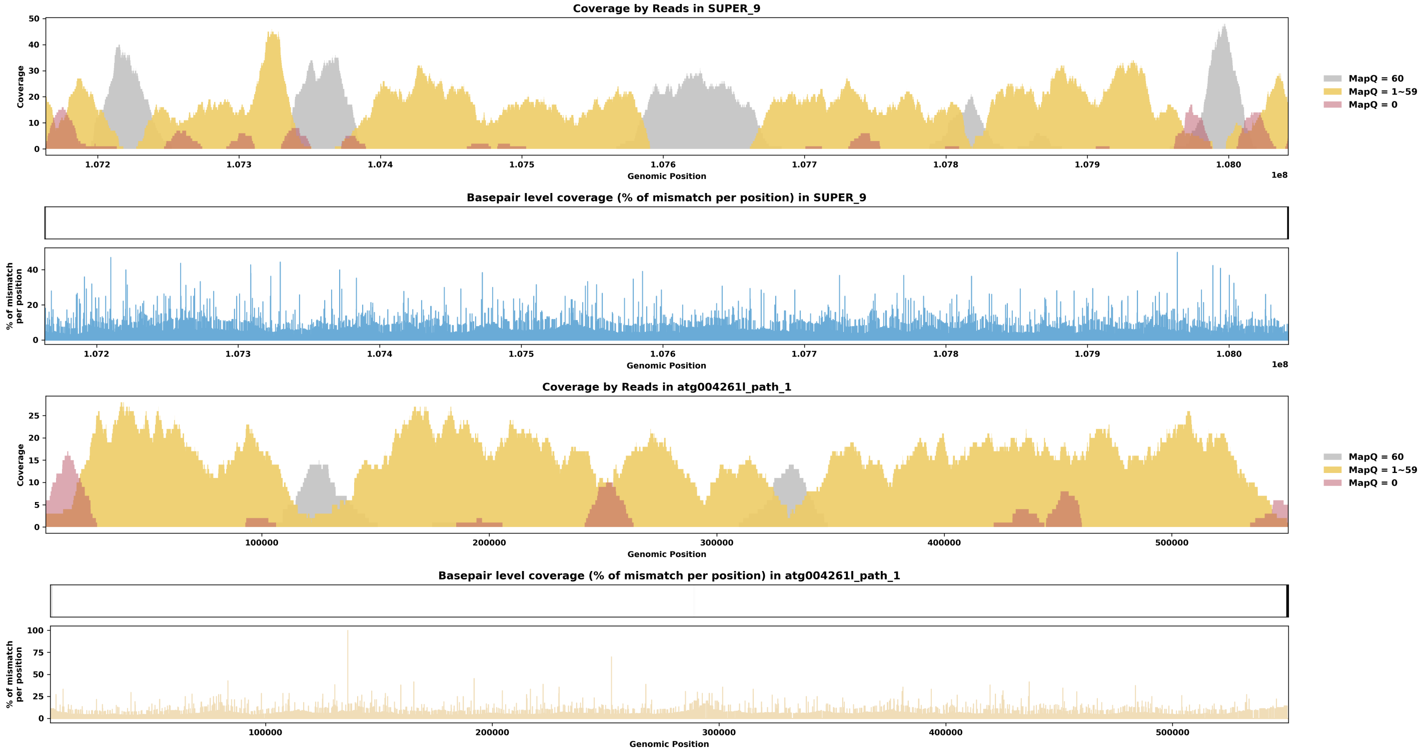

Species ID: mMusNiv1

Common Name: Least weasel

Scientific Name: Mustela nivalis

Assembly Type: Haplotype Resolved

Data Source: VGP

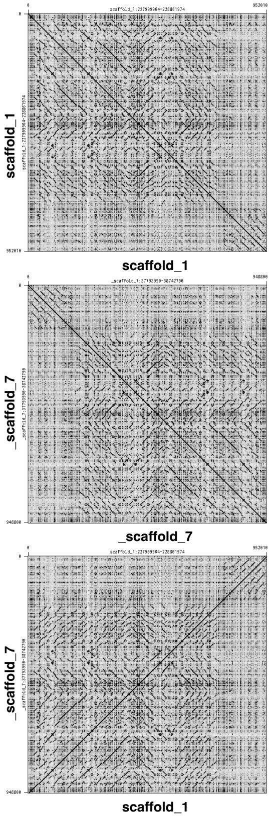


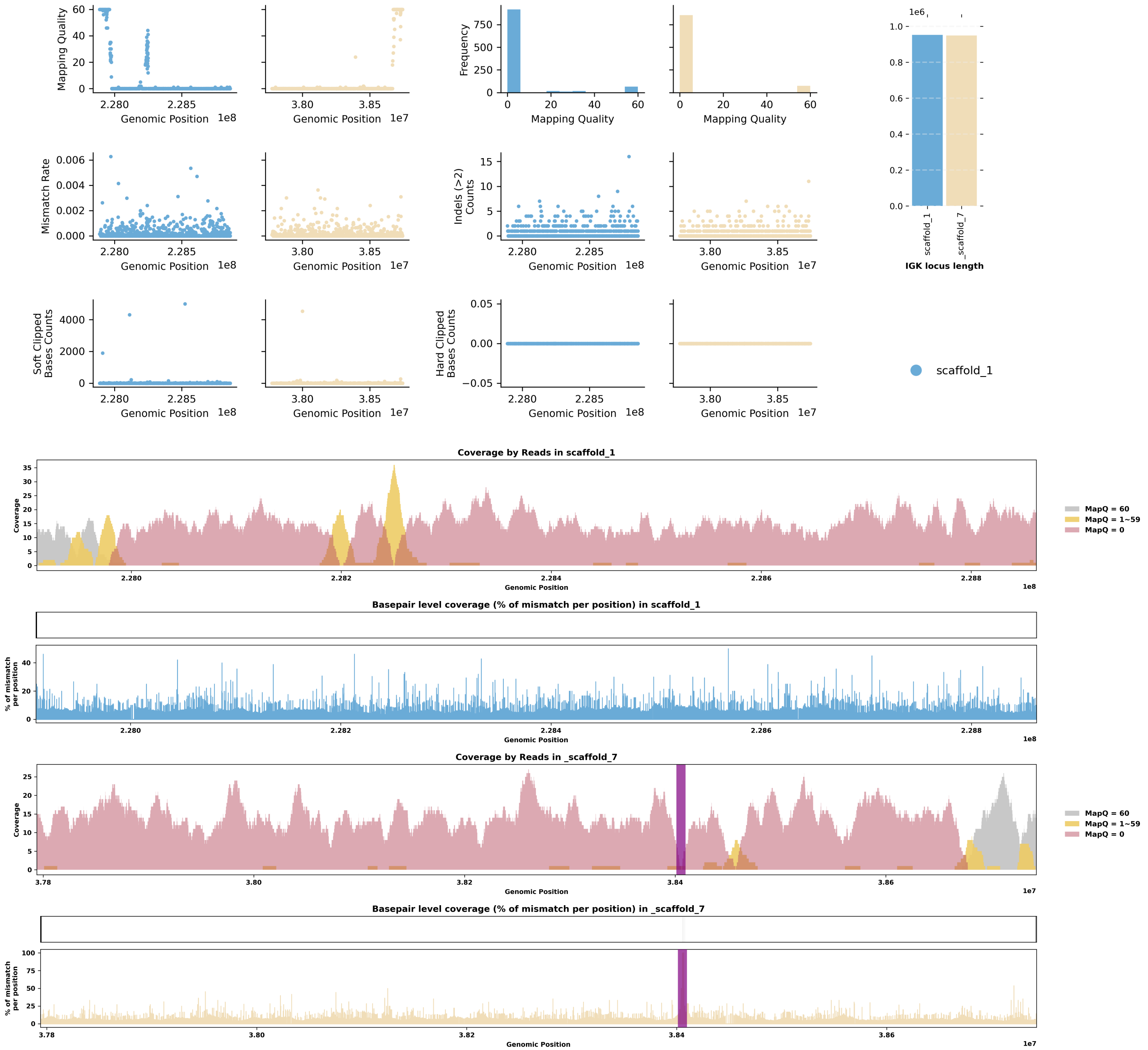

Species ID: mNeoNeb1

Common Name: Clouded Leopard

Scientific Name: Neofelis_nebulosa

Assembly Type: Not Haplotype Resolved

Data Source: VGP

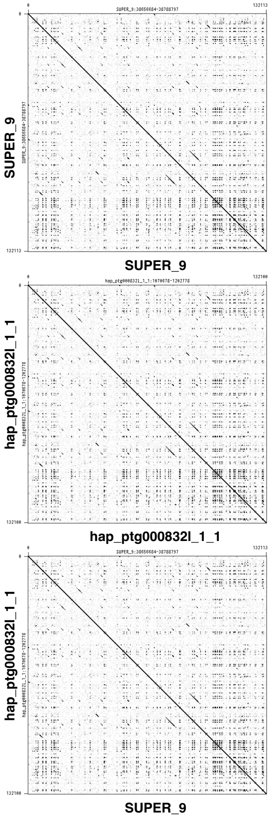


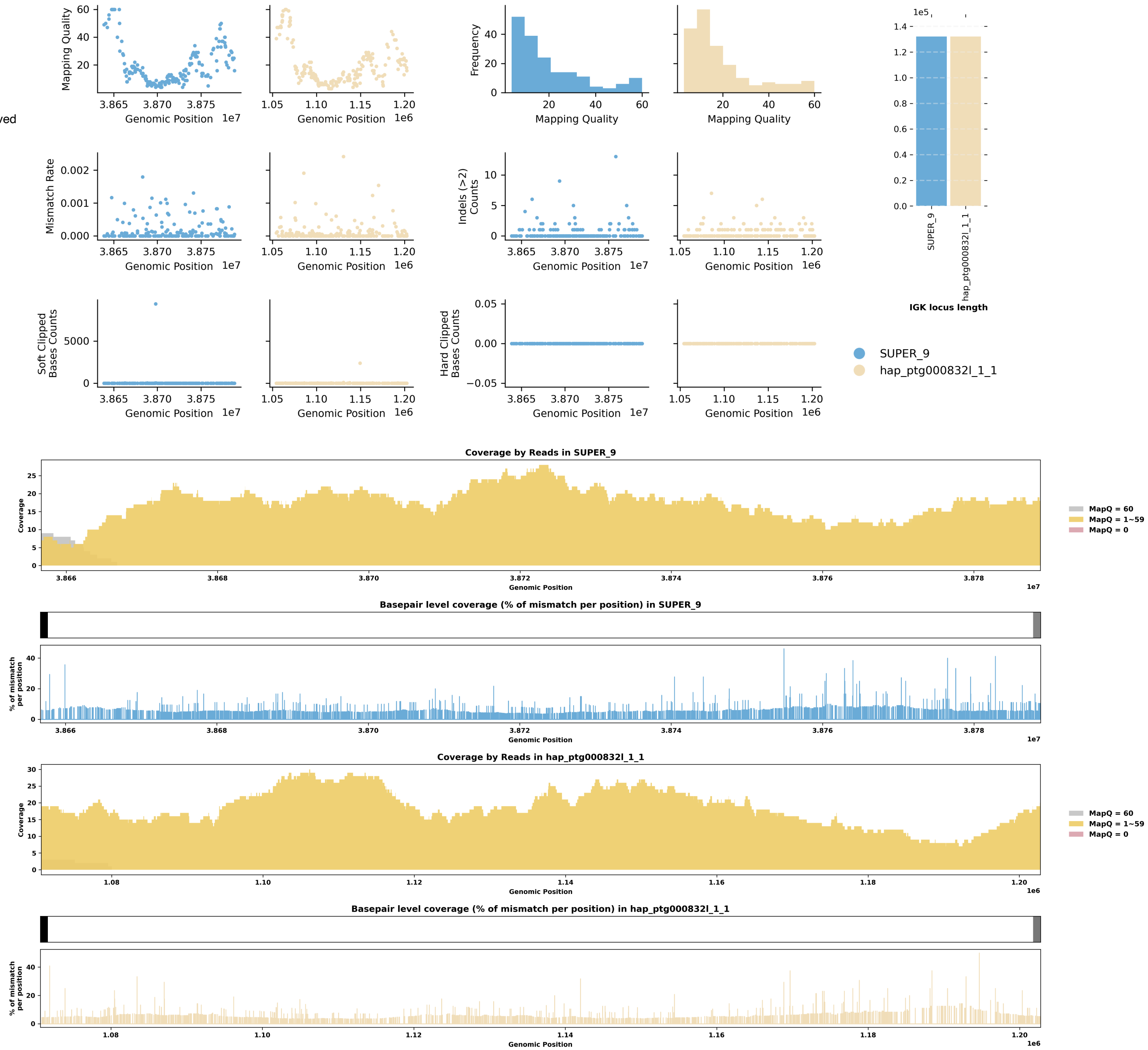

Species ID: mNycCou1

Common Name: slow loris

Scientific Name: Nycticebus coucang

Assembly Type: Not Haplotype Resolved

Data Source: VGP

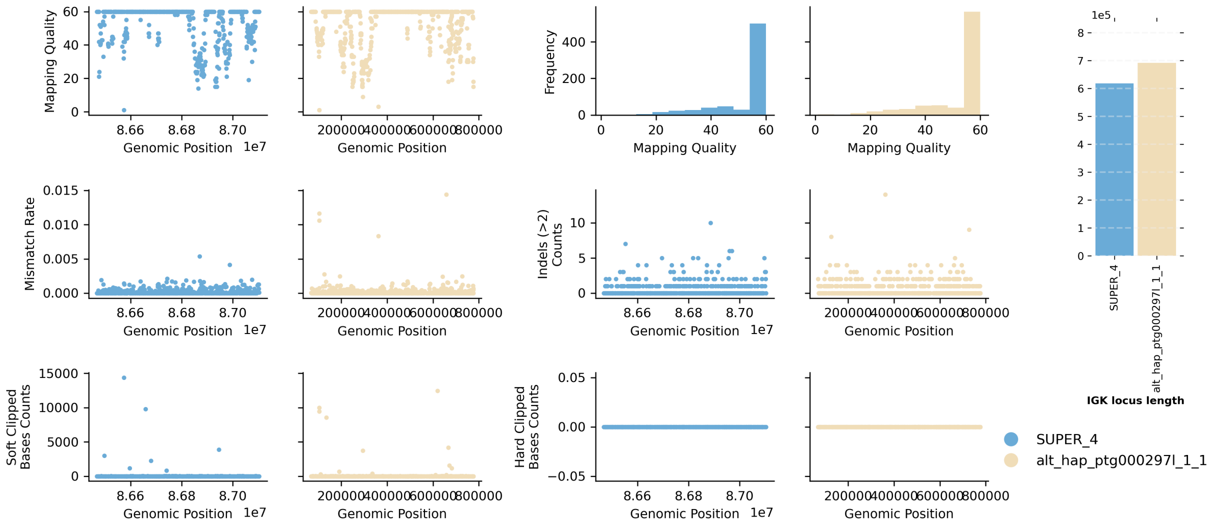


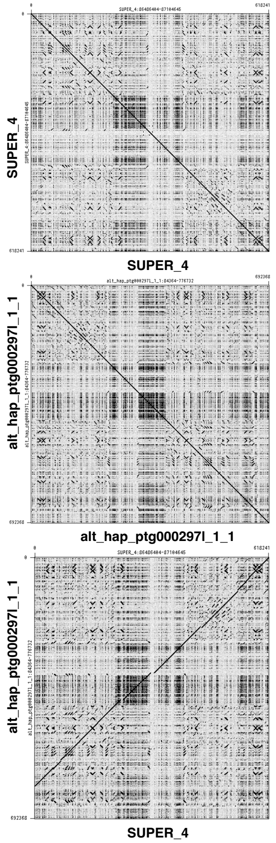


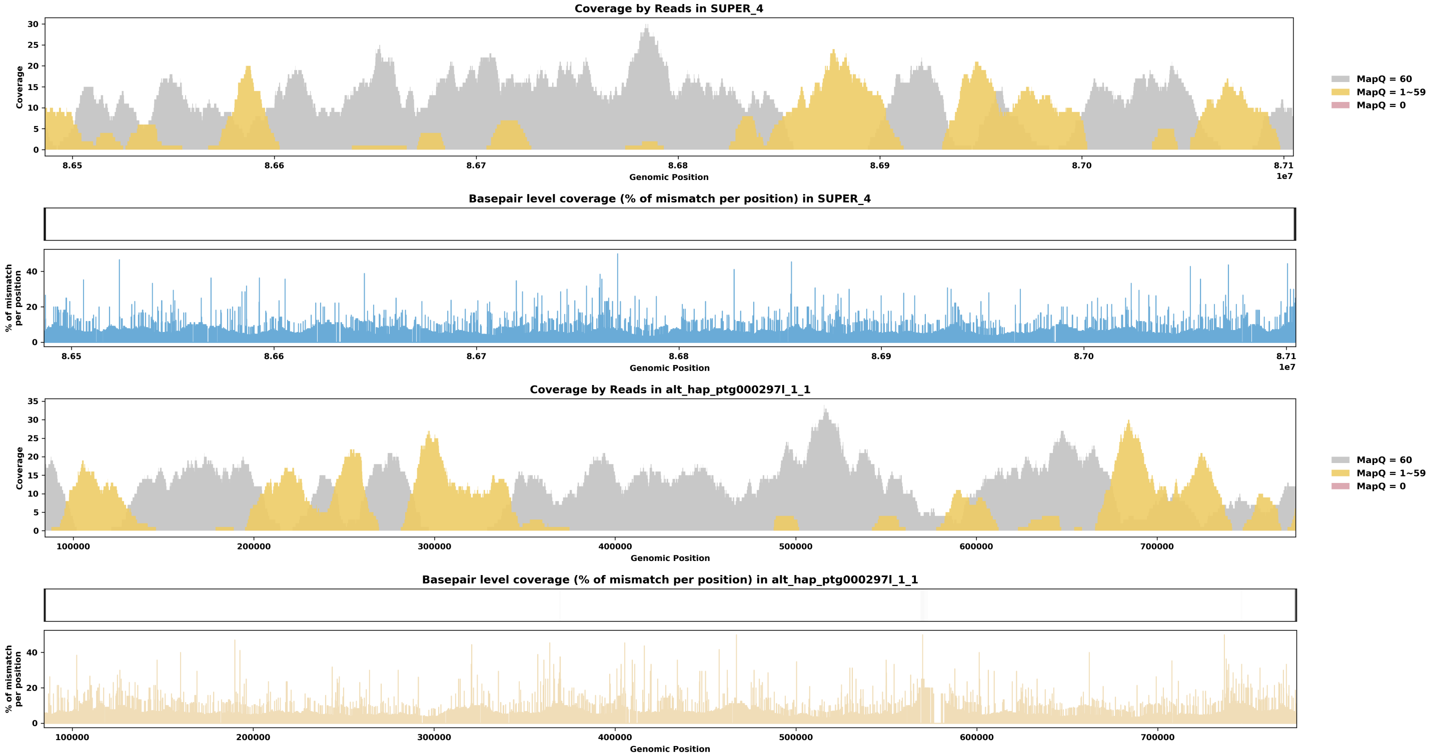

Species ID: mOrcOrc1

Common Name: killer whale

Scientific Name: Orcinus orca

Assembly Type: Not Haplotype Resolved

Data Source: VGP

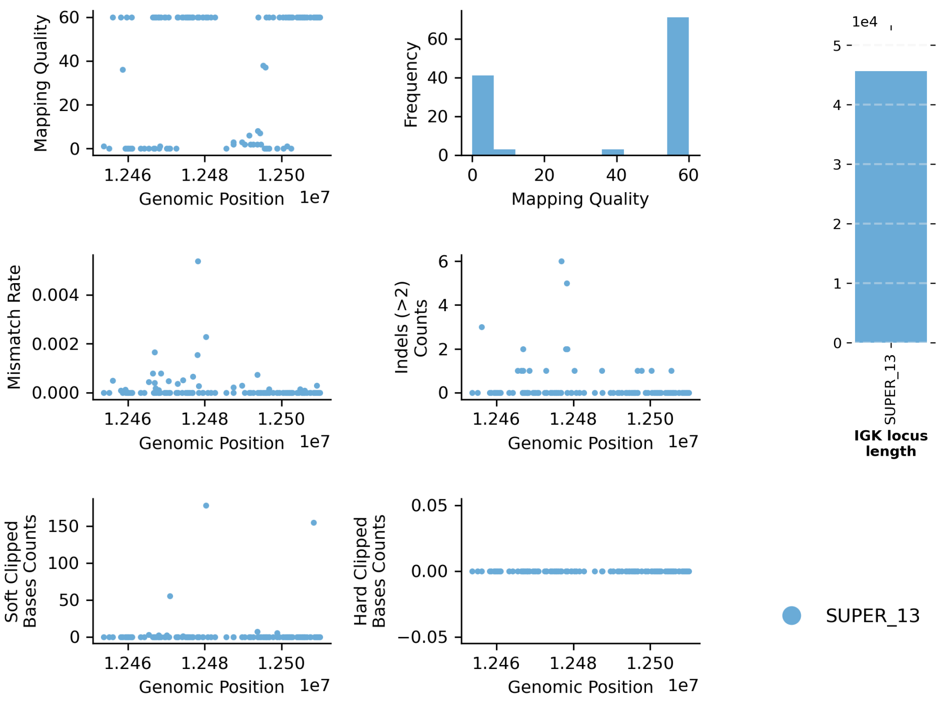


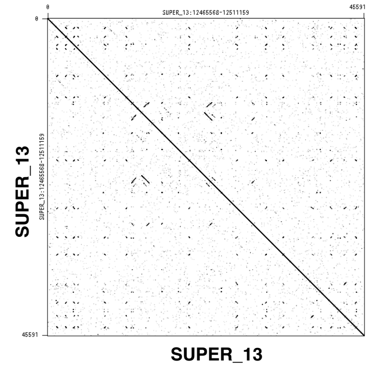


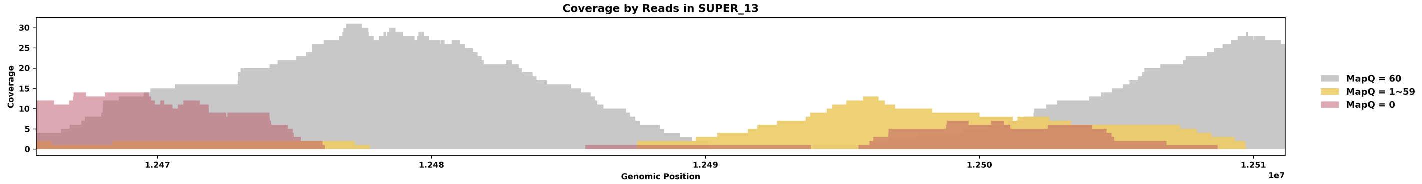


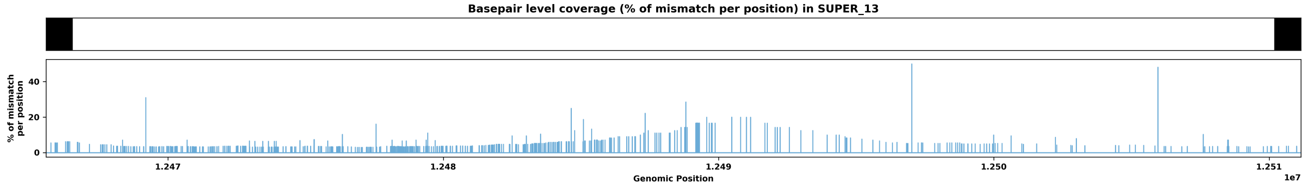

Species ID: mOryCun1

Common Name: rabbit

Scientific Name: Oryctolagus cuniculus

Assembly Type: Not Haplotype Resolved

Data Source: VGP

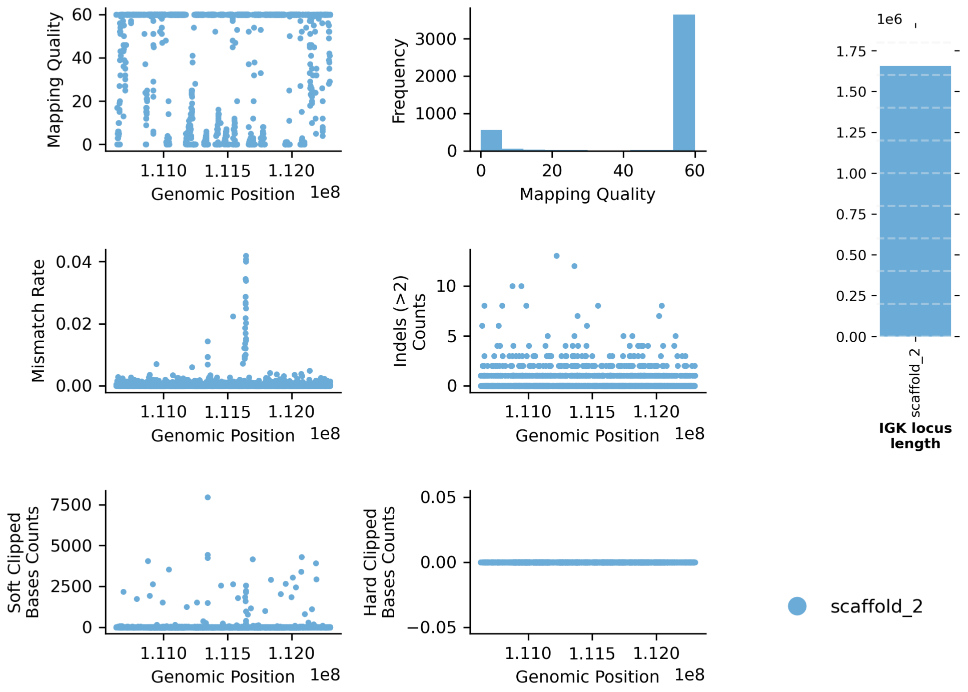


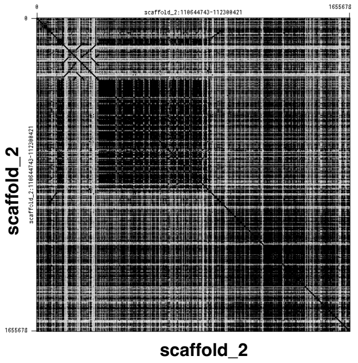


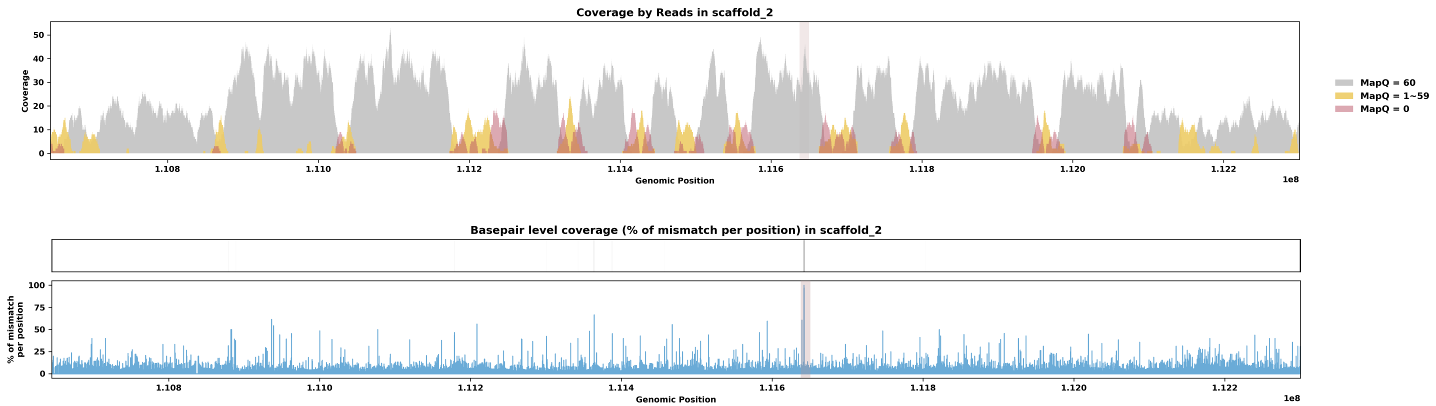

Species ID: mPanPan1

Common Name: Bonobo

Scientific Name: Pan paniscus

Assembly Type: Haplotype Resolved

Data Source: T2T Primate

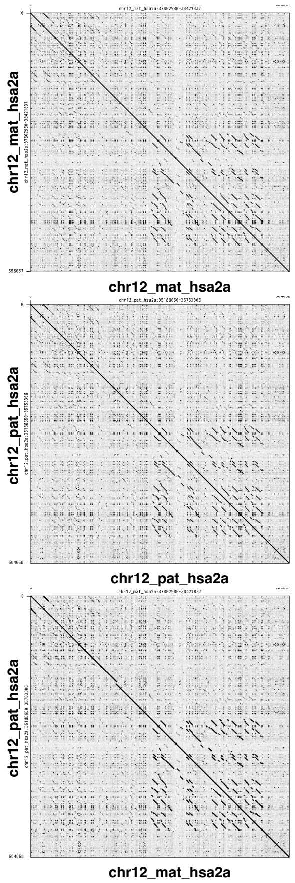


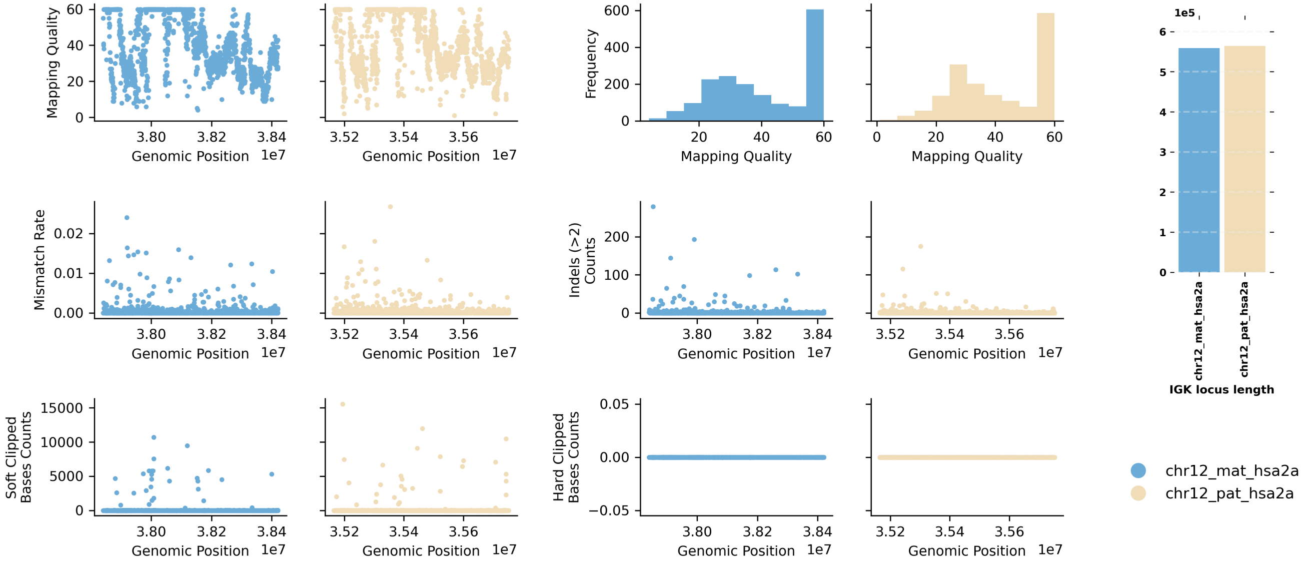


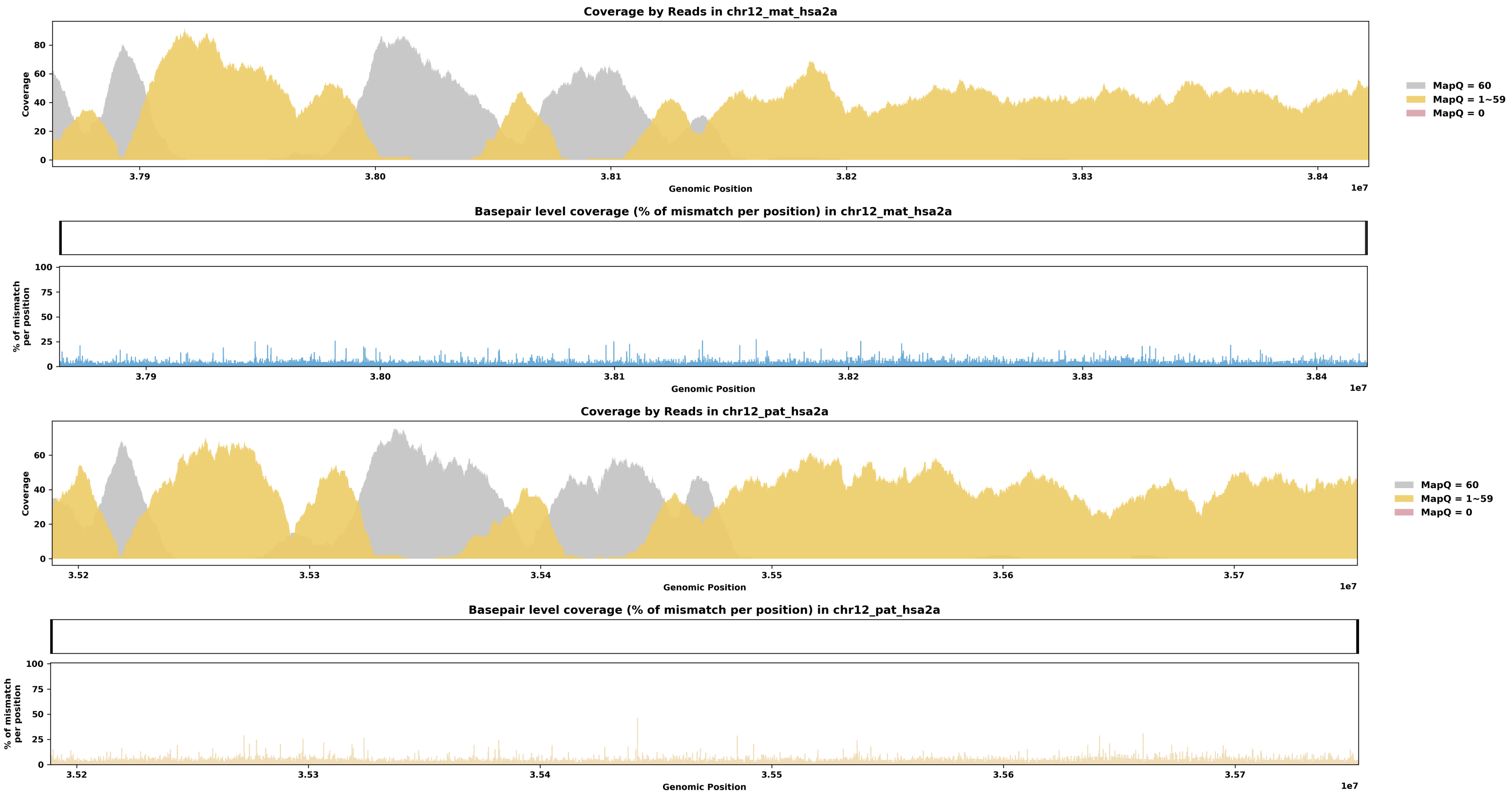

Species ID: mPerMan1

Common Name: deer mouse

Scientific Name: Peromyscus maniculatus

Assembly Type: Not Haplotype Resolved

Data Source: CCGP

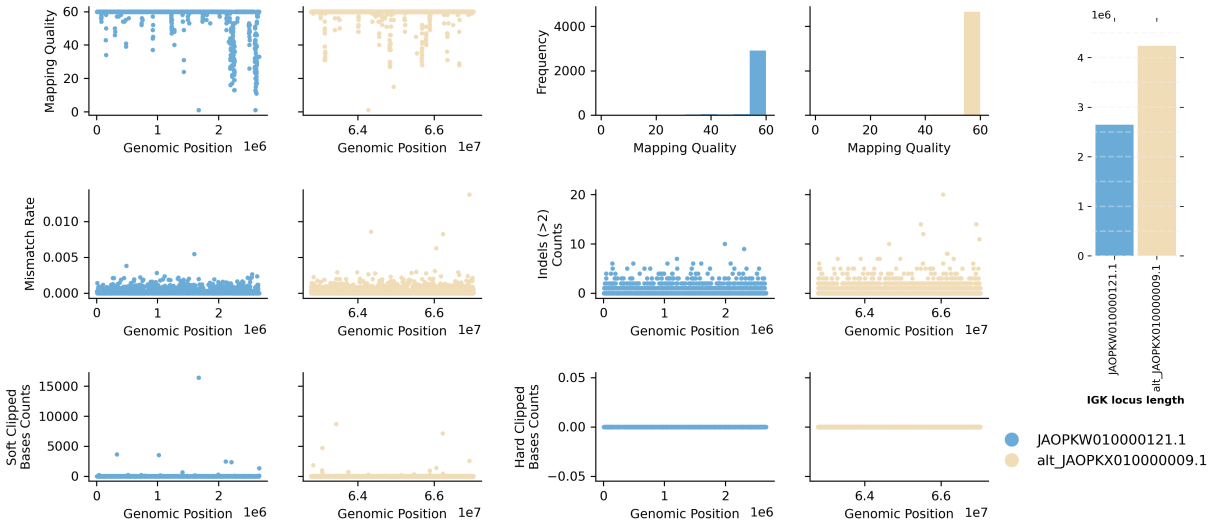


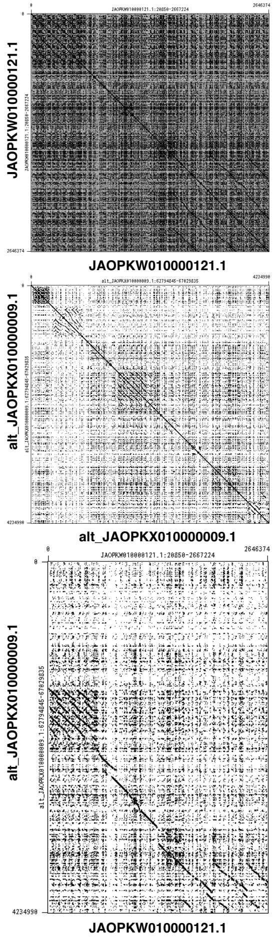


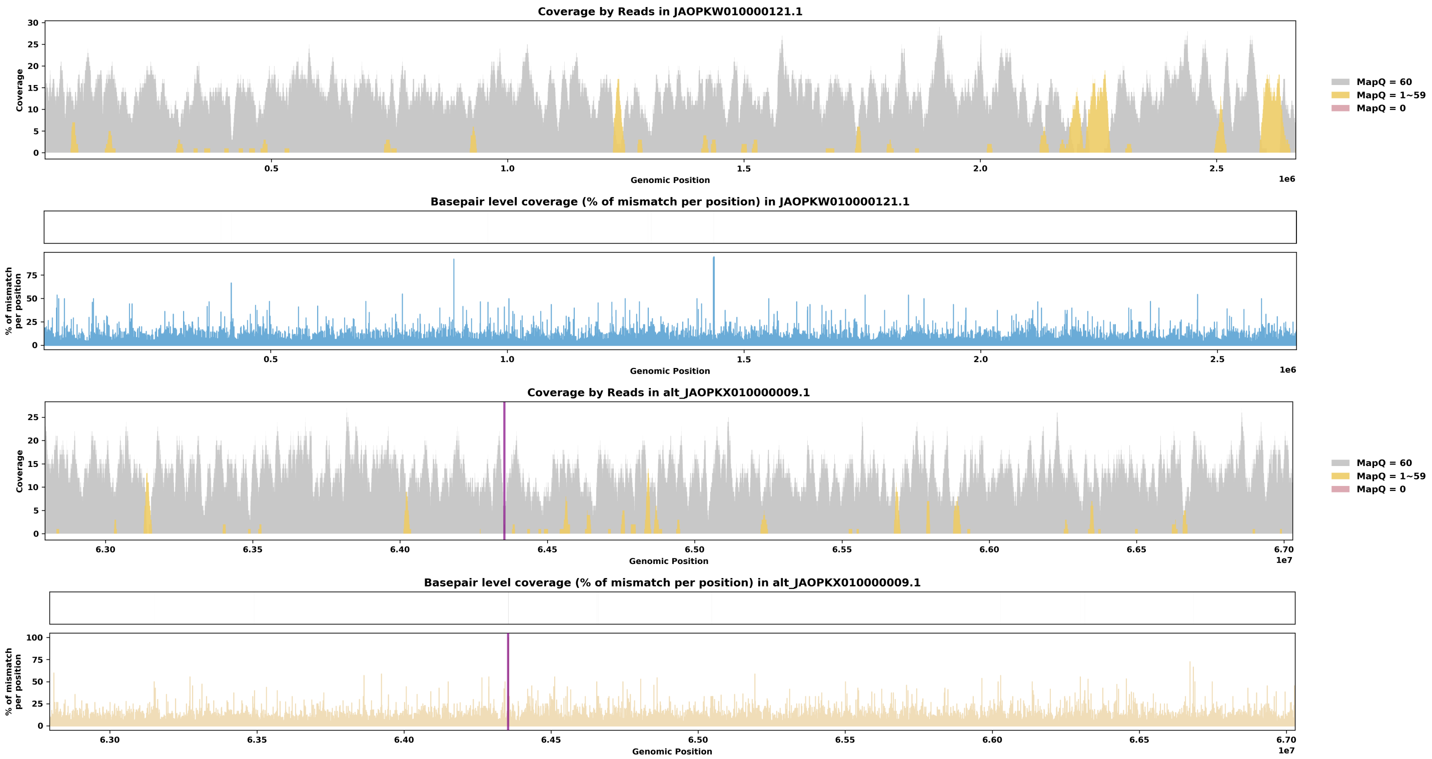

Species ID: mPhoPho1

Common Name: harbor porpoise

Scientific Name: Phocoena phocoena

Assembly Type: Not Haplotype Resolved

Data Source: VGP

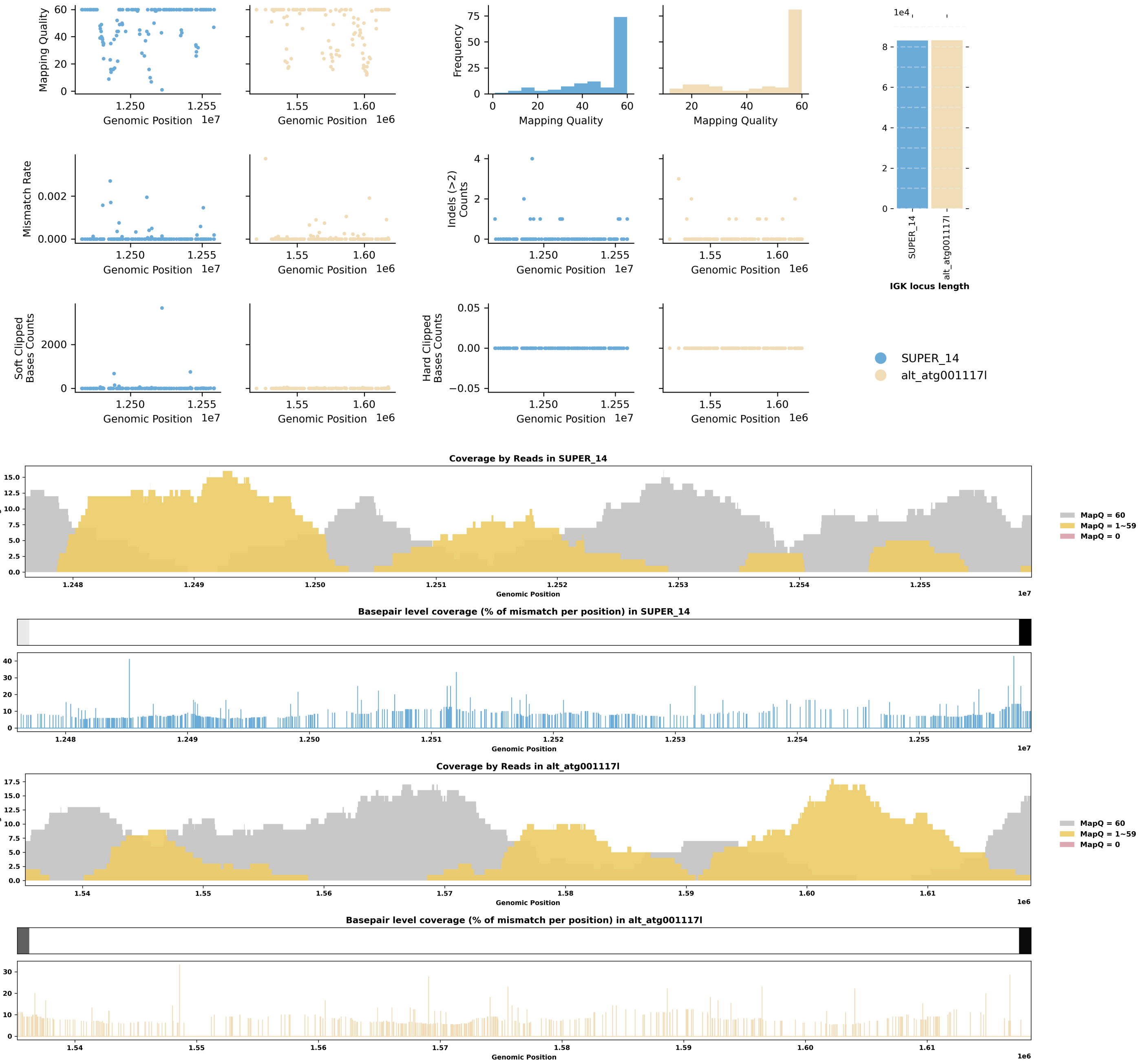


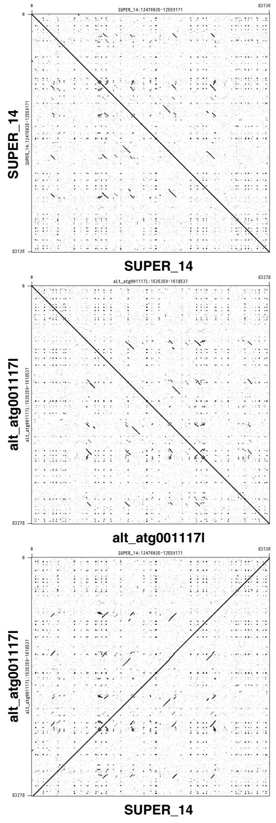

Species ID: mPonAbe1

Common Name: Sumatran orangutan

Scientific Name: Pongo_abelii

Assembly Type: Haplotype Resolved

Data Source: T2T Primate

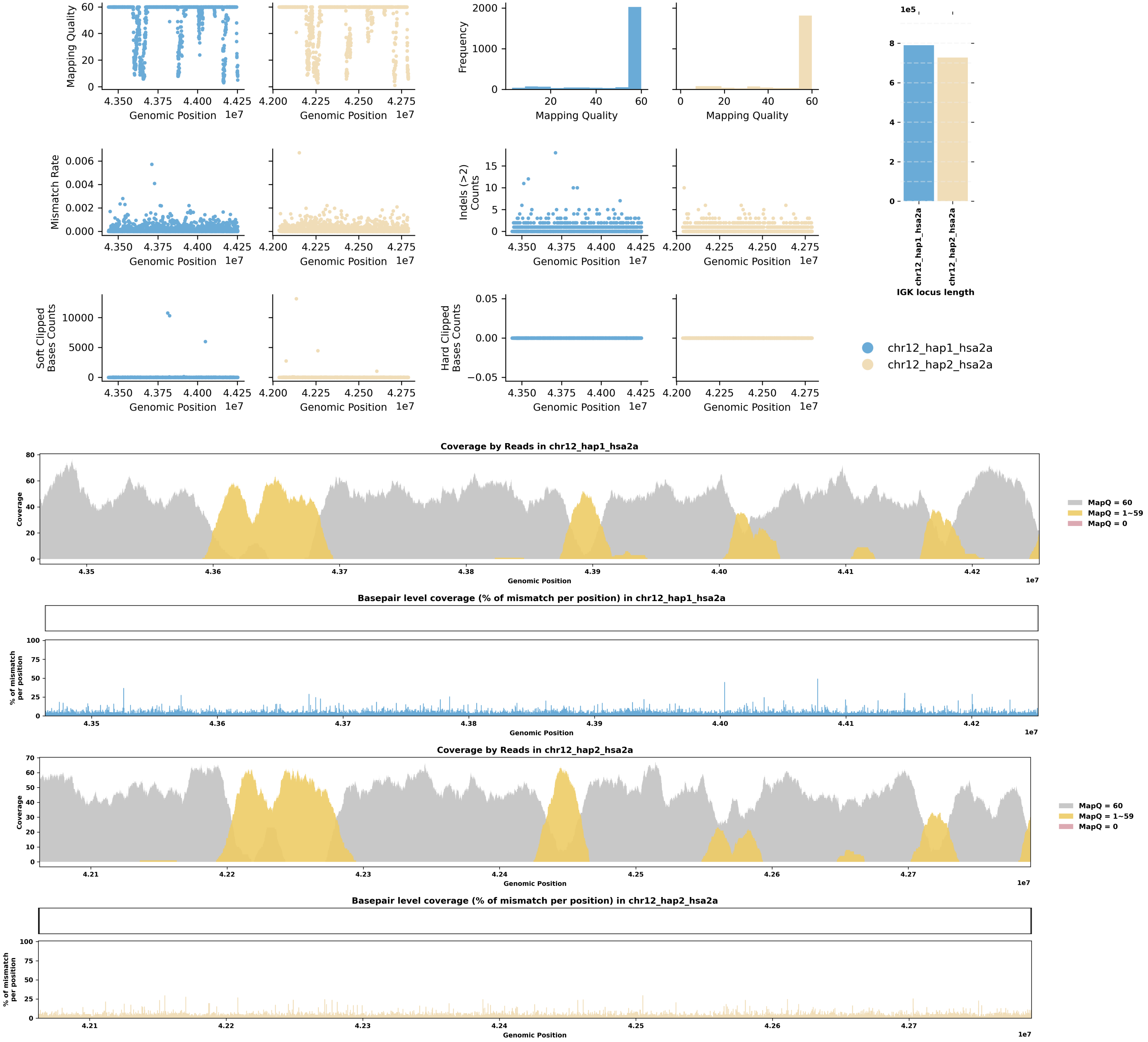


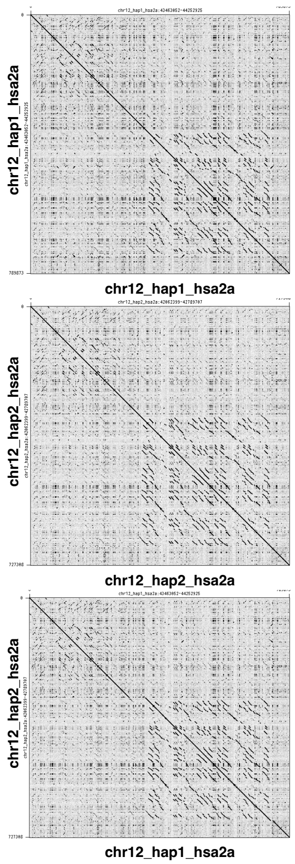

Species ID: mPonPyg2

Common Name: Bornean orangutan

Scientific Name: Pongo_pygmaeus

Assembly Type: Haplotype Resolved

Data Source: T2T Primate

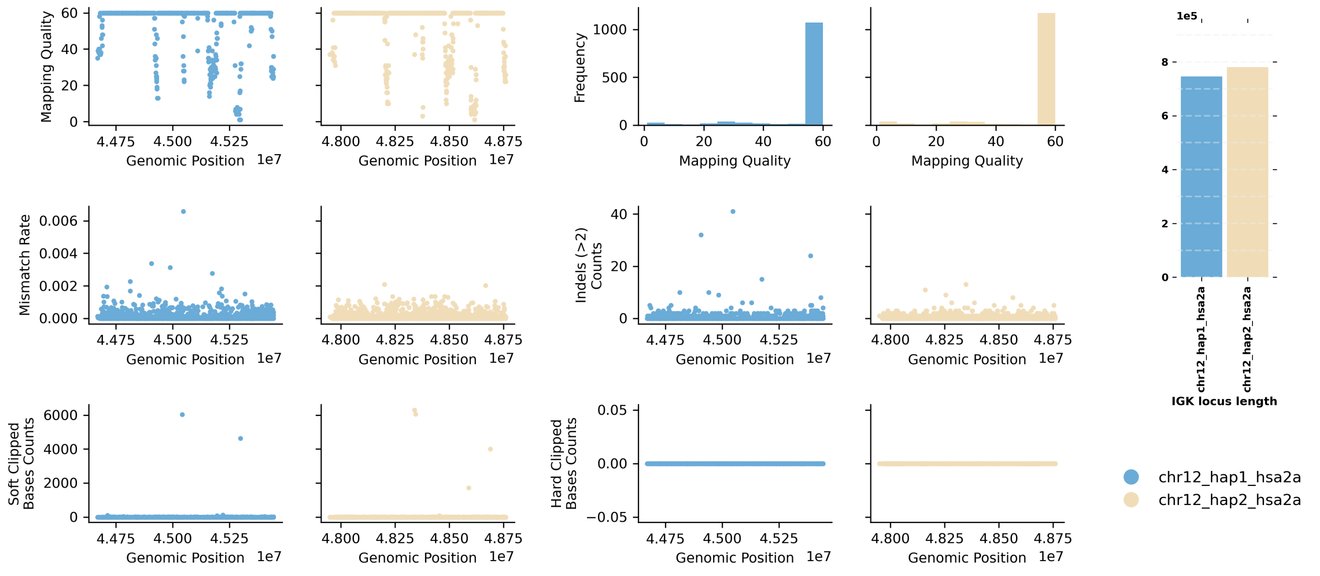


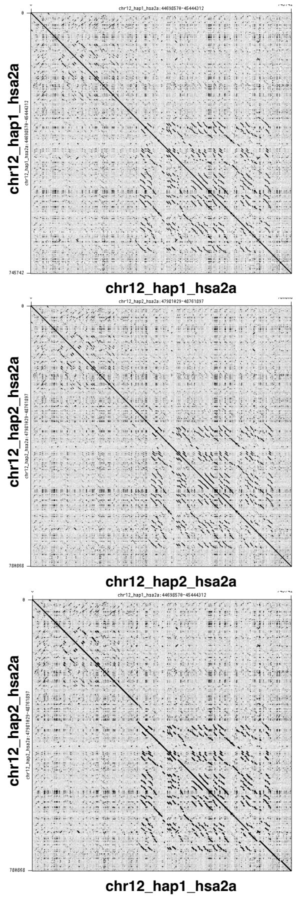


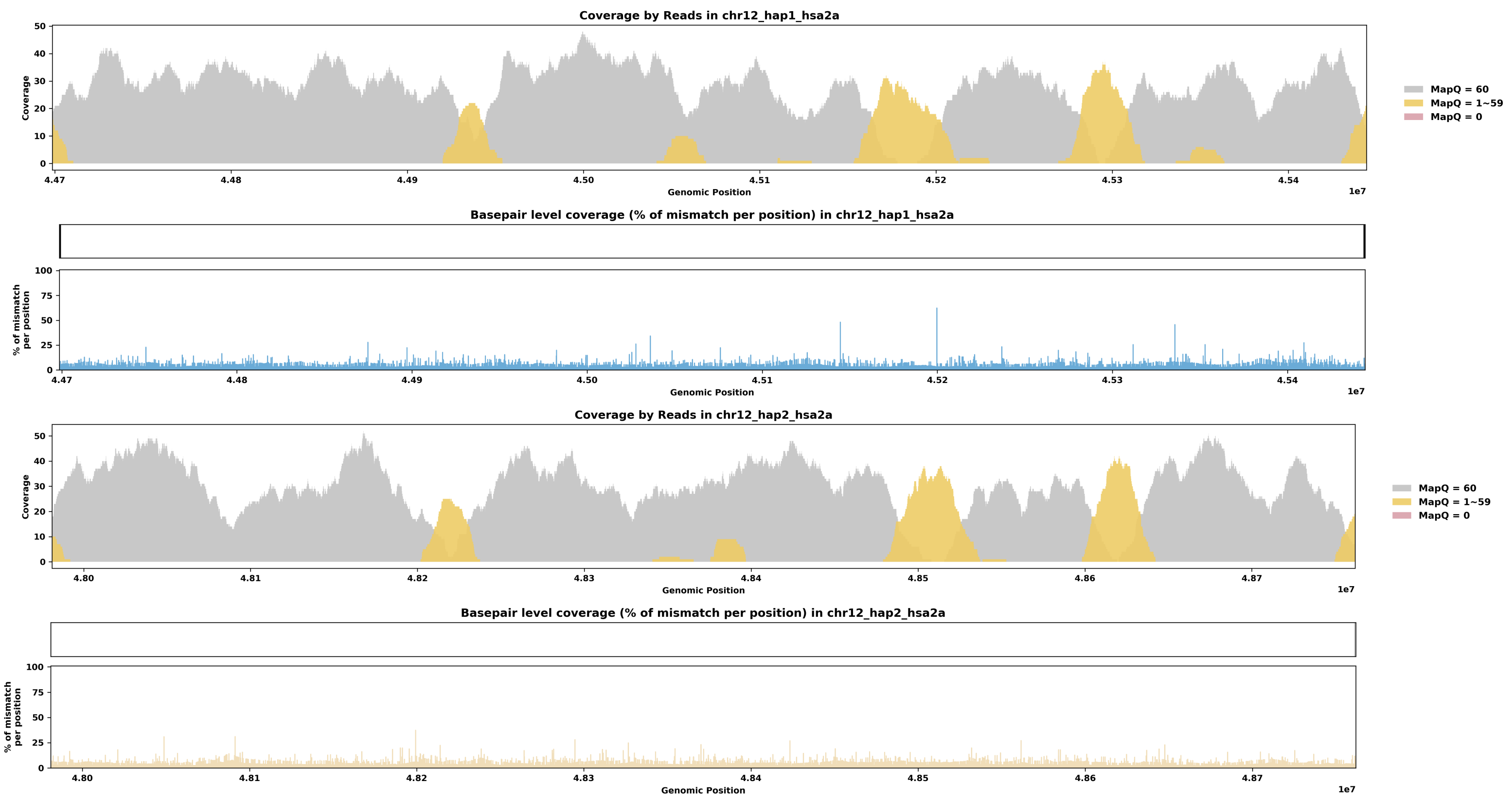

Species ID: mPseCra1

Common Name: false killer whale

Scientific Name: Pseudorca crassidens

Assembly Type: Haplotype Resolved

Data Source: VGP

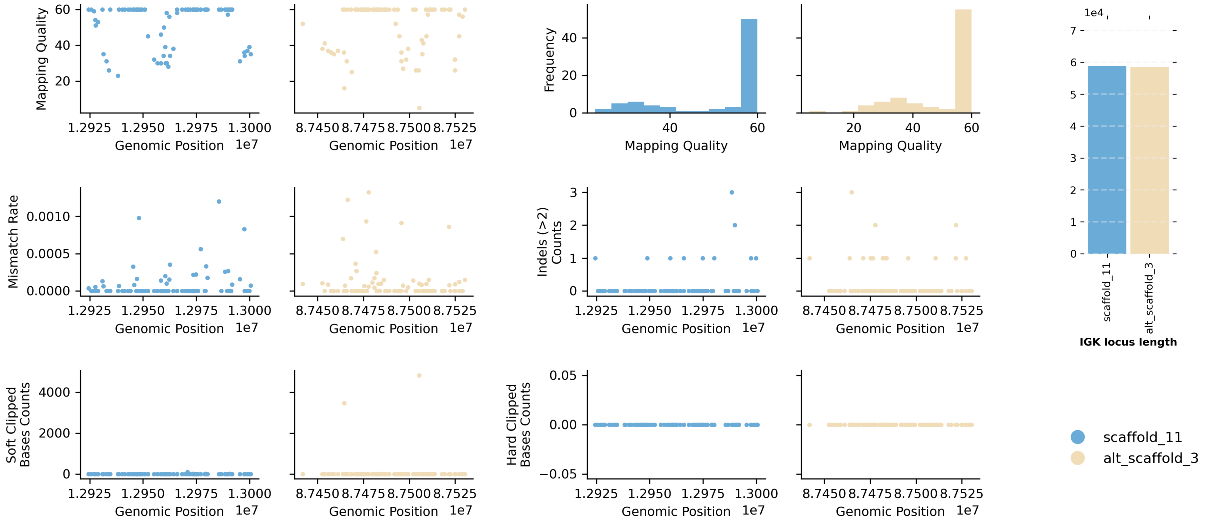


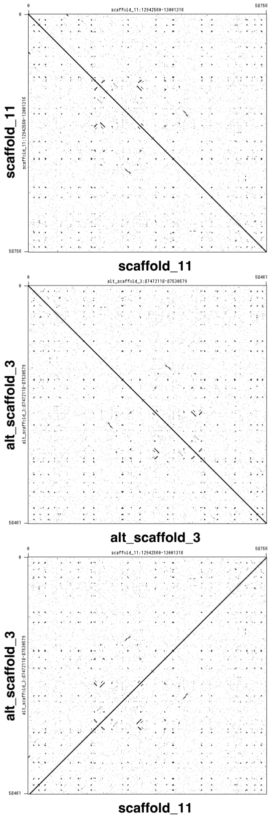


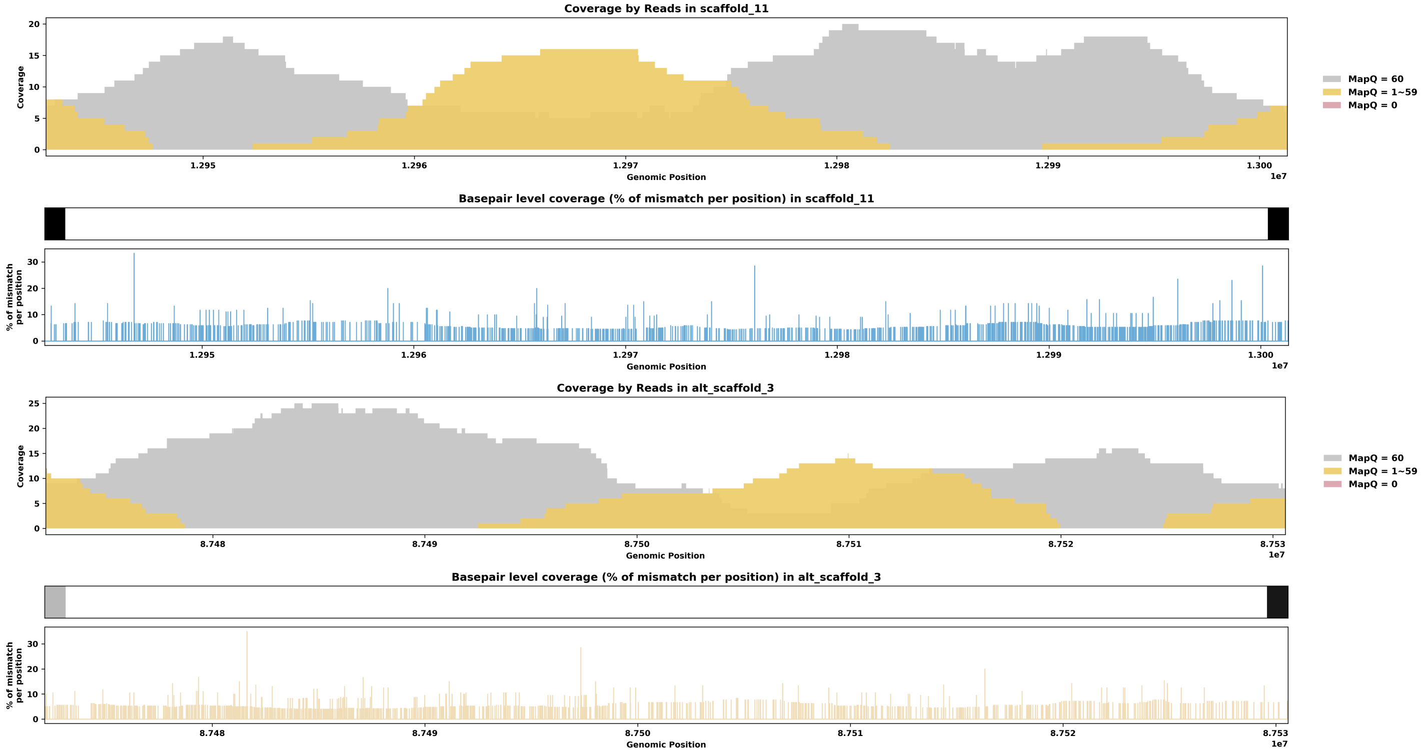

Species ID: mPumCon1

Common Name: Mountain Lion

Scientific Name: Puma concolor

Assembly Type: Haplotype Resolved

Data Source: CCGP

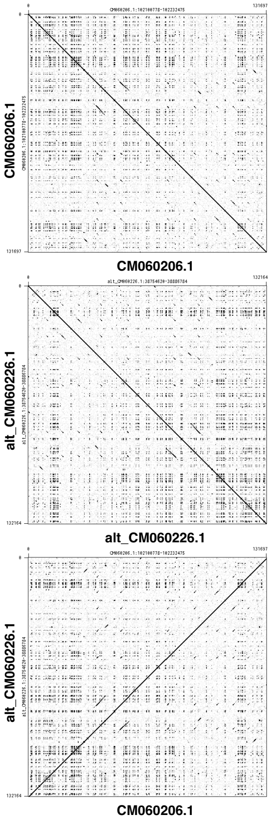


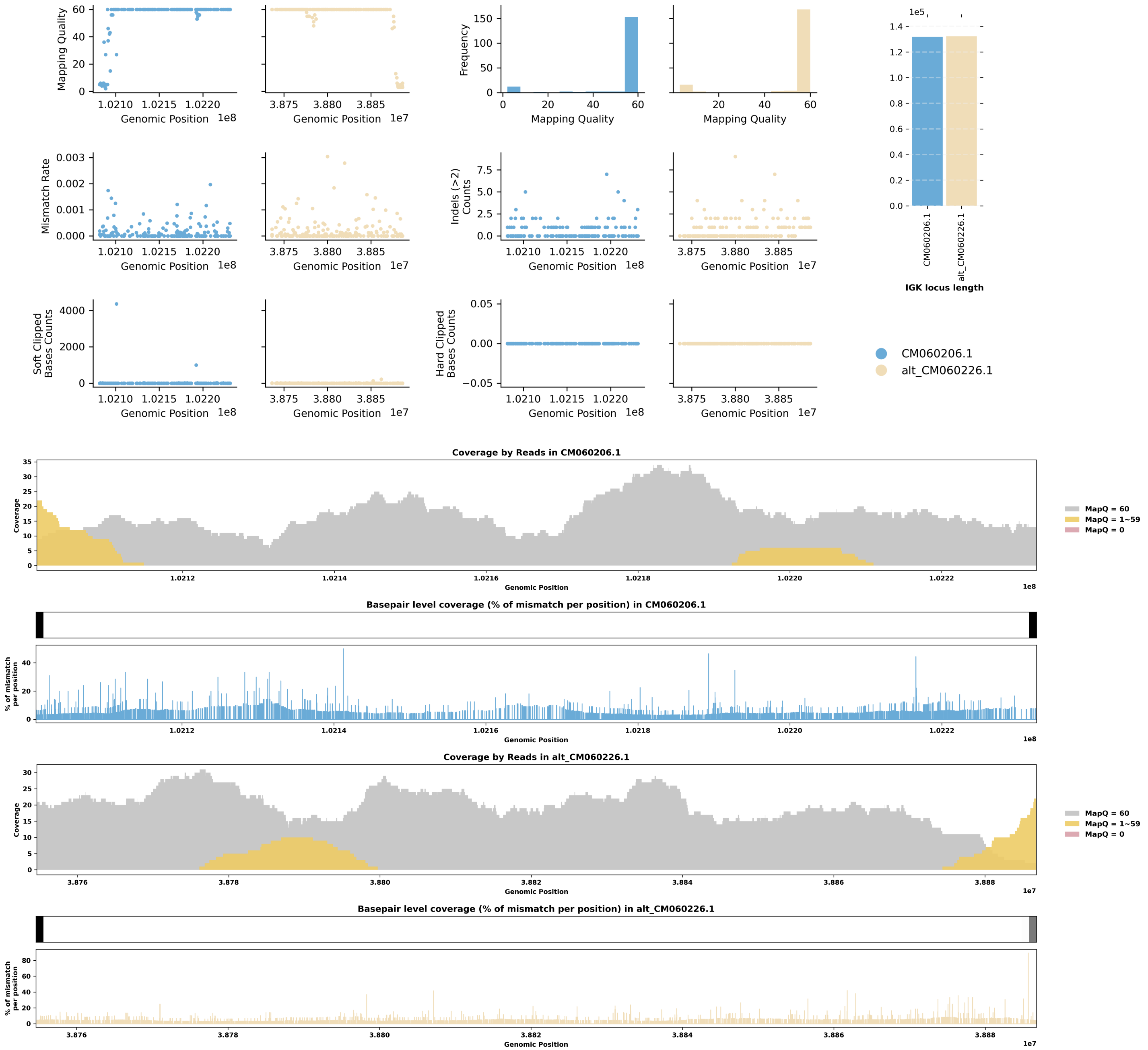

Species ID: mSorAra2

Common Name: Common shrew

Scientific Name: Sorex araneus

Assembly Type: Not Haplotype Resolved

Data Source: VGP

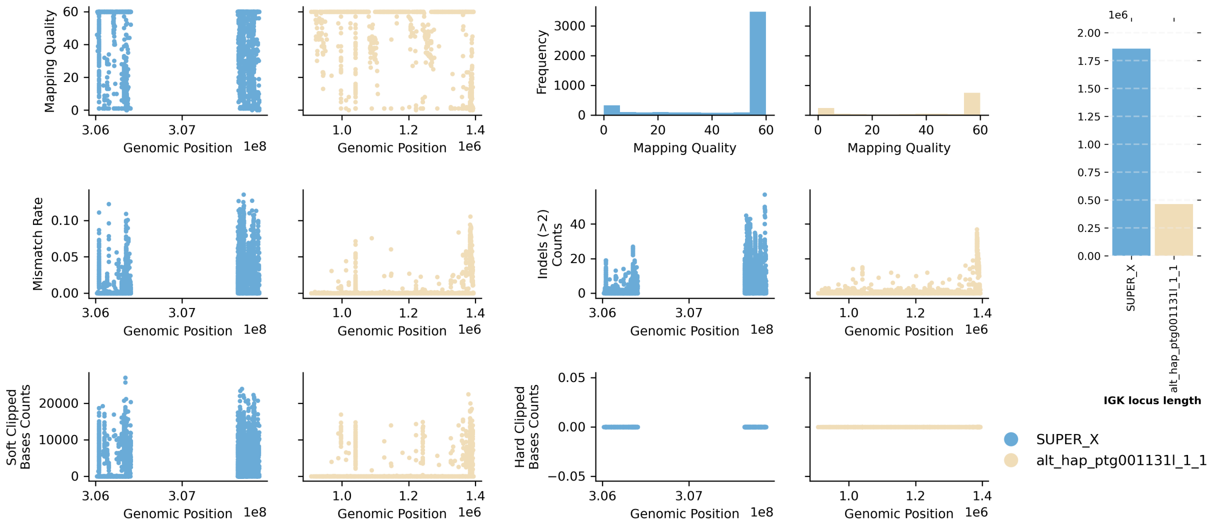


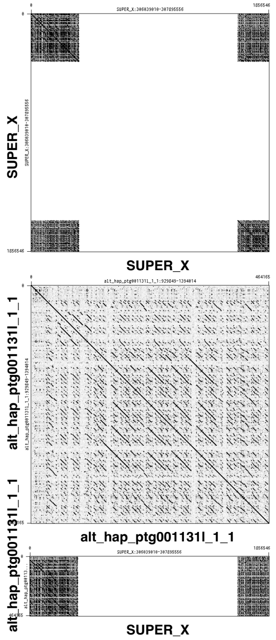


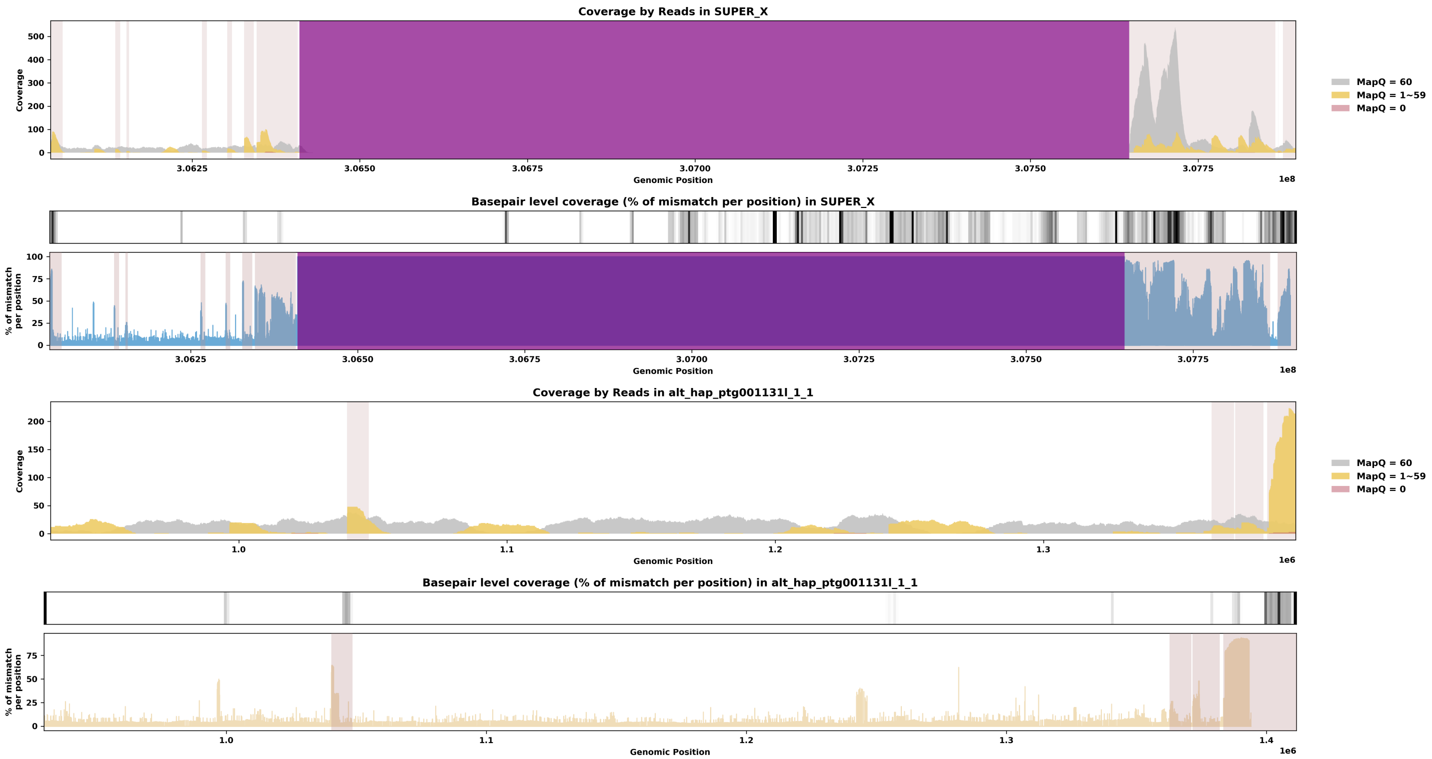

Species ID: mSteCoe1

Common Name: striped dolphin

Scientific Name: Stenella coeruleoalba

Assembly Type: Not Haplotype Resolved

Data Source: VGP

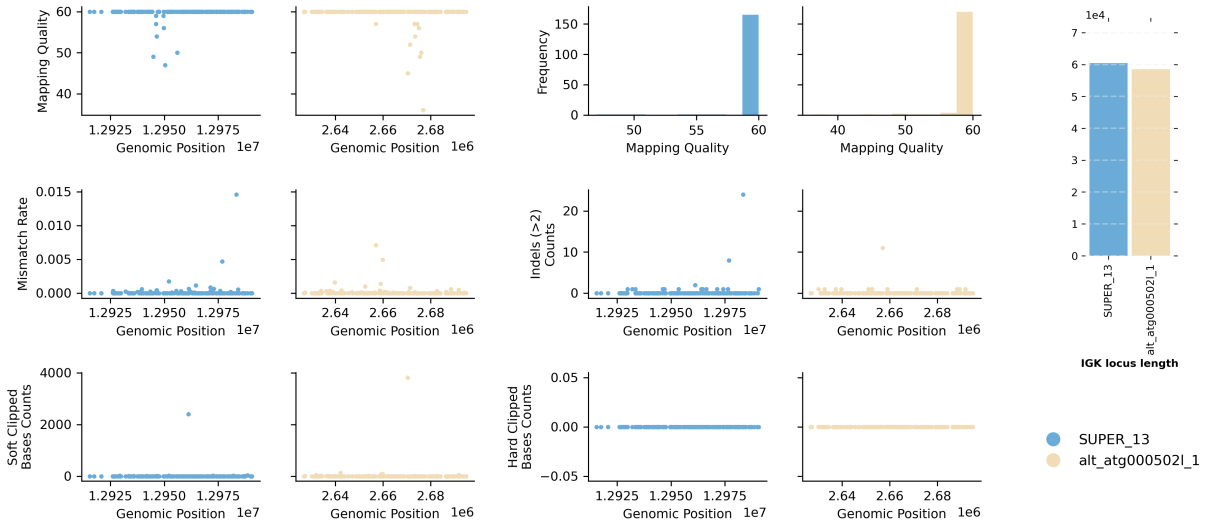


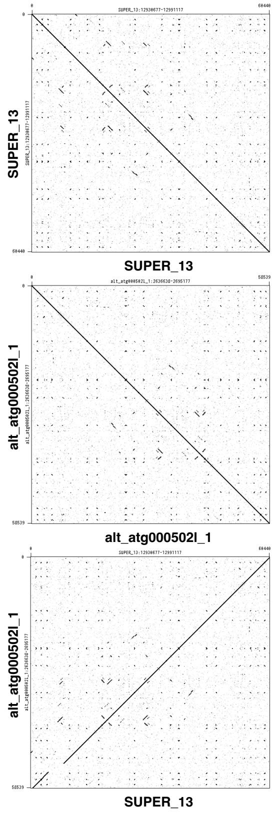


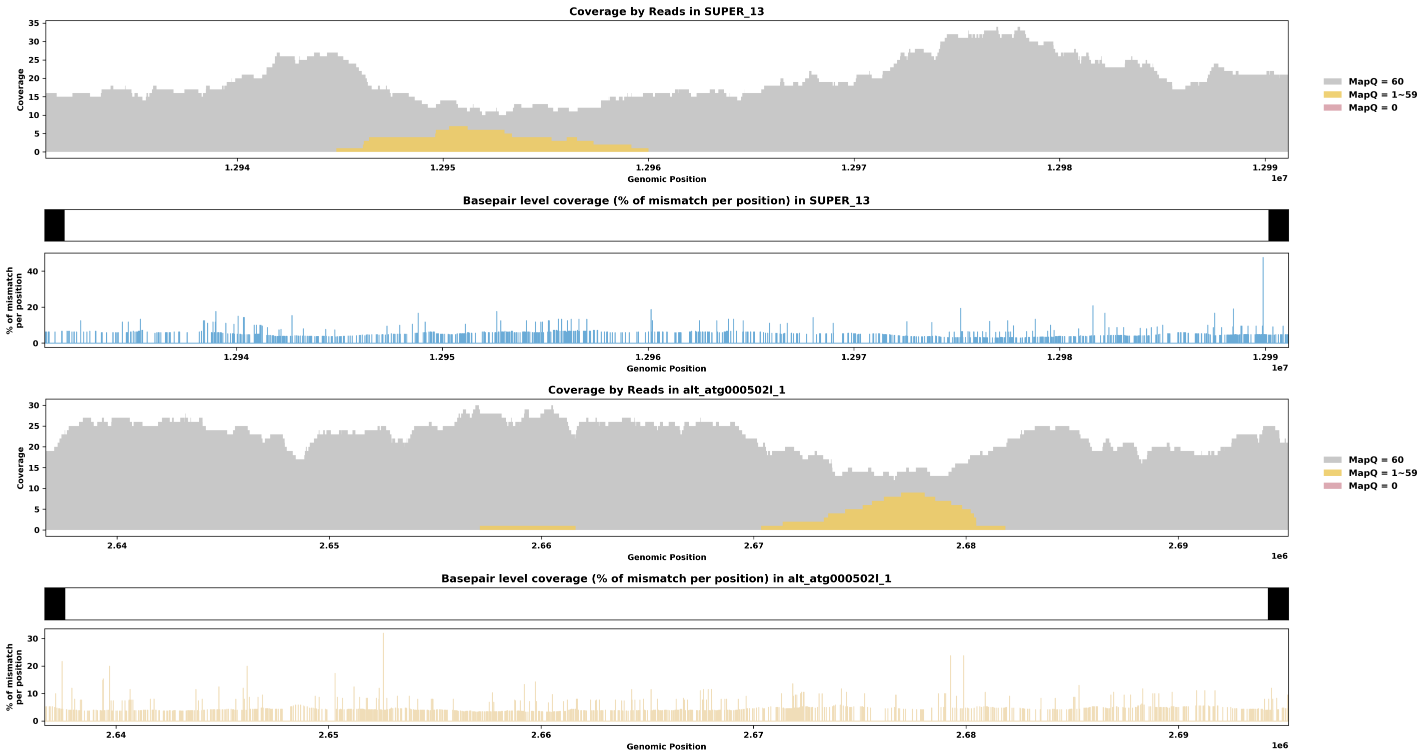

Species ID: mTalEur1

Common Name: European mole

Scientific Name: Talpa europaea

Assembly Type: Not Haplotype Resolved

Data Source: VGP

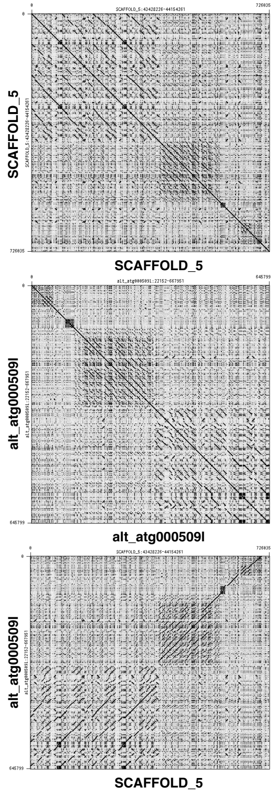


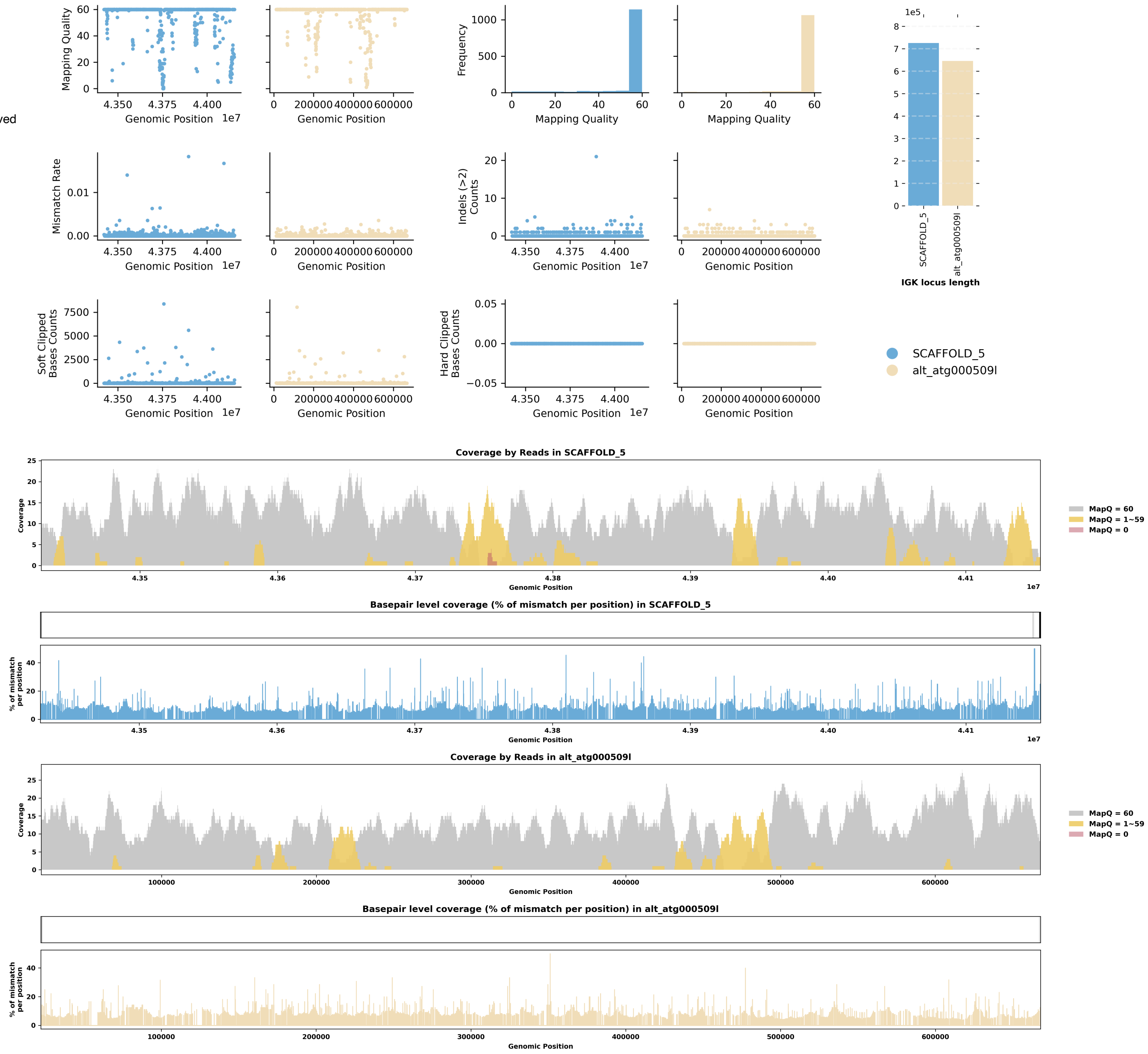

Species ID: mThoBot1

Common Name: Botta's pocket gopher

Scientific Name: Thomomys bottae

Assembly Type: Not Haplotype Resolved

Data Source: CCGP

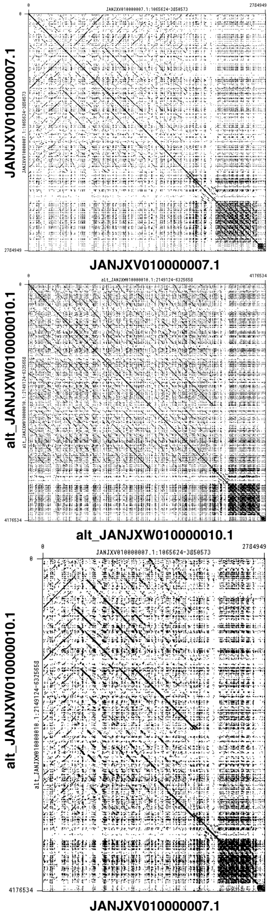


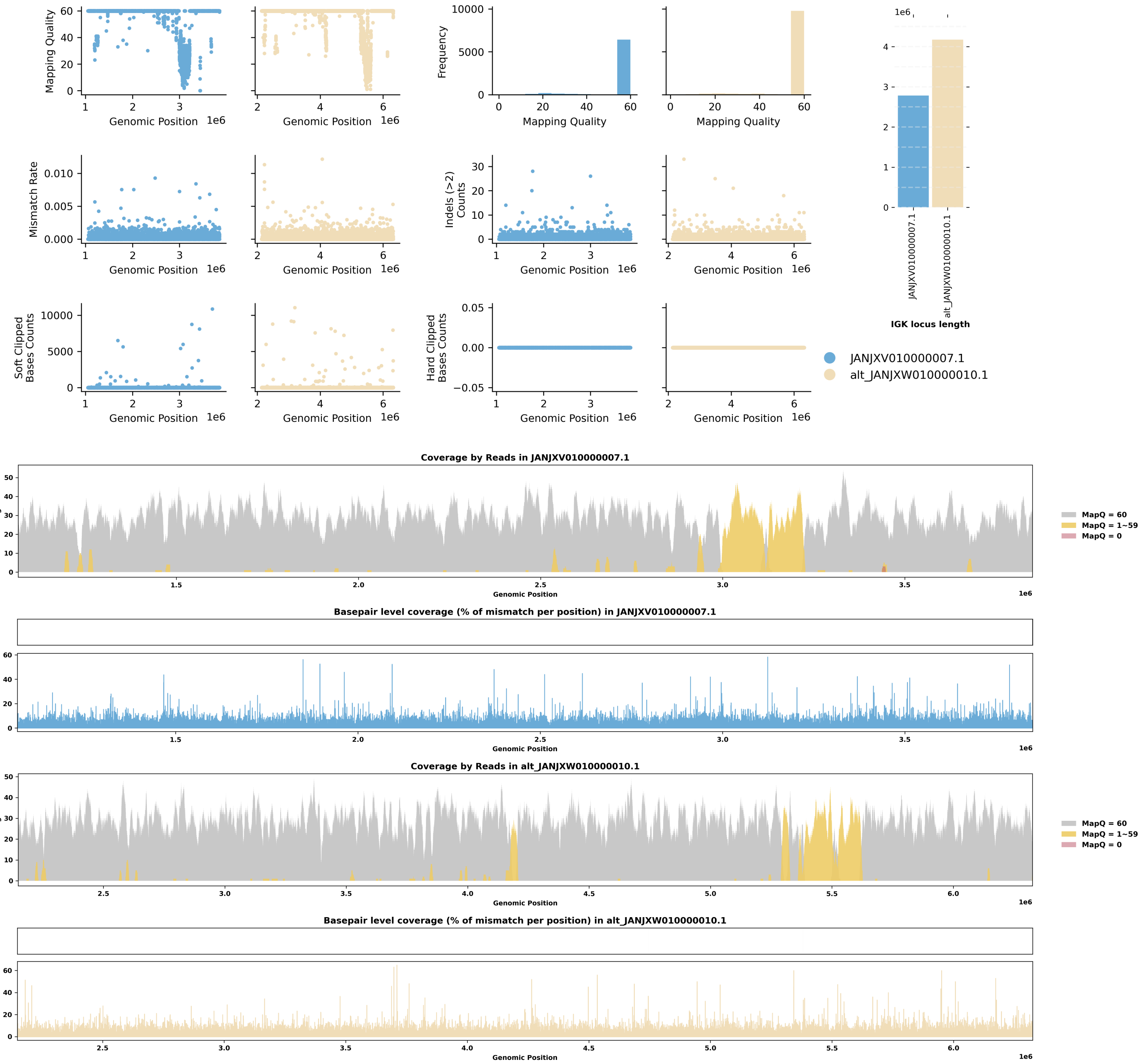

Species ID: mUrsAme1

Common Name: American black bear

Scientific Name: Ursus americanus

Assembly Type: Not Haplotype Resolved

Data Source: CCGP

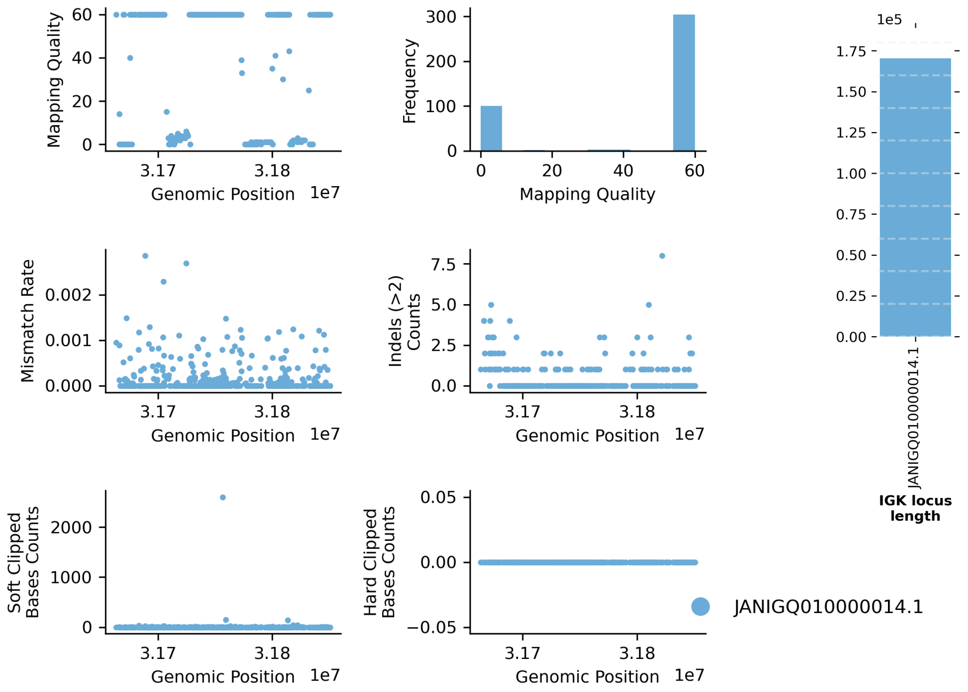


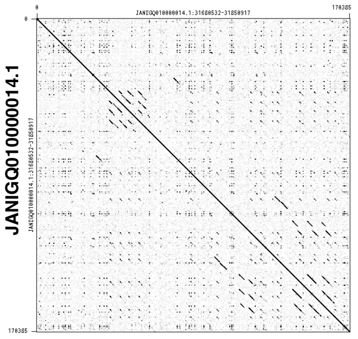


JANIGQ010000014.1

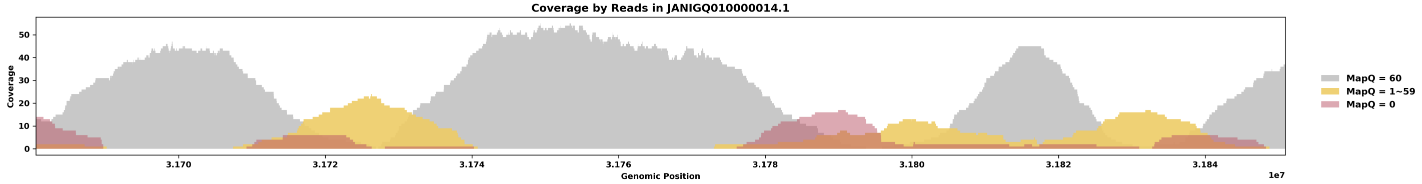


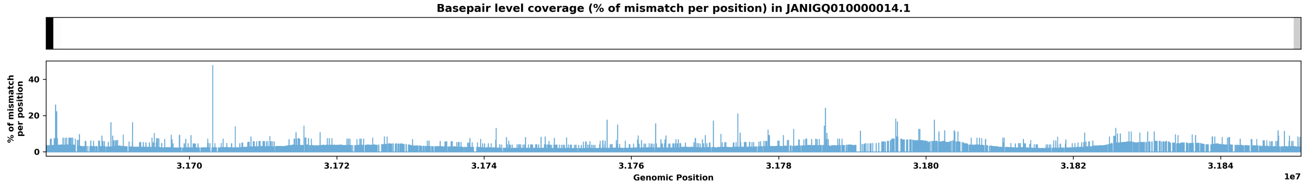

Species ID: mUrsArc2

Common Name: brown bear

Scientific Name: Ursus arctos

Assembly Type: Not Haplotype Resolved

Data Source: NCBI

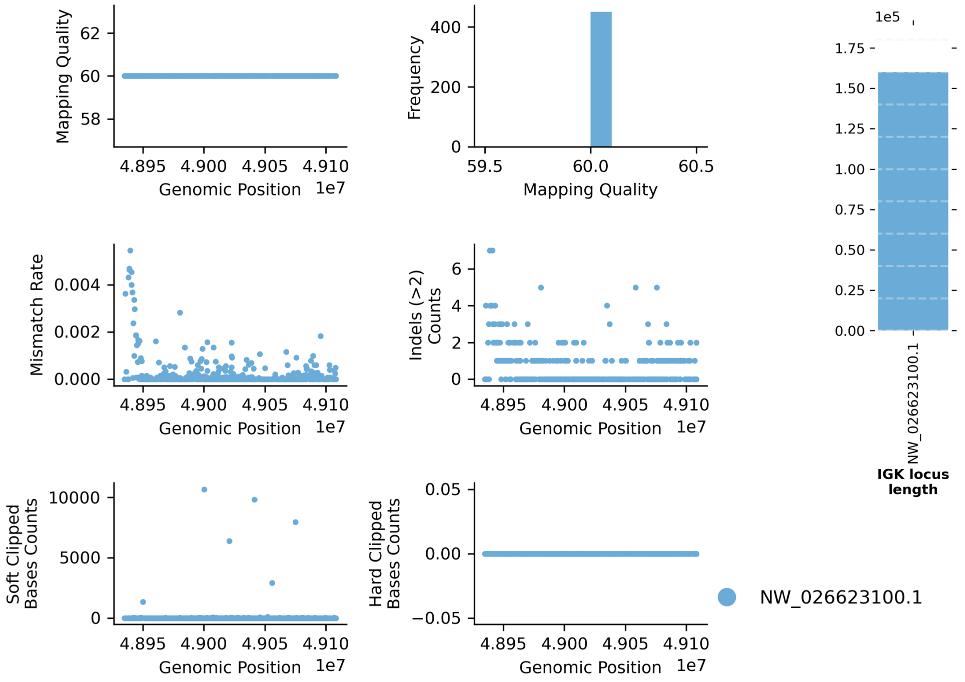


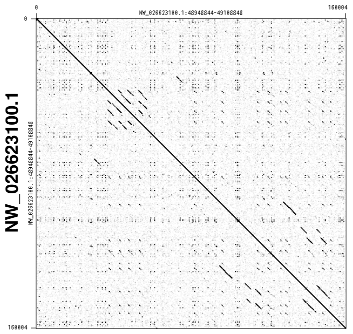


NW_026623100.1

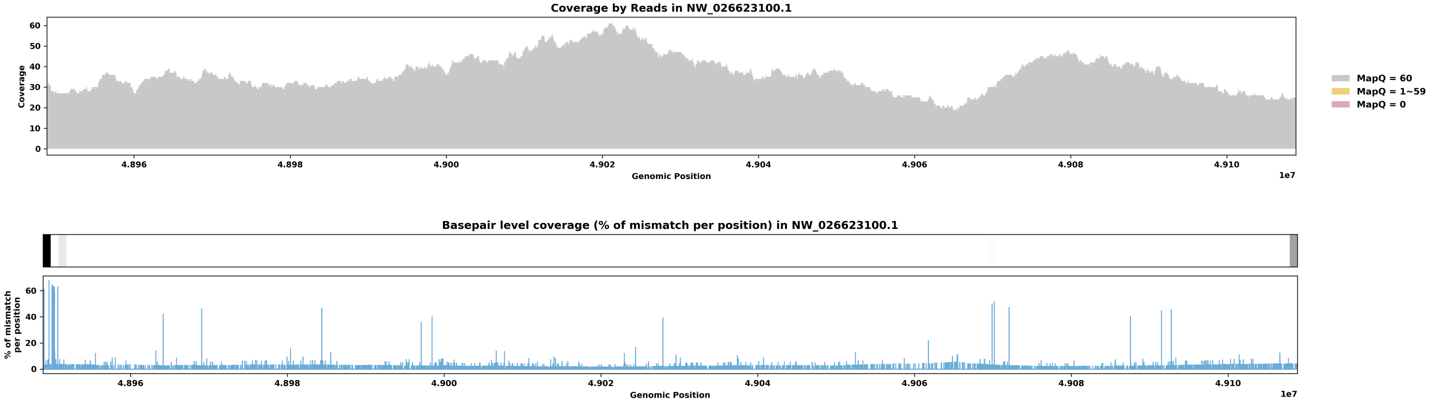

Species ID: rAllMis2

Common Name: American alligator

Scientific Name: Alligator mississippiensis

Assembly Type: Haplotype Resolved

Data Source: VGP

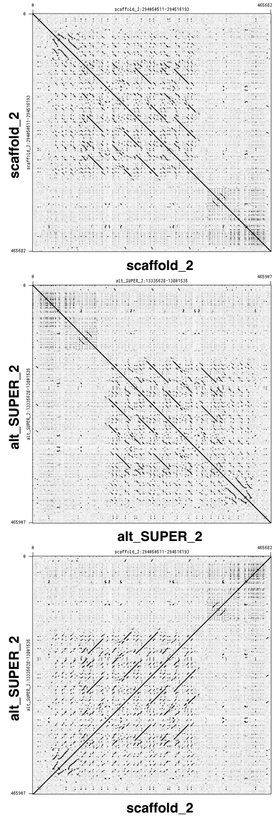


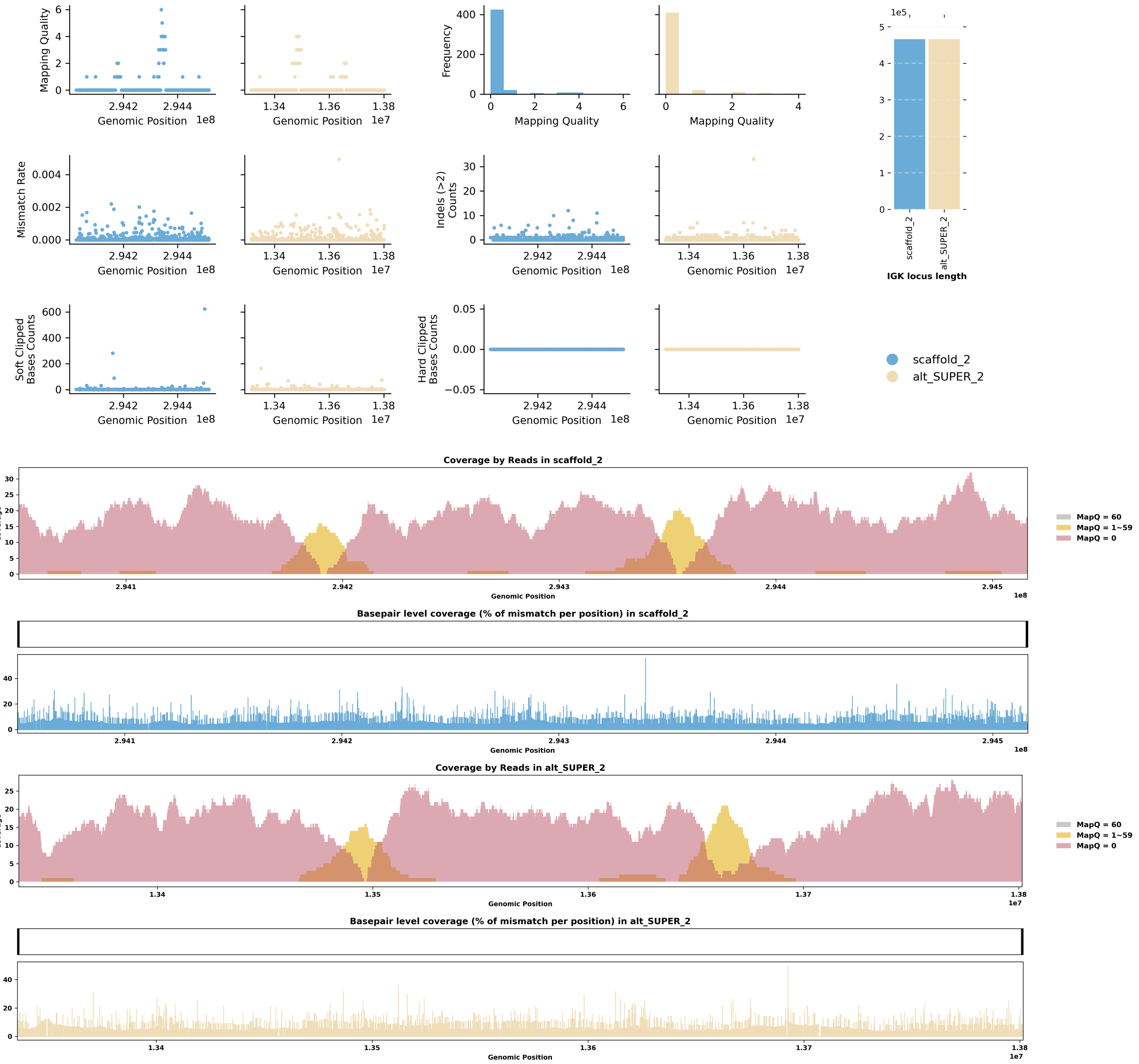

Species ID: rCarCar2

Common Name: Loggerhead turtle

Scientific Name: Caretta caretta

Assembly Type: Haplotype Resolved

Data Source: VGP

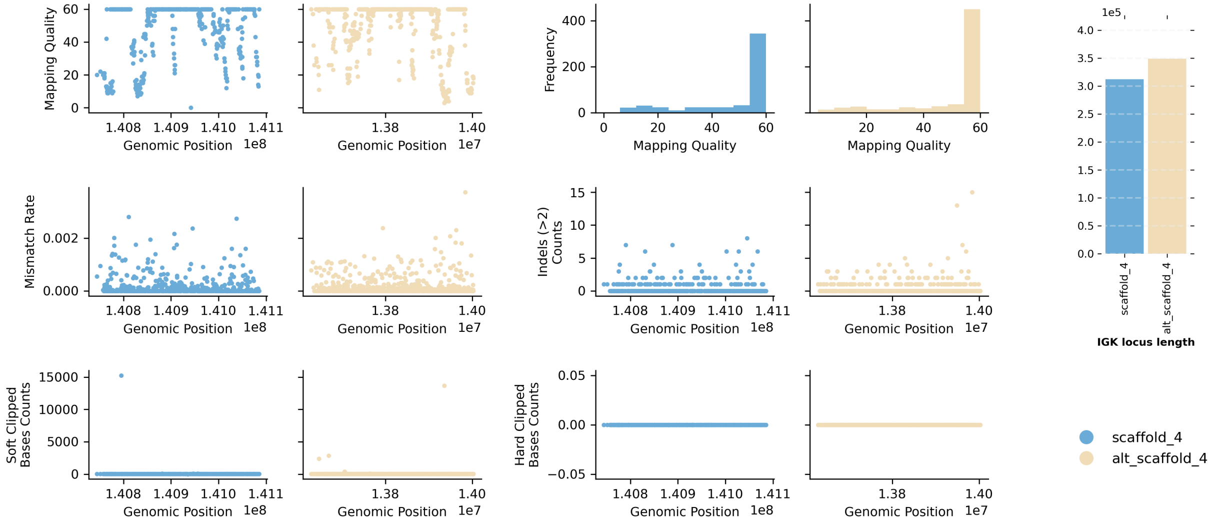


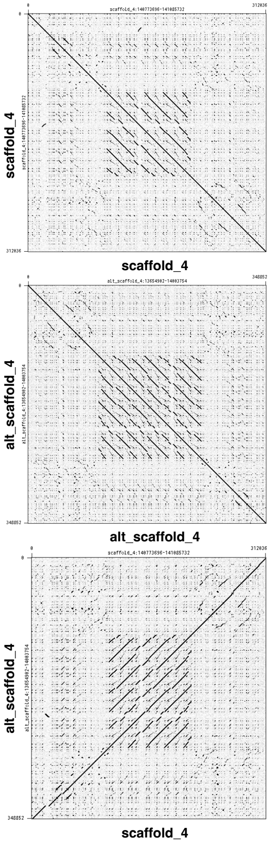


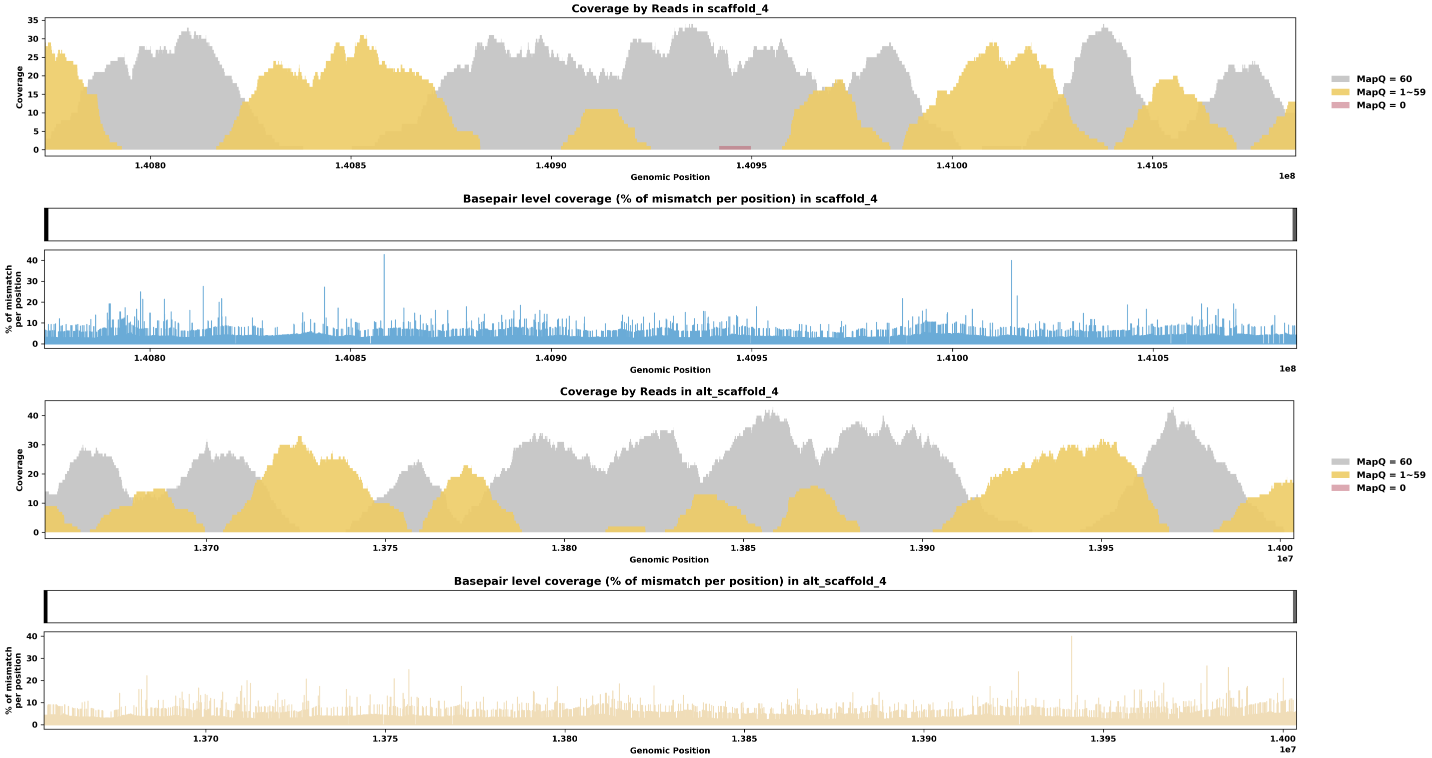

Species ID: rEmyOrb1

Common Name: European pond turtle

Scientific Name: Emys orbicularis

Assembly Type: Haplotype Resolved

Data Source: VGP

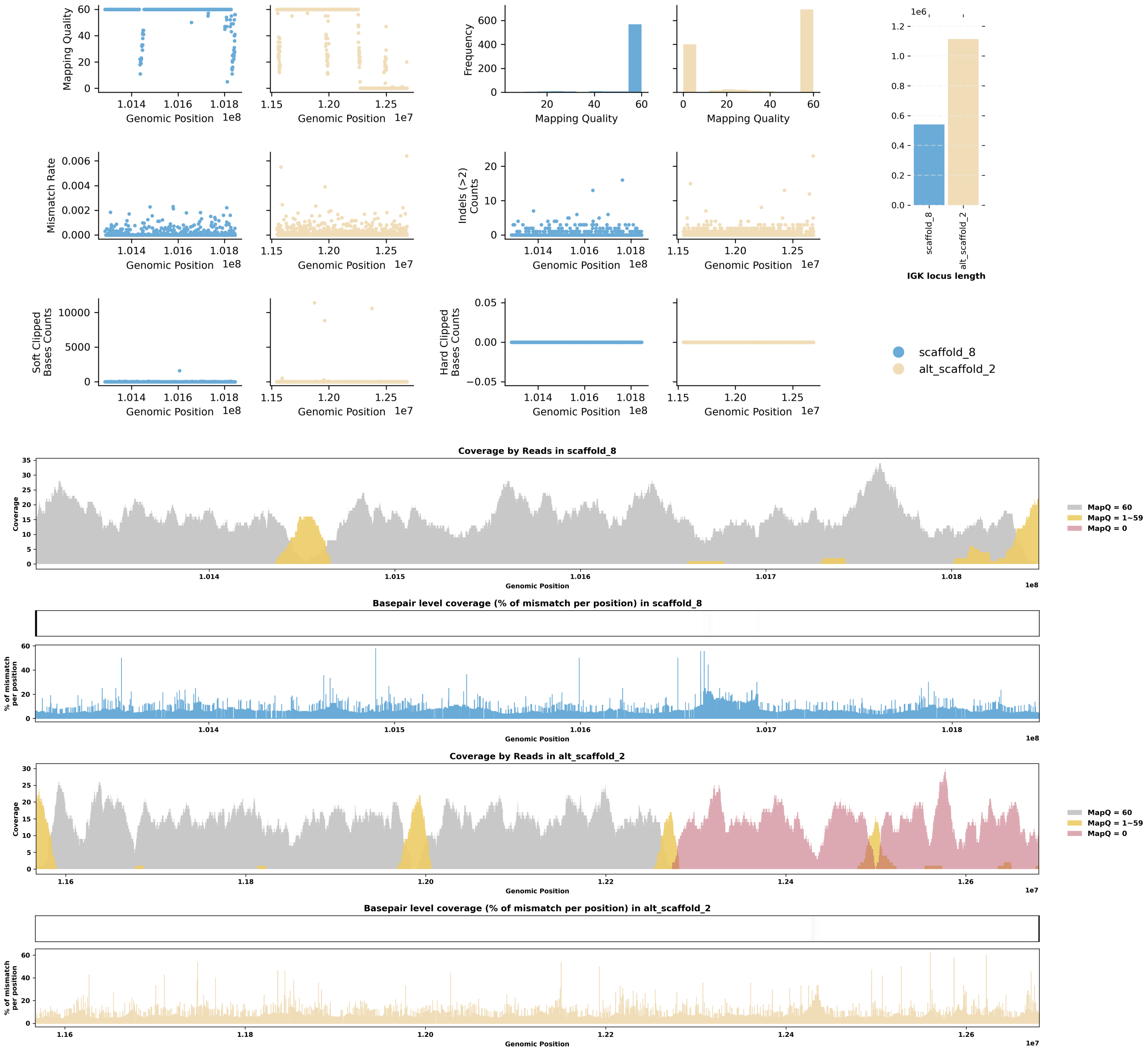


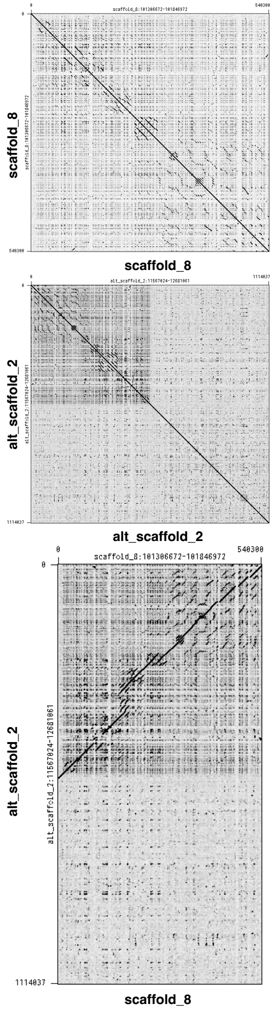

Species ID: rMalTer1

Common Name: diamondback terrapin

Scientific Name: Malaclemys terrapin

Assembly Type: Haplotype Resolved

Data Source: VGP

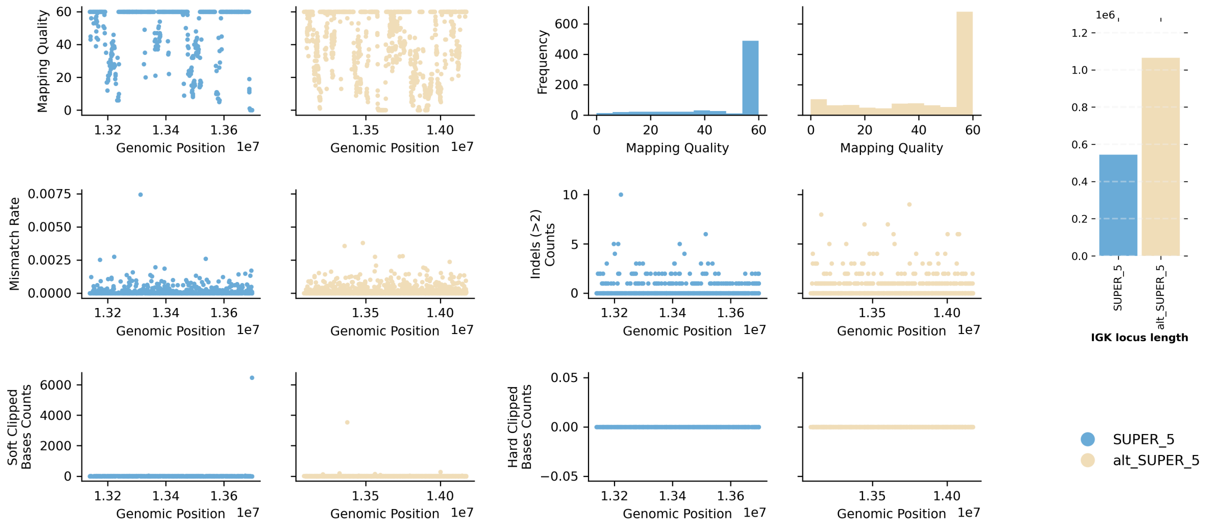


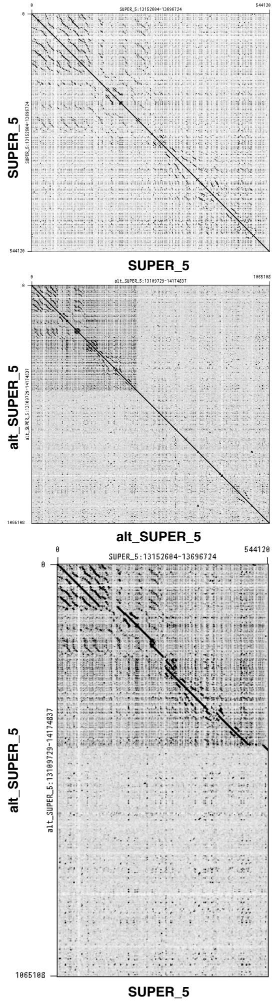


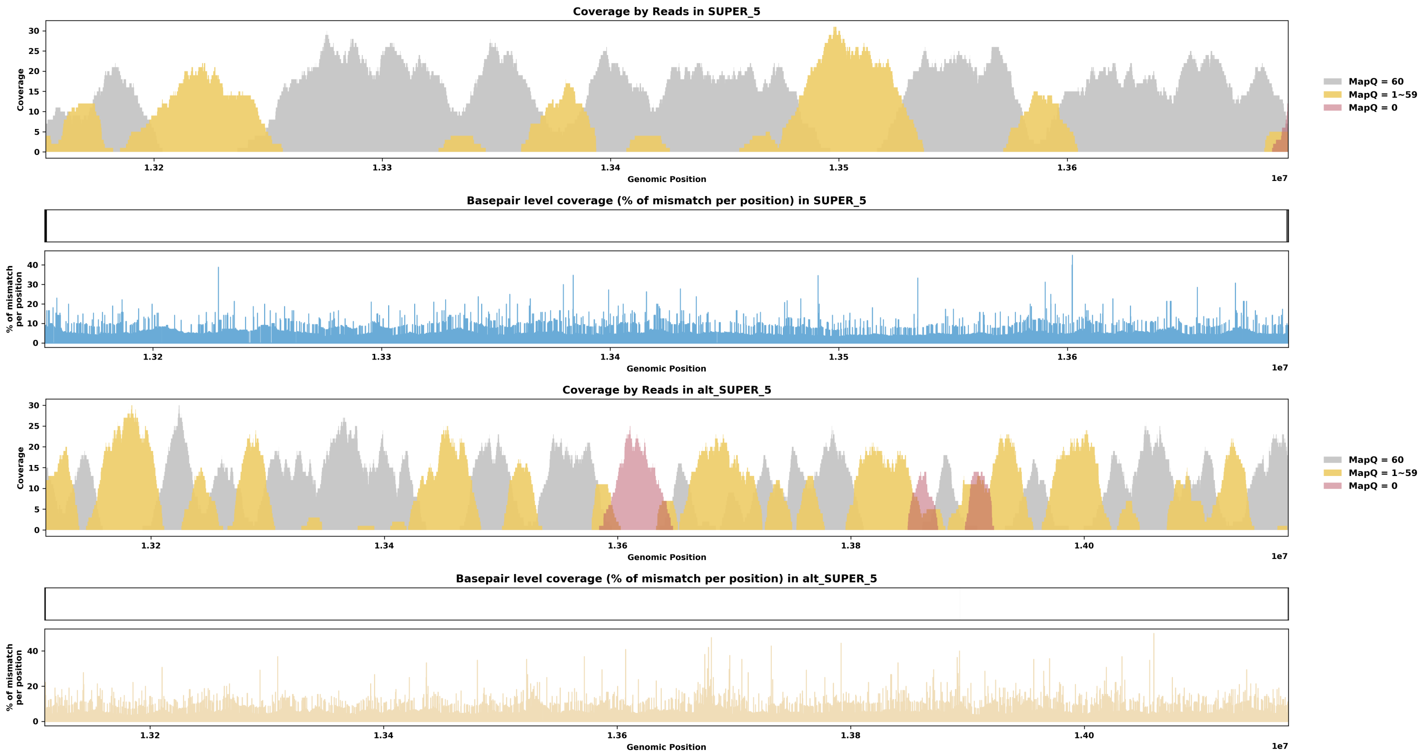
