## Supplementary material for "Assessing Assembly Errors in Immunoglobulin Loci: A Comprehensive Evaluation of Long-read Genome Assemblies Across Vertebrates": IGL Loci Evaluations

Table of Contents

1. mApoSyl1 - Apodemus sylvaticus - Wood mouse
2. mBalAcu1 - Balaenoptera acutorostrata - Minke whale
3. mCamDro1 - Camelus dromedarius - Dromedary
4. mCanLor1 - Canis lupus - Greenland Wolf
5. mCanLor2 - Canis lupus - Greenland Wolf
6. mCerEla1 - Cervus elaphus - Red Deer
7. mCorTow1.0 - Corynorhinus townsendii - Townsend's Big-eared Bat
8. mCynVol1 - Cynocephalus volans - Philippine flying lemur
9. mDasNov1 - Dasypus novemcinctus - Nine-banded armadillo
10. mDelDel1 - Delphinus delphis - Saddleback dolphin
11. mDicBic1 - Diceros bicornis - Black rhinoceros
12. mEleMax1 - Elephas maximus - Asiatic Elephant
13. mEptNil1 - Eptesicus nilssonii - Northern bat
14. mEriEur2 - Erinaceus europaeus - Western European hedgehog
15. mEscRob2 - Eschrichtius robustus - Grey whale
16. mEubGla1 - Eubalaena glacialis - North Atlantic right whale
17. mGloMel1 - Globicephala melas - Long-finned pilot whale
18. mGorGor1 - Gorilla gorilla - Gorilla
19. mHetBru1 - Heterohyrax brucei - Yellow-spotted hyrax
20. mHipAmp2 - Hippopotamus amphibius kiboko - Hippopotamus
21. mHypAmp2 - Hyperoodon ampullatus - Northern bottlenose whale
22. mLagAlb1 - Lagenorhynchus albirostris - White-beaked dolphin
23. mLemCat1 - Lemur catta - Ring-tailed lemur
24. mLynRuf1 - Lynx rufus - Bobcat
25. mMacEug1 - Macropus eugenii - Tammar wallaby
26. mManPen7 - Manis pentadactyla - Chinese pangolin
27. mMarMar1 - Martes martes - European pine marten
28. mMelMel3 - Meles meles - European badger
29. mMesDen1 - Mesoplodon densirostris - Blainville's beaked whale
30. mMicCal1.0 - Microtus californicus - California Vole
31. mMicMin1 - Micromys minutus - European harvest mouse
32. mMirAng1 - Mirounga angustirostris - Northern Elephant Seal
33. mMonDom1 - Monodelphis domestica - Gray short-tailed opossum
34. mMunRee1 - Muntiacus reevesi - Reeves' muntjac
35. mMusAve1 - Muscardinus avellanarius - Hazel dormouse
36. mMusLut2 - Mustela lutreola - European mink
37. mMusNiv1 - Mustela nivalis - Least weasel
38. mMyoDau2 - Myotis daubentonii - Daubenton's bat
39. mMyoYum1.0 - Myotis yumanensis - Yuma myotis
40. mNeoNeb1 - Neofelis nebulosa - Clouded Leopard
41. mNycCou1 - Nycticebus coucang - Slow loris
42. mOrcOrc1 - Orcinus orca - Killer whale
43. mOryCun1 - Oryctolagus cuniculus - Rabbit
44. mPanPan1 - Pan paniscus - Bonobo
45. mPerMan1 - Peromyscus maniculatus - Deer mouse
46. mPhoPho1 - Phocoena phocoena - Harbor porpoise
47. mPipPyg2 - Pipistrellus pygmaeus - Soprano pipistrelle
48. mPonAbe1 - Pongo abelii - Sumatran orangutan
49. mPonPyg2 - Pongo pygmaeus - Bornean orangutan
50. mPseCra1 - Pseudorca crassidens - False killer whale
51. mPumCon1.1 - Puma concolor - Mountain Lion
52. mSorAra2/1 - Sorex araneus - Common shrew
53. mSteCoe1 - Stenella coeruleoalba - Striped dolphin
54. mTalEur1 - Talpa europaea - European mole
55. mUrsAme1 - Ursus americanus - American black bear
56. mUrsArc2 - Ursus arctos - Brown bear
57. mVesMur1 - Vespertilio murinus - Particolored bat
58. rAllMis2 - Alligator mississippiensis - American alligator
59. rCarCar2 - Caretta caretta - Loggerhead turtle
60. rEmyOrb1 - Emys orbicularis - European pond turtle
61. rMalTer1 - Malaclemys terrapin - Diamondback terrapin

Note: This supplementary material only displays alternate IG loci if they are longer than one-quarter of the corresponding primary IG locus length. Loci shorter than this threshold are typically too fragmented to be shown. 18 species had alternate IGL loci shorter than this threshold and are not shown.

Species ID: mApoSyl1

Common Name: wood mouse

Scientific Name: Apodemus sylvaticus

Assembly Type: Not Haplotype Resolved

Data Source: VGP

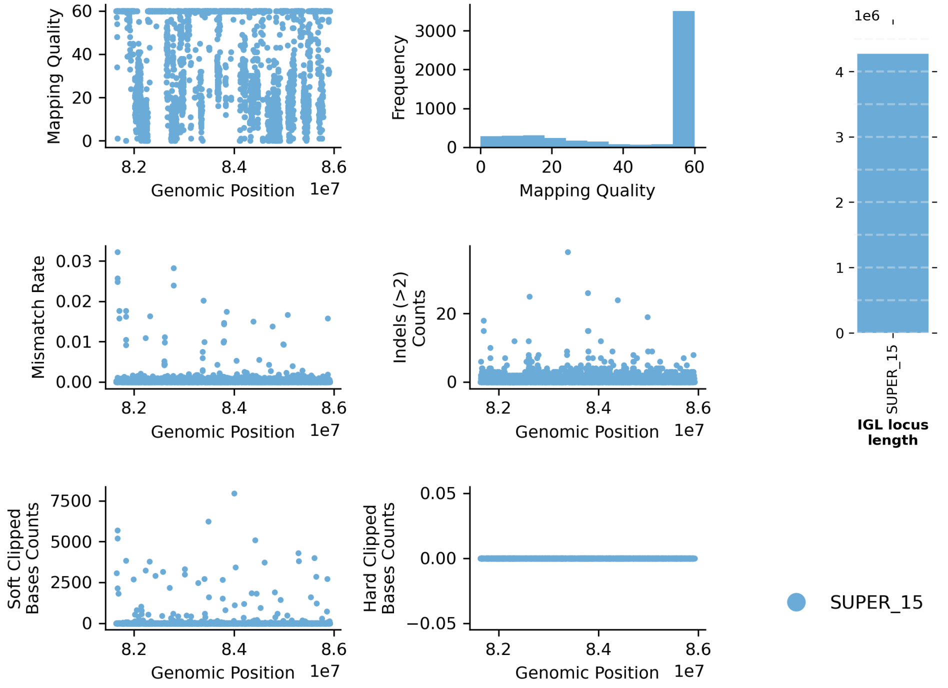


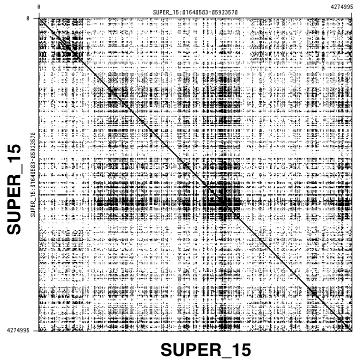


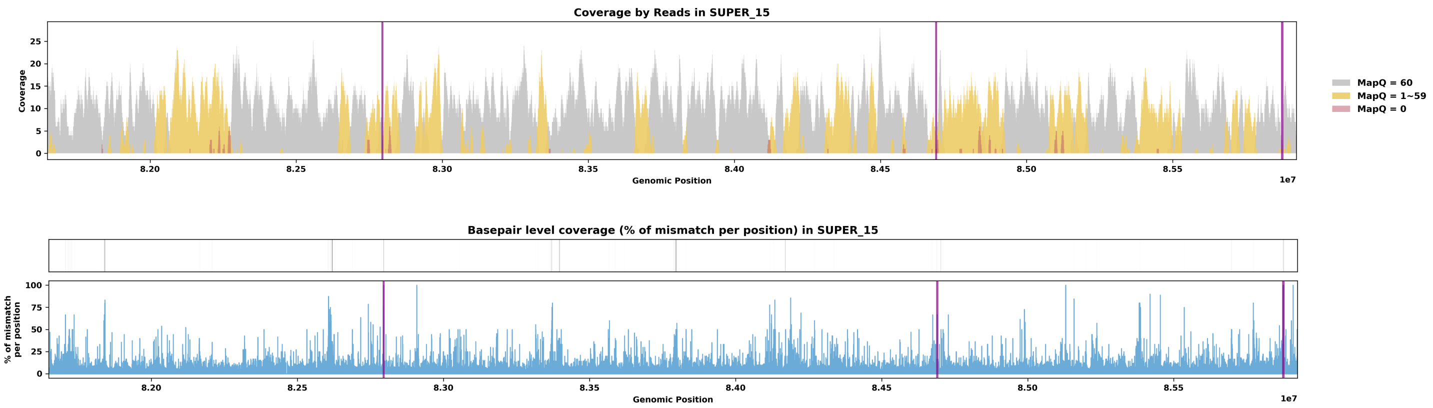

Species ID: mBalAcu1

Common Name: minke whale

Scientific Name: Balaenoptera_acutorostrata

Assembly Type: Not Haplotype Resolved

Data Source: VGP

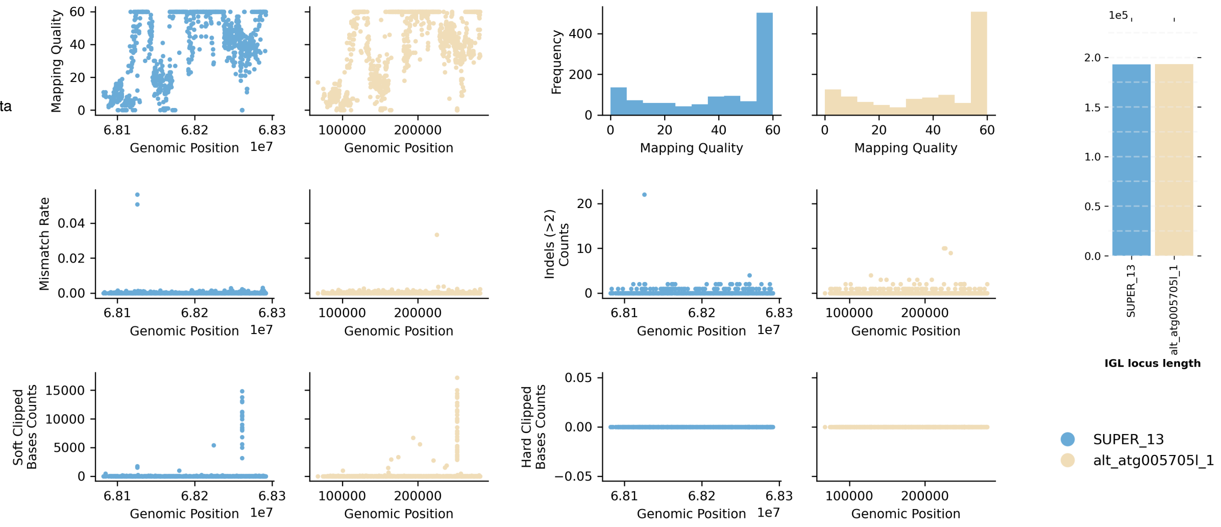


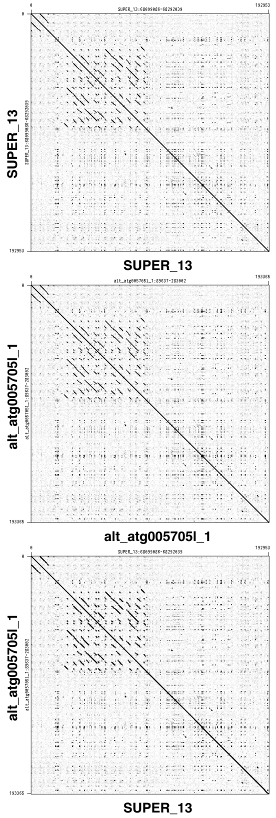


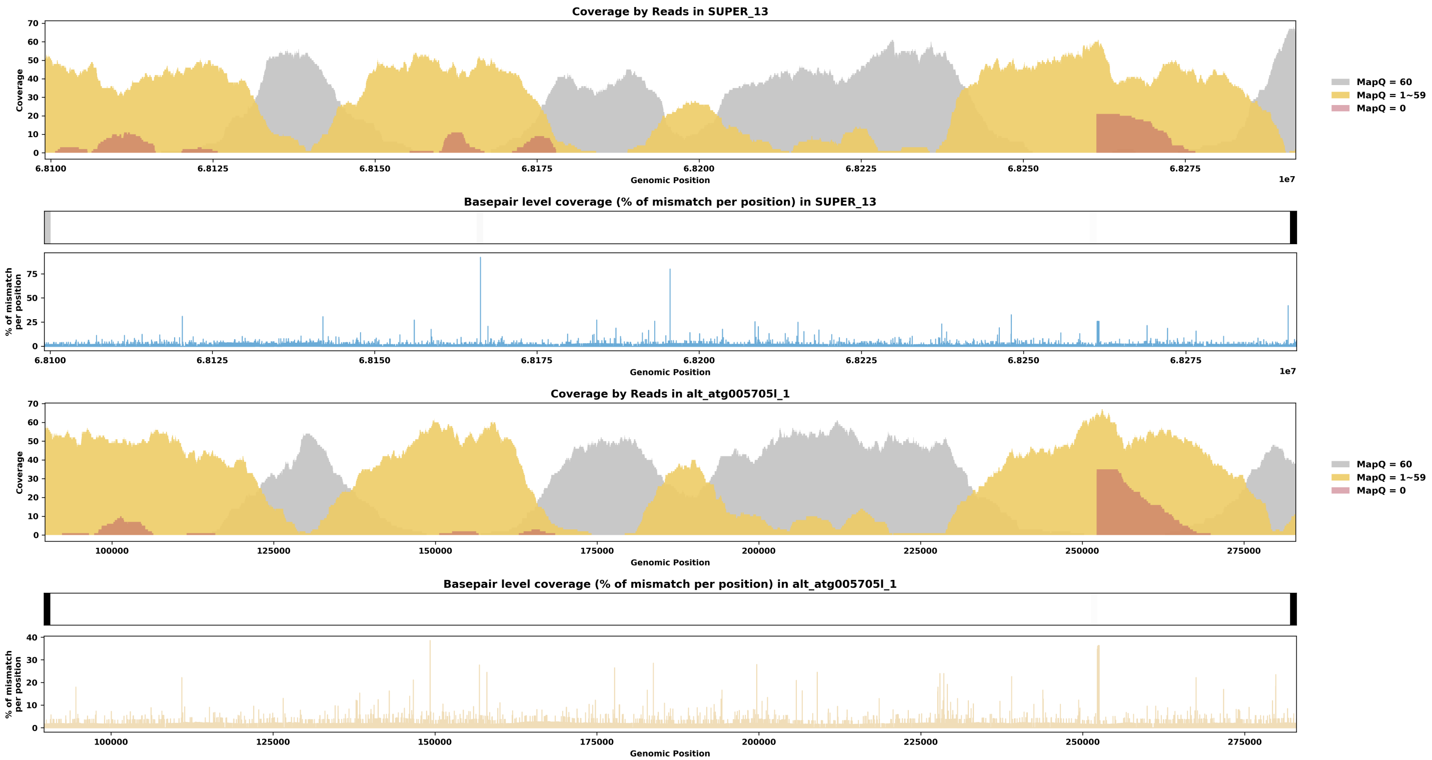

Species ID: mCamDro1

Common Name: dromedary

Scientific Name: Camelus dromedarius

Assembly Type: Haplotype Resolved

Data Source: VGP

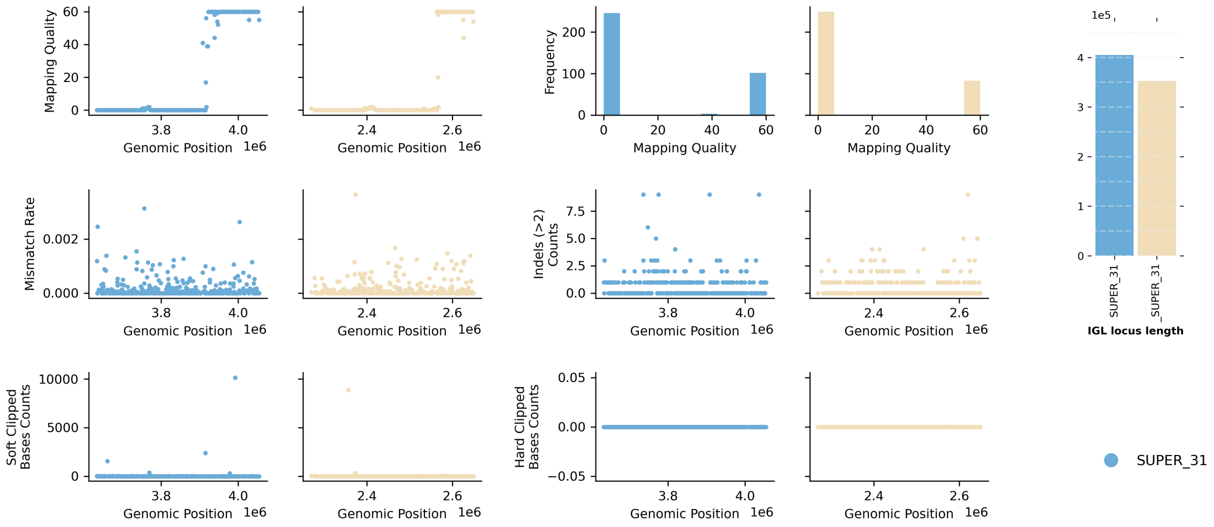


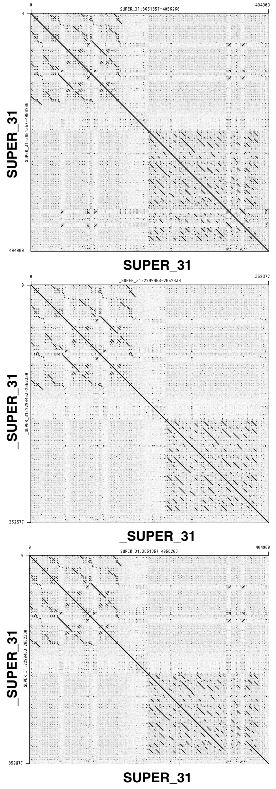


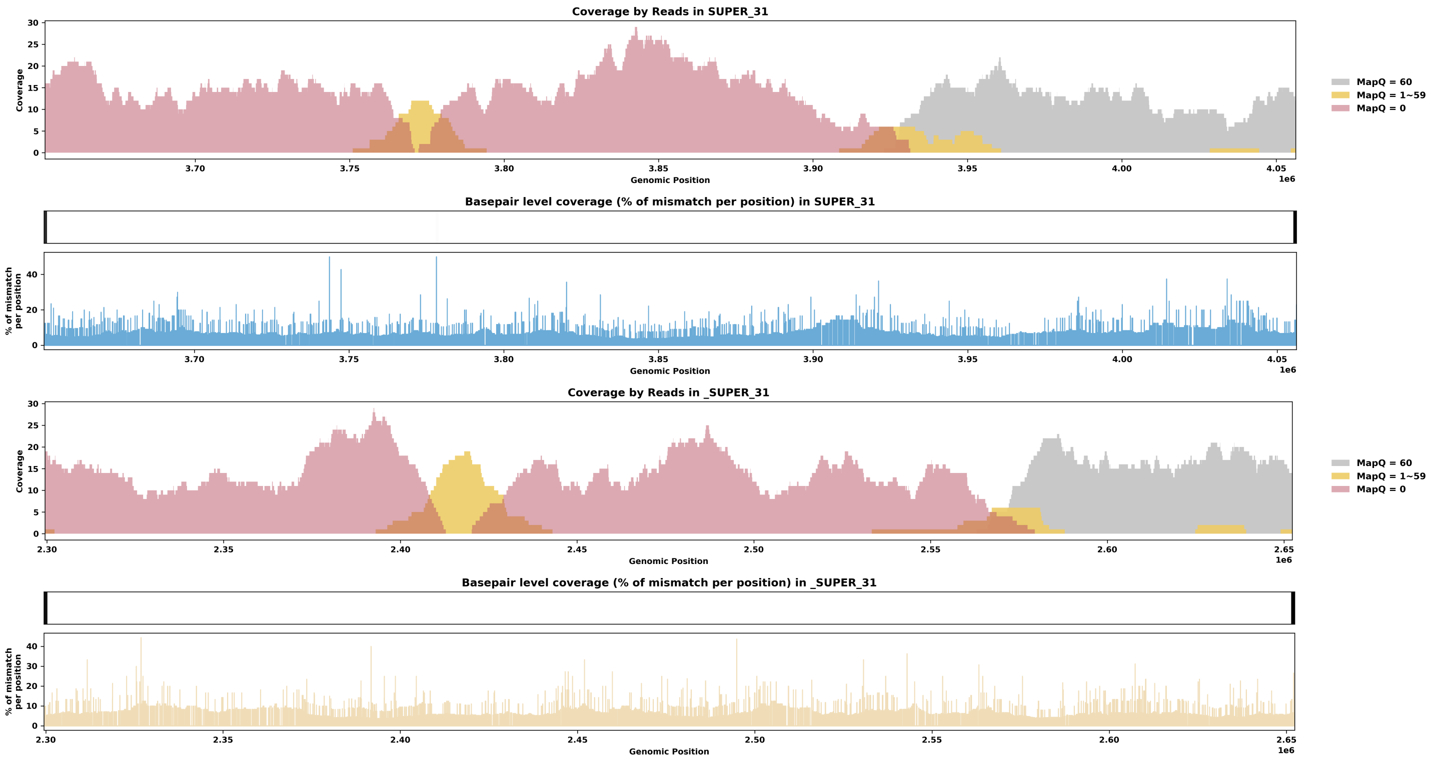

Species ID: mCanLor1

Common Name: Greenland Wolf

Scientific Name: Canis_lupus

Assembly Type: Not Haplotype Resolved

Data Source: VGP

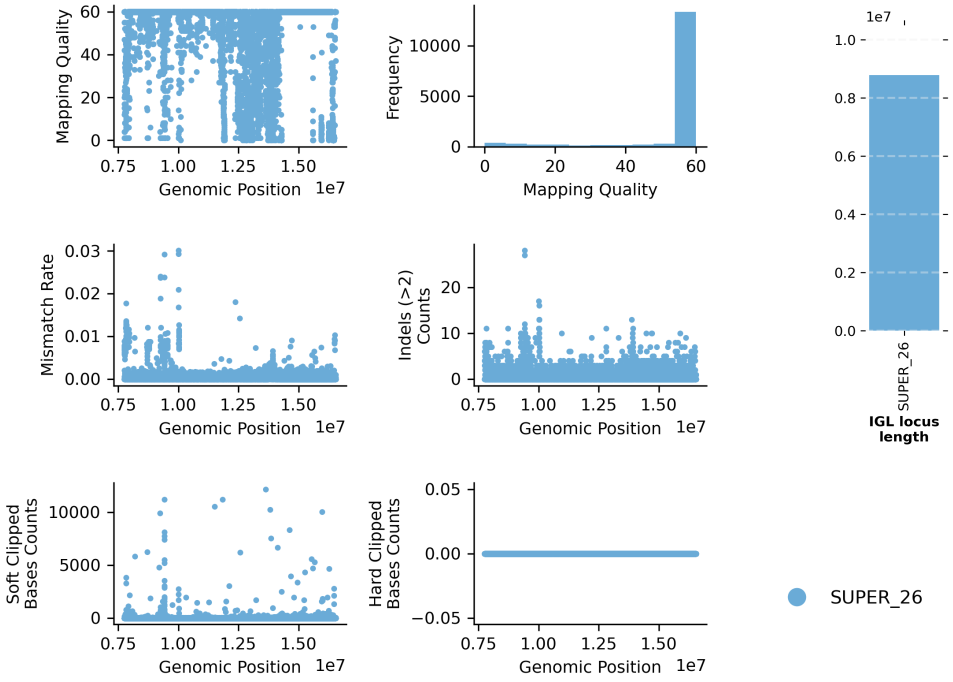


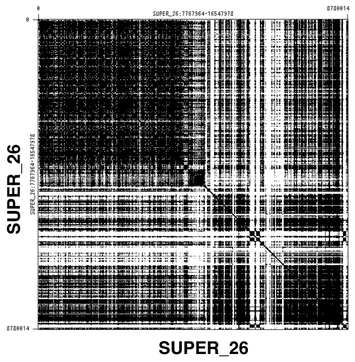


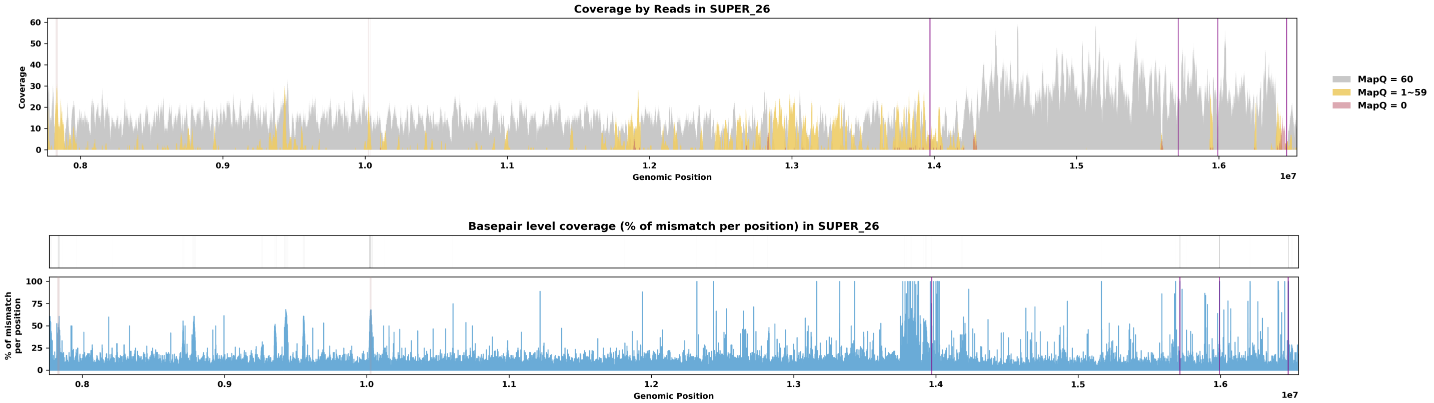

Species ID: mCanLor2

Common Name: Greenland Wolf

Scientific Name: Canis_lupus

Assembly Type: Haplotype Resolved

Data Source: VGP

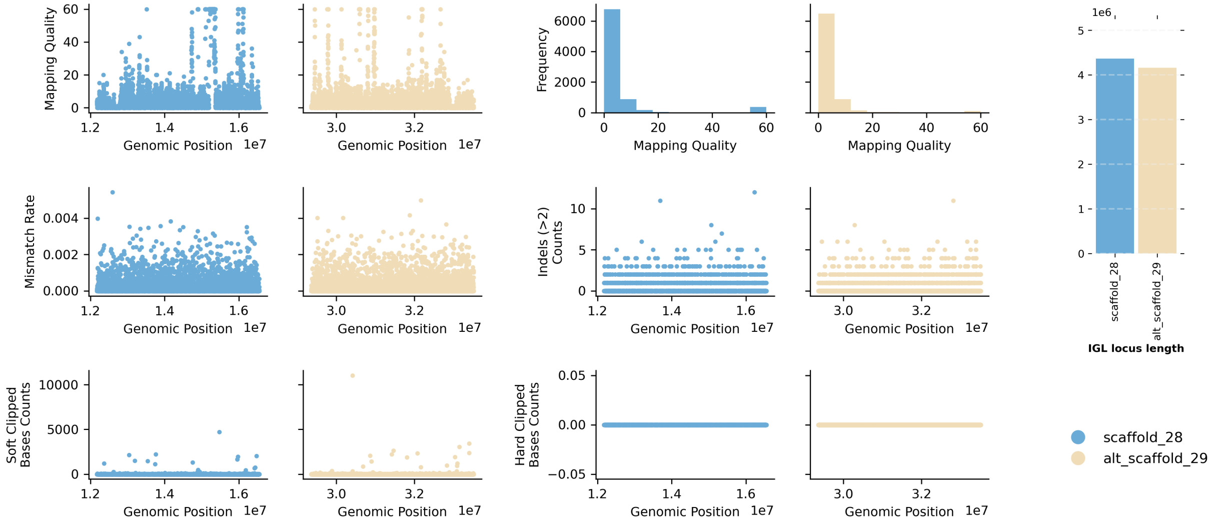


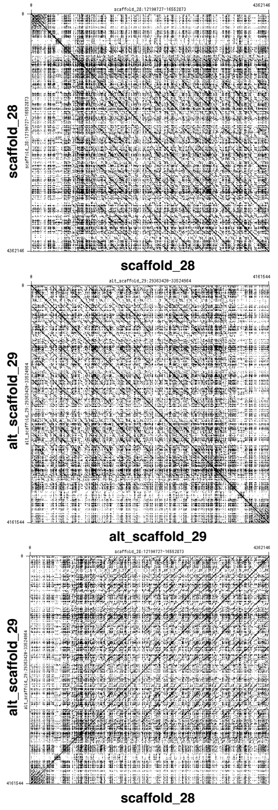


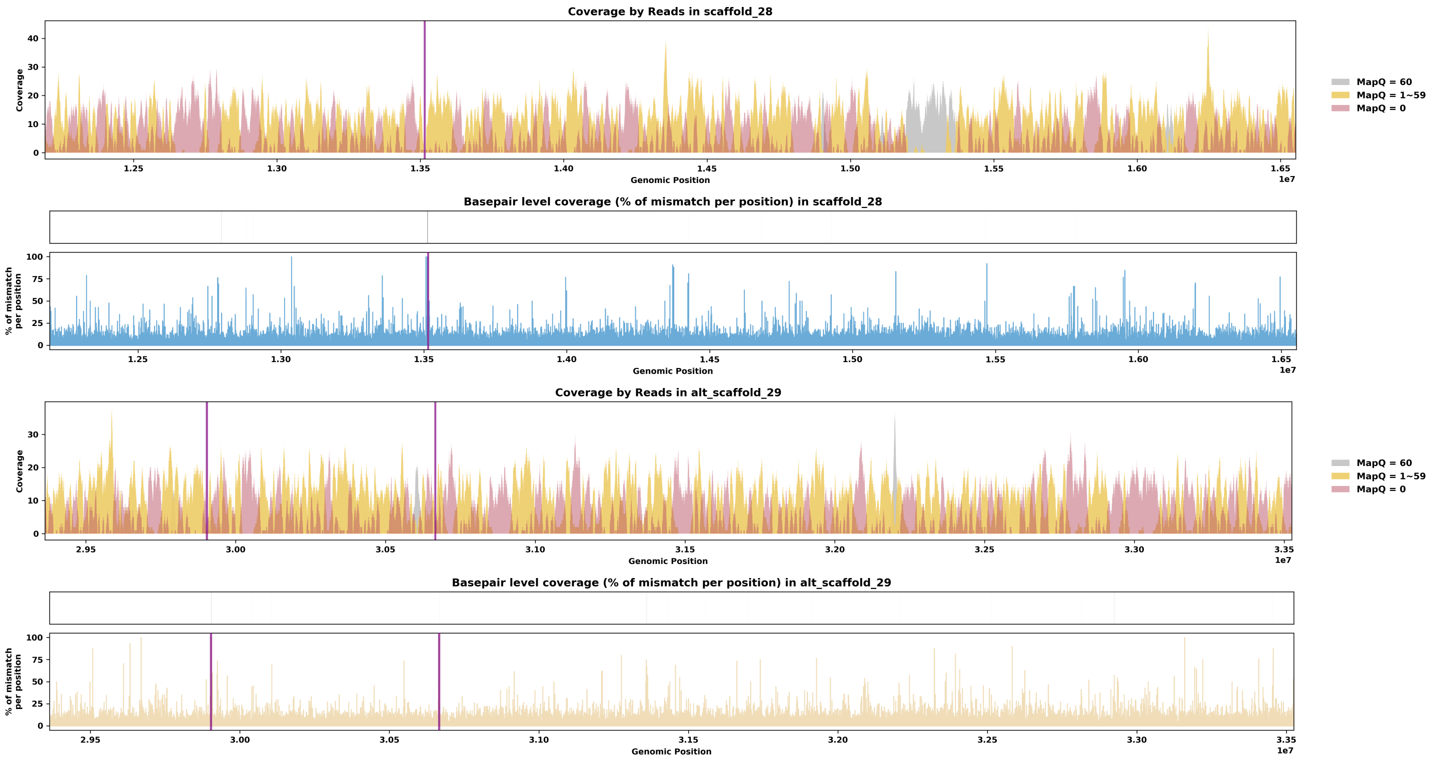

Species ID: mCerEla1

Common Name: Red Deer

Scientific Name: Cervus_elaphus

Assembly Type: Not Haplotype Resolved

Data Source: VGP

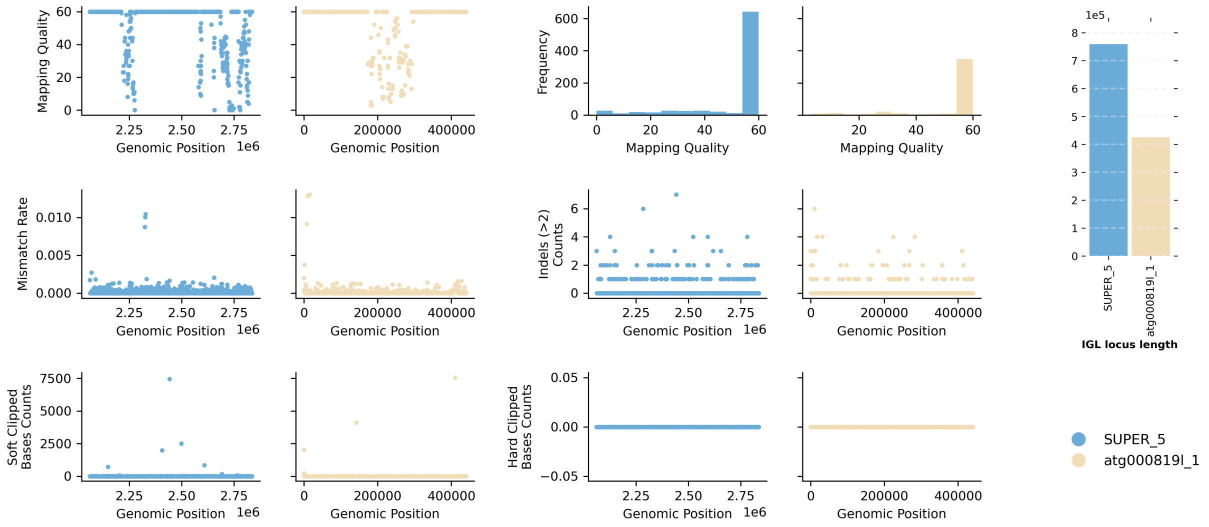


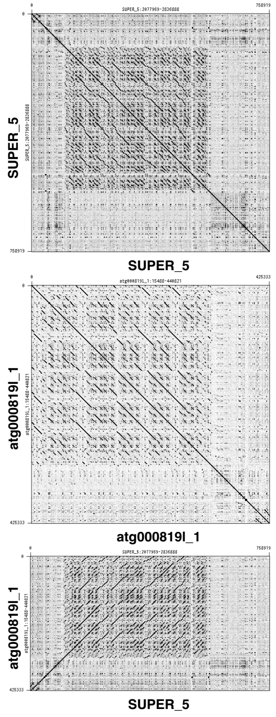


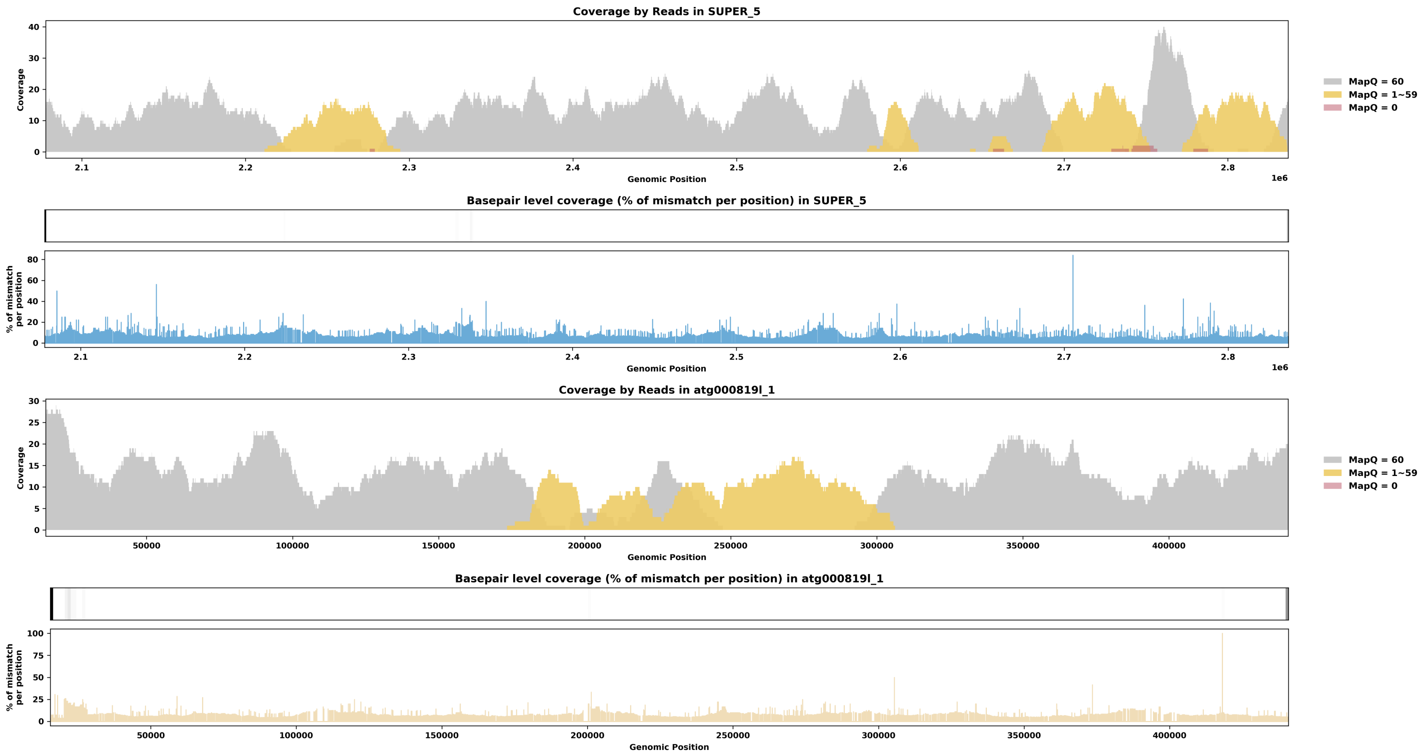

Species ID: mCorTow1

Common Name: Townsend's Big-eared Bat

Scientific Name: Corynorhinus townsendii

Assembly Type: Haplotype Resolved

Data Source: CCGP

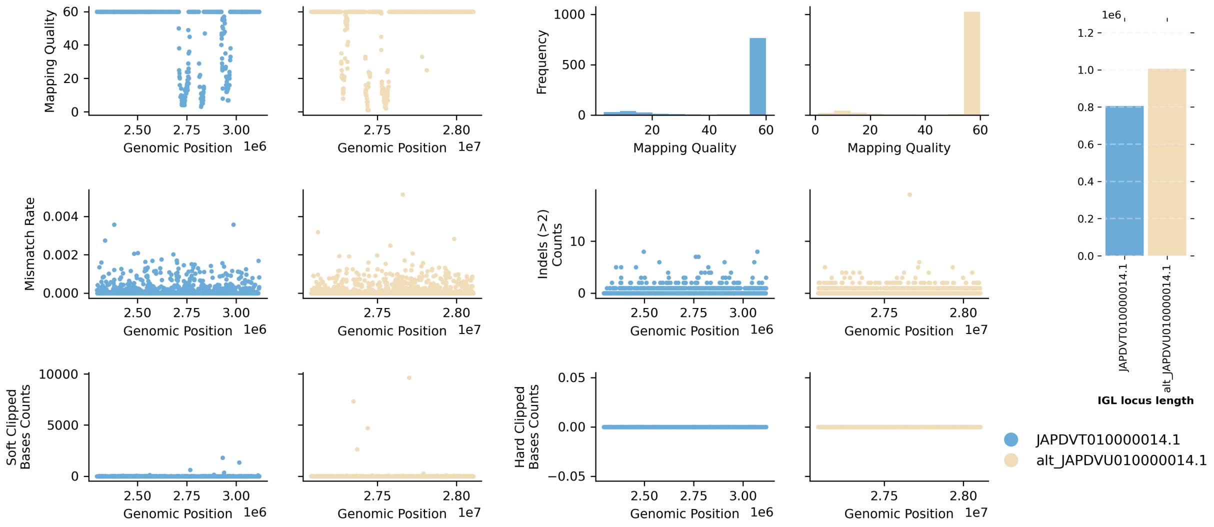


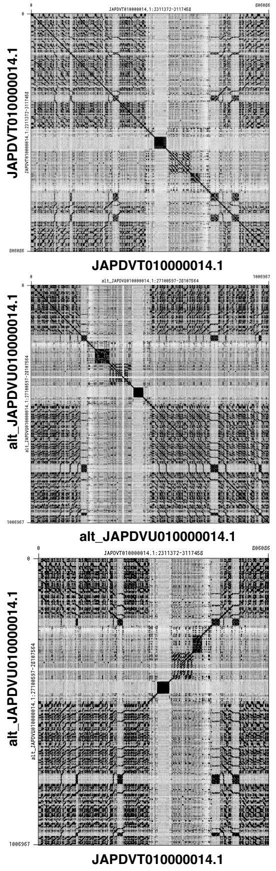


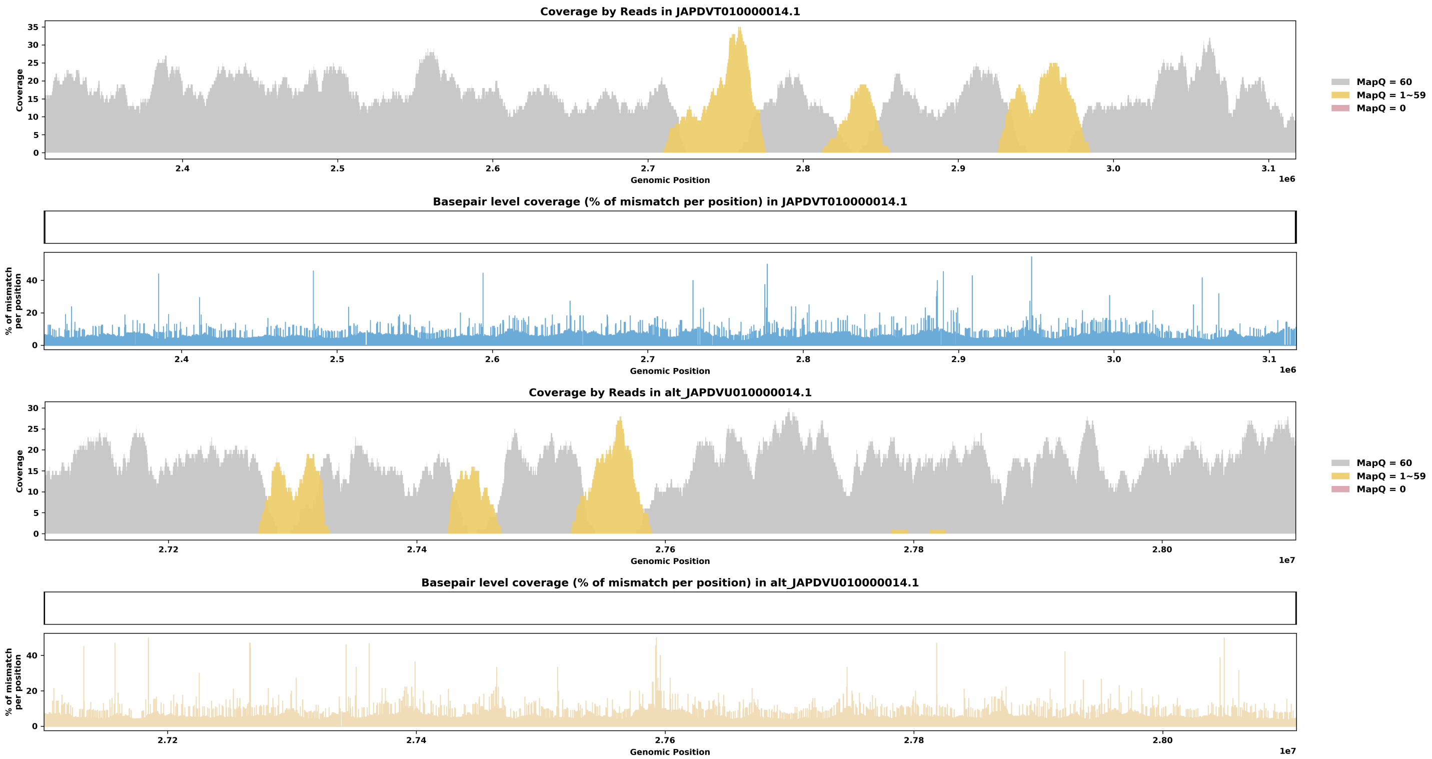

Species ID: mCynVol1

Common Name: Philiphine flying lemur

Scientific Name: Cynocephalus volans

Assembly Type: Not Haplotype Resolved

Data Source: VGP

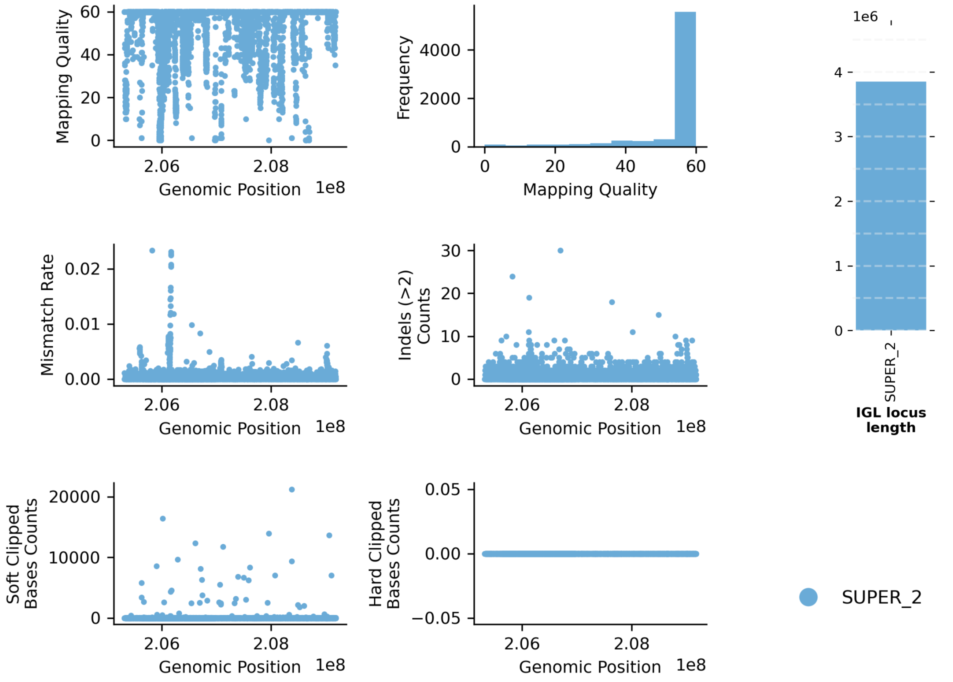


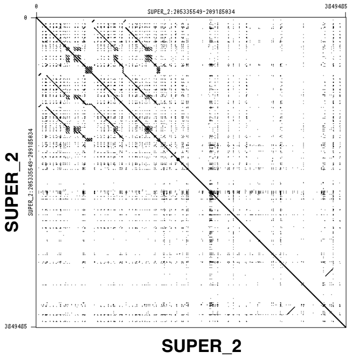


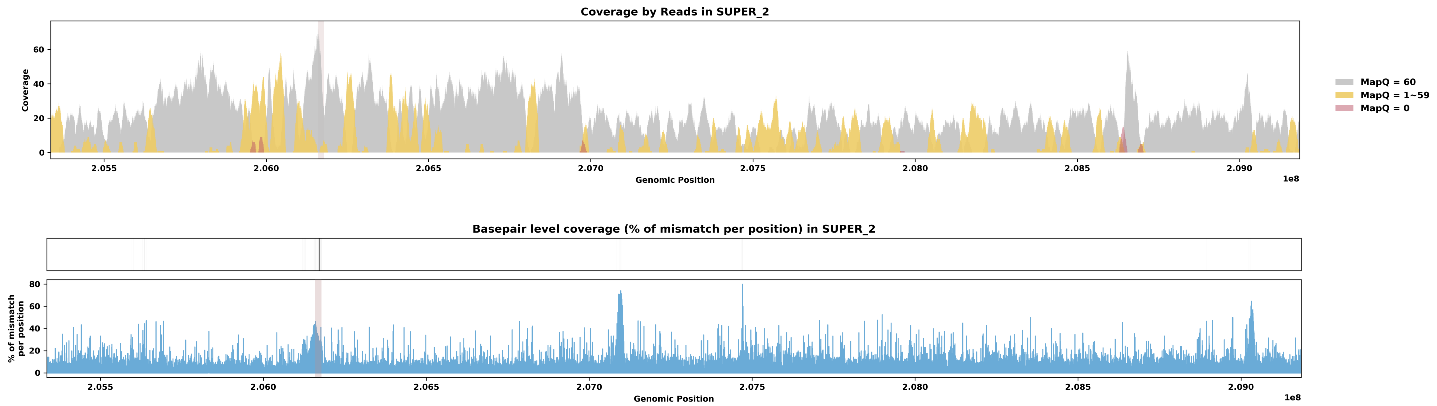

Species ID: mDasNov1

Common Name: nine-banded armadillo

Scientific Name: Dasypus novemcinctus

Assembly Type: Haplotype Resolved

Data Source: VGP

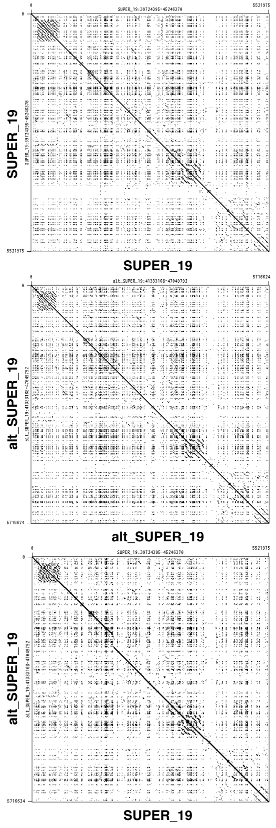


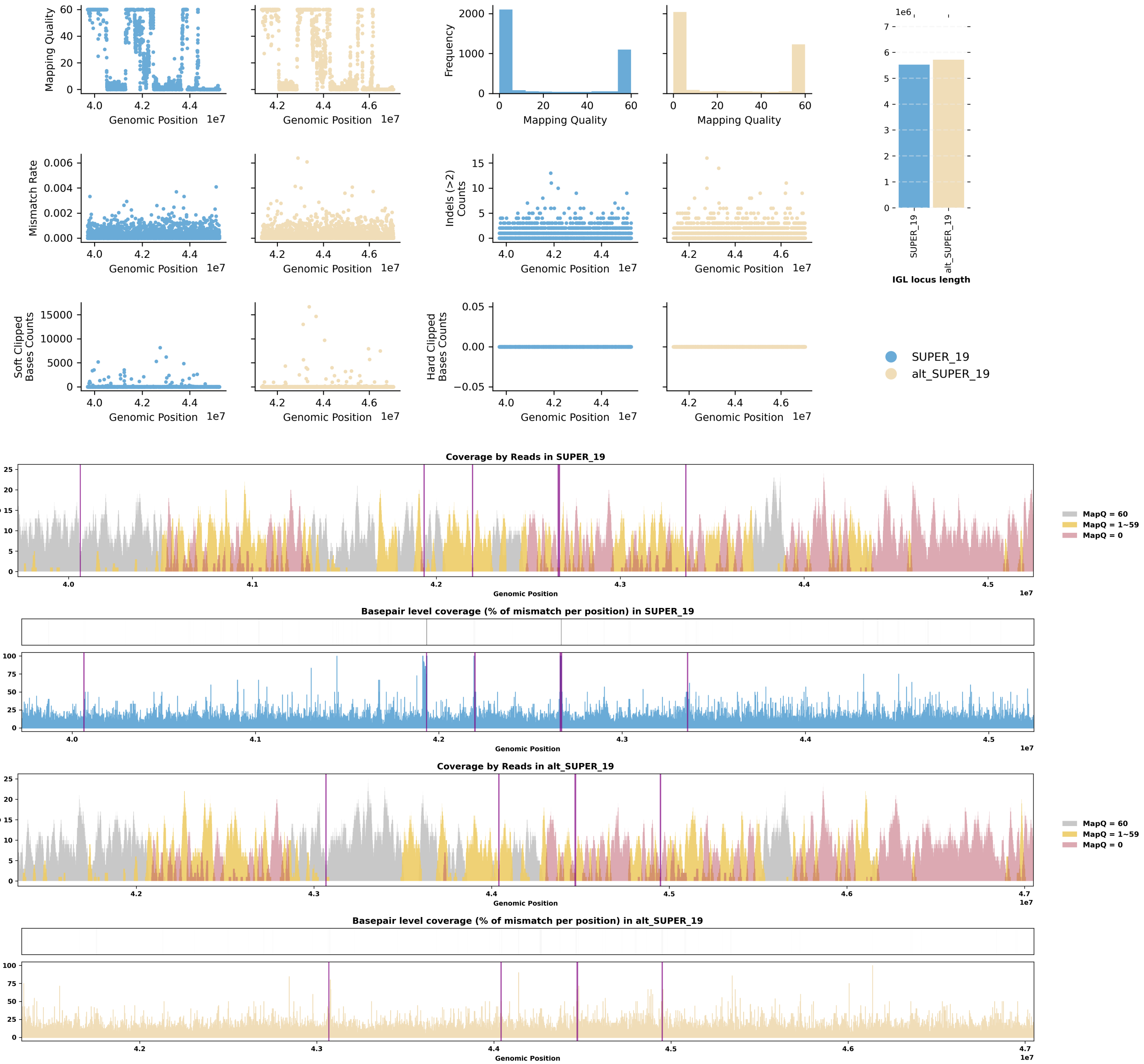

Species ID: mDelDel1

Common Name: saddleback dolphin

Scientific Name: Delphinus delphis

Assembly Type: Not Haplotype Resolved

Data Source: VGP

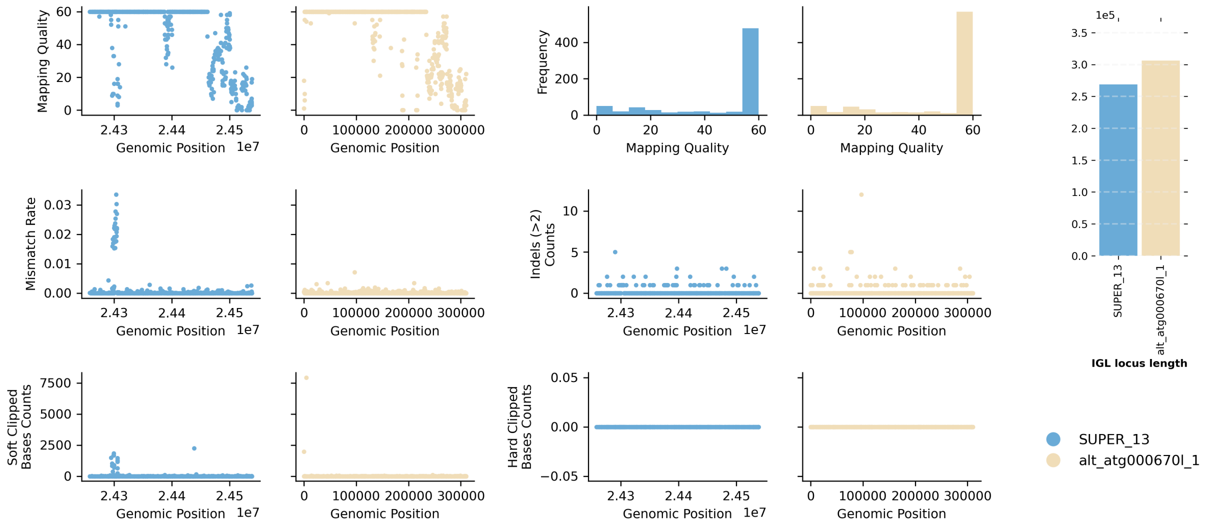


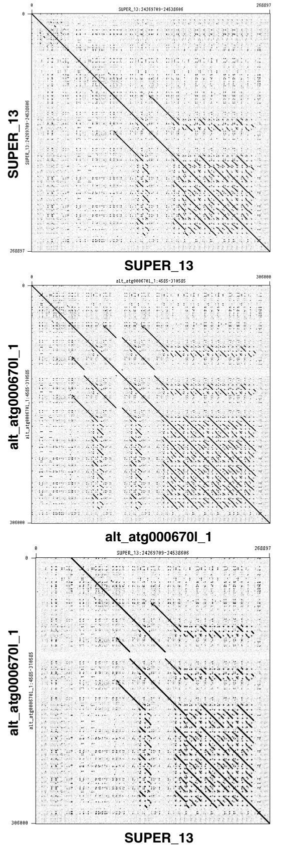


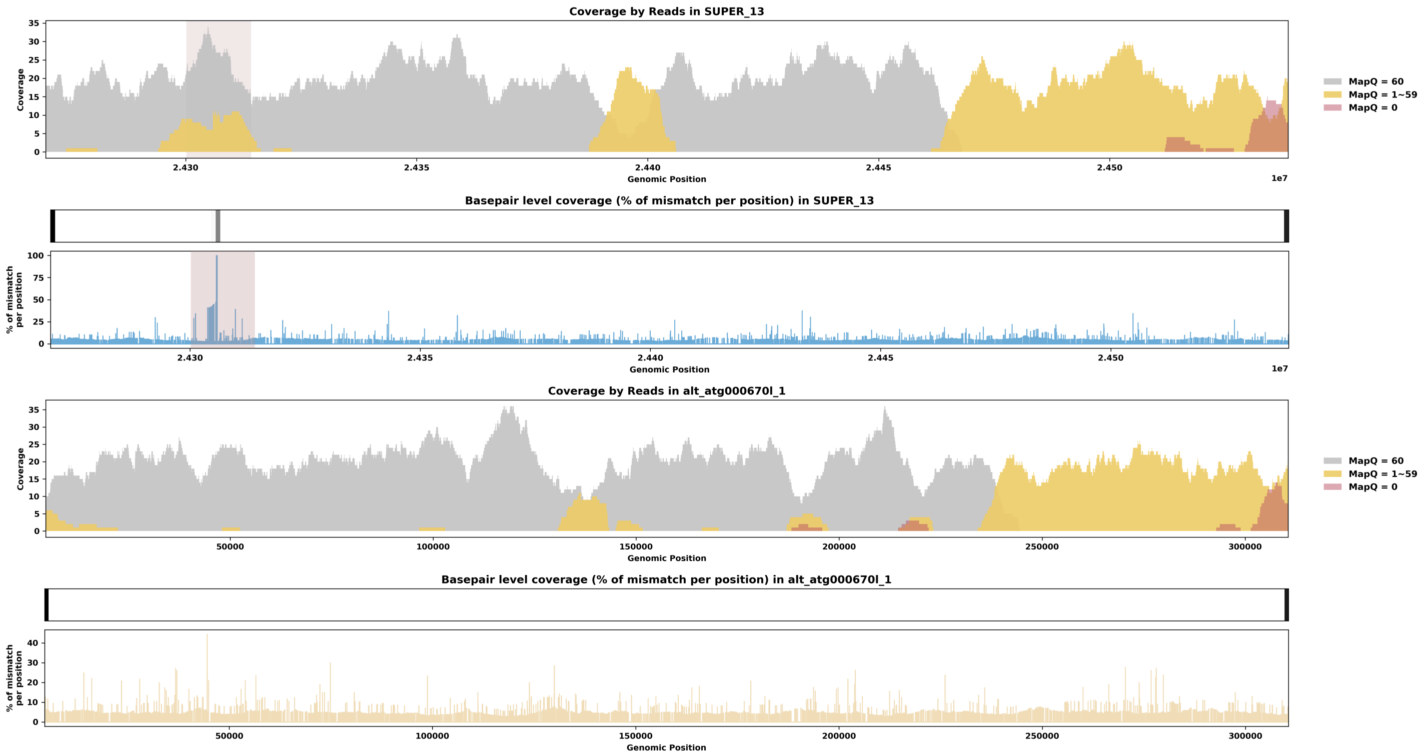

Species ID: mDicBic1

Common Name: black rhinoceros

Scientific Name: Diceros bicornis

Assembly Type: Haplotype Resolved

Data Source: VGP

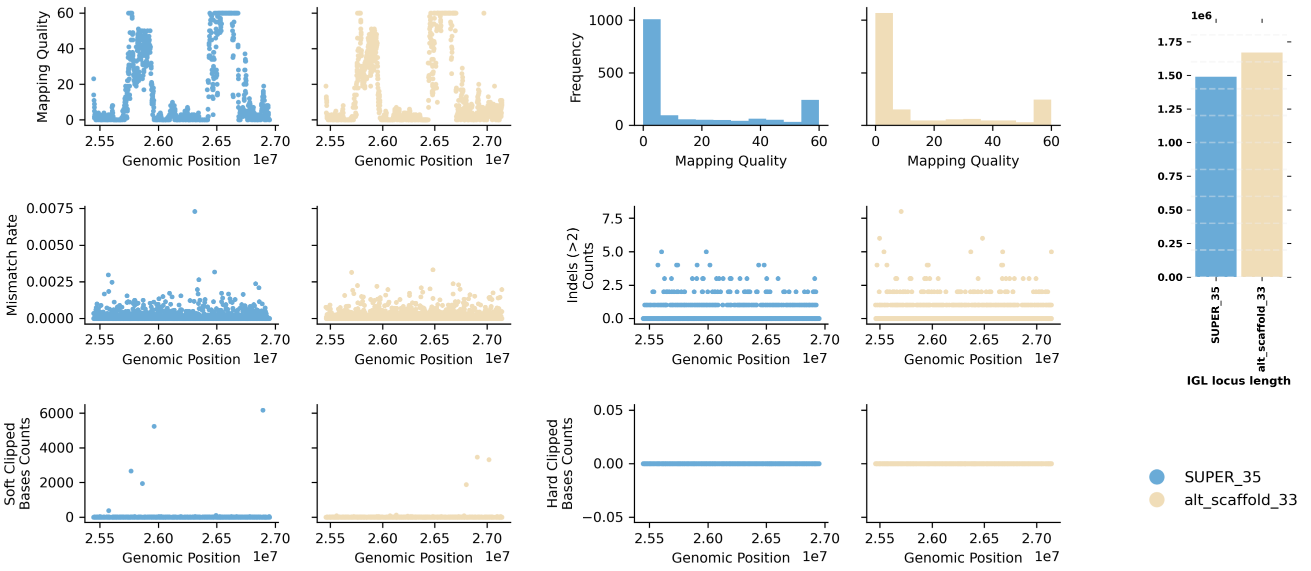


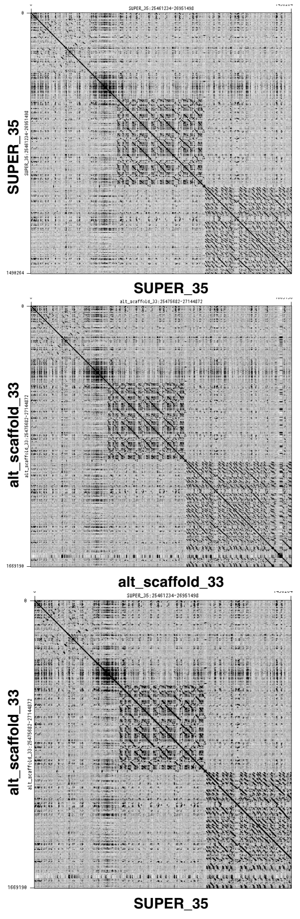


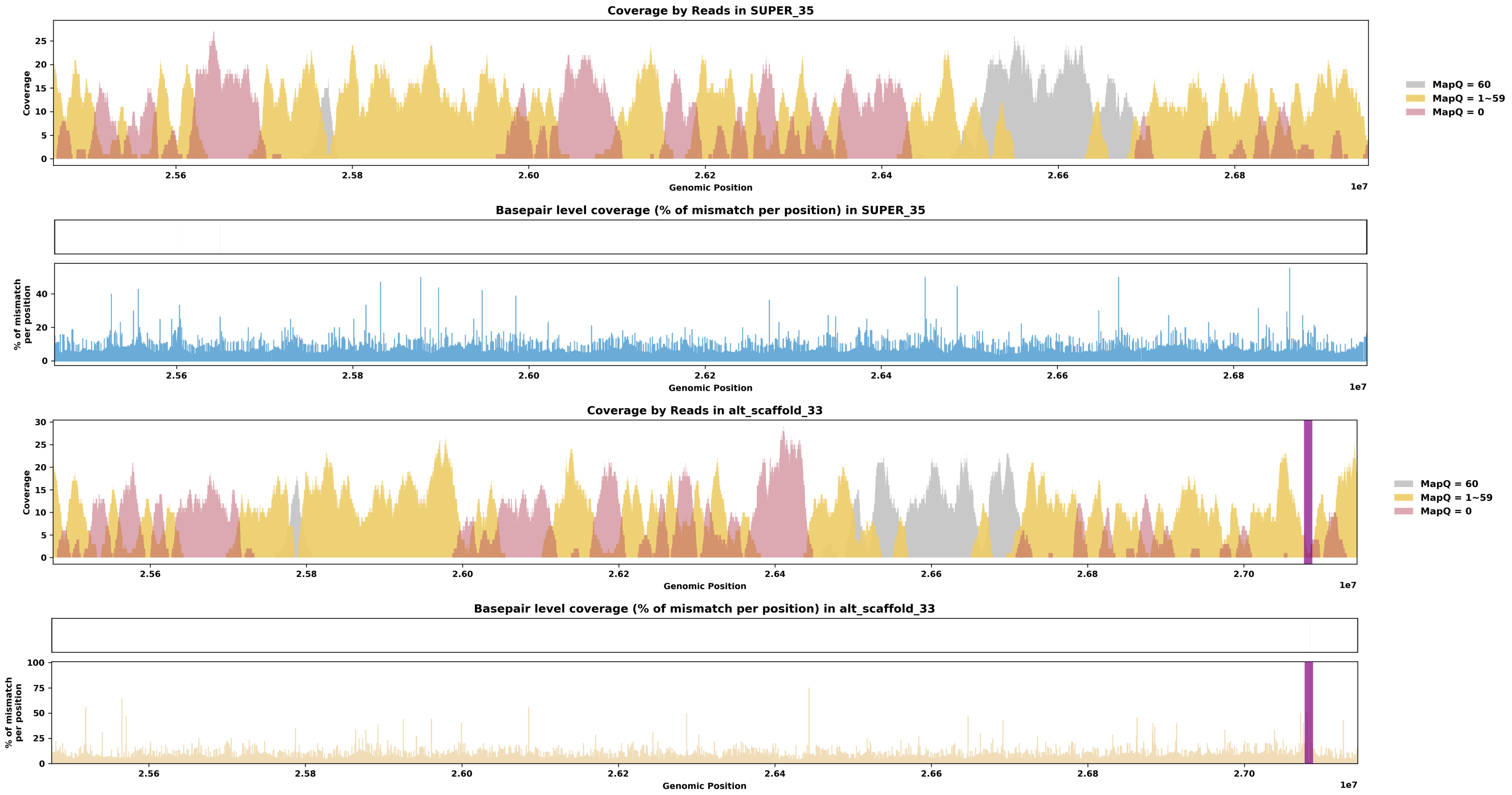

Species ID: mEleMax1

Common Name: Asiatic Elephant

Scientific Name: Elephas maximus

Assembly Type: Not Haplotype Resolved

Data Source: VGP

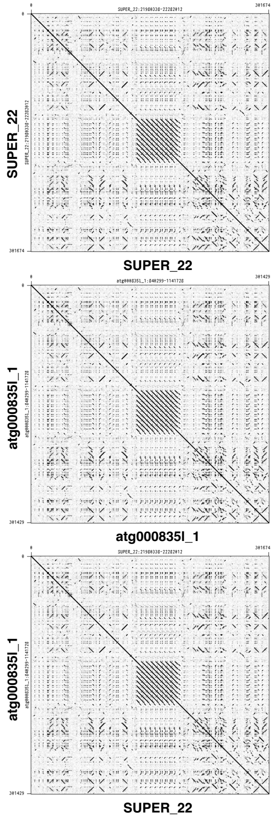


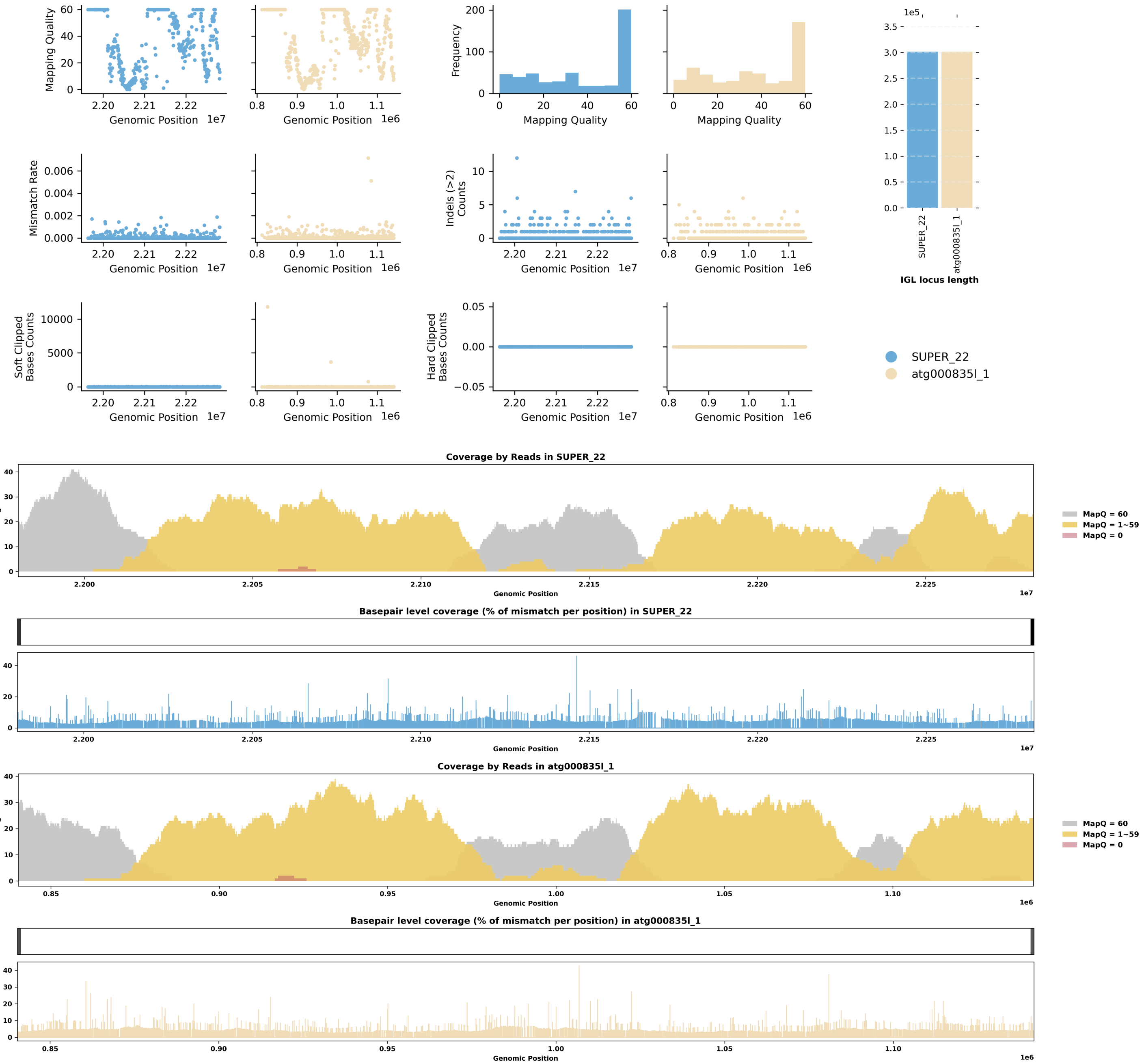

Species ID: mEptNil1

Common Name: northern bat

Scientific Name: Eptesicus nilssonii

Assembly Type: Not Haplotype Resolved

Data Source: VGP

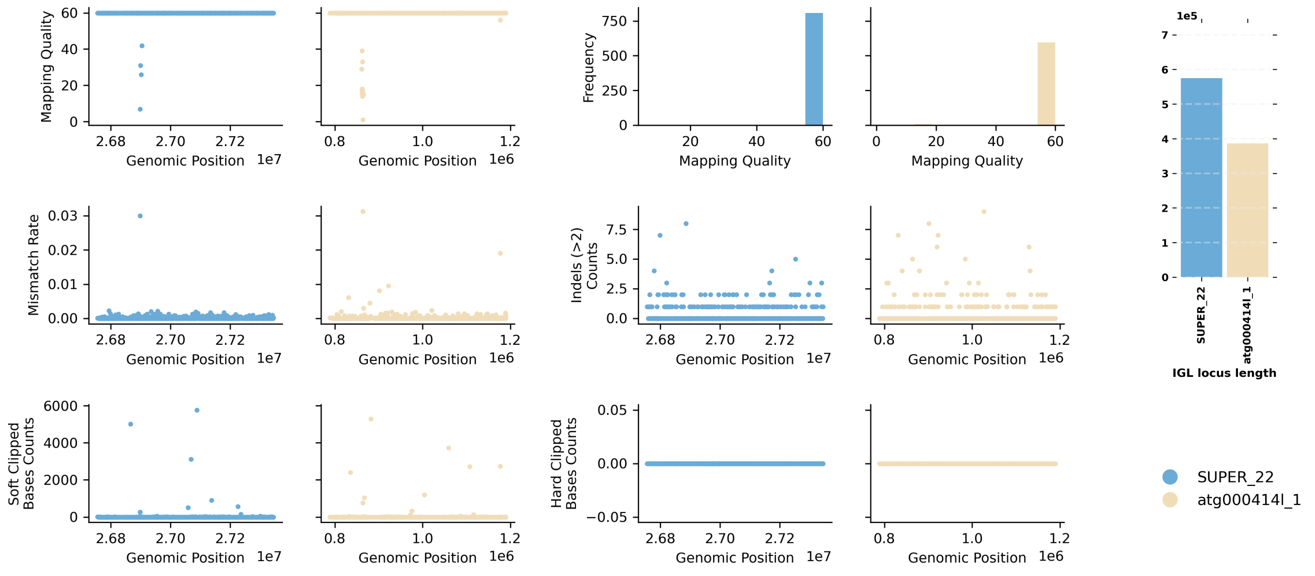


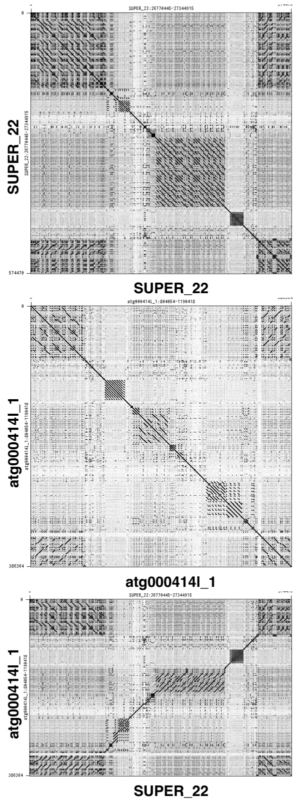


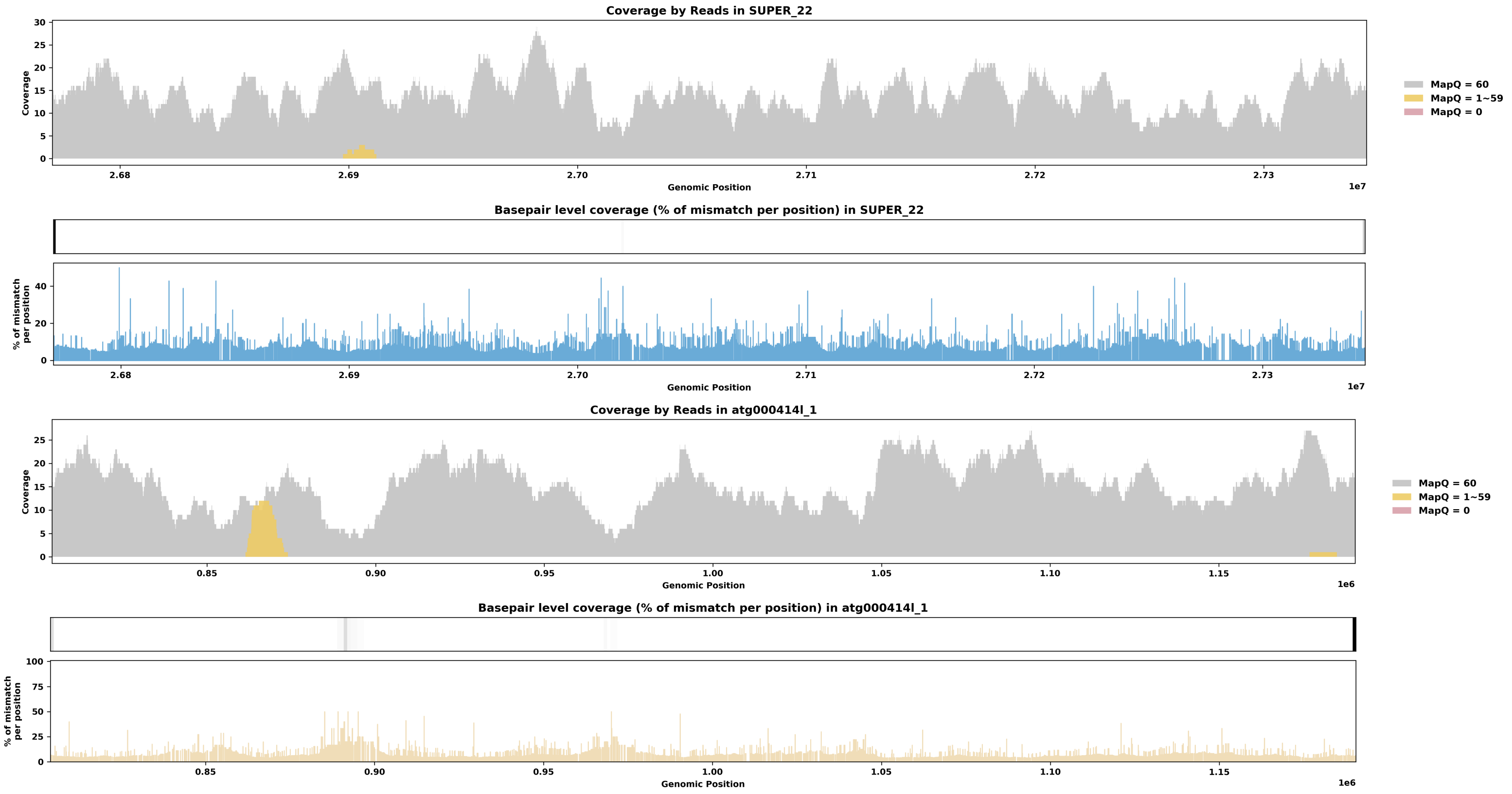

Species ID: mEriEur2

Common Name: western European hedgehog

Scientific Name: Erinaceus europaeus

Assembly Type: Not Haplotype Resolved

Data Source: VGP

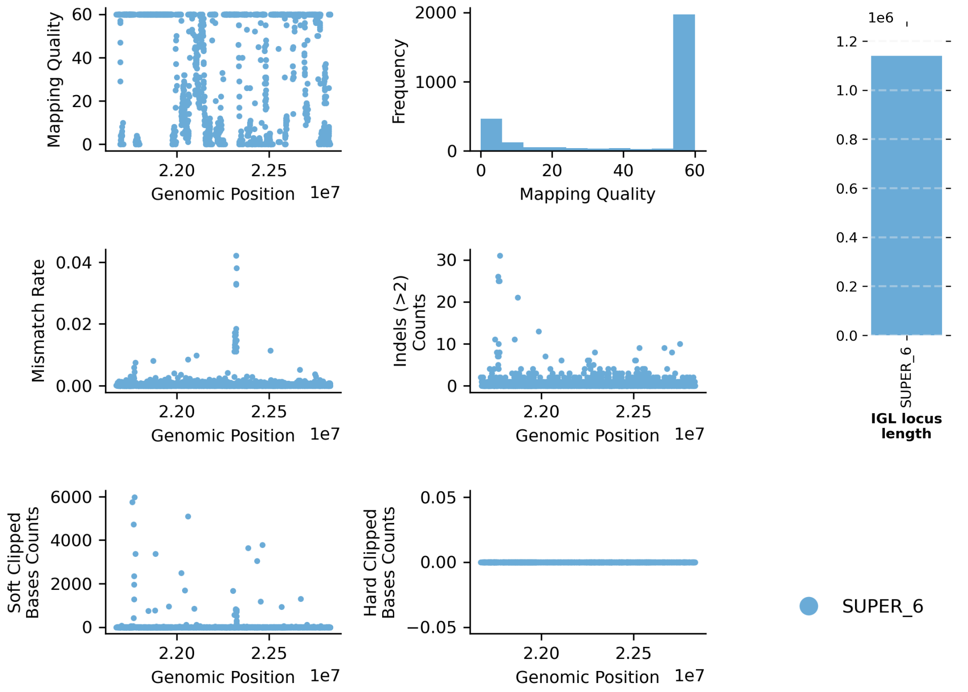


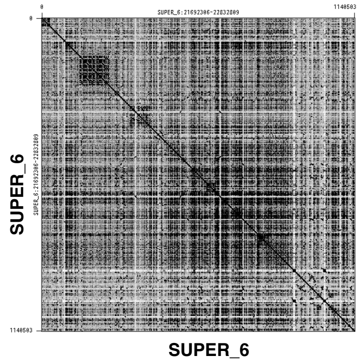


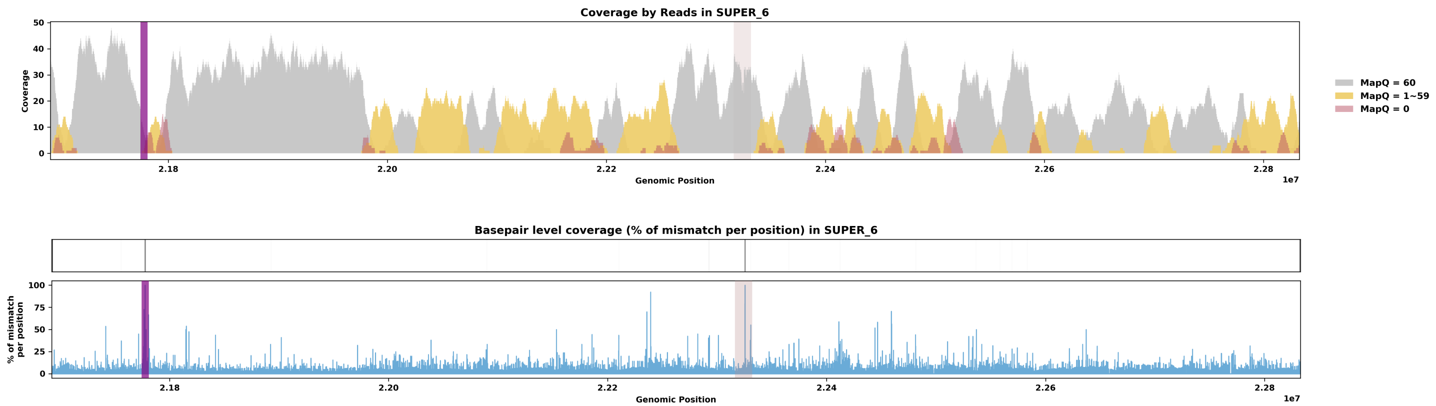

Species ID: mEscRob2

Common Name: grey whale

Scientific Name: Eschrichtius robustus

Assembly Type: Not Haplotype Resolved

Data Source: VGP

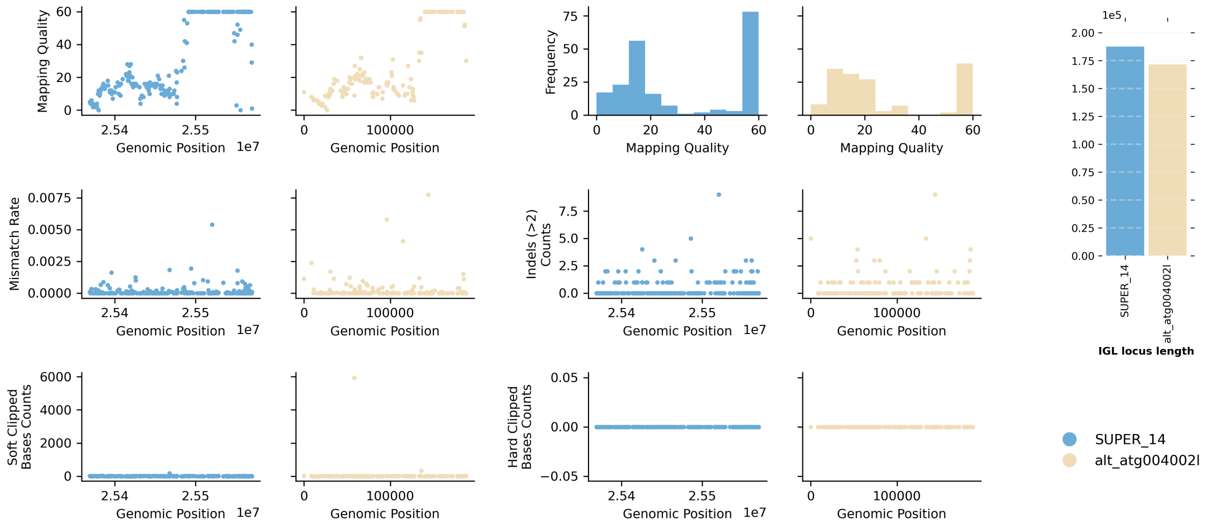


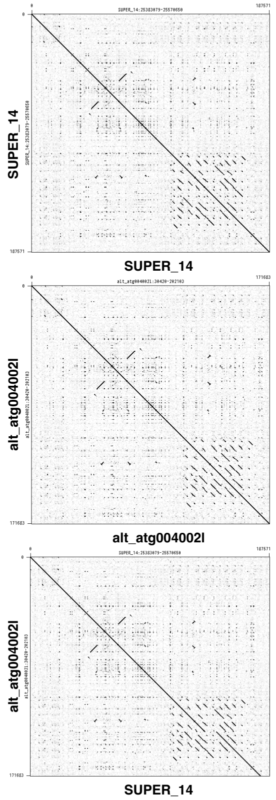


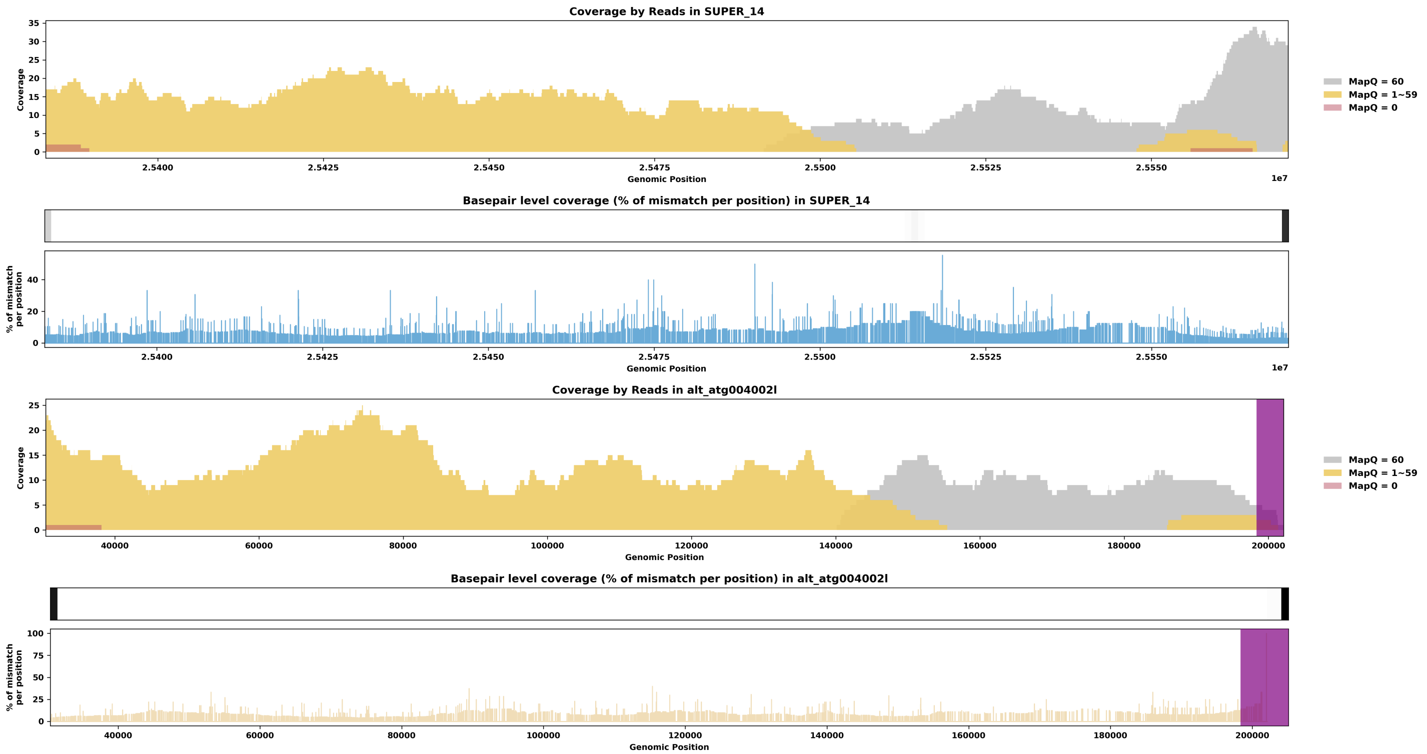

Species ID: mEubGla1

Common Name: North Atlantic right whale

Scientific Name: Eubalaena glacialis

Assembly Type: Haplotype Resolved

Data Source: VGP

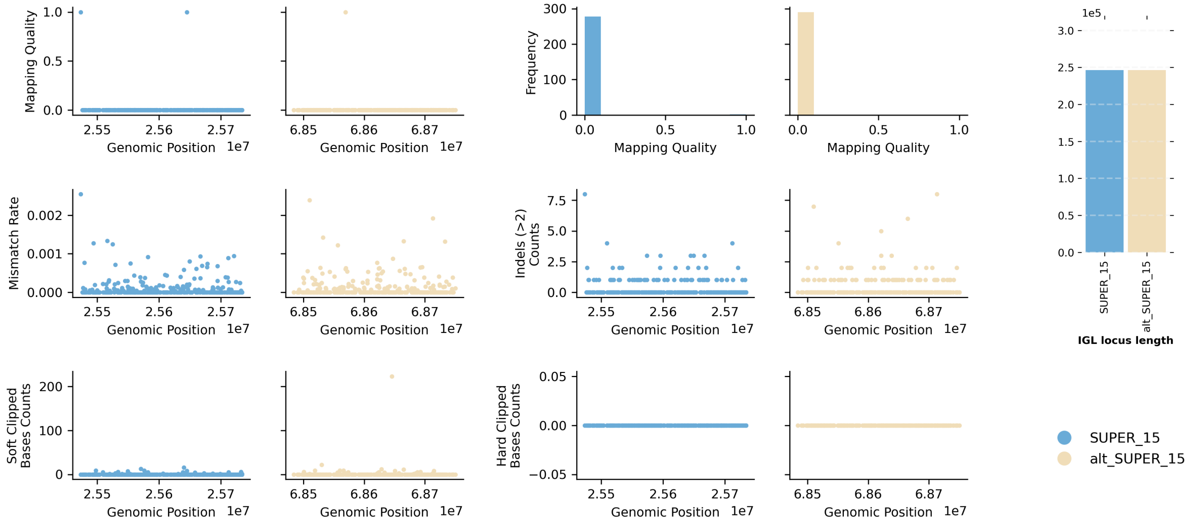


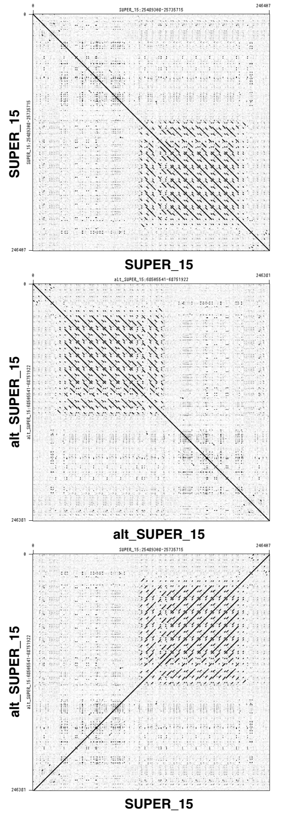


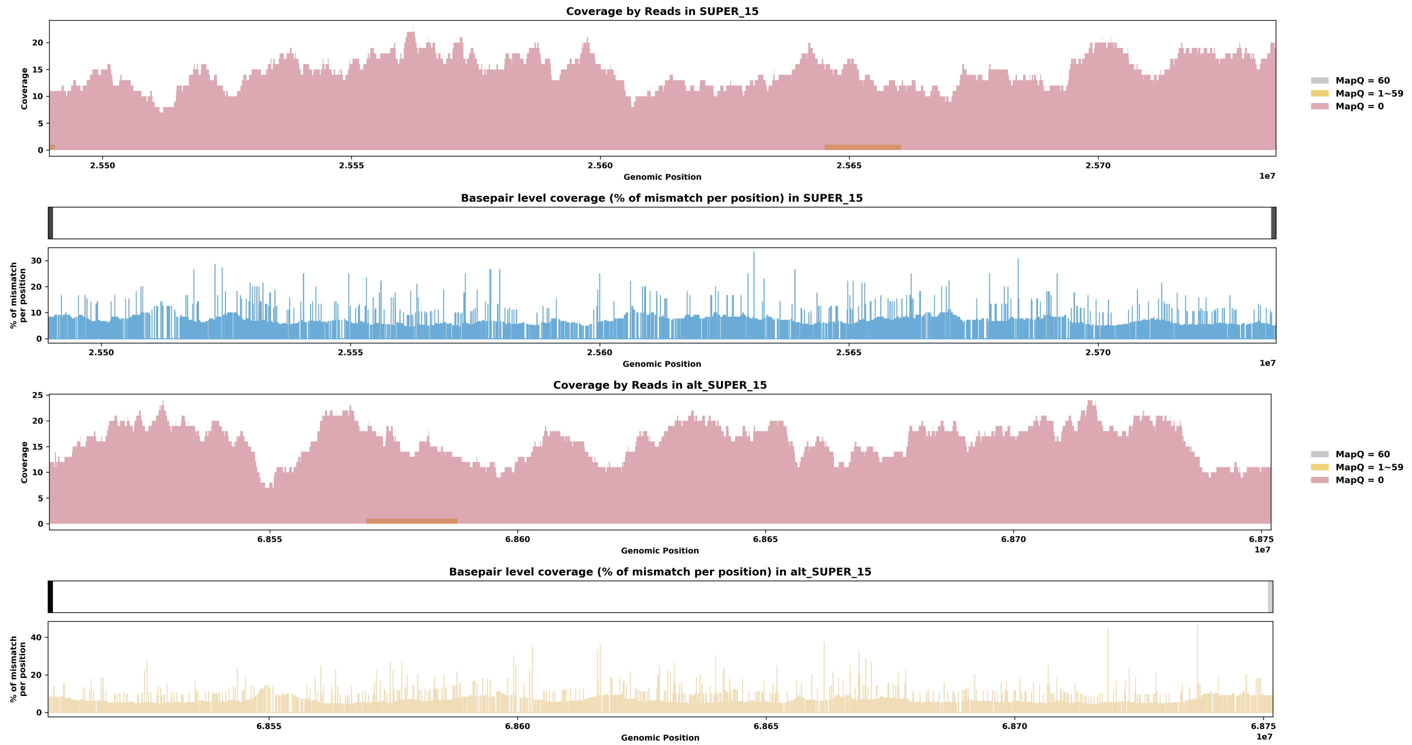

Species ID: mGloMel1

Common Name: long-finned pilot whale

Scientific Name: Globicephala melas

Assembly Type: Not Haplotype Resolved

Data Source: VGP

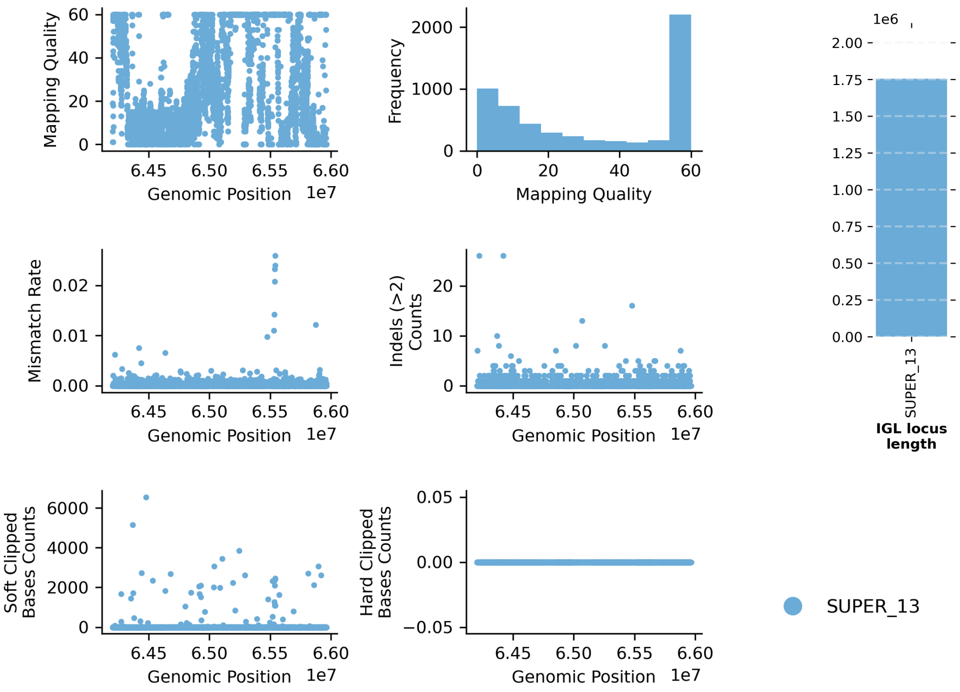


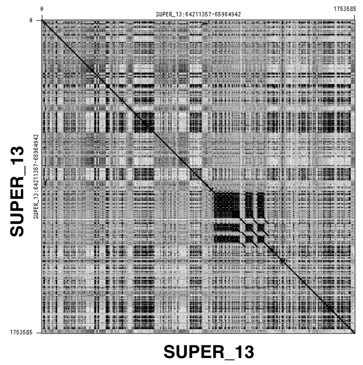


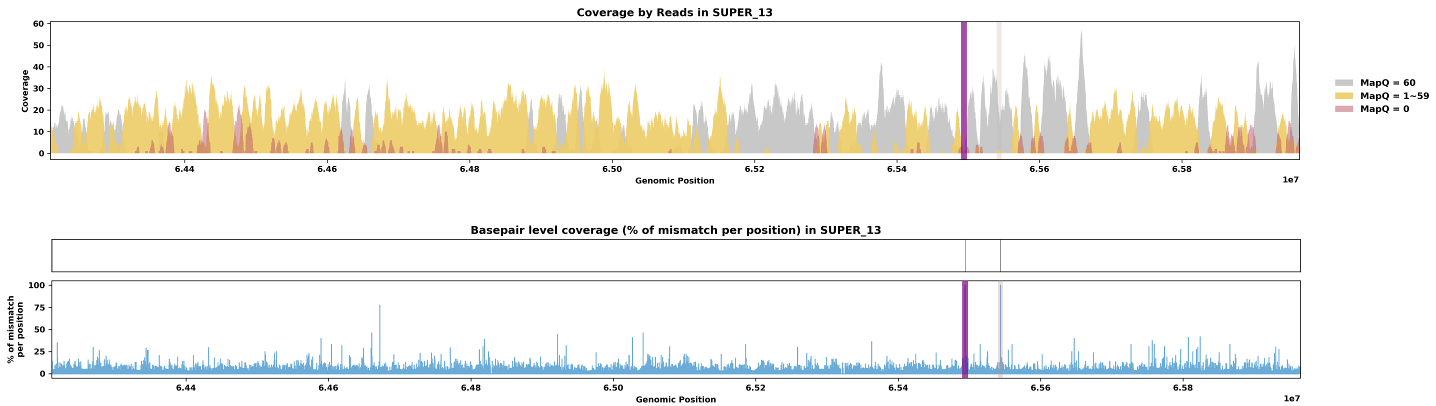

Species ID: mGorGor1

Common Name: Gorilla

Scientific Name: Gorilla_gorilla

Assembly Type: Haplotype Resolved

Data Source: T2T Primate

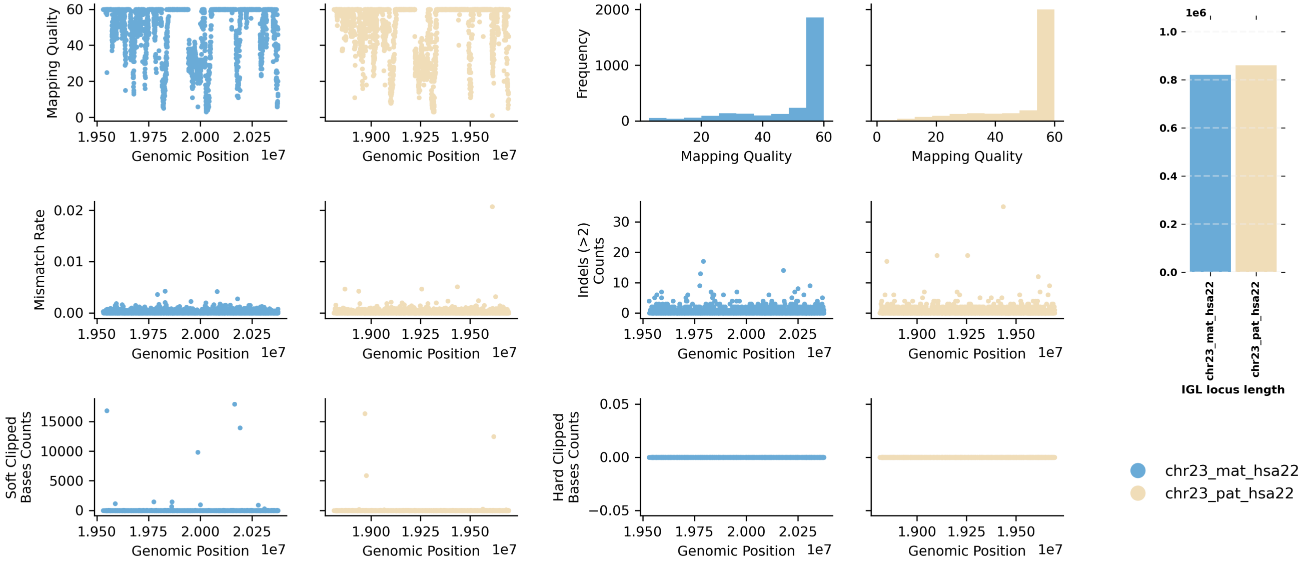


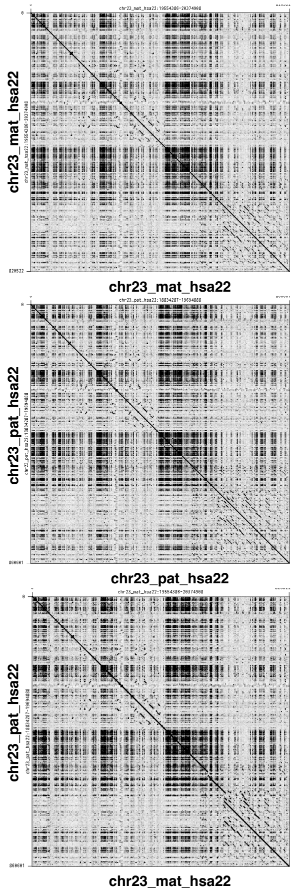


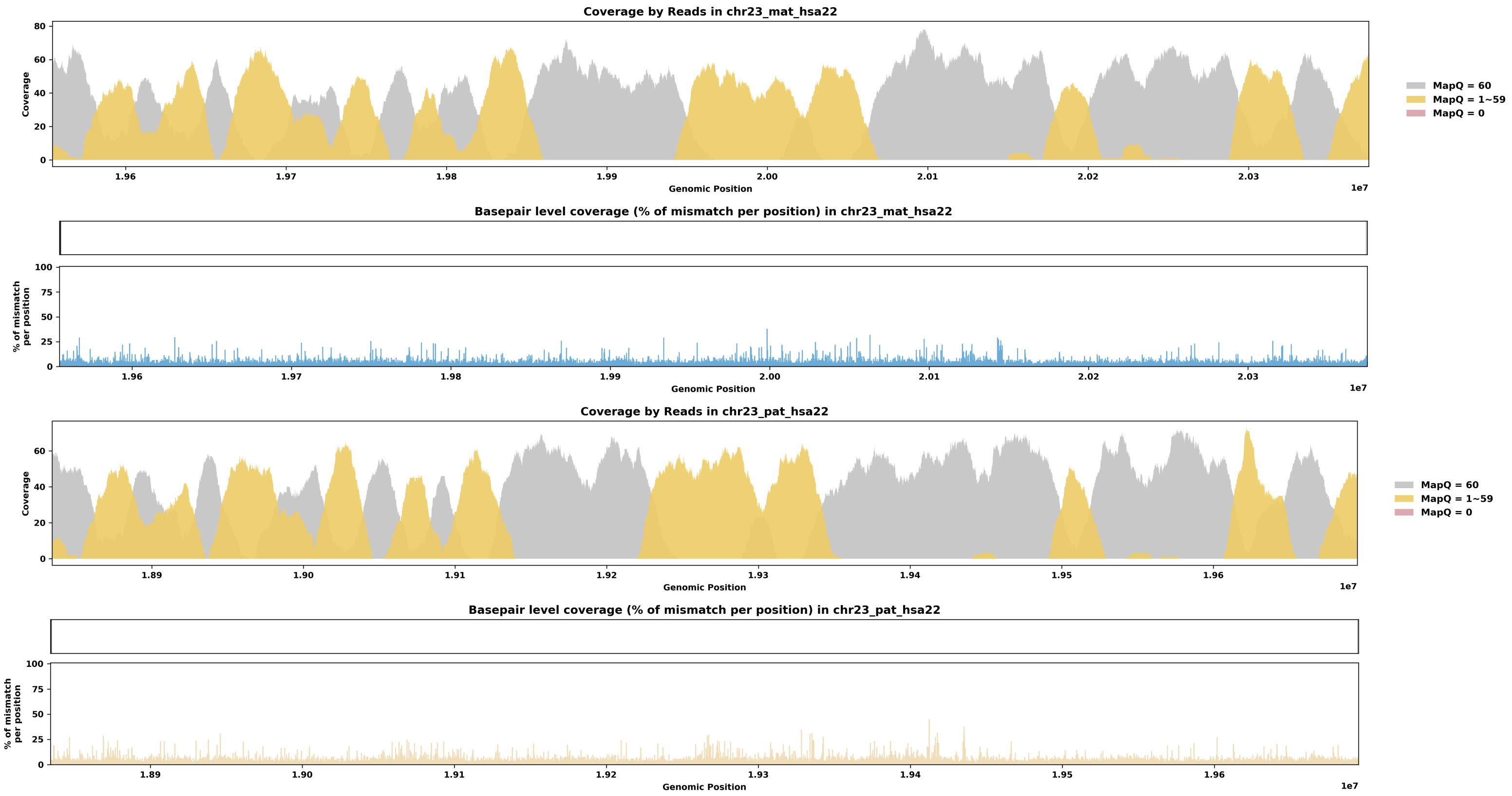

Species ID: mHetBru1

Common Name: Yellow-spotted hyrax

Scientific Name: Heterohyrax brucei

Assembly Type: Not Haplotype Resolved

Data Source: VGP

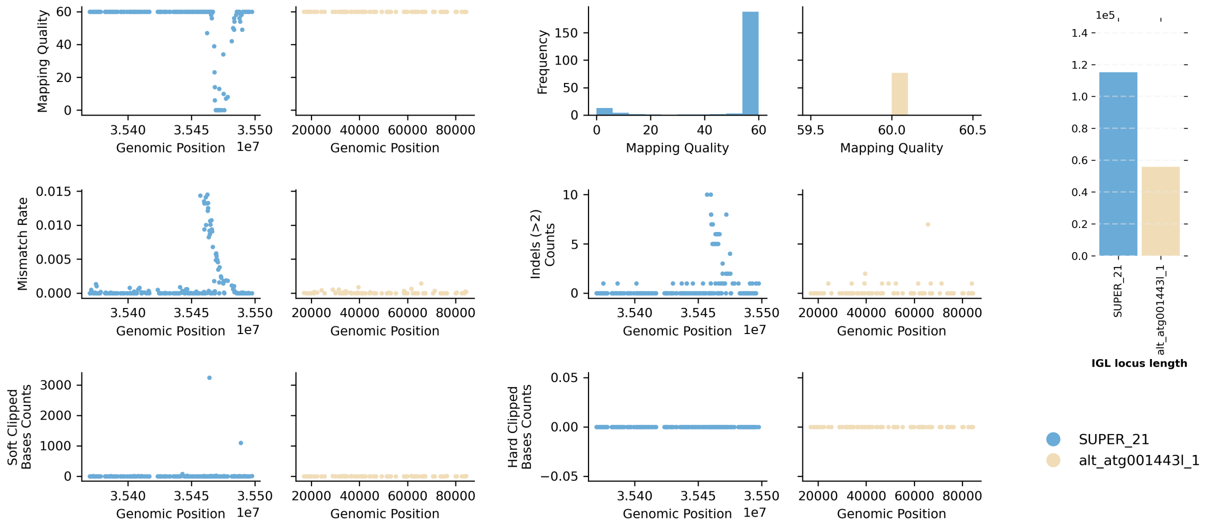


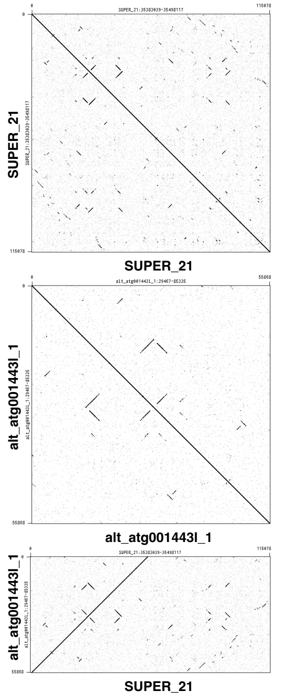


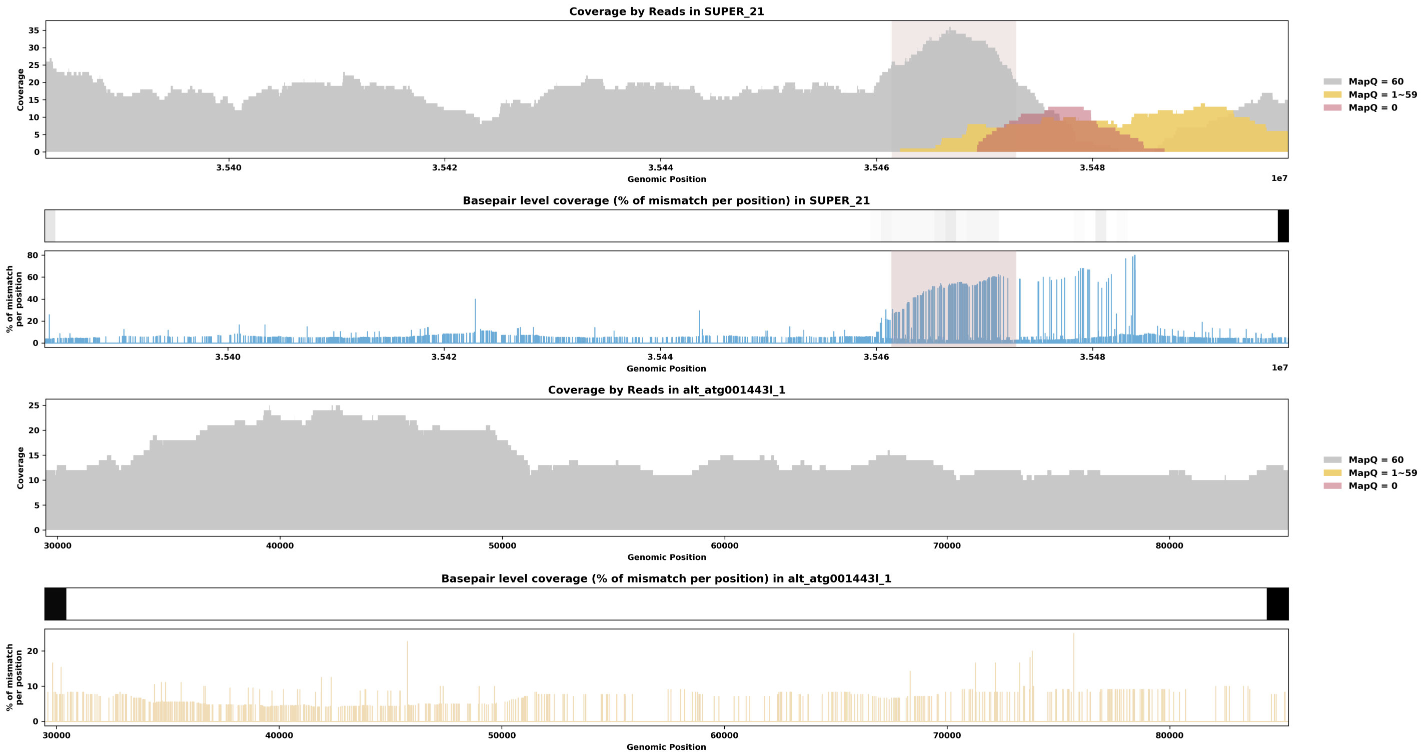

Species ID: mHipAmp2

Common Name: hippopotamus

Scientific Name: Hippopotamus amphibius kiboko

Assembly Type: Haplotype Resolved

Data Source: VGP

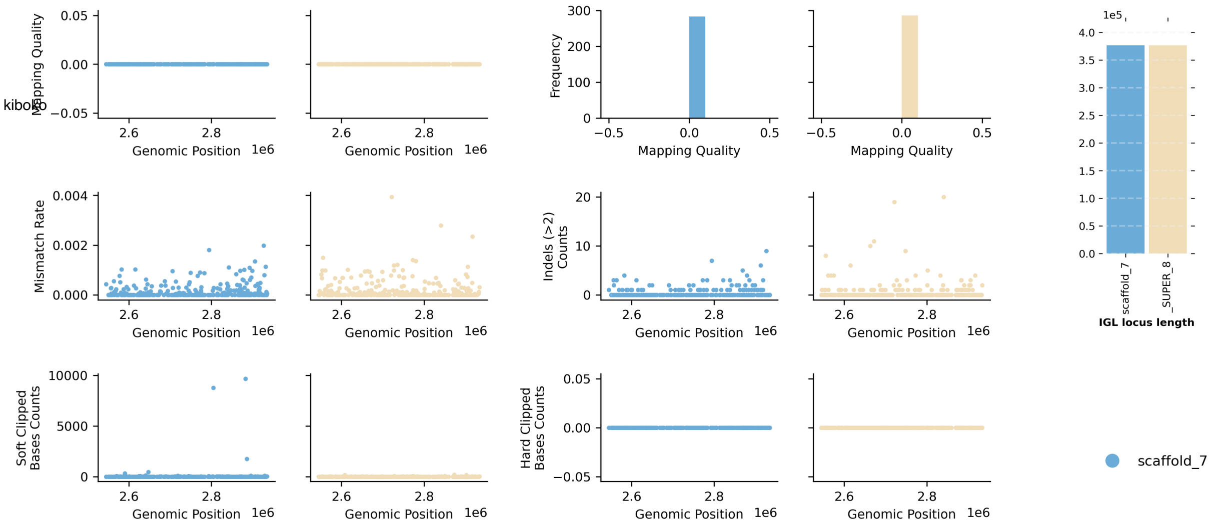


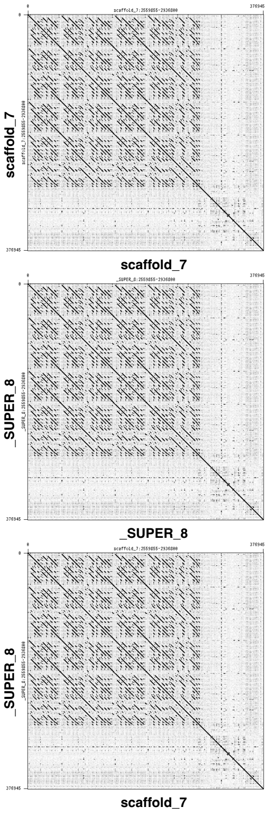


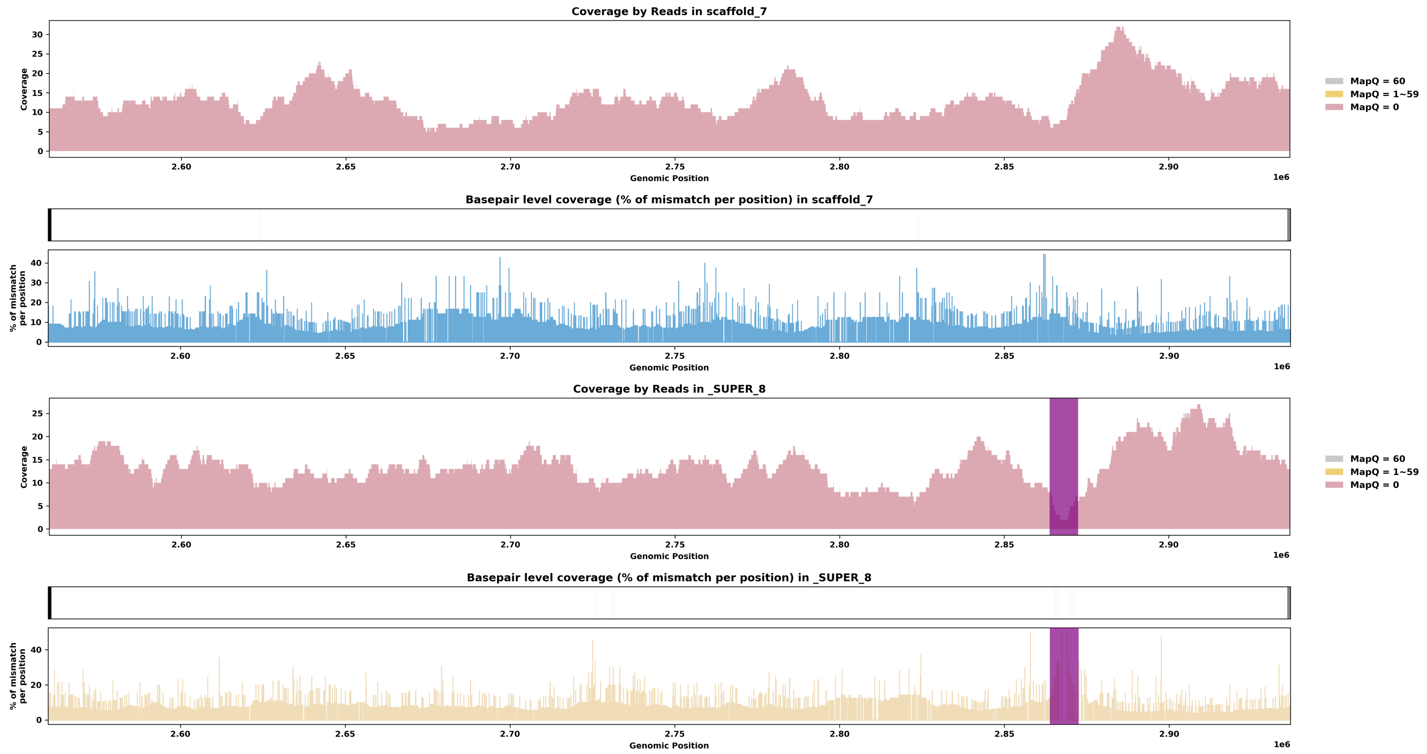

Species ID: mHypAmp2

Common Name: northern bottlenose whale

Scientific Name: Hyperoodon ampullatus

Assembly Type: Not Haplotype Resolved

Data Source: VGP

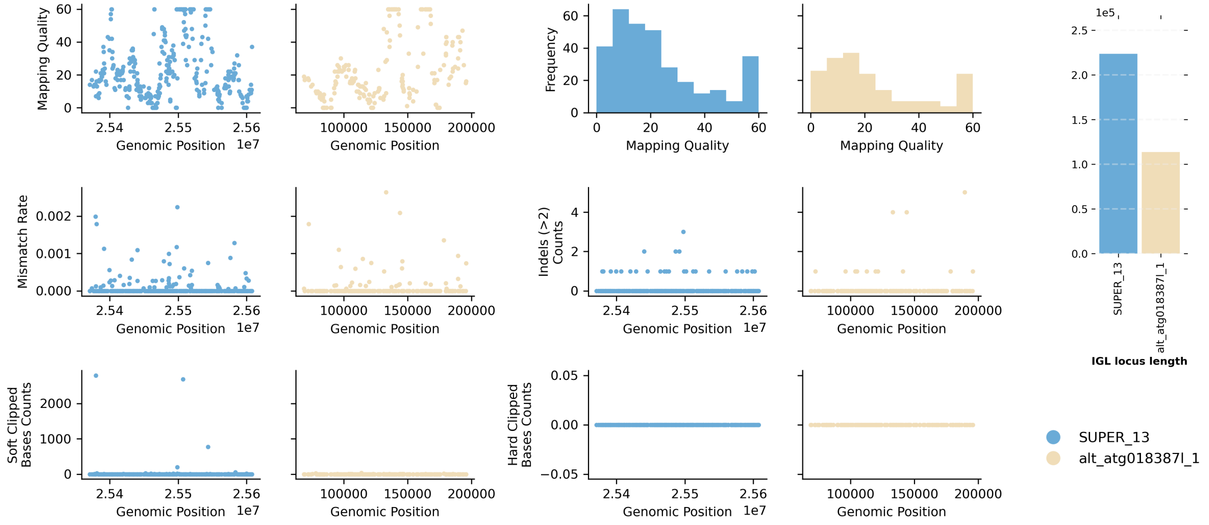


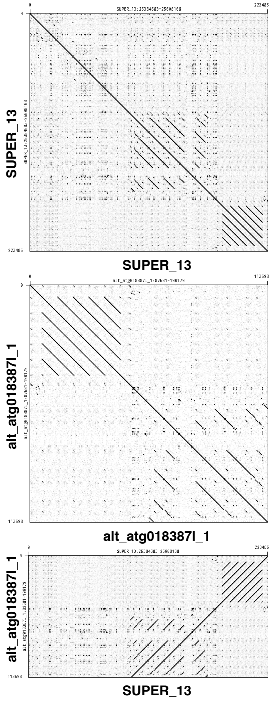


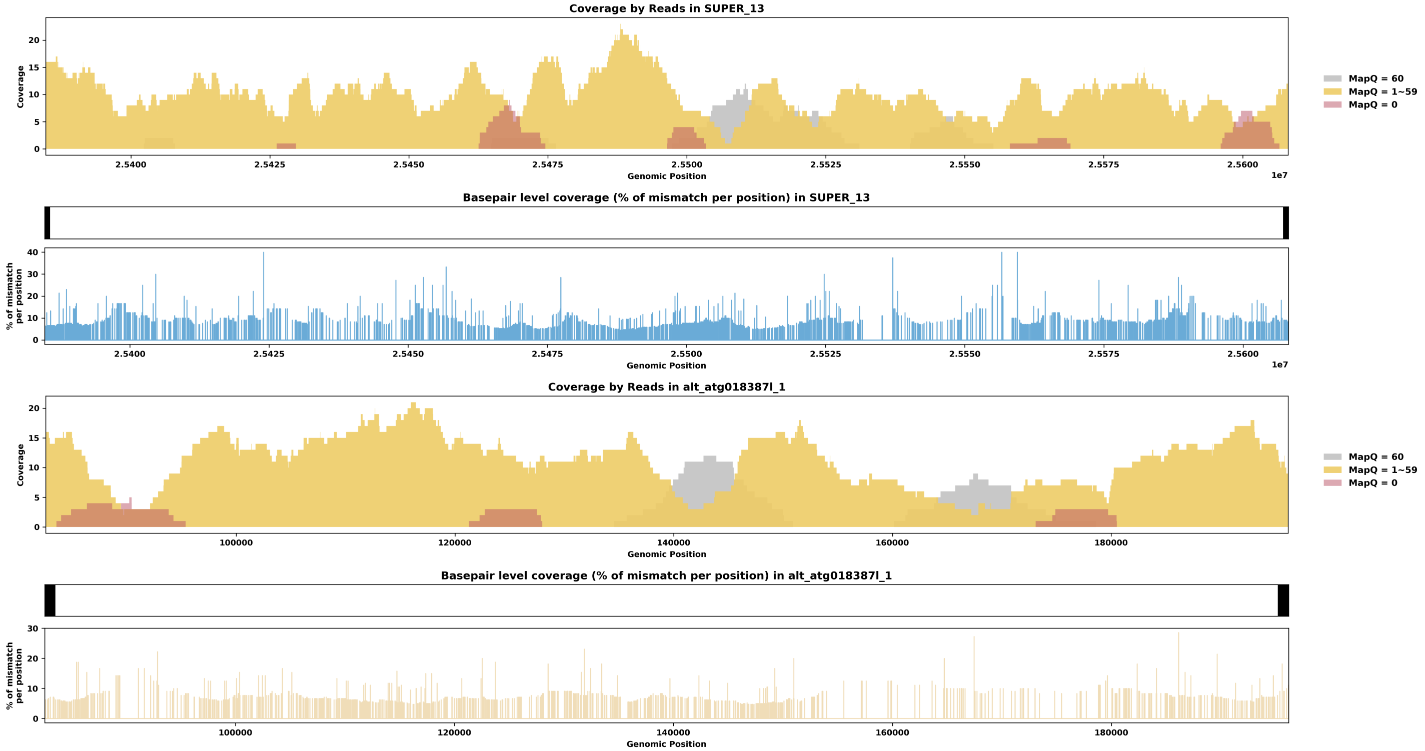

Species ID: mLagAlb1

Common Name: white-beaked dolphin

Scientific Name: Lagenorhynchus albirostris

Assembly Type: Not Haplotype Resolved

Data Source: VGP

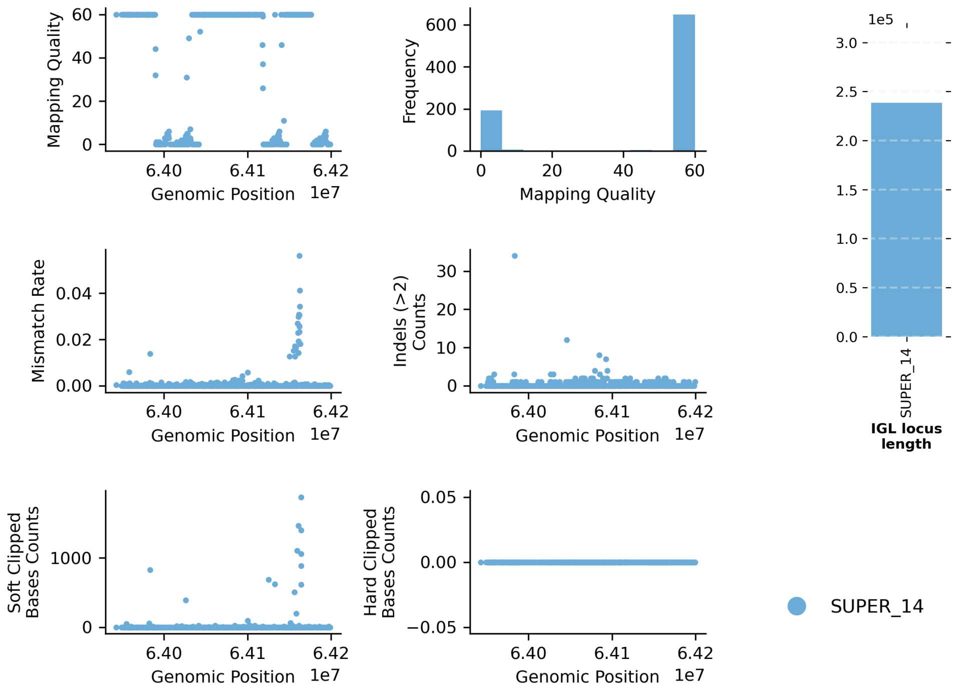


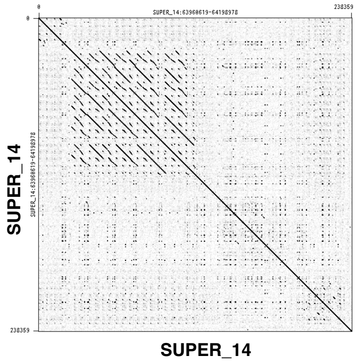


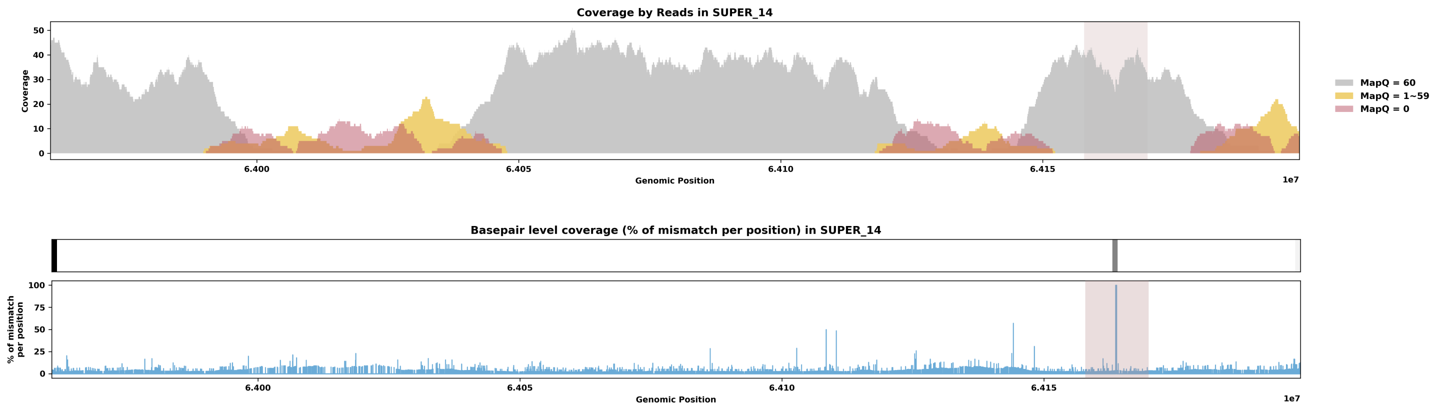

Species ID: mLemCat1

Common Name: Ring-tailed lemur

Scientific Name: Lemur catta

Assembly Type: Not Haplotype Resolved

Data Source: VGP

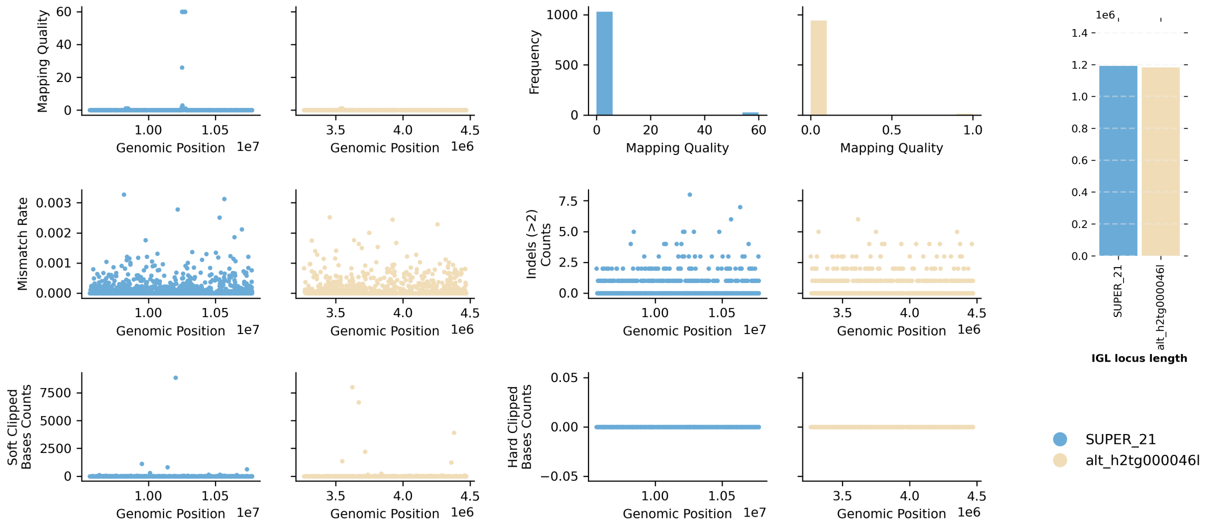


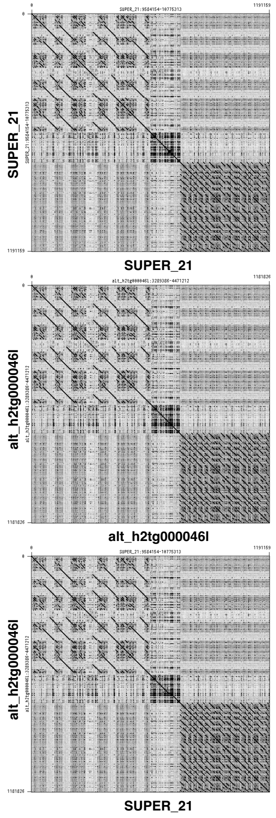


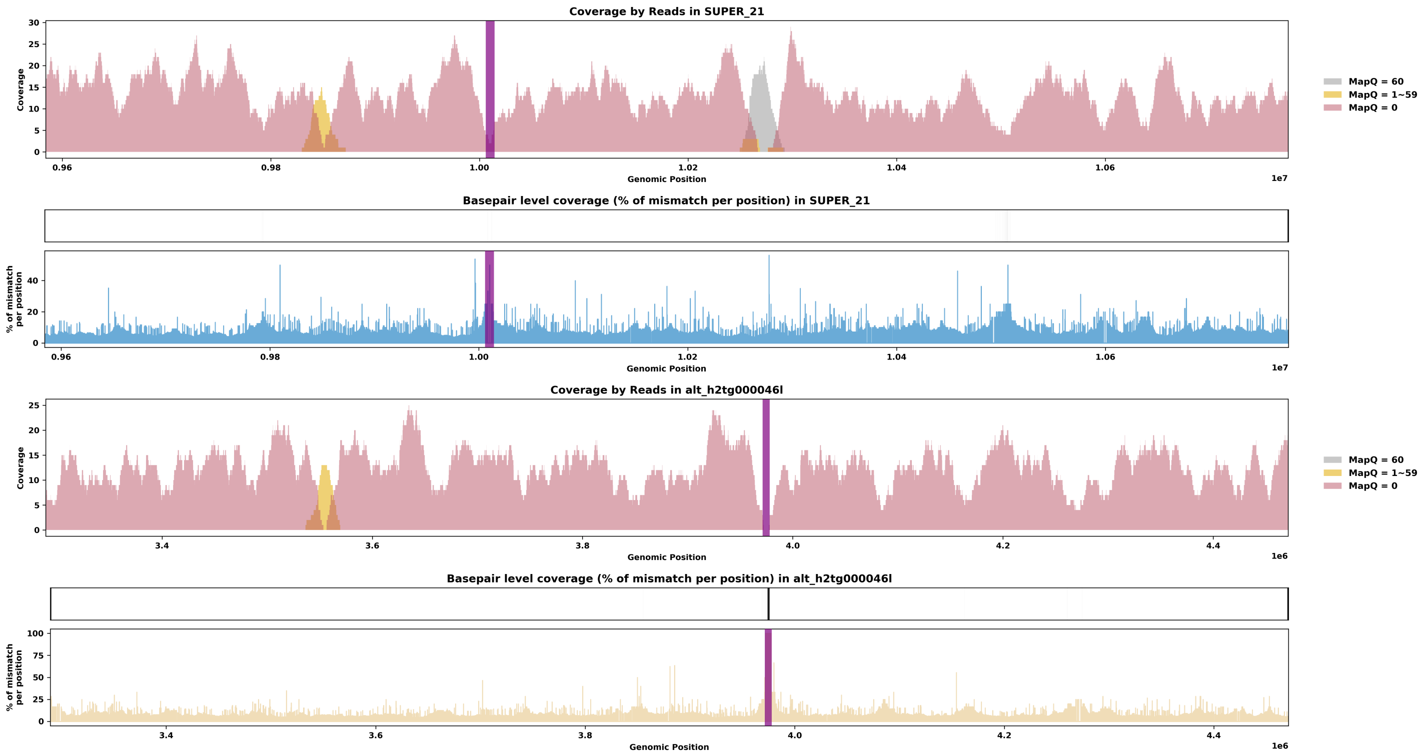

Species ID: mLynRuf1

Common Name: Bobcat

Scientific Name: Lynx rufus

Assembly Type: Not Haplotype Resolved

Data Source: CCGP

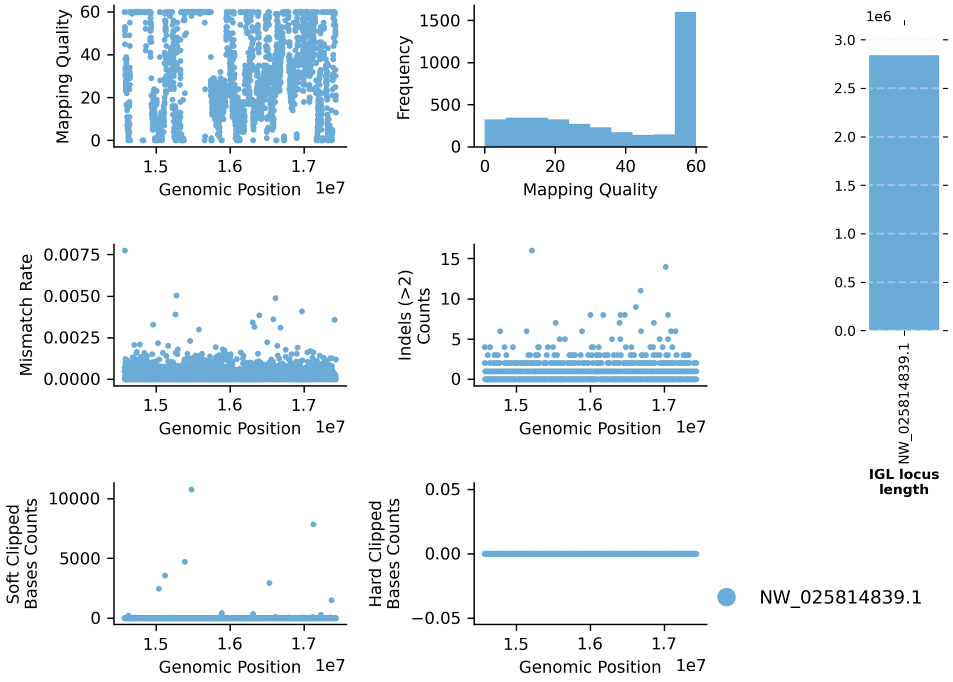


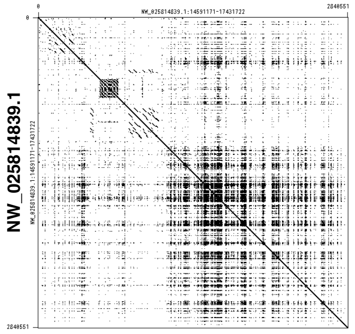

NW_025814839.1

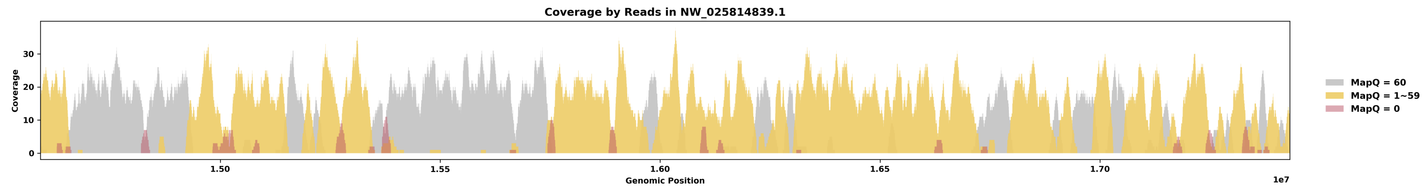


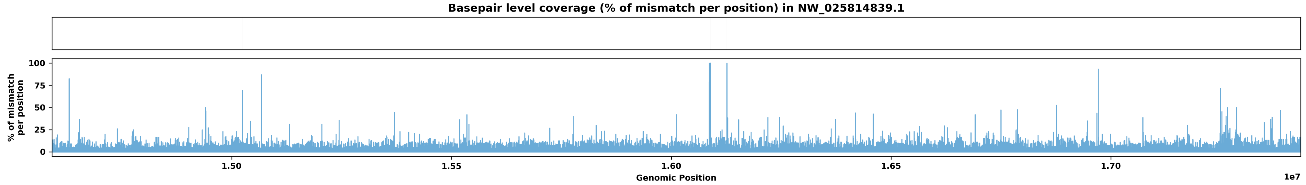

Species ID: mMacEug1

Common Name: tammar wallaby

Scientific Name: Macropus eugenii

Assembly Type: Not Haplotype Resolved

Data Source: VGP

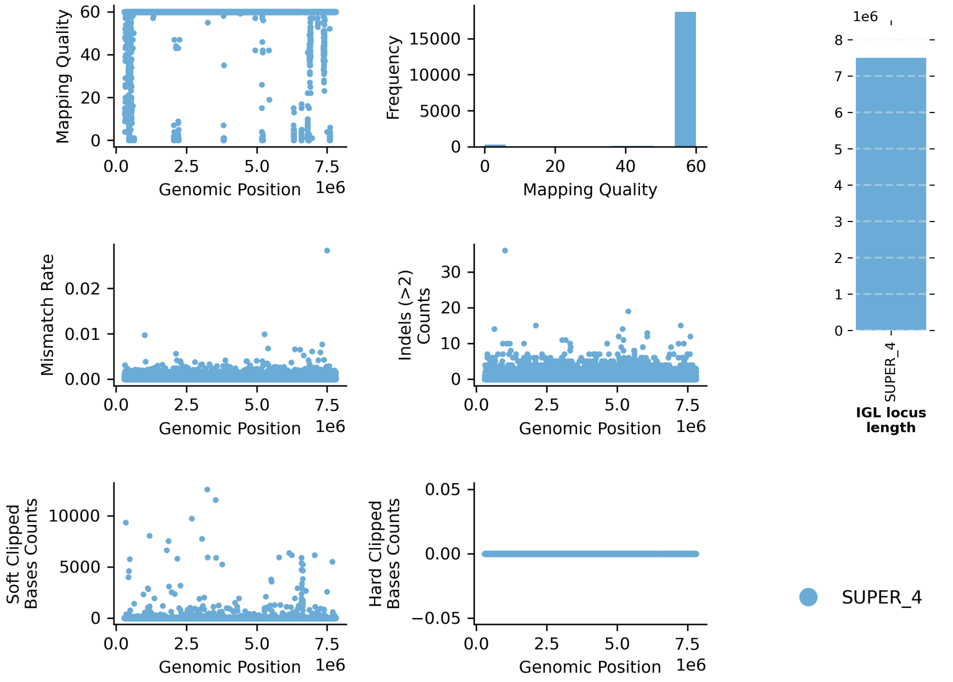


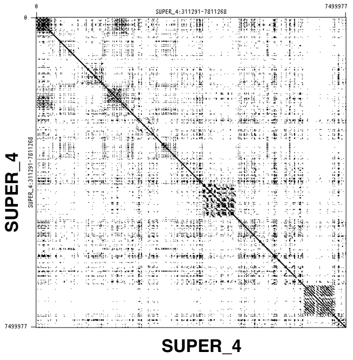


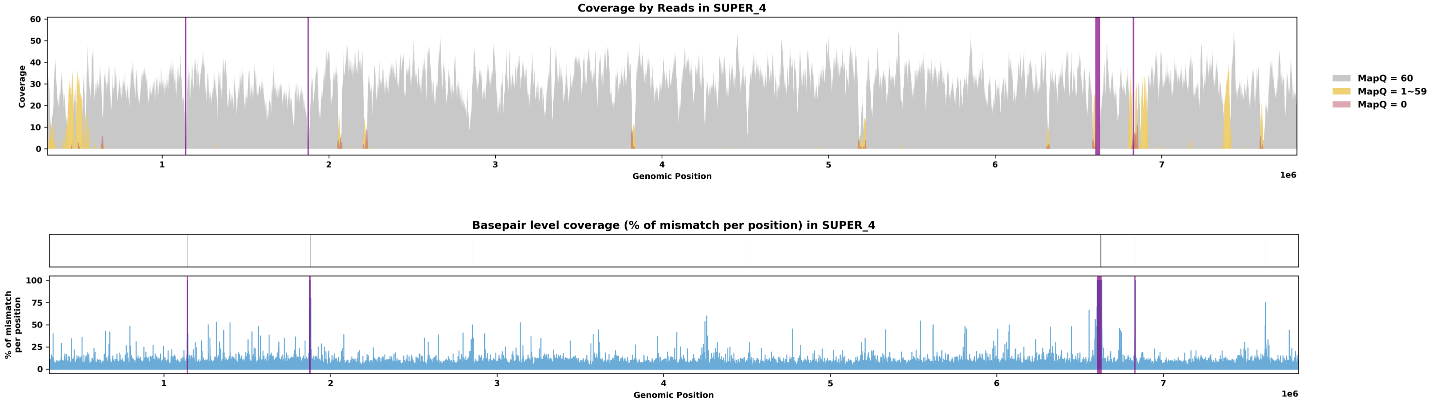

Species ID: mManPen7

Common Name: Chinese pangolin

Scientific Name: Manis pentadactyla

Assembly Type: Haplotype Resolved

Data Source: VGP

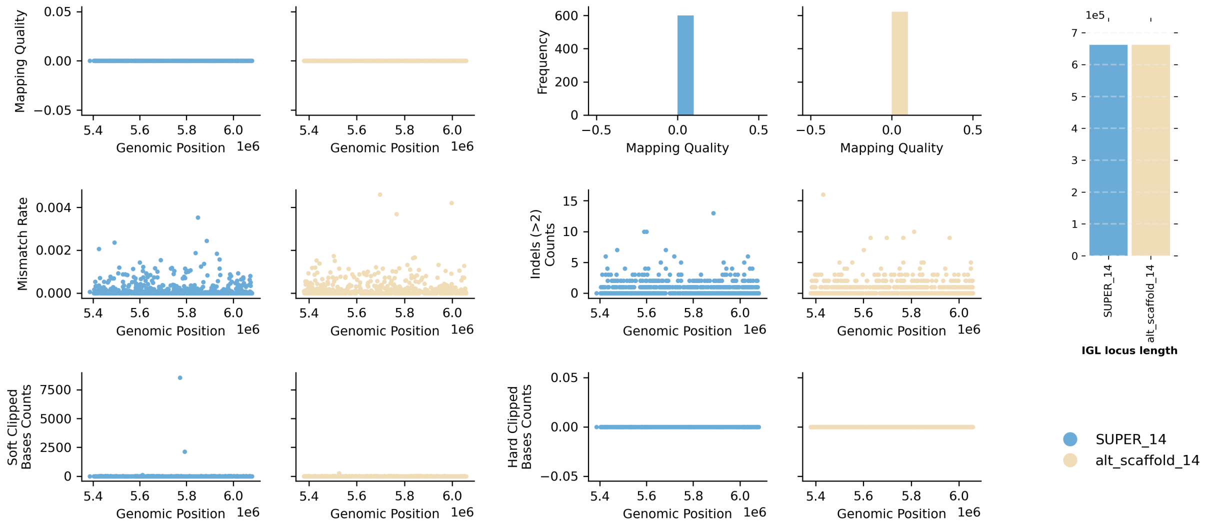


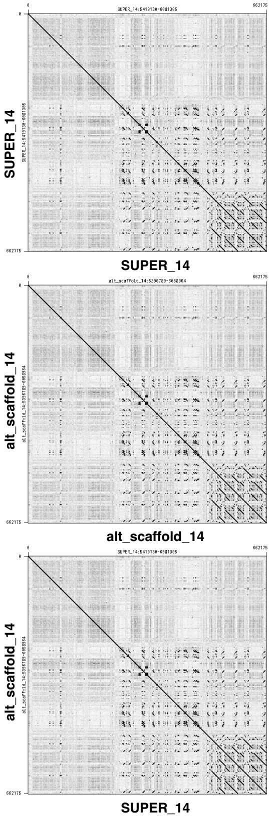


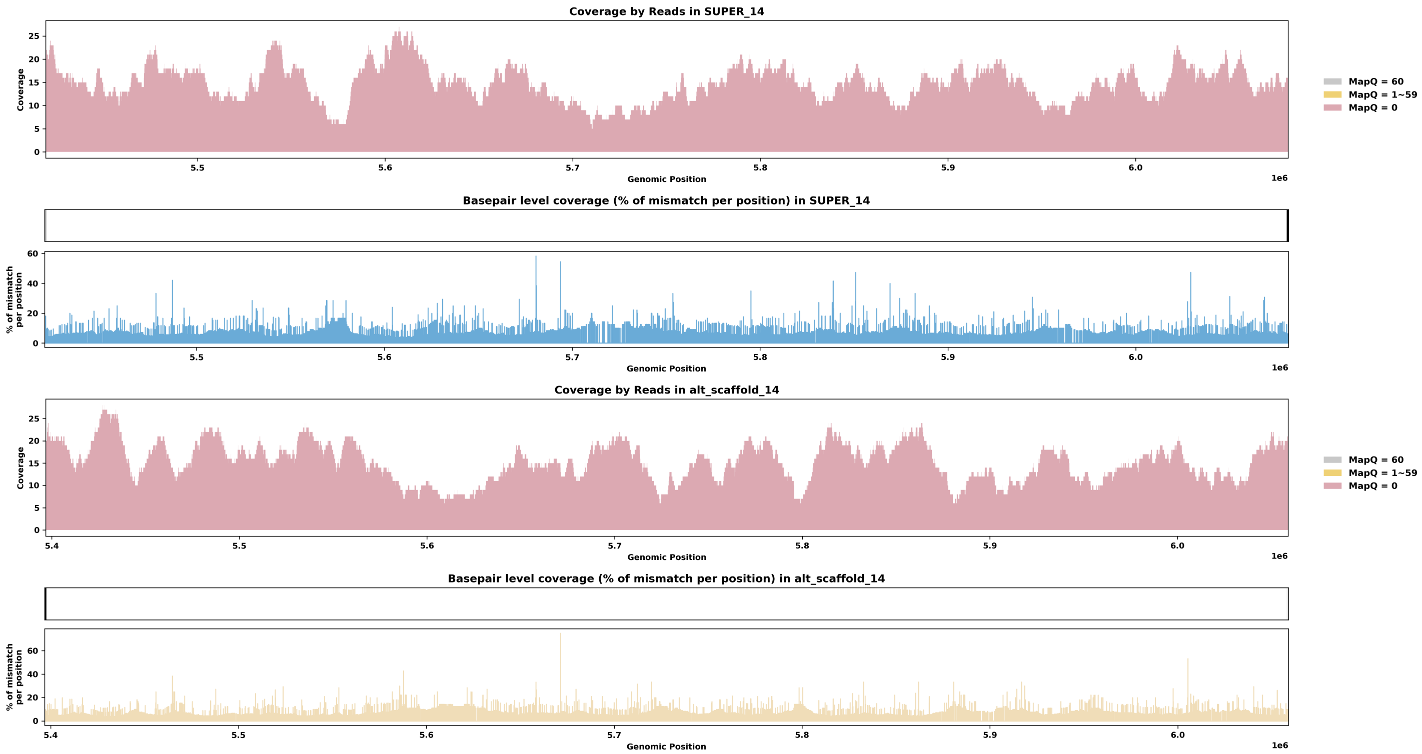

Species ID: mMarMar1

Common Name: European pine marten

Scientific Name: Martes martes

Assembly Type: Not Haplotype Resolved

Data Source: VGP

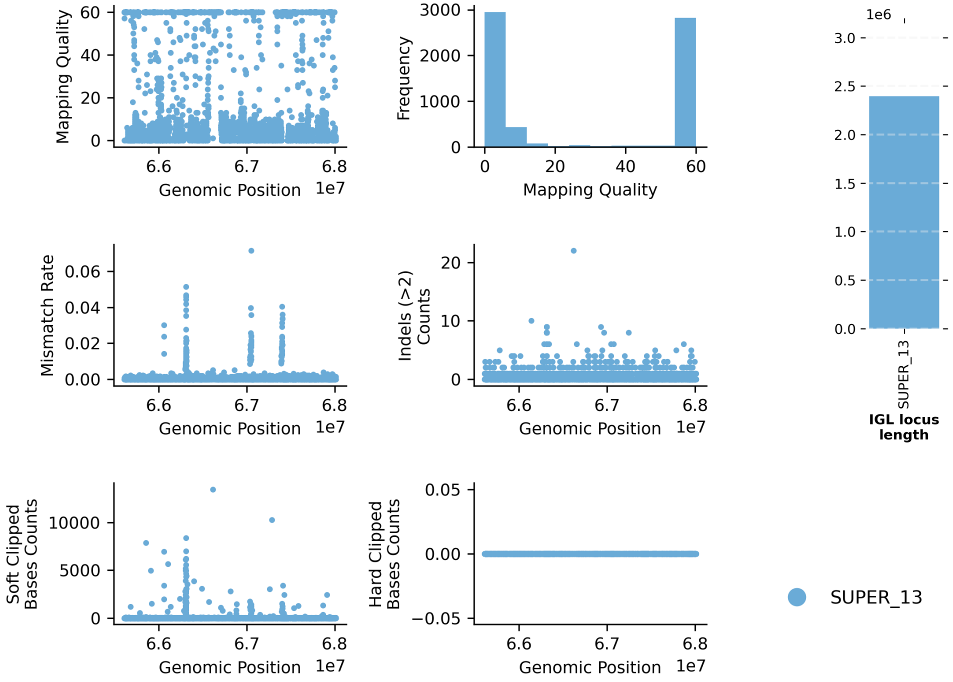


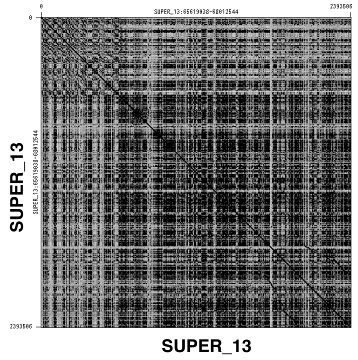


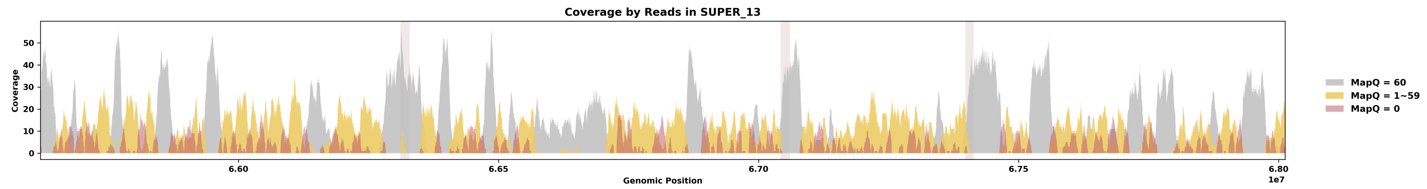


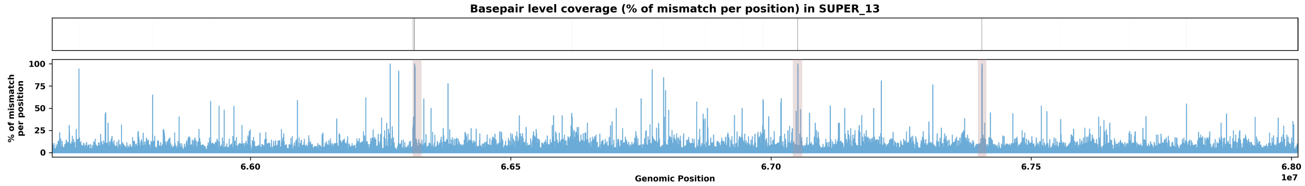

Species ID: mMelMel3

Common Name: European badger

Scientific Name: Meles meles

Assembly Type: Haplotype Resolved

Data Source: VGP

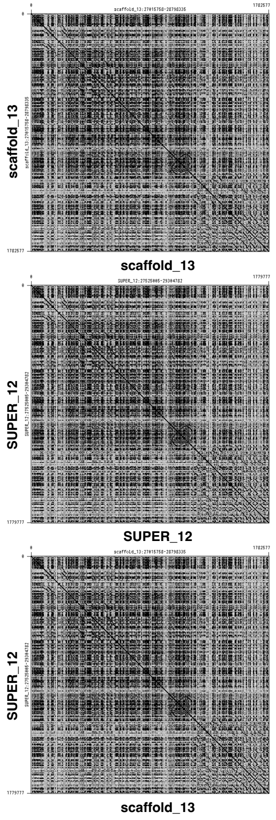


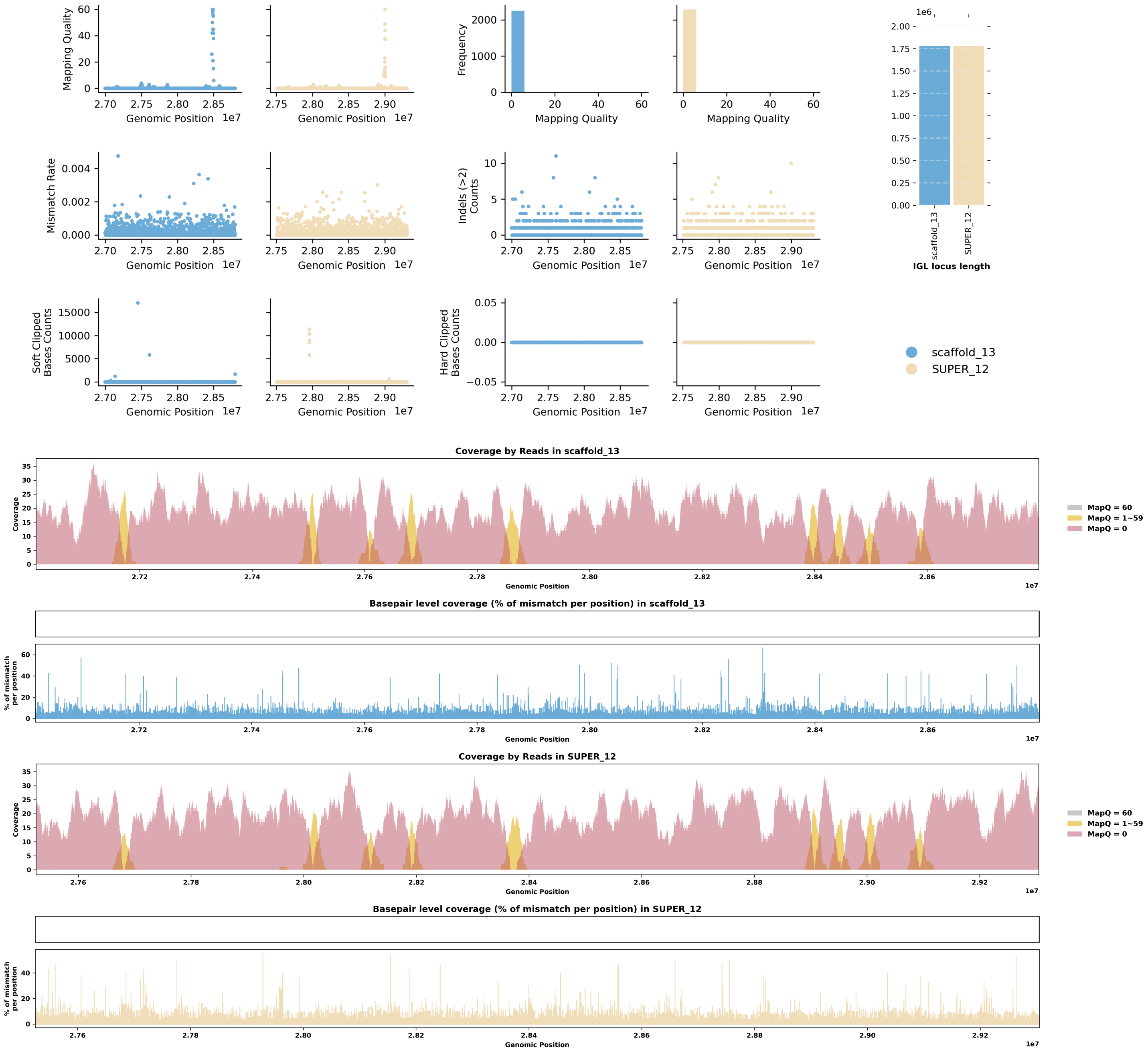

Species ID: mMesDen1

Common Name: Blainville's beaked whale

Scientific Name: Mesoplodon densirostris

Assembly Type: Not Haplotype Resolved

Data Source: VGP

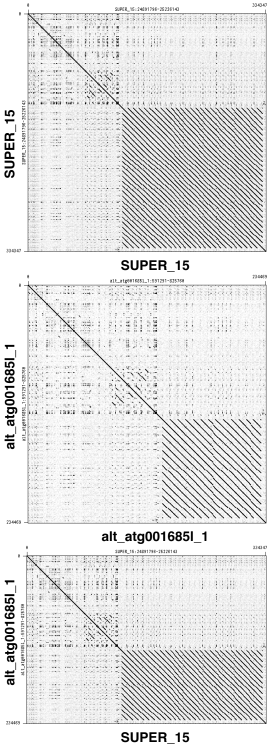


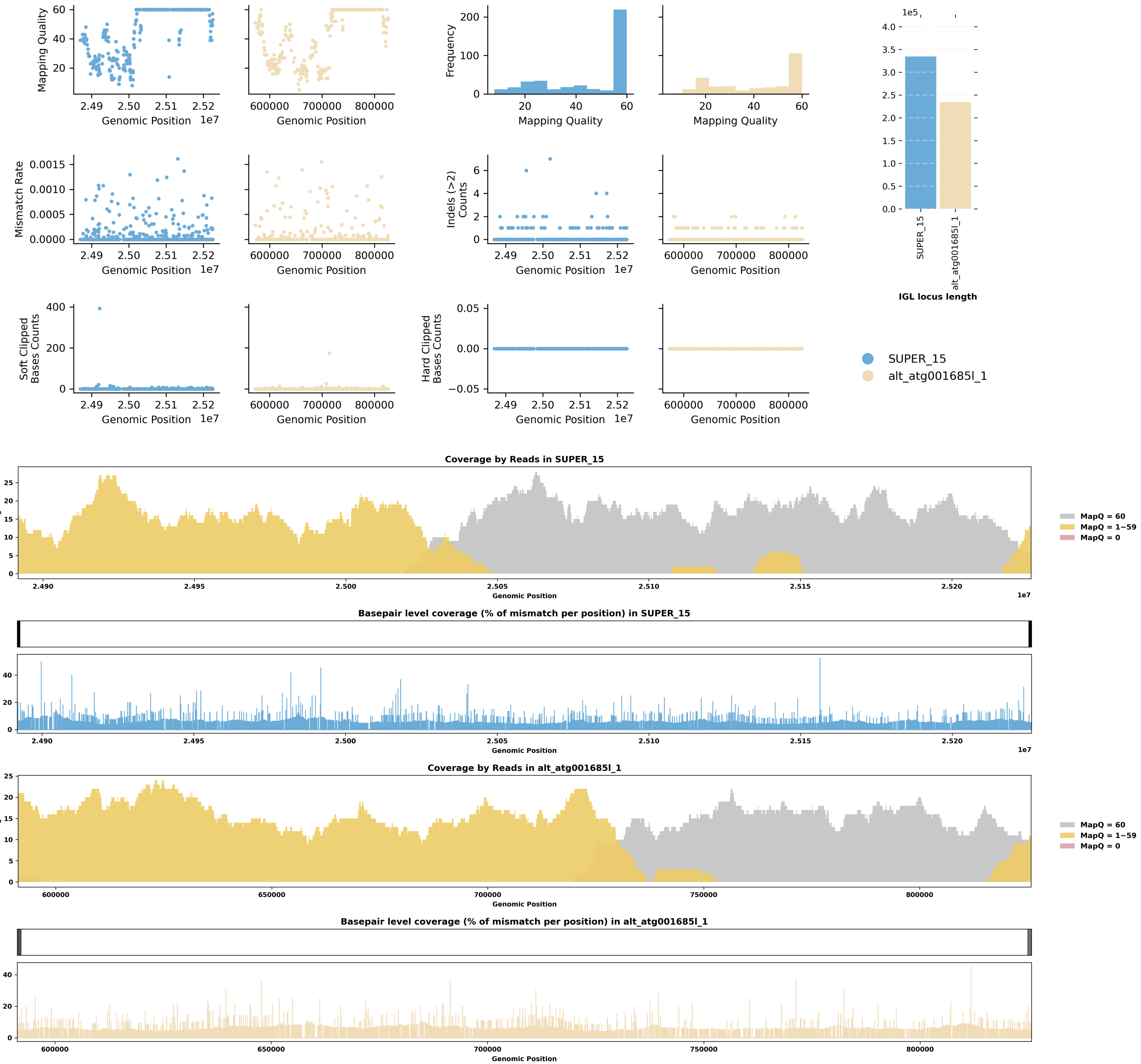

Species ID: mMicCal1

Common Name: California Vole

Scientific Name: Microtus californicus

Assembly Type: Haplotype Resolved

Data Source: CCGP

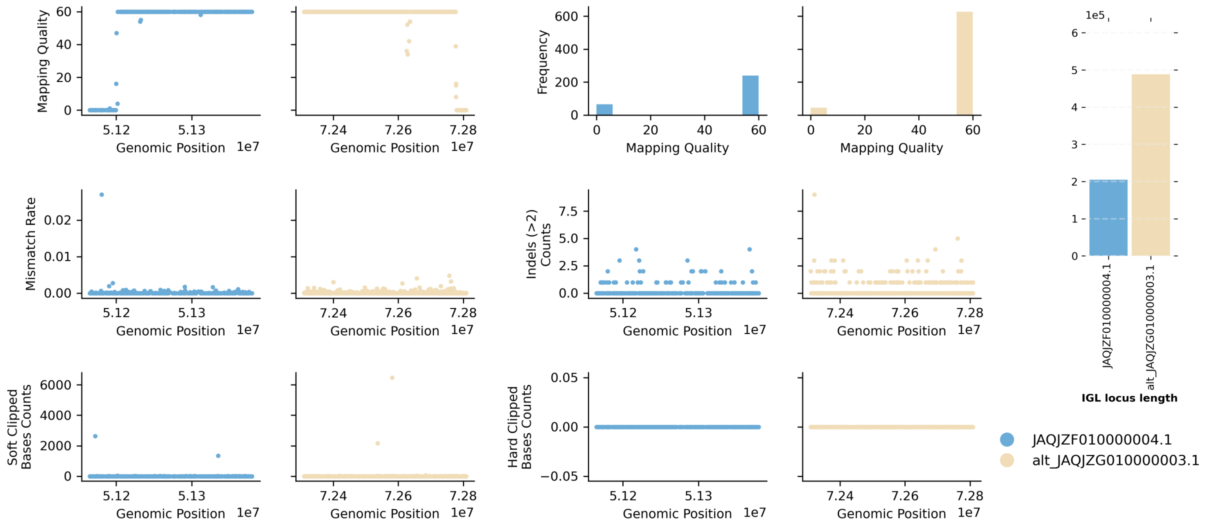


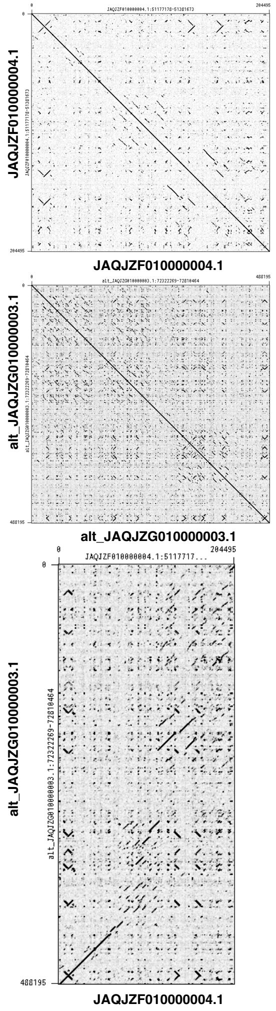


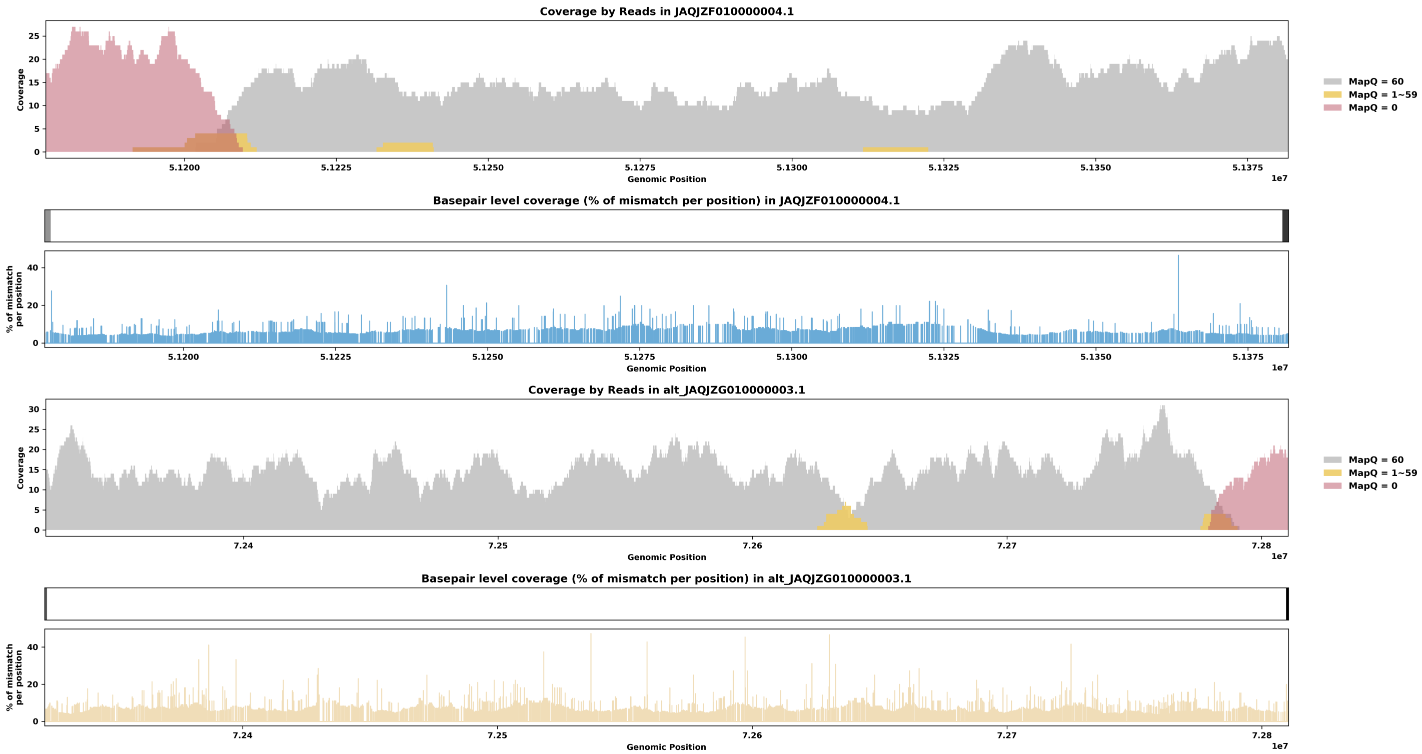

Species ID: mMicMin1

Common Name: European harvest mouse

Scientific Name: Micromys minutus

Assembly Type: Not Haplotype Resolved

Data Source: VGP

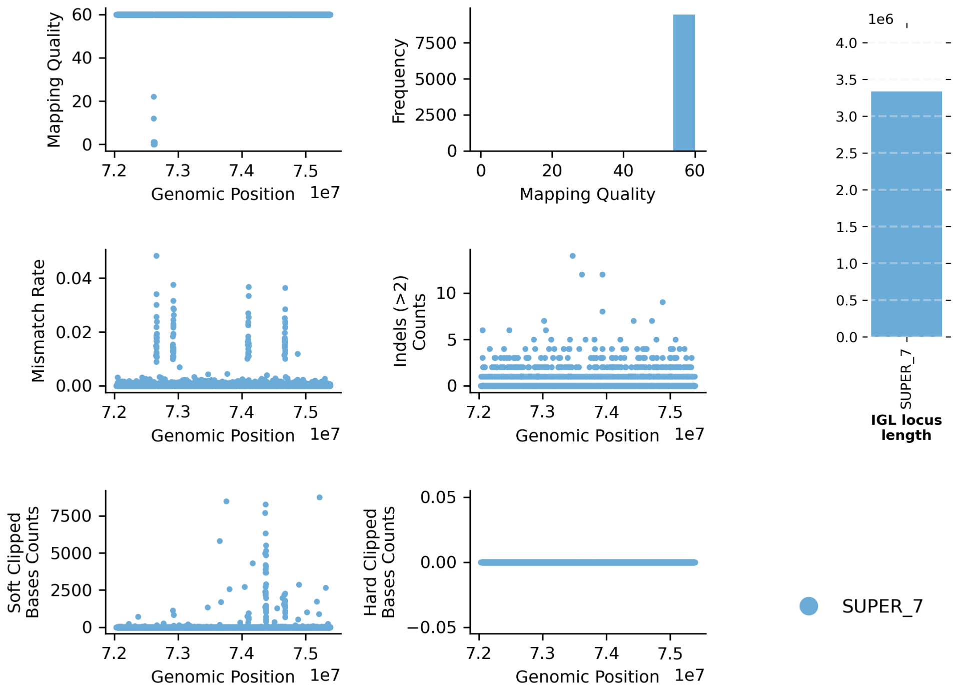


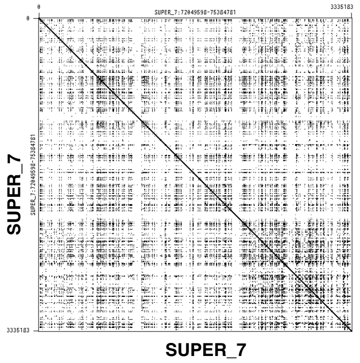


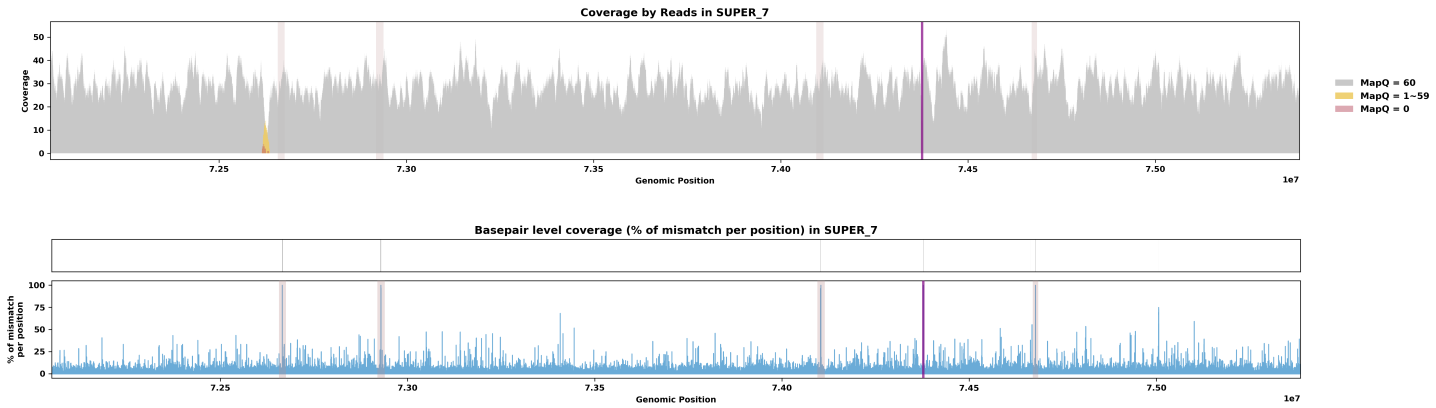

Species ID: mMirAng1

Common Name: Northern Elephant Seal

Scientific Name: Mirounga angustirostris

Assembly Type: Haplotype Resolved

Data Source: CCGP

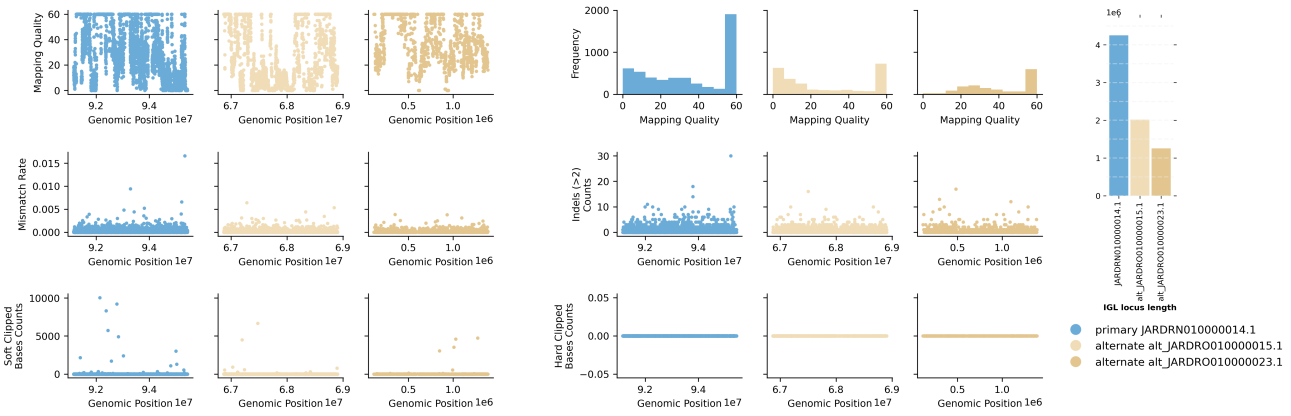


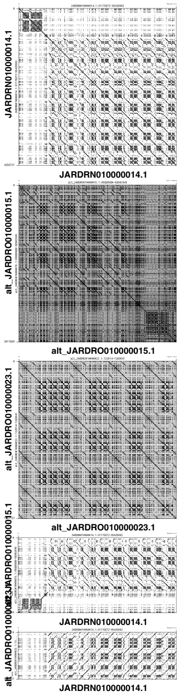


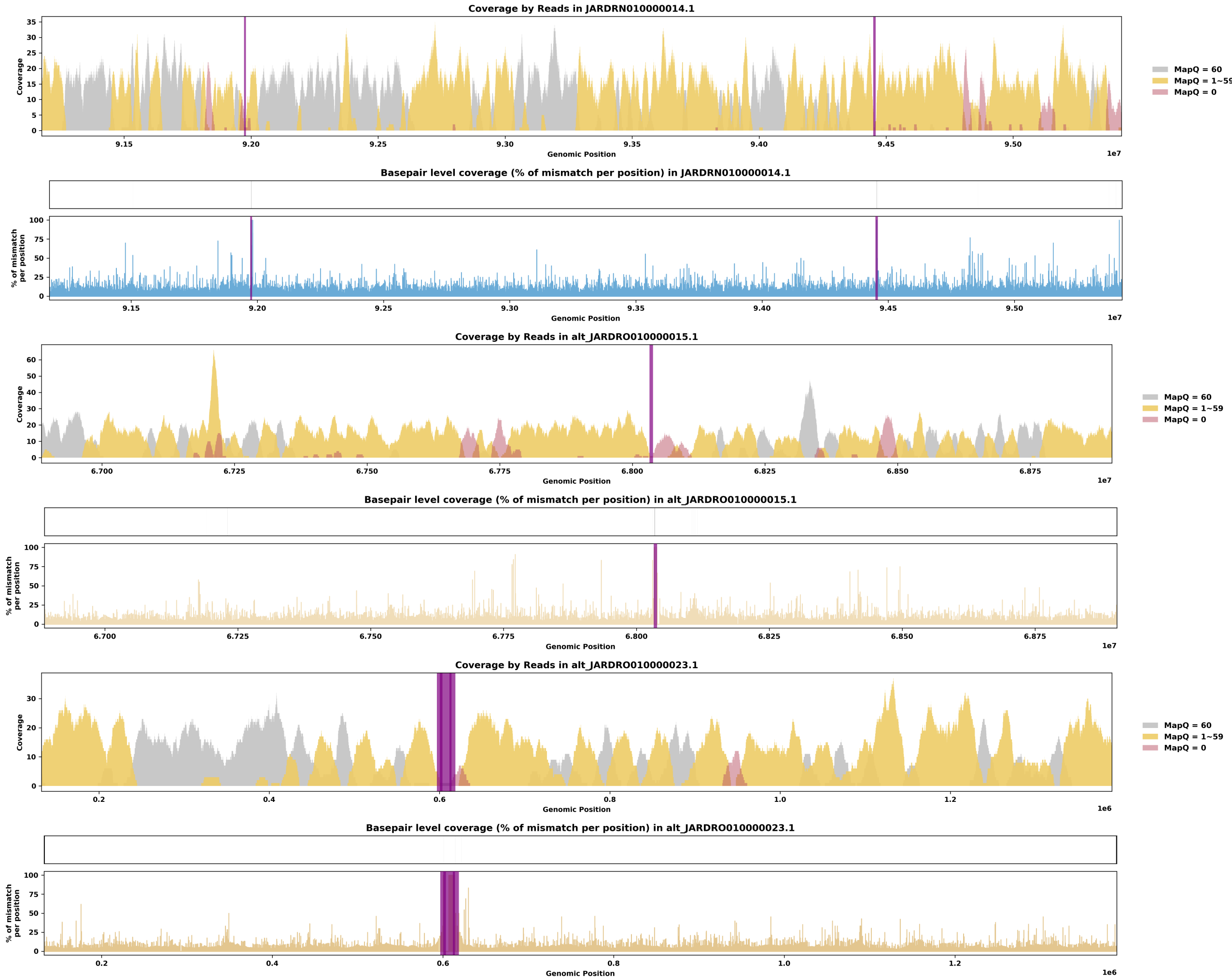

Species ID: mMonDom1

Common Name: gray short-tailed opossum

Scientific Name: Monodelphis_domestica

Assembly Type: Not Haplotype Resolved

Data Source: VGP

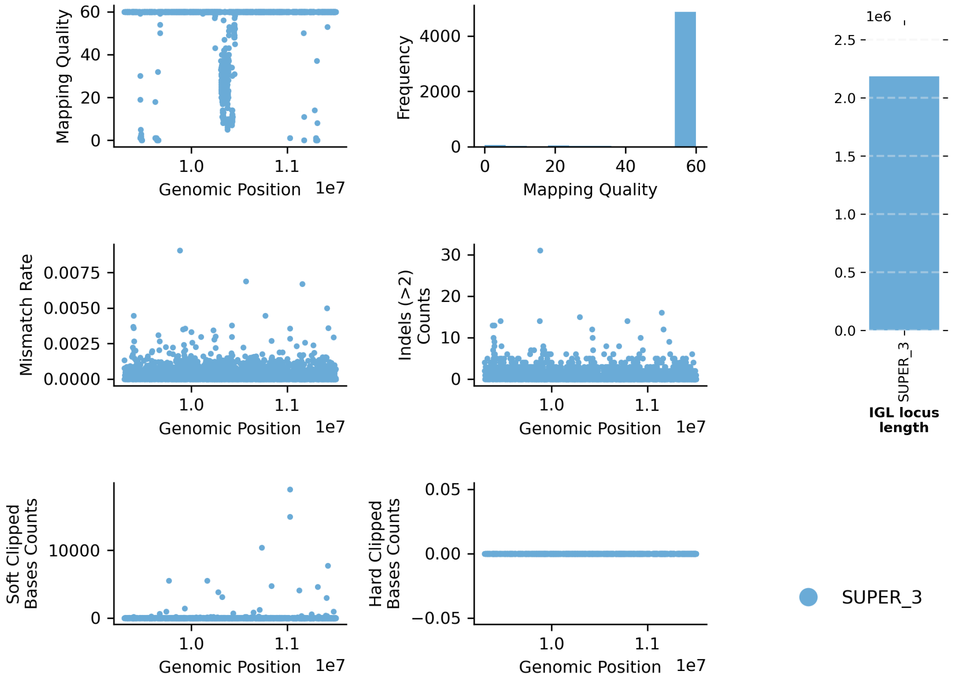


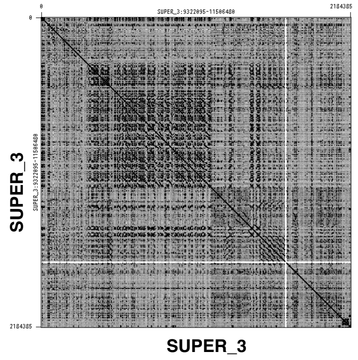


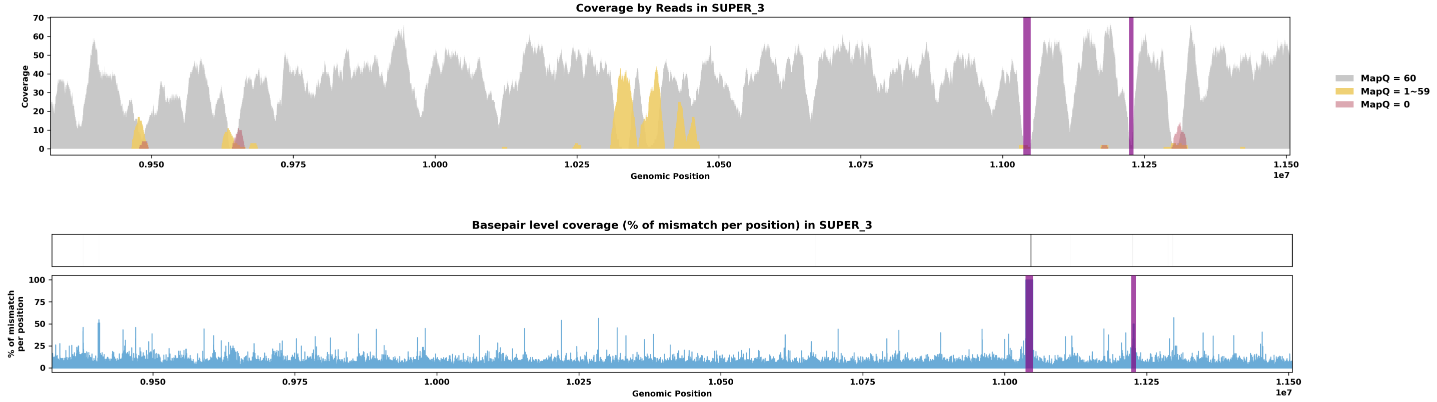

Species ID: mMunRee1

Common Name: Reeves' muntjac

Scientific Name: Muntiacus reevesi

Assembly Type: Not Haplotype Resolved

Data Source: VGP

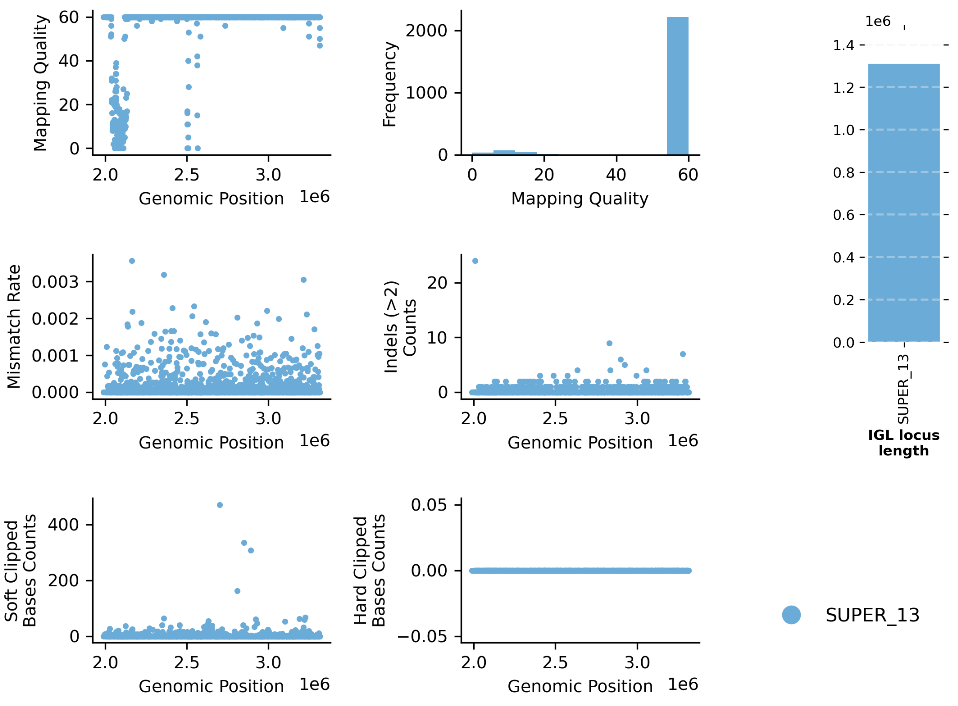


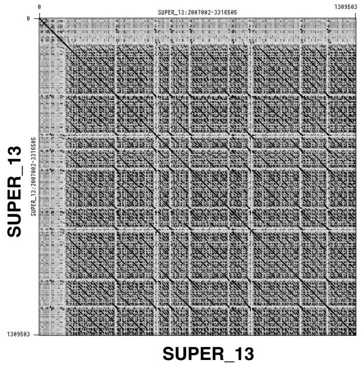


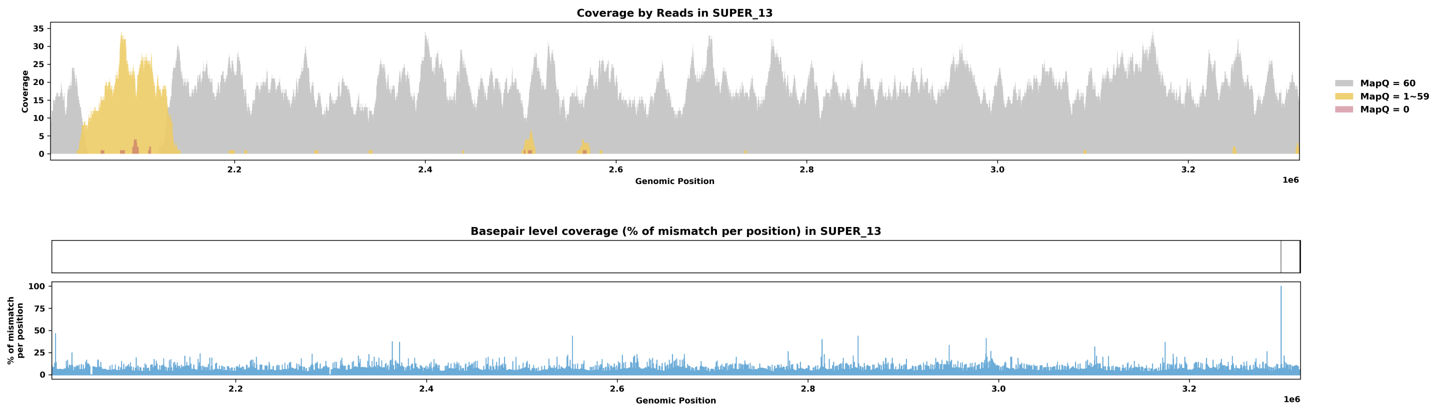

Species ID: mMusAve1

Common Name: hazel dormouse

Scientific Name: Muscardinus avellanarius

Assembly Type: Not Haplotype Resolved

Data Source: VGP

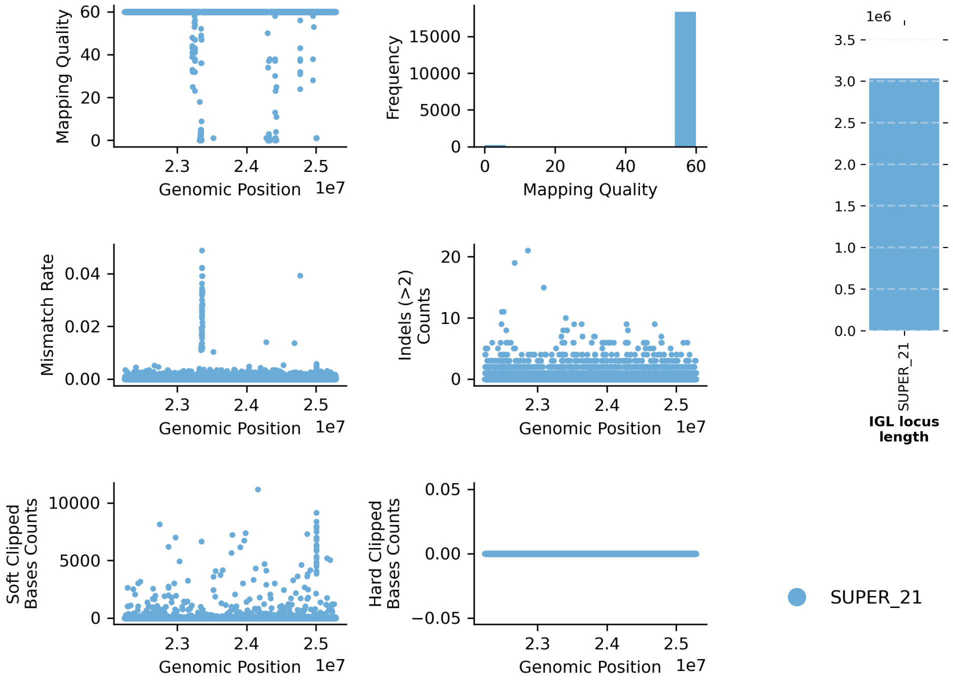


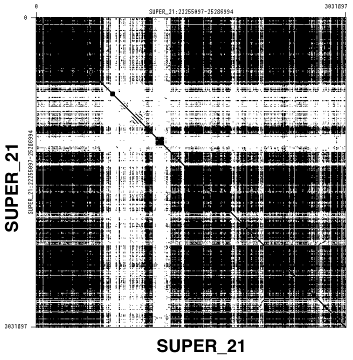


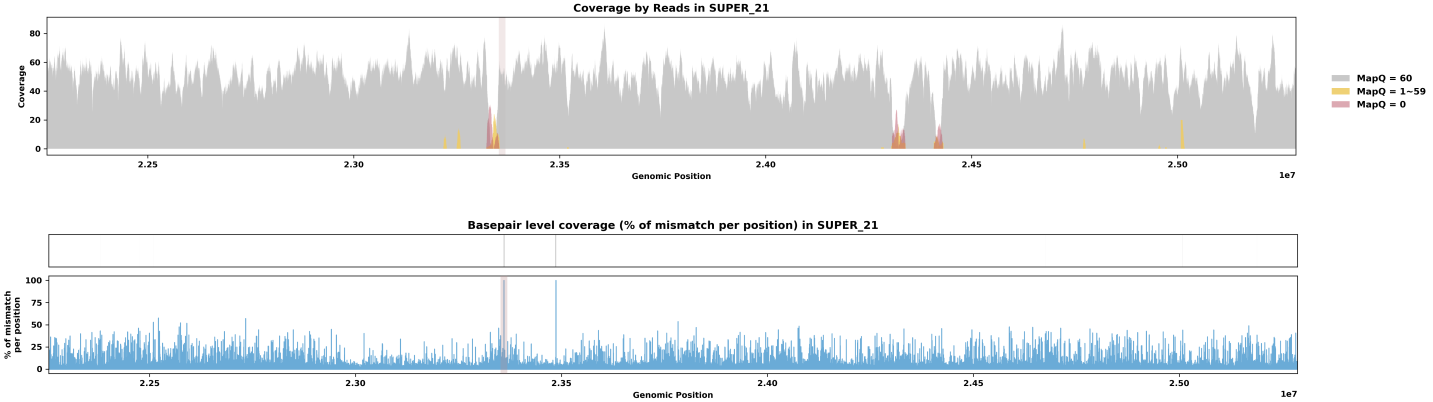

Species ID: mMusLut2

Common Name: European mink

Scientific Name: Mustela lutreola

Assembly Type: Not Haplotype Resolved

Data Source: VGP

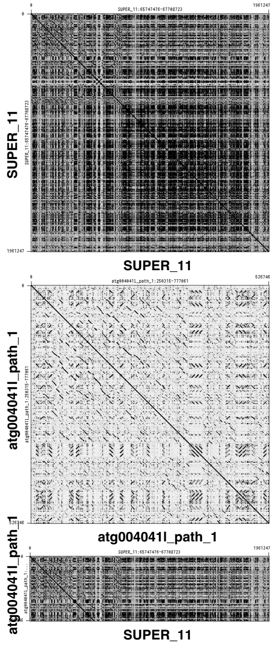


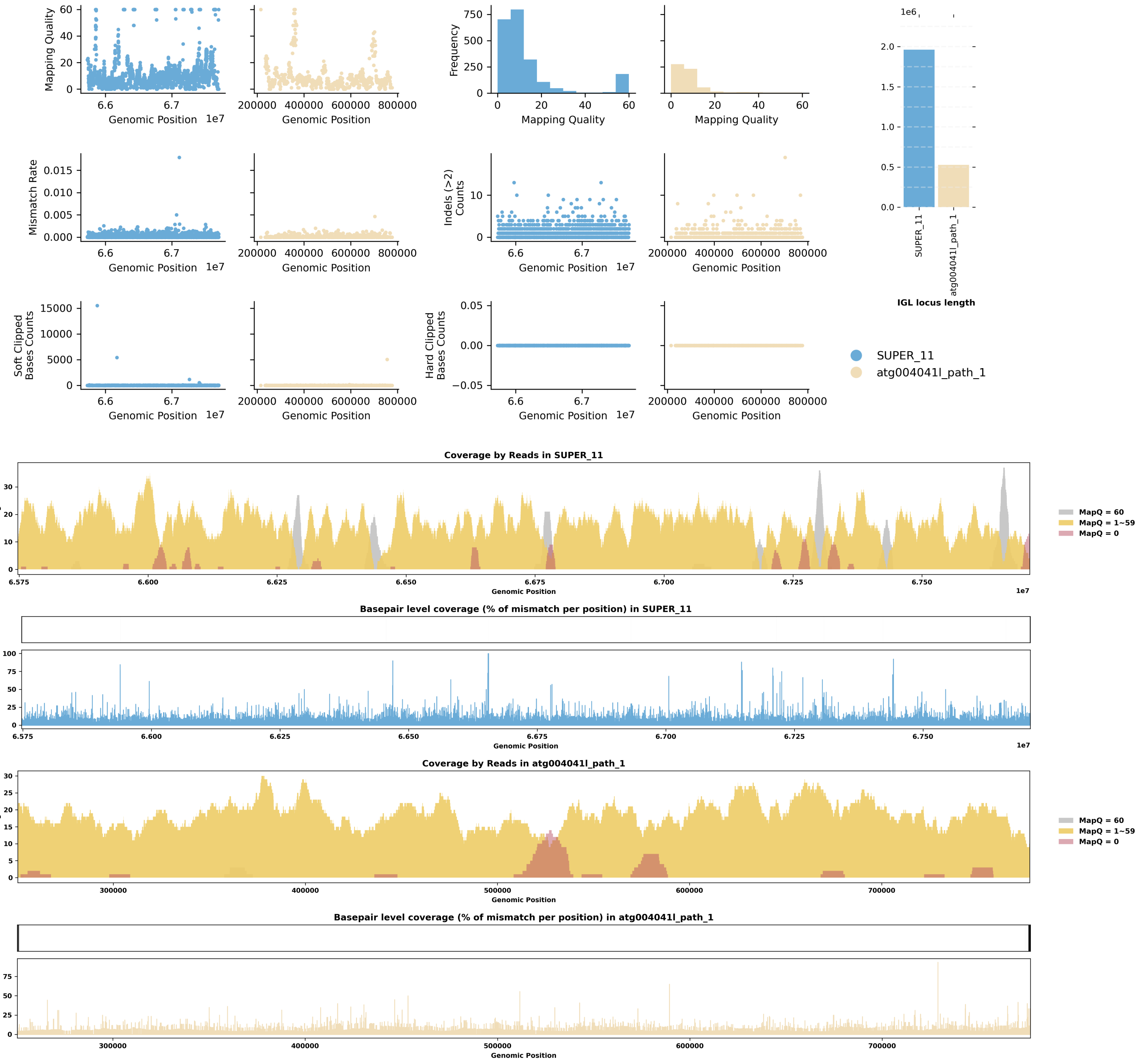

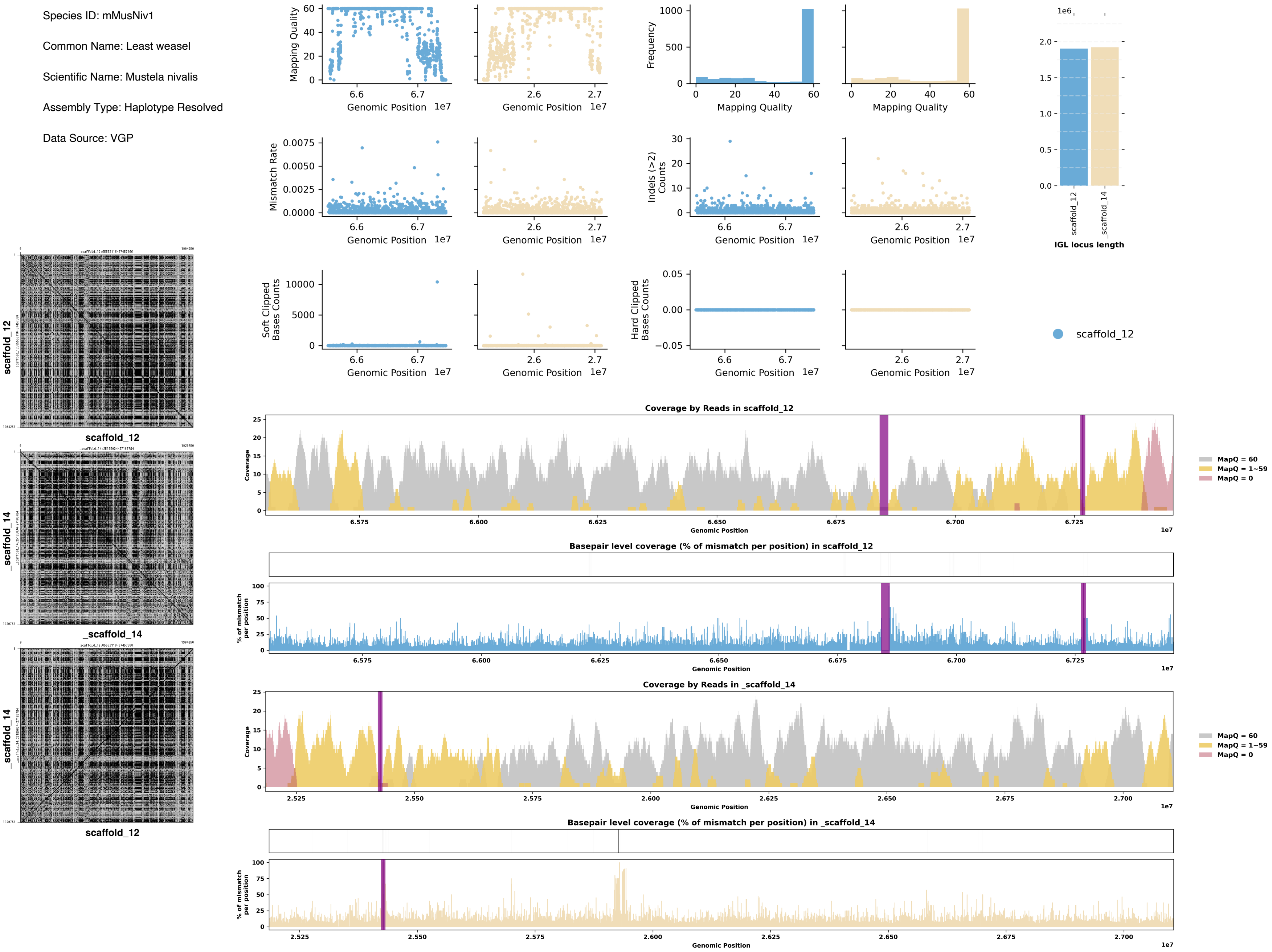
Species ID: mMusNiv1
Common Name: Least weasel
Scientific Name: Mustela nivalis
Assembly Type: Haplotype Resolved
Data Source: VGP
Mapping Quality
Genomic Position
Frequency
Mismatch Rate
Indels (>2) Counts
Soft Clipped Bases Counts
Hard Clipped Bases Counts
IGL locus length
scaffold_12
_scaffold_14
Coverage by Reads in scaffold_12
Basepair level coverage (% of mismatch per position) in scaffold_12
Coverage by Reads in _scaffold_14
Basepair level coverage (% of mismatch per position) in _scaffold_14
Coverage
% of mismatch per position
MapQ = 60
MapQ = 1~59
MapQ = 0

Species ID: mMyoDau2

Common Name: Daubenton's bat

Scientific Name: Myotis daubentonii

Assembly Type: Not Haplotype Resolved

Data Source: VGP

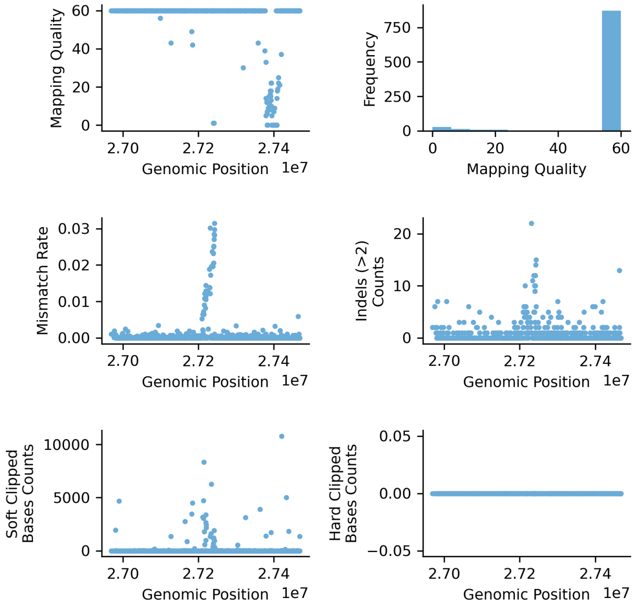


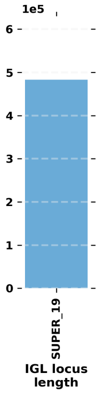


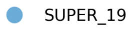


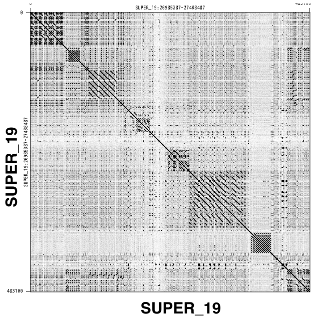


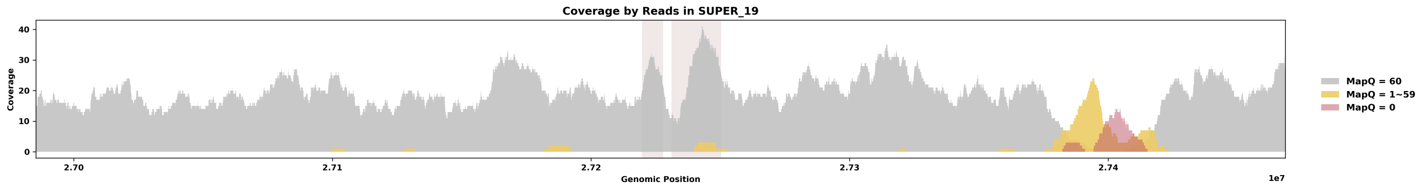


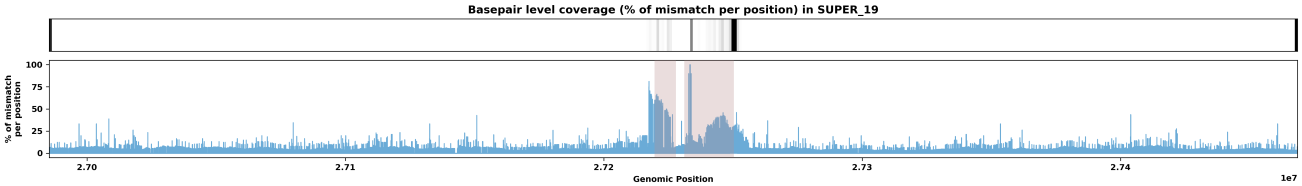

Species ID: mMyoYum1

Common Name: Yuma myotis

Scientific Name: Myotis yumanensis

Assembly Type: Haplotype Resolved

Data Source: CCGP

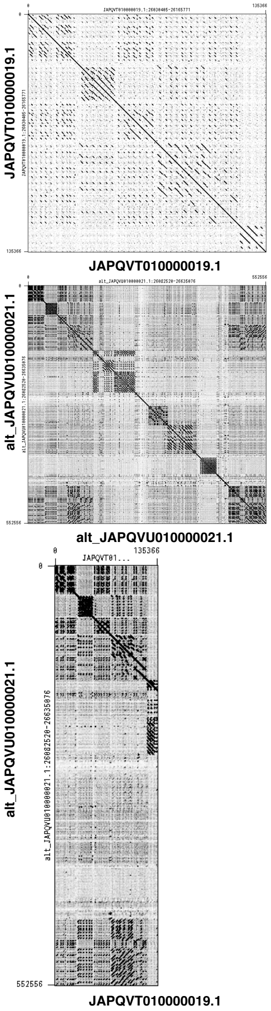


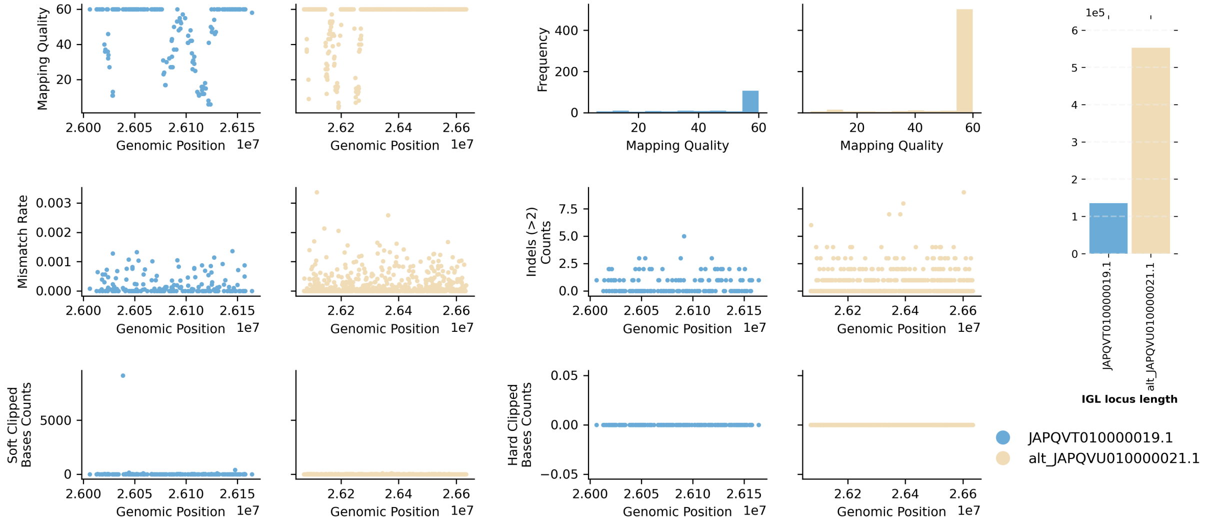


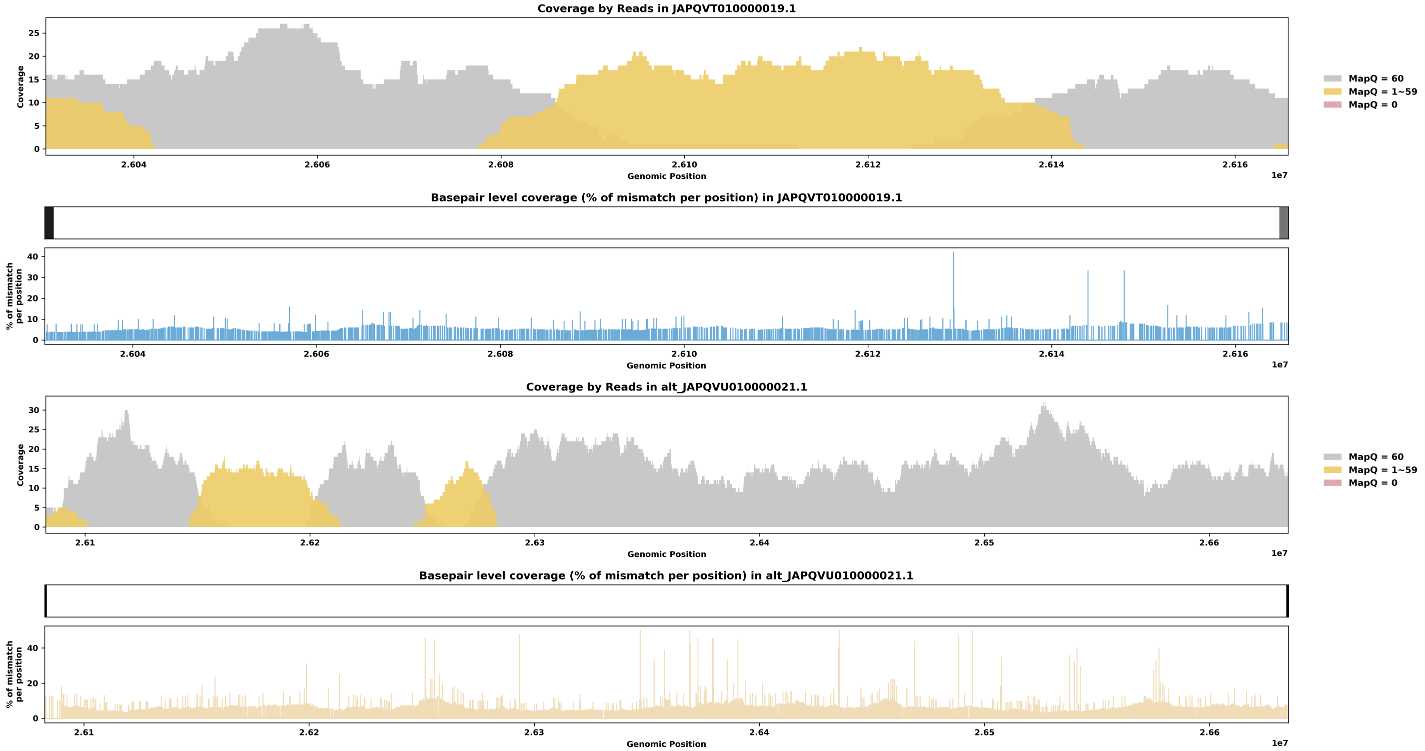

Species ID: mNeoNeb1

Common Name: Clouded Leopard

Scientific Name: Neofelis_nebulosa

Assembly Type: Not Haplotype Resolved

Data Source: VGP

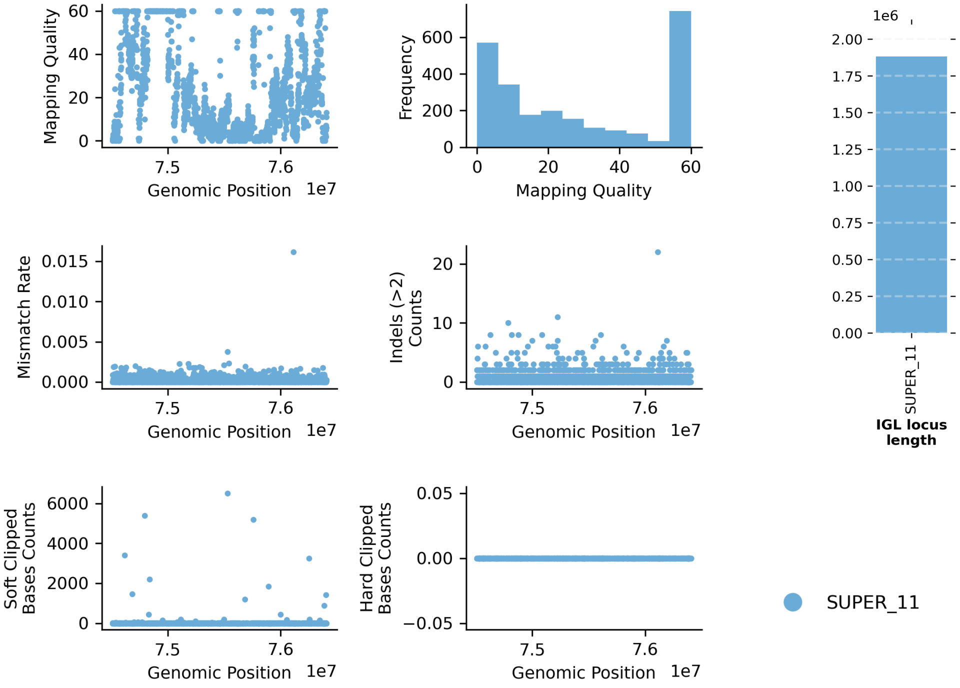


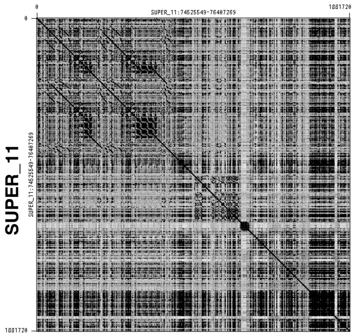


SUPER_11

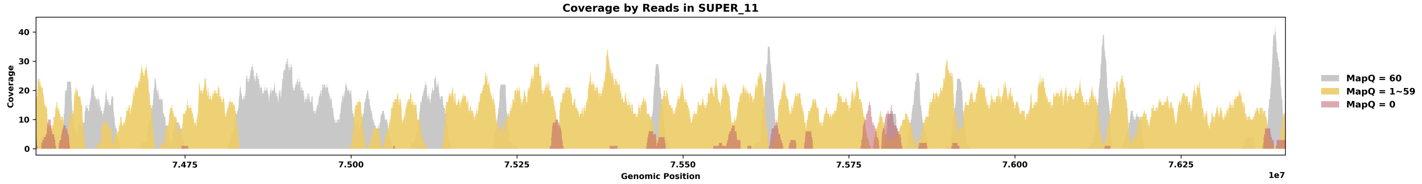


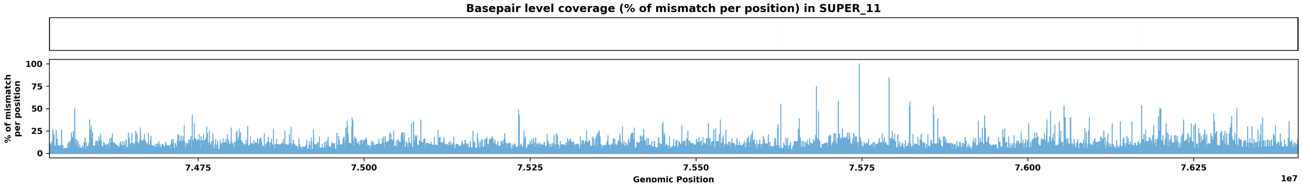

Species ID: mNycCou1

Common Name: slow loris

Scientific Name: Nycticebus coucang

Assembly Type: Not Haplotype Resolved

Data Source: VGP

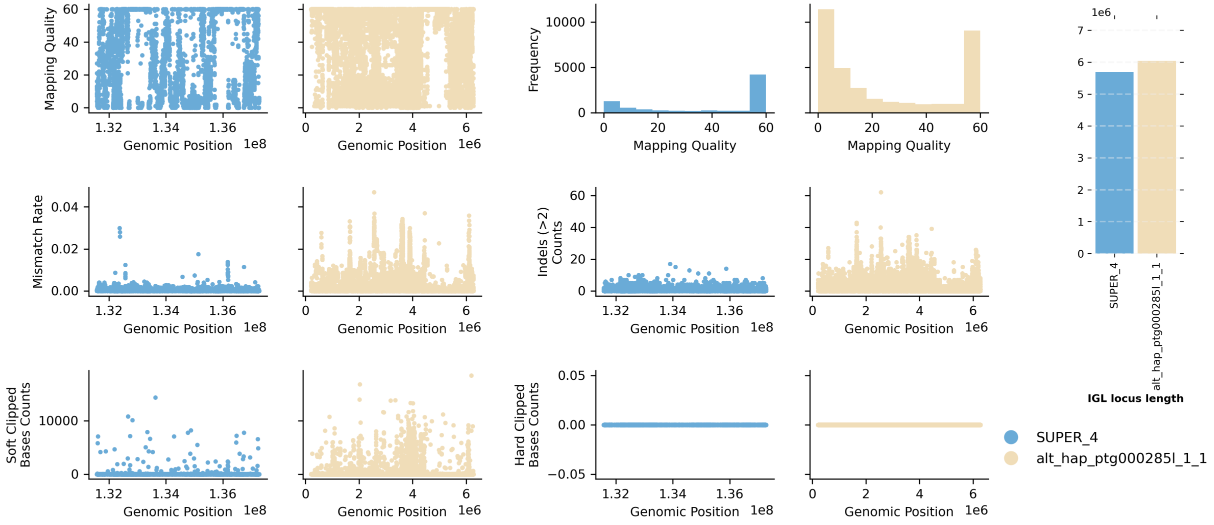


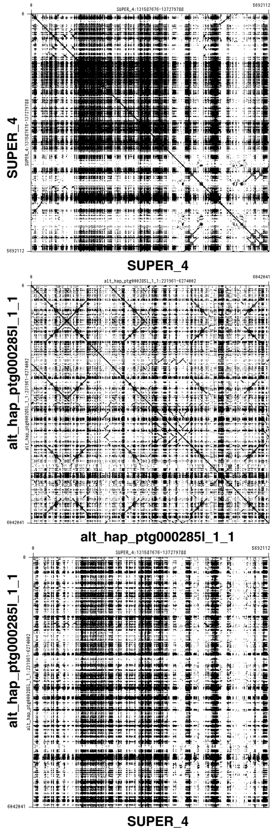


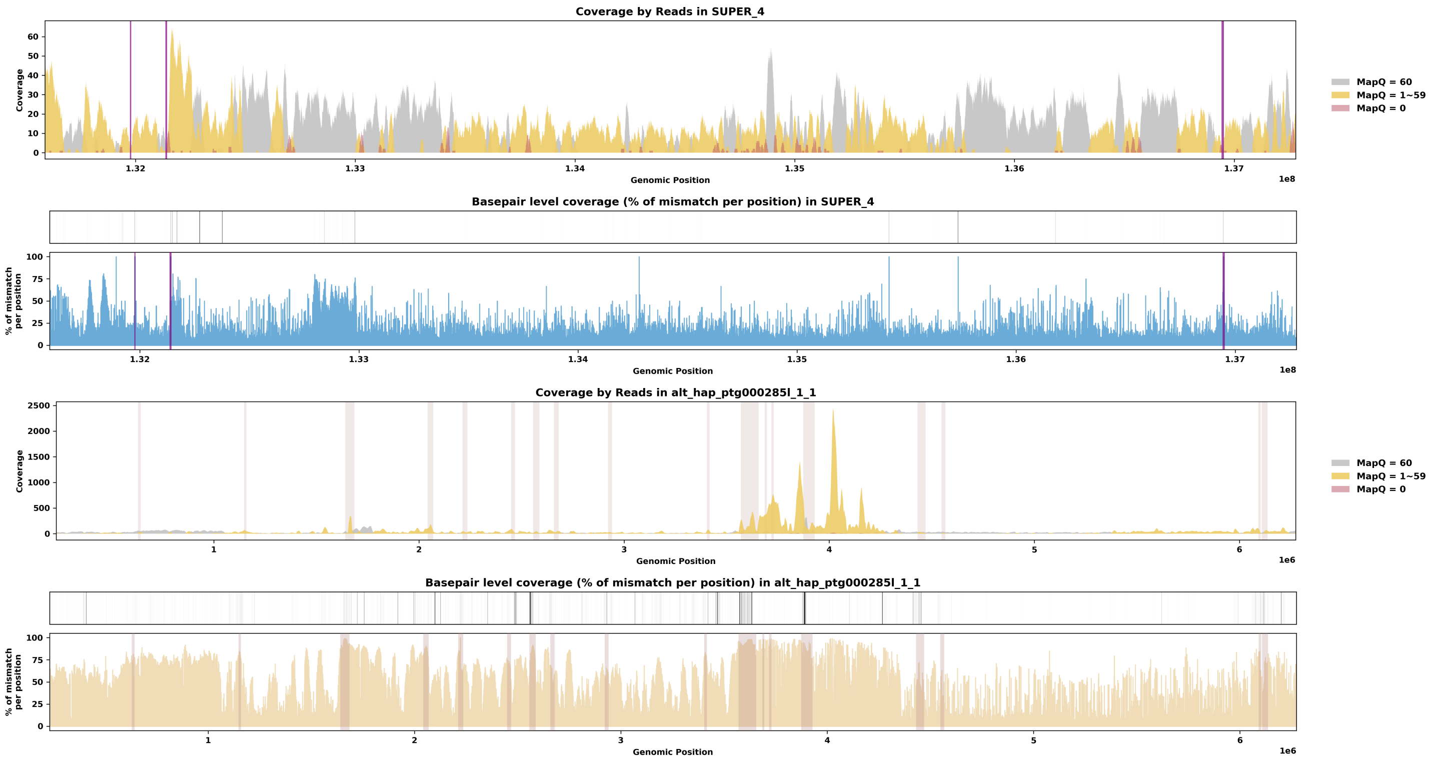

Species ID: mOrcOrc1

Common Name: killer whale

Scientific Name: Orcinus orca

Assembly Type: Not Haplotype Resolved

Data Source: VGP

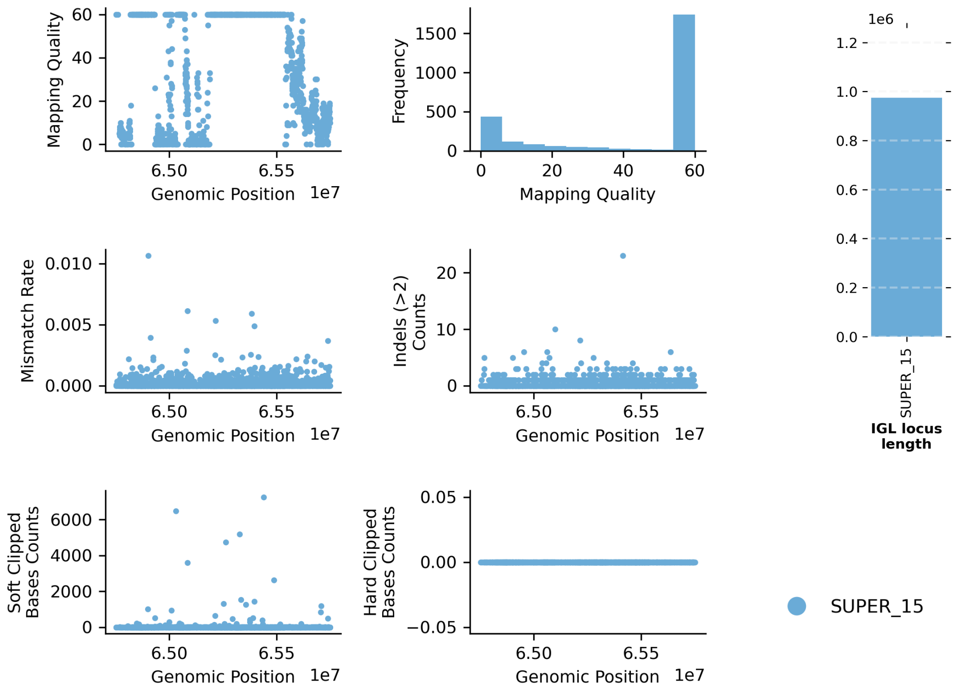


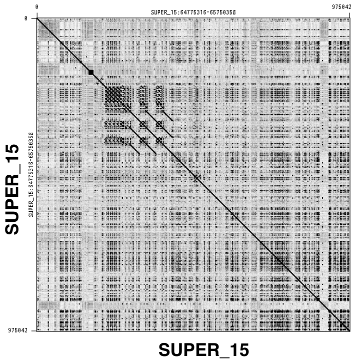


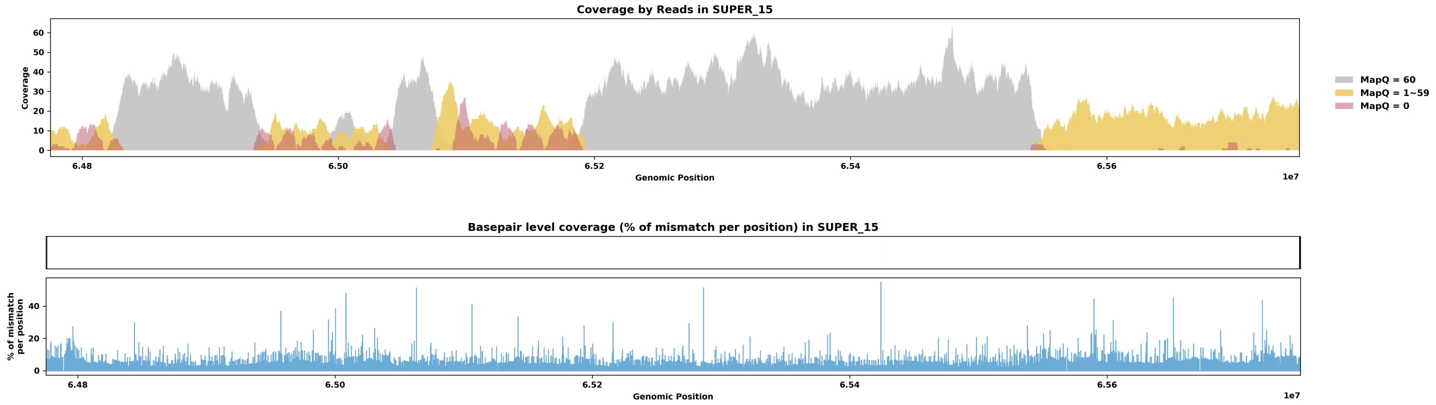

Species ID: mOryCun1

Common Name: rabbit

Scientific Name: Oryctolagus cuniculus

Assembly Type: Not Haplotype Resolved

Data Source: VGP

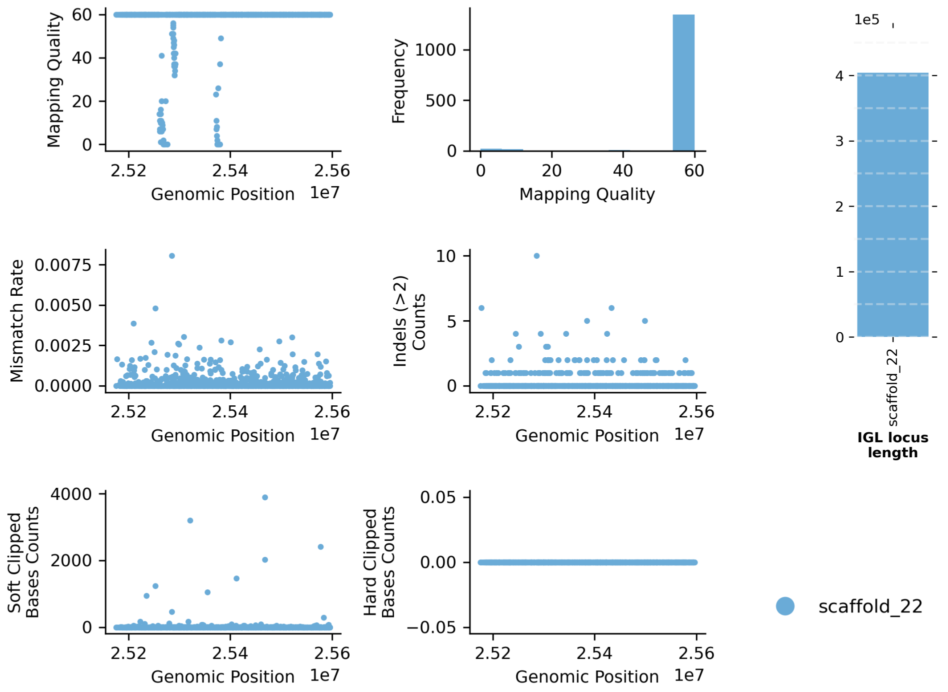


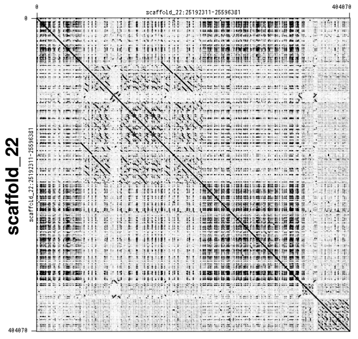


scaffold_22

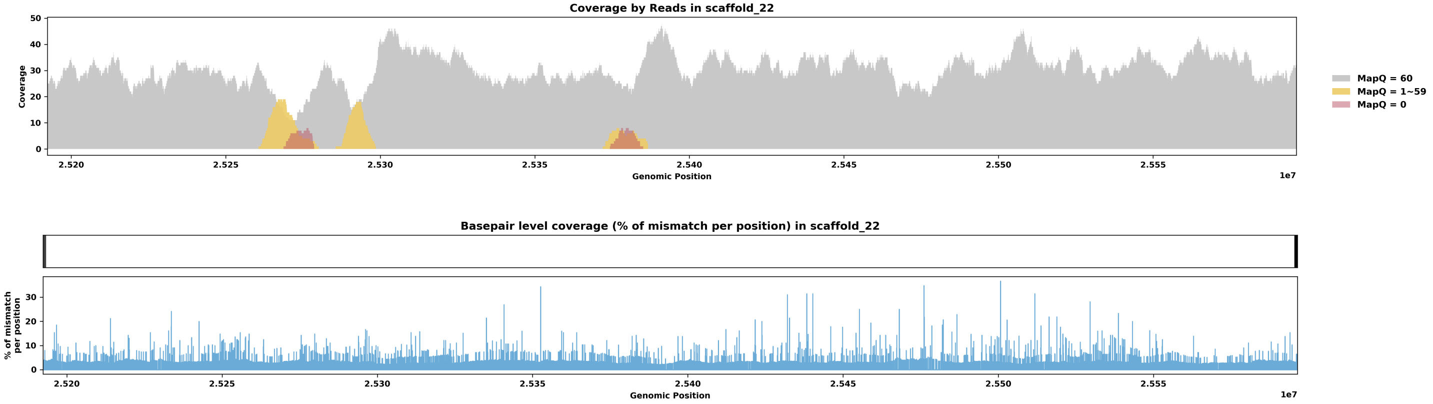

Species ID: mPanPan1

Common Name: Bonobo

Scientific Name: Pan paniscus

Assembly Type: Haplotype Resolved

Data Source: T2T Primate

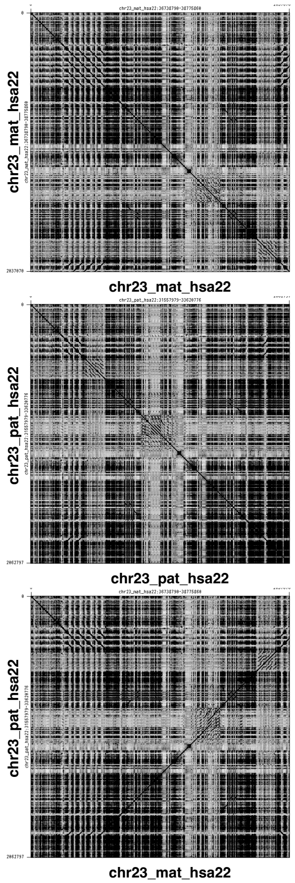


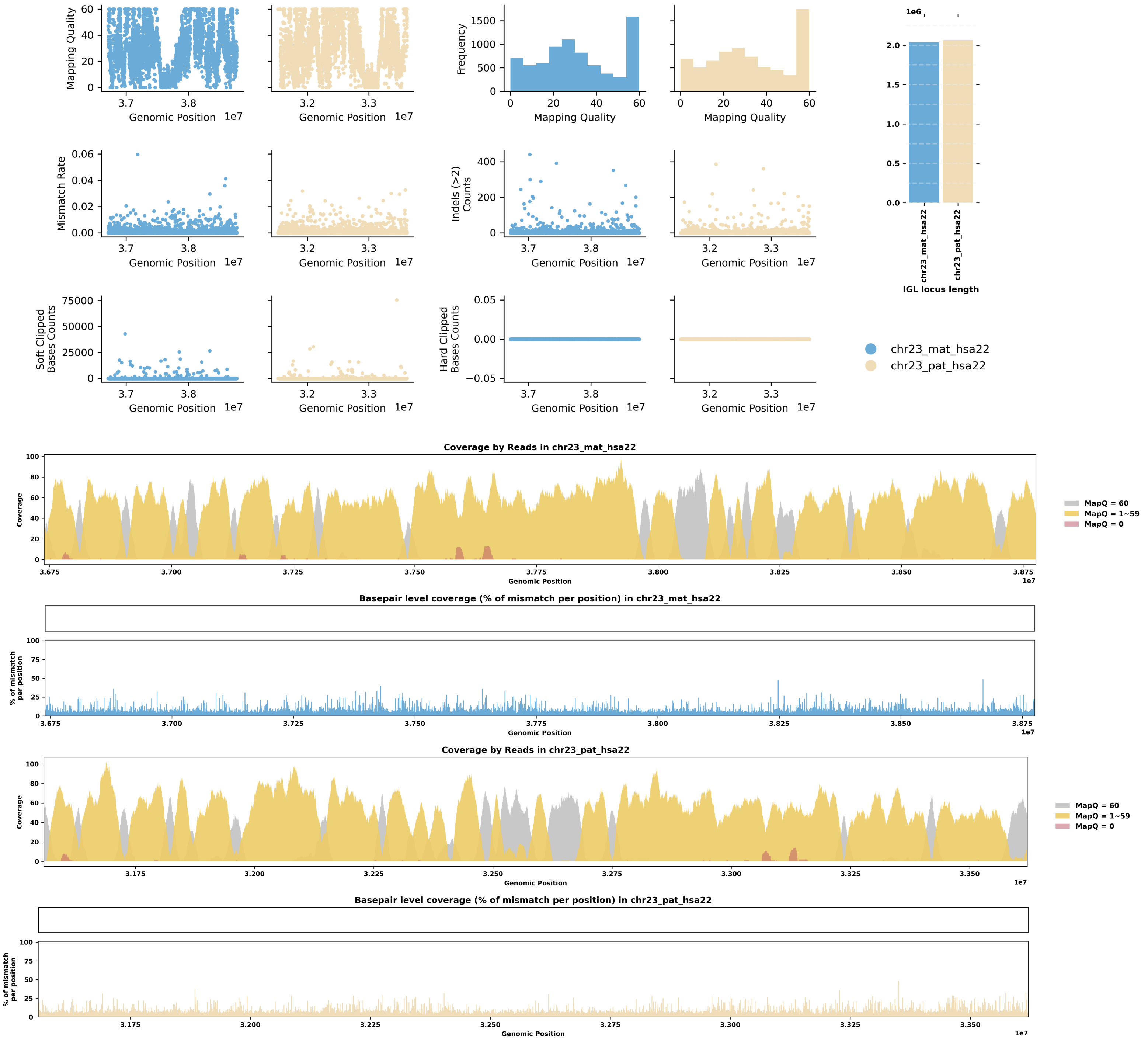

Species ID: mPerMan1

Common Name: deer mouse

Scientific Name: Peromyscus maniculatus

Assembly Type: Not Haplotype Resolved

Data Source: CCGP

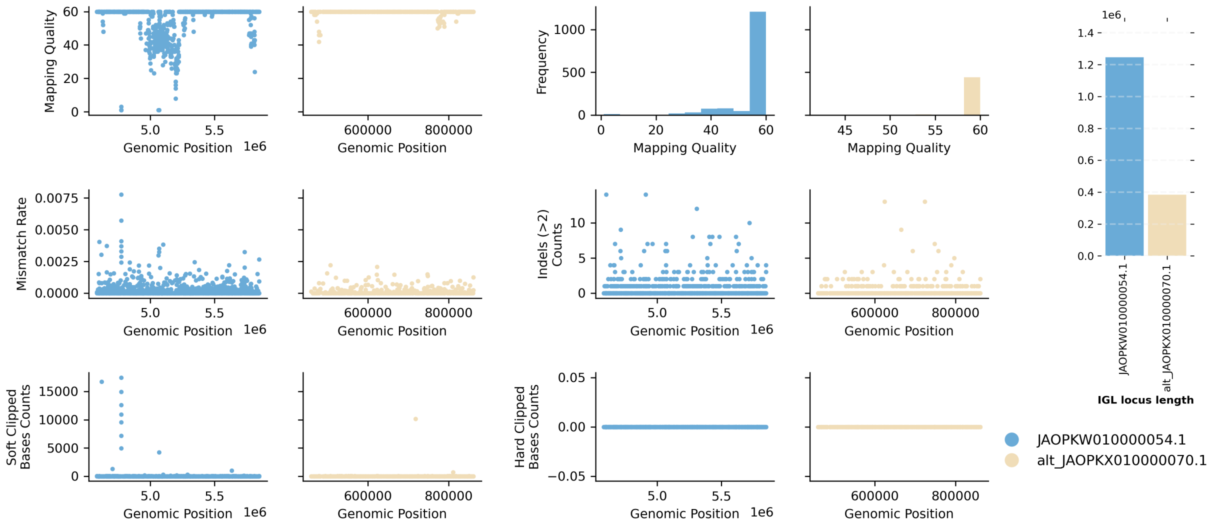


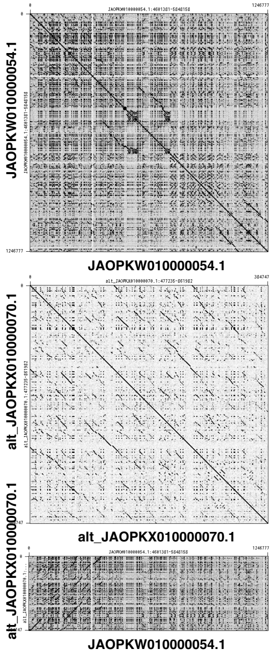


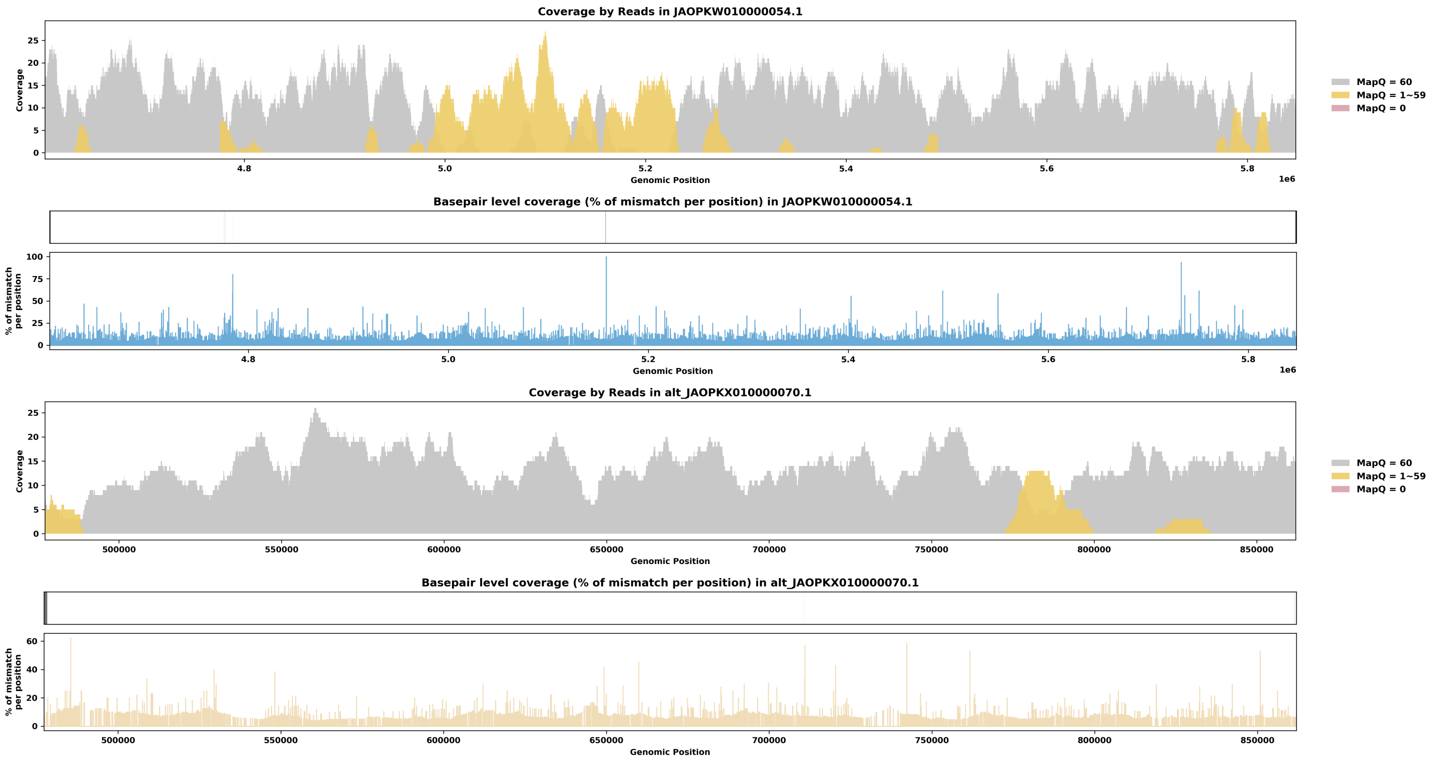

Species ID: mPhoPho1

Common Name: harbor porpoise

Scientific Name: Phocoena phocoena

Assembly Type: Not Haplotype Resolved

Data Source: VGP

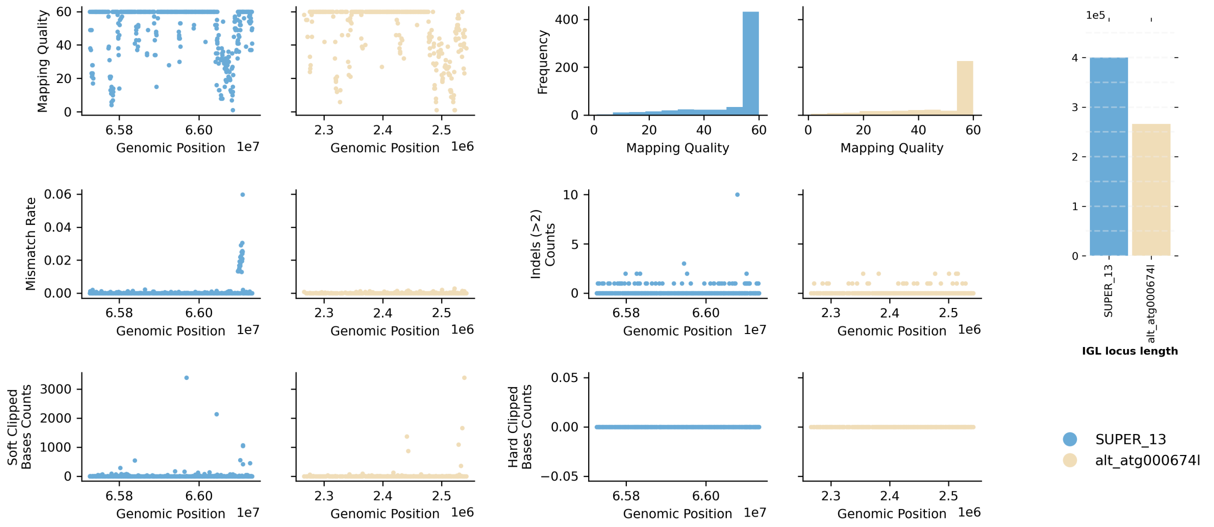


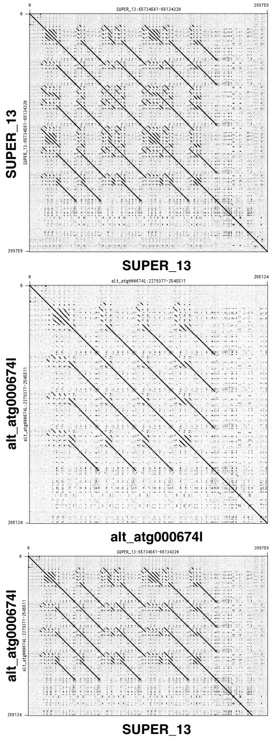


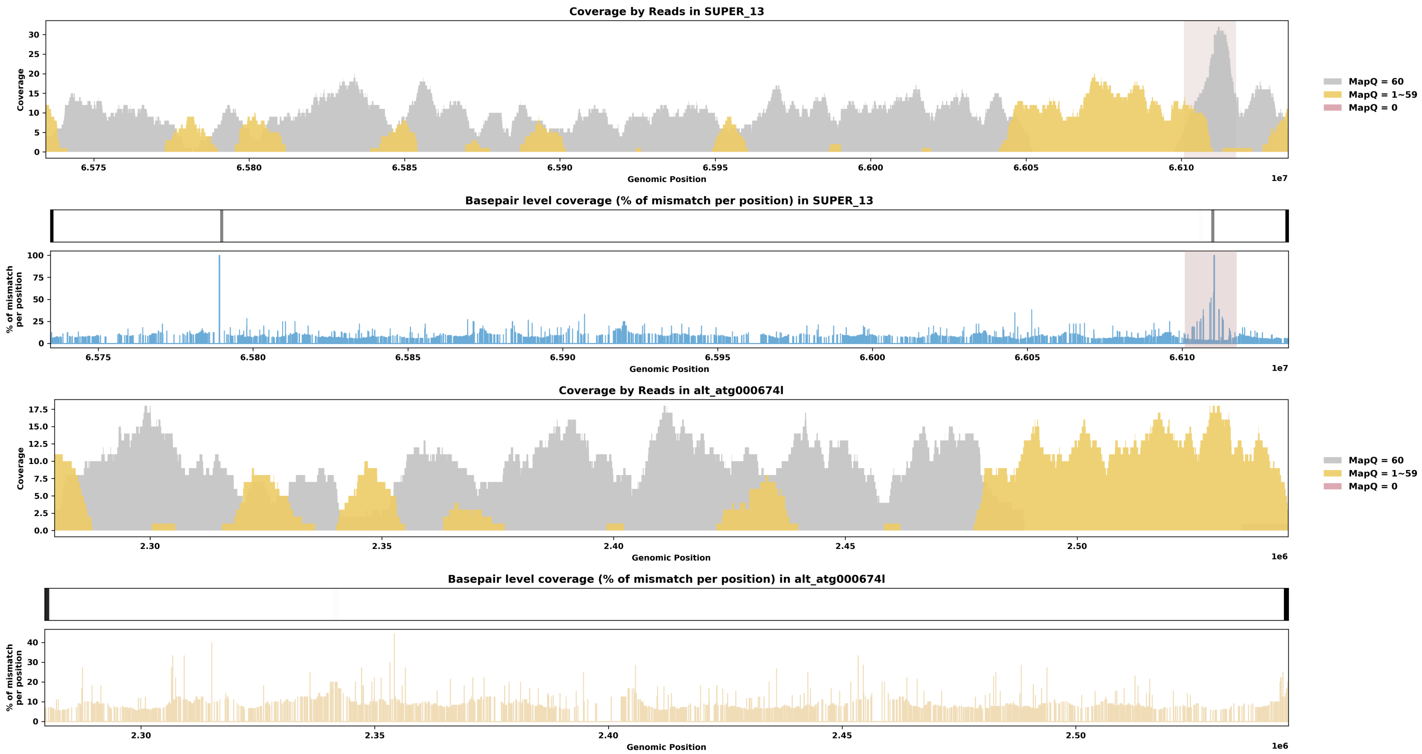

Species ID: mPipPyg2

Common Name: soprano pipistrelle

Scientific Name: Pipistrellus pygmaeus

Assembly Type: Not Haplotype Resolved

Data Source: VGP

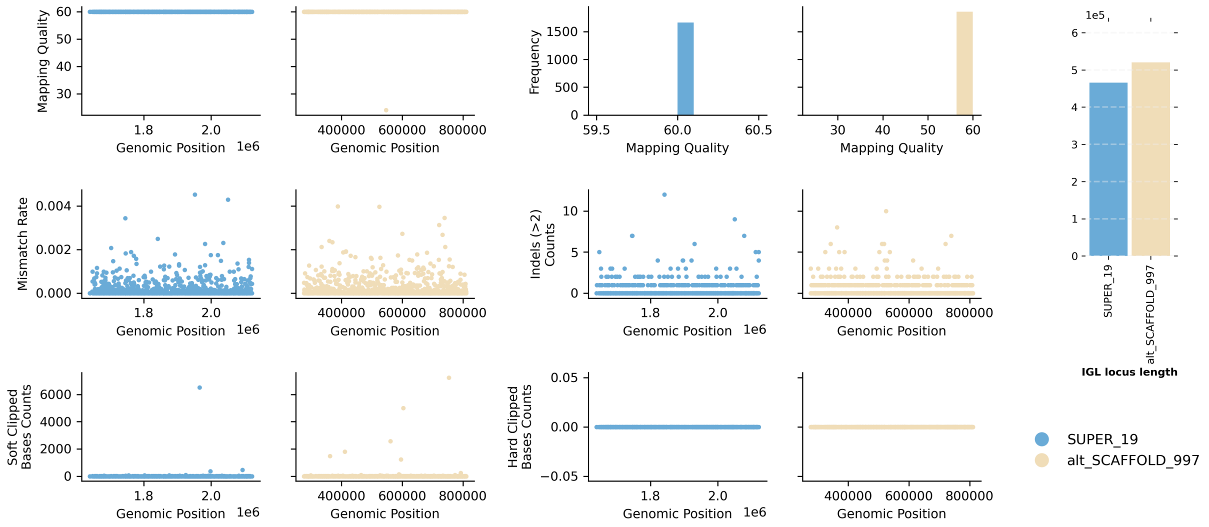


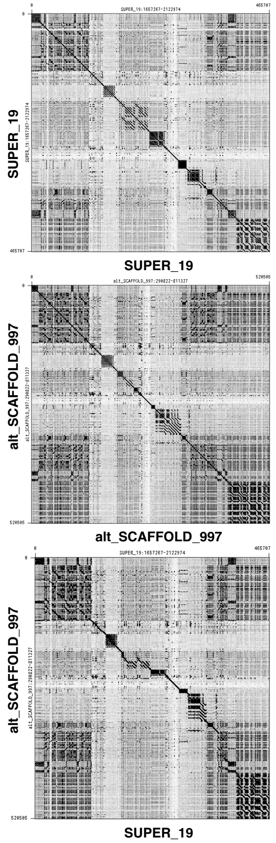


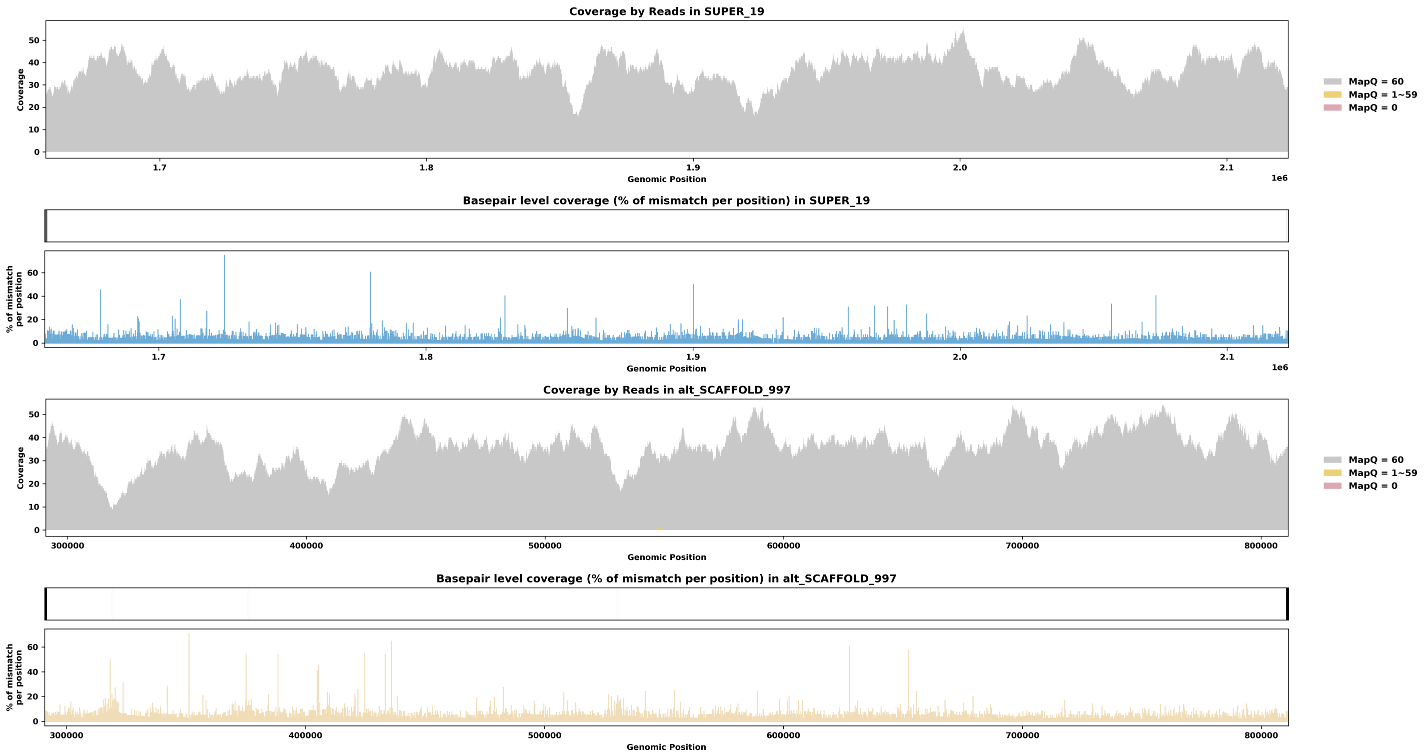

Species ID: mPonAbe1

Common Name: Sumatran orangutan

Scientific Name: Pongo_abelii

Assembly Type: Haplotype Resolved

Data Source: T2T Primate

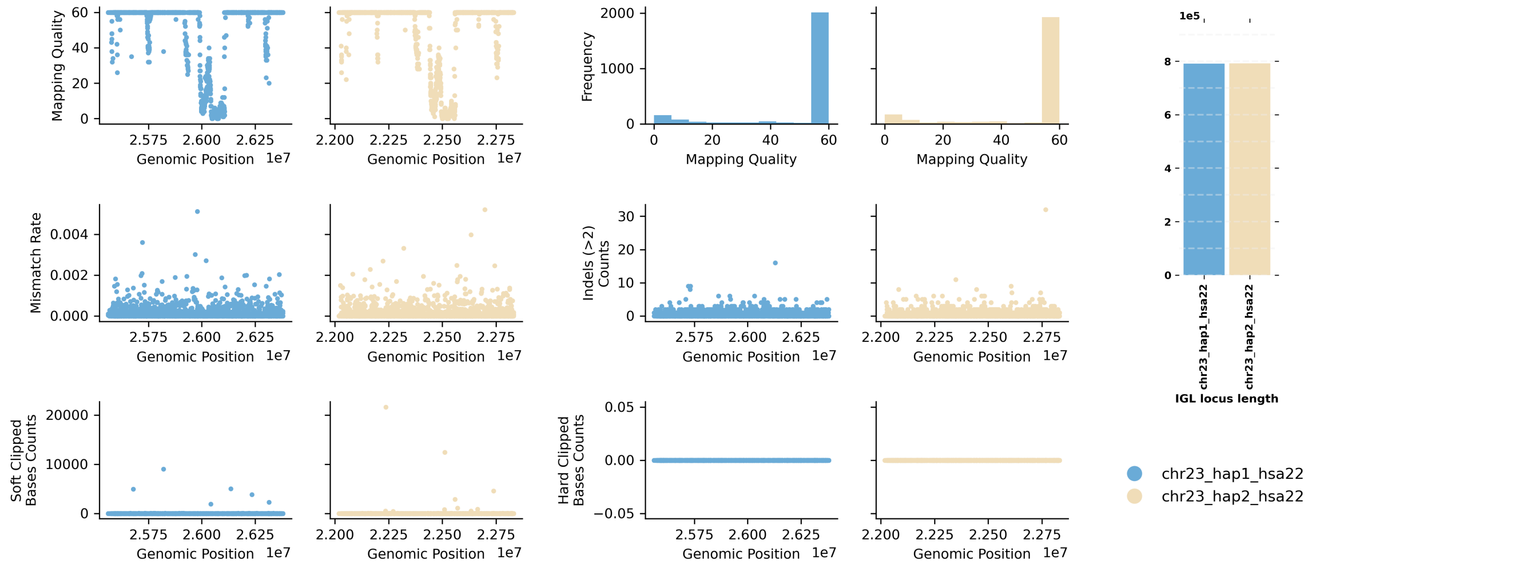


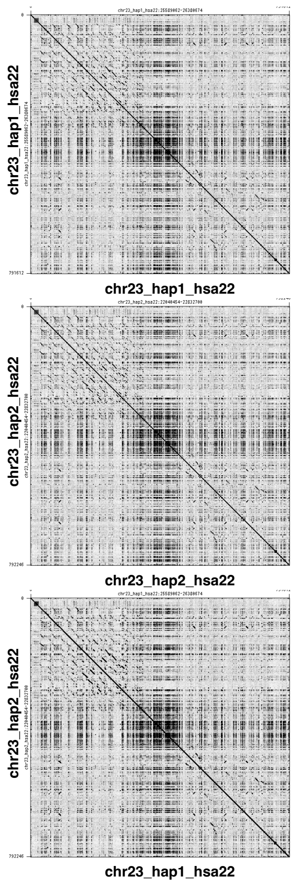


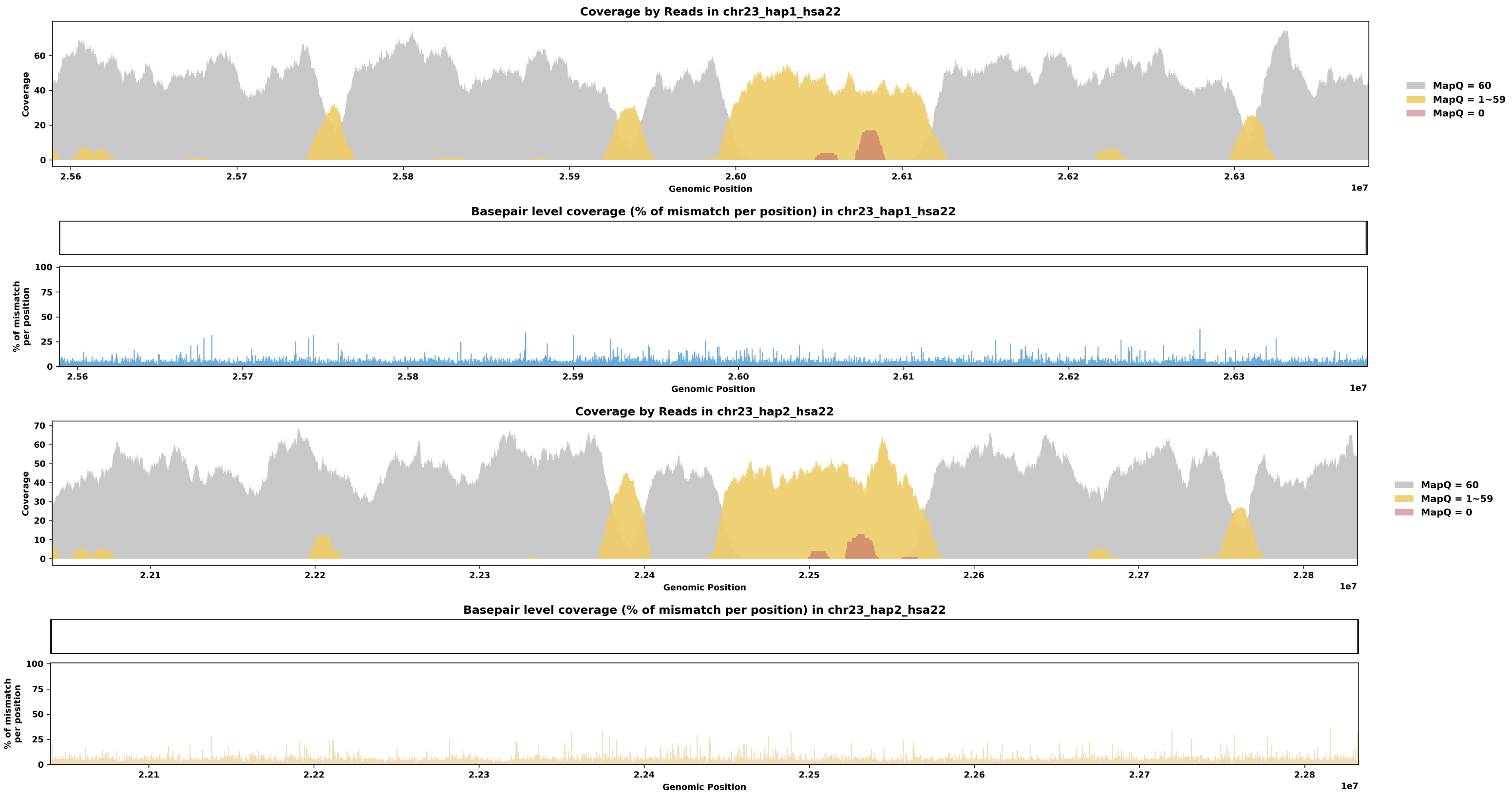

Species ID: mPonPyg2

Common Name: Bornean orangutan

Scientific Name: Pongo_pygmaeus

Assembly Type: Haplotype Resolved

Data Source: T2T Primate

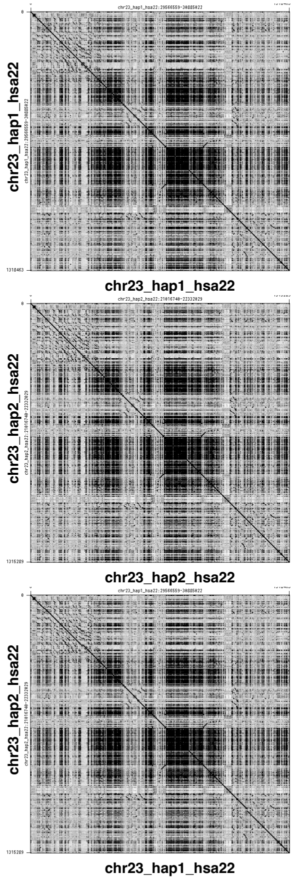


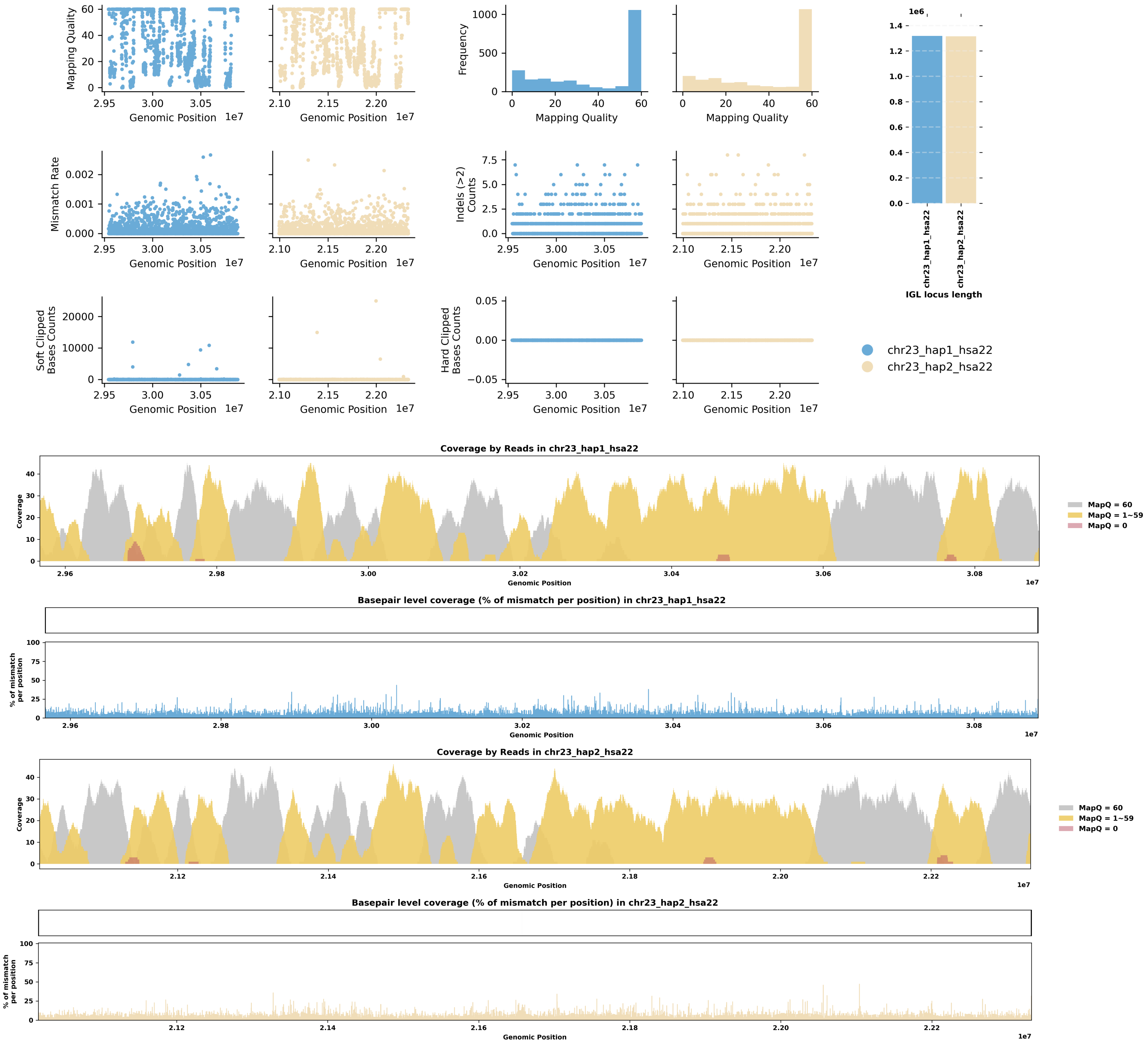

Species ID: mPseCra1

Common Name: false killer whale

Scientific Name: Pseudorca crassidens

Assembly Type: Haplotype Resolved

Data Source: VGP

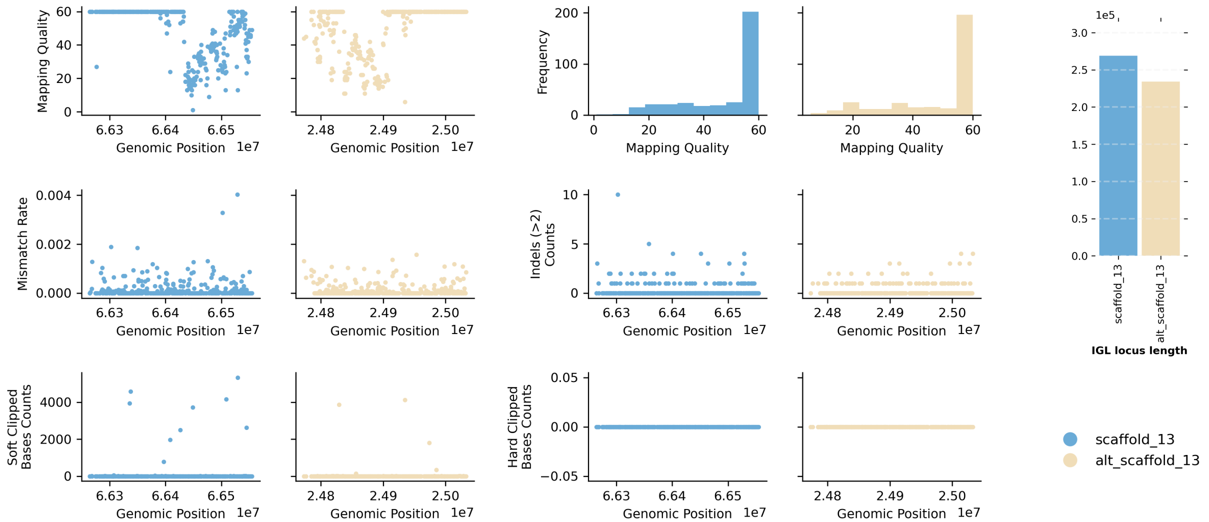


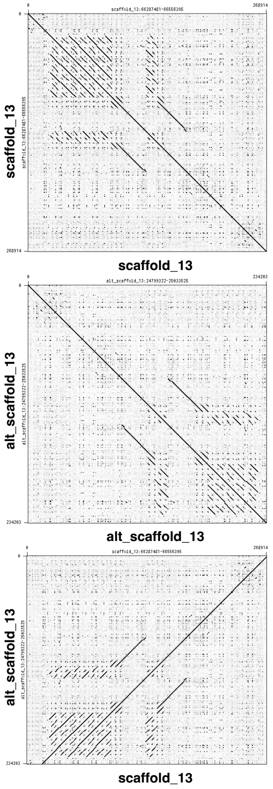


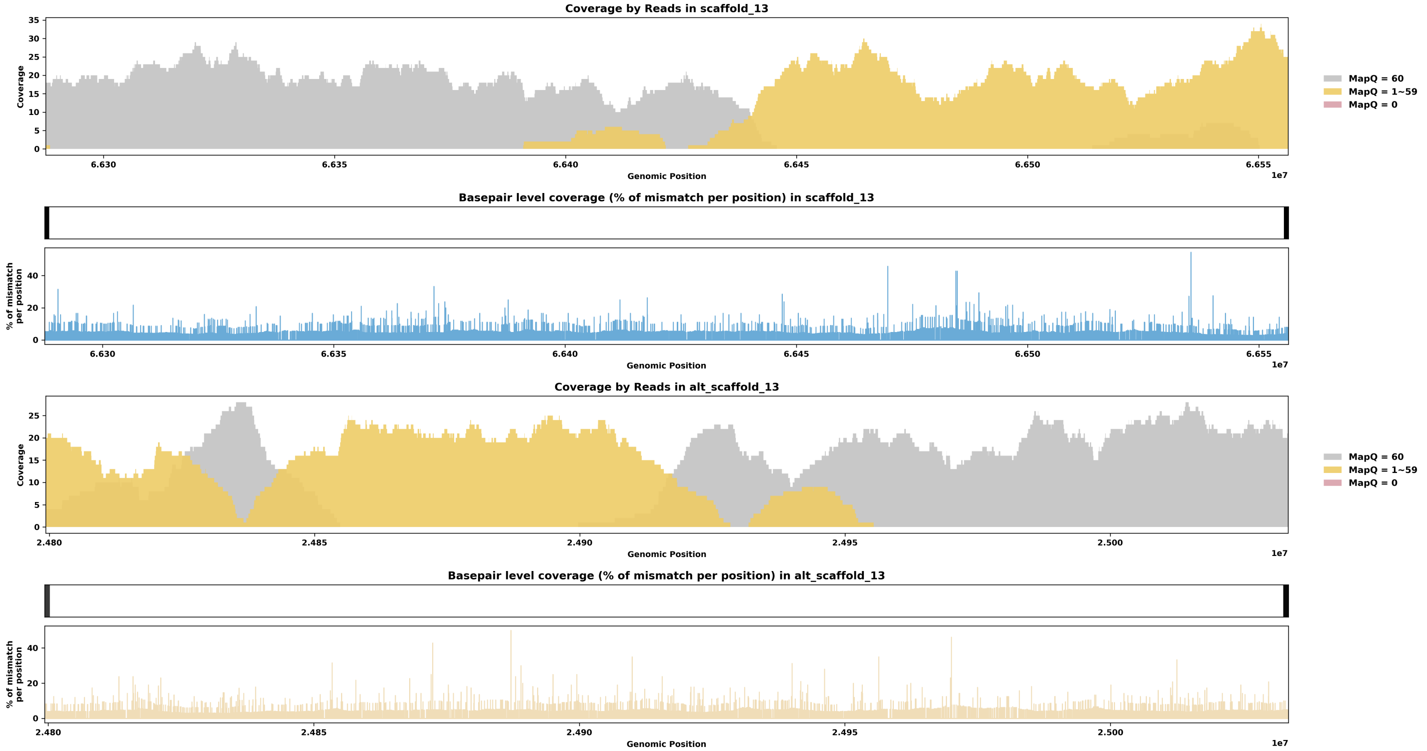

Species ID: mPumCon1

Common Name: Mountain Lion

Scientific Name: Puma concolor

Assembly Type: Haplotype Resolved

Data Source: CCGP

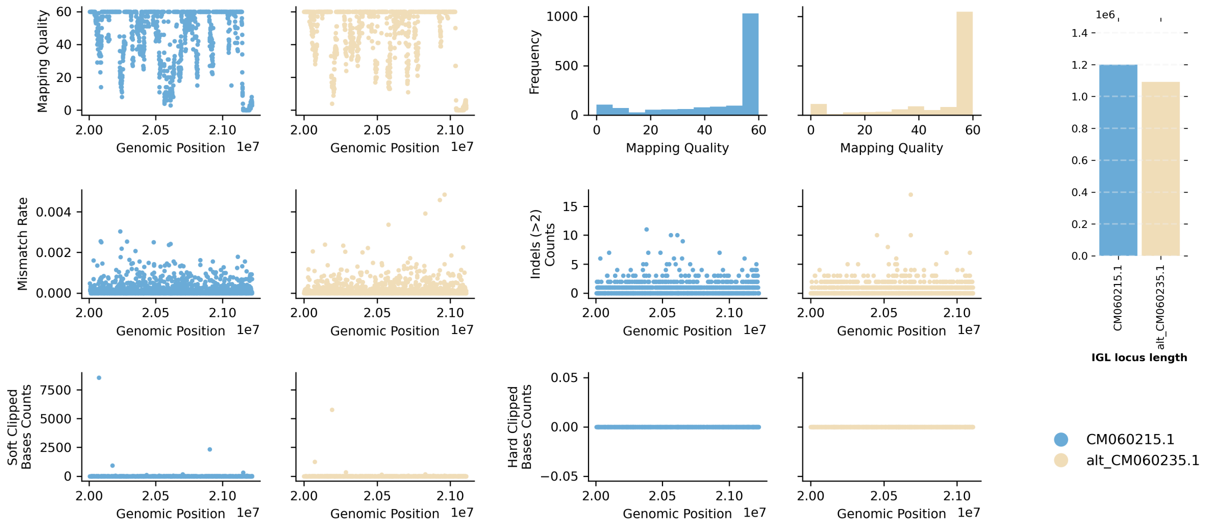


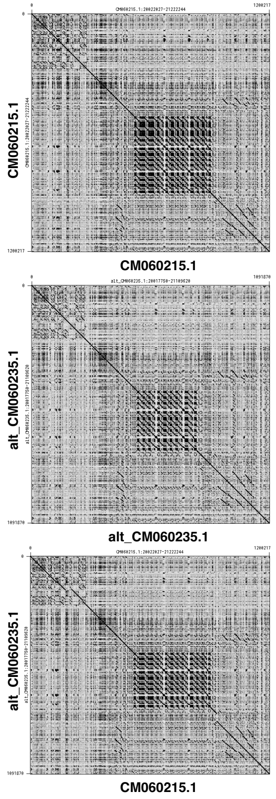


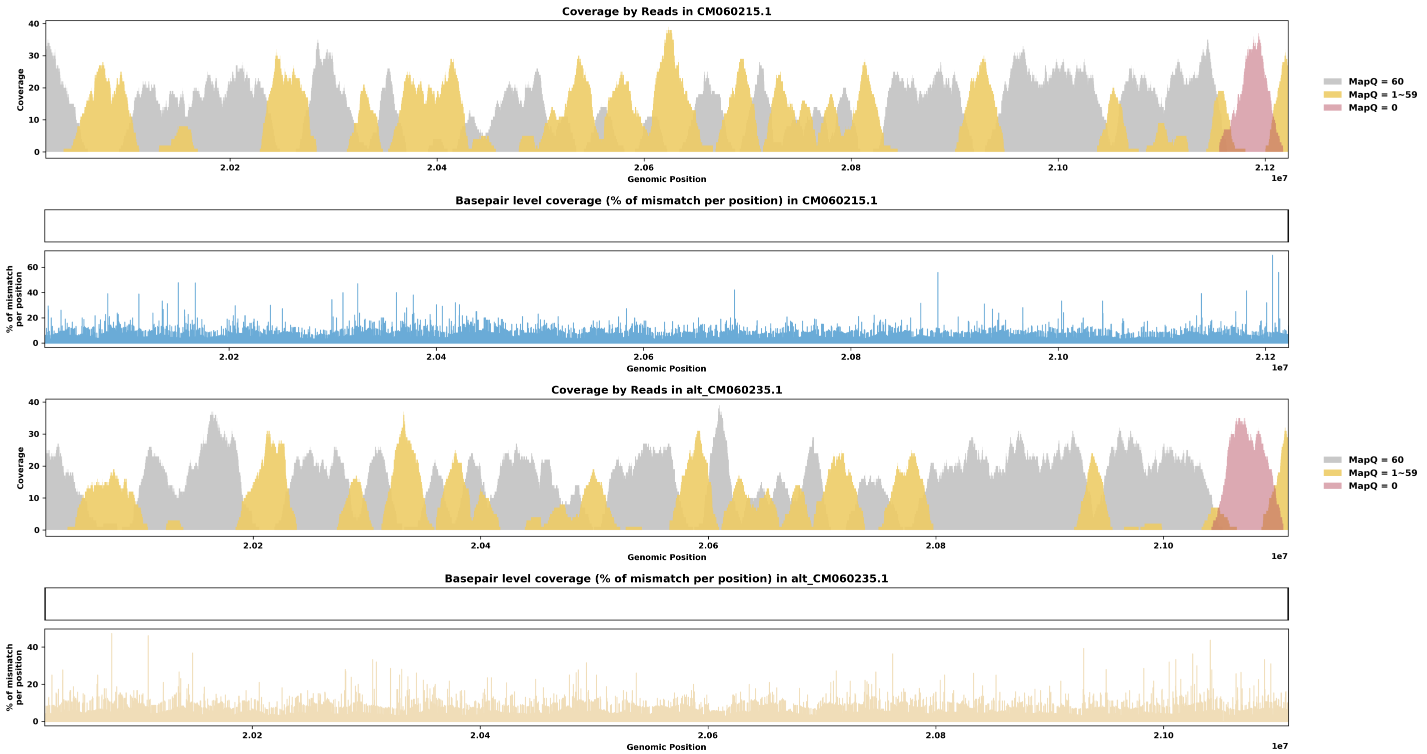

Species ID: mSorAra2

Common Name: Common shrew

Scientific Name: Sorex araneus

Assembly Type: Not Haplotype Resolved

Data Source: VGP

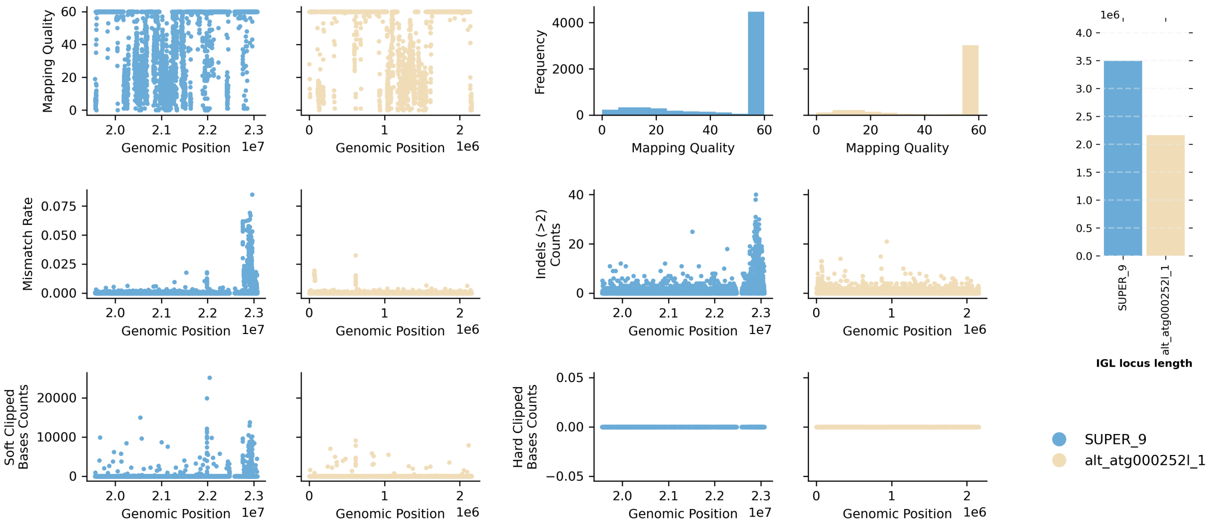


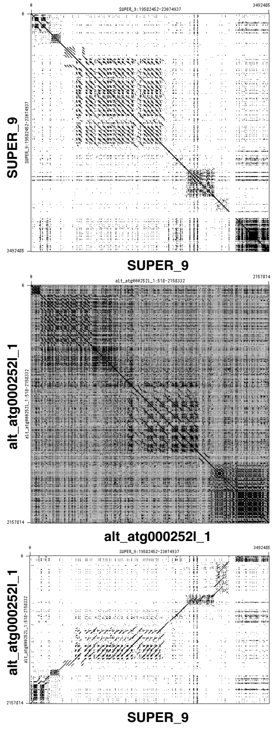


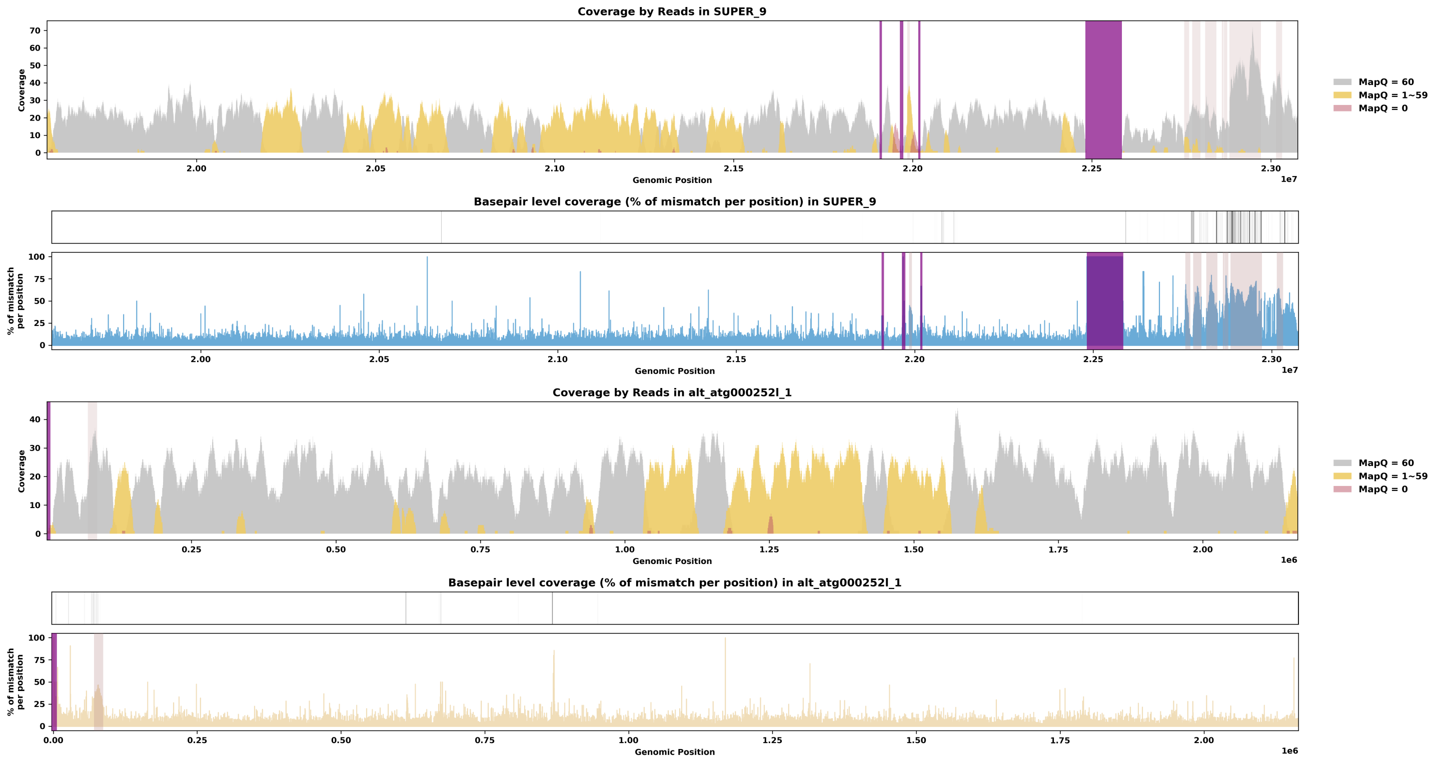

Species ID: mSteCoe1

Common Name: striped dolphin

Scientific Name: Stenella coeruleoalba

Assembly Type: Not Haplotype Resolved

Data Source: VGP

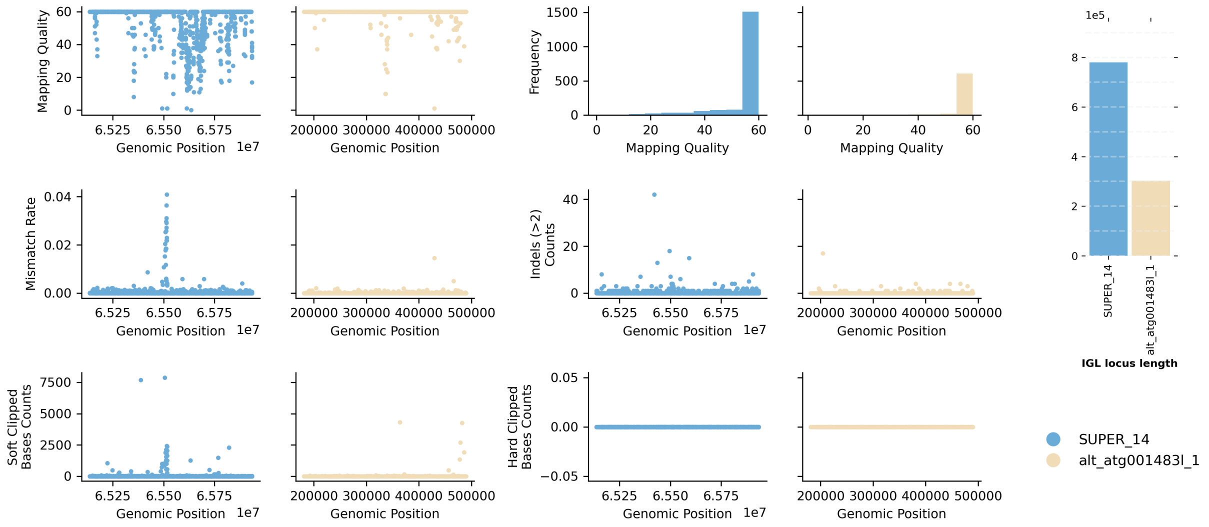


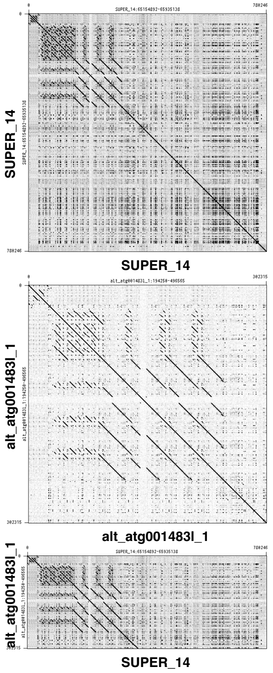


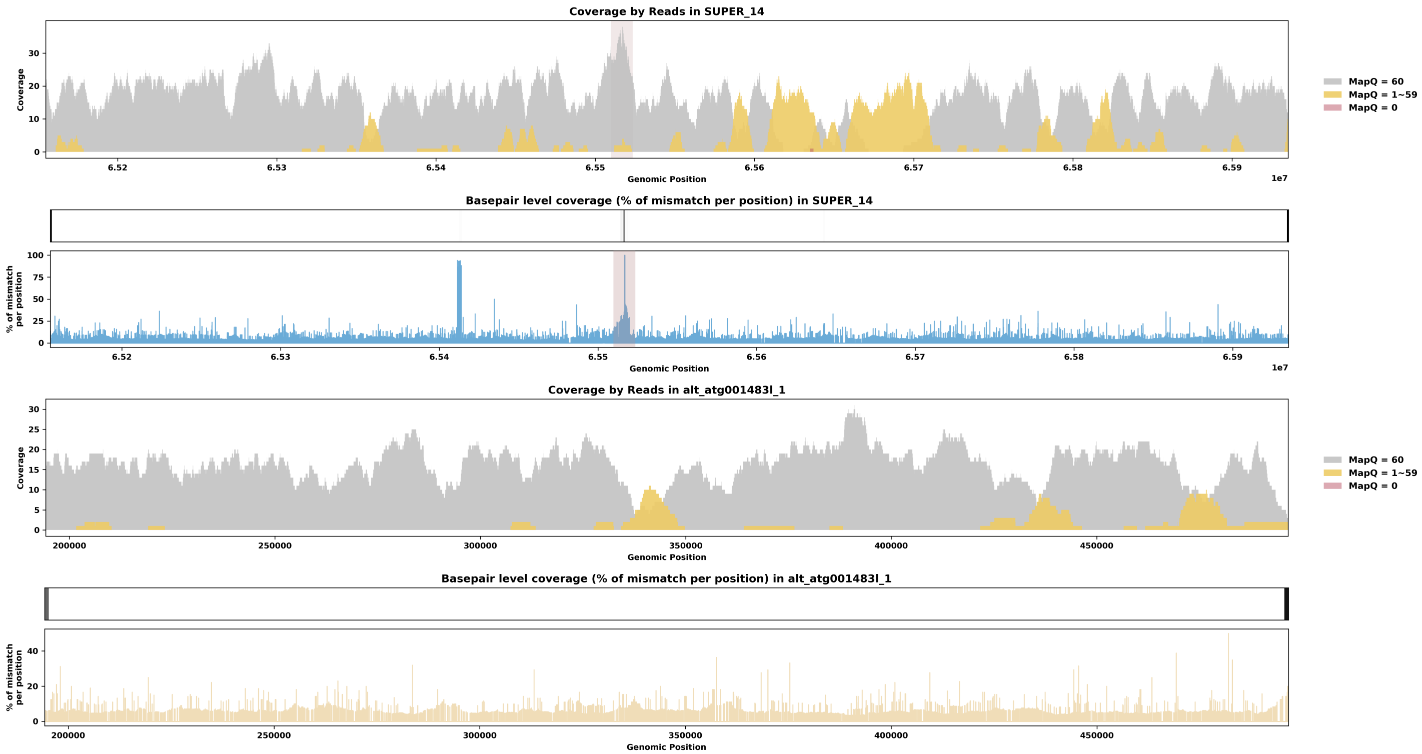

Species ID: mTalEur1

Common Name: European mole

Scientific Name: Talpa europaea

Assembly Type: Not Haplotype Resolved

Data Source: VGP

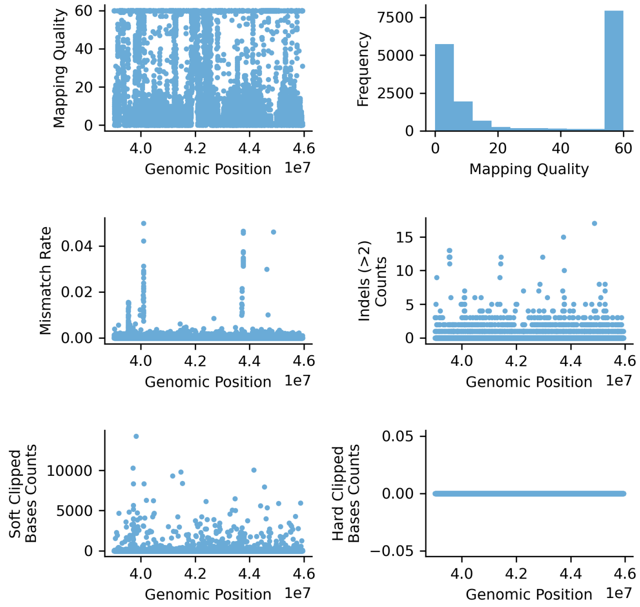


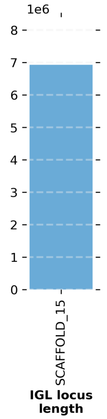


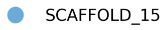


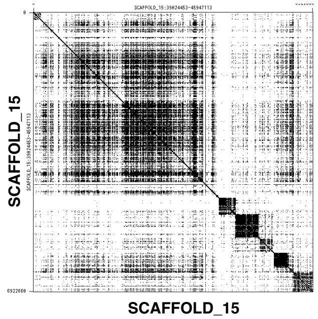


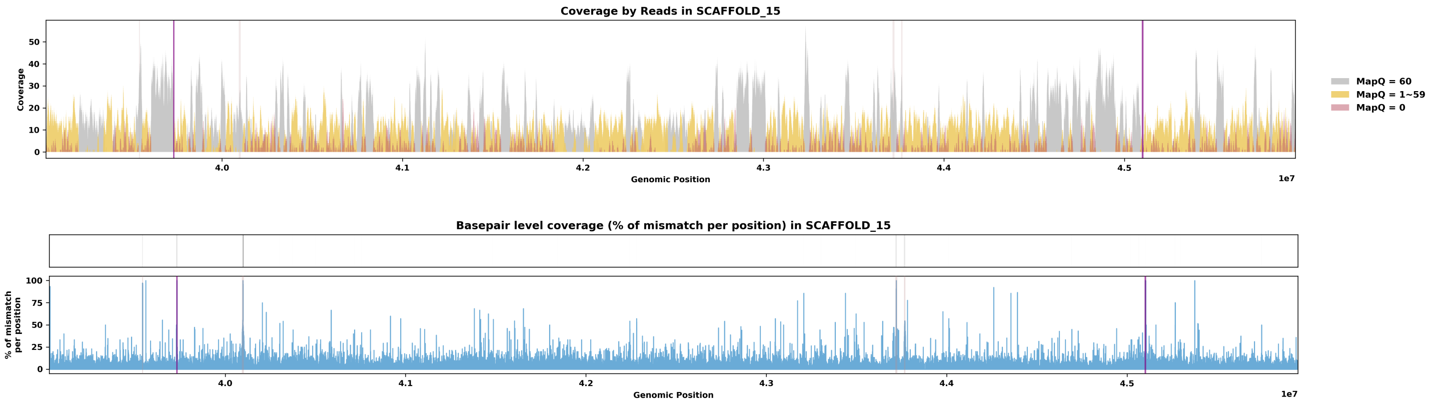

Species ID: mUrsAme1

Common Name: American black bear

Scientific Name: Ursus americanus

Assembly Type: Not Haplotype Resolved

Data Source: CCGP

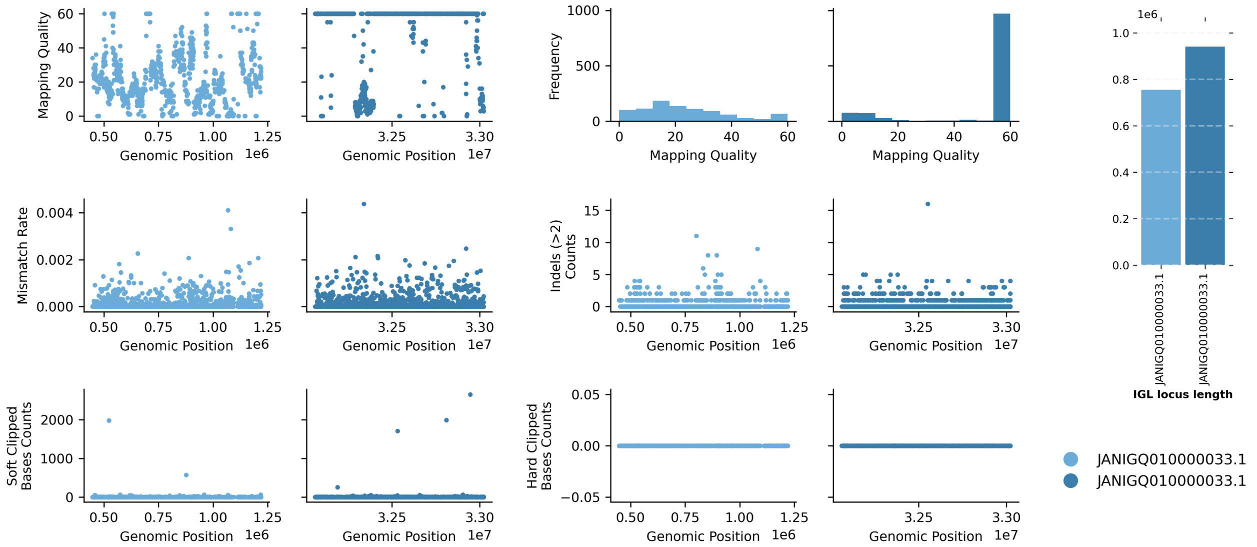


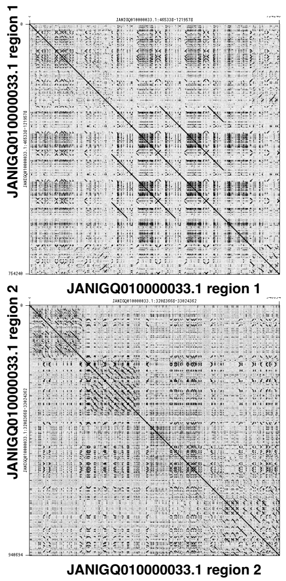


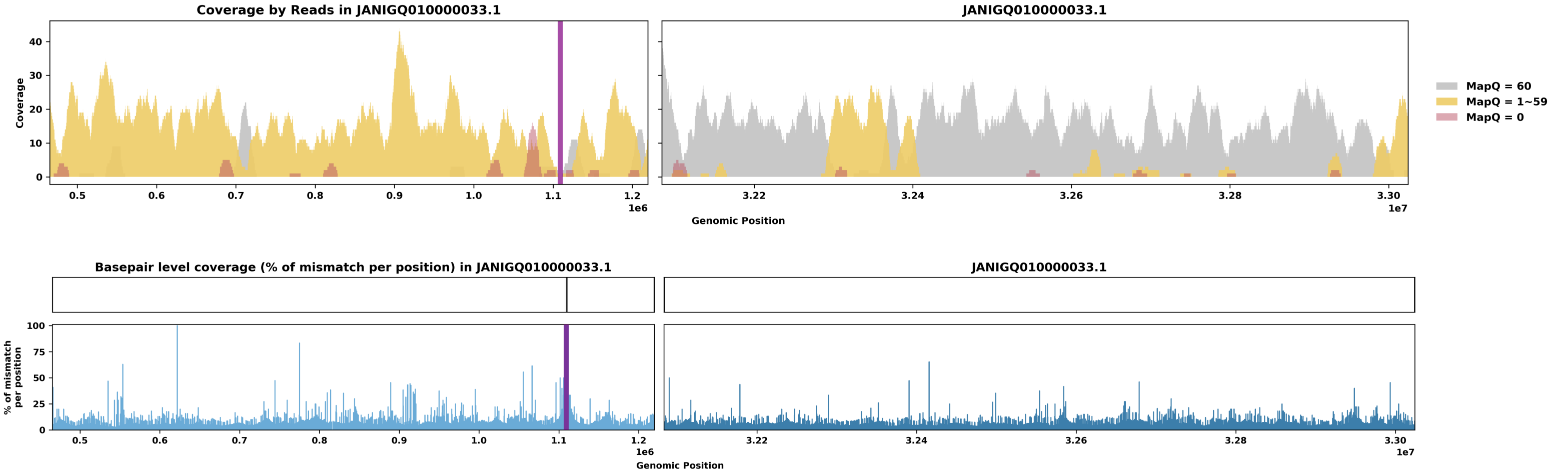

Species ID: mUrsArc2

Common Name: brown bear

Scientific Name: Ursus arctos

Assembly Type: Not Haplotype Resolved

Data Source: NCBI

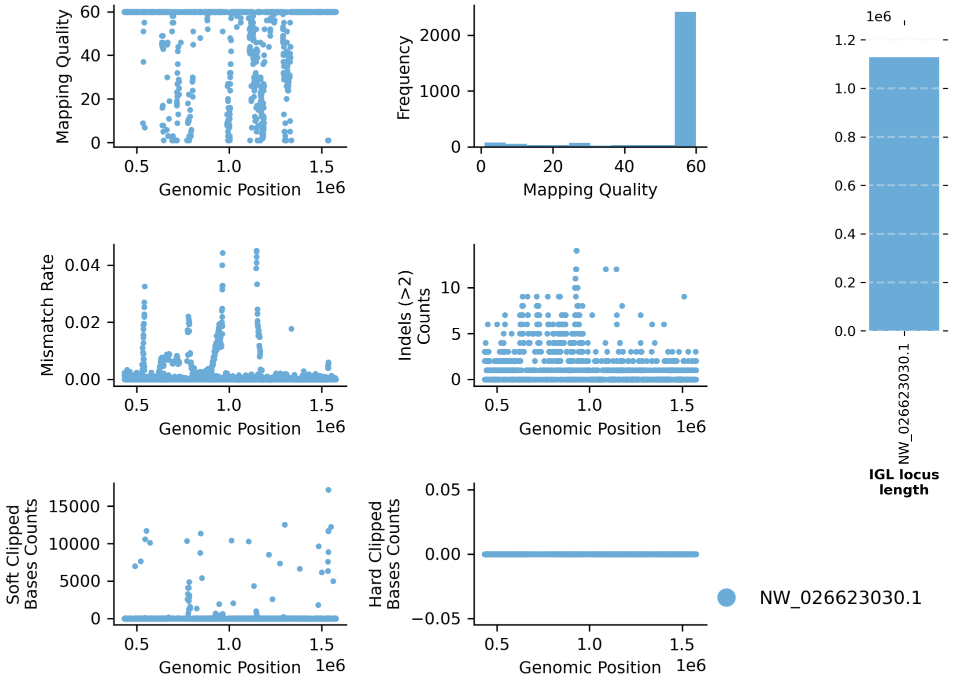


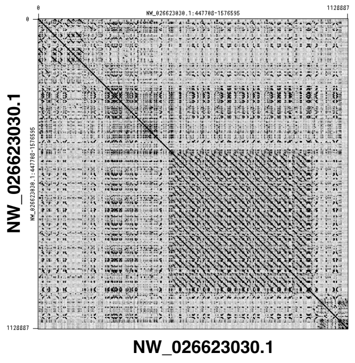


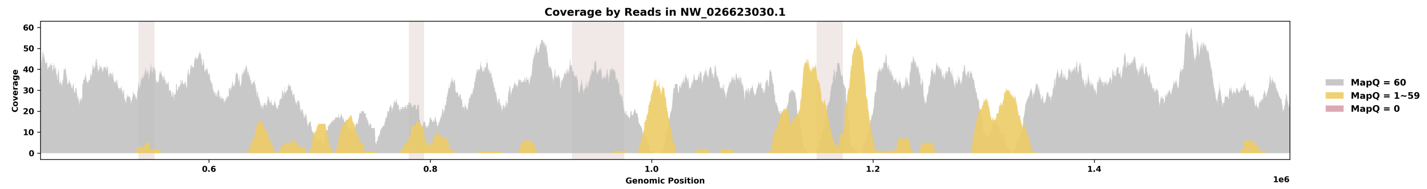


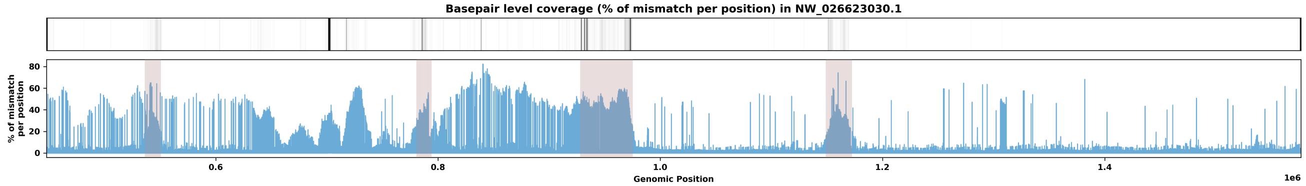

Species ID: mVesMur1

Common Name: particolored bat

Scientific Name: Vespertilio murinus

Assembly Type: Not Haplotype Resolved

Data Source: VGP

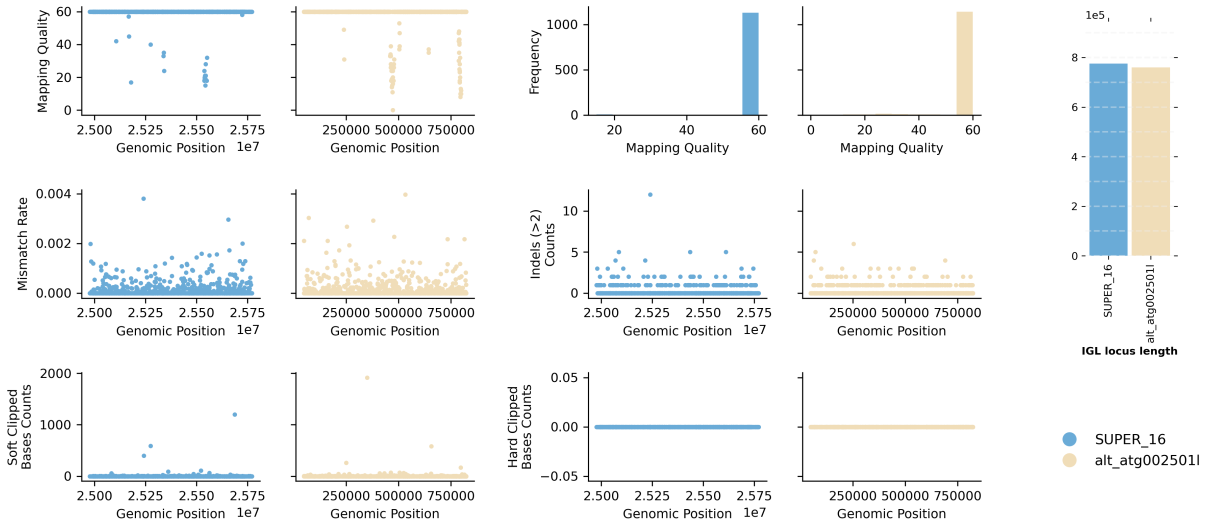


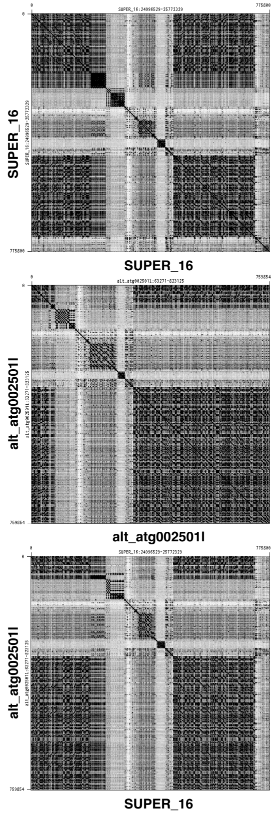


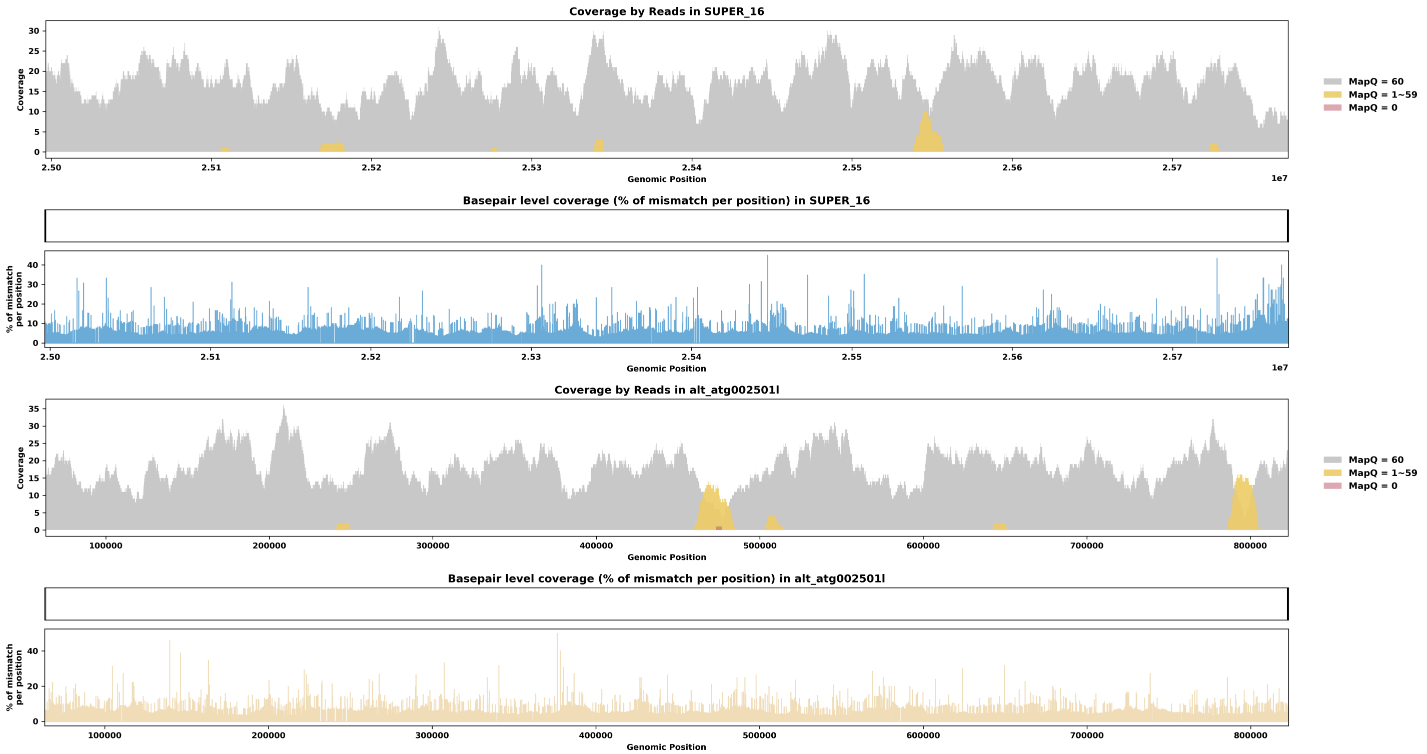

Species ID: rAllMis2

Common Name: American alligator

Scientific Name: Alligator mississippiensis

Assembly Type: Haplotype Resolved

Data Source: VGP

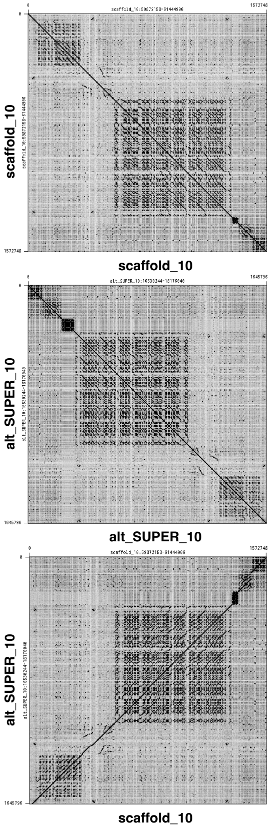


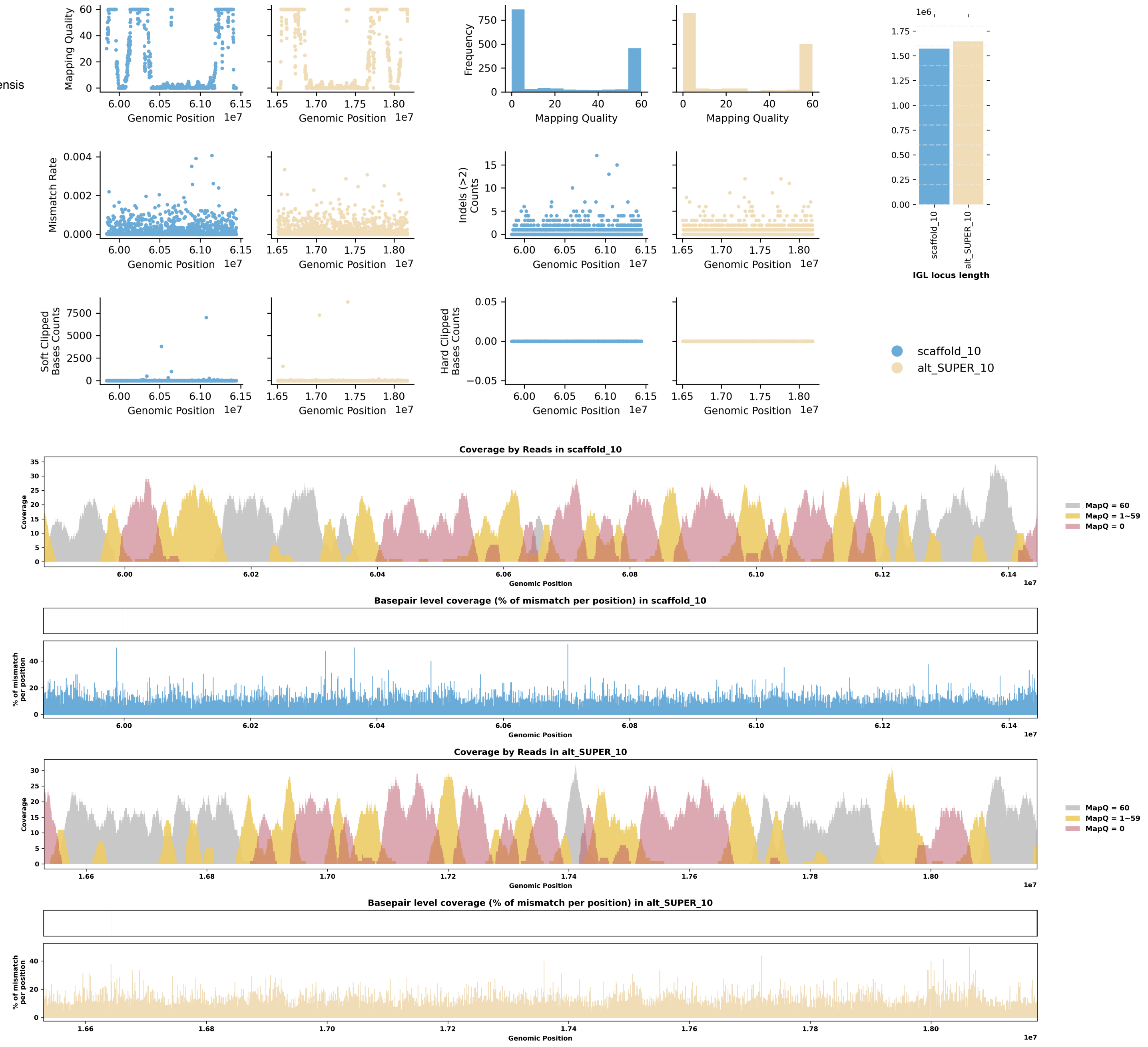

Species ID: rCarCar2

Common Name: Loggerhead turtle

Scientific Name: Caretta caretta

Assembly Type: Haplotype Resolved

Data Source: VGP

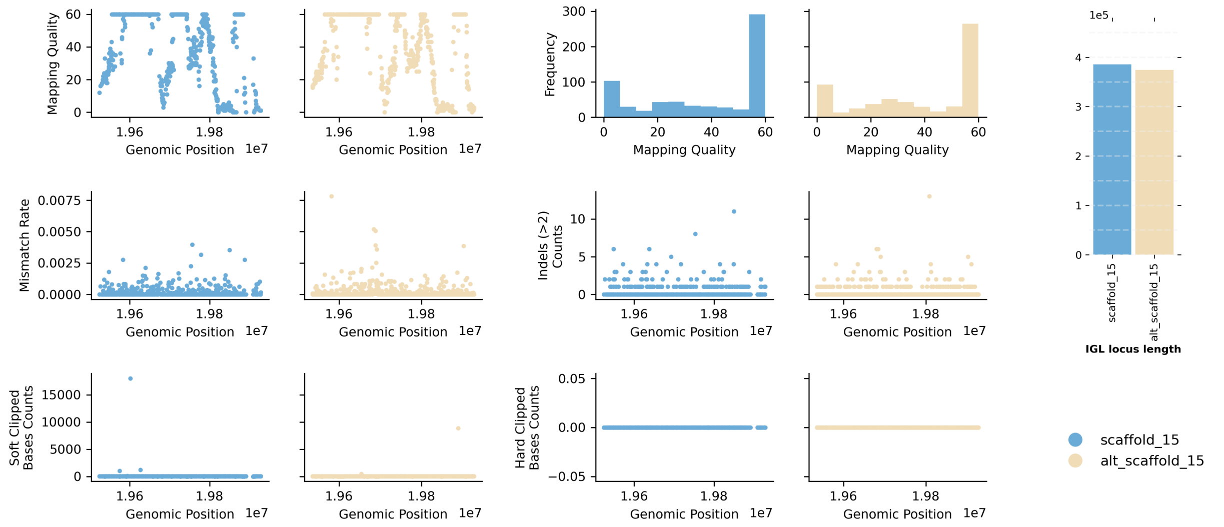


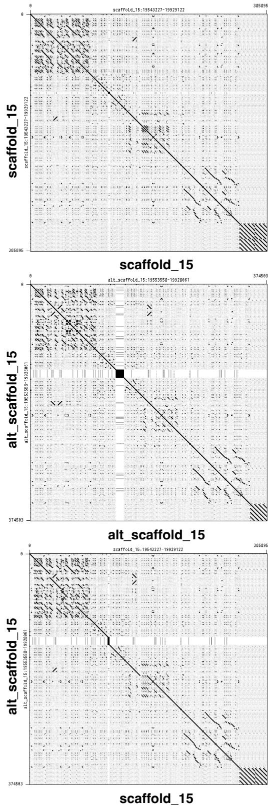


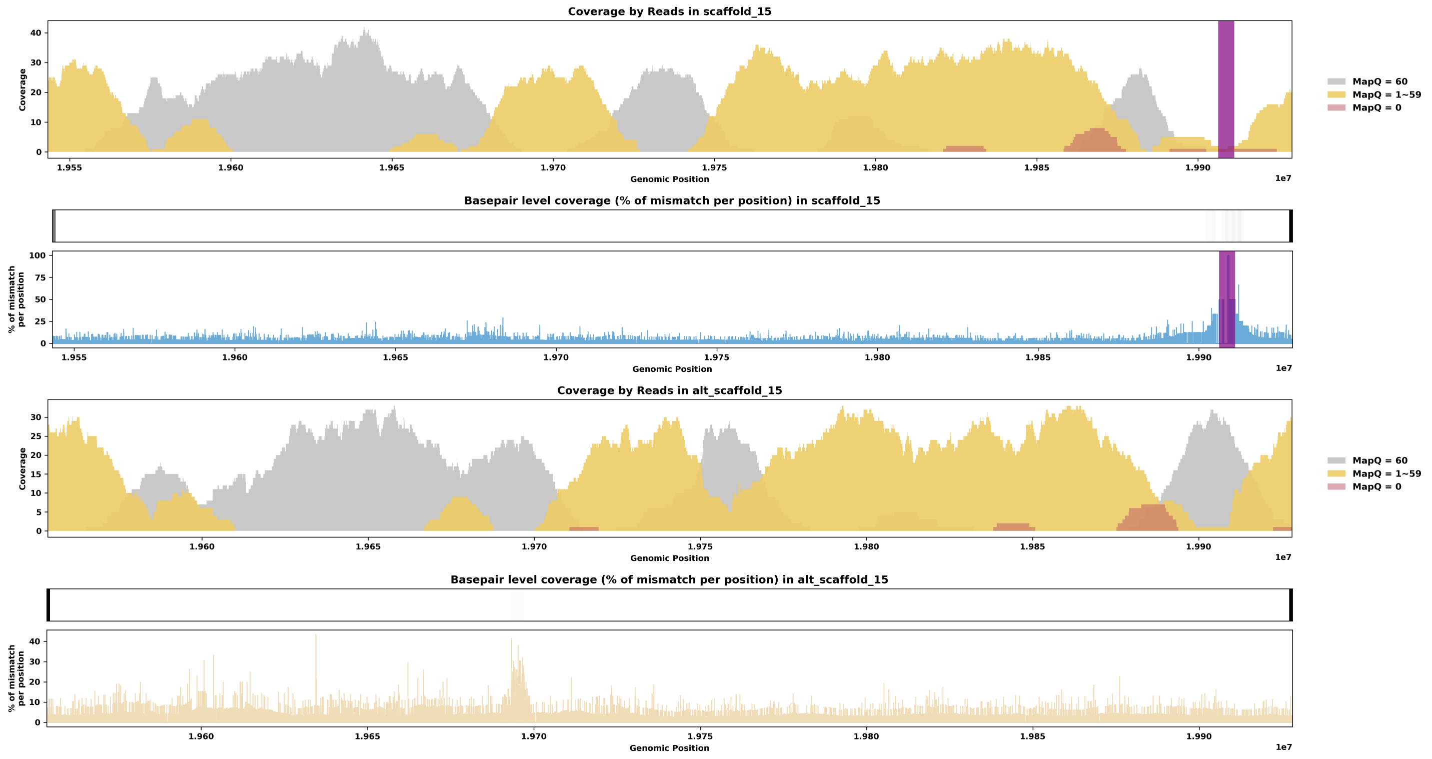

Species ID: rEmyOrb1

Common Name: European pond turtle

Scientific Name: Emys orbicularis

Assembly Type: Haplotype Resolved

Data Source: VGP

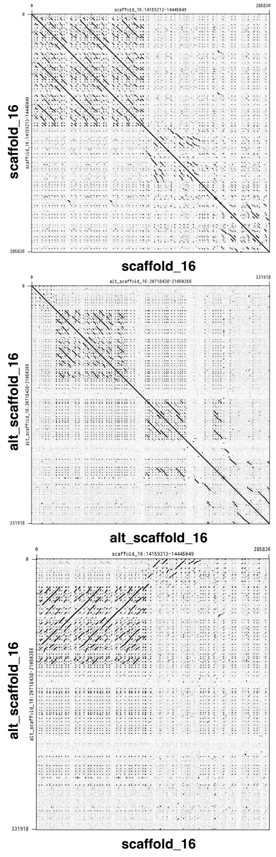


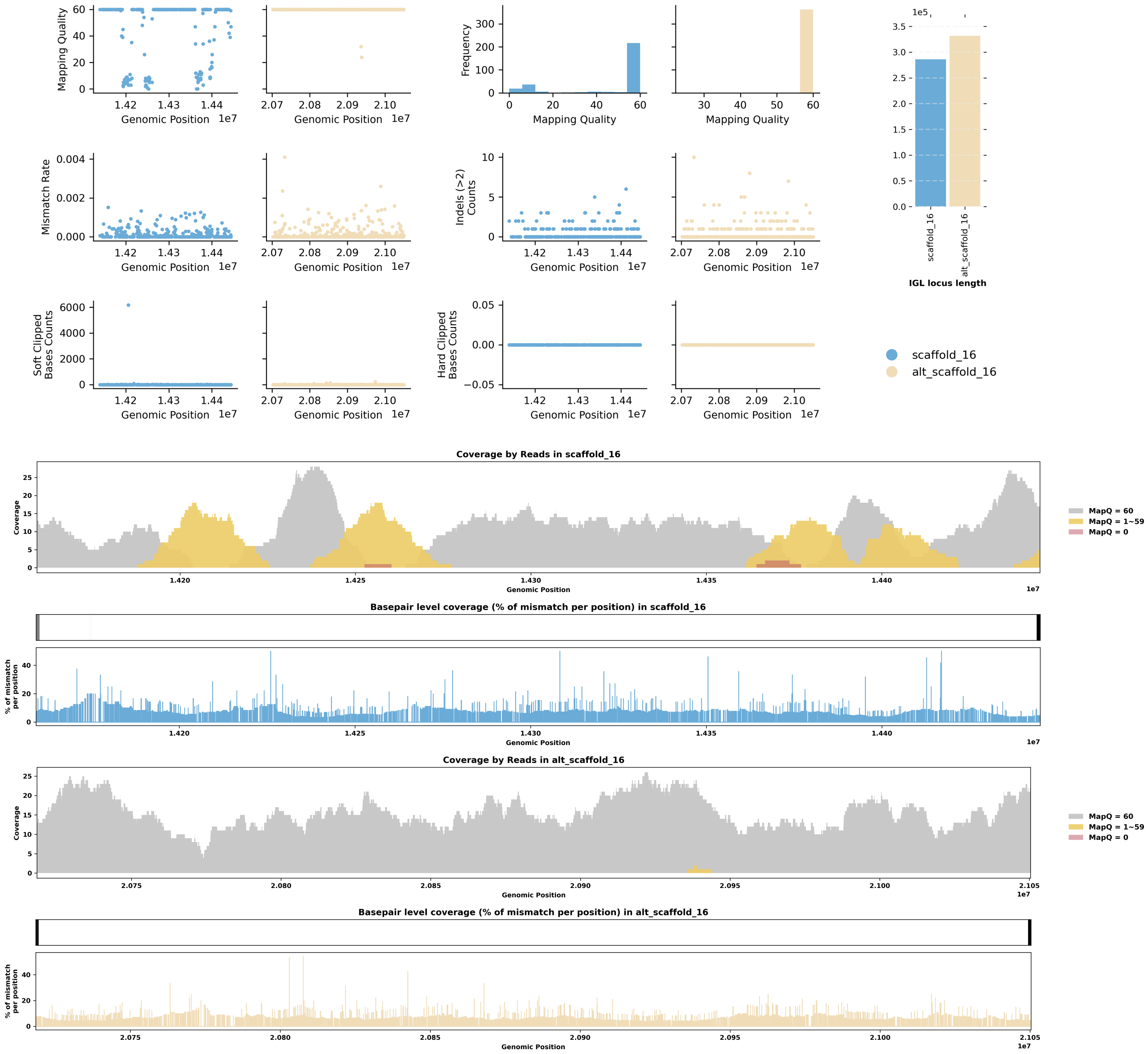

Species ID: rMalTer1

Common Name: diamondback terrapin

Scientific Name: Malaclemys terrapin

Assembly Type: Haplotype Resolved

Data Source: VGP

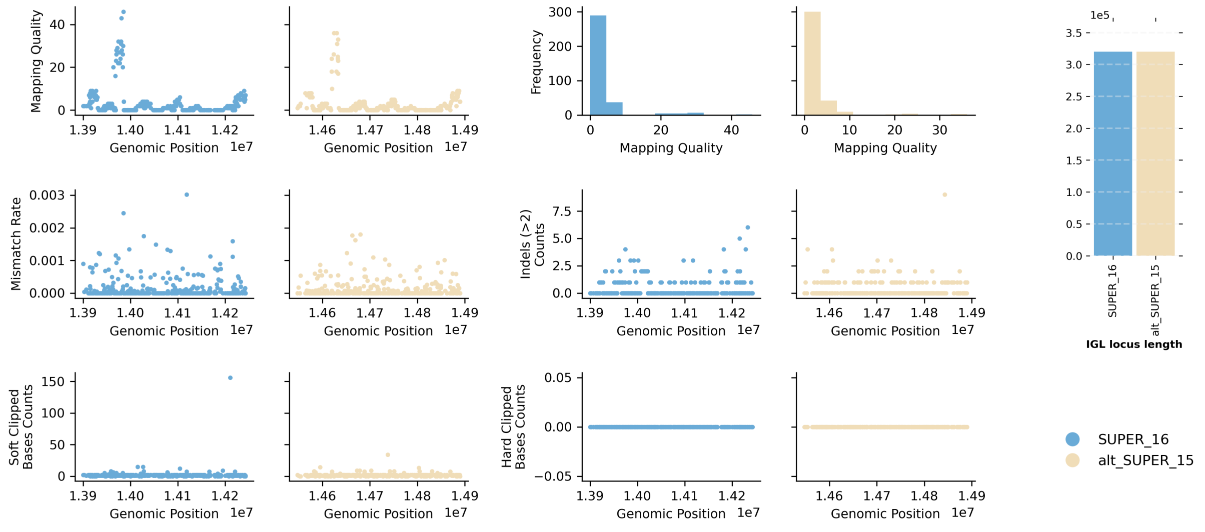


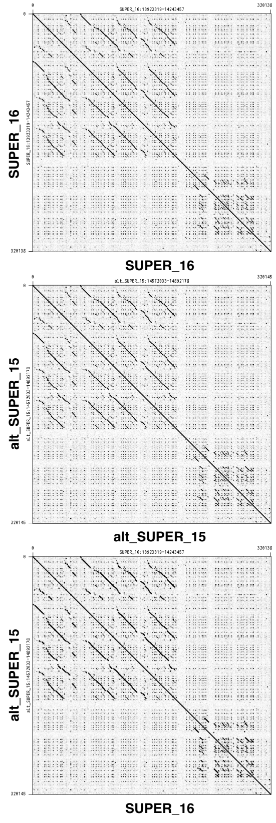


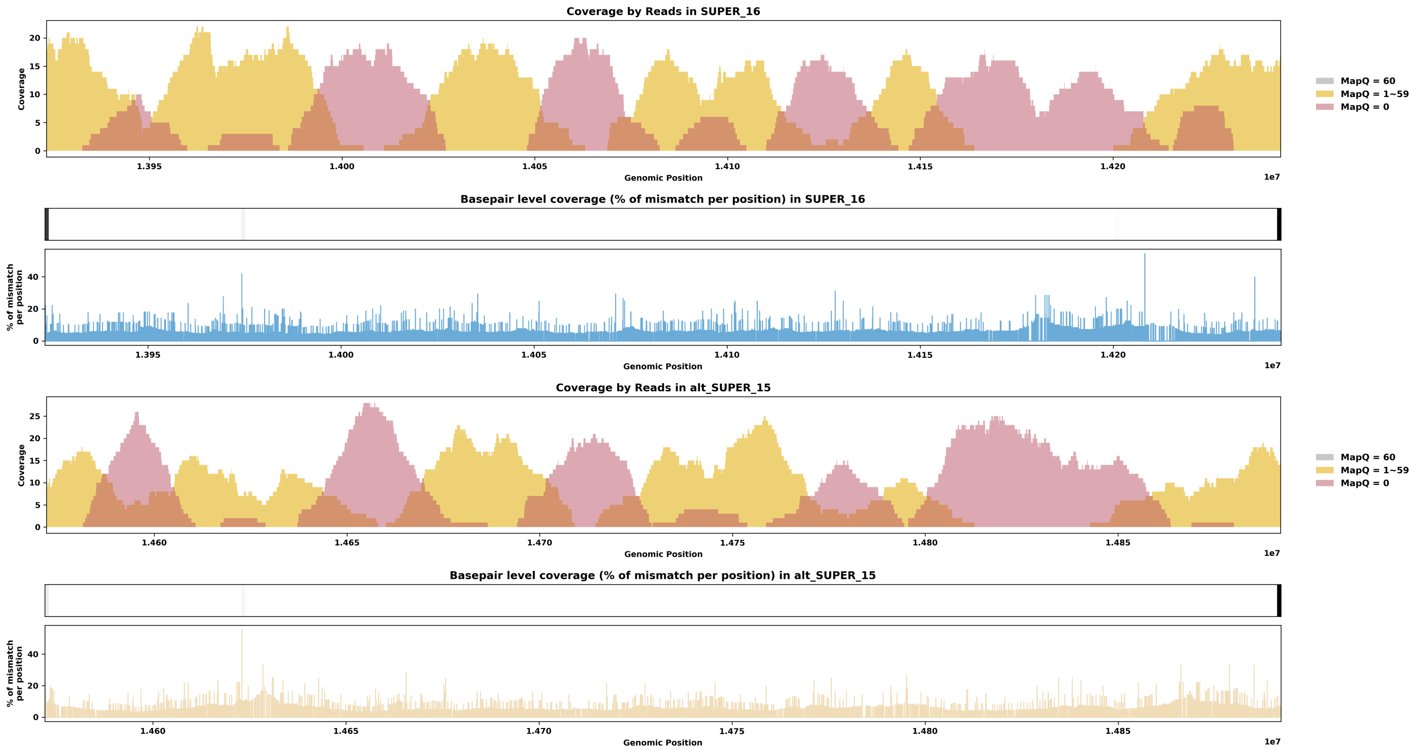
